# A portable N-terminal biphenyl motif in LL-37 enhances the activity of short human cathelicidin derivatives

**DOI:** 10.1101/2025.05.22.655619

**Authors:** John S. Albin, Dimuthu A. Vithanage, Corey Johnson, Bradley L. Pentelute

**Author notes:** Sanofi, Cambridge, MA 02141, USA. Amide Technologies, Waltham, MA 02451, USA.

## Abstract

Issues such as potency and stability limit our ability to realize the potential of host defense peptides (HDPs) to serve as new antibiotics. Informed by our prior structure-activity work, we demonstrate that transposition of an N-terminal biphenyl motif from full-length LL-37 onto the previously-described central activity region (residues 18-29) improves activity against gram-negative bacteria by >16-fold. We further improve upon this lead derivative, termed FF-14, using stabilizing modifications such as D-amino acids and C-terminal amidation as well as selected N-lipidation moieties, and demonstrate the transferability of biphenyl motif activity to longer cathelicidin scaffolds. Mechanistic interrogation reveals that the biphenyl pharmacophore increases both inner and outer membrane permeabilization while promoting helical structure in the 14-mer without directly impacting peptide stability. Inclusion of the LL-37 biphenyl motif is thus a portable strategy for augmenting short cathelicidin activity and better approximating the activity profile of LL-37 itself in short sequences.

## Introduction

Antimicrobial resistance is a growing threat to human health, accounting for 1.2 million deaths in 2019, particularly due to infections with resistant gram-negative bacteria^1^. This death toll is further expected to escalate in the coming decades^2^, highlighting the importance of developing new antibiotics. One approach to the pursuit of new antibiotics is to develop derivatives of host defense peptides (HDPs), which are structurally diverse peptides produced by all branches of life to defend against microbes^3,4^. Human HDPs, in particular, are attractive templates. Having evolved in the context of the human immune system, LL-37 and other human HDP templates are generally nonimmunogenic in their starting sequences, but may encode desirable immunomodulatory activities to enhance the clearance of an infection^5–8^. Moreover, because these templates are chemically simple, it is straightforward to diversify their structures through the application of noncanonical amino acids and combinatorial chemistry^9,10^.

Despite their promise, the clinical impact of HDPs has been limited by familiar issues such as stability, pharmacokinetics, toxicity, and potency^11^. Though significant, these barriers also present clear opportunities to apply well-characterized peptide-engineering strategies. Consider GLP-1 agonists, for example. Native GLP-1(7-37) is a short, helical peptide with poor stability and pharmacokinetics. The difference between this nonviable therapeutic lead and the transformative drug semaglutide is three amino acid changes aimed primarily at improving the stability and pharmacokinetics of the native peptide^12^. In principle, the same rules should apply to structurally similar helical peptides such as LL-37, which is active against gram-negative bacteria and demonstrates a wide range of proposed immunomodulatory functions that may be of therapeutic utility^13–19^.

There are a number of well-characterized approaches by which to modulate peptide stability through strategies such as stapling^20,21^, primary sequence changes^12^, and peptidomimetic alterations including the use of D-amino acids inert to native L-proteases^22^. Many of these have been directly applied to cathelicidin derivatives before, and it is likely that they will prove relevant to many distinct primary cathelicidin sequences. Similarly, a rich literature describes a multitude of strategies for improving the therapeutic properties of distinct peptides, and some such as lipidation or sequence modifications to control stability and pharmacokinetics through targeted binding of serum components may also be applicable to cathelicidins^23–26^.

Less obvious, however, is the path forward for addressing issues of cathelicidin toxicity and antimicrobial potency, which pose barriers beyond those common to endocrine peptides. Many prior reports describe the activities of diverse fragments of LL-37^27^, for example, but it is not always clear to what extent these short units remain structurally stable and mechanistically analogous to LL-37 – or why. A further limiting factor is the incomplete selectivity of cathelicidin derivatives between mammalian and bacterial membranes, which can be difficult to ablate even in highly optimized HDP sequences^28,29^. To address this toxicity gap, we have recently reported structure-activity relationships in full-length LL-37 that revealed a rational approach to separating activity against gram-negative and mammalian cell membranes by modulating the cathelicidin oligomerization interface^30,31^.

The study presented here builds on a separate observation from these structure-activity relationship studies, namely the loss of antimicrobial activity among some mutants with changes predominantly in the LL-37 N-terminus. This loss of activity with mutations linearly distant from the core activity region spanning residues 18-29 suggests that N-terminal residues may be important for the recapitulation of native LL-37-like activity against gram-negative bacteria^30^.

Developing this hypothesis, we report here that transposition of an N-terminal biphenyl motif to the previously described core activity region yields an enhanced short cathelicidin derivative, FF-14, with >16-fold improved potency against gram-negative bacteria compared with the core activity region alone. We further find that common stabilization strategies such as the use of D-amino acids and C-terminal amidation remain applicable in this template, and that selected N-lipidation moieties may also help to limit mammalian cell toxicity. Mechanistic interrogation reveals that gram-negative outer and inner membrane permeabilization increases with each sequential addition of Phe to the N-terminus, with permeabilization by FF-14 approximating that of LL-37. These N-terminal Phe residues also contribute to enhanced helicity in short derivatives, while distinct cathelicidins demonstrate unique serum stability profiles independent of the presence of the biphenyl motif. The studies described here thus highlight the importance and portability of the biphenyl motif for antimicrobial activity among human cathelicidin derivatives while laying the groundwork for integration with additional strategies for the rational detoxification and pharmacokinetic control of cathelicidin antimicrobials.

## Results

### Limited activity of short derivatives of LL-37 against gram-negative bacteria

By convention, peptide derivatives of the human cathelicidin *CAMP* gene are named according to their first two amino acids and their overall length – thus, LL-37 starts with the amino acids Leu-Leu and is 37 residues long^32–35^. A large number of studies have previously reported on the activity of distinct fragments of LL-37^27^. Among these, KR-12, a 12-residue fragment extending from residues 18-29 of the native LL-37 peptide, is generally considered the definitive minimal unit of LL-37 activity to which many peptidomimetic alterations have been previously applied^36–48^, and longer derivatives typically incorporate this same region^49–52^.

The initial description of KR-12 suggests only modest potency, with a minimum inhibitory concentration (MIC) of 40 µM against *Escherichia coli* K12, which in those studies was equivalent to the MICs seen with another derivative, FK-13^53,54^, as well as wild-type LL-37. A follow-up study revealed similarly modest antimicrobial activity of KR-12 itself at >50 µM against *E. coli* (identical to our findings below)^48^, though results can vary with the methods used for susceptibility testing^45,48^. By contrast, most approved antibiotics active against *Enterobacterales* have Clinical & Laboratory Standards Institute (CLSI) minimum inhibitory concentration (MIC) cutoffs in the single-digit micromolar range^55^. Thus, although KR-12 is the shortest unit of LL-37 activity, the overall potency of this primary sequence is somewhat lower than is likely to be required in order to eventually derive clinically useful antimicrobials from LL-37 templates. Even FK-13, which is slightly more potent than KR-12 in most studies but still relatively non-toxic, is borderline by this measure^56^. We therefore sought to identify short derivatives of LL-37 with greater potency, from which we might then apply structure-activity approaches such as those that we have used to detoxify full-length LL-37 while conserving antimicrobial activity^30,31^.

To revisit the subject of shorter LL-37 derivatives comparable in size to approved antimicrobial natural products such as daptomycin and colistin, we made a series of fragments of LL-37 based on prior studies and assessed their relative activity against *E. coli* and *Pseudomonas aeruginosa* as well as their hemolytic activity against defibrinated sheep blood, where we take hemolysis as an *in vitro* surrogate of activity against mammalian cell membranes. Among these fragments listed in **Figure 1A**, all except (#2) LL-25, (#4) GD-23, (#7) FK-13, and full-length (#1) LL-37 demonstrated MICs > 50 µM against *E. coli* and *P. aeruginosa*, indicative of generally low levels of antimicrobial activity (**Figure 1B**). Consistent with their low levels of antimicrobial activity, most derivatives demonstrated little to no hemolytic activity in **Figure 1B**, the major exception being control wild-type (#1) LL-37. Thus, while many (#1) LL-37 derivatives show low toxicity to mammalian cell membranes, these generally also lack potent activity against gram-negative bacteria, limiting their utility as starting points for the development of new antibiotics based on the (#1) LL-37 scaffold. The major exceptions to this rule are (#2) LL-25 and (#4) GD-23, each of which recapitulates the separation of hemolytic and antimicrobial activities that we have observed separately in selected mutants of full-length (#1) LL-37^30^ as demonstrated by their higher **Hemo/MIC** and **Fold Δ** values (**Figure 1B**). This may reflect the absence of residues that we have previously found to be important both for oligomerization and for global membrane permeabilization activity (see also **Discussion**)^30,31^. Thus, the best leads from **Figure 1** incorporate residues corresponding to the LL-37 N-terminus while truncating before the end of the central 18-29 activity region in (#10) KR-12.

**Figure 1.**
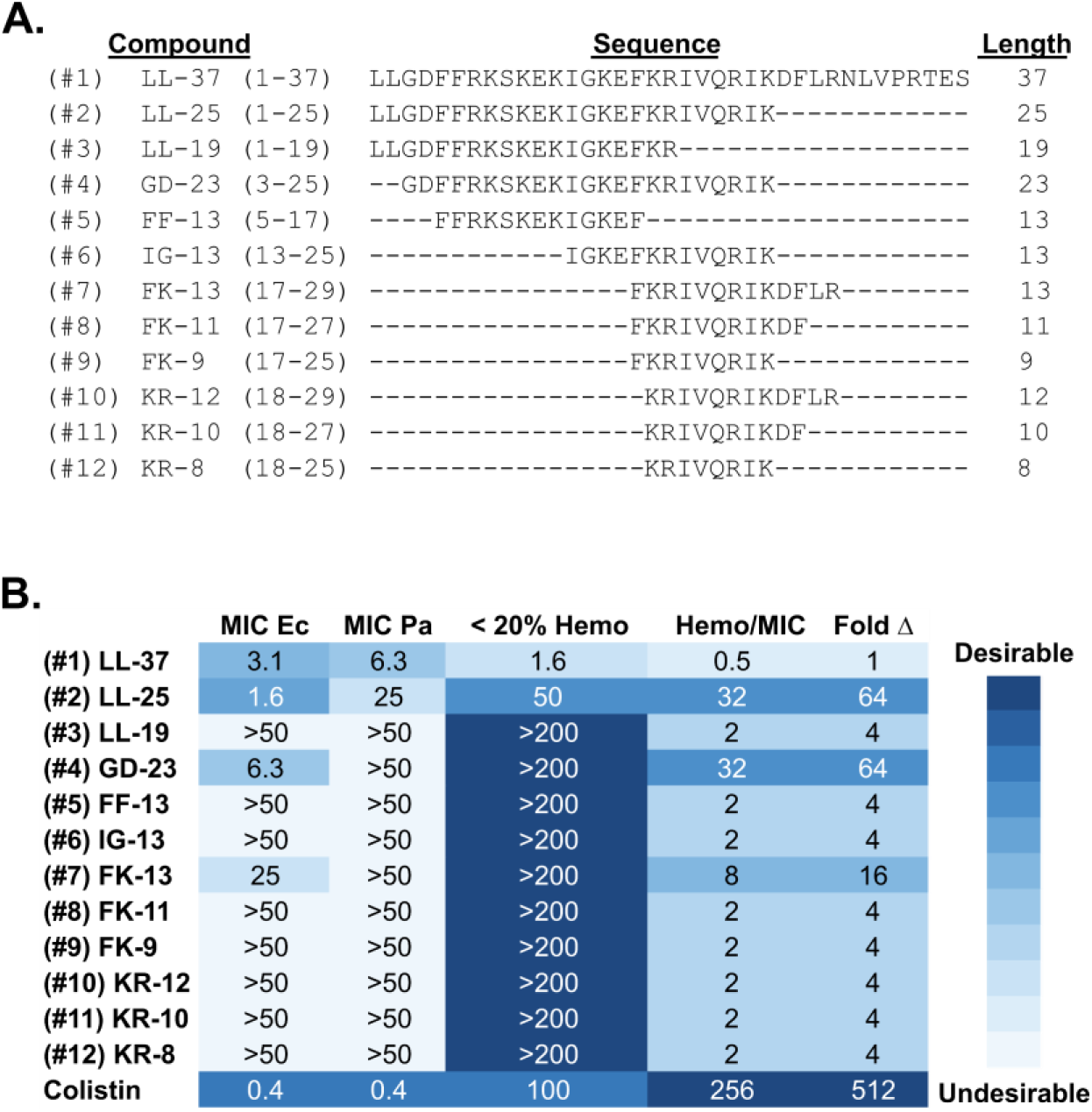
Limited activity of shorter derivatives of (#1) LL-37. **A.** Alignment summarizing the compounds tested. Each line reads from left-to-right with a compound number in parentheses, a compound name based on cathelicidin naming conventions, the numbering of each compound primary sequence relative to full-length LL-37, the primary sequence itself, and the sequence length. **B.** The table format presented here and in similar figures (**2B**, **4B, 5B**, **S89B**, **6B**) is a standardized description showing the MICs of each compound against *E. coli* (ATCC 25922) (**MIC Ec**) and *P. aeruginosa* (ATCC 27853) (**MIC Pa**) in the left two columns. The **< 20% Hemo** column shows the highest concentration of peptide in which hemolysis remains below a 20% threshold. Empiric derivation of these **MIC** and **< 20% Hemo** values is described in more detail in the **Experimental Section**. **Hemo/MIC** = (**< 20% Hemo**) / **MIC Ec,** where larger numbers represent higher concentrations required to induce hemolysis, lower concentrations required to kill *E. coli*, or both, and thus are considered desirable. **Fold Δ** = (**Hemo/MIC** of compound) / (**Hemo/MIC** of LL-37), which gives a measure of improvement in *in vitro* selectivity over the (#1) LL-37 template. Shading of the values within each column follows a convention by which desirable values (low **MIC Ec** or **MIC Pa** values, high **< 20% Hemo** thresholds, high **Hemo/MIC** ratios, or high **Fold Δ** ratios) are assigned darker shades, while less desirable values are assigned lighter shades. Data summarized represent the output of three independent susceptibility and hemolysis assays.

**Figure S1** provides a heatmap depiction of the data in **Figure 1B**; visual interpretation of these data are used to define the MIC cutoffs used in **Figure 1B** and similar tables throughout. **Figure S2** provides a histogram depiction of the same data including error bars; the hemolysis cutoff values used in **Figure 1B** and similar tables throughout derive from these analyses. **Figures S3-15** (antimicrobial susceptibility) and **S16-28** (hemolysis) show the kinetic growth and hemolysis curves associated with each experimental endpoint depicted in **Figure 1B**. In some peptides such as (#10) KR-12, bending of growth curves demonstrates modest inhibition below the MIC threshold applied in these studies (**Figure S12**).

### Mimicry of the (#1) LL-37 N-terminus improves the activity of short derivatives of (#1) LL-37 against gram-negative bacteria

Among the peptides in **Figure 1**, we noted that the most active candidates tended to include greater proportions of the (#1) LL-37 N-terminus. This is consistent with structure-activity mutagenesis that we have recently reported in full-length (#1) LL-37, in which grouped Ala mutagenesis of the (#1) LL-37 N-terminus ablates (#1) LL-37 activity against gram-negative bacteria despite being distant in linear sequence from the putative core activity region surrounding (#1) LL-37 residues 18-29^30^. One of these surface mutagenesis derivatives, in particular, has no mutations within the core 18-29 activity region, but is notable for containing two of the four Phe residues in (#1) LL-37 as a biphenyl motif. In wild-type (#1) LL-37, these four native Phe residues are thought to be primary mediators of membrane interaction, and conserved aromatic residues are also found among α defensins^36,57^. Finally, we noted that the inclusion of N-terminal Phe in (#7) FK-13 yielded improved activity over the (#10) KR-12 minimal unit (**Figure 1**). We therefore reasoned that transposition of the (#1) LL-37 N-terminal biphenyl motif, `FF[RK]`, to the N-terminus of (#10) KR-12, `--[KR]`, might yield a short derivative of (#1) LL-37 with improved antimicrobial activity.

To test this hypothesis, we made derivatives (#18) FF-14 and (#19) FF-14-RK, in which the N-terminal biphenyl motif of (#1) LL-37 was transposed onto the central 18-29 region as depicted in **Figure 2A**. As shown in **Figures 2B**, this results in 14-mer derivatives of (#1) LL-37 that more closely mimic the antimicrobial activities of wild-type LL-37, including potency >16-times greater than (#10) KR-12 (3.1 µM for (#18) FF-14 versus >50 µM for (#10) KR-12 against *E. coli*) and 4-times greater than (#7) FK-13, safely into low micromolar MIC ranges closer to those of many approved antibiotics. Although this comes at a cost of increased hemolysis relative to (#10) KR-12 / (#7) FK-13, hemolysis among these 14-mers remained 4-8 times less than (#1) LL-37 as in the **Fold Δ** column of **Figure 2B**; no substantial differences were noted based on the ordering of `[KR]` within this motif.

**Figure 2.**
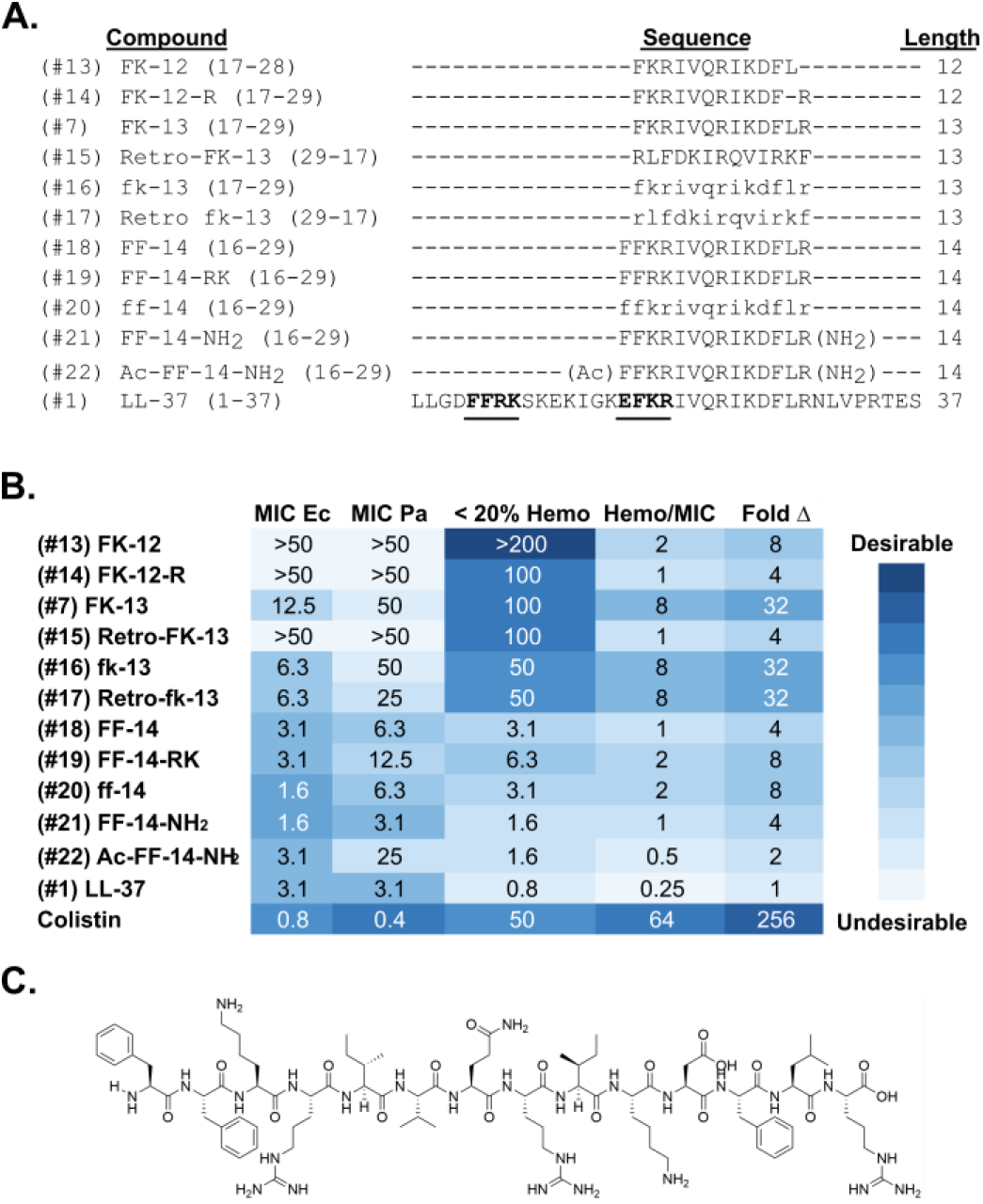
Mimicry of the (#1) LL-37 N-terminus yields a 14-mer with activity comparable to full-length (#1) LL-37. **A.** Alignment summarizing the compounds tested, as described under Figure 1. The FF[RK] region transposed to the central activity region on which (#10) KR-12 and (#7) FK-13 are built is shown in bold with highlighting. **B.** Compound properties including the MICs of each compound against *E. coli* (ATCC 25922) (**MIC Ec**) and *P. aeruginosa* (ATCC 27853) (**MIC Pa**) in the left two columns and the highest concentration of peptide in which hemolysis remains below a 20% threshold (**< 20% Hemo**). Calculated values include **Hemo/MIC** = (**< 20% Hemo**) / **MIC Ec,** where larger ratios indicate greater separation between the peptide concentrations required to induce hemolysis and the peptide concentrations required to kill *E. coli*. **Fold Δ** = (**Hemo/MIC** of compound) / (**Hemo/MIC** of LL-37), where larger ratios indicate greater degrees of improvement in hemolysis / killing separation in comparison with parent (#1) LL-37. Rationales for analysis and associated calculations are described in more detail under the **Experimental Section**. Shading of the values within each column follows a convention by which less desirable values are assigned lighter shades while more desirable values are assigned darker shades. Data summarized represent the output of three independent antimicrobial susceptibility assays and three independent hemolysis assays. **C.** Chemical structure of (#18) FF-14.

Mimicry of an (#1) LL-37 N-terminal sequence motif within (#18) FF-14 thus yields a scaffold with >16-fold improvements in activity, providing the basis for further modification of this scaffold toward the development of cathelicidin-derived antibiotics. The chemical structure of (#18) FF-14 is shown in **Figure 2C**.

### Compatibility of D-amino acids and C-terminal amidation with short cathelicidin templates

A number of peptidomimetic alterations have been demonstrated to stabilize and improve upon the native activities of HDP scaffolds such as D-amino acids and end modifications (*e.g.* ^38,53,58,59^ among examples with (#1) LL-37 and derivatives thereof). To determine whether similar alterations would prove compatible with (#18) FF-14, we made an expanded set of peptidomimetic derivatives in which (#18) FF-14 was either changed to an all-D configuration or amidated at the C-terminus. As shown in **Figure 2B**, (#20) enantiomeric ff-14 and (#21) C-amidated FF-14 each demonstrated similar-to-improved antimicrobial activity, particularly against *E. coli*, with similar shifts in fold-separation between hemolytic and antimicrobial activities in the **Hemo/MIC** column of **Figure 2B**. Synthetic leads based on the FF-14 template may therefore benefit from the inclusion of stabilizing mutations such as the incorporation of D-amino acids and C-terminal amidation. Of note, activity of FK-13 also improved in the all-D format (#16), with retention of the same **Hemo/MIC** ratio in **Figure 2B**. Although (#17) retro-inverso fk-13 also retained activity equivalent to (#16) all-D fk-13, (#15) retro-FK-13 lost all apparent antimicrobial activity as assessed by its MIC, though curve bending in **Figure S34** may be consistent with a weak inhibitory effect below the limits of MIC detection.

Thus, FF-14 is compatible with standard stabilization approaches such as D-amino acids and C-amidation. **Figure S29** provides heatmap depiction of the data in **Figure 2B**, while **Figure S30** provides a histogram depiction of the data in **Figure 2B** including error bars. **Figures S31-43** (antimicrobial susceptibility) and **S44-56** (hemolysis) show the kinetic growth and hemolysis curves associated with each experimental endpoint depicted in **Figure 2B**.

### Perfluoroaryl stapling does not reliably separate antimicrobial and hemolytic cathelicidin activities

Despite the improved activity of (#18) FF-14 and derivatives thereof against gram-negative bacteria, we noted substantial ongoing hemolytic activity that would likely preclude the use of these cathelicidin derivatives as lead therapeutics without additional detoxifying modifications. In related work, we have previously identified I24 and L28 as the primary determinants of mammalian cell toxicity in full-length (#1) LL-37^30^. Consistent with this finding, (#2) LL-25 and (#4) GD-23 in **Figure 1** omit L28 and demonstrate decreased hemolytic activity. Moreover, in separate mutagenesis of the (#18) FF-14 template, we have found that substitution of the I24 equivalent residue with aminoisobutyric acid (Aib) may contribute to diminished hemolytic activity^31^. Because equivalents of I24 and L28 occur within the (#10) KR-12 core (I9 and L13 in (#18) FF-14 numbering), we hypothesized that stapling strategies, particularly those involving I9 and L13, might result in similar ablation of toxicity within (#18) FF-14 while preserving antimicrobial activity.

To test this hypothesis, we made a comprehensive set of *i*,*i*+3, *i*,*i*+4, and *i*,*i*+7 perfluoroaryl stapled versions of the (#18) FF-14 hydrophobic surface as well as selected perfluoroaryl stapled versions of (#7) FK-13^20^. The general scheme for stapling at Cys residues under this methodology is shown in **Figure 3**, and **Figure 4A** shows Cys positioning within the primary (#7) FK-13 and (#18) FF-14 sequences to permit perfluoroarylation. As shown in **Figure 4B**, perfluoroaryl stapling of (#7) FK-13 and (#18) FF-14 derivatives generally proved neutral-to-detrimental to overall antimicrobial activity, and in most cases failed to substantially improve upon the characteristics of the associated primary sequence alone. In particular, compounds such as (#30), (#32), and (#33) that modify the position corresponding to critical residue I24 in (#1) LL-37^30^ do not separate antimicrobial activity from hemolytic activity. Similarly, compounds (#23) and (#26) with stapling at the position corresponding to critical residue L28^30^ and compound (#31) with stapling at the positions corresponding to both I24 and L28 also fail to achieve the separation of function seen in full-length LL-37. Pilot experiments with lactam and Pd arylation stapling chemistries did not improve upon these results (data not shown). Because separation of function in full-length (#1) LL-37 depends, in part, on decreasing hydrophobic character in the (#1) LL-37 dimerization interface, it may be that the perfluoroaryl staples used here are too hydrophobic to permit the separation of function that we have achieved with primary sequence modifications in full-length (#1) LL-37 and (#18) FF-14 in other work^30,31^. This hydrophobicity effect may also be reflected in the hemolysis kinetics among many of the hemolytically active stapled derivatives, which were often more rapid and complete than those seen with (#1) LL-37 and, to a lesser extent, (#18) FF-14 (**Figures S59-73**).

**Figure 3.**
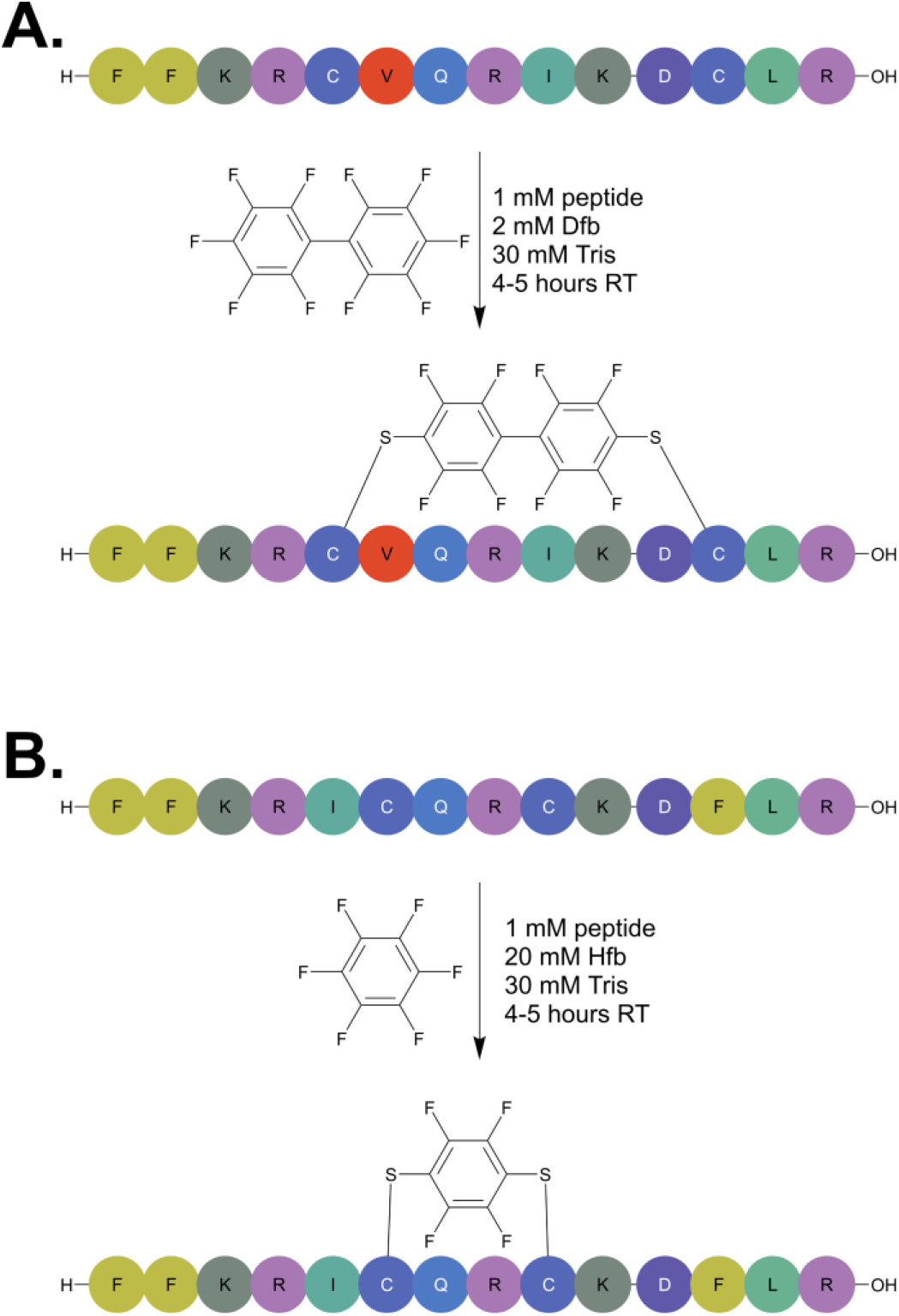
General scheme for cysteine perfluoroarylation. **A.** Schematic showing conditions for stapling of *i,i+7* positions with decafluorobiphenyl (Dfb). **B.** Schematic showing conditions for stapling of *i,i+3* or *i,i+4* positions with hexafluorobenzene (Hfb).

**Figure 4.**
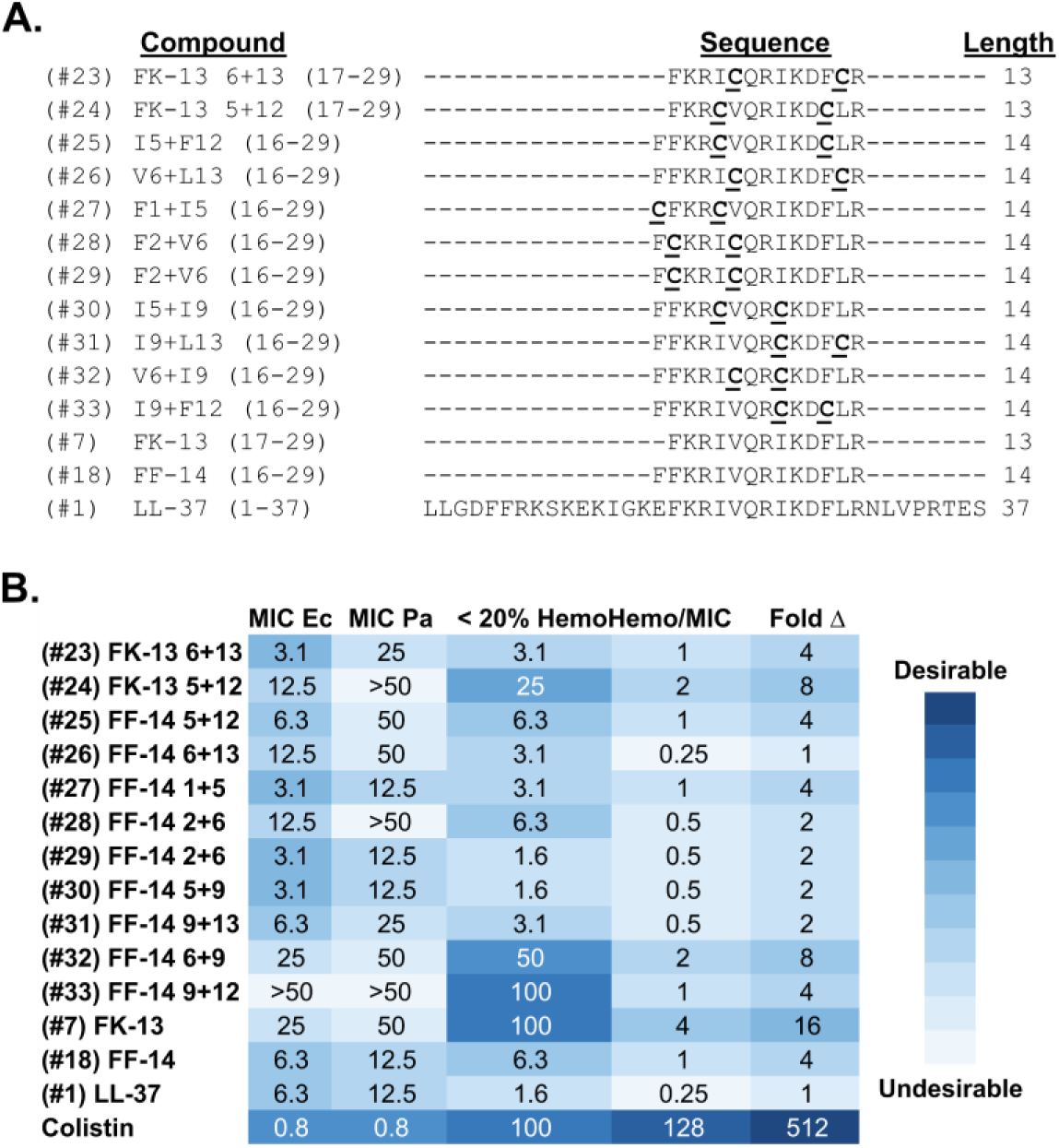
Antimicrobial activity and hemolysis remain correlated in stapled short cathelicidin derivatives. **A.** Alignment summarizing the compounds tested, in which Cys placements to allow for perfluoroarylation within the (#7) FK-13 or (#18) FF-14 primary sequences are bolded and underlined. Numbering of stapling positions is referenced to the primary sequence of (#18) FF-14 in the conventional names given, the first residue of which corresponds to the 16^th^ residue of (#1) LL-37. **B.** Compound properties including the MICs of each compound against *E. coli* (ATCC 25922) (**MIC Ec**) and *P. aeruginosa* (ATCC 27853) (**MIC Pa**) in the left two columns and the highest concentration of peptide in which hemolysis remains below a 20% threshold (**< 20% Hemo**). Calculated values include **Hemo/MIC** = (**< 20% Hemo**) / **MIC Ec**, where larger numbers indicate greater selectivity for bacterial over mammalian cells. **Fold Δ** = (**Hemo/MIC** of compound) / (**Hemo/MIC** of LL-37), where larger numbers indicate improvements in selectivity in comparison to the (#1) LL-37 parent peptide. Rationales for analysis strategies are discussed further under the **Experimental Section**. Shading of the values within each column follows a convention by which less desirable values are assigned lighter shades while more desirable values are assigned darker shades. Data summarized represent the output of three independent antimicrobial susceptibility assays and three independent hemolysis assays.

Because hydrophobicity is common to many stapling strategies, and because our separate efforts focused on primary sequence changes met with more success, we did not further pursue alternative stapling strategies here.

In summary, although stapling might prove useful in the context of primary human cathelicidin sequences with lower intrinsic activity such as (#10) KR-12 as previously reported^42^, or perhaps when using chemistries with more hydrophilic character, our data favor changes in primary sequence as a more productive approach to modulating human cathelicidin toxicity^30,31^. **Figure S57** provides a heatmap depiction of the data in **Figure 4**, while **Figure S58** provides a histogram depiction of the data in **Figure 4** including error bars. **Figures S59-73** (antimicrobial susceptibility) and **S74-88** (hemolysis) show the kinetic growth and hemolysis curves associated with each experimental endpoint depicted in **Figure 4**.

### N-lipidation modulates FF-14 activity

A lipid tail is important for the activity of certain antibiotics such as colistin and daptomycin. Along these lines, prior studies have indicated that some lipid tails can improve the activity of the (#10) KR-12 template^48,60,61^, though often at a cost of toxicity. Moreover, the potential impact of N-lipidation is implied by the hydrophobic character of the (#1) LL-37 N-terminus, where two Leu residues precede the native biphenyl motif – L_1_LGDFF_6_. We therefore reasoned that N-lipidation might also affect the activity of (#18) FF-14 or (#7) FK-13 and proceeded to test the effects of lipid moieties of several structural types on the antimicrobial and hemolytic activities in (#18) FF-14, with an emphasis on lipid moieties found in approved antimicrobials (*e.g.*, 5-dimethylamino pentanoic acid, 10-methylundecanoic acid, and chlorobiphenyl in lipoglycopeptides) or in promising experimental scaffolds (*e.g.*, biphenyl moieties^62^). Of note, these are largely distinct from those previously reported in the context of (#10) KR-12^48,60,61^. Moieties used in in the context of (#18) FF-14 in **Figure 5** are shown in **Figure 5A**, while those used in the context of (#7) FK-13 in **Figure S89** are shown in **Figure S89A**.

**Figure 5.**
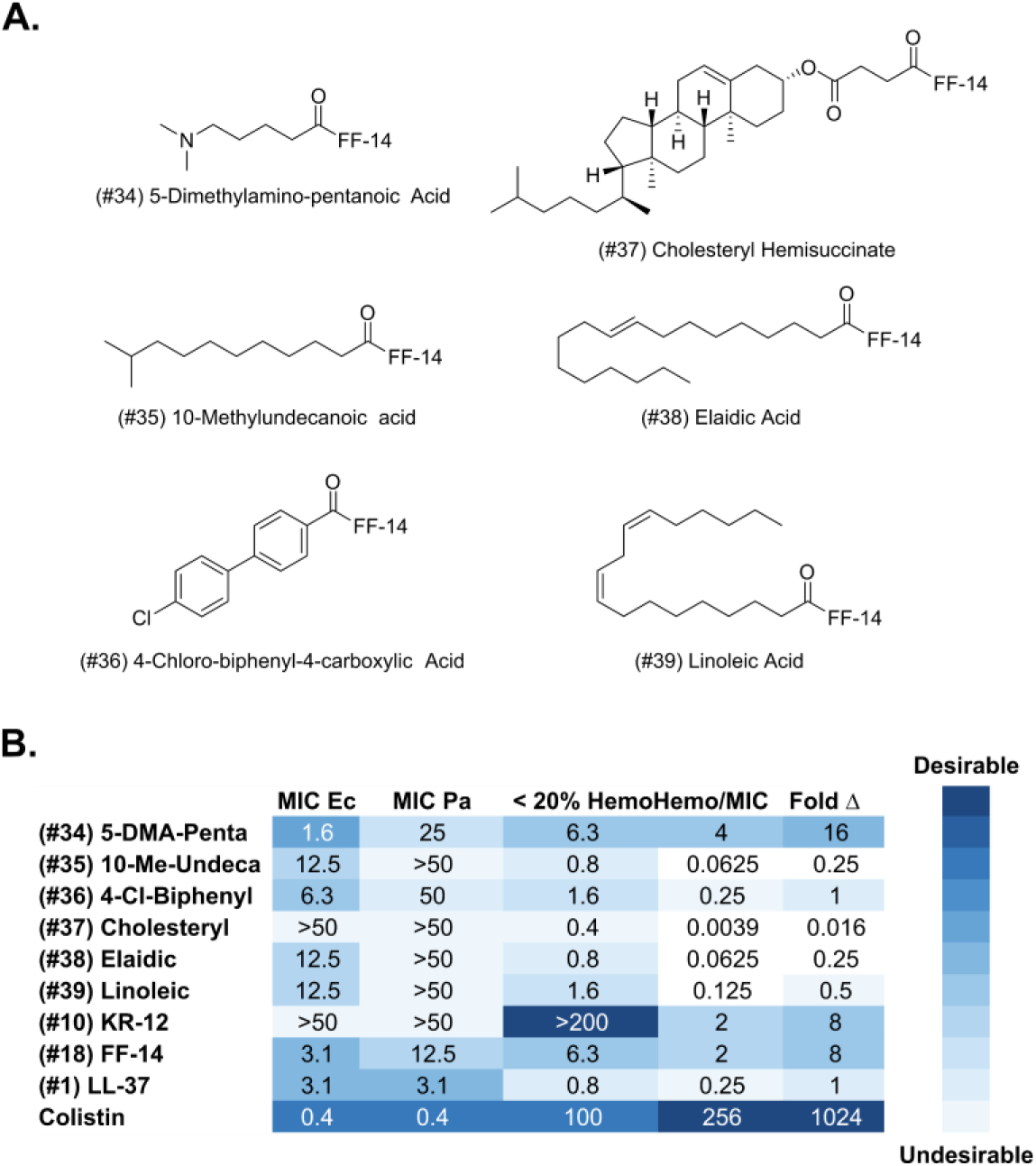
Modulation of (#18) FF-14 activities via N-lipidation. **A.** Structures are shown of the lipid tails appended to the N-terminus of (#18) FF-14. 5-DMA-Penta, 10-Me-Undeca, and 4-Cl-Biphenyl derive from lipoglycopeptide antibiotics, cholesteryl is taken as a sterol representative, and elaidic and linoleic acid represent longer chain mono- and poly-unsaturated tails, respectively. **B.** Compound properties including the MICs of each compound against *E. coli* (ATCC 25922) (**MIC Ec**) and *P. aeruginosa* (ATCC 27853) (**MIC Pa**) in the left two columns and the highest concentration of peptide in which hemolysis remains below a 20% threshold (**< 20% Hemo**). Calculated values include **Hemo/MIC** = (**< 20% Hemo**) / **MIC Ec** and **Fold Δ** = (**Hemo/MIC** of compound) / (**Hemo/MIC** of LL-37), where larger numbers indicate peptides with improved antimicrobial activity, cell type selectivity, or both. Rationales for analysis strategies are discussed further under the **Experimental Section**. Shading of the values within each column follows a convention by which less desirable values are assigned lighter shades while more desirable values are assigned darker shades. Data summarized represent the output of three independent antimicrobial susceptibility assays and three independent hemolysis assays.

As shown in **Figure 5B**, the addition of multiple moieties to the FF-14 N-terminus results in improved antimicrobial activity, though only 5-dimethylamino-pentanoic acid yielded neutral-to-improved potency by halving the (#18) FF-14 **MIC Ec** while maintaining the same **< 20% Hemo** toxicity threshold. Despite its activity against *E. coli*, this and most lipidated derivatives demonstrated diminished activity against *P. aeruginosa* as shown in the **Figure 5B MIC Pa** column. Similar trends were seen with lipid tails tested in the context of (#7) FK-13 on lower purity compounds, with the notable exception of octanoic acid in compound (#43) (**Figure S89**).

**Figures 5** and **S89** thus suggest that the additive value of N-lipidation within the (#18) FF-14 template is largely limited to 5-dimethylamino pentanoic acid or octanoic acid, though future work on placement elsewhere within cathelicidin templates may prove fruitful. In particular, appending further phenyl groups to the N-terminus beyond the biphenyl motif itself as in (#36) 4-Cl-Bph-FF-14 did not yield additional increases in activity. As in the case of stapling, our data support the principle that the most efficient route to improved cathelicidin properties runs through primary sequence modifications. **Figures S90-91** provides a heatmap depiction of the data in **Figures 5** and **S89**, respectively, while **Figures 92-93** provide histogram depictions of the data in **Figures 5** and **S89,** including error bars. **Figures S94-109** (antimicrobial susceptibility) and **S110-125** (hemolysis) show the kinetic growth and hemolysis curves associated with each experimental endpoint depicted in **Figure 5** and **Figure S89**. As in the case of our stapled peptides, hemolysis curves in this series of experiments were notable for more rapid and complete hemolysis among some lipidated derivatives, which may reflect increased hydrophobicity.

### The biphenyl motif is a portable enhancer of antimicrobial activity in longer cathelicidin-derived sequences

Based on the results in **Figure 5** and **Figure S89**, we elected to further examine three candidate N-lipidation moieties – octanoic acid (Oct), 5-dimethylamino pentanoic acid (DMA), and the quaternary amine equivalent of DMA, δ valerobetaine (DVB), the structures of which are shown in **Figure 6A**. We further expanded the application of these moieties to the longer cathelicidin derivatives (#2) LL-25 and (#4) GD-23 noted in **Figure 1** that partially separate antimicrobial and hemolytic activity. Finally, to test the impact of recapitulating the (#18) FF-14 sequence within these longer cathelicidin scaffolds, we included derivatives of (#2) LL-25 and (#4) GD-23 with duplicate biphenyl motifs – the native sequence at the N-terminus, and the FF-14 sequence within the central activity region. Test compounds associated with these experiments are shown in **Figure 6B**.

**Figure 6.**
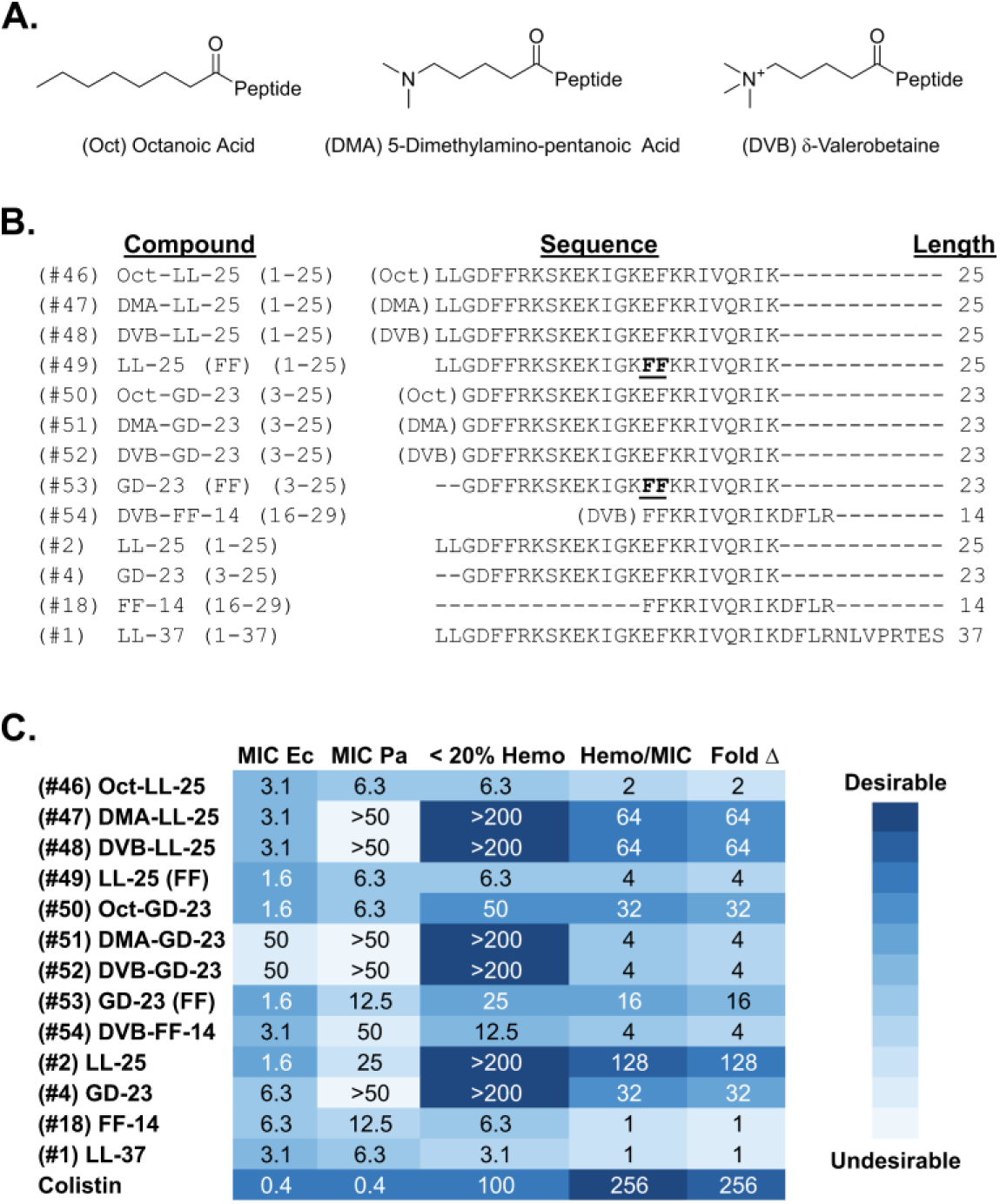
Portability of the N-terminal biphenyl motif for the enhancement of antimicrobial activity. **A.** Structures are shown of the lipid tails appended to the N-terminus of (#2) LL-25, (#4) GD-23, or (#18) FF-14 based on the experiments in Figures 5 and **S89**. **B.** Primary sequences of the compounds tested in Figure 6. **C.** Compound properties including the MICs of each compound against *E. coli* (ATCC 25922) (**MIC Ec**) and *P. aeruginosa* (ATCC 27853) (**MIC Pa**) in the left two columns and the highest concentration of peptide in which hemolysis remains below a 20% threshold (**< 20% Hemo**). Calculated values include **Hemo/MIC** = (**< 20% Hemo**) / **MIC Ec** and **Fold Δ** = (**Hemo/MIC** of compound) / (**Hemo/MIC** of LL-37), where larger numbers are preferable as described in **Methods**. Shading of the values within each column follows a convention by which less desirable values are assigned lighter shades while more desirable values are assigned darker shades. Data summarized represent the output of three independent antimicrobial susceptibility assays and three independent hemolysis assays.

As shown in the **MIC Ec** column of **Figure 6C**, most N-lipidated derivatives of (#2) LL-25 and (#4) GD-23 demonstrated even-to-diminished antimicrobial activity compared to the wild-type primary sequence. Activity of N-lipidated derivatives against *P. aeruginosa* was entirely ablated for these sequences in the cases of DMA and DVB. (#50) Oct-GD-23 demonstrated improved antimicrobial activity, though at some cost of hemolytic activity. Consistent with the effects of DMA in **Figure 5**, however, DVB did yield four-fold separation between antimicrobial and hemolytic activity in the context of FF-14 (#54). Among the modifications tested, however, the greatest impact on antimicrobial activity was observed when including a second biphenyl motif to recapitulate the (#18) FF-14 sequence within the longer (#2) LL-25 and (#4) GD-23 scaffolds. This was particularly true for activity against *P. aeruginosa*, where placement of the biphenyl pharmacophore within (#2) LL-25 and (#4) GD-23 improved MICs from 25 µM to 6.3 µM in (#49) LL-25 (FF) and from >50 µM to 12.5 µM in (#53) GD-23 (FF).

With the possible exceptions of (#34) DMA or (#54) DVB in the context of FF-14, lipidation generally did not improve upon the characteristics of the native sequence in shorter cathelicidin derivatives such as (#2) LL-25 or (#4) GD-23. Duplication of the biphenyl motif to recapitulate the sequence of (#18) FF-14 within the longer scaffolds (#2) LL-25 and (#4) GD-23, however, did improve antimicrobial activity against *P. aeruginosa* while maintaining or improving upon activity against *E. coli*, albeit at a cost of increased hemolytic activity.

Thus, the (#1) LL-37 N-terminal biphenyl motif is portable also to longer cathelicidin scaffolds beyond (#18) FF-14. Moreover, our results again demonstrate that for cathelicidin templates that are already highly active, the expected contributions of peptidomimetic modifications to the primary sequence itself are generally marginal. Efforts to modulate cathelicidin antimicrobial activity and toxicity may more efficiently address these goals by focusing on primary sequence modifications. For example, it may be possible to merge activity-enhancing modifications such as the portable biphenyl motif described here with orthogonal toxicity-reducing modifications such as modulation of the cathelicidin dimerization interface to achieve improved cell type selectivity^30,31^.

**Figure S126** renders the primary data in **Figure 6** as heatmaps, while **Figure S127** provides a histogram depiction of the same data including error bars. **Figures S128-141** (antimicrobial susceptibility) and **S142-155** (hemolysis) show the kinetic growth and hemolysis curves associated with each experimental endpoint depicted in **Figure 6**.

### The (#18) FF-14 biphenyl motif contributes to increased outer and inner membrane permeabilization in gram-negative bacteria

To better characterize the mechanism underlying the contributions of the biphenyl motif to (#18) FF-14 antimicrobial activity, we carried out kinetic dual membrane permeabilization assays in *E. coli* and *P. aeruginosa* comparing the relative permeabilization of the outer (OM) and inner (IM) gram-negative membranes as measured by N-phenyl-naphthylamine (NPN) and propidium iodide (PI) staining over time in the presence of a range of concentrations of each of (#10) KR-12, (#7) FK-13, (#18) FF-14, and (#1) LL-37. This assay allows for the combination of both kinetics and raw levels of permeabilization into a single composite metric of activity, as further explained in the **Experimental Section**. As shown in **Figure 7A-B**, permeabilization was modest in the (#10) KR-12 derivative and increased sequentially through the addition of one ((#7) FK-13) and then two ((#18) FF-14) Phe residues to the N-terminus for both the OM and IM in both *E. coli* and *P. aeruginosa*, with (#18) FF-14 permeabilization activity approaching that of (1) LL-37 itself. Thus, mimicry of the LL-37 N-terminus as in the placement of the biphenyl motif in (#18) FF-14 results in improved membrane permeabilization that more closely recapitulates that observed with (#1) LL-37.

**Figure 7.**
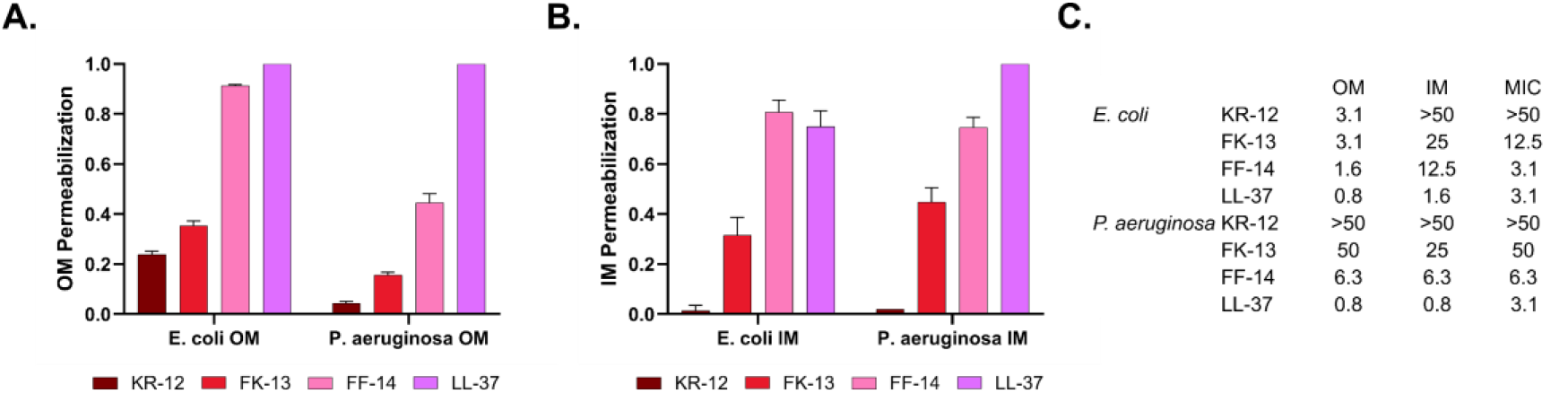
Relative outer and inner membrane permeabilization of short cathelicidin derivatives. Dual membrane permeabilization assays were completed by incubating *E. coli* (ATCC 25922) or *P. aeruginosa* (ATCC 27853) in the presence of N-phenyl-naphthylamine as a marker of outer membrane (OM) permeabilization (**A**) and propidium iodide as a marker of inner membrane (IM) permeabilization (**B**) in the presence of varying concentrations of each indicated peptide. Permeabilization was then quantified over the course of 240 minutes relative to (#1) LL-37. The single-point permeabilization index presented represents the product of the peak permeabilization signal relative to (#1) LL-37 multiplied by the proportion of the 240-minute incubation required to surpass 80% of the maximum signal observed for each peptide. Full kinetic curves associated with each condition are shown in **Figures S156-159. C.** Comparison between typical peptide MICs in this study and peptide concentrations associated with observable OM and IM permeabilization. Threshold values presented are the lowest concentrations at which a given peptide achieves at least 10% of the peak (#1) LL-37 permeabilization signal. This threshold is chosen based on the observation that the peptide concentration at which LL-37 achieves at least 10% of its maximum permeabilization signal is usually at or below the LL-37 MIC. The 10% threshold thus serves as a marker of permeabilization levels that in the control (#1) LL-37 condition are associated with killing. Data summarized represent the output of three independent experiments.

While the single-point composite measure of peak permeabilization and kinetics used in **Figures 7A-B** gives a sense of the maximum permeabilization potential of each peptide, we were also interested in assessing permeabilization behavior at the functionally relevant limits of assay detection. Because the IM permeabilization of (#1) LL-37 consistently achieves at least 10% of its maximum value at concentrations at or below the corresponding MIC, we chose to define this 10% threshold as an internally calibrated minimum threshold of functionally relevant permeabilization.

On comparing the 10% threshold values for OM and IM permeabilization for each peptide with their corresponding MICs as in **Figure 7C**, we noted that, while all OM thresholds fall at or below the corresponding typical MIC, the ratio between the 10% threshold and the MIC is 8-fold greater for (#18) FF-14 than for (#1) LL-37 in *E. coli*. That is, while (#18) FF-14 and (#1) LL-37 both usually display similar *E. coli* MICs in our studies, the FF-14 IM threshold value is 12.5 µM, while that for LL-37 is 1.6 µM. We have observed even larger gaps among other (#1) LL-37 derivatives in *E. coli* in other studies^63^. Similarly, the 10% threshold values for IM permeabilization observed with (#18) FF-14 are 8-fold higher than those observed with (#1) LL-37 in both *E. coli* (12.5 vs. 1.6 µM) and *P. aeruginosa* (6.3 versus 0.8 µM), again despite similar capacities for killing as measured by their MICs. Thus, despite the correlation between increased permeabilization in (#18) FF-14 and lower MICs compared with (#10) KR-12 or (#7) FK-13, there is a discrepancy between detectable permeabilization and killing in (#18) FF-14 when compared with (#1) LL-37. This is also visually apparent in the kinetic permeabilization curves presented in **Figures S156-157**, where FF-14 and LL-37 are both highly efficient at permeabilizing the OM, while FF-14 demonstrates only modest IM permeabilization. This may imply mechanisms of cathelicidin killing of gram-negative bacteria more complex than gross membrane perturbation, which is a subject of ongoing research in our group beyond the scope of the current study^63^. Kinetic curves associated with each condition are shown in **Figures S156-S160.**

### Contributions of the biphenyl motif to helical structure

It is commonly thought that cathelicidin-derived antimicrobials must take on a helical structure in order to disrupt membranes^36,56,64,65^, in which case increased helical propensity may be a driver for enhanced membrane permeabilization and/or functional activity in cathelicidin derivatives carrying the portable biphenyl motif. To ascertain the relative helical propensity of key peptides developed in the course of these studies, we dissolved (#10) KR-12, (#7) FK-13, (#18) FF-14 and (#1) LL-37 as well as (#4) LL-25, (#49) LL-25 (FF), (#4) GD-23, and (#53) GD-23 (FF) in water with 20% trifluoroethanol (TFE) and collected their circular dichroism spectra at 25 °C and 37 °C.

As shown in **Figures 8A-D**, sequential addition of Phe residues to the 18-29 central activity region of (#10) KR-12 as in (#7) FK-13 and then (#18) FF-14 resulted in the associated peptides going from disordered ((#10) KR-12) structures to moderate helices ((#18) FF-14), where helical content is estimated using BeStSel^66^. In the context of longer sequences such as (#2) LL-25 and (#4) GD-23, however, gains in estimated helical content were lower with addition of the biphenyl motif in (#49) LL-25 (FF) and (#53) GD-23 (FF) (**Figures 8E-H**). Thus, the N-terminal biphenyl motif may help to support helical structure in shorter peptides with less inherent helical propensity than longer scaffolds such as (#2) LL-25 and (#4) GD-23.

**Figure 8.**
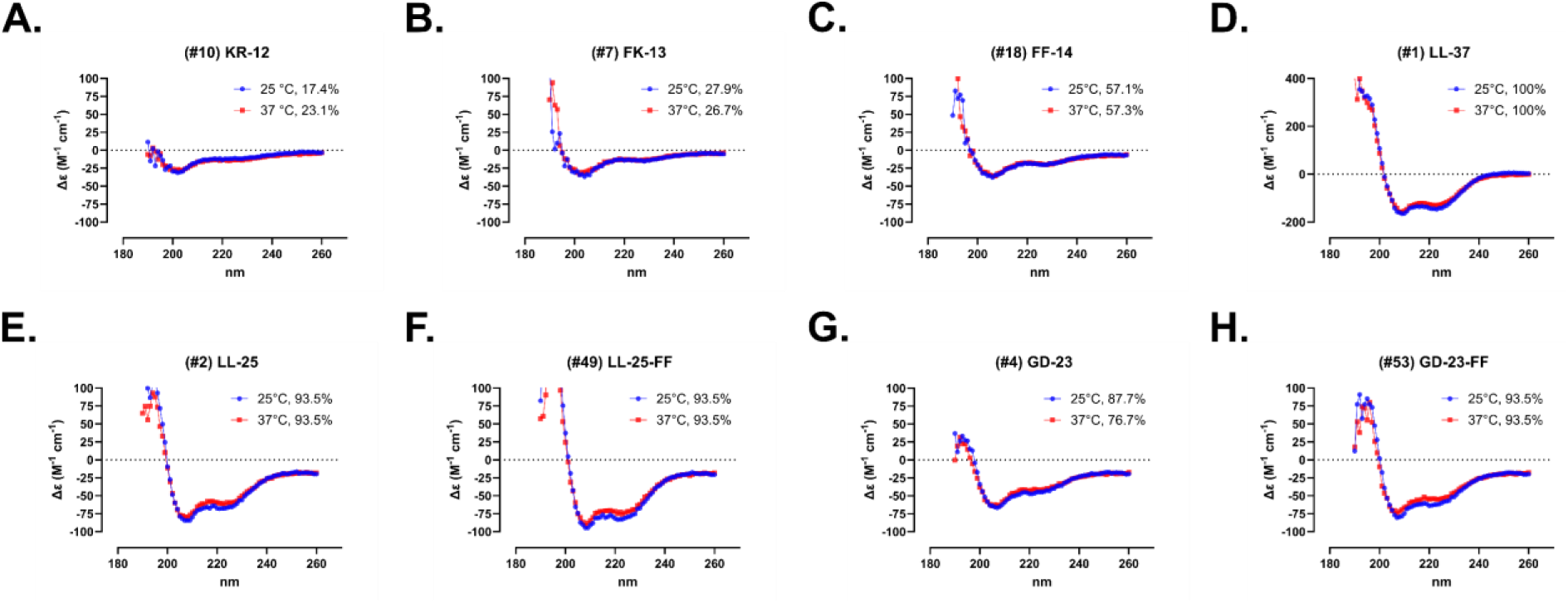
Effect of a portable biphenyl motif on peptide secondary structure. The indicated peptides in panels **A-H** were dissolved to 0.5 mg/mL in water + 20% TFE followed by collection of CD spectra from 260-190 nm at 25 °C and 37 °C. Helical content was estimated using BeStSel and is provided with the figure legends in each panel.

### Serum recovery and stability vary among cathelicidin derivatives

Although the immediate goal of the work described here was to identify features in short cathelicidins that promote enhanced activity against gram-negative bacteria, a long-term goal is to adapt such findings to the development of antimicrobial therapeutics. To determine whether the biphenyl motif might affect stability to human proteases, we carried out serum stability studies on (#10) KR-12, (#18) FF-14, (#4) LL-25, (#49) LL-25 (FF), and (#1) LL-37.

To do this, we first sought to establish a baseline for cathelicidin recovery from serum. For example, it has been previously reported that (#1) LL-37 binds to serum components such as apolipoprotein A-I, with some authors describing immediate loss of (#1) LL-37 on addition to serum, suggesting rapid degradation and/or sequestration with serum components during serum precipitation^67,68^. To distinguish these possibilities, we hypothesized that it might be possible to improve the recovery of (#1) LL-37 and other cathelicidins through the addition of a heating step to disrupt and denature potential serum interaction surfaces during precipitation. As shown in **Figure 9A**, heating to 70 °C during the first 10 minutes of a 20-minute trichloroacetic acid (TCA) precipitation resulted in marked increases in (#1) LL-37 recovery, suggesting possible heat-sensitive interactions between LL-37 and serum components as a contributor to the rapid loss of LL-37 signal in serum stability assays. In contrast, heat was either neutral or detrimental to the recovery of the other cathelicidins in our matrix, such that optimal recovery of a given cathelicidin may require the use of peptide-specific conditions.

**Figure 9.**
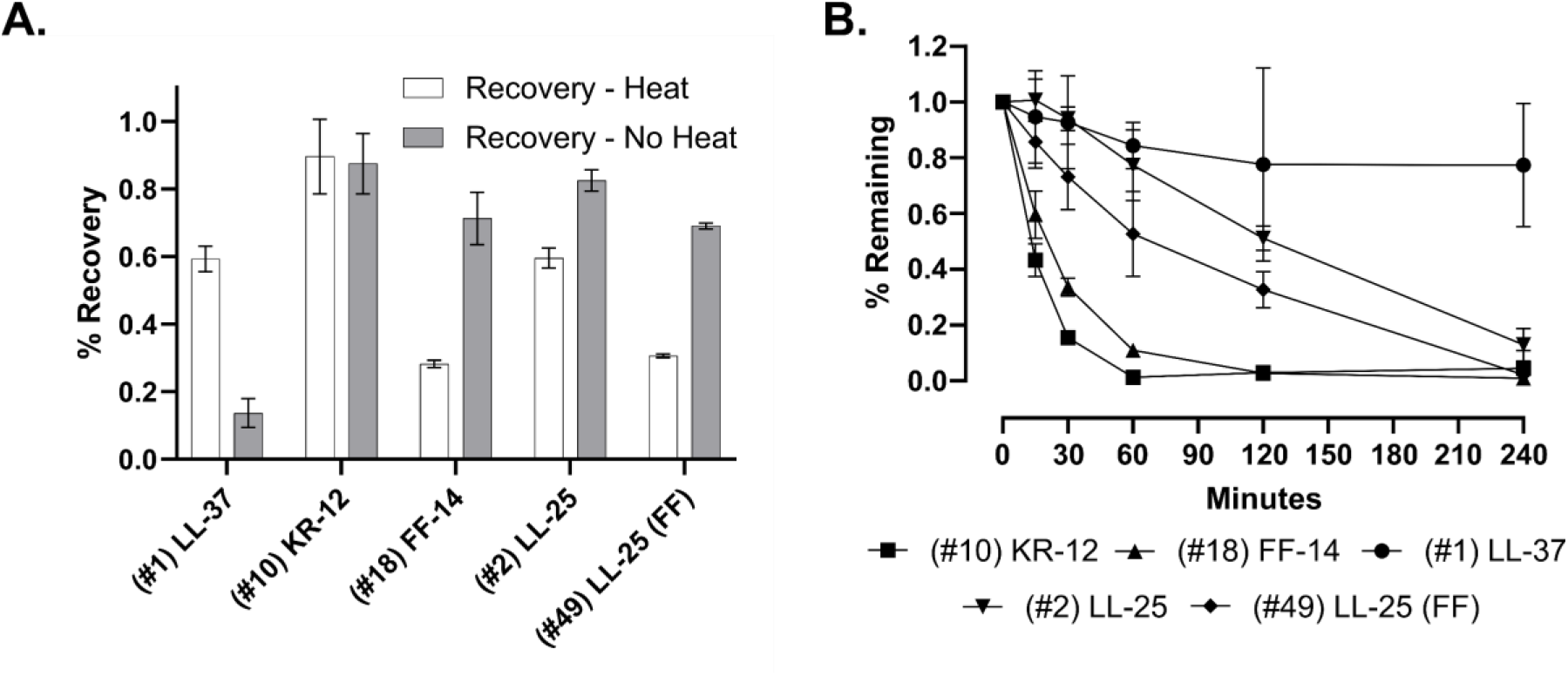
Serum recovery and stability of cathelicidin derivatives. **A.** The indicated peptides were added to both PBS and PBS + 10% human serum conditions and then immediately precipitated with 1% TCA followed by incubation either at 70 °C for 10 minutes followed by room temperature for 10 minutes (Heat) or at room temperature for 20 minutes (No Heat). Peptide recovery was then assayed by LCMS, with recovery calculated as the ratio of the extracted ion chromatogram (EIC) areas detected from the PBS + 10% Serum condition to the same EIC areas detected from the PBS condition – (PBS + 10% Serum) / PBS. **B.** Serum stability of the indicated peptides was assessed over 240 minutes by dividing the recovery ratio at 0 minutes (set to 1) by the same recovery ratios collected at each time point. 0 minute recovery ratios in **A** were collected under both Heat and No Heat conditions, while stability measurements in **B** were processed solely under Heat precipitation conditions. Data shown are the mean and standard deviation of three independent experiments.

With the goal of better characterizing (#1) LL-37 serum stability while also ascertaining the effects of the biphenyl pharmacophore on short cathelicidin stability, we next proceeded to incubate each of the above peptides over 240 minutes at 37 °C with subsequent precipitation and recovery using our (#1) LL-37-optimized heat disruption protocol. As expected, both (#10) KR-12 and (#18) FF-14 were rapidly degraded with half-lives of approximately 15 minutes in PBS supplemented with 10% human serum (**Figure 9**). In contrast, the longer and more structured (see **Figure 8**) peptides (#4) LL-25 and (#49) LL-25 (FF) demonstrated intermediate stability with half-lives on the order of 120 minutes. Surprisingly, full-length (#1) LL-37 was the most stable peptide in this group, with approximately 80% remaining at 240 minutes. Thus, within the limitations of the modest number of peptides tested, we observed a correlation between cathelicidin serum stability and peptide length and helical propensity (**Figure 8**), but no apparent effect of the biphenyl motif on serum stability.

## Discussion

In this study, we build on our prior structure-activity work in full-length (#1) LL-37 to demonstrate that transposition of an N-terminal biphenyl motif onto the central LL-37 (18-29) activity region improves activity against gram-negative bacteria >16-fold. This is consistent with structural data from the same landmark study that first described LL-37 (18-29), also known as (#10) KR-12, which indicated that the four Phe residues of (#1) LL-37 are critical for interaction with model membranes^36^. The resultant short derivative, (#18) FF-14, better recapitulates the gram-negative and hemolytic activities of (#1) LL-37, which may make this a useful lead scaffold from which to build out peptidomimetic derivatives. Further work identifies stabilizing interventions such as the use of D-amino acids and C-amidation as well as selected N-lipidation moieties as potentially useful activity modulation strategies in the more active (#18) FF-14, as well as the portability of the antimicrobial biphenyl motif to longer cathelicidin scaffolds.

With respect to mechanism, the data presented in **Figure 7** demonstrate increased OM and IM permeabilization associated with the biphenyl pharmacophore, which may contribute to increased antimicrobial activity in peptides containing this motif. We also note, however, that there is an 8-fold discrepancy between the lower limits of detectable IM permeabilization between (#18) FF-14 at 12.5 µM and (#1) LL-37 at 1.6 µM in **Figure 7C** despite their similar MICs. It is therefore possible that mechanisms underpinning the observed functional readouts and antimicrobial activity more broadly may be more complex than membrane disruption alone, a hypothesis that we have investigated in more detail in related work^30,31,63^.

In addition to the importance of the (#1) LL-37 N-terminus for activity against gram-negative bacteria, our data suggest that primary sequence drives overall potency and toxicity in cathelicidin derivatives. Stabilizing modifications may prove necessary in downstream therapeutics, but neither these nor N-lipidation result in large changes in (#18) FF-14 activity. It is possible, however, that chemical modification may be more important for the modulation of primary sequences with lower intrinsic activity such as (#10) KR-12. Of note, while the studies here focus on the effects of N-terminal lipidation on the activity profiles of short cathelicidin derivatives, a separate question of interest is whether lipidation moieties placed at activity-neutral sites in these templates might yield favorable pharmacokinetic properties as in GLP-1 agonists. We have separately carried out a comprehensive mutagenesis to identify structure-promoting primary sequence modifications in FF-14, which may help to address this question^31^. Independent of pharmacokinetic engineering strategies based on the use of albumin binding moieties, however, our serum stability studies in **Figure 9** suggest that longer cathelicidins may already have natural interactions with serum components that may be leveraged to influence half-life and pharmacokinetics. Better understanding and exploiting these heat-labile interactions between cathelicidins and serum components to yield more drug-like cathelicidin derivatives is a focus of our development efforts moving forward.

Although the main focus of this study was on developing short derivatives like (#10) KR-12 with improved antimicrobial activity, a surprising finding shown here is that derivatives (#2) LL-25 and, to a lesser extent, (#4) GD-23 are both potent and substantially less hemolytic than (#1) LL-37. (#2) LL-25 and (#4) GD-23 have both been described previously, but the separation between antimicrobial and hemolytic activity in these two peptides is, to our knowledge, a new finding^59^. Both (#2) LL-25 and (#4) GD-23 truncate prior to L28, which in our recent structure-activity studies in full-length (#1) LL-37 is one of the two critical residues required for oligomerization – the other being I24. Because I24 is retained in (#2) LL-25, (#4) GD-23, and (#18) FF-14, it may be possible to develop these derivatives as lead therapeutics with better cell-type selectivity through further modifications around I24^30,31^.

## Conclusions

In summary, the portable biphenyl motif described here offers a rational design strategy for improving cathelicidin activity against gram-negative bacteria. Development approaches are further supported by the compatibility of cathelicidins with stabilizing interventions such as the use of D-amino acids as well as binding of some cathelicidins to serum components. Combined with our separate work on improving the cell type selectivity of cathelicidins via modulation of the dimerization interface, we are now positioned to apply natural cathelicidin biology and rational design principles to the goal of developing cathelicidin therapeutics built to address common modes of failure in peptide therapeutics.

## Experimental Section

### Antimicrobial susceptibility testing

Susceptibility testing was carried out based on previously described adaptations of CLSI methods modified for use with antimicrobial peptides. This includes the use of polypropylene plates as well as unadjusted Mueller Hinton Broth^69^. Methods were further miniaturized for use in 384-well plates at total volumes of either 35 or 50 µL. In brief, a 10x concentration of peptide was plated followed by the addition of a concentrated medium and cell mixture to yield a 1x mixture of peptide, medium, and cells. For example, if plating 50 µL, one would plate 5 µL of a 10x concentration of peptide followed by 45 µL of a 10/9 strength MHB and cell mixture. Plates were then monitored for absorbance at 600 nm for 18 hours at 37 °C at 30-minute intervals on a Tecan Spark plate reader, during which time they were housed in a large Tecan humidity cassette to minimize evaporation over the course of the experiment. Endpoint measures as depicted are the final time point at 18 hours. Growth curves associated with these experiments can be found in the **Supporting Information** and may be utilized to identify peptides or particular peptide concentrations that are partially inhibitory without achieving a true MIC. Strains used throughout are antimicrobial susceptibility testing quality control strains, namely *E. coli* ATCC 25922 and *P. aeruginosa* ATCC 27853. Data within **Figures 1-2**, **4**, **5 / S89**, and **6** are from three independent experiments each. In heatmap format, data show the median of these experiments, which is chosen as a central measure due to the standard CLSI practice of accepting a quality control result within one dilution of an expected MIC value as valid. For example, the colistin MIC against *P. aeruginosa* in these studies is typically 0.4 µM, but one can sometimes see late outgrowth at this concentration in the associated growth curves (*e.g.*, **Figure S43D**). Use of the median excludes such an outlier and focuses the visual interpretation of MICs in the heatmaps on a more typical value. The histogram depictions show the mean and standard deviation of each endpoint measure of OD600 and may be utilized to visualize spread in the raw data.

### Hemolysis assays

Hemolysis assays were carried out by treating defibrinated sheep blood (Hardy Diagnostics) with each test peptide. In brief, peptides were prepared at a concentration of 500 µM in water and then diluted to 400 µM with 5x PBS. These were then serially diluted to the levels shown and plated at 25 µL per well in 384-well plates. To these peptides, we then added 25 µL of sheep blood in PBS washed to supernatant clearance for a final concentration of 0.5% v/v. Wells were then monitored at OD600 in a fashion similar to the methods used for antimicrobial susceptibility testing over 18 hours, which provides the kinetic curves shown in the **Supporting Information**. At the end of 18 hours, 25 µL of PBS was added to each well followed by spinning of the 384-well plates for 10 minutes at 800 rcf. 25 µL of supernatant was then removed and added to a fresh 384-well plate for detection of free hemoglobin at 414 nm. Hemolysis controls throughout were thrice freeze-thawed blood from the same preparation used in plating, which were set to 100%. Data within **Figures 1-2**, **4**, **5 / S89**, and **6** are from three independent experiments each. The heatmap depictions as above show the median of three experiments. The histogram depictions of these data in the **Supporting Information** show the mean and standard deviation of three experiments and are used to identify the highest peptide concentrations at which < 20% of total maximum hemolysis signal is detected, which is the hemolysis threshold value applied to the main text tables. This measure is chosen because it represents a value at which hemolysis can be consistently qualitatively assessed as present or absent when completing these experiments with colistin as an antibiotic comparator control.

### Table calculated values

As discussed in the above sections, MIC cutoffs used in the **MIC Ec** and **MIC Pa** columns of the main text tables are derived by visual interpretation of the heatmaps presented in the **Supporting Information**. Similarly, hemolysis threshold values in the **< 20% Hemo** columns of these tables are derived by determining the highest concentration of an individual peptide at which the hemolysis signal is < 20% of the highest assay value. Two calculated values based on these empiric observations are presented in the main text tables. **Hemo/MIC** = (**< 20% Hemo**) / **MIC Ec**, where larger numbers indicate higher peptide concentrations required to induce hemolysis, lower peptide concentrations required to kill *E. coli*, or both. Higher numbers are thus rated as desirable. **Fold Δ** = (**Hemo/MIC** of compound) / (**Hemo/MIC** of LL-37), where larger numbers represent greater separation between the quantities of peptide associated with hemolysis and gram-negative killing in comparison with the parent peptide LL-37. Higher numbers are thus rated as desirable. When making these calculations, MICs >50 µM are treated as 100 µM, while hemolysis cutoffs of >200 µM are treated as 200 µM.

### Dual membrane permeabilization assays

Dual membrane permeabilization assays based on prior protocols (*e.g.* ^70^) were carried out in M9+glucose with divalent cation concentrations adjusted to the levels present in the unadjusted MHB used for antimicrobial susceptibility testing in this study. In brief, cells were grown to an OD600 of approximately 0.5 and then washed and resuspended in M9+glucose. To this cell suspension, we then added a final concentration of 10 µM N-phenyl-naphthylamine (NPN) and 5 µM propidium iodide (PI). We then added 50 µL of this mixture to 5 µL of 10x concentrated peptide over a range of concentrations in 384-well black, transparent bottom plates and monitored NPN (outer membrane permeabilization) at Ex 350 / Em 420 nm and PI (inner membrane permeabilization) at Ex 535 / Em 617 nm at 15-minute intervals over 4 hours at 37 °C in a Tecan Spark plate reader using a large humidity cassette to minimize evaporation. Quantification of fluorescence was scaled to that observed for (#1) LL-37. In brief, the average of cell background with no fluorophores added was subtracted from all wells. Fold-changes over no-peptide controls for each concentration of each peptide were then calculated. Finally, the highest fold-change within LL-37 was set to 100%, with remaining values scaled to this reference. These scaled values are shown in **Figures S155-159**. The single values shown in **Figure 7** represent a permeabilization index that is the product of the maximum permeabilization observed for each peptide and the proportion of the assay time required for that peptide to demonstrate at least 80% of its maximum level of permeabilization. Several of the high peptide wells in these experiments, particularly for (#18) FF-14, yielded signals over assay at early timepoints. Where this occurs, the over assay timepoint was substituted with the next within-assay timepoint. The general effect of this is to underestimate the peaks of (#18) FF-14 permeabilization; no effect is expected on quantification of the lower limits of (#18) FF-14 permeabilization.

### Circular dichroism

Peptides were dissolved to 0.5 mg/mL in water with 20% TFE. Spectra were collected on an Aviv 420 circular dichroism spectrometer from 260-190 nm at 25 °C and 37 °C. Estimation of secondary structure content was completed using BeStSel^66^ and is presented as a measure of relative helical propensity among peptides.

### Serum stability

Master tubes corresponding to each peptide containing either PBS or PBS with 10% pooled human serum (Innovative Research) were prepared in LoBind tubes (Eppendorf). The serum condition was pre-cleared by spinning at 21,380 rcf for 10 minutes at 4 °C in a Hermle z326k centrifuge and collecting 80% of the supernatant for experiments. Peptide was then added to 50 µM in both PBS and PBS + 10% serum conditions and incubated at 37 °C with harvesting at 0 (immediate harvest after addition of peptide), 15, 30, 60, 120, or 240 minutes into polypropylene tubes. At the time of harvest, TCA was added to a final concentration of 1% v/v, and tubes were incubated at 70 °C for 10 minutes followed by room temperature for 10 minutes under the “Heat” condition collected for all timepoints versus 20 minutes at room temperature under the “No Heat” condition collected only at 0 minutes to estimate recovery independent of degradation. Peptides were then spun as above for 5 minutes. 80% of supernatant was harvested to a fresh polypropylene tube and again spun as above for 5 minutes. 80% of the resultant supernatant was then taken for LCMS analysis on an Agilent 6550 QTOF with a 10 minute 1-91 %B water / acetonitrile gradient with 0.1% formic acid on a Zorbax 2.1 x 150 mm 300SB-C3 column.

Data were analyzed by integrating the peak areas of an extracted ion chromatogram (EIC) covering 2 m/z centered on the base peak of each peptide. To calculate recovery as in **Figure 9A**, the EIC area from a given PBS + 10% serum condition at a given timepoint was divided by the same area from the PBS only condition. To calculate stability over time as in **Figure 9B**, this initial 0 minute recovery ratio set to 1 was divided by the (10% serum + PBS) / PBS recovery ratio collected at each time point. Data in **Figure 9** are the mean and standard deviation of three independent experiments.

### Peptide synthesis

Peptides used in these studies were synthesized on automated flow synthesizers in a manner similar to those previously described^71–74^. Resin was 4-(4-hydroxymethyl-3-methoxyphenoxy)butyric acid (HMPB, ChemMatrix) except in the case of C-amidated peptides, where Rink Amide (ChemMatrix) was used. Cleavage was generally carried out by treating resin with Reagent K (82.5 % trifluoroacetic acid (TFA), 5% water, 5% phenol, 2.5% 1,2-ethanedithiol) for 2 hours at room temperature followed by precipitation with cold ether. Precipitated peptide was then resuspended in 50% water + 0.1% TFA / 50% acetonitrile + 0.1% TFA and lyophilized. Crude material was purified most commonly by reverse phase flash purification on a Biotage Selekt flash chromatography instrument with selected peptides purified by reverse phase high performance liquid chromatography. Purity was determined by HPLC and is annotated with each compound in the **Supporting Information**.

Analytical data associated with these syntheses are shown in **Figures S160-171**(associated with **Figure 1**), **172-181** (associated with **Figure 2**), **182-192** (associated with **Figure 4**), **193-198** (associated with **Figure 5**), **199-204** (associated with **Figure S89**), and **205-214** (associated with **Figure 6**). Peptides that are reused as controls across more than one experiment are grouped with the first figure in which they appear. Single-pass purifications were used throughout. All compounds demonstrated ≥95% purity by HPLC with the following exceptions among stapled or lipidated compounds: i) stapled peptides in **Figure 4** (range 84.5%-94.6%, as well as a tested major side-product of the stapling reaction in **Figure S187** at 54.4%); ii) (#39) (76%, **Figure S198**) in **Figure 5**; iii) (#47) (94%, **Figure S206**), (#48) and (#52) (both 87%, **Figures S207** and **S211**) as well as (#50) (71%, **Figure S209**) in **Figure 6**; iv) all compounds in **Figure S89** (range 43-82%, **Figures S199-204**). No additional purification or testing of compounds with <95% purity was pursued, as initial testing did not indicate a biological signal of interest and the compounds in question were not relevant to the description of the biphenyl motif that is the focus of this paper.

### Perfluoroaryl stapling

Methods for stapling are based on those previously reported (*e.g.*, ^20^). Intervals of *i,i+3* or *i,i+4* were stapled using hexafluorobenzene, while intervals of *i,i+7* were stapled with decafluorobiphenyl. In brief, peptide was dissolved to 1 mM in dimethylformamide (DMF) with the addition of stapling agent (2 mM final concentration for decafluorobiphenyl; 20 mM final concentration for hexafluorobenzene) and Tris to a final concentration of 30 mM. Reactions were then allowed to proceed for approximately 4-5 hours prior to dilution in water and purification by reverse phase flash chromatography.

### Lipidation

Lipidation was generally carried out via standard peptide bond formation procedures either in batch or by semi-automated flow synthesis^74^ between the on-resin protected peptide and the carboxylic acid on each of the tails used in these studies (**Figures 5-6** and **S89**).

Cleavage then proceeded as per above for most moieties with selected exceptions for which alternatives such as Reagent B (88% TFA, 5% water, 5% phenol, 2% triisopropylsilane) were substituted.

## Ancillary Information

Supporting information is available free of charge at the provided address. This includes heatmap and histogram depictions as well as growth and hemolysis curves associated with **Figures 1-2** and **4-6**, kinetic permeabilization curves associated with **Figure 7**, and analytical data for all presented peptides.

## Author Contributions

JSA designed and conducted experiments and wrote the paper. CJ and DAV conducted experiments. BLP guided experiments and wrote the paper.

## Acknowledgements

This work has been funded by the National Institute of Allergy and Infectious Diseases (T32 AI007061 and K08 AI166345 to JSA; U19 AI142780 to BLP) and by the Cystic Fibrosis Foundation (ALBIN19F0, ALBIN21Q0, ALBIN22A0-KB to JSA).

## Table of Contents Graphic

This graphic is adapted from PDB 2K6O as reported by Wang^36^, used under CC0 1.0.

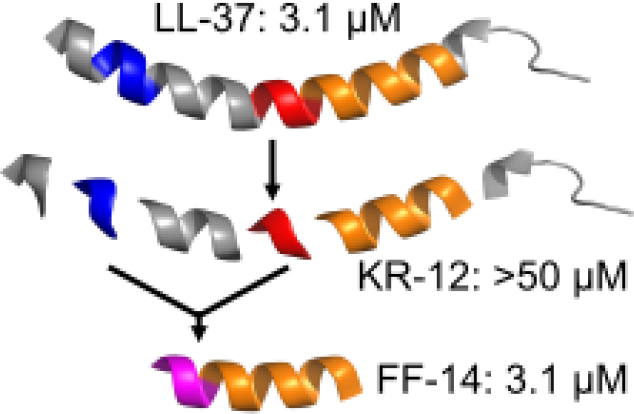

