## Supplemental_Information for "A portable N-terminal biphenyl motif in LL-37 enhances the activity of short human cathelicidin derivatives"

**Table of Contents:**


| | |
|---|---|
| Figure S1 Heatmaps associated with Figure 1 | S3-S4 |
| Figure S2 Histograms associated with Figure 1 | S5-S6 |
| Figures S3-S15 Growth curves associated with Figure 1 | S7-S32 |
| Figures S16-S28 Hemolysis curves associated with Figure 1 | S33-S58 |
| Figure S29 Heatmaps associated with Figure 2 | S59-S60 |
| Figure S30 Histograms associated with Figure 2 | S61-S62 |
| Figures S31-S43 Growth curves associated with Figure 2 | S63-S88 |
| Figures S44-S56 Hemolysis curves associated with Figure 2 | S89-S114 |
| Figure S57 Heatmaps associated with Figure 4 | S115-S116 |
| Figure S58 Histograms associated with Figure 4 | S117-S118 |
| Figures S59-S73 Growth curves associated with Figure 4 | S119-S148 |
| Figures S74-S88 Hemolysis curves associated with Figure 4 | S149-S178 |
| Figure S89 Structure activity of FK-13 lipidation | S179-S180 |
| Figures S90-S91 Heatmaps associated with Figures 5 and S89 | S181-S184 |
| Figures S92-S93 Histograms associated with Figures 5 and S89 | S185-S188 |
| Figures S94-S109 Growth curves associated with Figures 5 and S89 | S189-S220 |
| Figures S110-S125 Hemolysis curves associated with Figures 5 and S89 | S221-S252 |
| Figure S126 Heatmaps associated with Figure 6 | S253-S254 |
| Figure S127 Histograms associated with Figure 6 | S255-S256 |
| Figures S128-S141 Growth curves associated with Figure 6 | S257-S284 |
| Figures S142-S155 Hemolysis curves associated with Figure 7 | S285-S312 |
| Figures S156-S159 Kinetic membrane permeabilization from Figure 8 | S313-S320 |
| Figures S160-S171 Analytical figures associated with Figure 1 | S321-S344 |
| Figures S172-S181 Analytical figures associated with Figure 2 | S345-S364 |
| Figures S182-S192 Analytical figures associated with Figure 4 | S365-S386 |
| Figures S193-S198 Analytical figures associated with Figure 5 | S387-S398 |
| Figures S199-S204 Analytical figures associated with Figure S89 | S399-S410 |
| Figures S205-S214 Analytical figures associated with Figure 6 | S411-S430 |

A.

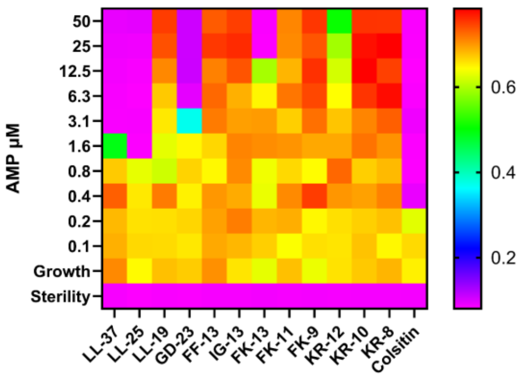


B.

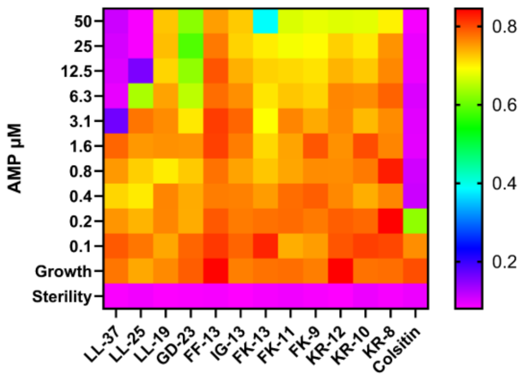


C.

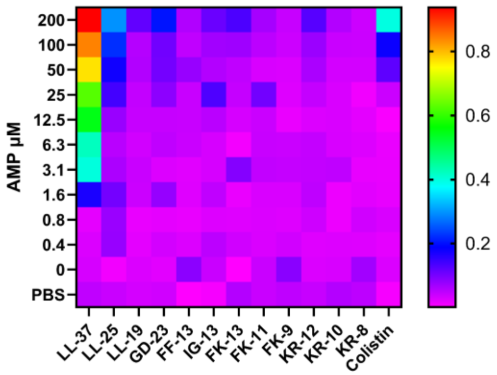


D.

```
LL-37    LLGDFFRKSKEKIGKEFKRIVQRIKDFLRNLVPRTES   37
LL-25    LLGDFFRKSKEKIGKEFKRIVQRIK------------   25
LL-19    LLGDFFRKSKEKIGKEFKR------------------   19
GD-23    --GDFFRKSKEKIGKEFKRIVQRIK------------   23
FF-13    ----FFRKSKEKIGKEF--------------------   13
IG-13    ------------IGKEFKRIVQRIK------------   13
FK-13    ----------------FKRIVQRIKDFLR--------   13
FK-11    ----------------FKRIVQRIKDF----------   11
FK-9     ----------------FKRIVQRIK------------   9
KR-12    -----------------KRIVQRIKDFLR--------   12
KR-10    -----------------KRIVQRIKDF----------   10
KR-8     -----------------KRIVQRIK------------   8
```

**Figure S1 – Heatmap depiction of the antimicrobial and hemolytic activities of shorter derivatives of LL-37. A.** Antimicrobial susceptibility testing showing the activity of each derivative as well as a colistin control for reference against *E. coli* (ATCC 25922). In the heatmap plate depiction, cooler colors indicate low or absent bacterial growth as detected by the OD600 measured at 18 hours of incubation at 37 °C. **B.** As in **A.**, but with *P. aeruginosa* (ATCC 27853). **C.** Hemolysis assays in which defibrinated sheep blood in PBS was incubated with the indicated peptide for 18 hours at 37 °C with subsequent quantification of hemoglobin released into the supernatant at 414 nm. Cool colors indicate little or no release of hemoglobin, where all levels are scaled to 100% hemolysis freeze-thaw controls. Levels throughout are the median across three independent experiments as explained further in **Methods**. **D.** Alignment showing the sequence of LL-37 in comparison with shorter derivatives tested in initial studies, which have been described previously.

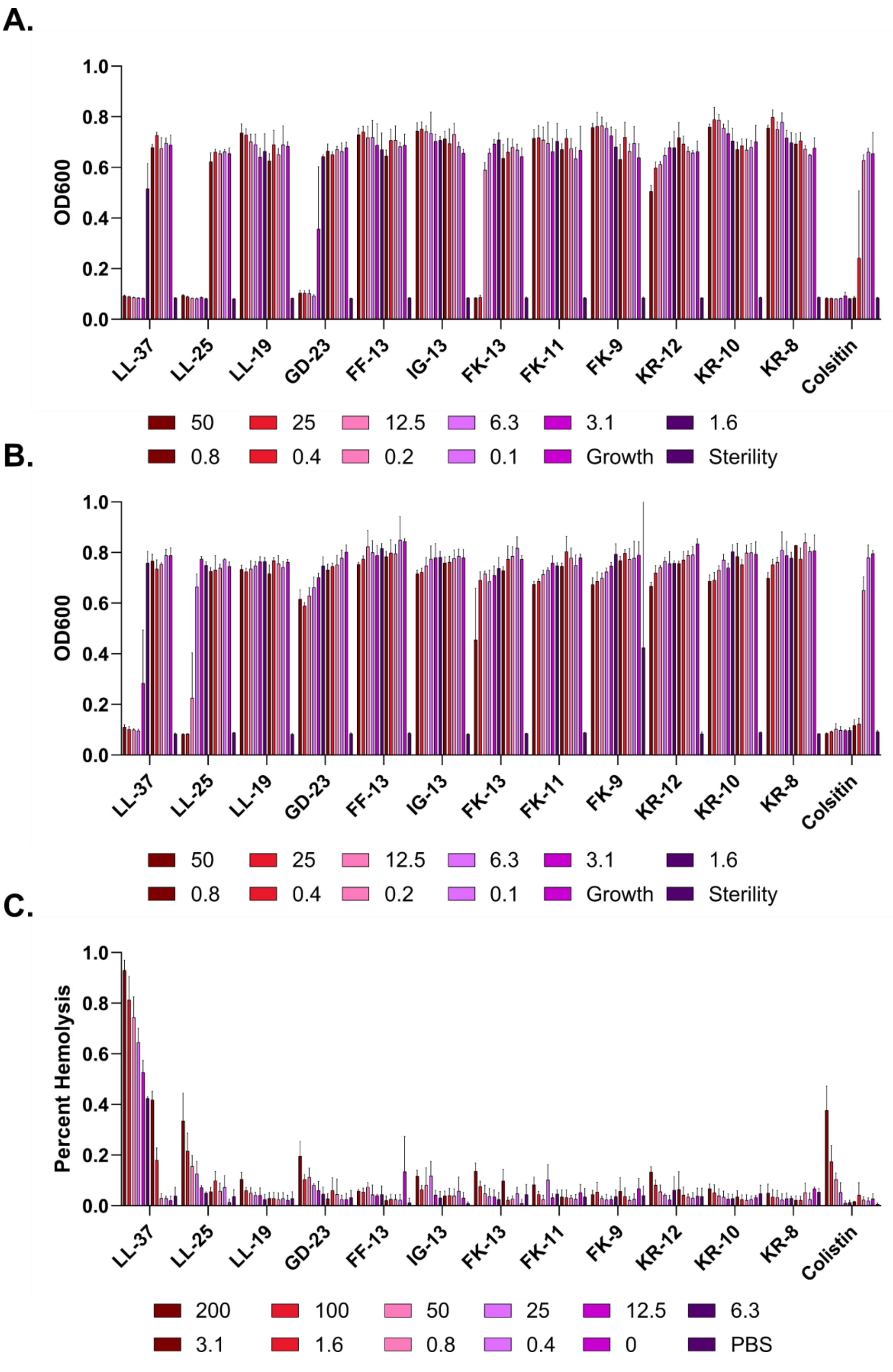
A.
OD600
1.0
0.8
0.6
0.4
0.2
0.0
LL-37
LL-25
LL-19
GD-23
FF-13
IG-13
FK-13
FK-11
FK-9
KR-12
KR-10
KR-8
Colsitin
50
25
12.5
6.3
3.1
1.6
0.8
0.4
0.2
0.1
Growth
Sterility
B.
OD600
1.0
0.8
0.6
0.4
0.2
0.0
LL-37
LL-25
LL-19
GD-23
FF-13
IG-13
FK-13
FK-11
FK-9
KR-12
KR-10
KR-8
Colsitin
50
25
12.5
6.3
3.1
1.6
0.8
0.4
0.2
0.1
Growth
Sterility
C.
Percent Hemolysis
1.0
0.8
0.6
0.4
0.2
0.0
LL-37
LL-25
LL-19
GD-23
FF-13
IG-13
FK-13
FK-11
FK-9
KR-12
KR-10
KR-8
Colsitin
200
100
50
25
12.5
6.3
3.1
1.6
0.8
0.4
0
PBS

**Figure S2 – Histograms corresponding to Figure 1**. **A.** Growth of *E. coli* (ATCC 25922) by OD600 at 18 hours at 37 °C. Error bars indicate the standard deviation of three independent experiments. **B.** As in **A.**, but with *P. aeruginosa* (ATCC 27853). **C.** Hemolysis assays in which sheep RBCs in PBS were incubated with the indicated peptide for 18 hours at 37 °C with subsequent quantification of hemoglobin released into the supernatant at 414 nm relative to a 100% hemolysis freeze-thaw control. Error bars indicate the standard deviation of three independent experiments. All peptide levels given in μM.

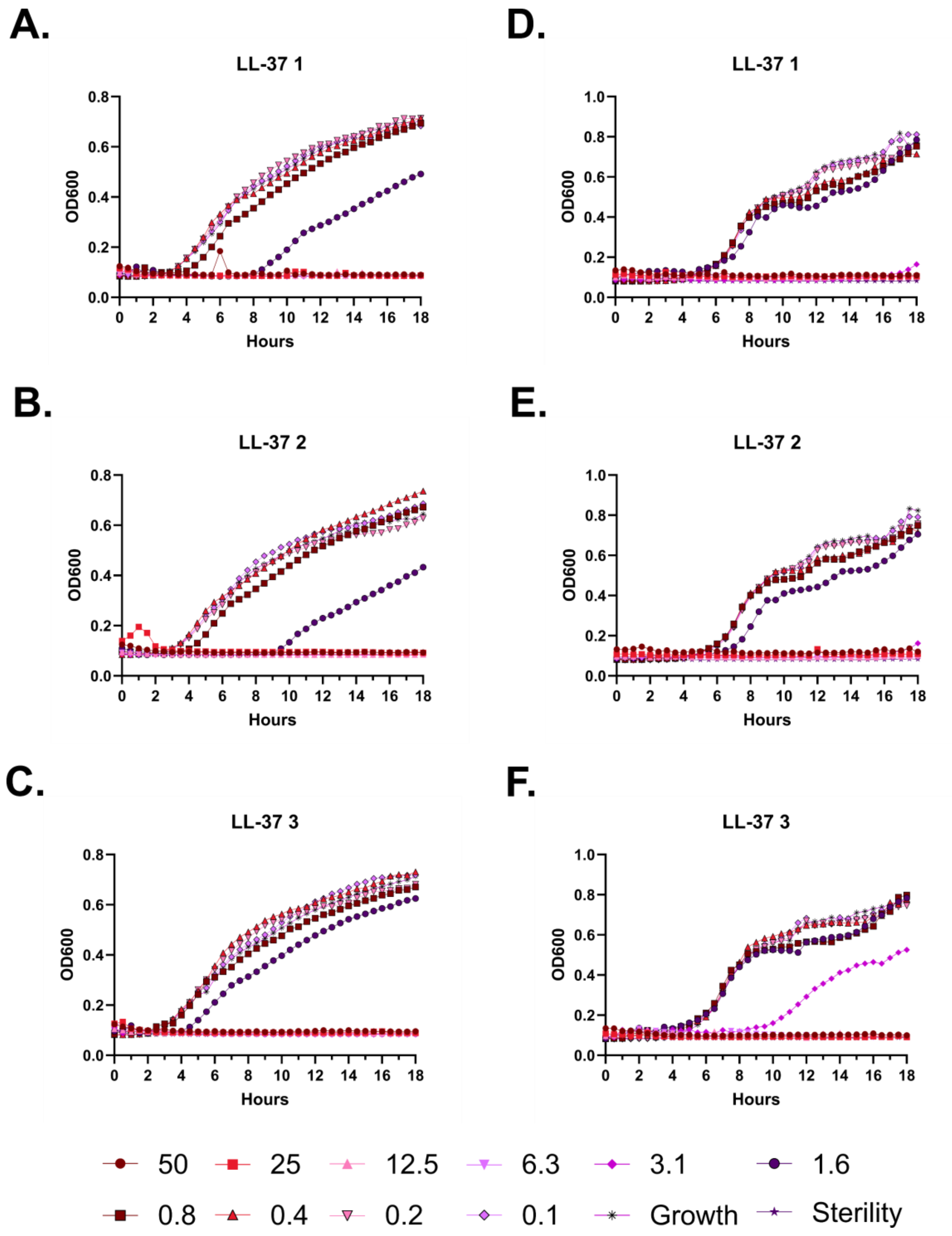
A.
LL-37 1
D.
LL-37 1
B.
LL-37 2
E.
LL-37 2
C.
LL-37 3
F.
LL-37 3
OD600
Hours
50
25
12.5
6.3
3.1
1.6
0.8
0.4
0.2
0.1
Growth
Sterility

**Figure S3 – Antimicrobial susceptibility curves for (#1) LL-37. A-C.** Independent experiments with *E. coli*. **D-F.** Independent experiments with *P. aeruginosa.* Growth over 18 hours at 37 °C with a gradient of peptide concentrations given in μM.

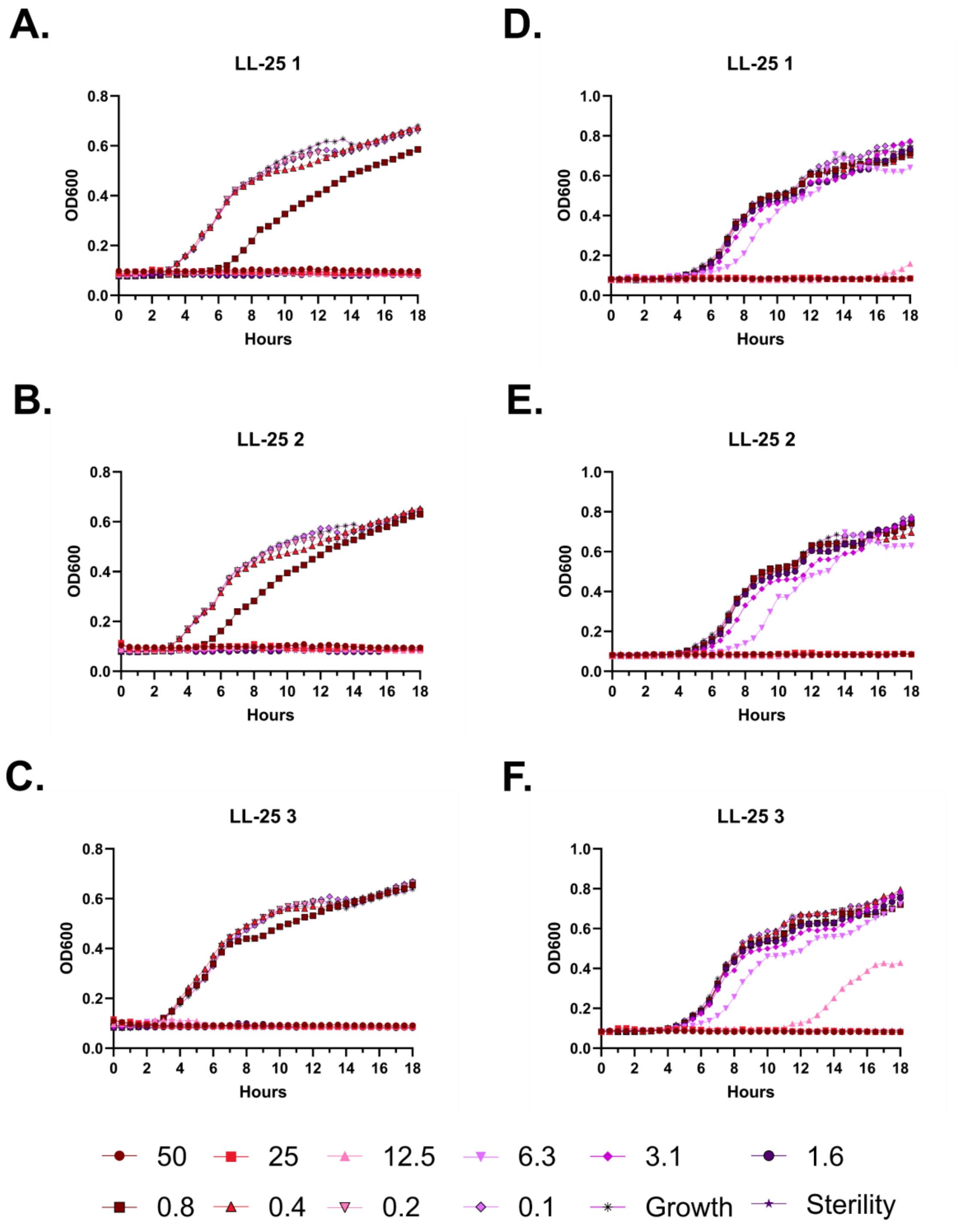
A.
LL-25 1
B.
LL-25 2
C.
LL-25 3
D.
LL-25 1
E.
LL-25 2
F.
LL-25 3
OD600
Hours
50
25
12.5
6.3
3.1
1.6
0.8
0.4
0.2
0.1
Growth
Sterility

**Figure S4 – Antimicrobial susceptibility curves for (#2) LL-25. A-C.** Independent experiments with *E. coli*. **D-F.** Independent experiments with *P. aeruginosa.* Growth over 18 hours at 37 °C with a gradient of peptide concentrations given in μM.

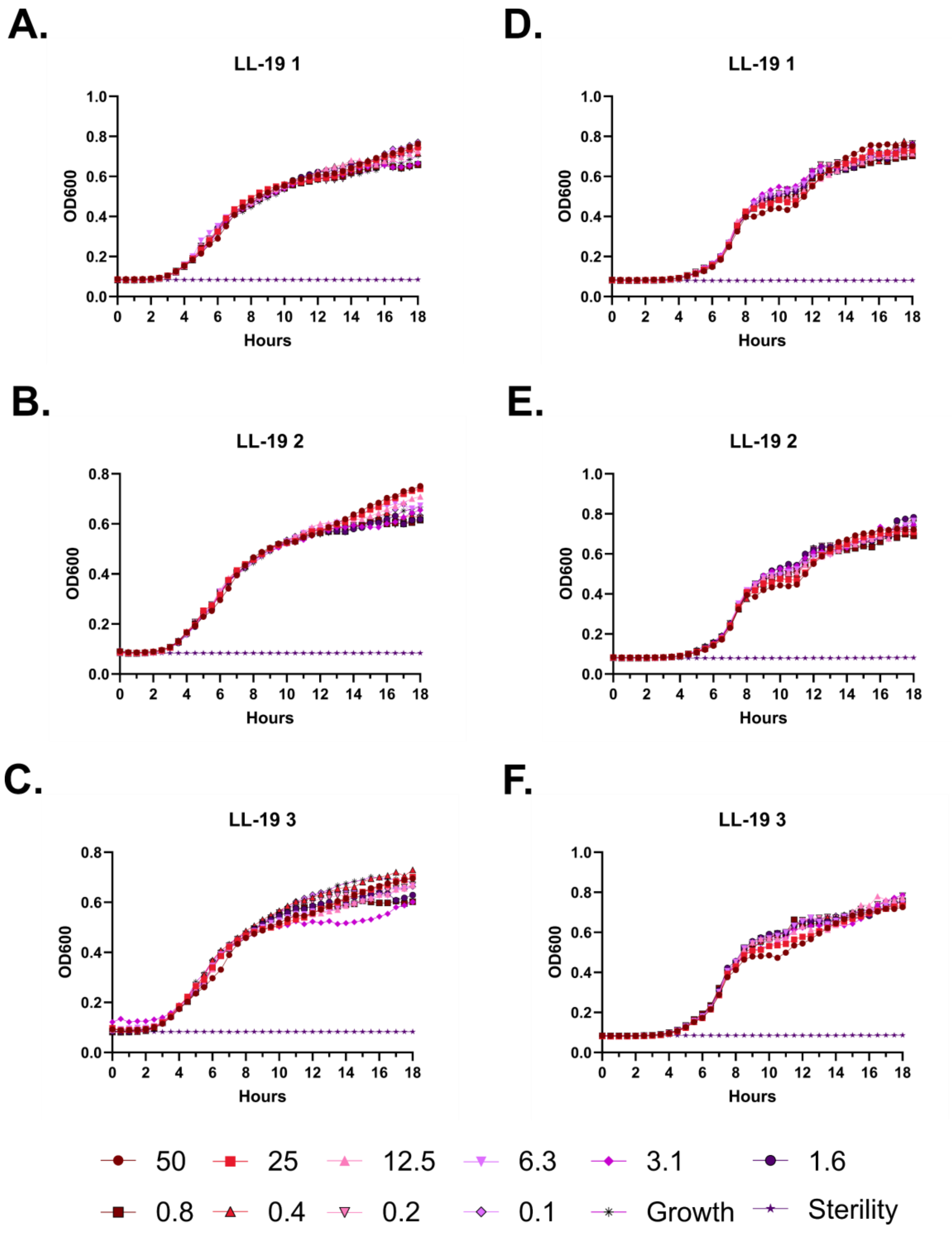
A.
LL-19 1
D.
LL-19 1
B.
LL-19 2
E.
LL-19 2
C.
LL-19 3
F.
LL-19 3
OD600
Hours
50
25
12.5
6.3
3.1
1.6
0.8
0.4
0.2
0.1
Growth
Sterility

**Figure S5 – Antimicrobial susceptibility curves for (#3) LL-19. A-C.** Independent experiments with *E. coli*. **D-F.** Independent experiments with *P. aeruginosa.* Growth over 18 hours at 37 °C with a gradient of peptide concentrations given in μM.

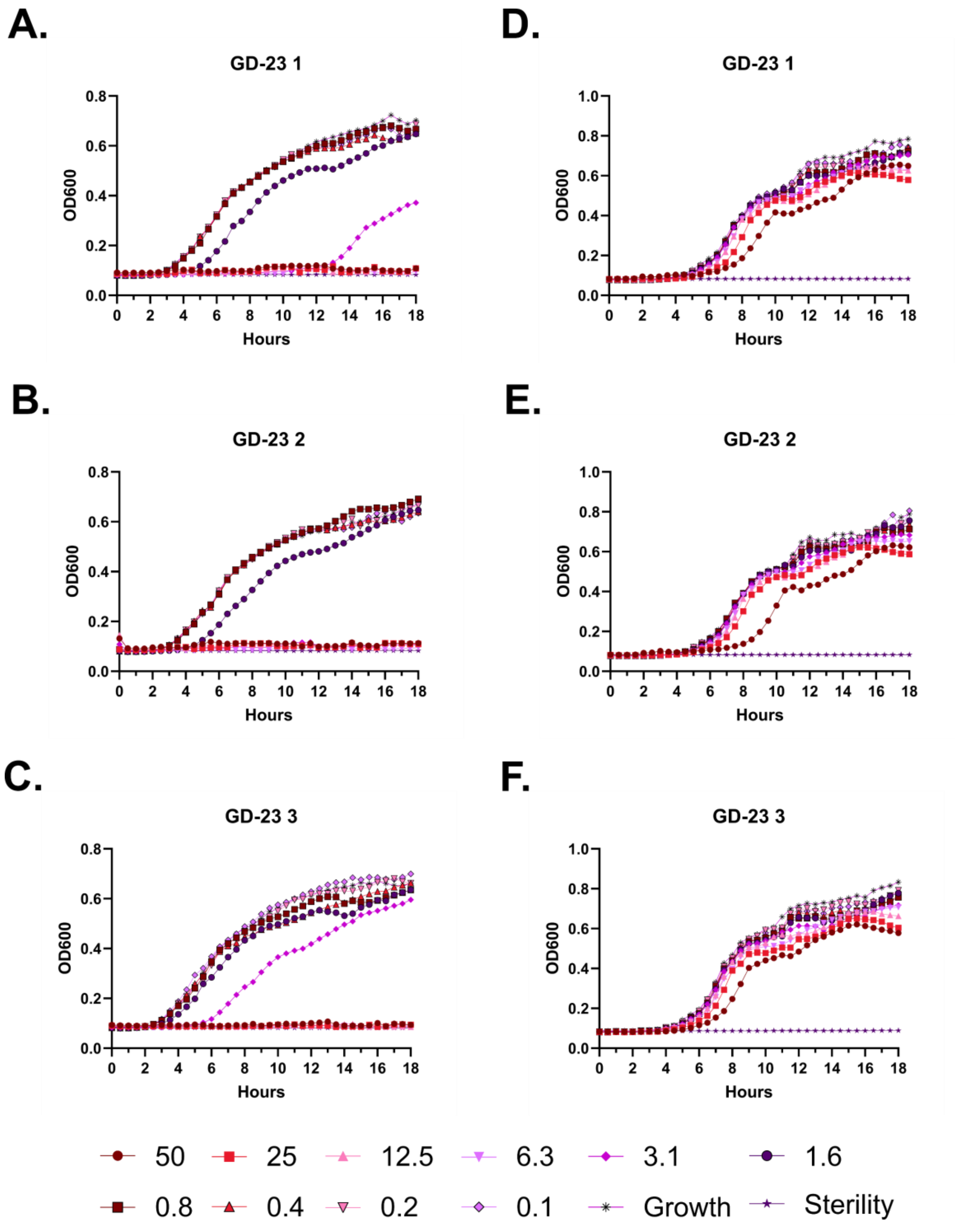
A.
GD-23 1
B.
GD-23 2
C.
GD-23 3
D.
GD-23 1
E.
GD-23 2
F.
GD-23 3
OD600
Hours
50
25
12.5
6.3
3.1
1.6
0.8
0.4
0.2
0.1
Growth
Sterility

**Figure S6 – Antimicrobial susceptibility curves for (#4) GD-23. A-C.** Independent experiments with *E. coli*. **D-F.** Independent experiments with *P. aeruginosa.* Growth over 18 hours at 37 °C with a gradient of peptide concentrations given in μM.

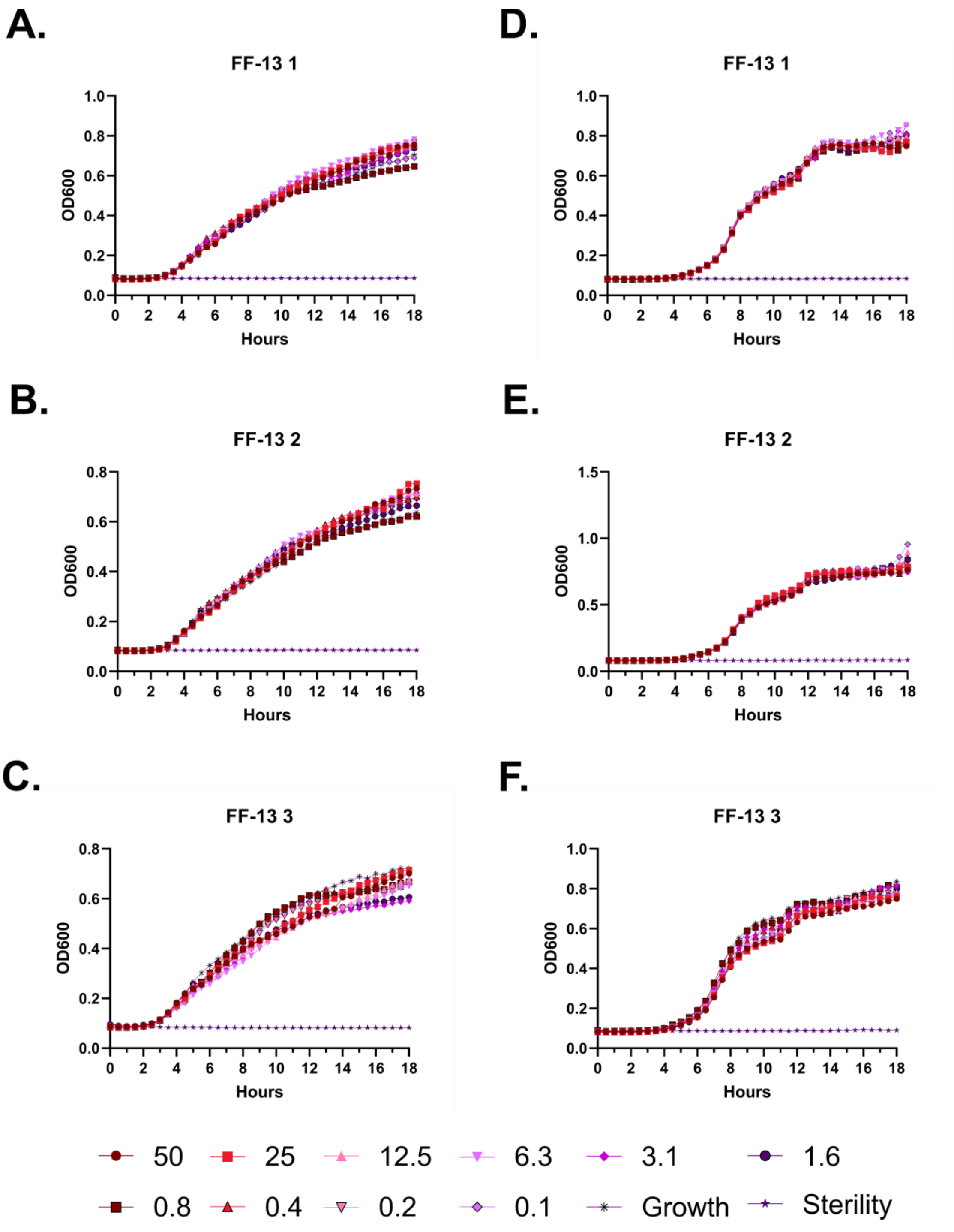

A.
FF-13 1
OD600
Hours
B.
FF-13 2
OD600
Hours
C.
FF-13 3
OD600
Hours
D.
FF-13 1
OD600
Hours
E.
FF-13 2
OD600
Hours
F.
FF-13 3
OD600
Hours
50
25
12.5
6.3
3.1
1.6
0.8
0.4
0.2
0.1
Growth
Sterility

**Figure S7 – Antimicrobial susceptibility curves for (#5) FF-13. A-C.** Independent experiments with *E. coli*. **D-F.** Independent experiments with *P. aeruginosa.* Growth over 18 hours at 37 °C with a gradient of peptide concentrations given in μM.

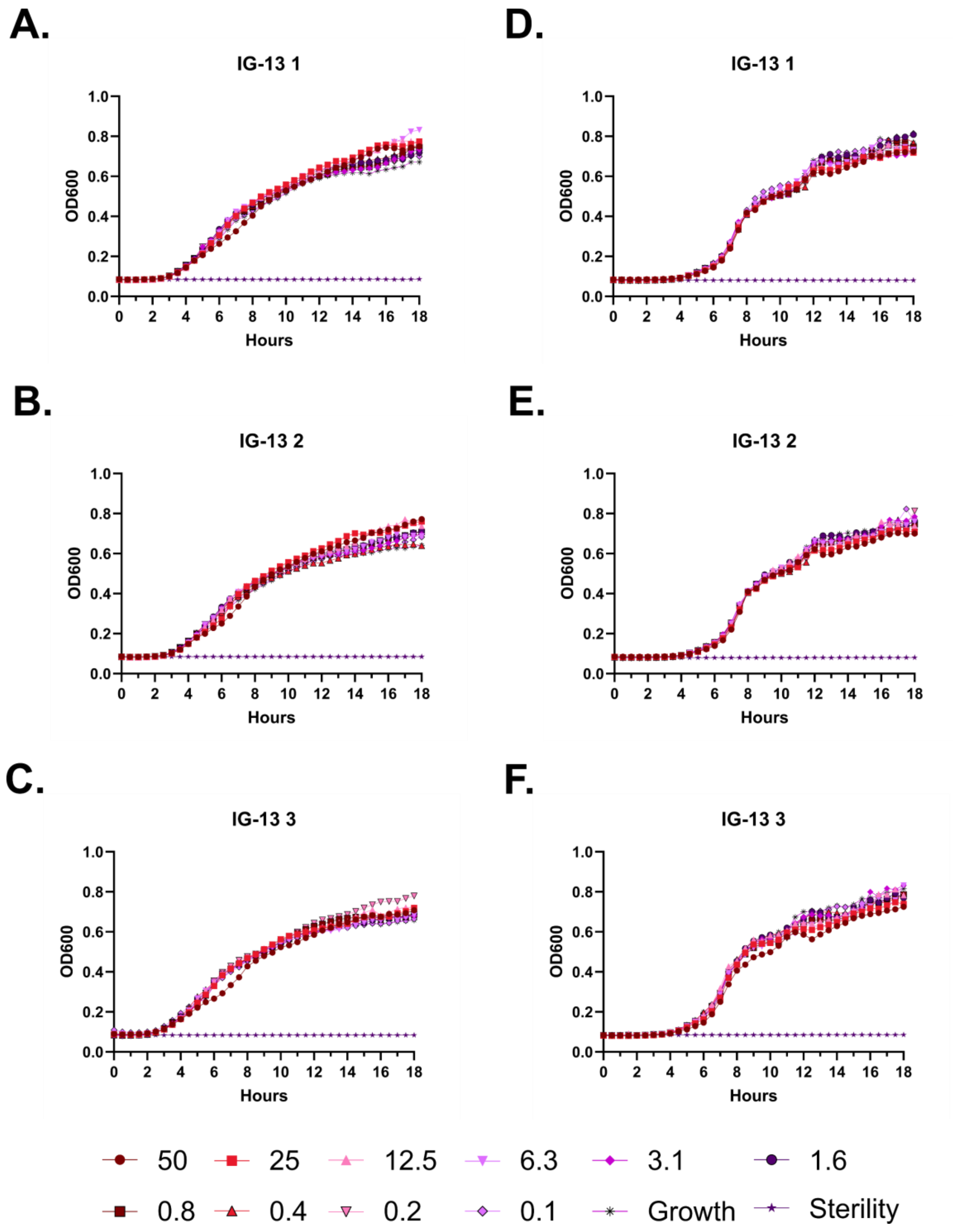
A.
IG-13 1
D.
IG-13 1
B.
IG-13 2
E.
IG-13 2
C.
IG-13 3
F.
IG-13 3
OD600
Hours
50
25
12.5
6.3
3.1
1.6
0.8
0.4
0.2
0.1
Growth
Sterility

**Figure S8 – Antimicrobial susceptibility curves for (#6) IG-13. A-C.** Independent experiments with *E. coli*. **D-F.** Independent experiments with *P. aeruginosa.* Growth over 18 hours at 37 °C with a gradient of peptide concentrations given in μM.

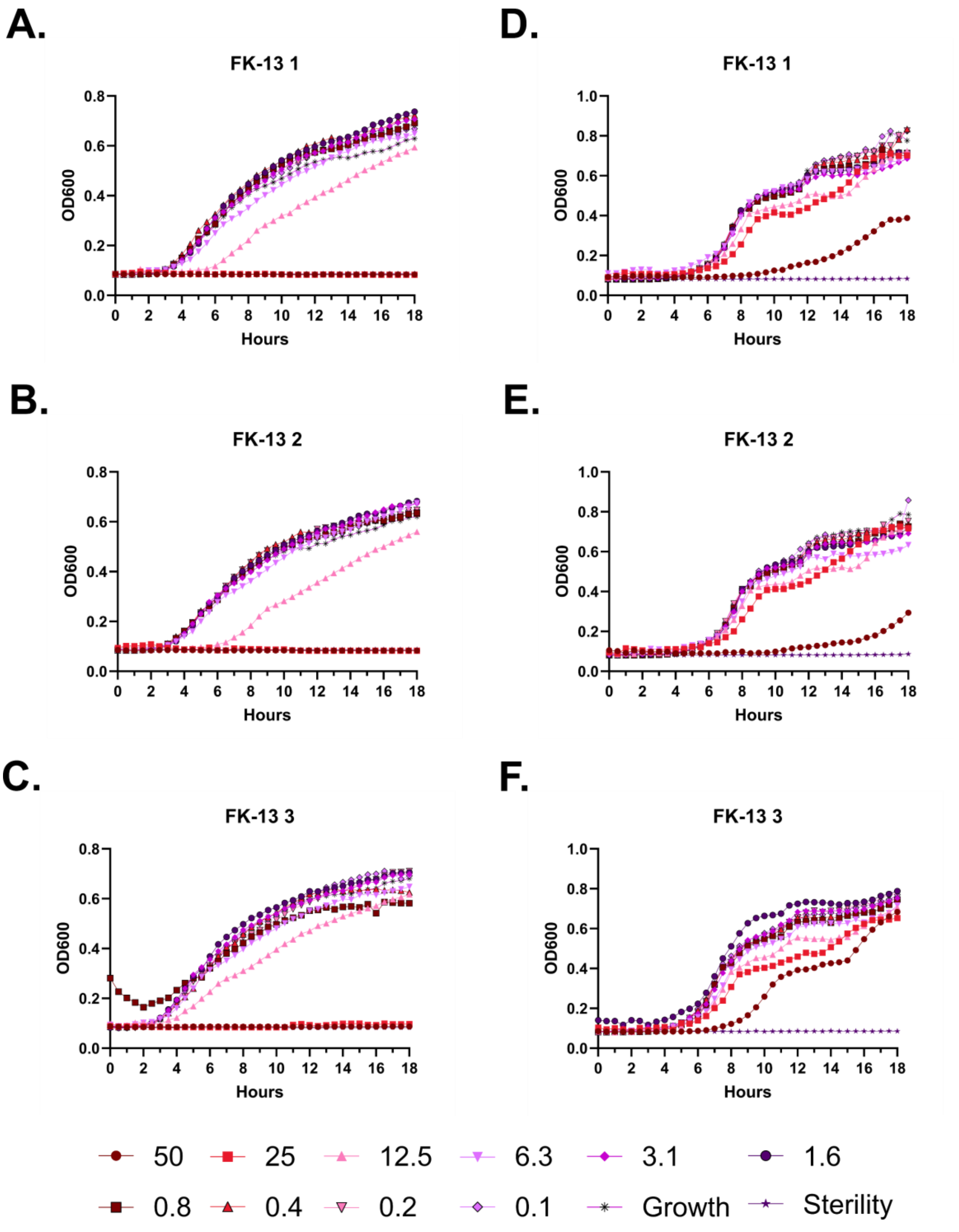
A.
FK-13 1
B.
FK-13 2
C.
FK-13 3
D.
FK-13 1
E.
FK-13 2
F.
FK-13 3
OD600
Hours
0 2 4 6 8 10 12 14 16 18
0.0 0.2 0.4 0.6 0.8
0.0 0.2 0.4 0.6 0.8 1.0
50
25
12.5
6.3
3.1
1.6
0.8
0.4
0.2
0.1
Growth
Sterility

**Figure S9 – Antimicrobial susceptibility curves for (#7) FK-13. A-C.** Independent experiments with *E. coli*. **D-F.** Independent experiments with *P. aeruginosa.* Growth over 18 hours at 37 °C with a gradient of peptide concentrations given in μM.

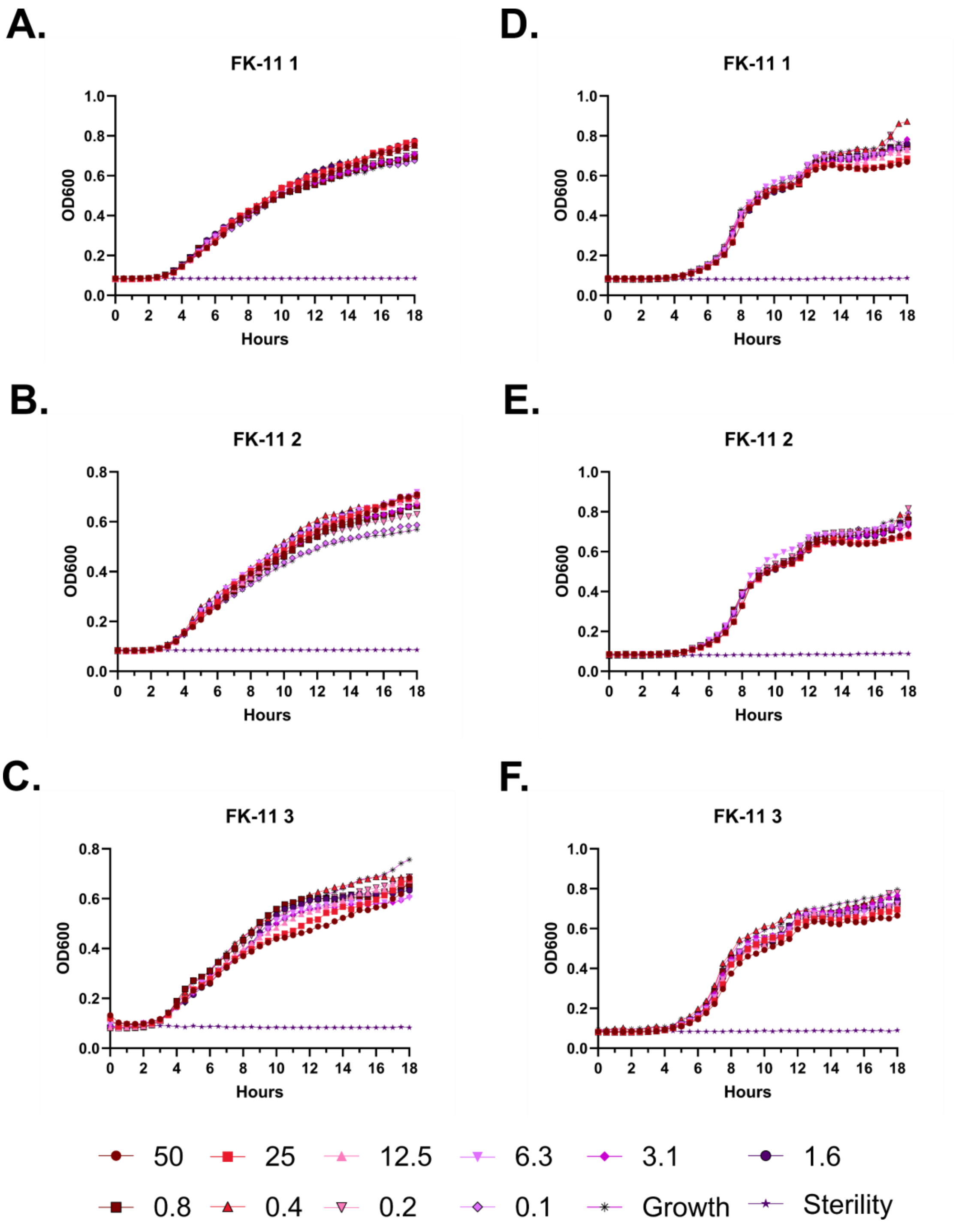

A.
FK-11 1
OD600
Hours
B.
FK-11 2
OD600
Hours
C.
FK-11 3
OD600
Hours
D.
FK-11 1
OD600
Hours
E.
FK-11 2
OD600
Hours
F.
FK-11 3
OD600
Hours
50
25
12.5
6.3
3.1
1.6
0.8
0.4
0.2
0.1
Growth
Sterility

**Figure S10 – Antimicrobial susceptibility curves for (#8) FK-11. A-C.** Independent experiments with *E. coli*. **D-F.** Independent experiments with *P. aeruginosa.* Growth over 18 hours at 37 °C with a gradient of peptide concentrations given in μM.

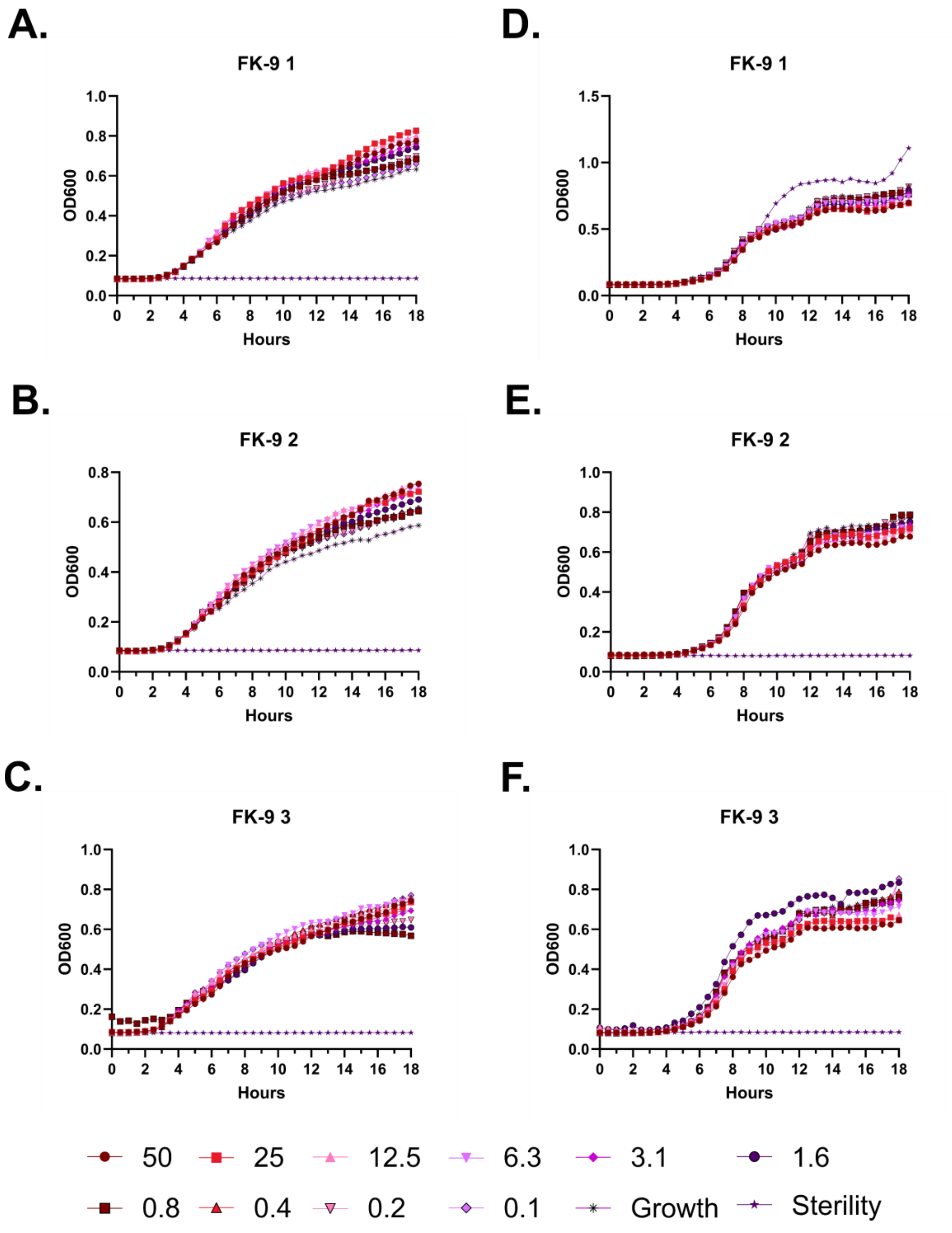

A.
FK-9 1
OD600
Hours
D.
FK-9 1
B.
FK-9 2
E.
FK-9 2
C.
FK-9 3
F.
FK-9 3
50
25
12.5
6.3
3.1
1.6
0.8
0.4
0.2
0.1
Growth
Sterility

**Figure S11 – Antimicrobial susceptibility curves for (#9) FK-9. A-C.** Independent experiments with *E. coli*. **D-F.** Independent experiments with *P. aeruginosa.* Growth over 18 hours at 37 °C with a gradient of peptide concentrations given in μM.

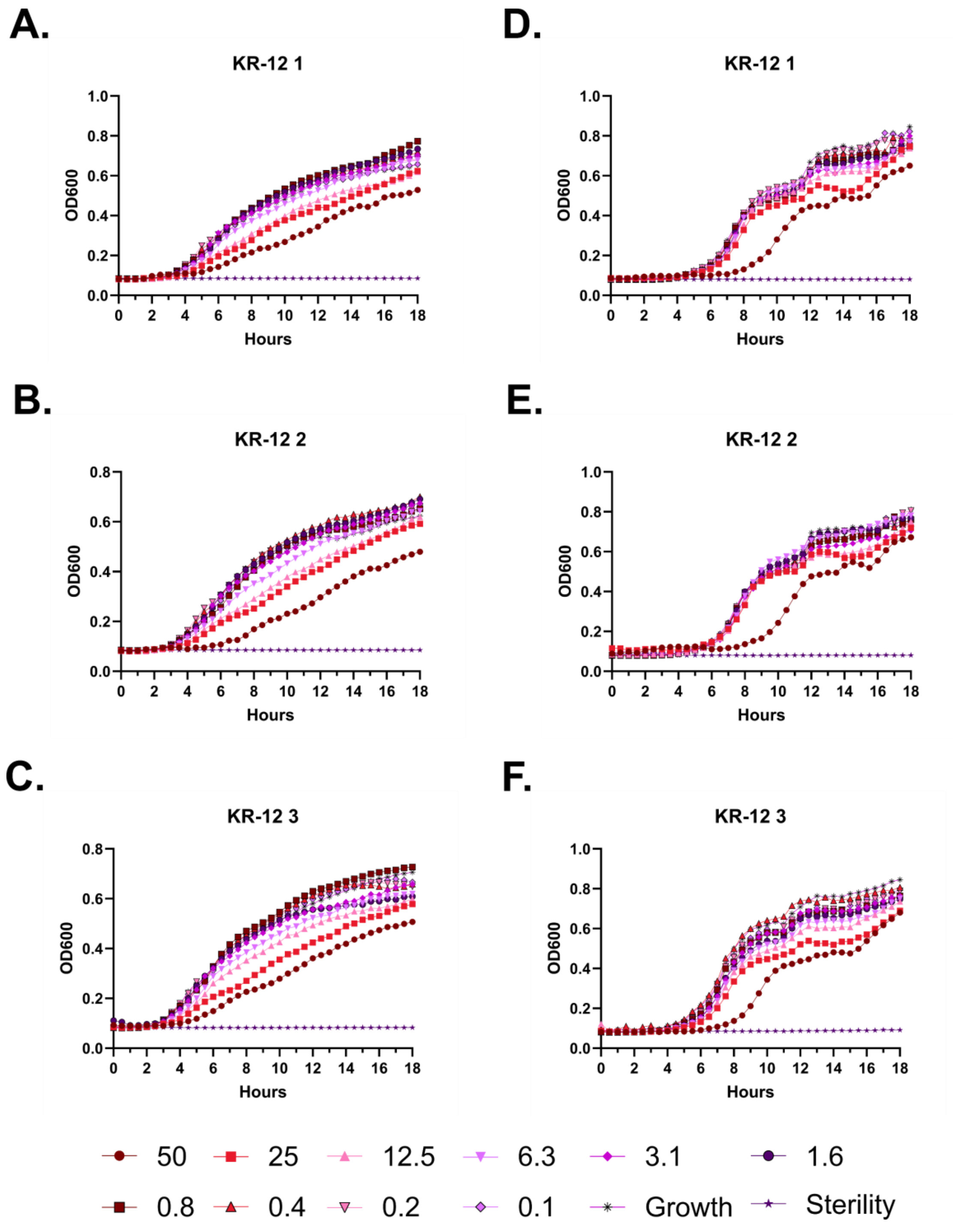
A.
KR-12 1
D.
KR-12 1
B.
KR-12 2
E.
KR-12 2
C.
KR-12 3
F.
KR-12 3
OD600
Hours
50
25
12.5
6.3
3.1
1.6
0.8
0.4
0.2
0.1
Growth
Sterility

**Figure S12 – Antimicrobial susceptibility curves for (#10) KR-12. A-C.** Independent experiments with *E. coli*. **D-F.** Independent experiments with *P. aeruginosa.* Growth over 18 hours at 37 °C with a gradient of peptide concentrations given in μM.

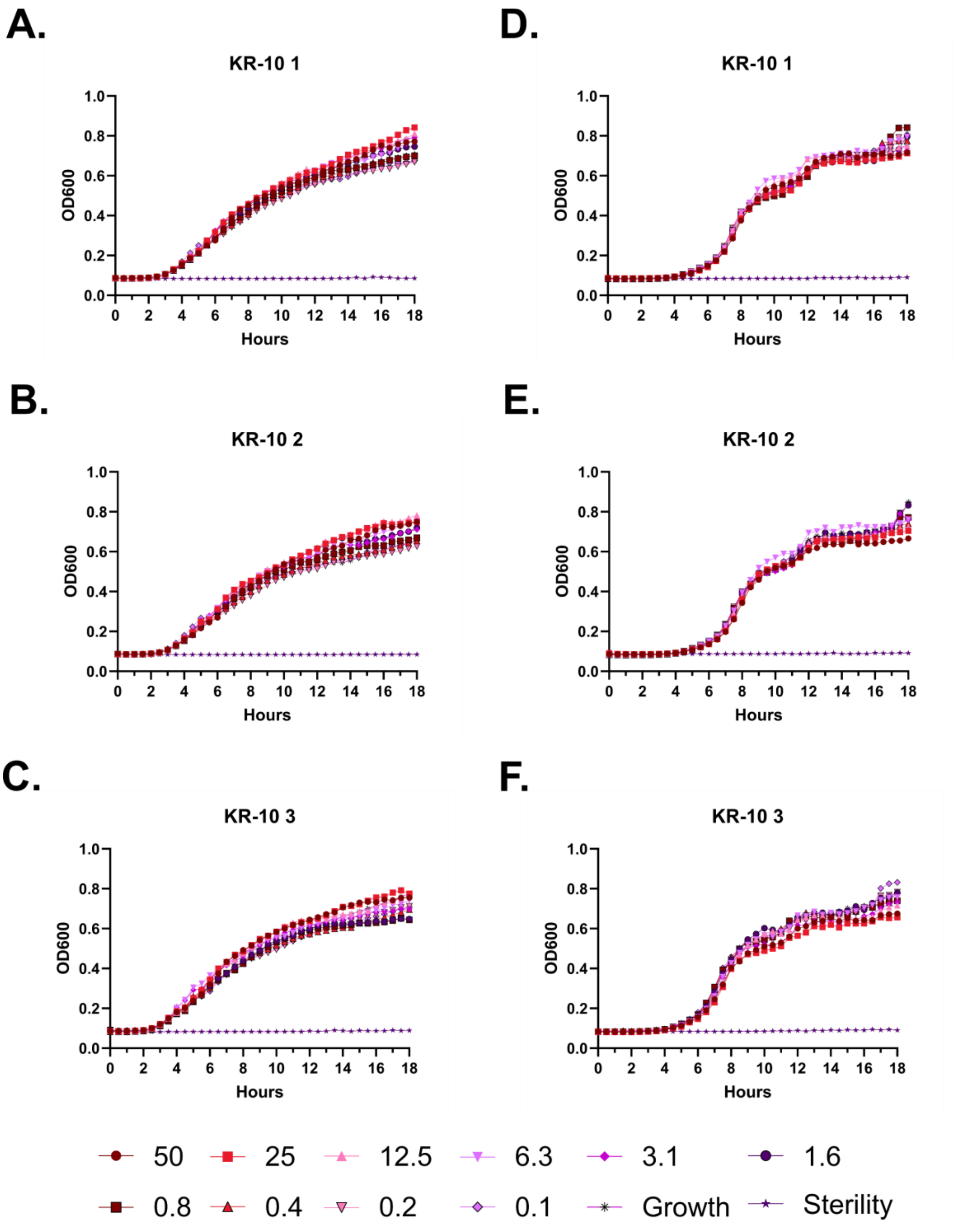
A.
KR-10 1
D.
KR-10 1
B.
KR-10 2
E.
KR-10 2
C.
KR-10 3
F.
KR-10 3
OD600
Hours
50
25
12.5
6.3
3.1
1.6
0.8
0.4
0.2
0.1
Growth
Sterility

**Figure S13 – Antimicrobial susceptibility curves for (#11) KR-10. A-C.** Independent experiments with *E. coli*. **D-F.** Independent experiments with *P. aeruginosa.* Growth over 18 hours at 37 °C with a gradient of peptide concentrations given in μM.

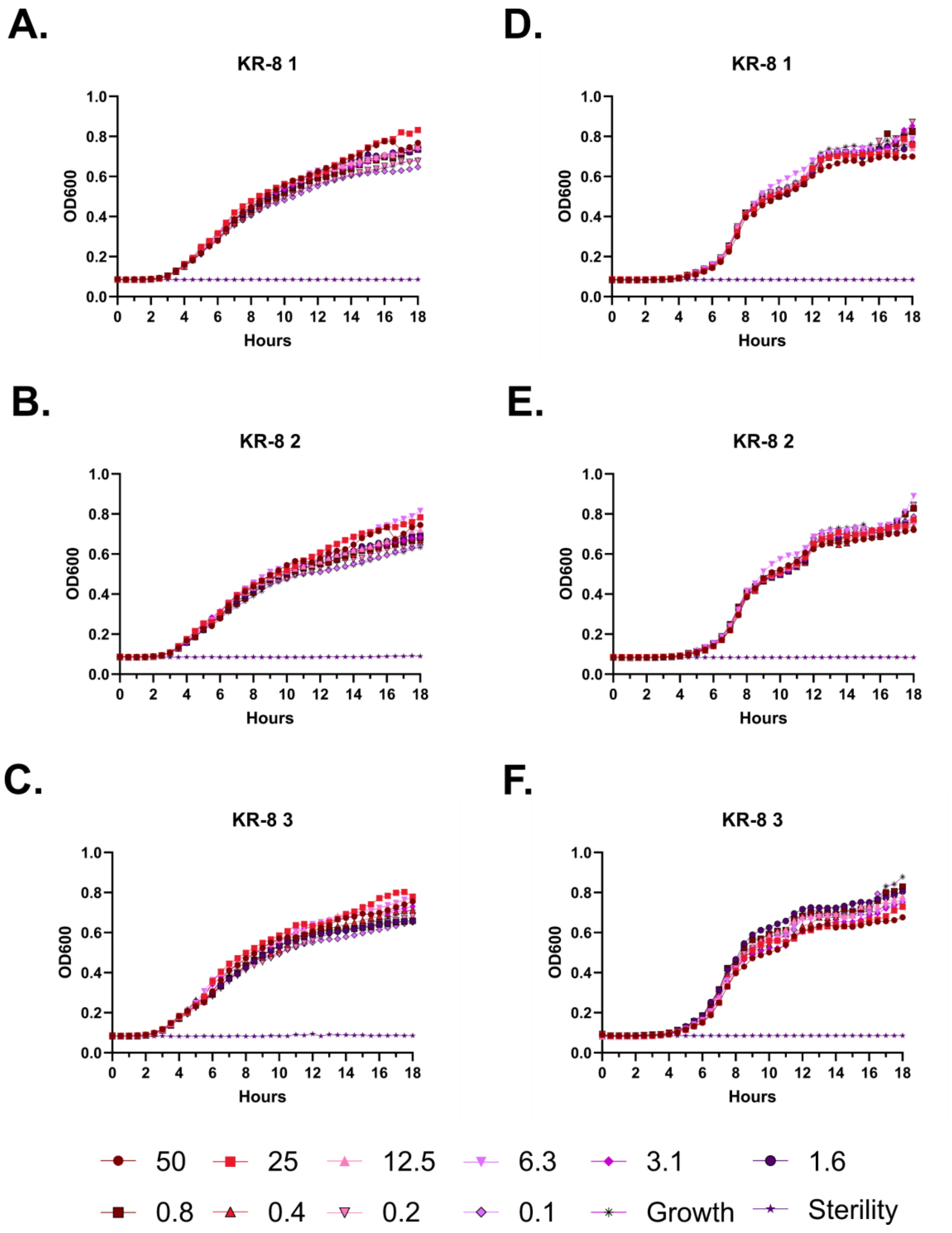
A.
KR-8 1
B.
KR-8 2
C.
KR-8 3
D.
KR-8 1
E.
KR-8 2
F.
KR-8 3
OD600
Hours
50
25
12.5
6.3
3.1
1.6
0.8
0.4
0.2
0.1
Growth
Sterility

**Figure S14 – Antimicrobial susceptibility curves for (#12) KR-8. A-C.** Independent experiments with *E. coli*. **D-F.** Independent experiments with *P. aeruginosa.* Growth over 18 hours at 37 °C with a gradient of peptide concentrations given in μM.

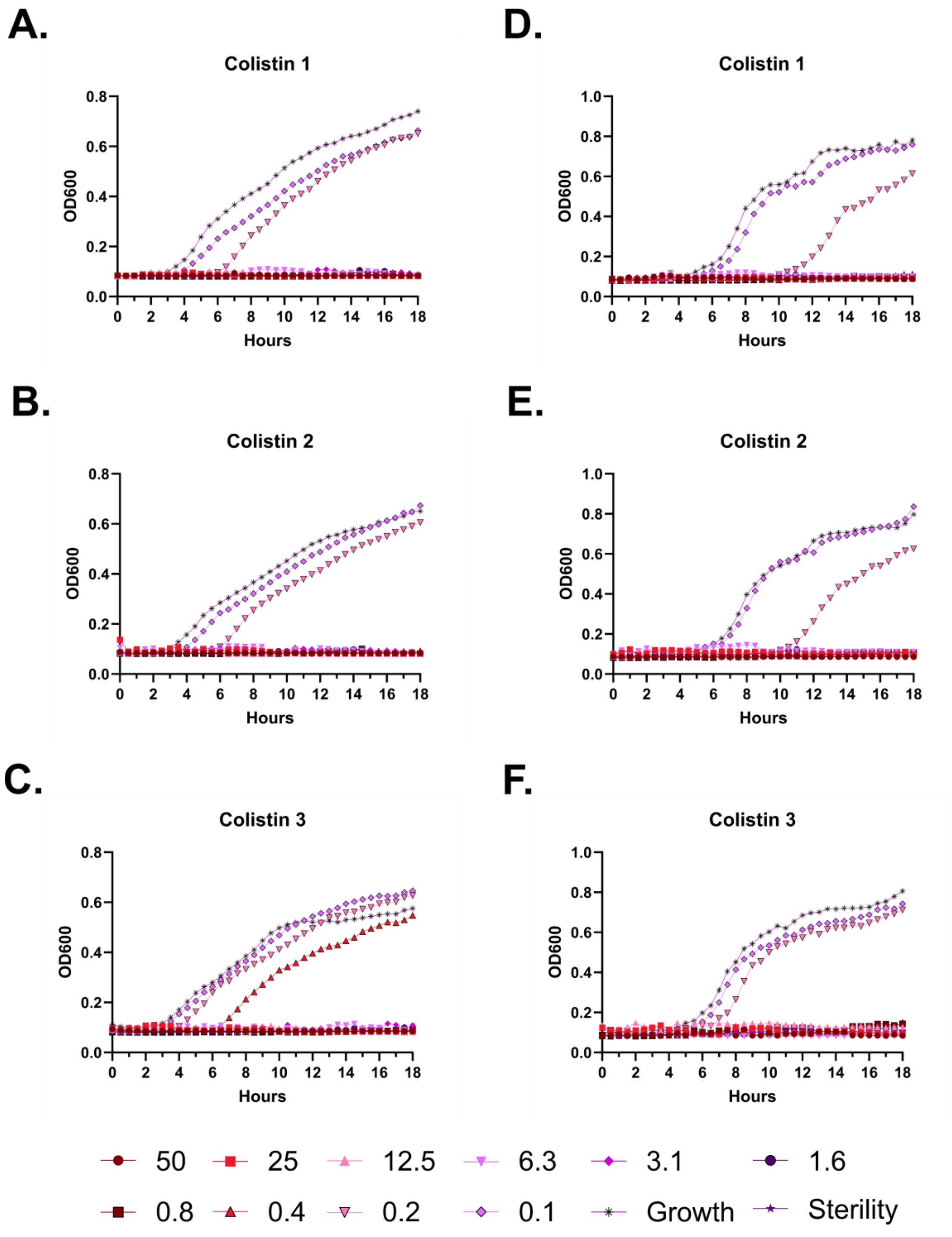
A.
Colistin 1
OD600
Hours
D.
Colistin 1
OD600
Hours
B.
Colistin 2
OD600
Hours
E.
Colistin 2
OD600
Hours
C.
Colistin 3
OD600
Hours
F.
Colistin 3
OD600
Hours
50
25
12.5
6.3
3.1
1.6
0.8
0.4
0.2
0.1
Growth
Sterility

**Figure S15 – Antimicrobial susceptibility curves for Colistin. A-C.** Independent experiments with *E. coli*. **D-F.** Independent experiments with *P. aeruginosa.* Growth over 18 hours at 37 °C with a gradient of peptide concentrations given in μM.

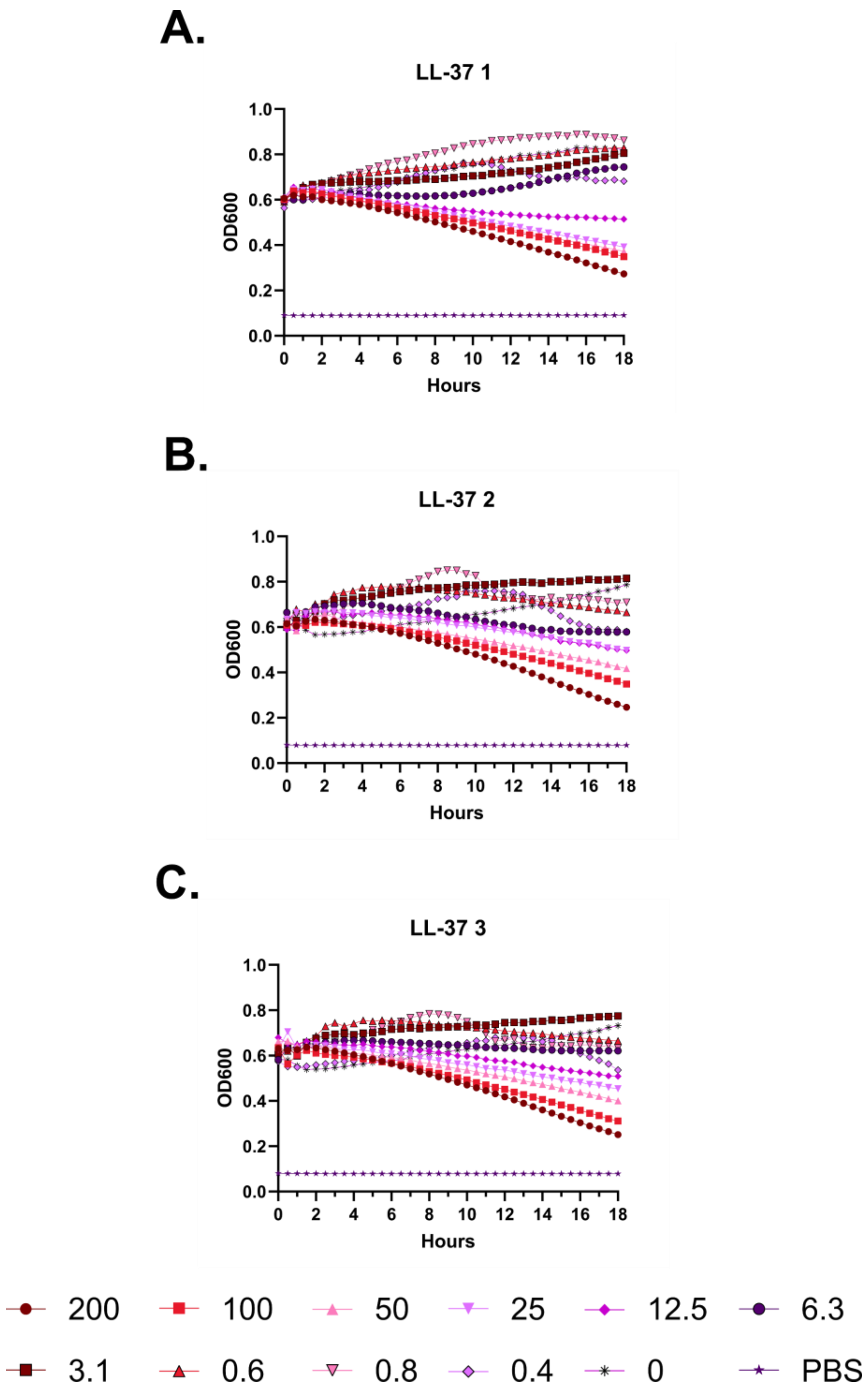
A.
LL-37 1
B.
LL-37 2
C.
LL-37 3
OD600
Hours
1.0
0.8
0.6
0.4
0.2
0.0
0
2
4
6
8
10
12
14
16
18
200
100
50
25
12.5
6.3
3.1
0.6
0.8
0.4
0
PBS

**Figure S16 – (#1) LL-37 kinetic hemolysis assay. A.** Defibrinated sheep blood was incubated in the presence of the indicated peptide concentrations at 37 °C with monitoring of OD600 over 18 hours, where decreasing OD600 indicates hemolysis and each panel indicates an independent experiment. Peptide concentrations are in μM.

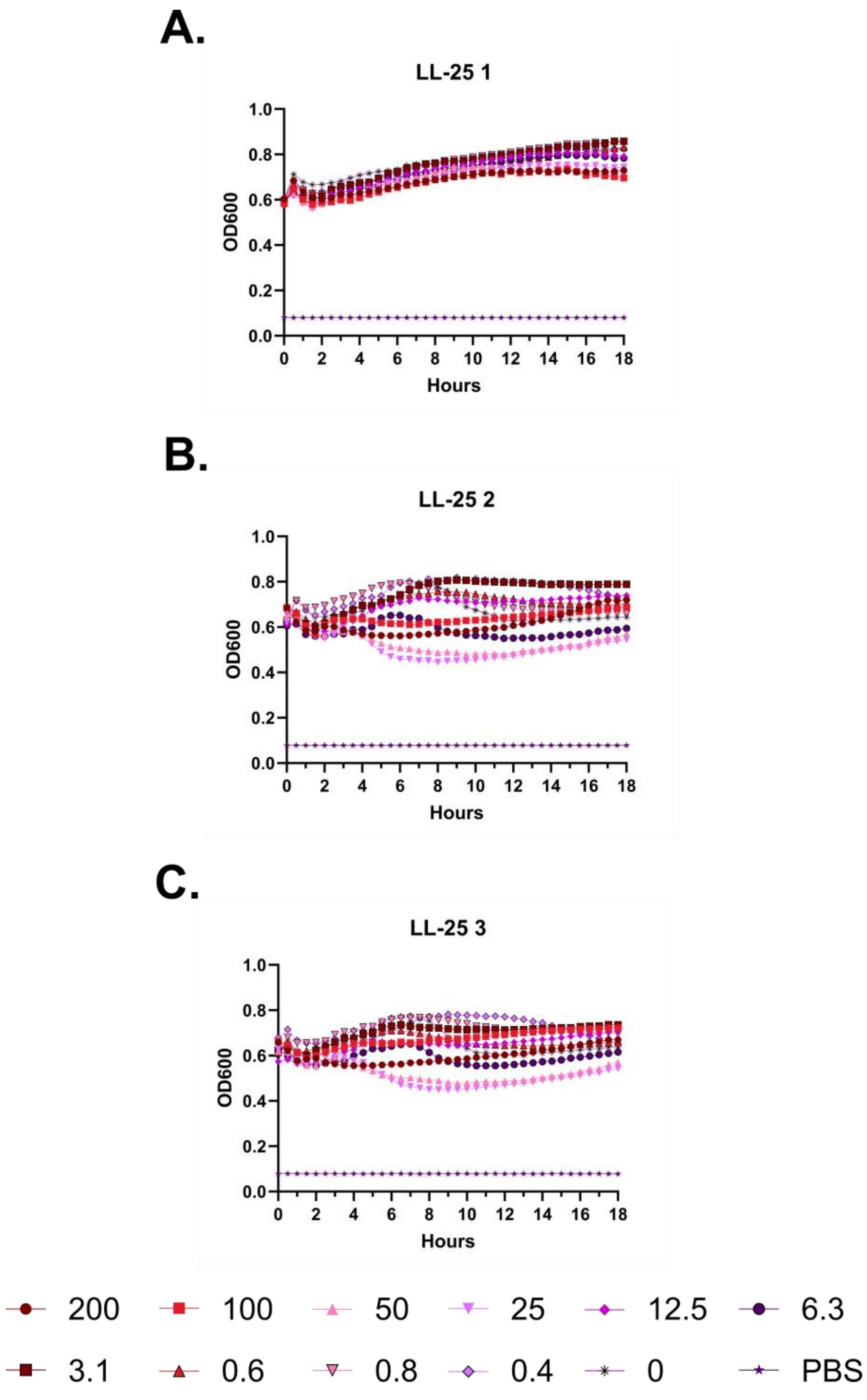
A.
LL-25 1
OD600
Hours
B.
LL-25 2
OD600
Hours
C.
LL-25 3
OD600
Hours
200
100
50
25
12.5
6.3
3.1
0.6
0.8
0.4
0
PBS

**Figure S17 –(#2) LL-25 kinetic hemolysis assay. A.** Defibrinated sheep blood was incubated in the presence of the indicated peptide concentrations at 37 °C with monitoring of OD600 over 18 hours, where decreasing OD600 indicates hemolysis and each panel indicates an independent experiment. Peptide concentrations are in μM.

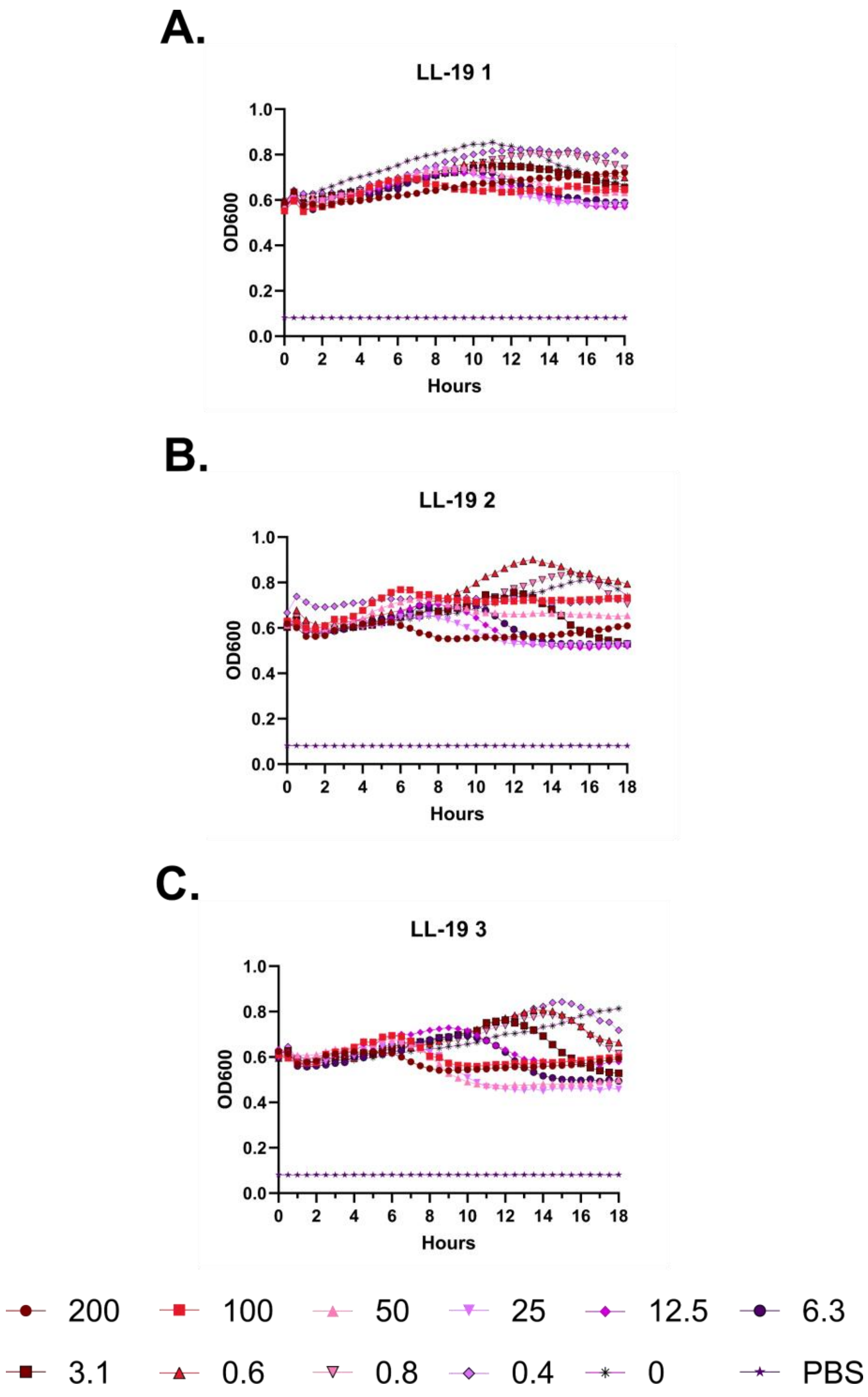
A.
LL-19 1
OD600
Hours
B.
LL-19 2
OD600
Hours
C.
LL-19 3
OD600
Hours
200
100
50
25
12.5
6.3
3.1
0.6
0.8
0.4
0
PBS

**Figure S18 – (#3) LL-19 kinetic hemolysis assay. A.** Defibrinated sheep blood was incubated in the presence of the indicated peptide concentrations at 37 °C with monitoring of OD600 over 18 hours, where decreasing OD600 indicates hemolysis and each panel indicates an independent experiment. Peptide concentrations are in μM.

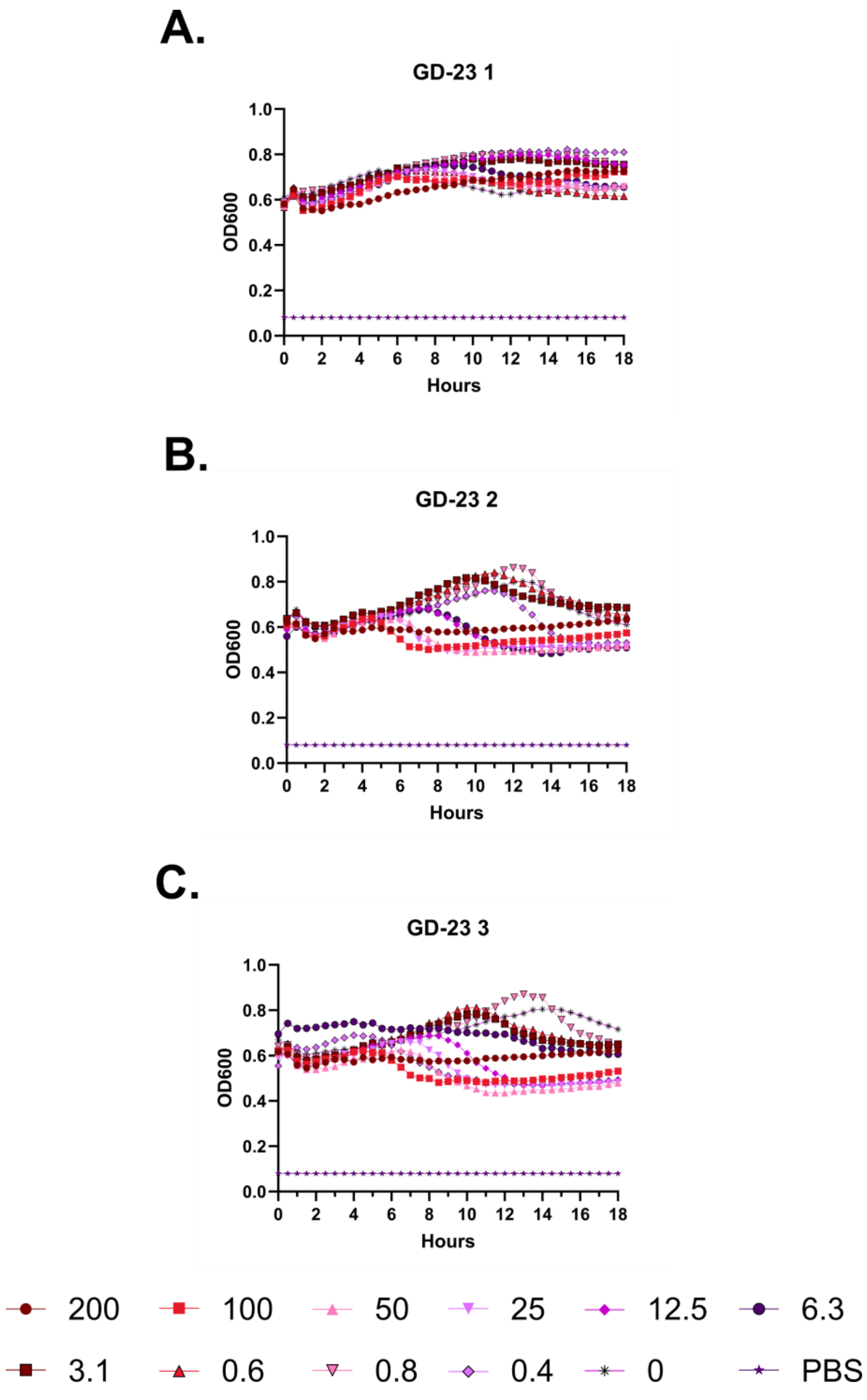
A.
GD-23 1
B.
GD-23 2
C.
GD-23 3
OD600
Hours
1.0
0.8
0.6
0.4
0.2
0.0
0
2
4
6
8
10
12
14
16
18
200
100
50
25
12.5
6.3
3.1
0.6
0.8
0.4
0
PBS

**Figure S19 – (#4) GD-23 kinetic hemolysis assay. A.** Defibrinated sheep blood was incubated in the presence of the indicated peptide concentrations at 37 °C with monitoring of OD600 over 18 hours, where decreasing OD600 indicates hemolysis and each panel indicates an independent experiment. Peptide concentrations are in μM.

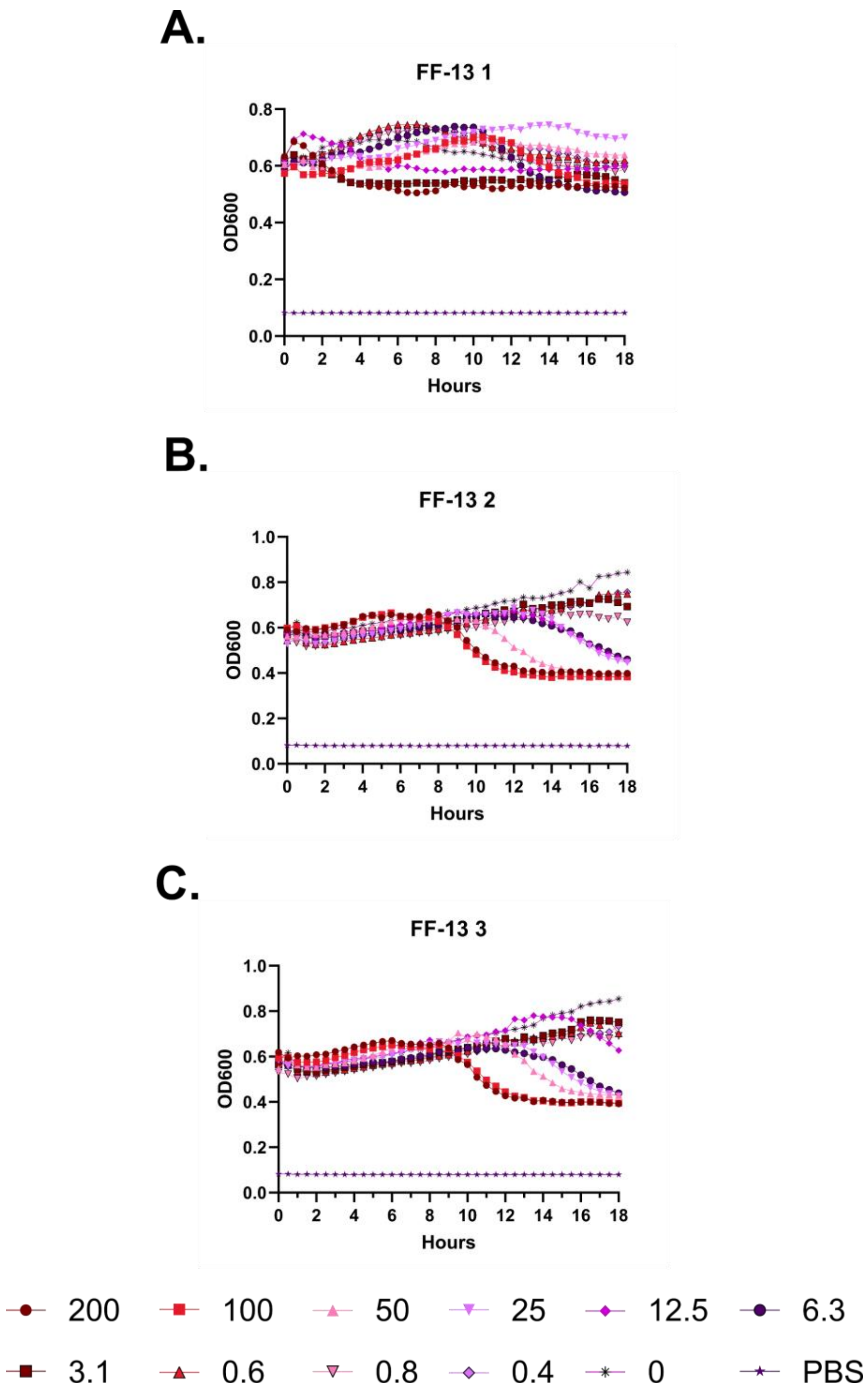
A.
FF-13 1
OD600
Hours
B.
FF-13 2
OD600
Hours
C.
FF-13 3
OD600
Hours
200
100
50
25
12.5
6.3
3.1
0.6
0.8
0.4
0
PBS

**Figure S20 – (#5) FF-13 kinetic hemolysis assay. A.** Defibrinated sheep blood was incubated in the presence of the indicated peptide concentrations at 37 °C with monitoring of OD600 over 18 hours, where decreasing OD600 indicates hemolysis and each panel indicates an independent experiment. Peptide concentrations are in μM.

A.

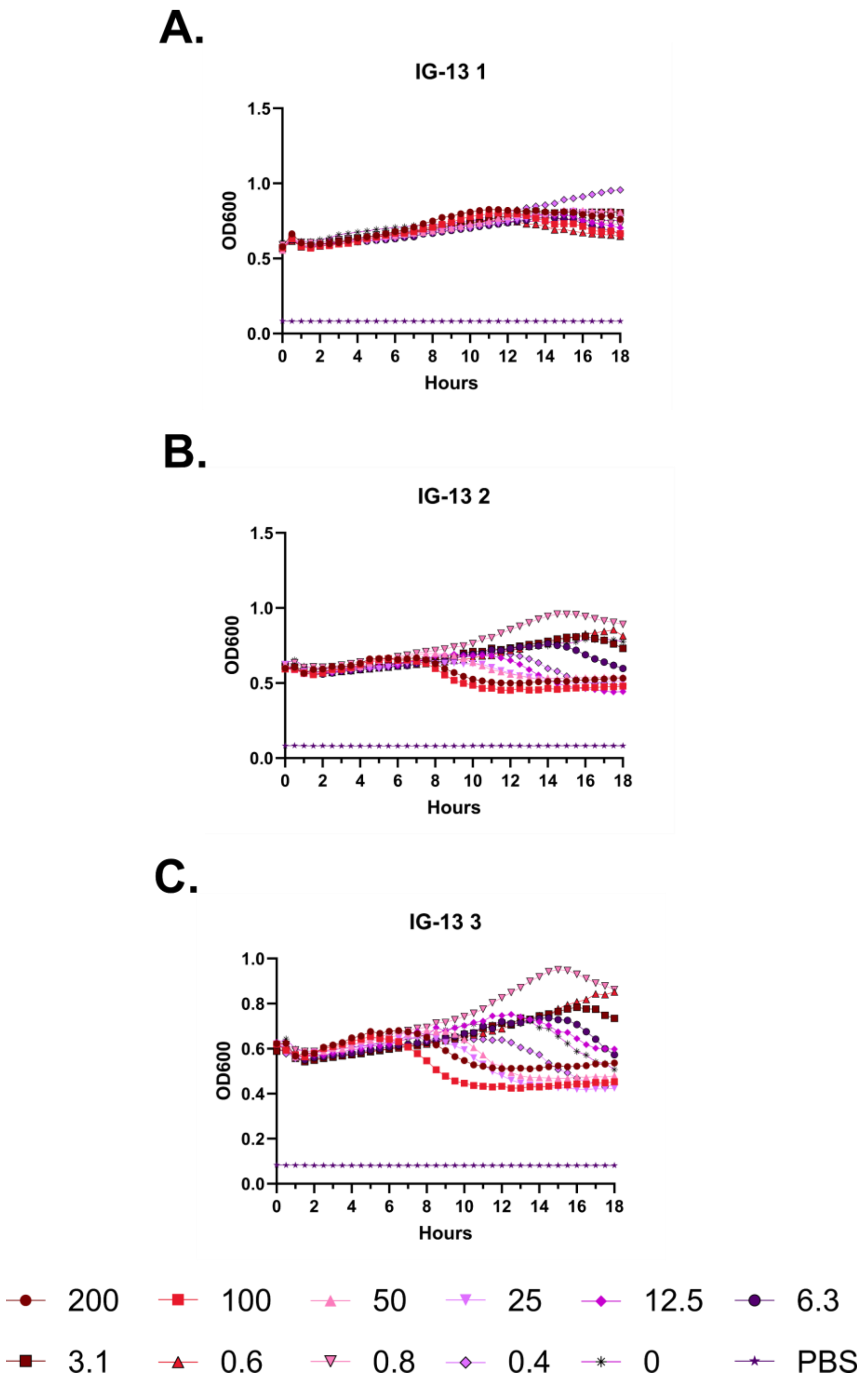
A.
IG-13 1
OD600
Hours
B.
IG-13 2
OD600
Hours
C.
IG-13 3
OD600
Hours
200
100
50
25
12.5
6.3
3.1
0.6
0.8
0.4
0
PBS

**Figure S21 – (#6) IG-13 kinetic hemolysis assay. A.** Defibrinated sheep blood was incubated in the presence of the indicated peptide concentrations at 37 °C with monitoring of OD600 over 18 hours, where decreasing OD600 indicates hemolysis and each panel indicates an independent experiment. Peptide concentrations are in μM.

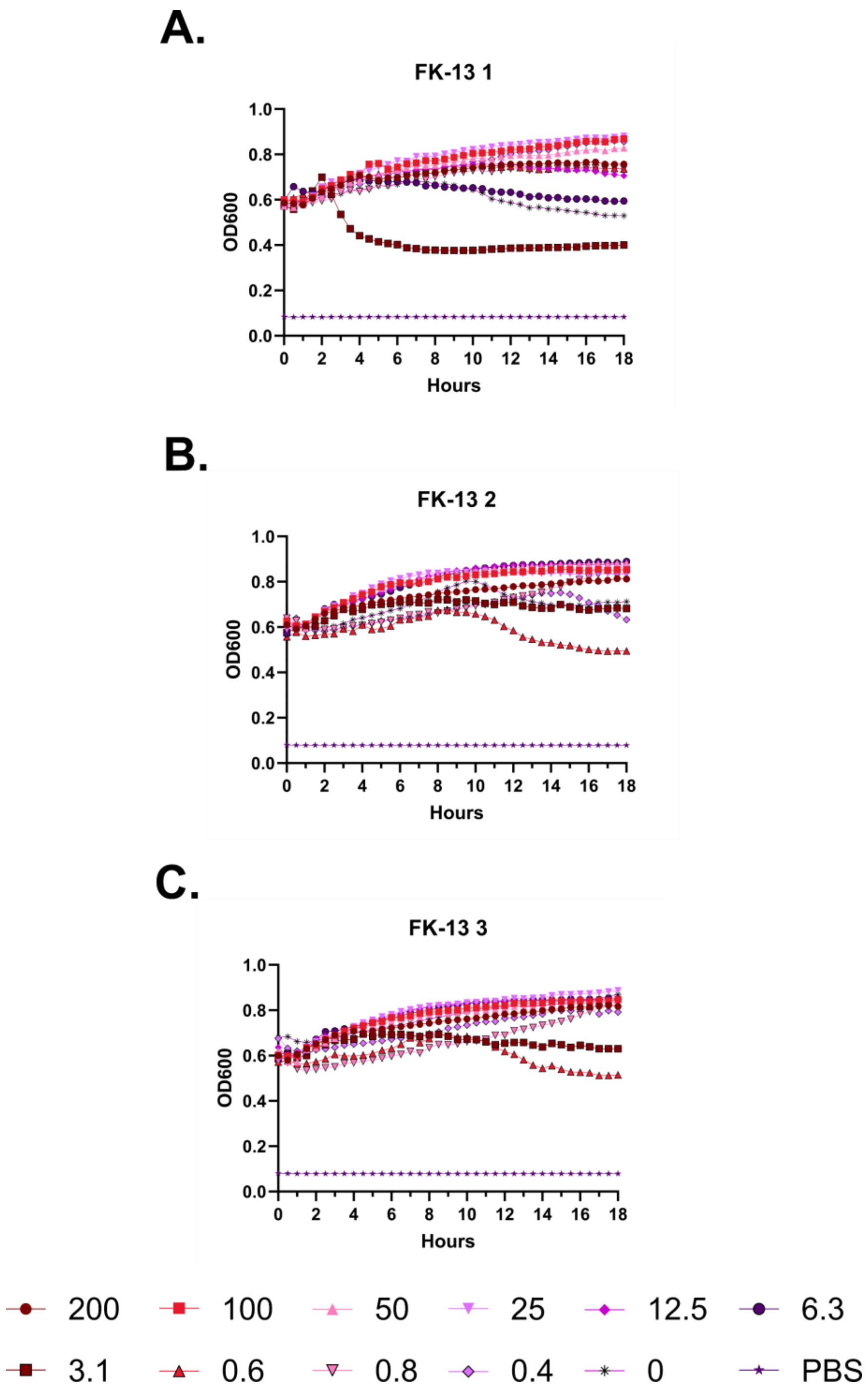
A.
FK-13 1
OD600
Hours
B.
FK-13 2
OD600
Hours
C.
FK-13 3
OD600
Hours
200
100
50
25
12.5
6.3
3.1
0.6
0.8
0.4
0
PBS

**Figure S22 – (#7) FK-13 kinetic hemolysis assay. A.** Defibrinated sheep blood was incubated in the presence of the indicated peptide concentrations at 37 °C with monitoring of OD600 over 18 hours, where decreasing OD600 indicates hemolysis and each panel indicates an independent experiment. Peptide concentrations are in μM.

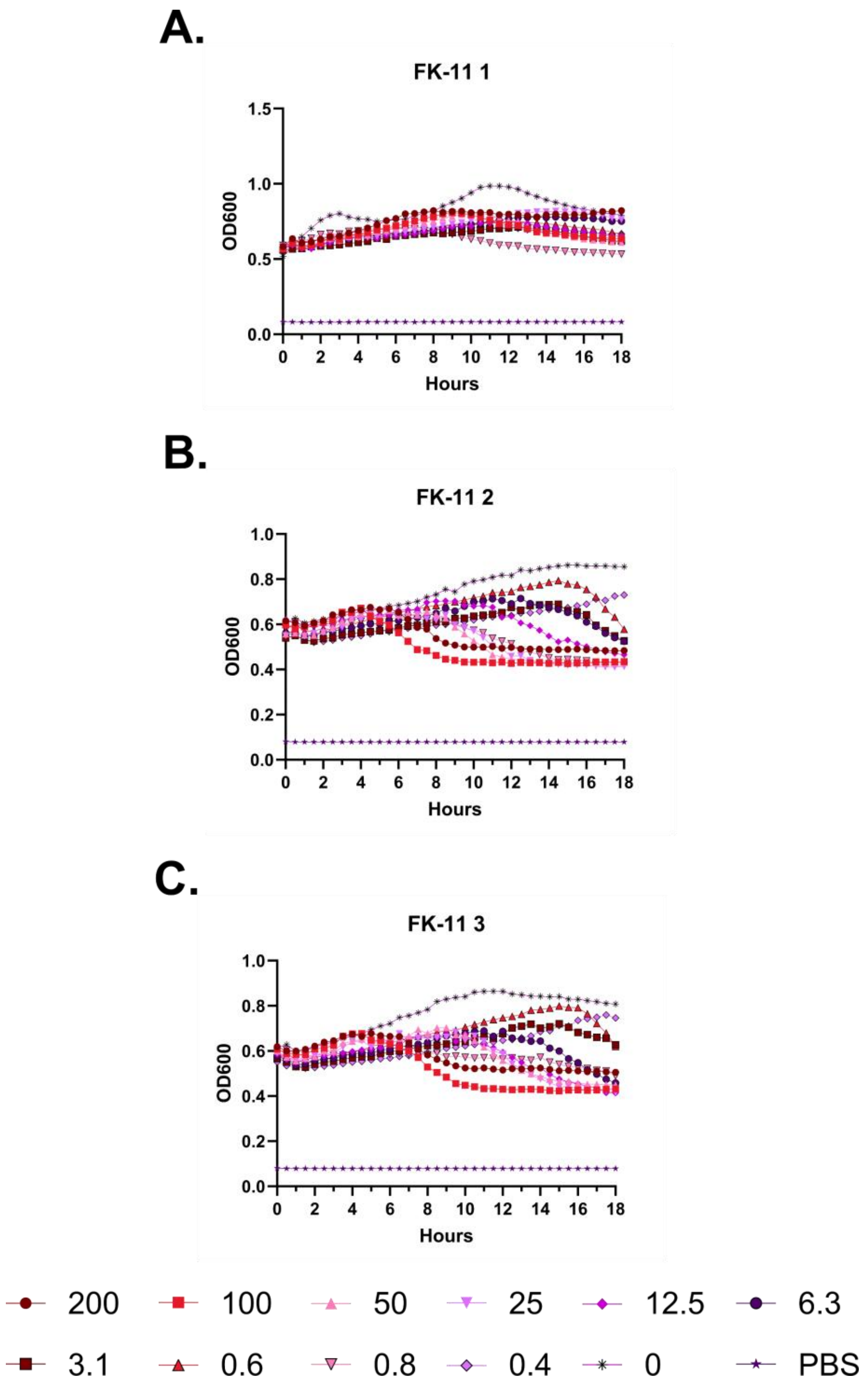
A.
FK-11 1
1.5
1.0
0.5
0.0
OD600
0
2
4
6
8
10
12
14
16
18
Hours
B.
FK-11 2
1.0
0.8
0.6
0.4
0.2
0.0
OD600
0
2
4
6
8
10
12
14
16
18
Hours
C.
FK-11 3
1.0
0.8
0.6
0.4
0.2
0.0
OD600
0
2
4
6
8
10
12
14
16
18
Hours
200
100
50
25
12.5
6.3
3.1
0.6
0.8
0.4
0
PBS

**Figure S23 – (#8) FK-11 kinetic hemolysis assay. A.** Defibrinated sheep blood was incubated in the presence of the indicated peptide concentrations at 37 °C with monitoring of OD600 over 18 hours, where decreasing OD600 indicates hemolysis and each panel indicates an independent experiment. Peptide concentrations are in μM.

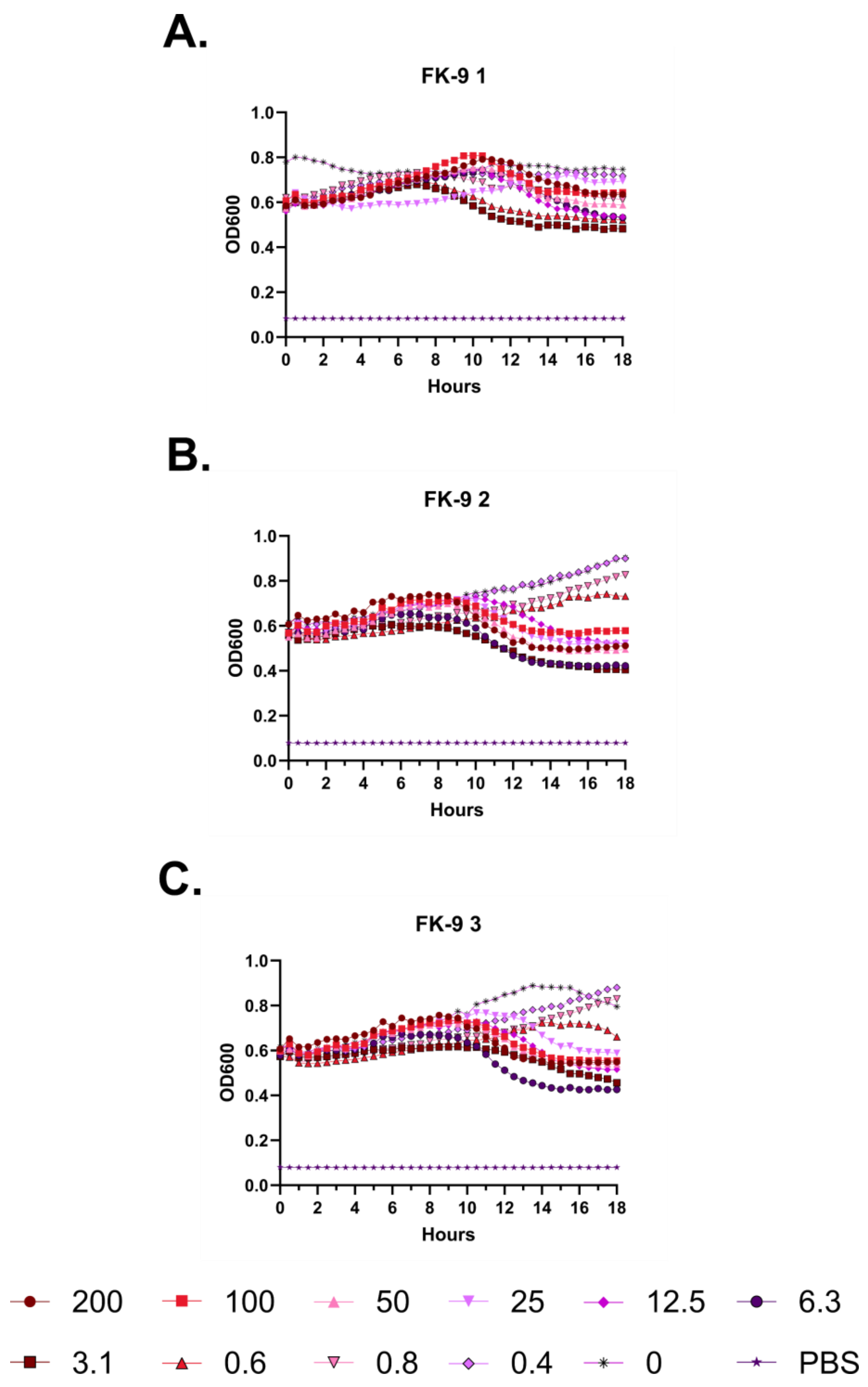
A.
FK-9 1
OD600
Hours
B.
FK-9 2
OD600
Hours
C.
FK-9 3
OD600
Hours
200
100
50
25
12.5
6.3
3.1
0.6
0.8
0.4
0
PBS

**Figure S24 – (#9) FK-9 kinetic hemolysis assay. A.** Defibrinated sheep blood was incubated in the presence of the indicated peptide concentrations at 37 °C with monitoring of OD600 over 18 hours, where decreasing OD600 indicates hemolysis and each panel indicates an independent experiment. Peptide concentrations are in μM.

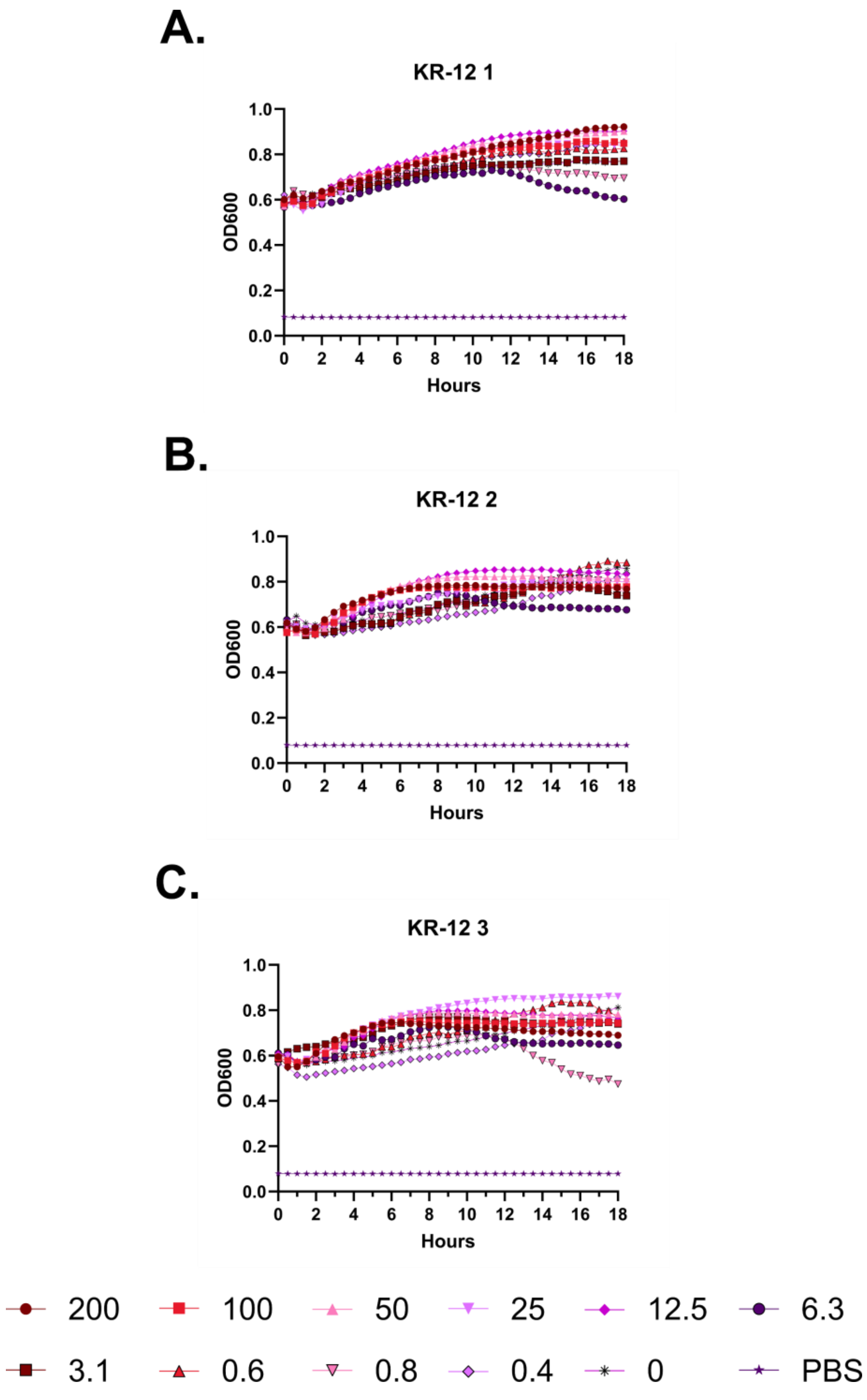
A.
KR-12 1
OD600
Hours
B.
KR-12 2
OD600
Hours
C.
KR-12 3
OD600
Hours
200
100
50
25
12.5
6.3
3.1
0.6
0.8
0.4
0
PBS

**Figure S25 – (#10) KR-12 kinetic hemolysis assay. A.** Defibrinated sheep blood was incubated in the presence of the indicated peptide concentrations at 37 °C with monitoring of OD600 over 18 hours, where decreasing OD600 indicates hemolysis and each panel indicates an independent experiment. Peptide concentrations are in µM.

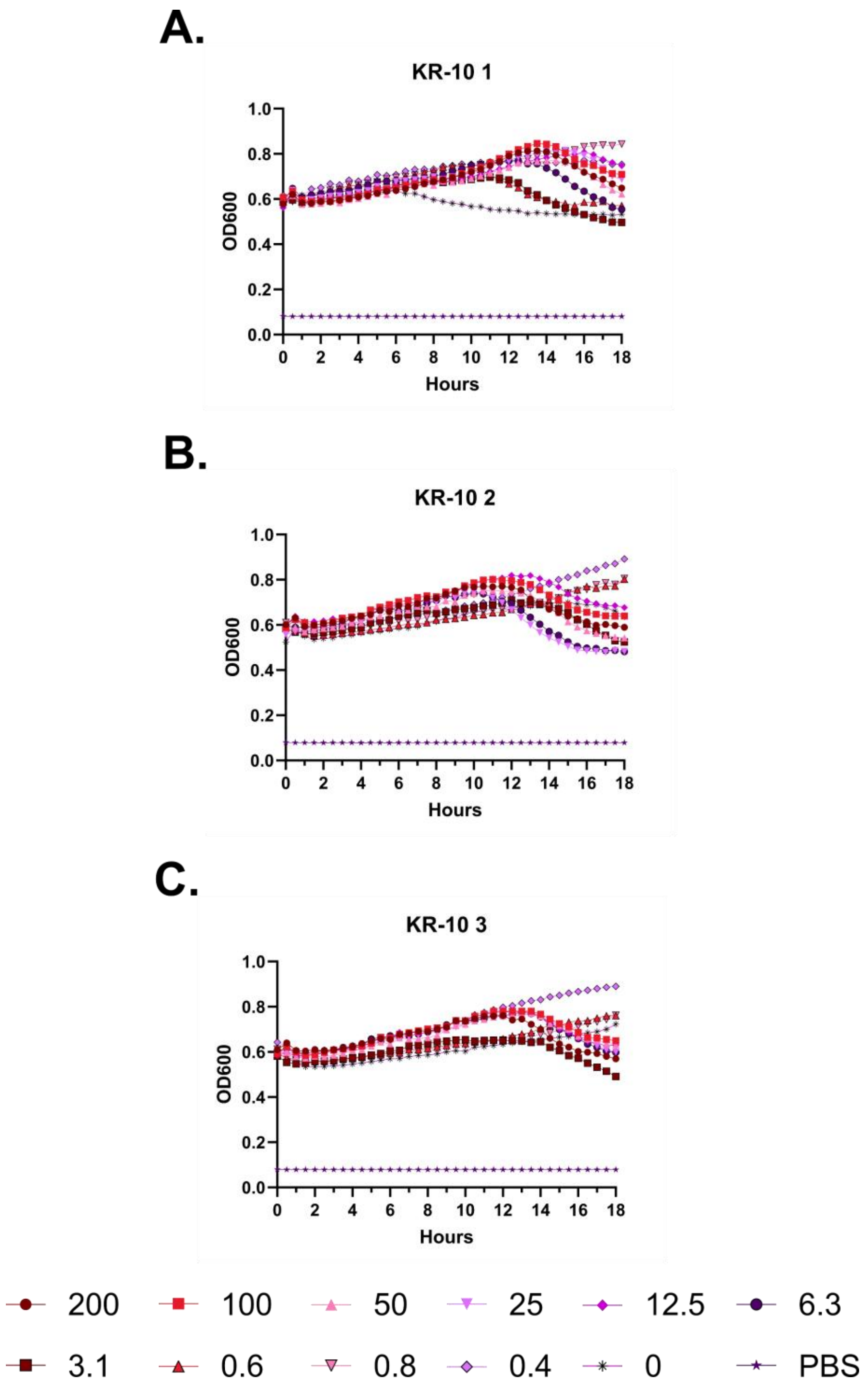
A.
KR-10 1
OD600
Hours
B.
KR-10 2
OD600
Hours
C.
KR-10 3
OD600
Hours
1.0
0.8
0.6
0.4
0.2
0.0
0
2
4
6
8
10
12
14
16
18
200
100
50
25
12.5
6.3
3.1
0.6
0.8
0.4
0
PBS

**Figure S26 – (#11) KR-10 kinetic hemolysis assay. A.** Defibrinated sheep blood was incubated in the presence of the indicated peptide concentrations at 37 °C with monitoring of OD600 over 18 hours, where decreasing OD600 indicates hemolysis and each panel indicates an independent experiment. Peptide concentrations are in µM.

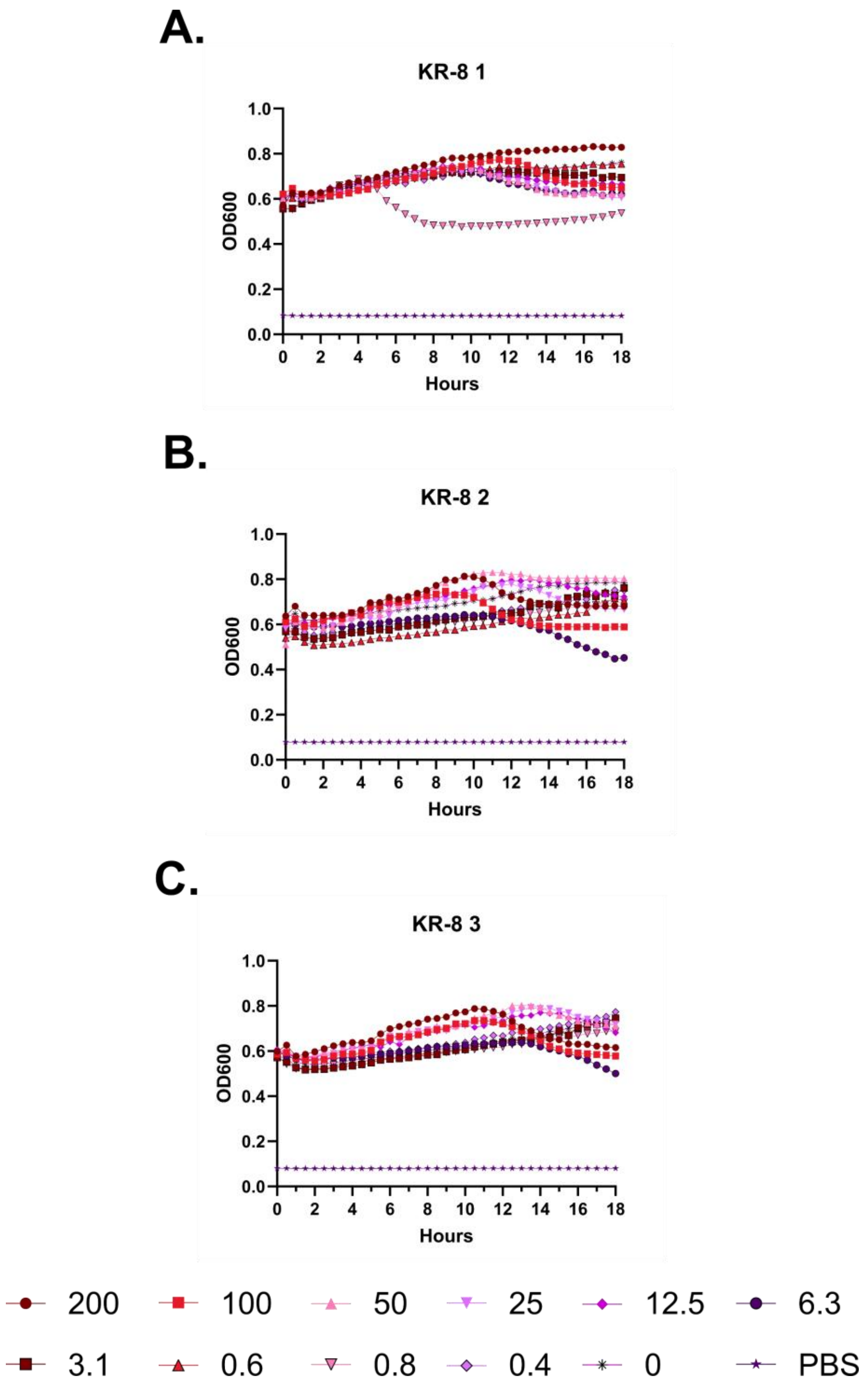
A.
KR-8 1
OD600
Hours
B.
KR-8 2
OD600
Hours
C.
KR-8 3
OD600
Hours
200
100
50
25
12.5
6.3
3.1
0.6
0.8
0.4
0
PBS

**Figure S27 – (#12) KR-8 kinetic hemolysis assay. A.** Defibrinated sheep blood was incubated in the presence of the indicated peptide concentrations at 37 °C with monitoring of OD600 over 18 hours, where decreasing OD600 indicates hemolysis and each panel indicates an independent experiment. Peptide concentrations are in μM.

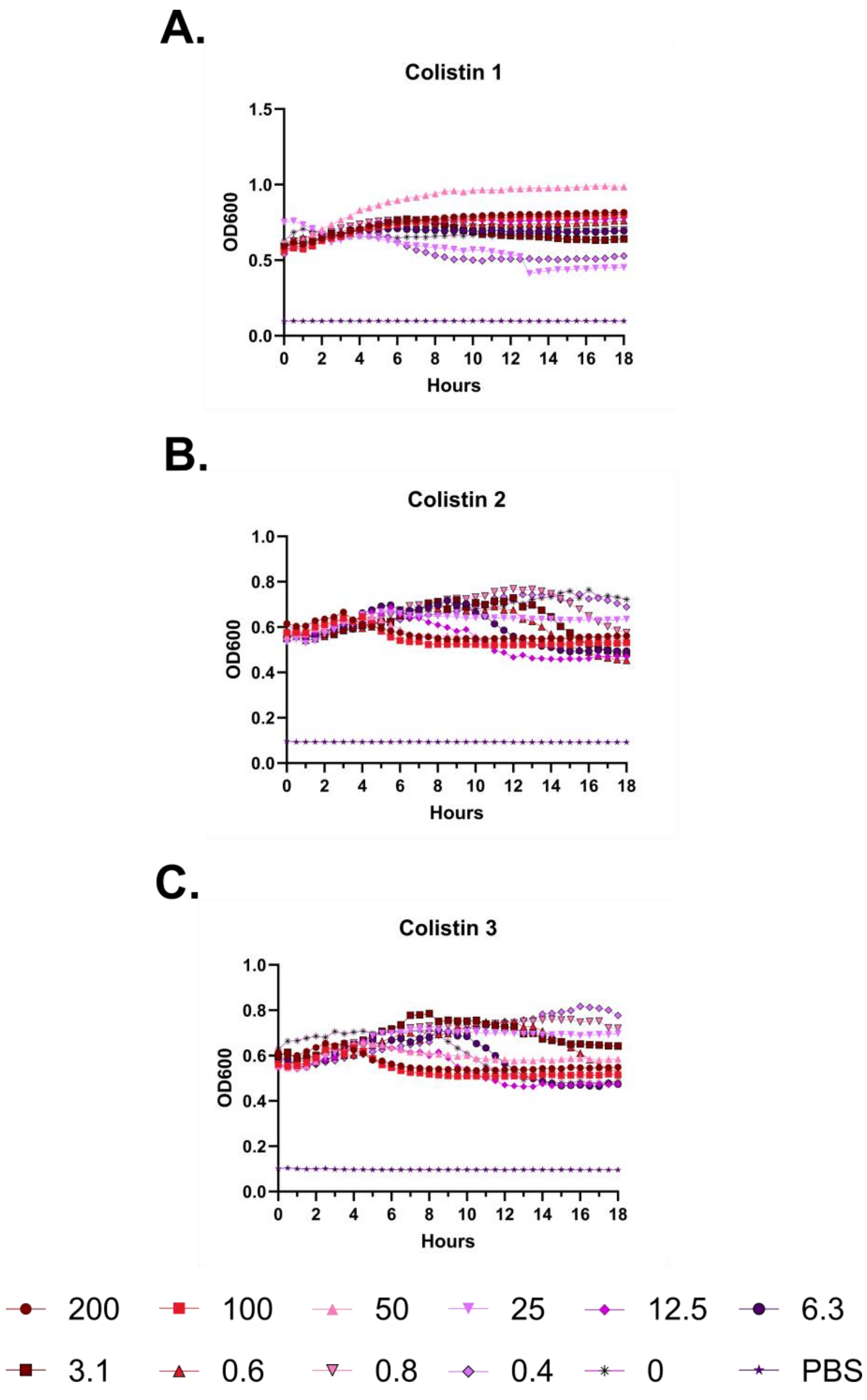
A.
Colistin 1
1.5
1.0
0.5
0.0
OD600
0 2 4 6 8 10 12 14 16 18
Hours
B.
Colistin 2
1.0
0.8
0.6
0.4
0.2
0.0
OD600
0 2 4 6 8 10 12 14 16 18
Hours
C.
Colistin 3
1.0
0.8
0.6
0.4
0.2
0.0
OD600
0 2 4 6 8 10 12 14 16 18
Hours
200
100
50
25
12.5
6.3
3.1
0.6
0.8
0.4
0
PBS

**Figure S28 – Colistin kinetic hemolysis assay. A.** Defibrinated sheep blood was incubated in the presence of the indicated peptide concentrations at 37 °C with monitoring of OD600 over 18 hours, where decreasing OD600 indicates hemolysis and each panel indicates an independent experiment. Peptide concentrations are in μM.

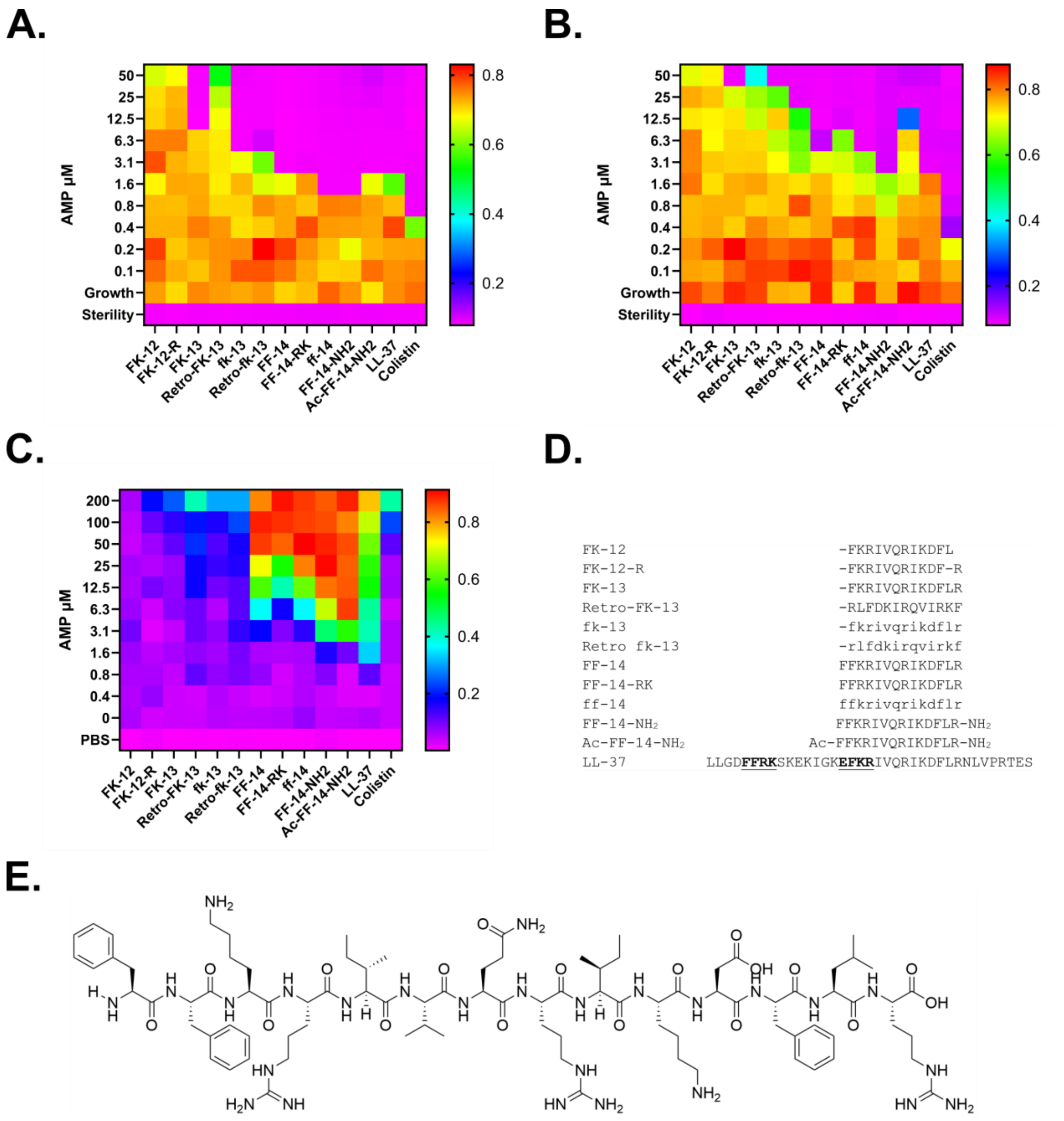
A.
B.
C.
D.
E.
AMP µM
50
25
12.5
6.3
3.1
1.6
0.8
0.4
0.2
0.1
Growth
Sterility
200
100
0
PBS
FK-12
FK-12-R
FK-13
Retro-FK-13
fk-13
Retro-fk-13
FF-14
FF-14-RK
ff-14
FF-14-NH2
Ac-FF-14-NH2
LL-37
Colistin
FK-12 -FKRIVQRIKDFL
FK-12-R -FKRIVQRIKDF-R
FK-13 -FKRIVQRIKDFLR
Retro-FK-13 -RLFDKIRQVIRKF
fk-13 -fkrivqrikdflr
Retro fk-13 -rlfdkirqvirkf
FF-14 FFKRIVQRIKDFLR
FF-14-RK FFRKIVQRIKDFLR
ff-14 ffkrivqrikdflr
FF-14-NH2 FFKRIVQRIKDFLR-NH2
Ac-FF-14-NH2 Ac-FFKRIVQRIKDFLR-NH2
LL-37 LLGDFFRKSKEKIGKEFKRIVQRIKDFLRNLVPRTES

**Figure S29 – Heatmap depiction of functional data in Figure 2. A.** Antimicrobial susceptibility testing showing the activity of each derivative as well as a colistin control against *E. coli* (ATCC 25922). In the heatmap plate depiction, cooler colors indicate low or absent bacterial growth as detected by the OD600 measure at 18 hours of incubation at 37 °C. **B.** As in **A.**, but with *P. aeruginosa* (ATCC 27853). **C.** Endpoint hemolysis assays with defibrinated sheep blood following incubation with the indicated peptide for 18 hours at 37 °C with subsequent quantification of supernatant free hemoglobin at 414 nm. Cool colors indicate little or no release of hemoglobin. Levels throughout show the median across three independent experiments. **D.** Alignment showing the sequence of (#1) LL-37 in comparison with shorter derivatives tested in these studies. The FF[RK] region transposed to the central activity region on which (#10) KR-12 and (#7) FK-13 are built is shown in bold with highlighting. **E.** Structure of (#18) FF-14.

A.

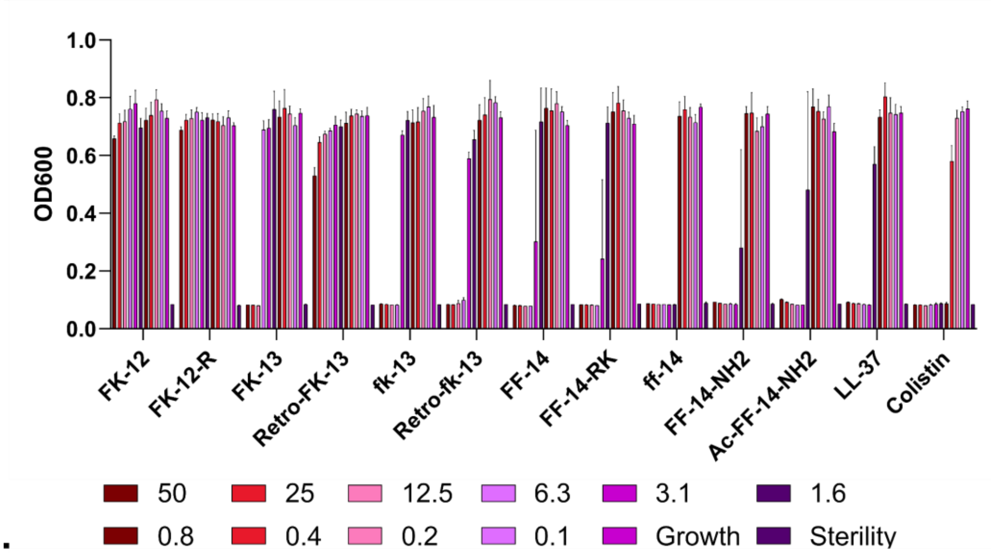
OD600
1.0
0.8
0.6
0.4
0.2
0.0
FK-12
FK-12-R
FK-13
Retro-FK-13
fk-13
Retro-fk-13
FF-14
FF-14-RK
ff-14
FF-14-NH2
Ac-FF-14-NH2
LL-37
Colistin
50
25
12.5
6.3
3.1
1.6
0.8
0.4
0.2
0.1
Growth
Sterility

B.

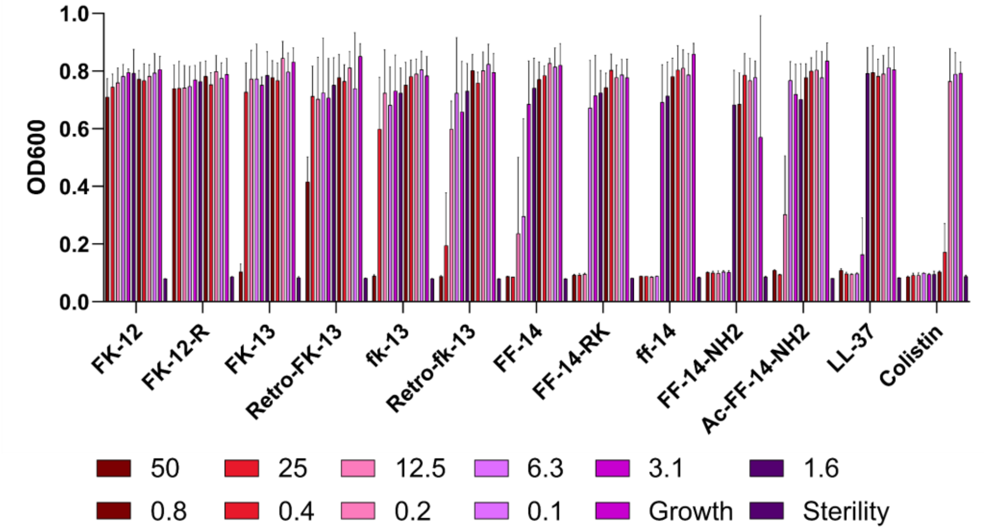
OD600
1.0
0.8
0.6
0.4
0.2
0.0
FK-12
FK-12-R
FK-13
Retro-FK-13
fk-13
Retro-fk-13
FF-14
FF-14-RK
ff-14
FF-14-NH2
Ac-FF-14-NH2
LL-37
Colistin
50
25
12.5
6.3
3.1
1.6
0.8
0.4
0.2
0.1
Growth
Sterility

C.

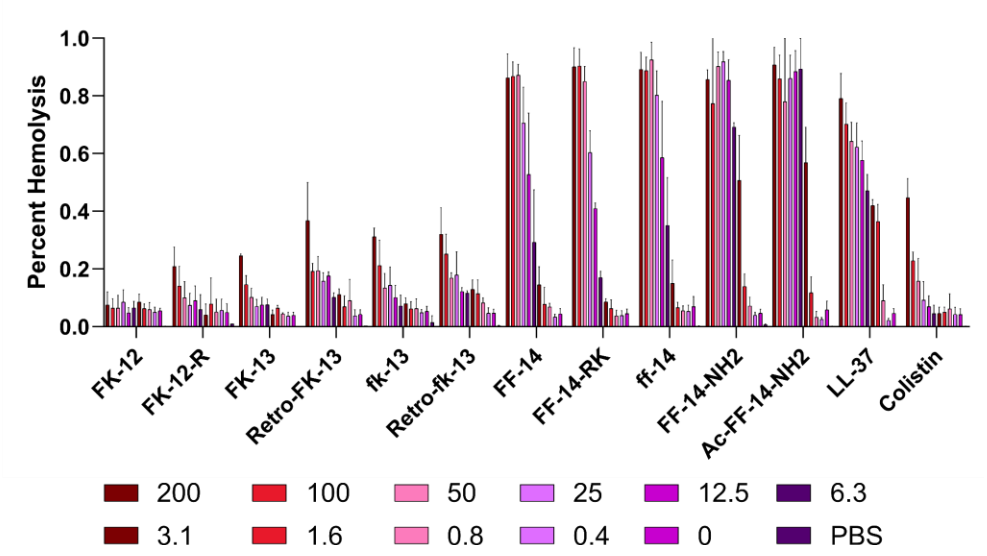
Percent Hemolysis
1.0
0.8
0.6
0.4
0.2
0.0
FK-12
FK-12-R
FK-13
Retro-FK-13
fk-13
Retro-fk-13
FF-14
FF-14-RK
ff-14
FF-14-NH2
Ac-FF-14-NH2
LL-37
Colistin
200
100
50
25
12.5
6.3
3.1
1.6
0.8
0.4
0
PBS

**Figure S30 – Histograms corresponding to Figure 2**. **A.** Growth of *E. coli* (ATCC 25922) by OD600 at 18 hours at 37 °C. Error bars indicate the standard deviation of three independent experiments. **B.** As in **A.**, but with *P. aeruginosa* (ATCC 27853). **C.** Endpoint hemolysis assays after incubation for 18 hours at 37 °C with the indicated peptide. Error bars indicate the standard deviation of three independent experiments. All peptide levels are given in μM.

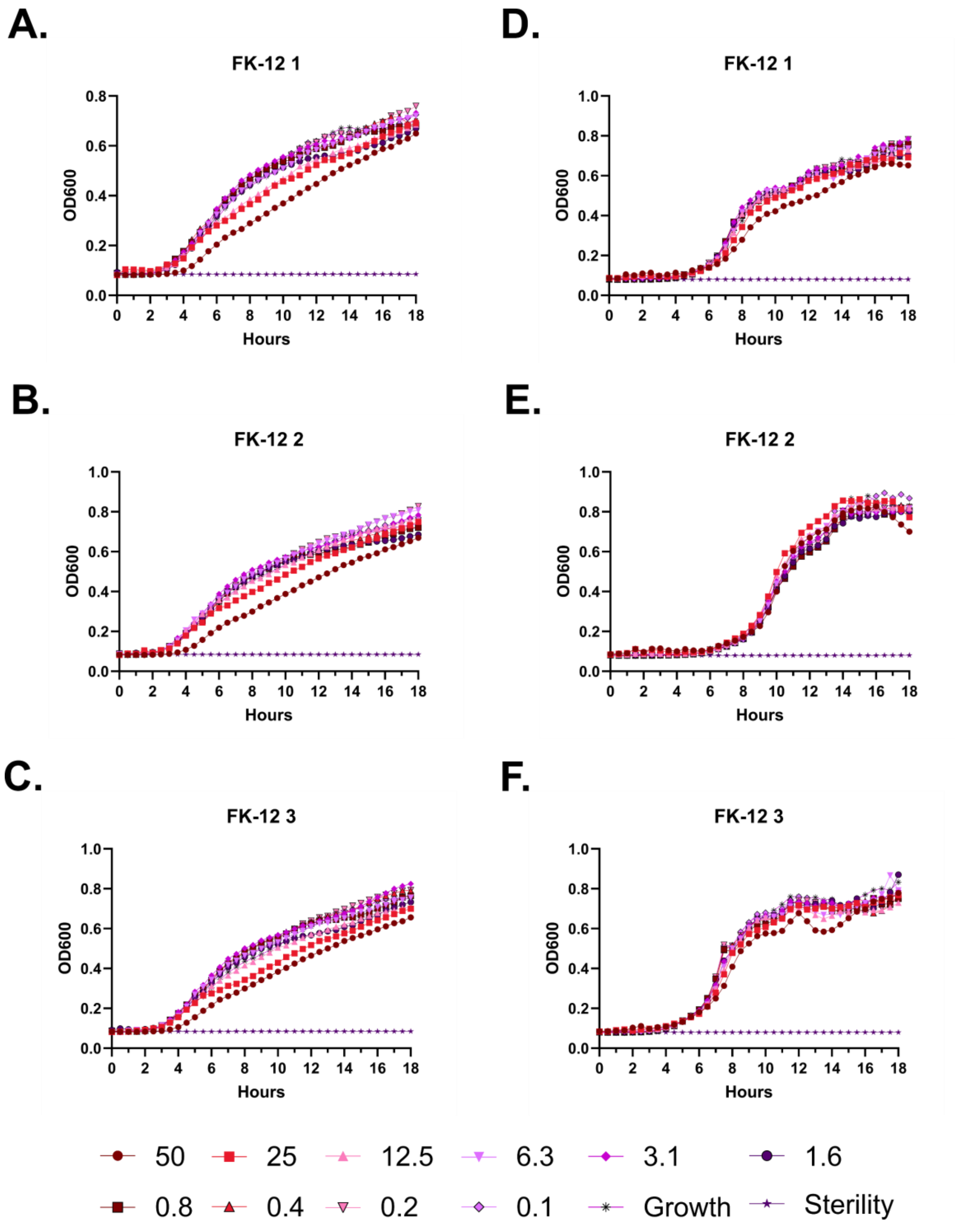
A.
FK-12 1
D.
FK-12 1
B.
FK-12 2
E.
FK-12 2
C.
FK-12 3
F.
FK-12 3
OD600
Hours
50
25
12.5
6.3
3.1
1.6
0.8
0.4
0.2
0.1
Growth
Sterility

**Figure S31 – Antimicrobial susceptibility curves for (#13) FK-12. A-C.** Independent experiments with *E. coli*. **D-F.** Independent experiments with *P. aeruginosa.* Growth over 18 hours at 37 °C with a gradient of peptide concentrations given in μM.

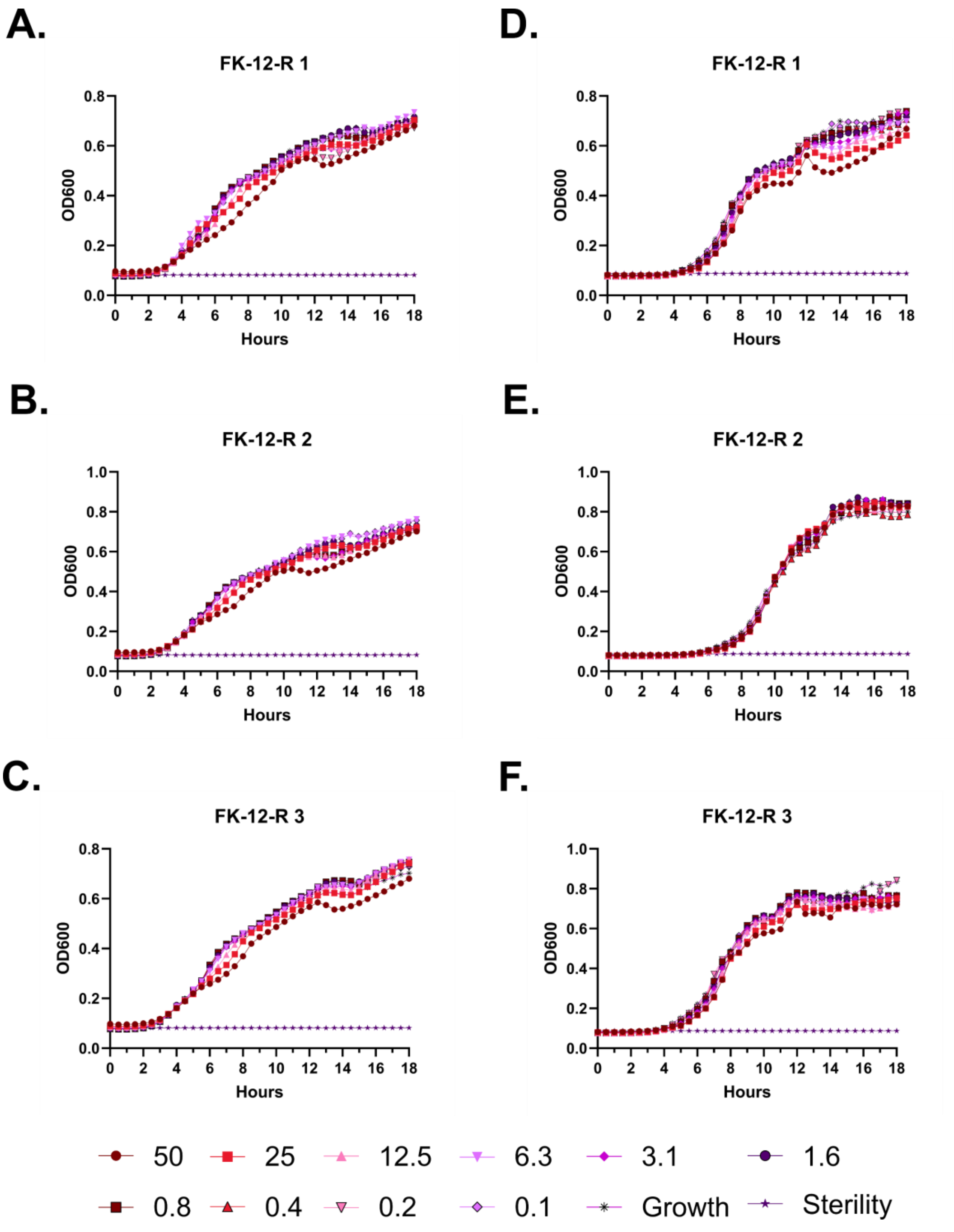
A.
FK-12-R 1
OD600
Hours
D.
FK-12-R 1
B.
FK-12-R 2
E.
FK-12-R 2
C.
FK-12-R 3
F.
FK-12-R 3
50
25
12.5
6.3
3.1
1.6
0.8
0.4
0.2
0.1
Growth
Sterility

**Figure S32 – Antimicrobial susceptibility curves for (#14) FK-12-R. A-C.** Independent experiments with *E. coli*. **D-F.** Independent experiments with *P. aeruginosa*.. Growth over 18 hours at 37 °C with a gradient of peptide concentrations given in μM.

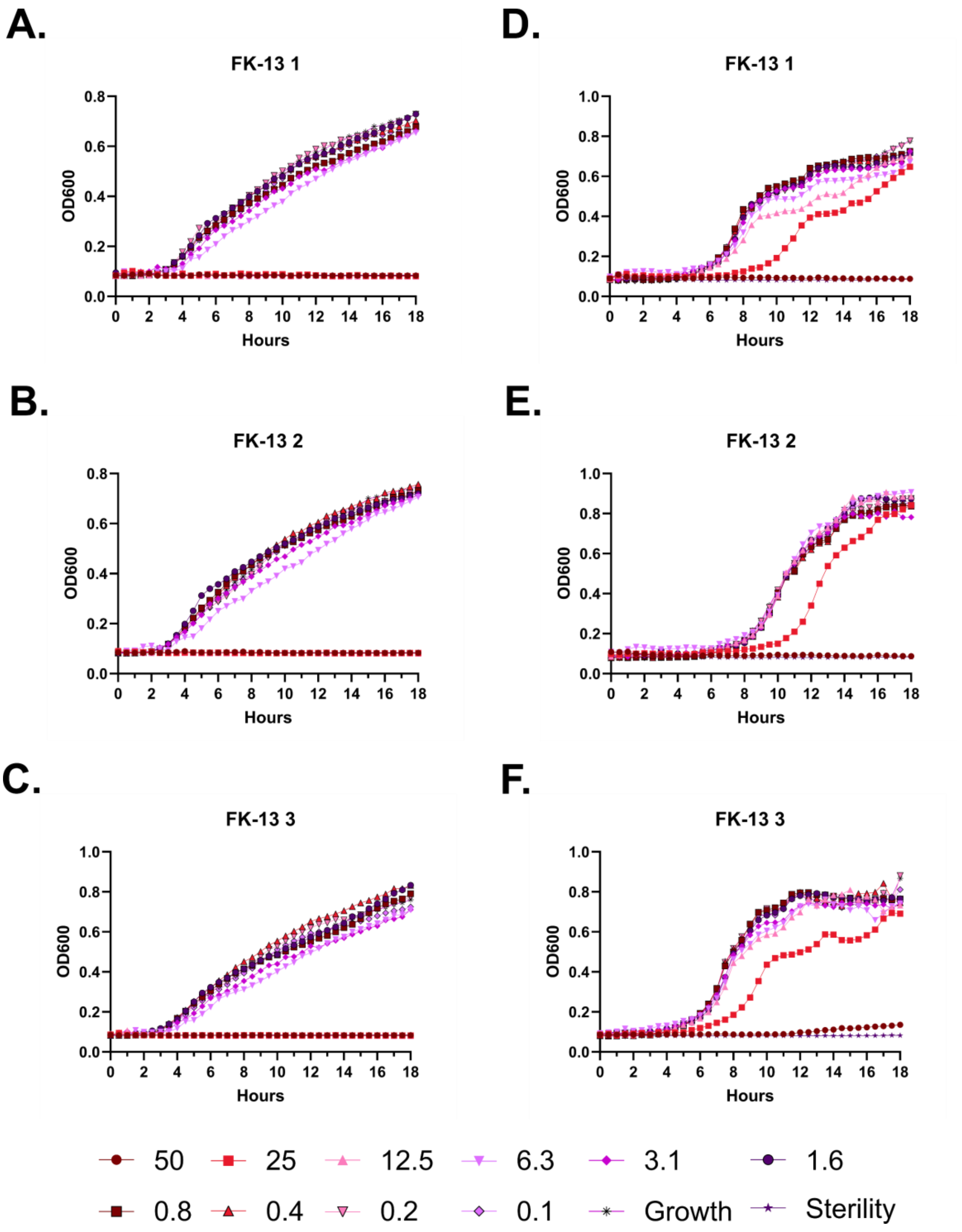
A.
FK-13 1
D.
FK-13 1
B.
FK-13 2
E.
FK-13 2
C.
FK-13 3
F.
FK-13 3
OD600
Hours
50
25
12.5
6.3
3.1
1.6
0.8
0.4
0.2
0.1
Growth
Sterility

**Figure S33 – Antimicrobial susceptibility curves for (#7) FK-13. A-C.** Independent experiments with *E. coli*. **D-F.** Independent experiments with *P. aeruginosa*.. Growth over 18 hours at 37 °C with a gradient of peptide concentrations given in μM.

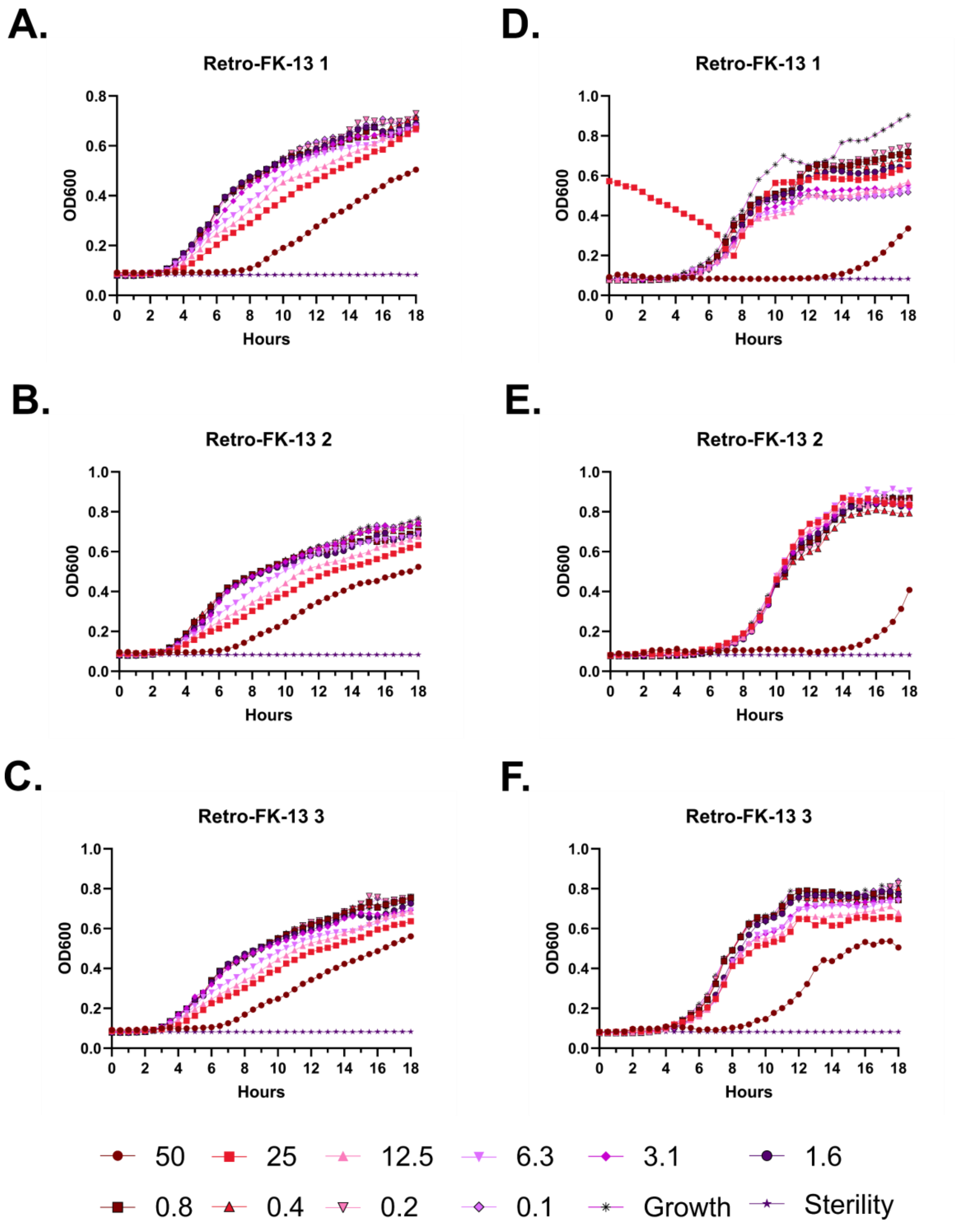
A.
Retro-FK-13 1
B.
Retro-FK-13 2
C.
Retro-FK-13 3
D.
Retro-FK-13 1
E.
Retro-FK-13 2
F.
Retro-FK-13 3
OD600
Hours
50
25
12.5
6.3
3.1
1.6
0.8
0.4
0.2
0.1
Growth
Sterility

**Figure S34 – Antimicrobial susceptibility curves for (#15) Retro-FK-13. A-C.** Independent experiments with *E. coli*. **D-F.** Independent experiments with *P. aeruginosa.*. Growth over 18 hours at 37 °C with a gradient of peptide concentrations given in μM.

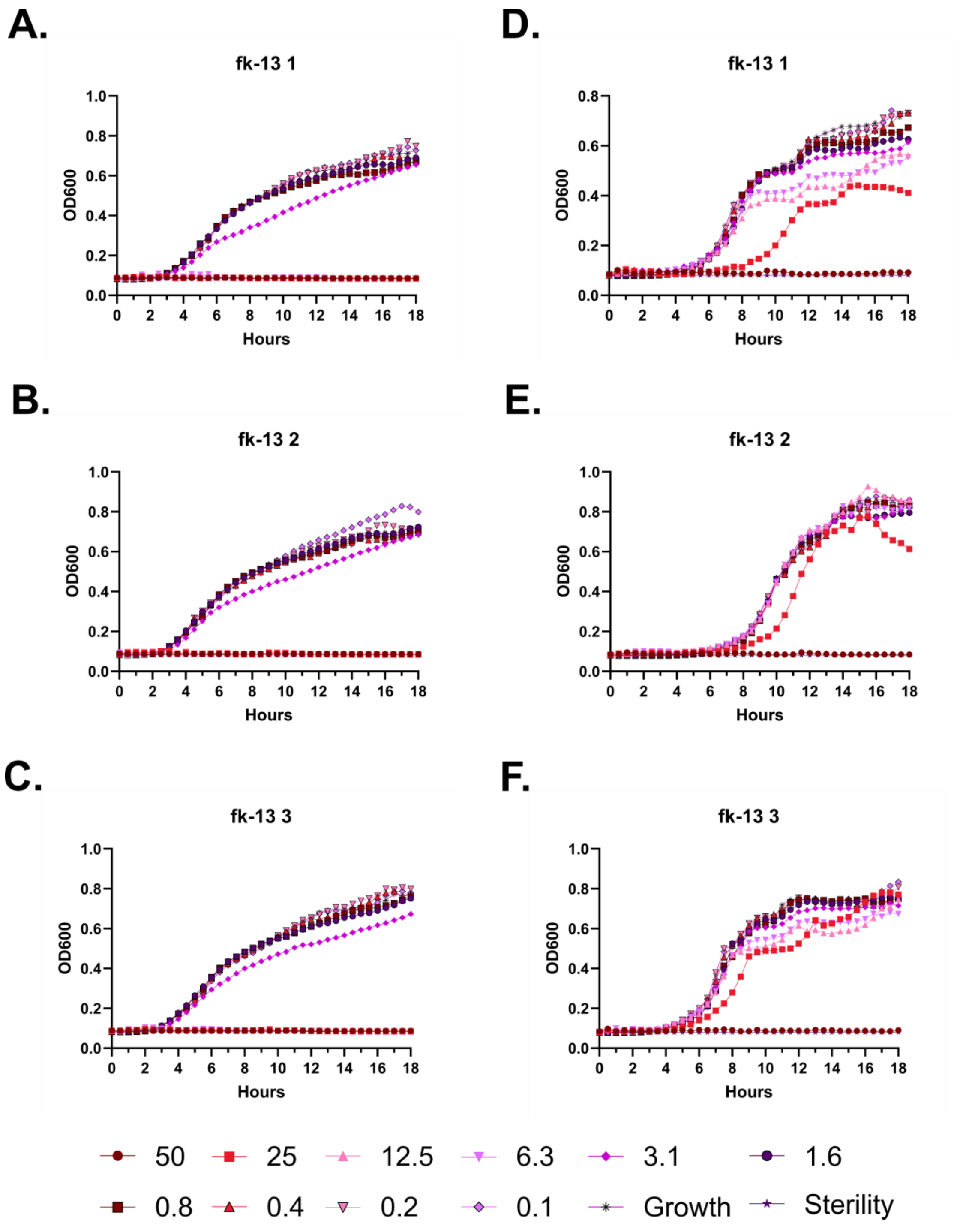

A.
fk-13 1
OD600
Hours
B.
fk-13 2
C.
fk-13 3
D.
fk-13 1
E.
fk-13 2
F.
fk-13 3
50
25
12.5
6.3
3.1
1.6
0.8
0.4
0.2
0.1
Growth
Sterility

**Figure S35 – Antimicrobial susceptibility curves for (#16) fk-13. A-C.** Independent experiments with *E. coli*. **D-F.** Independent experiments with *P. aeruginosa*.. Growth over 18 hours at 37 °C with a gradient of peptide concentrations given in μM.

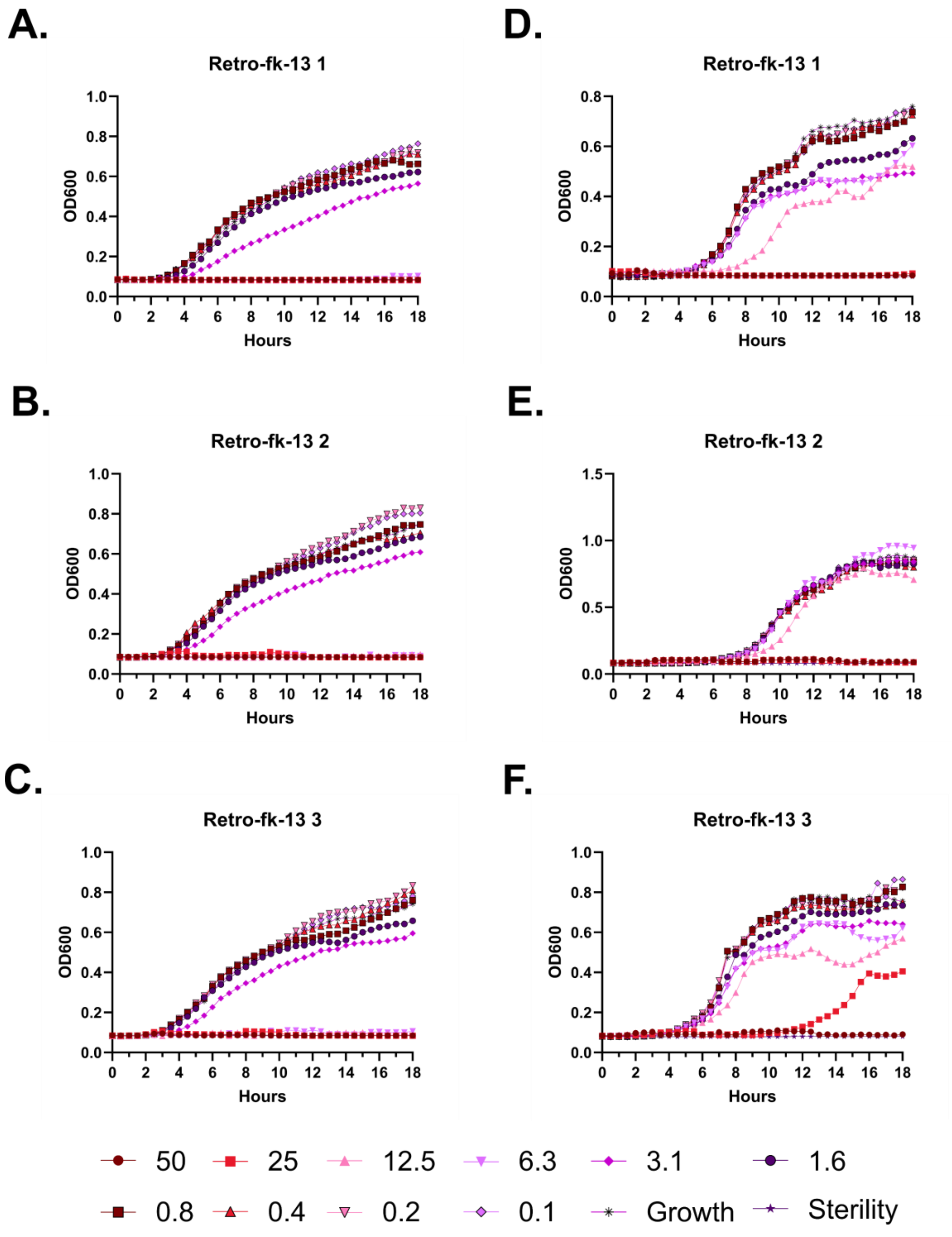
A.
Retro-fk-13 1
OD600
Hours
D.
Retro-fk-13 1
OD600
Hours
B.
Retro-fk-13 2
OD600
Hours
E.
Retro-fk-13 2
OD600
Hours
C.
Retro-fk-13 3
OD600
Hours
F.
Retro-fk-13 3
OD600
Hours
50
25
12.5
6.3
3.1
1.6
0.8
0.4
0.2
0.1
Growth
Sterility

**Figure S36 – Antimicrobial susceptibility curves for (#17) Retro-fk-13. A-C.** Independent experiments with *E. coli*. **D-F.** Independent experiments with *P. aeruginosa*.. Growth over 18 hours at 37 °C with a gradient of peptide concentrations given in μM.

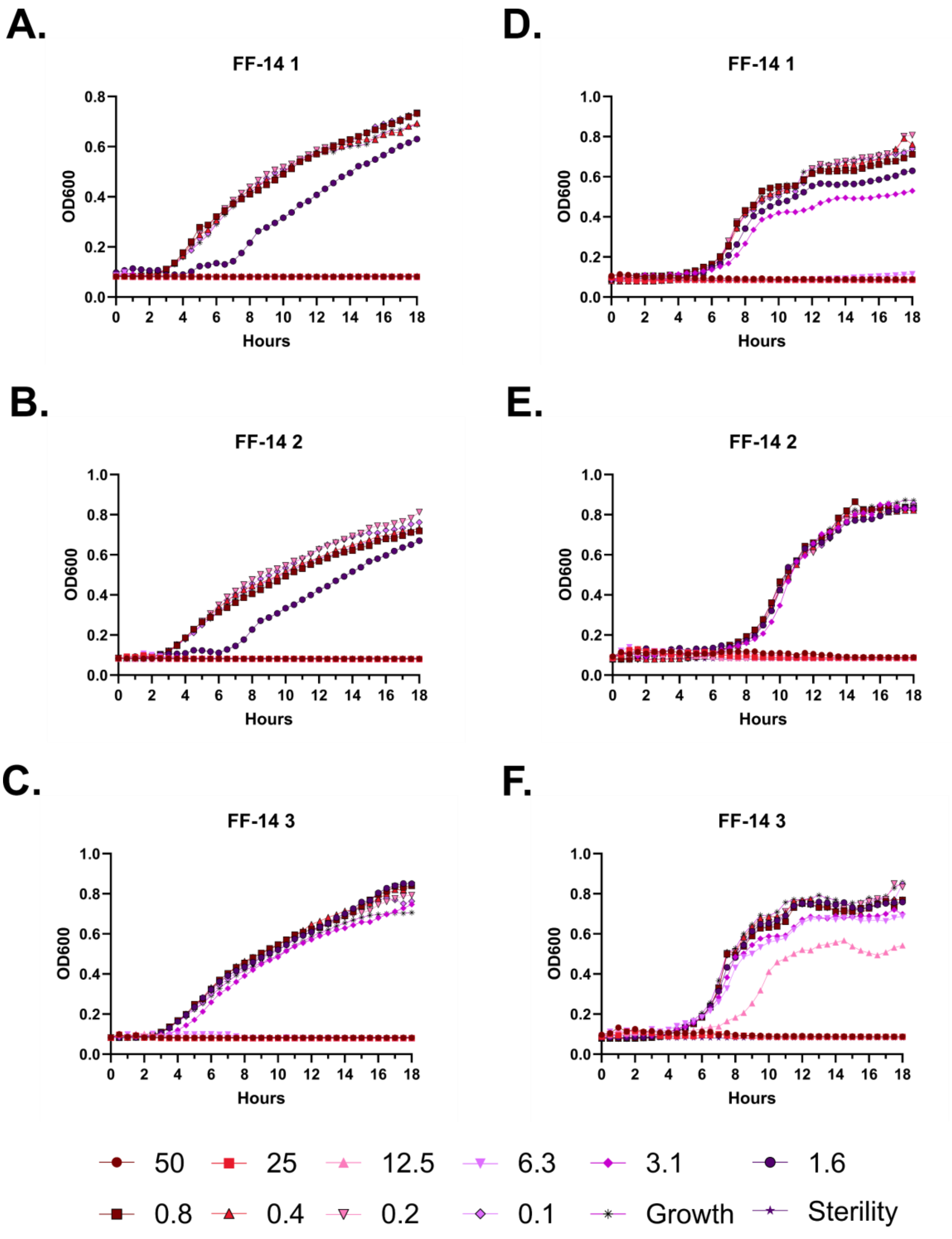
A.
FF-14 1
OD600
Hours
D.
FF-14 1
OD600
Hours
B.
FF-14 2
OD600
Hours
E.
FF-14 2
OD600
Hours
C.
FF-14 3
OD600
Hours
F.
FF-14 3
OD600
Hours
50
25
12.5
6.3
3.1
1.6
0.8
0.4
0.2
0.1
Growth
Sterility

**Figure S37 – Antimicrobial susceptibility curves for (#18) FF-14. A-C.** Independent experiments with *E. coli*. **D-F.** Independent experiments with *P. aeruginosa.* Growth over 18 hours at 37 °C with a gradient of peptide concentrations given in μM.

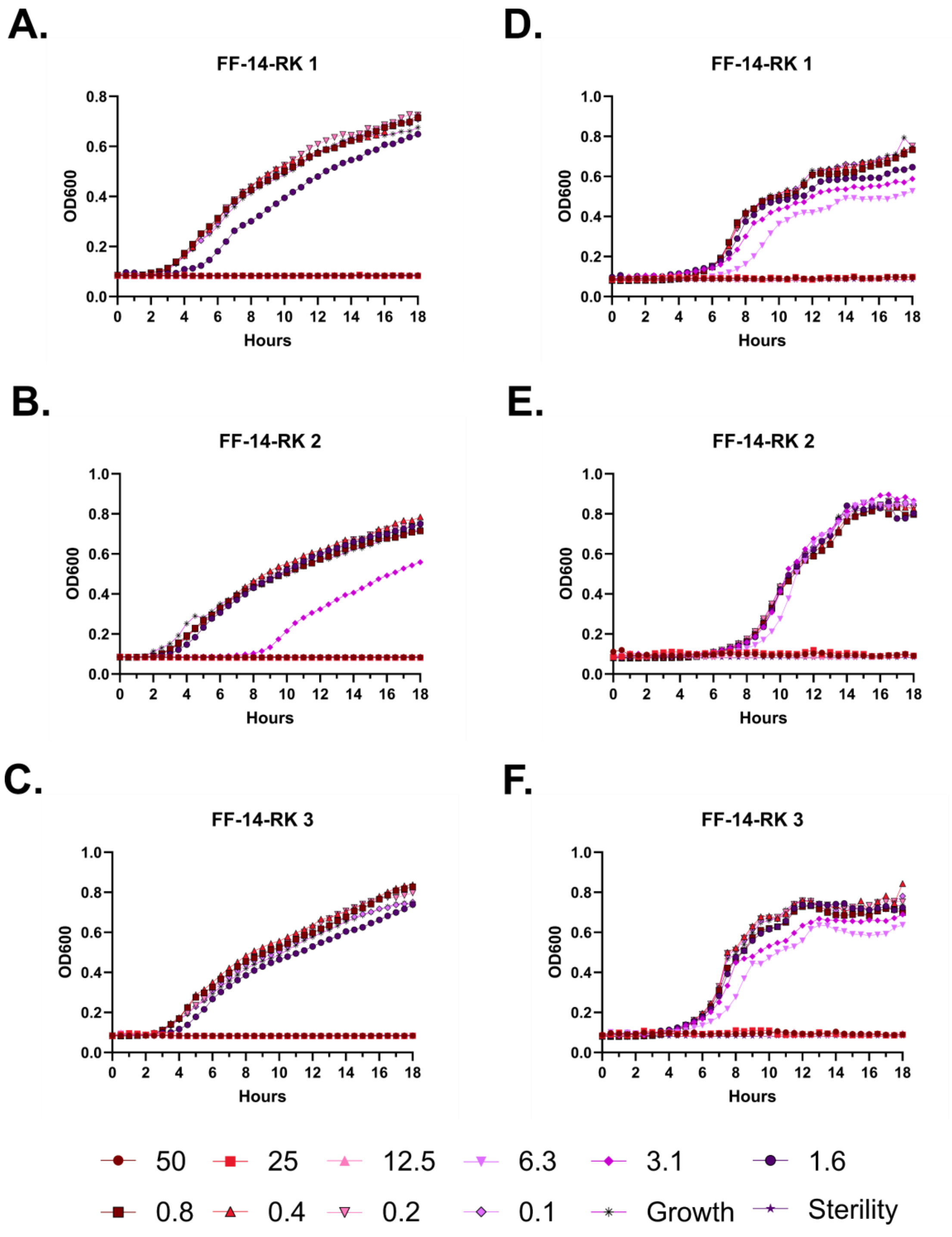
A.
FF-14-RK 1
D.
FF-14-RK 1
B.
FF-14-RK 2
E.
FF-14-RK 2
C.
FF-14-RK 3
F.
FF-14-RK 3
OD600
Hours
50
25
12.5
6.3
3.1
1.6
0.8
0.4
0.2
0.1
Growth
Sterility

**Figure S38 – Antimicrobial susceptibility curves for (#19) FF-14-RK. A-C.** Independent experiments with *E. coli*. **D-F.** Independent experiments with *P. aeruginosa.* Growth over 18 hours at 37 °C with a gradient of peptide concentrations given in μM.

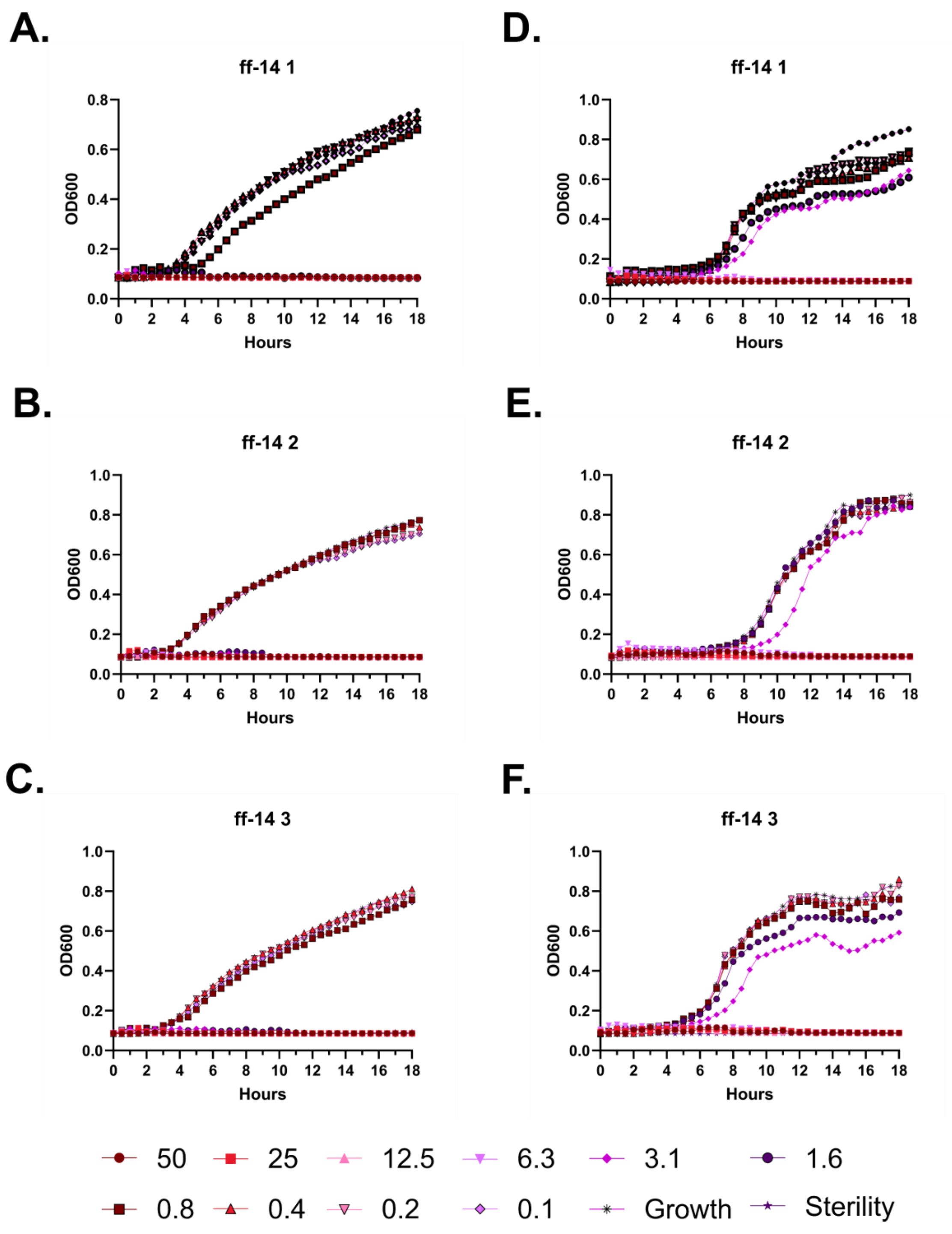
A.
ff-14 1
OD600
Hours
D.
ff-14 1
OD600
Hours
B.
ff-14 2
OD600
Hours
E.
ff-14 2
OD600
Hours
C.
ff-14 3
OD600
Hours
F.
ff-14 3
OD600
Hours
50
25
12.5
6.3
3.1
1.6
0.8
0.4
0.2
0.1
Growth
Sterility

**Figure S39 – Antimicrobial susceptibility curves for (#20) ff-14. A-C.** Independent experiments with *E. coli*. **D-F.** Independent experiments with *P. aeruginosa.* Growth over 18 hours at 37 °C with a gradient of peptide concentrations given in μM.

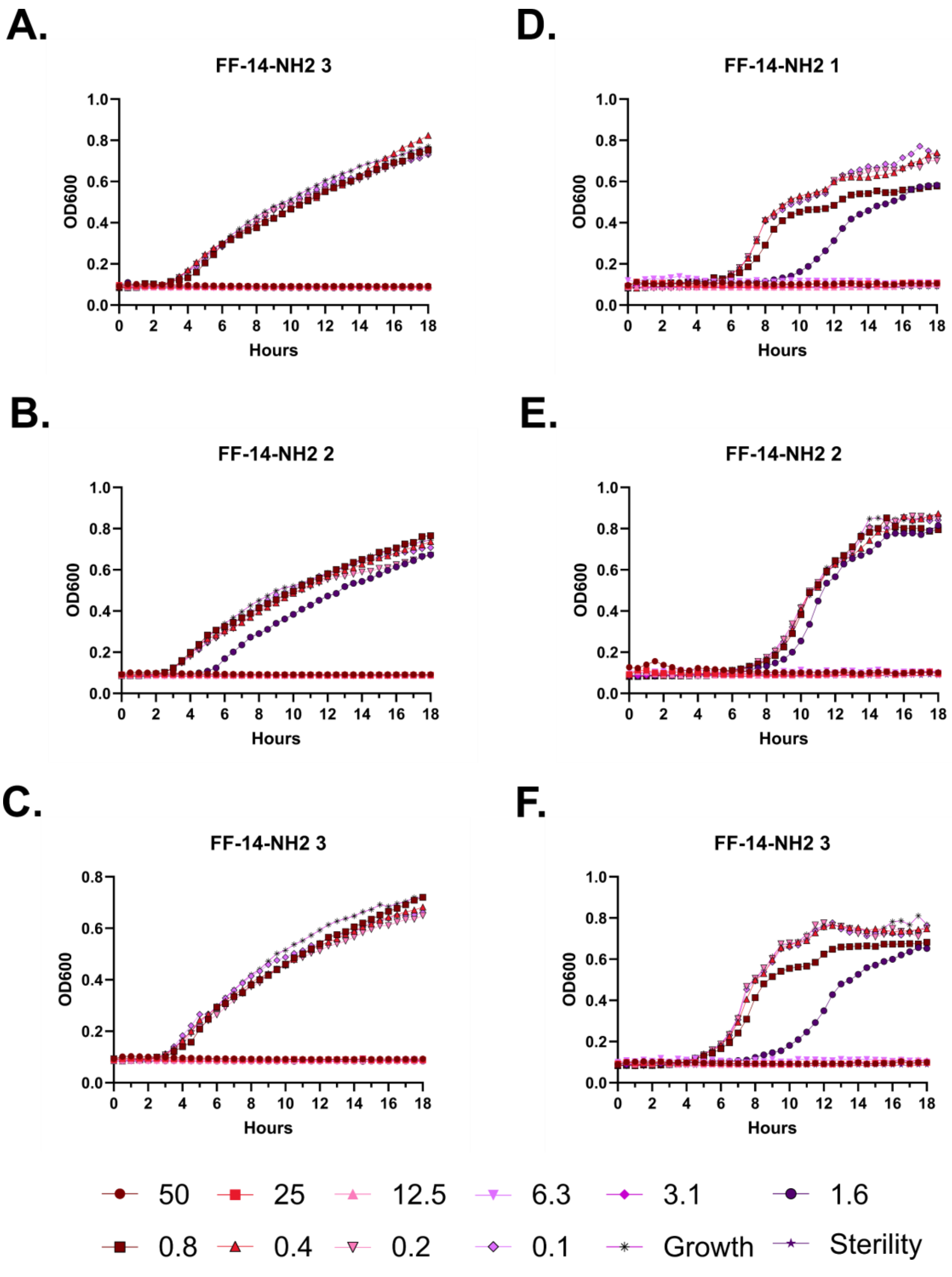
A.
FF-14-NH2 3
D.
FF-14-NH2 1
B.
FF-14-NH2 2
E.
FF-14-NH2 2
C.
FF-14-NH2 3
F.
FF-14-NH2 3
OD600
Hours
50
25
12.5
6.3
3.1
1.6
0.8
0.4
0.2
0.1
Growth
Sterility

**Figure S40 – Antimicrobial susceptibility curves for (#21) FF-14-$NH_2$. A-C.** Independent experiments with *E. coli*. **D-F.** Independent experiments with *P. aeruginosa.* Growth over 18 hours at 37 °C with a gradient of peptide concentrations given in μM.

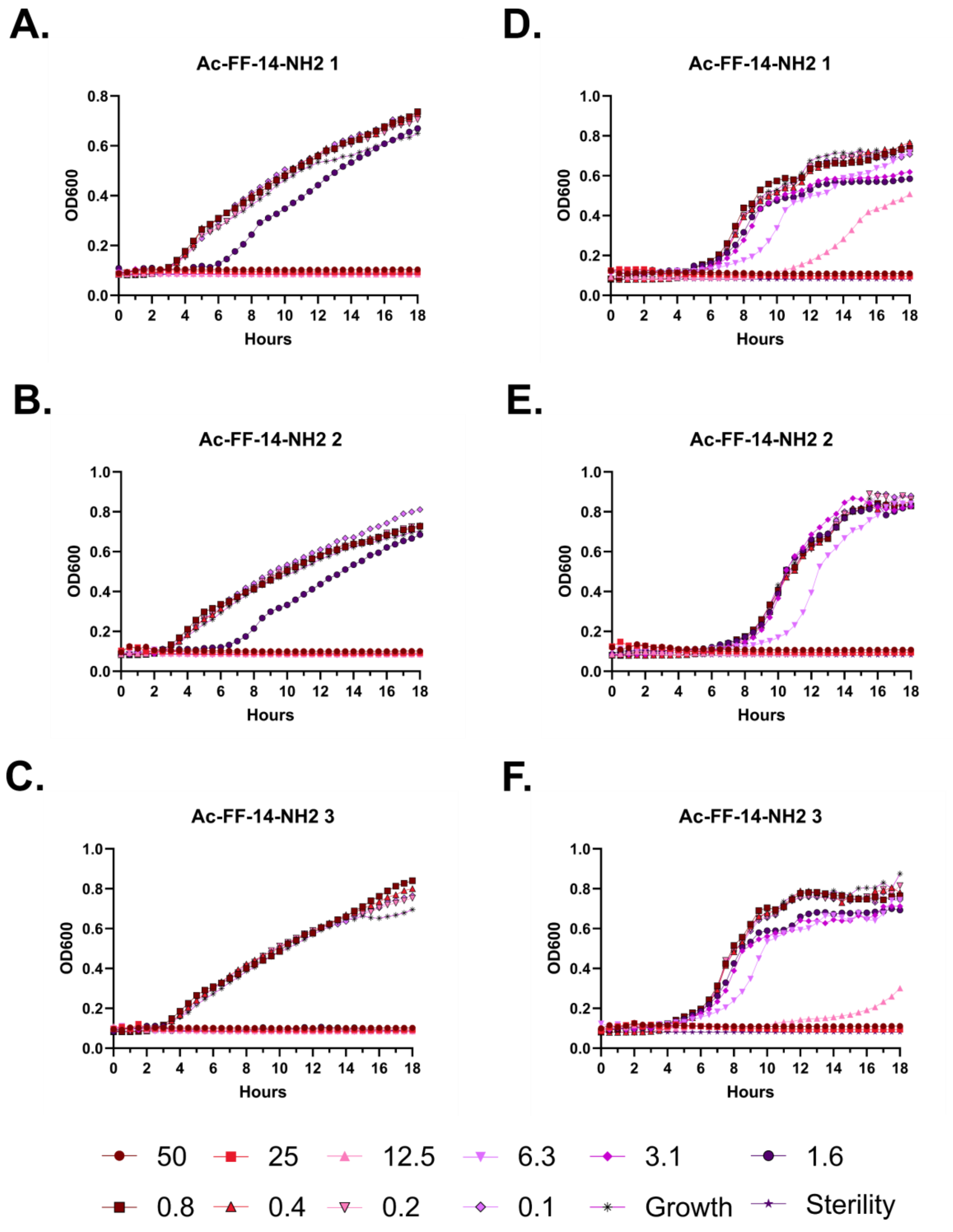
A.
Ac-FF-14-NH2 1
D.
Ac-FF-14-NH2 1
B.
Ac-FF-14-NH2 2
E.
Ac-FF-14-NH2 2
C.
Ac-FF-14-NH2 3
F.
Ac-FF-14-NH2 3
OD600
Hours
50
25
12.5
6.3
3.1
1.6
0.8
0.4
0.2
0.1
Growth
Sterility

**Figure S41 – Antimicrobial susceptibility curves for (#22) Ac-FF-14-$NH_2$. A-C.** Independent experiments with *E. coli*. **D-F.** Independent experiments with *P. aeruginosa.* Growth over 18 hours at 37 °C with a gradient of peptide concentrations given in μM.

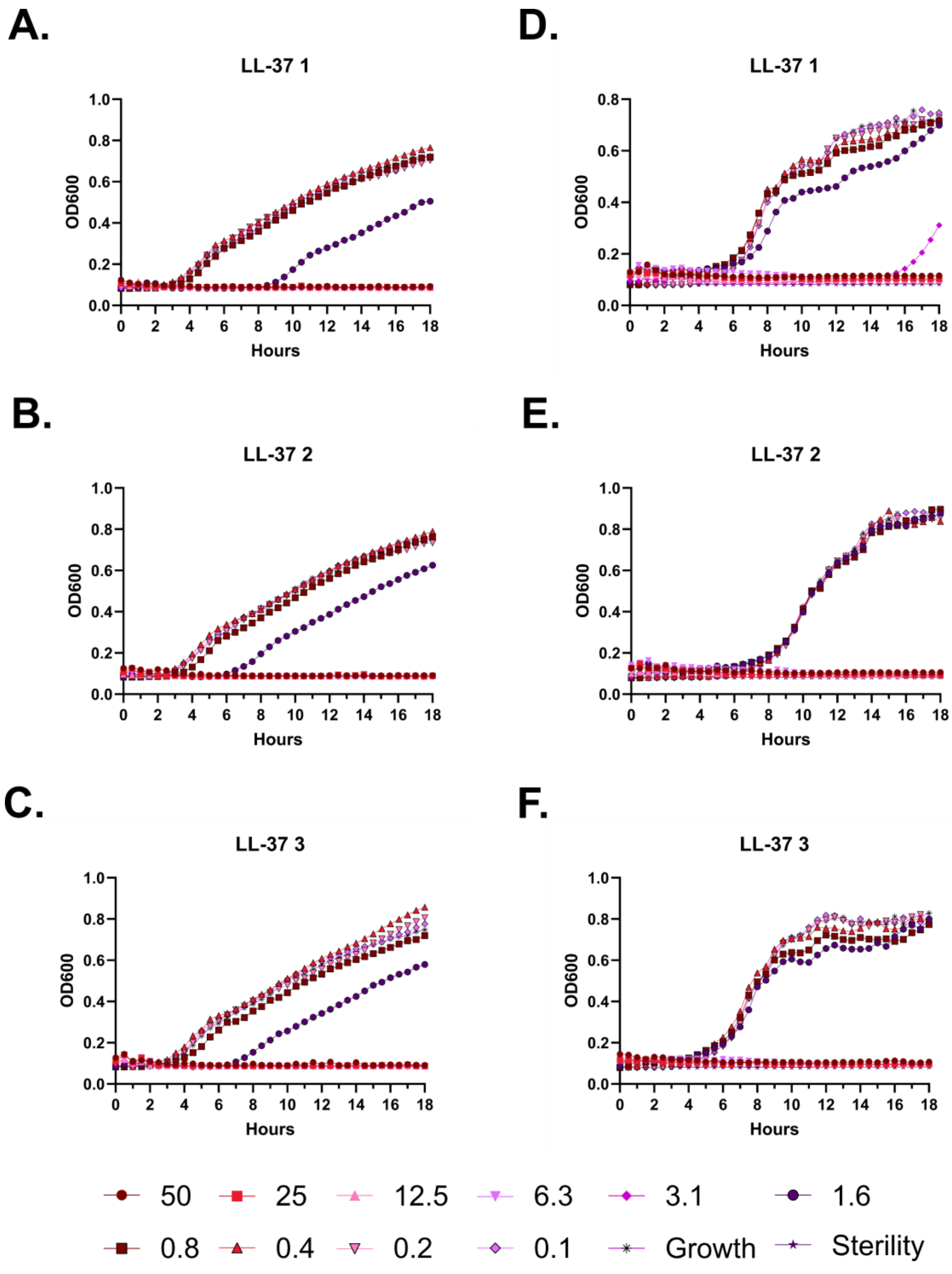
A.
LL-37 1
D.
LL-37 1
B.
LL-37 2
E.
LL-37 2
C.
LL-37 3
F.
LL-37 3
OD600
Hours
50
25
12.5
6.3
3.1
1.6
0.8
0.4
0.2
0.1
Growth
Sterility

**Figure S42 – Antimicrobial susceptibility curves for (#1) LL-37. A-C.** Independent experiments with *E. coli*. **D-F.** Independent experiments with *P. aeruginosa.* Growth over 18 hours at 37 °C with a gradient of peptide concentrations given in μM.

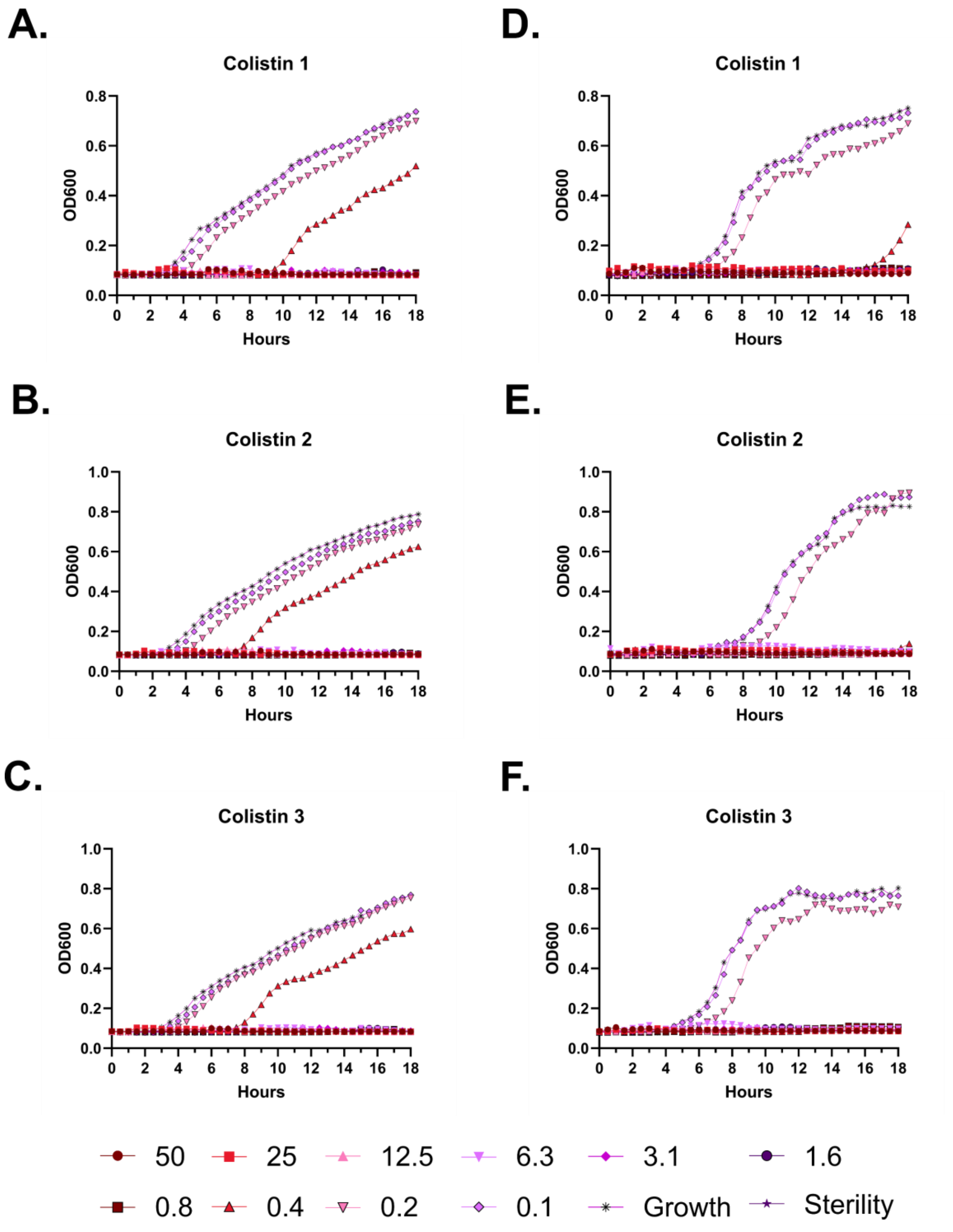
A.
Colistin 1
OD600
Hours
B.
Colistin 2
OD600
Hours
C.
Colistin 3
OD600
Hours
D.
Colistin 1
OD600
Hours
E.
Colistin 2
OD600
Hours
F.
Colistin 3
OD600
Hours
50
25
12.5
6.3
3.1
1.6
0.8
0.4
0.2
0.1
Growth
Sterility

**Figure S43 – Antimicrobial susceptibility curves for colistin. A-C.** Independent experiments with *E. coli*. **D-F.** Independent experiments with *P. aeruginosa.* Growth over 18 hours at 37 °C with a gradient of peptide concentrations given in μM.

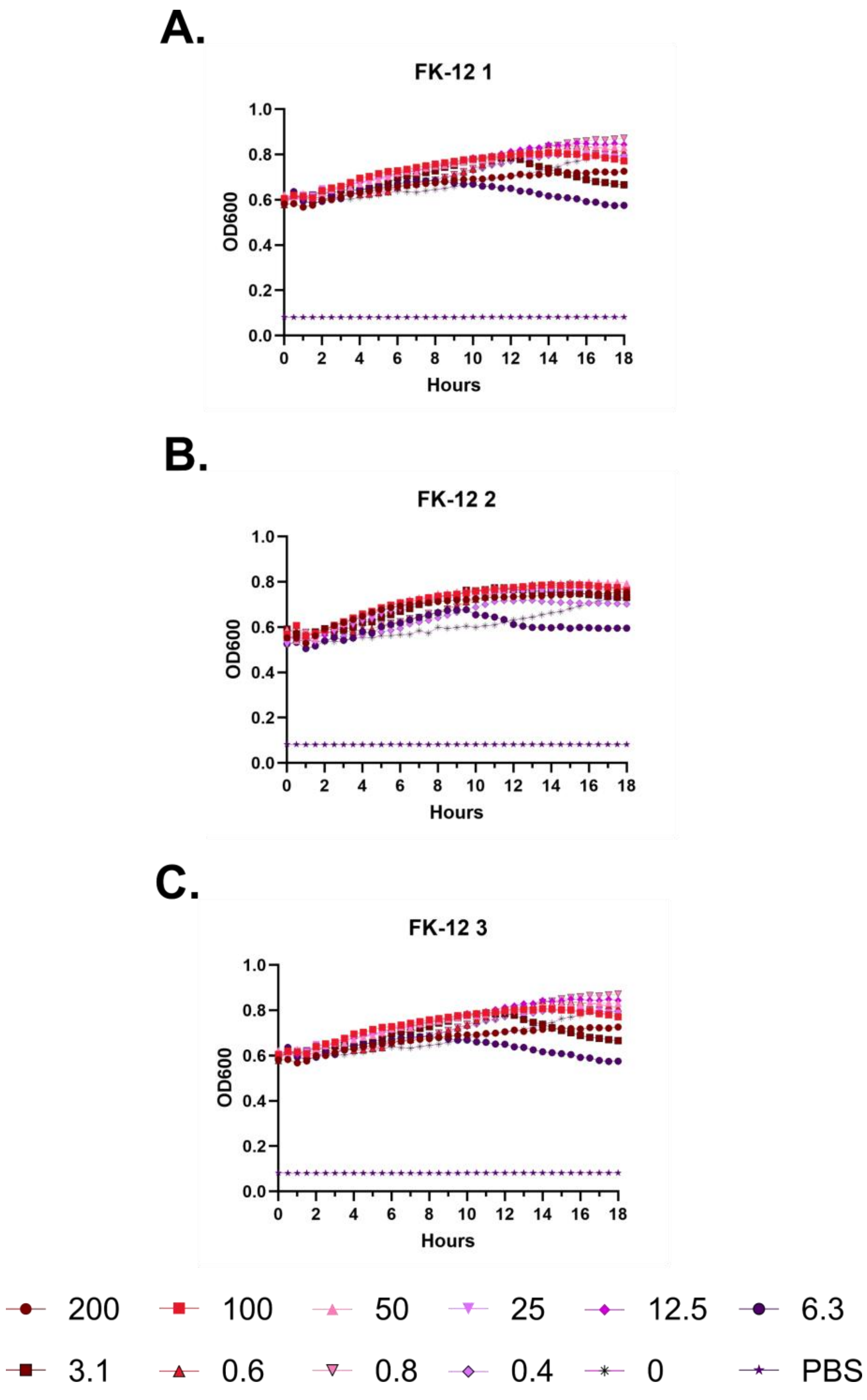
A.
FK-12 1
OD600
Hours
B.
FK-12 2
OD600
Hours
C.
FK-12 3
OD600
Hours
200
100
50
25
12.5
6.3
3.1
0.6
0.8
0.4
0
PBS

**Figure S44 – (#13) FK-12 kinetic hemolysis assay. A.** Defibrinated sheep blood was incubated in the presence of the indicated peptide concentrations at 37 °C with monitoring of OD600 over 18 hours, where decreasing OD600 indicates hemolysis and each panel indicates an independent experiment. Peptide concentrations are in μM.

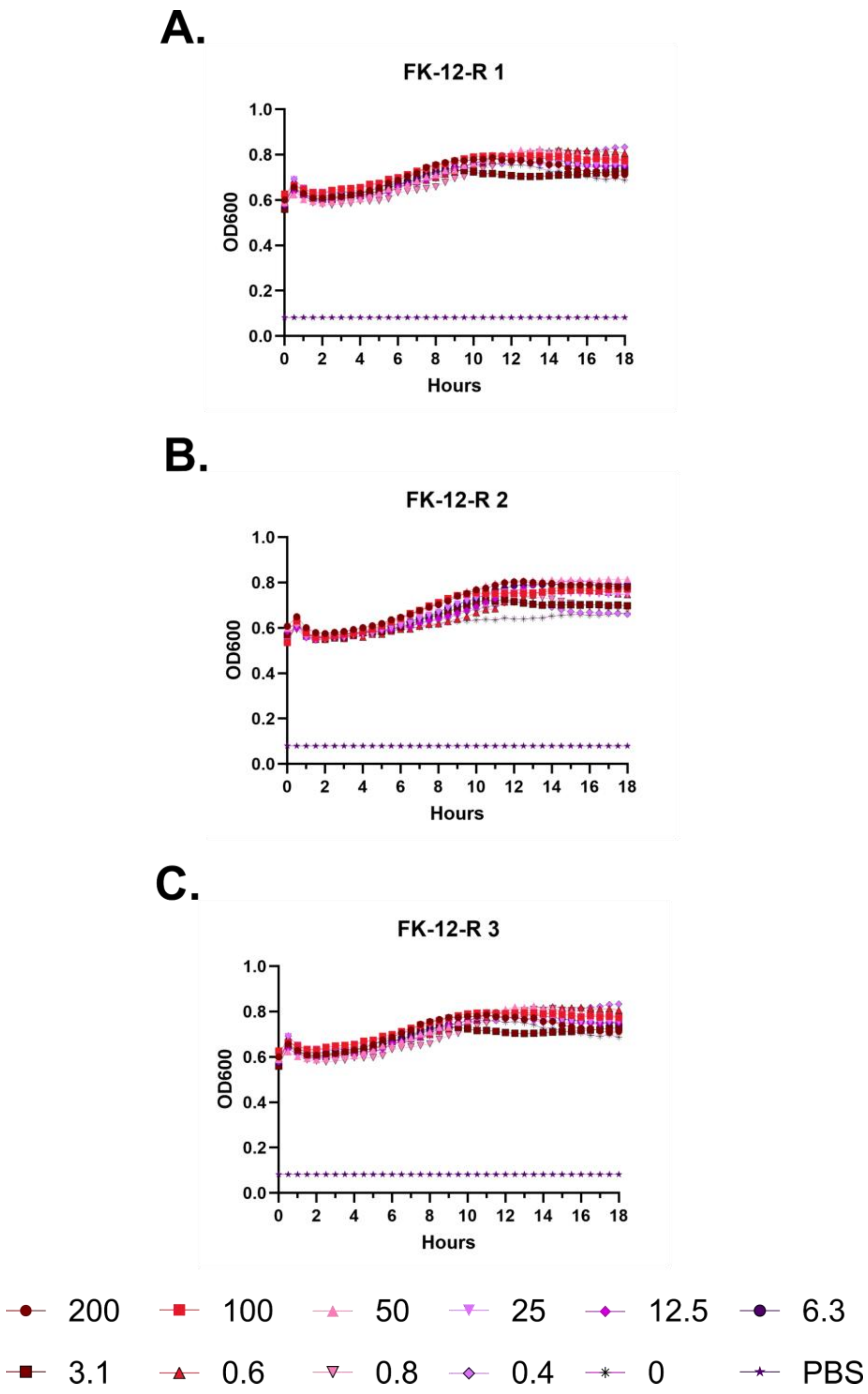
A.
FK-12-R 1
OD600
Hours
B.
FK-12-R 2
OD600
Hours
C.
FK-12-R 3
OD600
Hours
200
100
50
25
12.5
6.3
3.1
0.6
0.8
0.4
0
PBS

**Figure S45 – (#14) FK-12-R kinetic hemolysis assay. A.** Defibrinated sheep blood was incubated in the presence of the indicated peptide concentrations at 37 °C with monitoring of OD600 over 18 hours, where decreasing OD600 indicates hemolysis and each panel indicates an independent experiment. Peptide concentrations are in µM.

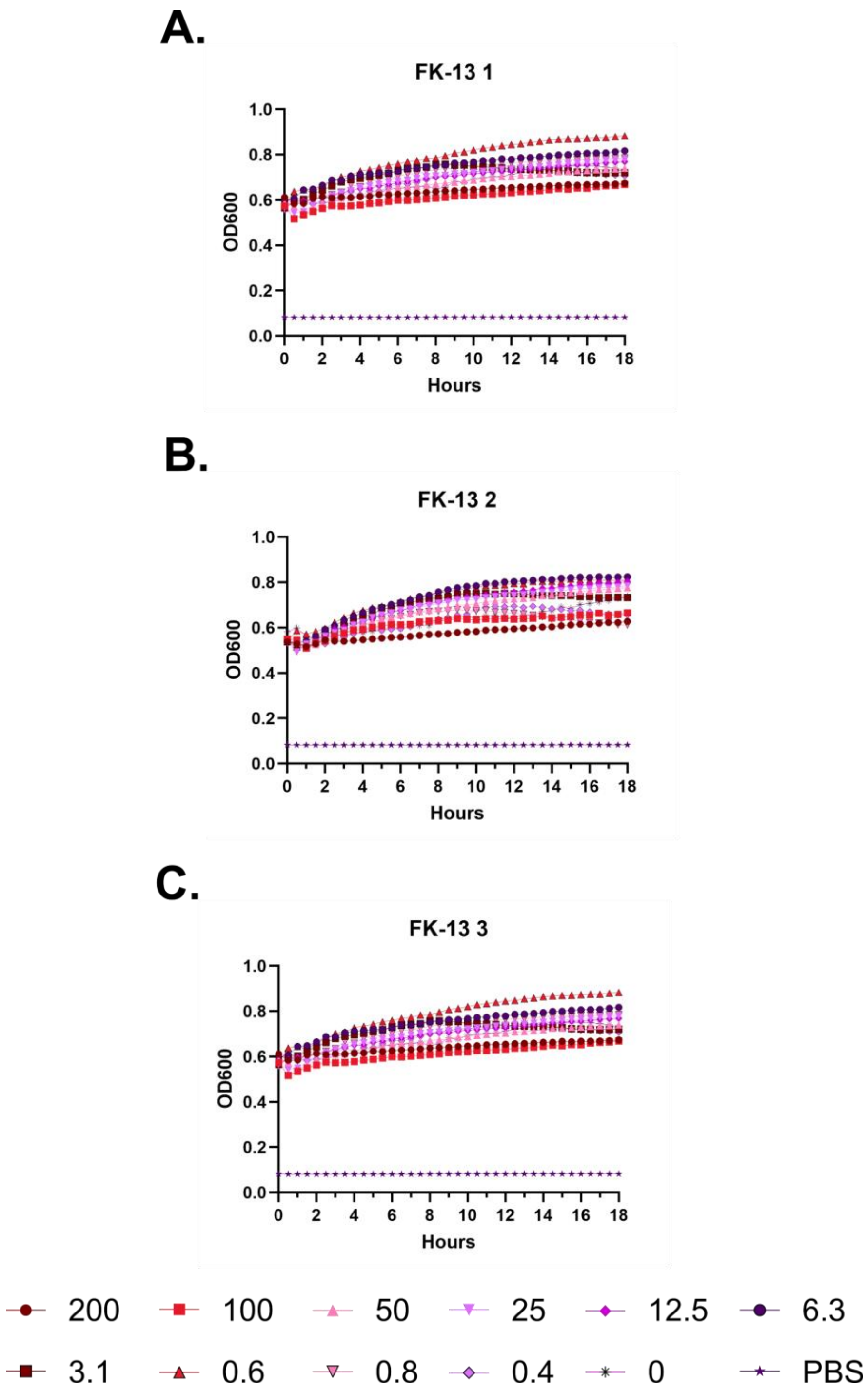
A.
FK-13 1
OD600
Hours
B.
FK-13 2
OD600
Hours
C.
FK-13 3
OD600
Hours
200
100
50
25
12.5
6.3
3.1
0.6
0.8
0.4
0
PBS

**Figure S46 – (#7) FK-13 kinetic hemolysis assay. A.** Defibrinated sheep blood was incubated in the presence of the indicated peptide concentrations at 37 °C with monitoring of OD600 over 18 hours, where decreasing OD600 indicates hemolysis and each panel indicates an independent experiment. Peptide concentrations are in μM.

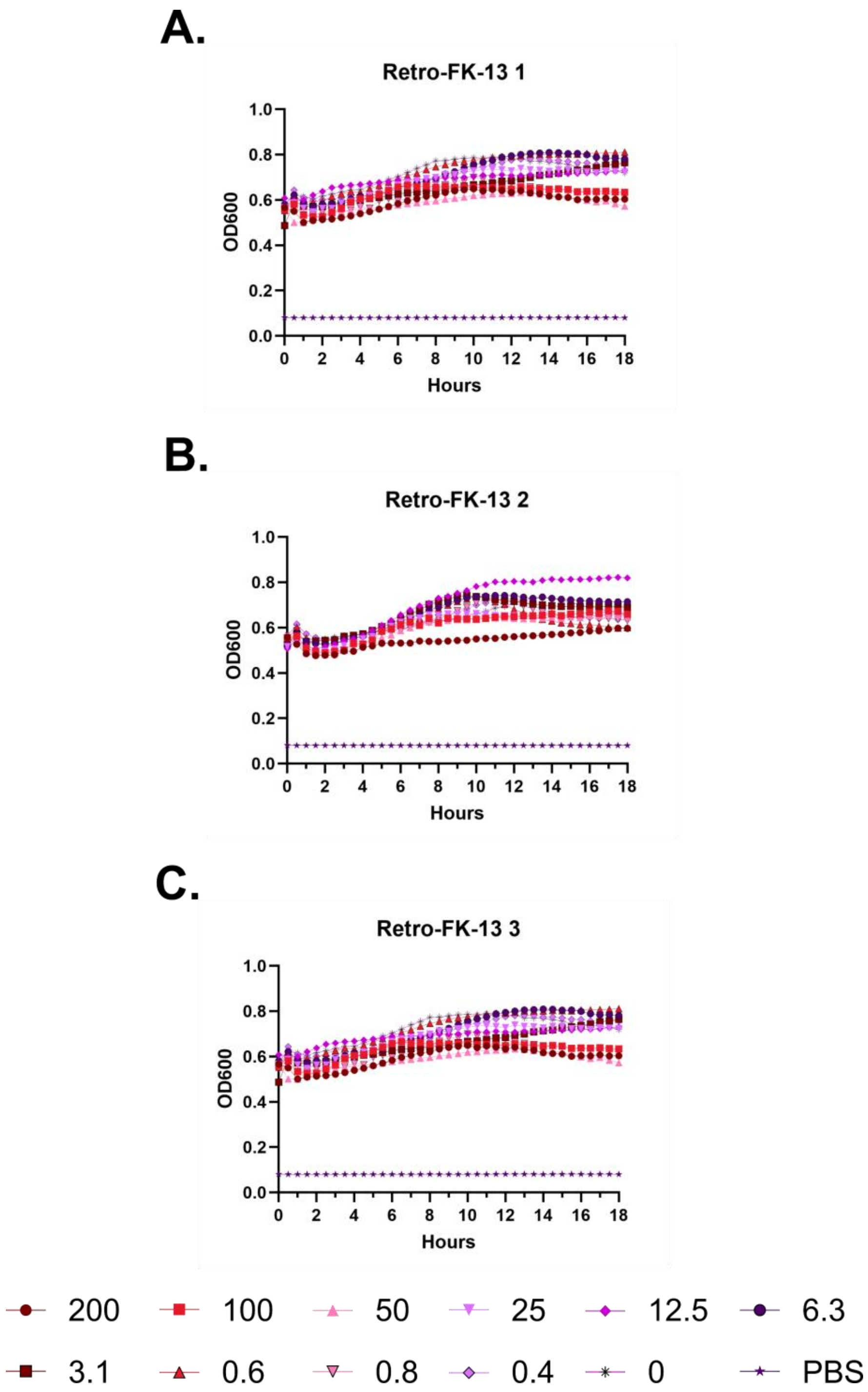
A.
Retro-FK-13 1
OD600
Hours
B.
Retro-FK-13 2
OD600
Hours
C.
Retro-FK-13 3
OD600
Hours
200
100
50
25
12.5
6.3
3.1
0.6
0.8
0.4
0
PBS

**Figure S47 – (#15) Retro-FK-13 kinetic hemolysis assay. A.** Defibrinated sheep blood was incubated in the presence of the indicated peptide concentrations at 37 °C with monitoring of OD600 over 18 hours, where decreasing OD600 indicates hemolysis and each panel indicates an independent experiment. Peptide concentrations are in µM.

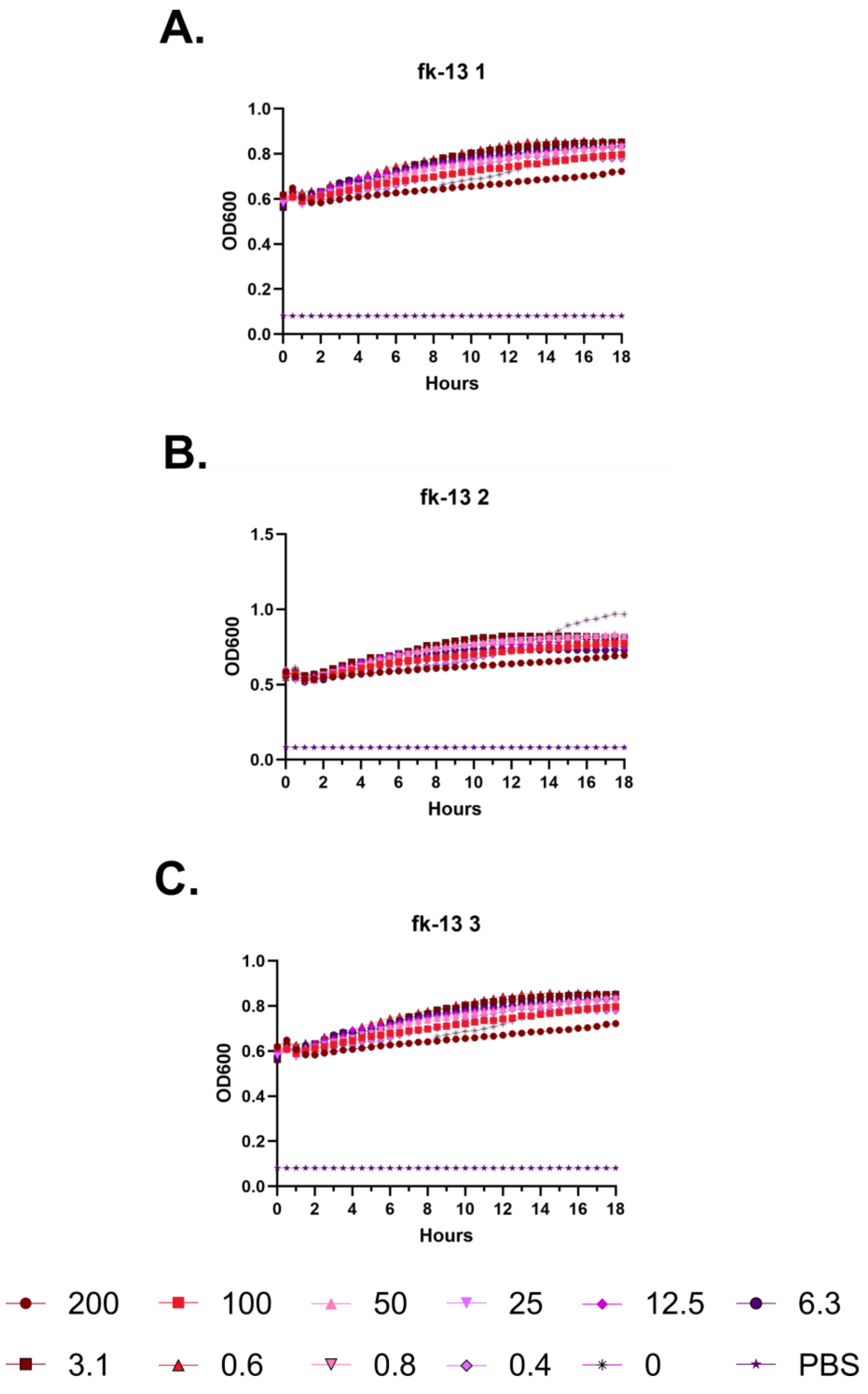
A.
fk-13 1
OD600
1.0
0.8
0.6
0.4
0.2
0.0
0 2 4 6 8 10 12 14 16 18
Hours
B.
fk-13 2
OD600
1.5
1.0
0.5
0.0
0 2 4 6 8 10 12 14 16 18
Hours
C.
fk-13 3
OD600
1.0
0.8
0.6
0.4
0.2
0.0
0 2 4 6 8 10 12 14 16 18
Hours
200
100
50
25
12.5
6.3
3.1
0.6
0.8
0.4
0
PBS

**Figure S48 – (#16) fk-13 kinetic hemolysis assay. A.** Defibrinated sheep blood was incubated in the presence of the indicated peptide concentrations at 37 °C with monitoring of OD600 over 18 hours, where decreasing OD600 indicates hemolysis and each panel indicates an independent experiment. Peptide concentrations are in μM.

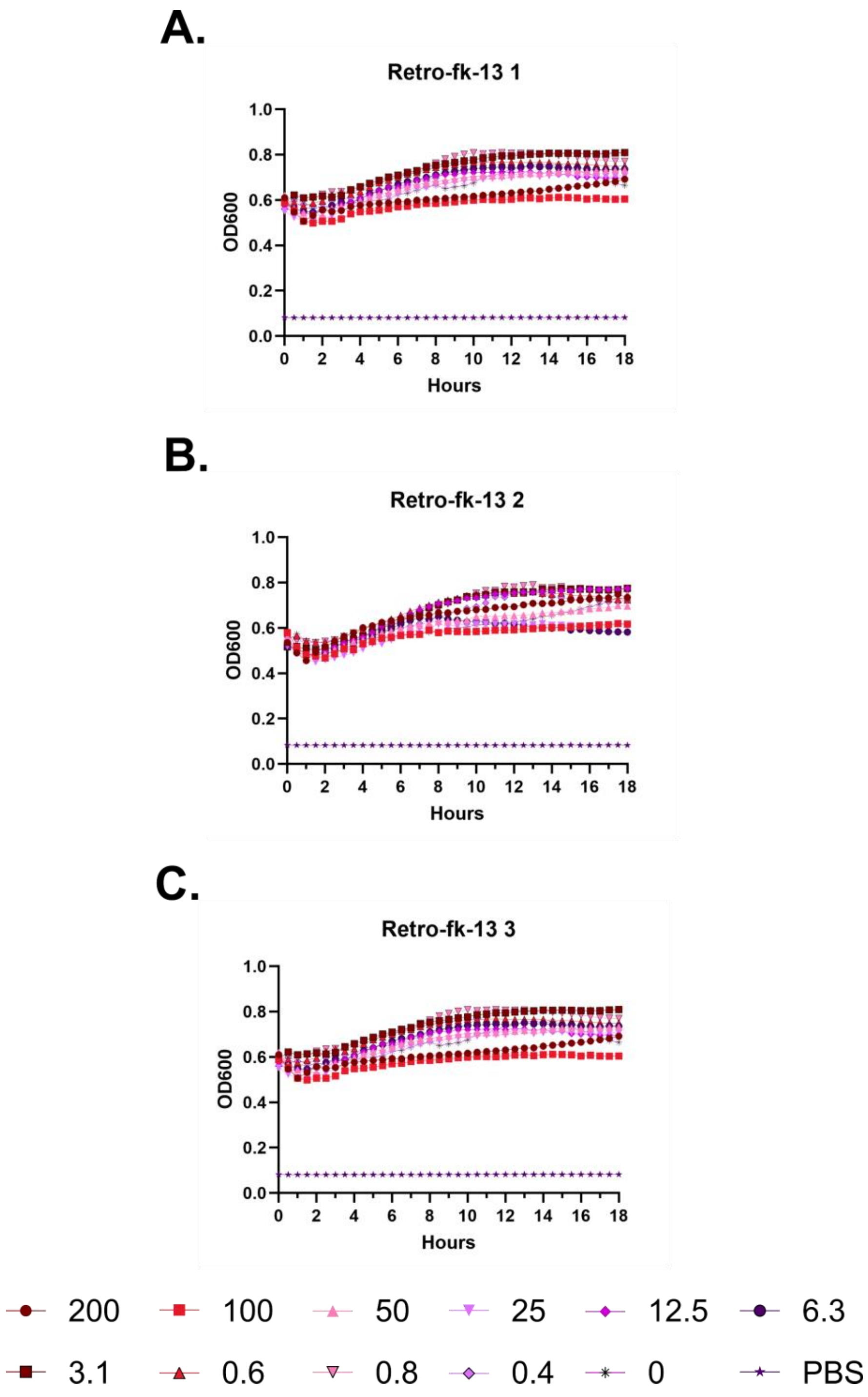
A.
Retro-fk-13 1
B.
Retro-fk-13 2
C.
Retro-fk-13 3
OD600
Hours
200
100
50
25
12.5
6.3
3.1
0.6
0.8
0.4
0
PBS

**Figure S49 – (#17) Retro-fk-13 kinetic hemolysis assay. A.** Defibrinated sheep blood was incubated in the presence of the indicated peptide concentrations at 37 °C with monitoring of OD600 over 18 hours, where decreasing OD600 indicates hemolysis and each panel indicates an independent experiment. Peptide concentrations are in μM.

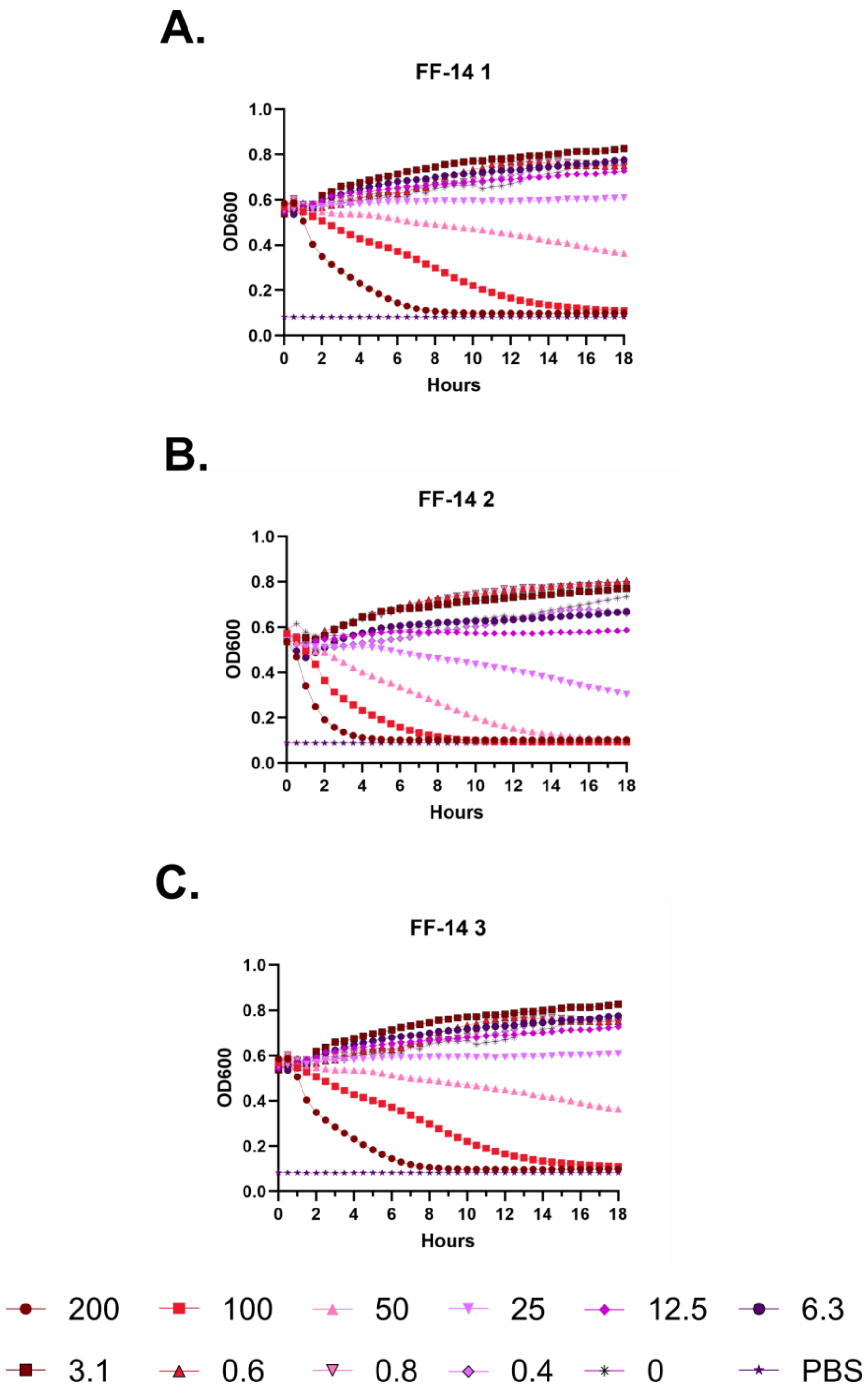

A.
FF-14 1
OD600
Hours
B.
FF-14 2
OD600
Hours
C.
FF-14 3
OD600
Hours
200
100
50
25
12.5
6.3
3.1
0.6
0.8
0.4
0
PBS

**Figure S50 – (#18) FF-14 kinetic hemolysis assay. A.** Defibrinated sheep blood was incubated in the presence of the indicated peptide concentrations at 37 °C with monitoring of OD600 over 18 hours, where decreasing OD600 indicates hemolysis and each panel indicates an independent experiment. Peptide concentrations are in µM.

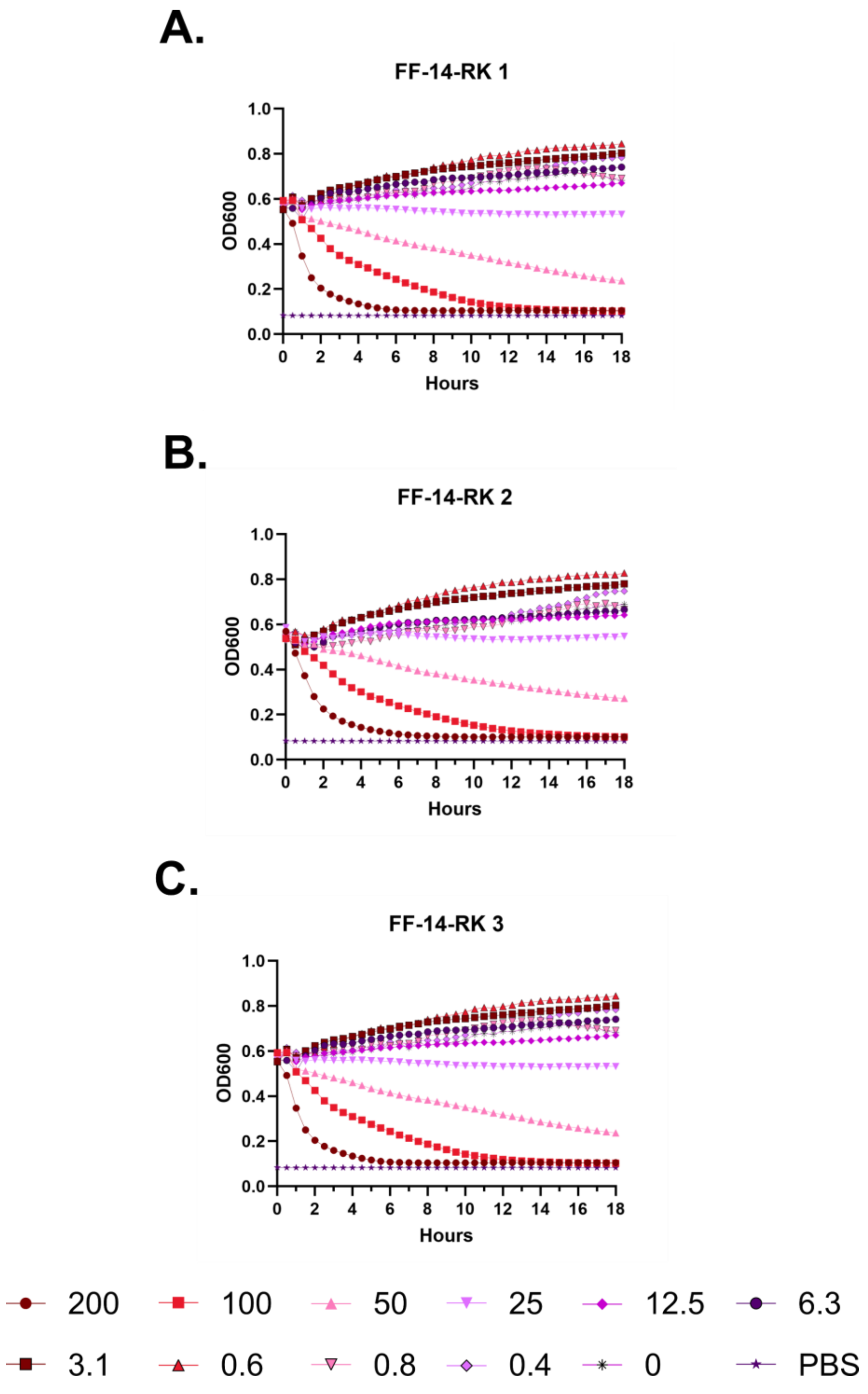
A.
FF-14-RK 1
OD600
Hours
B.
FF-14-RK 2
OD600
Hours
C.
FF-14-RK 3
OD600
Hours
200
100
50
25
12.5
6.3
3.1
0.6
0.8
0.4
0
PBS

**Figure S51 – (#19) FF-14-RK kinetic hemolysis assay. A.** Defibrinated sheep blood was incubated in the presence of the indicated peptide concentrations at 37 °C with monitoring of OD600 over 18 hours, where decreasing OD600 indicates hemolysis and each panel indicates an independent experiment. Peptide concentrations are in μM.

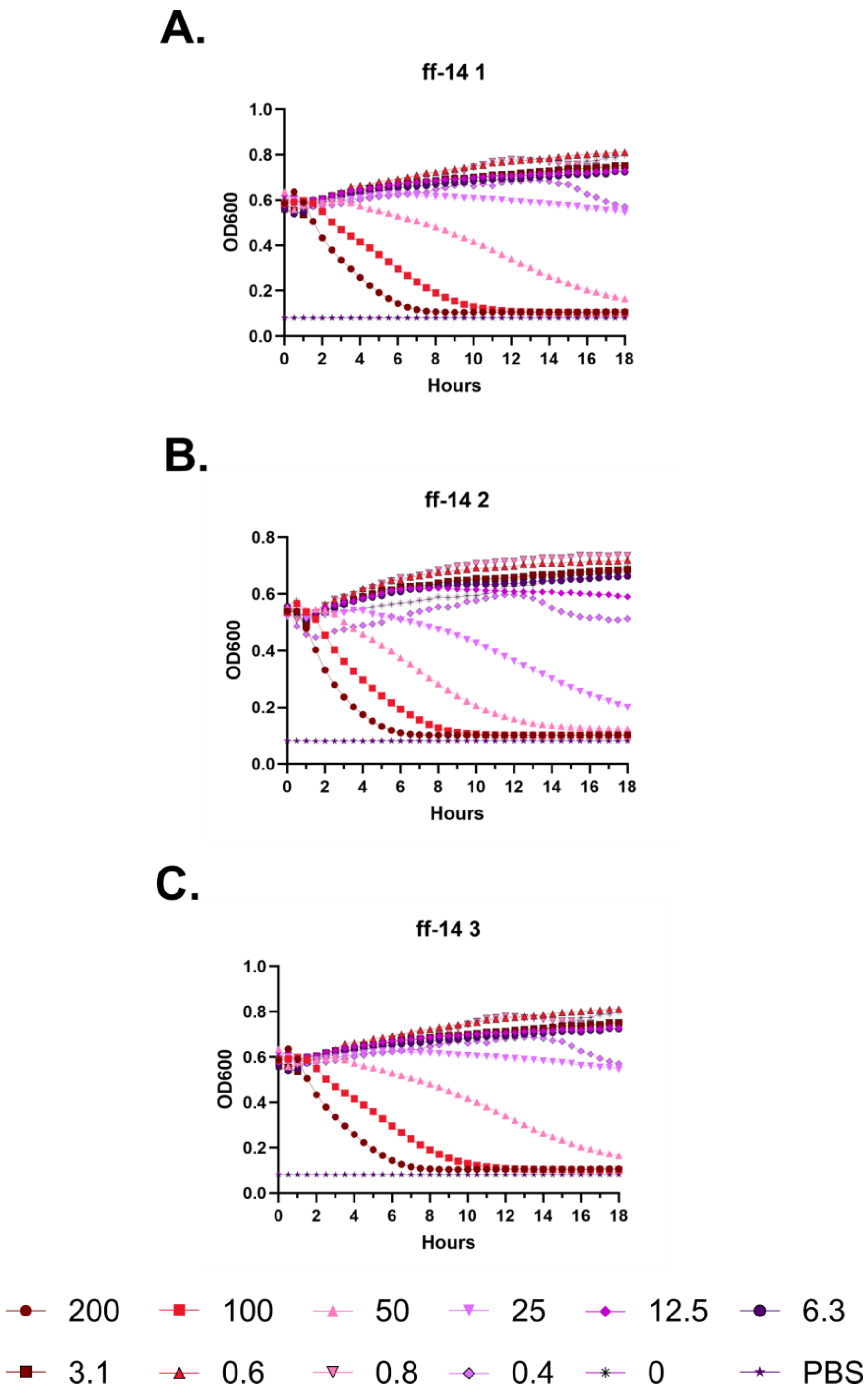
A.
ff-14 1
OD600
Hours
B.
ff-14 2
OD600
Hours
C.
ff-14 3
OD600
Hours
200
100
50
25
12.5
6.3
3.1
0.6
0.8
0.4
0
PBS

**Figure S52 – (#20) ff-14 kinetic hemolysis assay. A.** Defibrinated sheep blood was incubated in the presence of the indicated peptide concentrations at 37 °C with monitoring of OD600 over 18 hours, where decreasing OD600 indicates hemolysis and each panel indicates an independent experiment. Peptide concentrations are in μM.

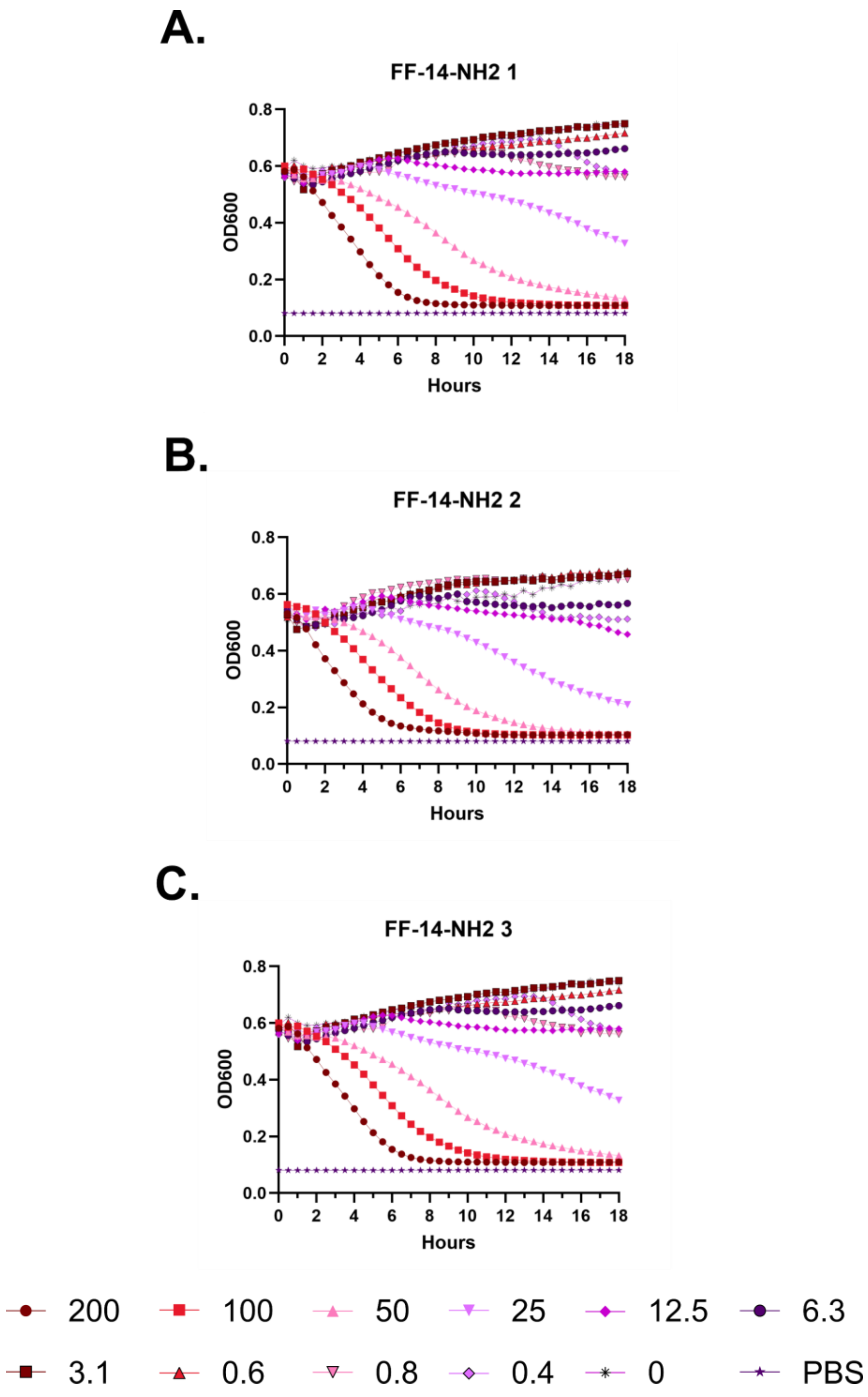
A.
FF-14-NH2 1
B.
FF-14-NH2 2
C.
FF-14-NH2 3
OD600
Hours
0.0
0.2
0.4
0.6
0.8
0
2
4
6
8
10
12
14
16
18
200
100
50
25
12.5
6.3
3.1
0.6
0.8
0.4
0
PBS

**Figure S51 – (#21) FF-14-$NH_2$ kinetic hemolysis assay. A.** Defibrinated sheep blood was incubated in the presence of the indicated peptide concentrations at 37 °C with monitoring of OD600 over 18 hours, where decreasing OD600 indicates hemolysis and each panel indicates an independent experiment. Peptide concentrations are in μM.

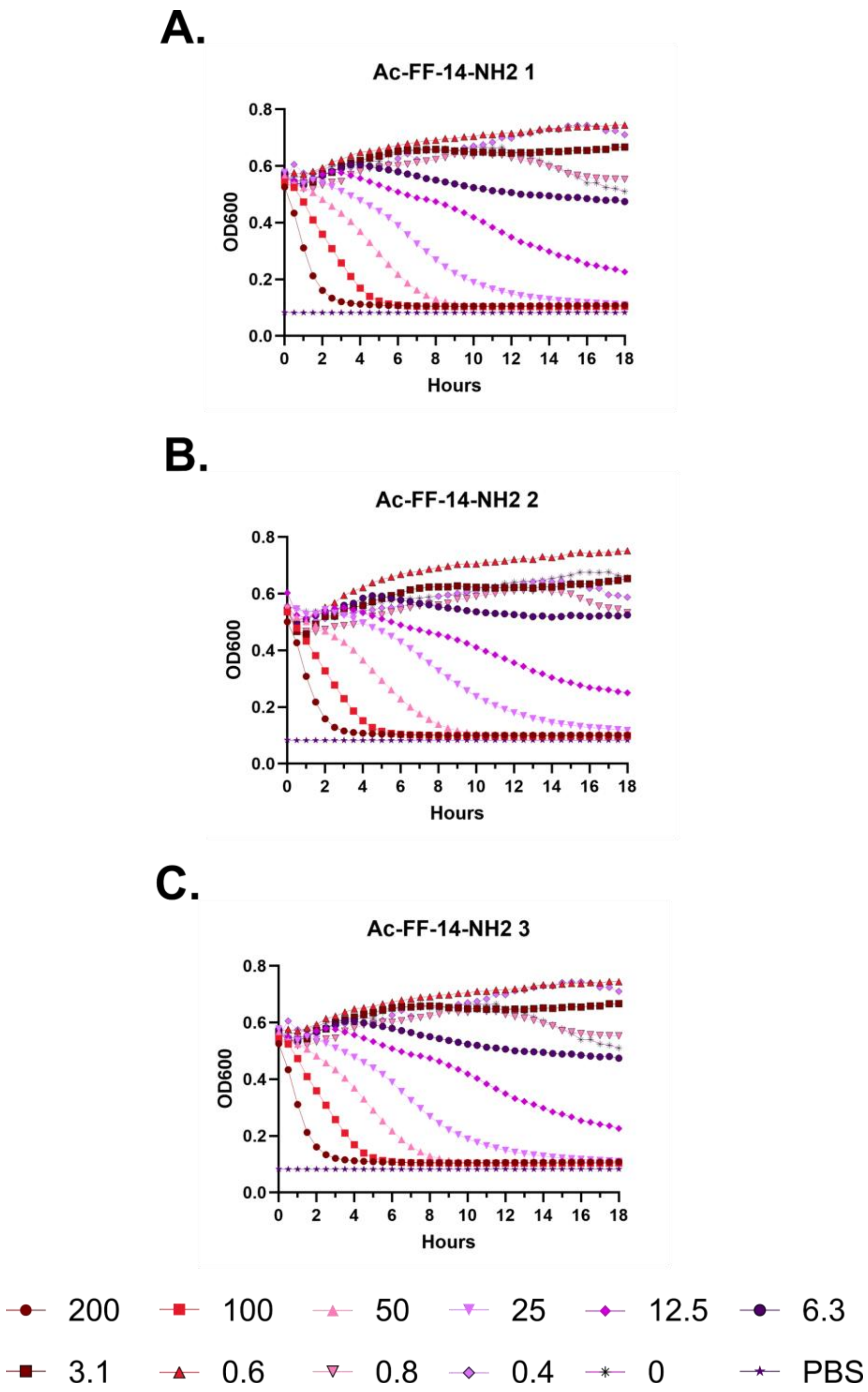
A.
Ac-FF-14-NH2 1
OD600
Hours
0.0
0.2
0.4
0.6
0.8
0
2
4
6
8
10
12
14
16
18
B.
Ac-FF-14-NH2 2
OD600
Hours
C.
Ac-FF-14-NH2 3
OD600
Hours
200
100
50
25
12.5
6.3
3.1
0.6
0.8
0.4
0
PBS

**Figure S54 – (#22) Ac-FF-14-$NH_2$ kinetic hemolysis assay. A.** Defibrinated sheep blood was incubated in the presence of the indicated peptide concentrations at 37 °C with monitoring of OD600 over 18 hours, where decreasing OD600 indicates hemolysis and each panel indicates an independent experiment. Peptide concentrations are in µM.

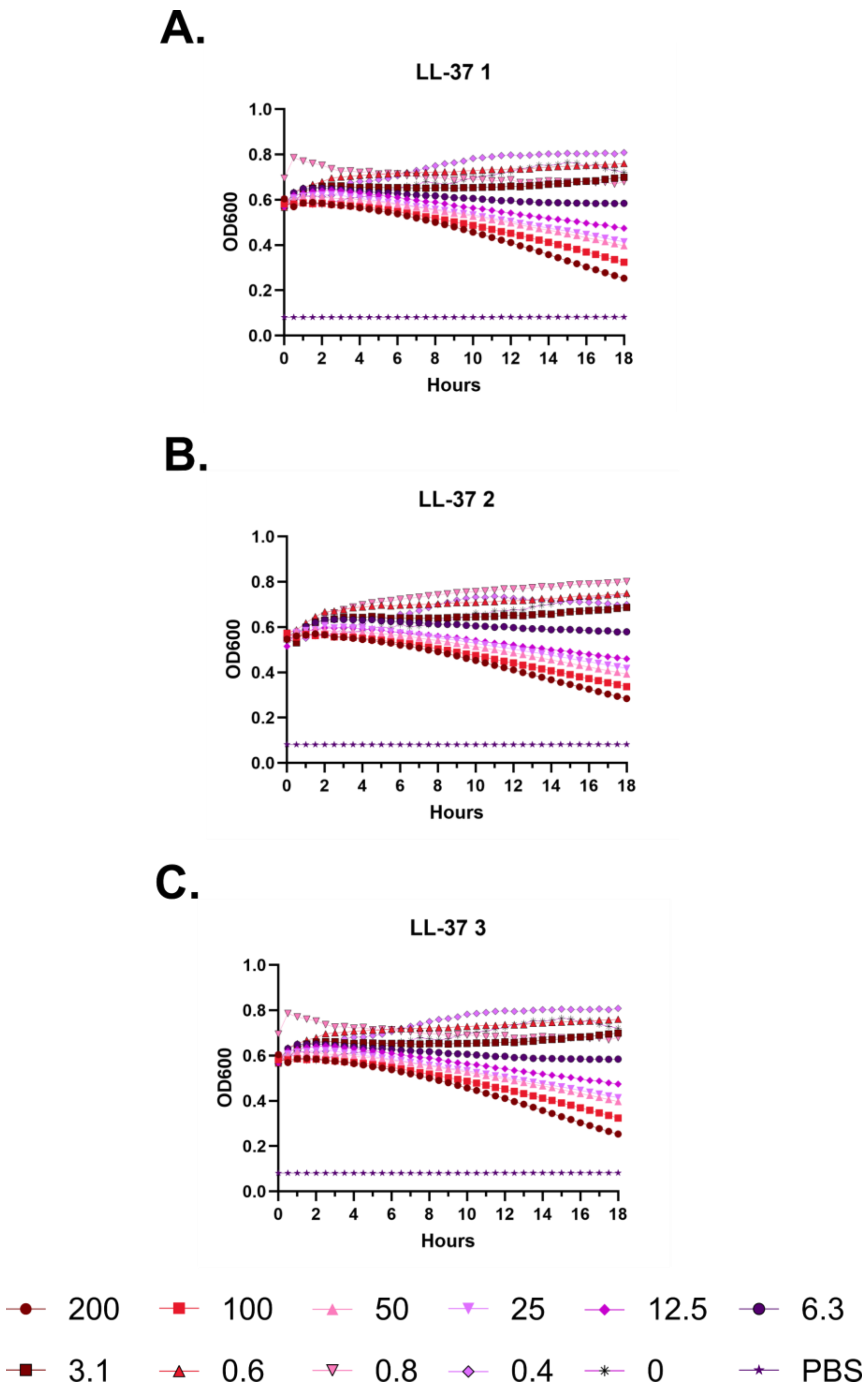
A.
LL-37 1
OD600
Hours
B.
LL-37 2
OD600
Hours
C.
LL-37 3
OD600
Hours
200
100
50
25
12.5
6.3
3.1
0.6
0.8
0.4
0
PBS

**Figure S55 – (#1) LL-37 kinetic hemolysis assay. A.** Defibrinated sheep blood was incubated in the presence of the indicated peptide concentrations at 37 °C with monitoring of OD600 over 18 hours, where decreasing OD600 indicates hemolysis and each panel indicates an independent experiment. Peptide concentrations are in μM.

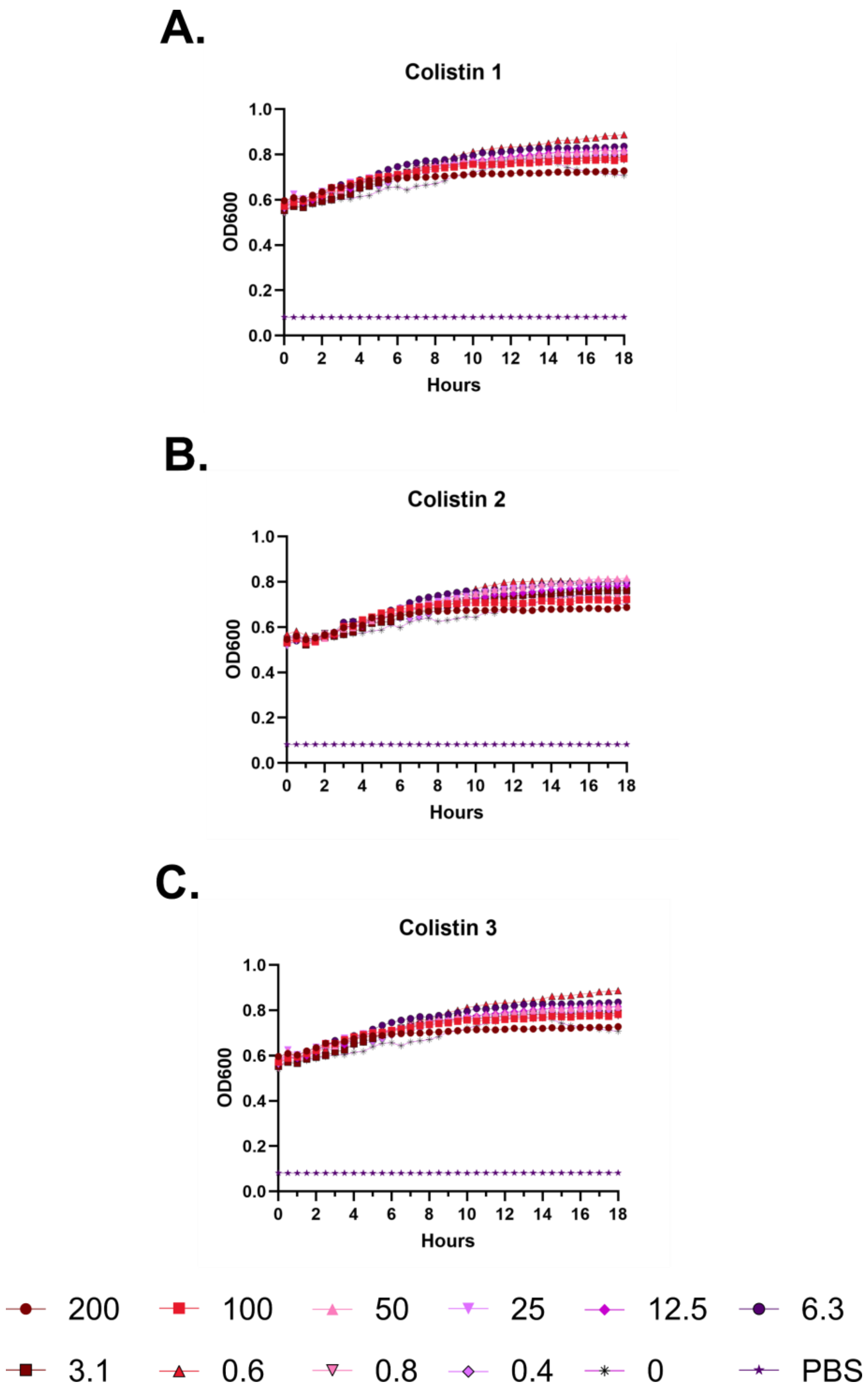
A.
Colistin 1
OD600
Hours
B.
Colistin 2
OD600
Hours
C.
Colistin 3
OD600
Hours
200
100
50
25
12.5
6.3
3.1
0.6
0.8
0.4
0
PBS

**Figure S56 – Colistin kinetic hemolysis assay. A.** Defibrinated sheep blood was incubated in the presence of the indicated peptide concentrations at 37 °C with monitoring of OD600 over 18 hours, where decreasing OD600 indicates hemolysis and each panel indicates an independent experiment. Peptide concentrations are in μM.

**A.**

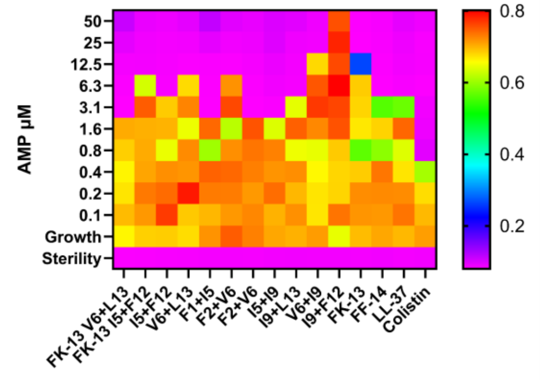


**B.**

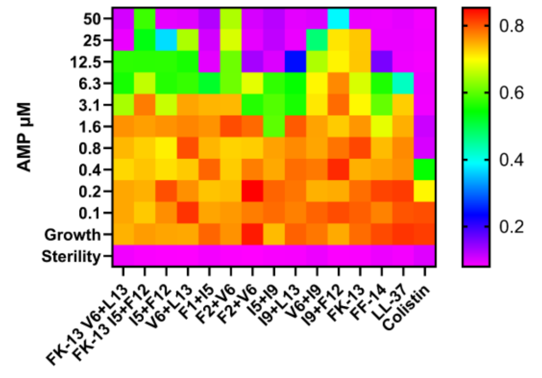


**C.**

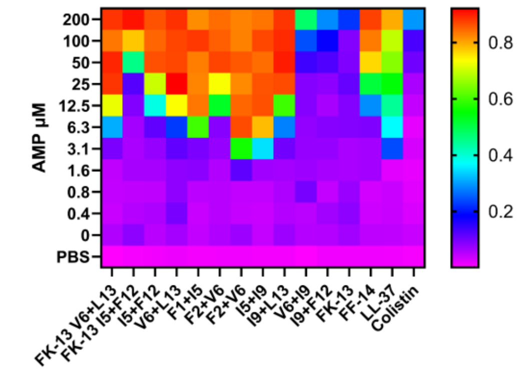


**D.**

| | |
|---|---|
| FK-13 6+13 | -FKRI**C**QRIKDF**C**R |
| FK-13 5+12 | -FKR**C**VQRIKD**C**LR |
| I5+F12 | FFKR**C**VQRIKD**C**LR |
| V6+L13 | FFKRI**C**QRIKDF**C**R |
| F1+I5 | **C**FKR**C**VQRIKDFLR |
| F2+V6 | F**C**KRI**C**QRIKDFLR |
| F2+V6 | F**C**KRI**C**QRIKDFLR |
| I5+I9 | FFKR**C**VQR**C**KDFLR |
| I9+L13 | FFKRIVQR**C**KDF**C**R |
| V6+I9 | FFKRI**C**QR**C**KDFLR |
| I9+F12 | FFKRIVQR**C**KD**C**LR |
| FK-13 | -FKRIVQRIKDFLR |
| FF-14 | FFKRIVQRIKDFLR |

**Figure S57 – Heatmap depiction of antimicrobial and hemolytic activity associated with perfluoroaryl stapled derivatives. A.** Antimicrobial susceptibility testing showing the activity of each derivative against *E. coli* (ATCC 25922). In the heatmap plate depiction, cooler colors indicate low or absent bacterial growth as detected by the OD600 measure at 18 hours of incubation at 37 °C. **B.** As in **A.**, but with *P. aeruginosa* (ATCC 27853). **C.** Endpoint hemolysis assays in which sheep RBCs in PBS were incubated with the indicated peptide for 18 hours at 37 °C. Cool colors indicate little or no release of hemoglobin as measured at 414 nm, where all levels are scaled to 100% hemolysis freeze-thaw controls. Levels throughout are the median across three independent experiments. **D.** Alignment showing the stapled Cys residues used in each derivative.

A.

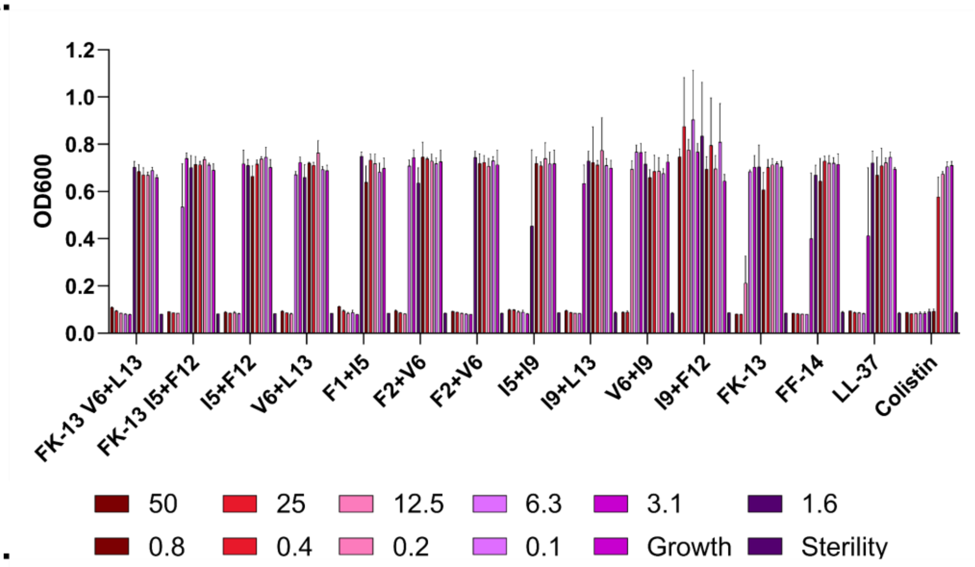
OD600
1.2
1.0
0.8
0.6
0.4
0.2
0.0
FK-13 V6+L13
FK-13 I5+F12
I5+F12
V6+L13
F1+I5
F2+V6
F2+V6
I5+I9
I9+L13
V6+I9
I9+F12
FK-13
FF-14
LL-37
Colistin
50
25
12.5
6.3
3.1
1.6
0.8
0.4
0.2
0.1
Growth
Sterility

B.

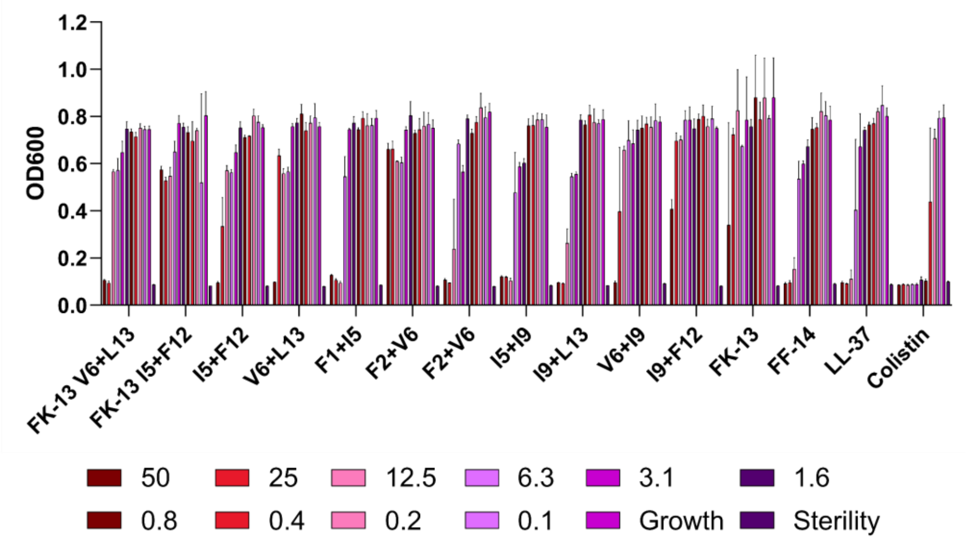
OD600
1.2
1.0
0.8
0.6
0.4
0.2
0.0
FK-13 V6+L13
FK-13 I5+F12
I5+F12
V6+L13
F1+I5
F2+V6
F2+V6
I5+I9
I9+L13
V6+I9
I9+F12
FK-13
FF-14
LL-37
Colistin
50
25
12.5
6.3
3.1
1.6
0.8
0.4
0.2
0.1
Growth
Sterility

C.

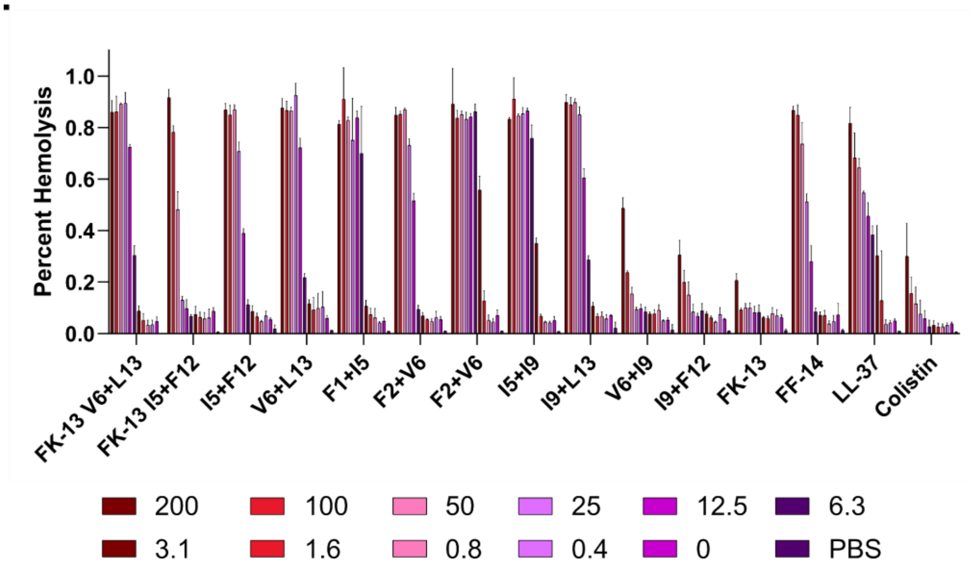
Percent Hemolysis
1.0
0.8
0.6
0.4
0.2
0.0
FK-13 V6+L13
FK-13 I5+F12
I5+F12
V6+L13
F1+I5
F2+V6
F2+V6
I5+I9
I9+L13
V6+I9
I9+F12
FK-13
FF-14
LL-37
Colistin
200
100
50
25
12.5
6.3
3.1
1.6
0.8
0.4
0
PBS

**Figure S58 – Histograms corresponding to Figure 4**. **A.** Growth of *E. coli* (ATCC 25922) by OD600 at 18 hours at 37 °C. Error bars indicate the standard deviation of three independent experiments. **B.** As in **A.**, but with *P. aeruginosa* (ATCC 27853). **C.** Endpoint hemolysis assays following incubation of defibrinated sheep blood with the indicated peptide for 18 hours at 37 °C with subsequent quantification of hemoglobin released into the supernatant at 414 nm relative to a 100% hemolysis freeze-thaw control. Error bars indicate the standard deviation of three independent experiments. All peptide levels are given in μM.

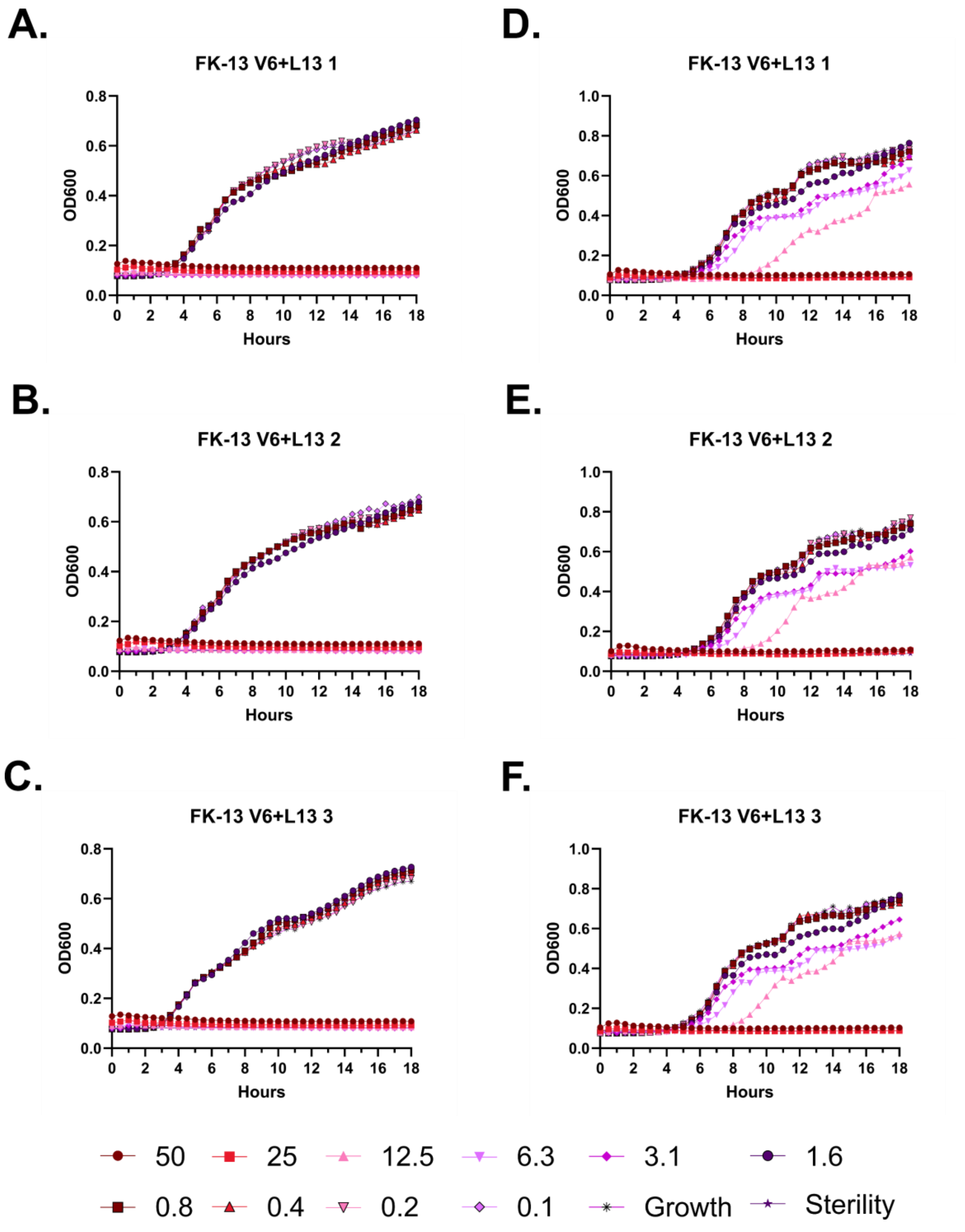

A.
FK-13 V6+L13 1
B.
FK-13 V6+L13 2
C.
FK-13 V6+L13 3
D.
FK-13 V6+L13 1
E.
FK-13 V6+L13 2
F.
FK-13 V6+L13 3
OD600
Hours
50
25
12.5
6.3
3.1
1.6
0.8
0.4
0.2
0.1
Growth
Sterility

**Figure S59 – Antimicrobial susceptibility curves for (#23) FK-13 V6+L13. A-C.** Independent experiments with *E. coli*. **D-F.** Independent experiments with *P. aeruginosa*. Growth over 18 hours at 37 °C with a gradient of peptide concentrations given in μM.

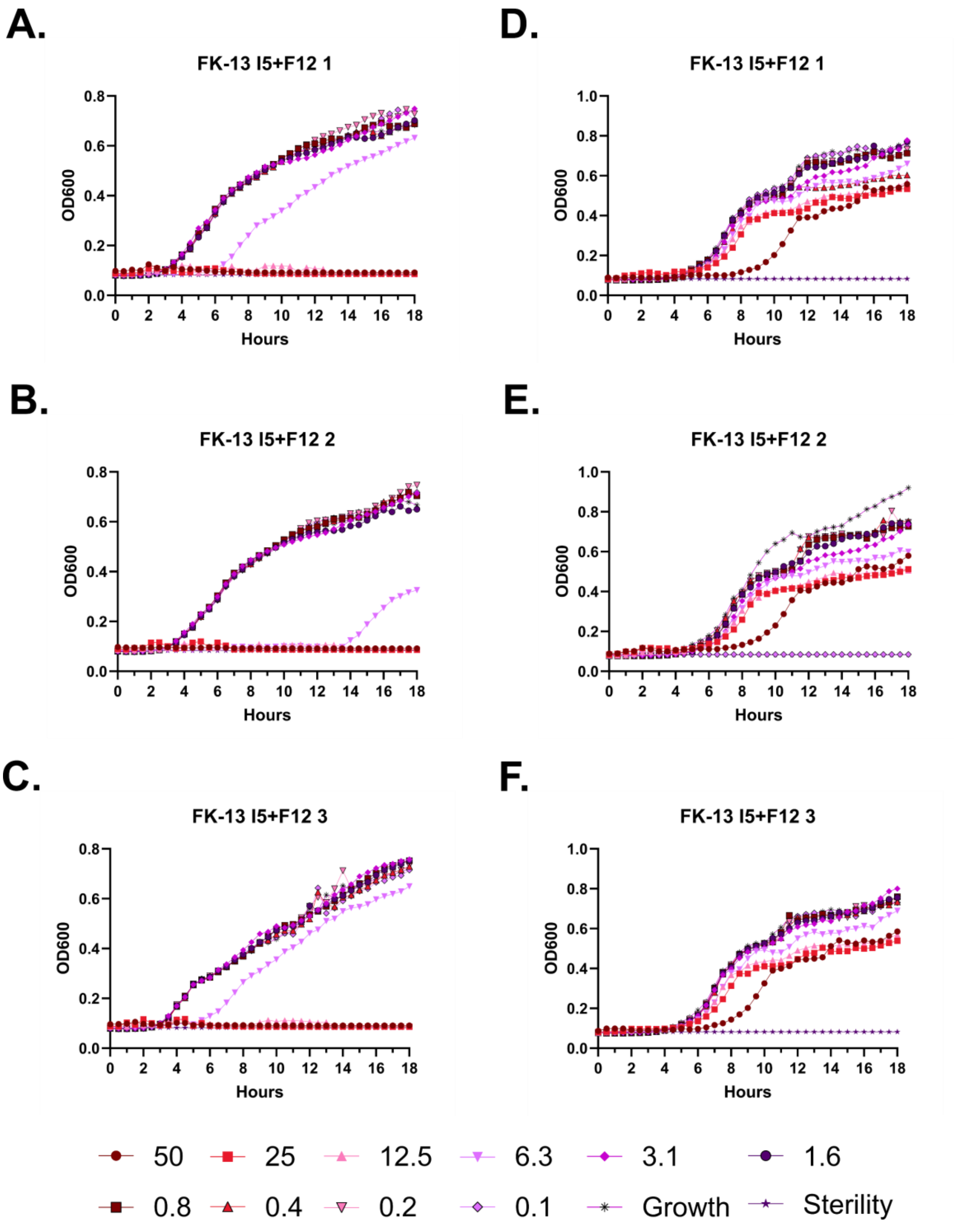
A.
FK-13 I5+F12 1
D.
FK-13 I5+F12 1
B.
FK-13 I5+F12 2
E.
FK-13 I5+F12 2
C.
FK-13 I5+F12 3
F.
FK-13 I5+F12 3
OD600
Hours
50
25
12.5
6.3
3.1
1.6
0.8
0.4
0.2
0.1
Growth
Sterility

**Figure S60 – Antimicrobial susceptibility curves for (#24) FK-13 I5+F12. A-C.** Independent experiments with *E. coli*. **D-F.** Independent experiments with *P. aeruginosa.* Growth over 18 hours at 37 °C with a gradient of peptide concentrations given in μM.

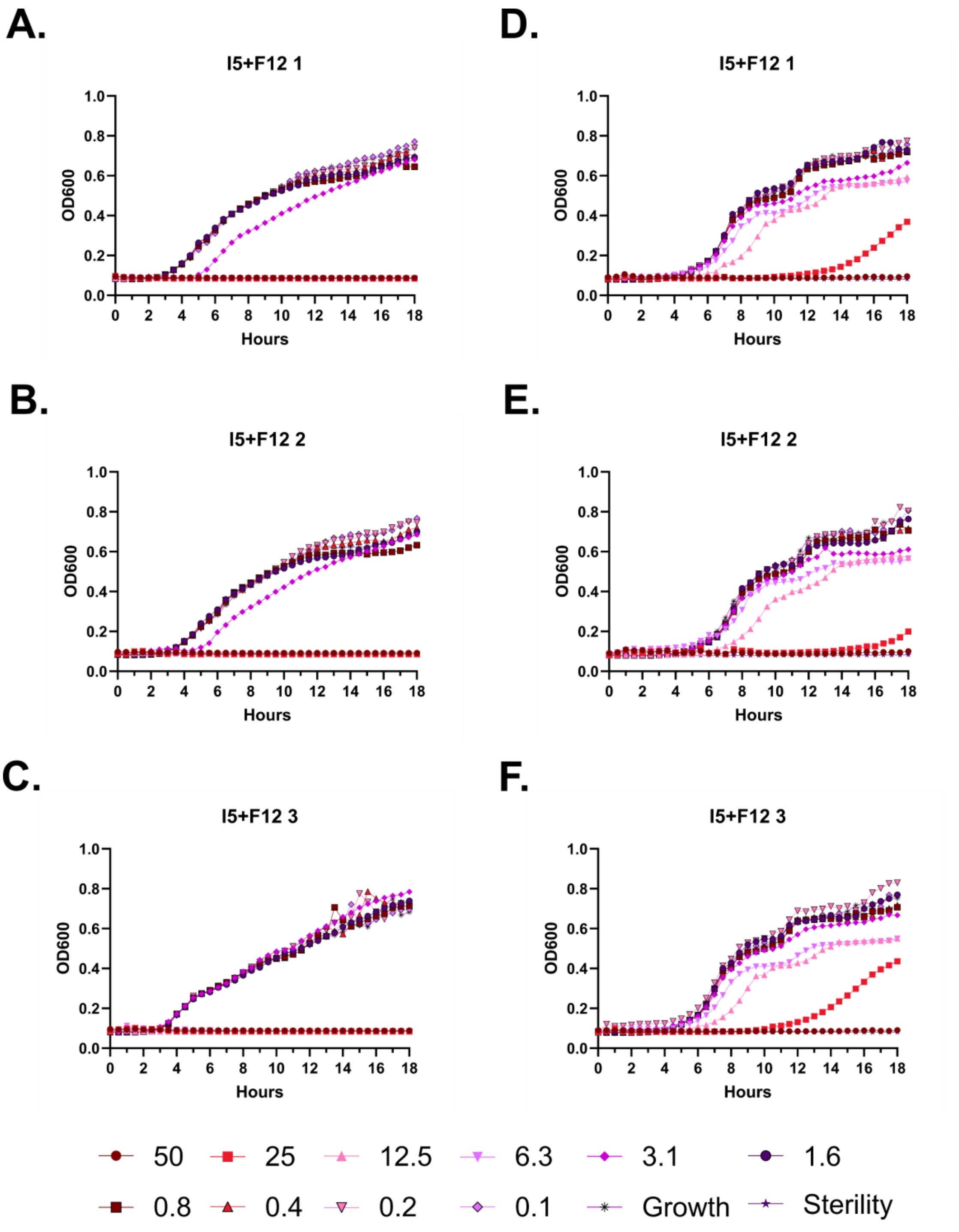
A.
I5+F12 1
B.
I5+F12 2
C.
I5+F12 3
D.
I5+F12 1
E.
I5+F12 2
F.
I5+F12 3
OD600
Hours
0.0
0.2
0.4
0.6
0.8
1.0
0
2
4
6
8
10
12
14
16
18
50
25
12.5
6.3
3.1
1.6
0.8
0.4
0.2
0.1
Growth
Sterility

**Figure S61 – Antimicrobial susceptibility curves for (#25) FF-14 I5+F12. A-C.** Independent experiments with *E. coli*. **D-F.** Independent experiments with *P. aeruginosa.* Growth over 18 hours at 37 °C with a gradient of peptide concentrations given in μM.

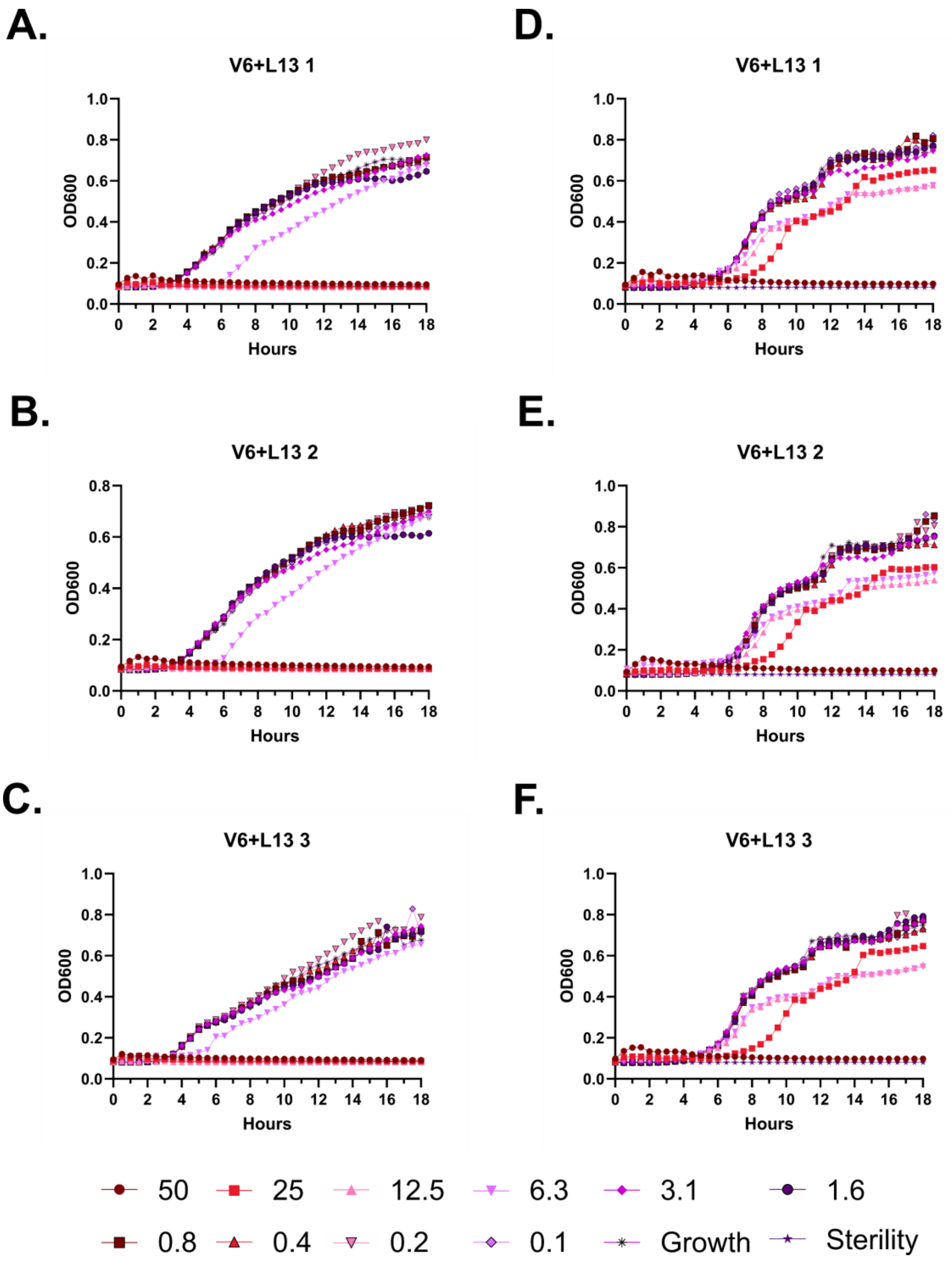
A.
V6+L13 1
D.
V6+L13 1
B.
V6+L13 2
E.
V6+L13 2
C.
V6+L13 3
F.
V6+L13 3
OD600
Hours
50
25
12.5
6.3
3.1
1.6
0.8
0.4
0.2
0.1
Growth
Sterility

**Figure S62 – Antimicrobial susceptibility curves for (#26) FF-14 V6+L13. A-C.** Independent experiments with *E. coli*. **D-F.** Independent experiments with *P. aeruginosa.* Growth over 18 hours at 37 °C with a gradient of peptide concentrations given in μM.

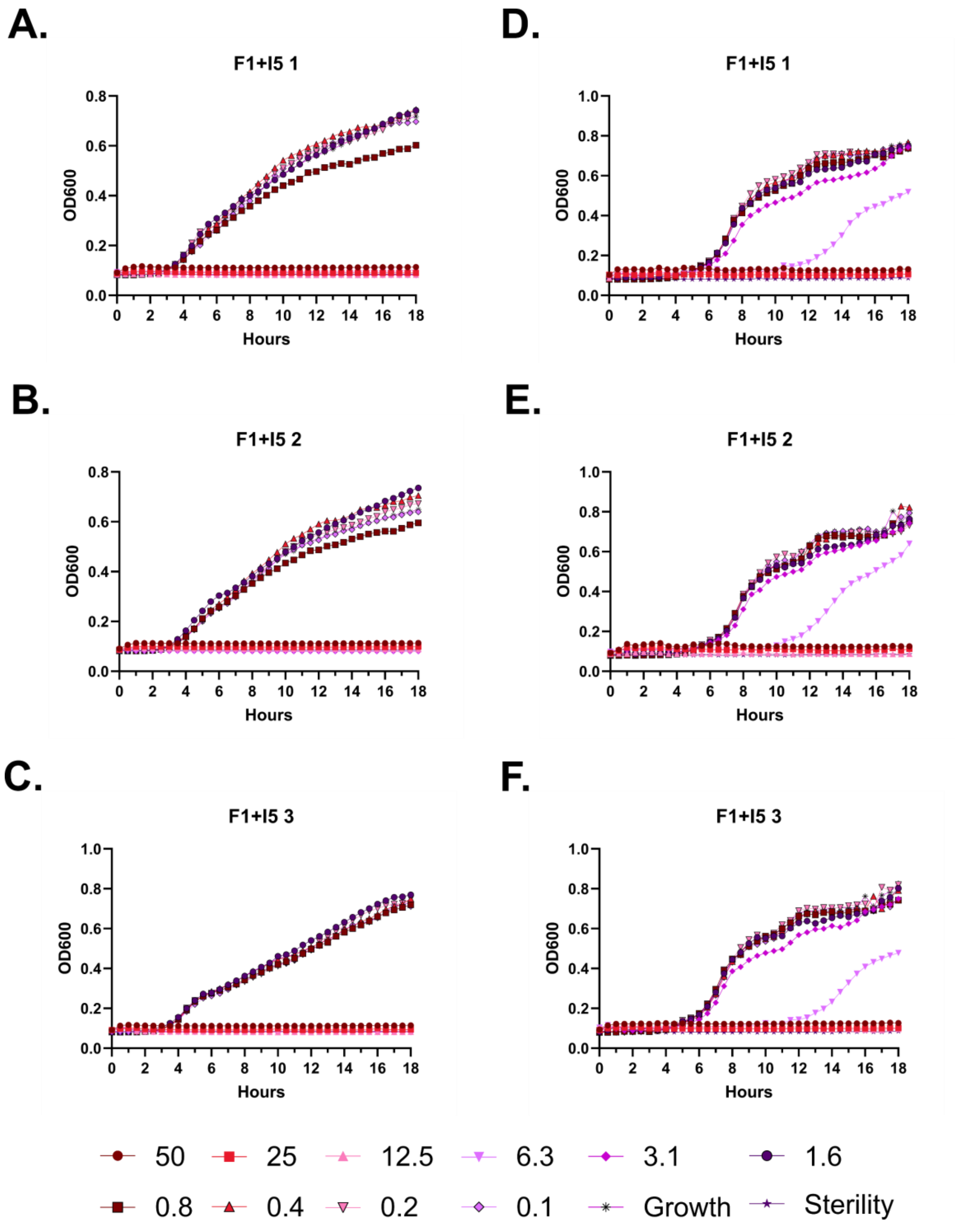
A.
F1+I5 1
OD600
Hours
B.
F1+I5 2
OD600
Hours
C.
F1+I5 3
OD600
Hours
D.
F1+I5 1
OD600
Hours
E.
F1+I5 2
OD600
Hours
F.
F1+I5 3
OD600
Hours
50
25
12.5
6.3
3.1
1.6
0.8
0.4
0.2
0.1
Growth
Sterility

**Figure S63 – Antimicrobial susceptibility curves for (#27) FF-14 F1+I5. A-C.** Independent experiments with *E. coli*. **D-F.** Independent experiments with *P. aeruginosa.* Growth over 18 hours at 37 °C with a gradient of peptide concentrations given in μM.

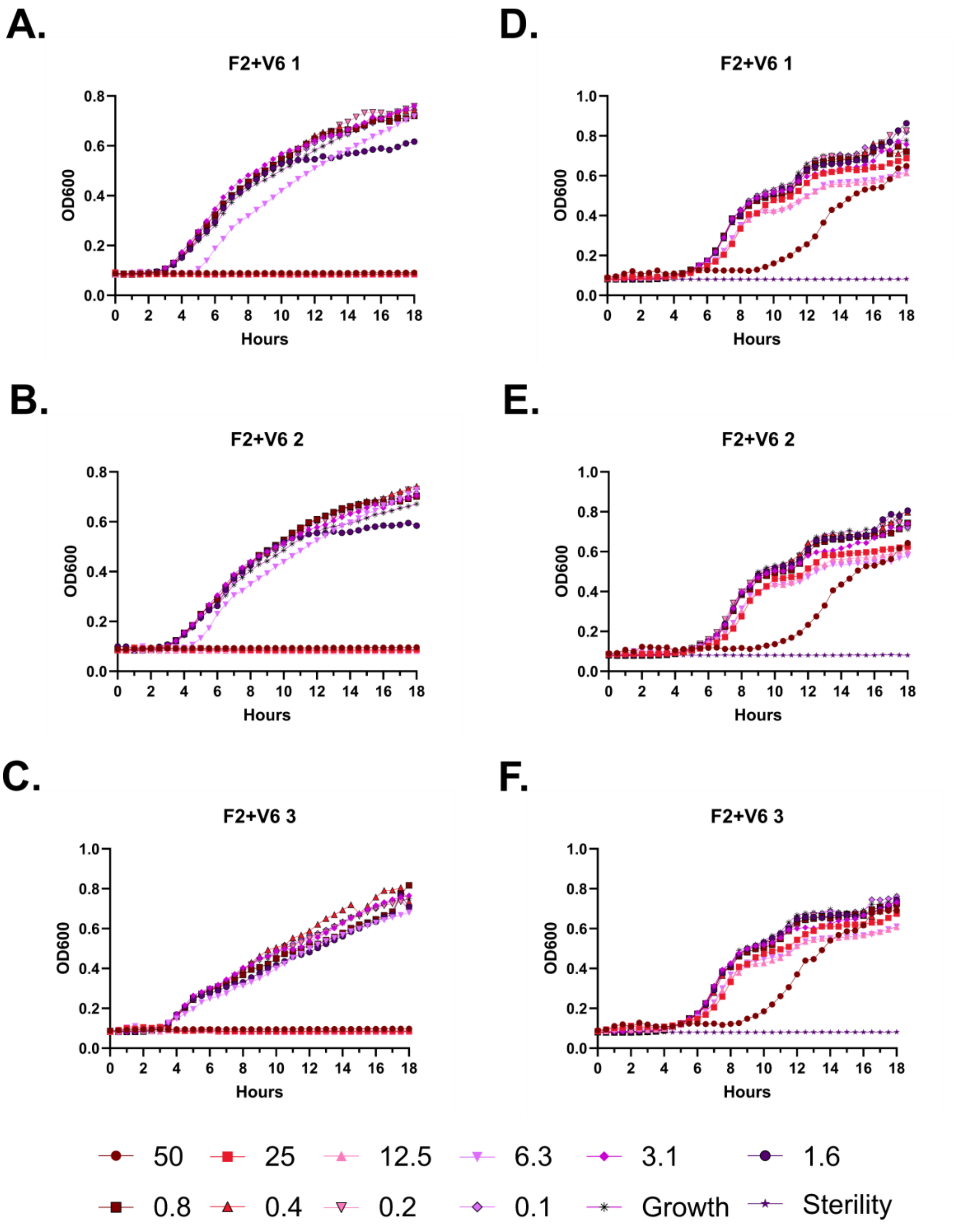
A.
F2+V6 1
OD600
Hours
D.
F2+V6 1
OD600
Hours
B.
F2+V6 2
OD600
Hours
E.
F2+V6 2
OD600
Hours
C.
F2+V6 3
OD600
Hours
F.
F2+V6 3
OD600
Hours
50
25
12.5
6.3
3.1
1.6
0.8
0.4
0.2
0.1
Growth
Sterility

**Figure S64 – Antimicrobial susceptibility curves for (#28) FF-14 F2+V6. A-C.** Independent experiments with *E. coli*. **D-F.** Independent experiments with *P. aeruginosa.* Growth over 18 hours at 37 °C with a gradient of peptide concentrations given in μM.

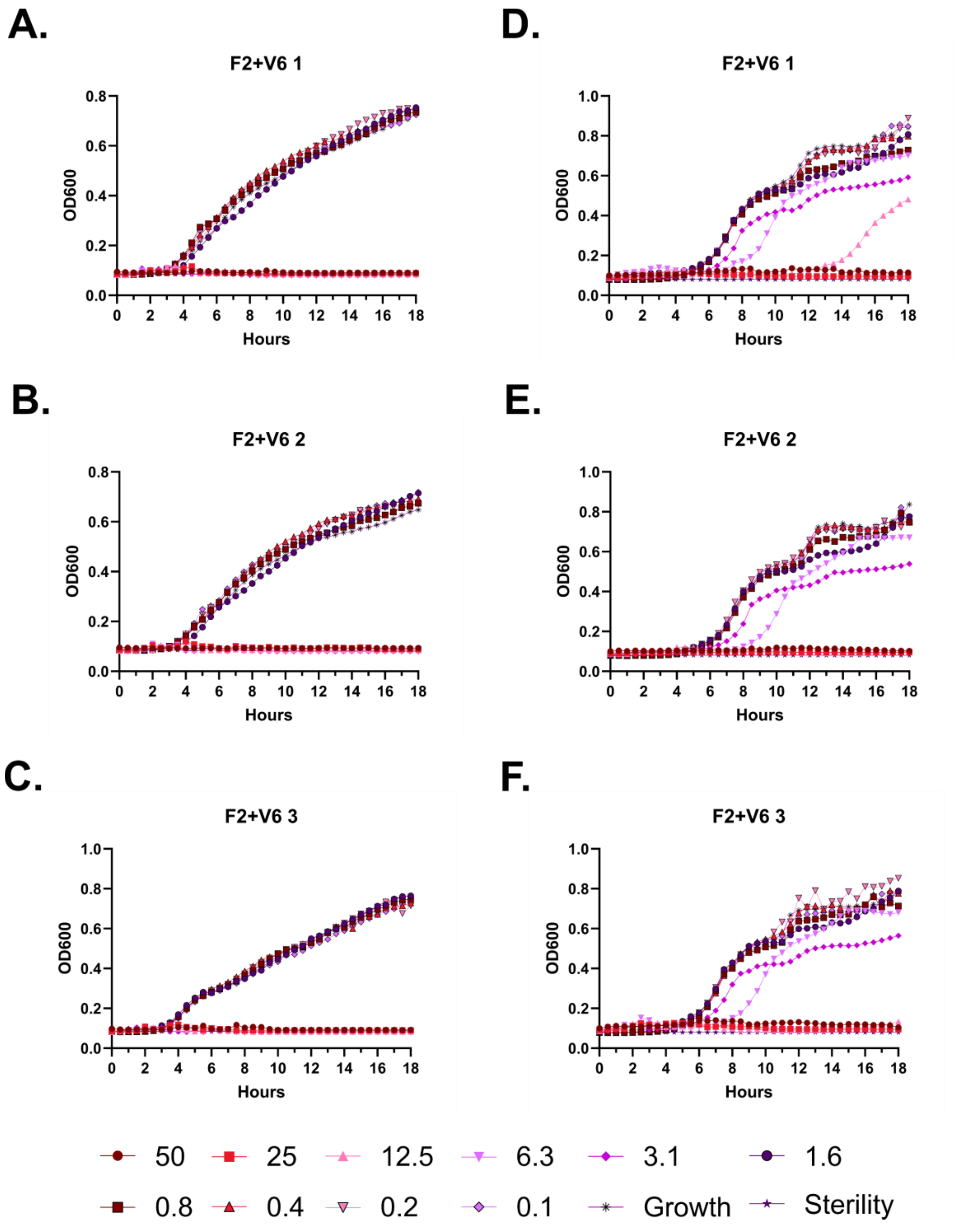

A.
F2+V6 1
OD600
Hours
B.
F2+V6 2
OD600
Hours
C.
F2+V6 3
OD600
Hours
D.
F2+V6 1
OD600
Hours
E.
F2+V6 2
OD600
Hours
F.
F2+V6 3
OD600
Hours
50
25
12.5
6.3
3.1
1.6
0.8
0.4
0.2
0.1
Growth
Sterility

**Figure S65 – Antimicrobial susceptibility curves for (#29) FF-14 F2+V6. A-C.** Independent experiments with *E. coli*. **D-F.** Independent experiments with *P. aeruginosa.* Growth over 18 hours at 37 °C with a gradient of peptide concentrations given in μM.

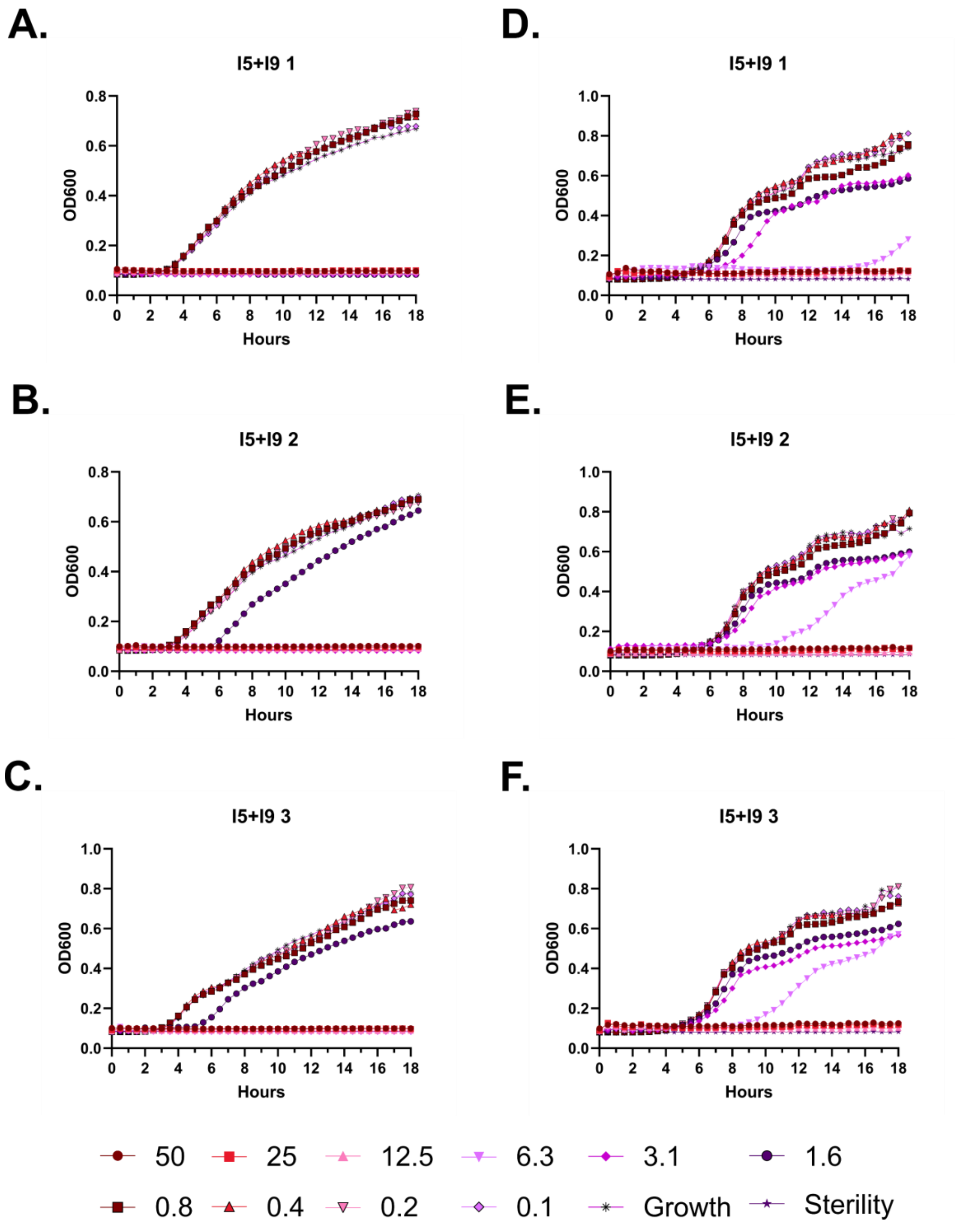
A.
I5+I9 1
D.
I5+I9 1
B.
I5+I9 2
E.
I5+I9 2
C.
I5+I9 3
F.
I5+I9 3
OD600
Hours
50
25
12.5
6.3
3.1
1.6
0.8
0.4
0.2
0.1
Growth
Sterility

**Figure S66 – Antimicrobial susceptibility curves for (#30) FF-14 I5+I9. A-C.** Independent experiments with *E. coli*. **D-F.** Independent experiments with *P. aeruginosa.* Growth over 18 hours at 37 °C with a gradient of peptide concentrations given in μM.

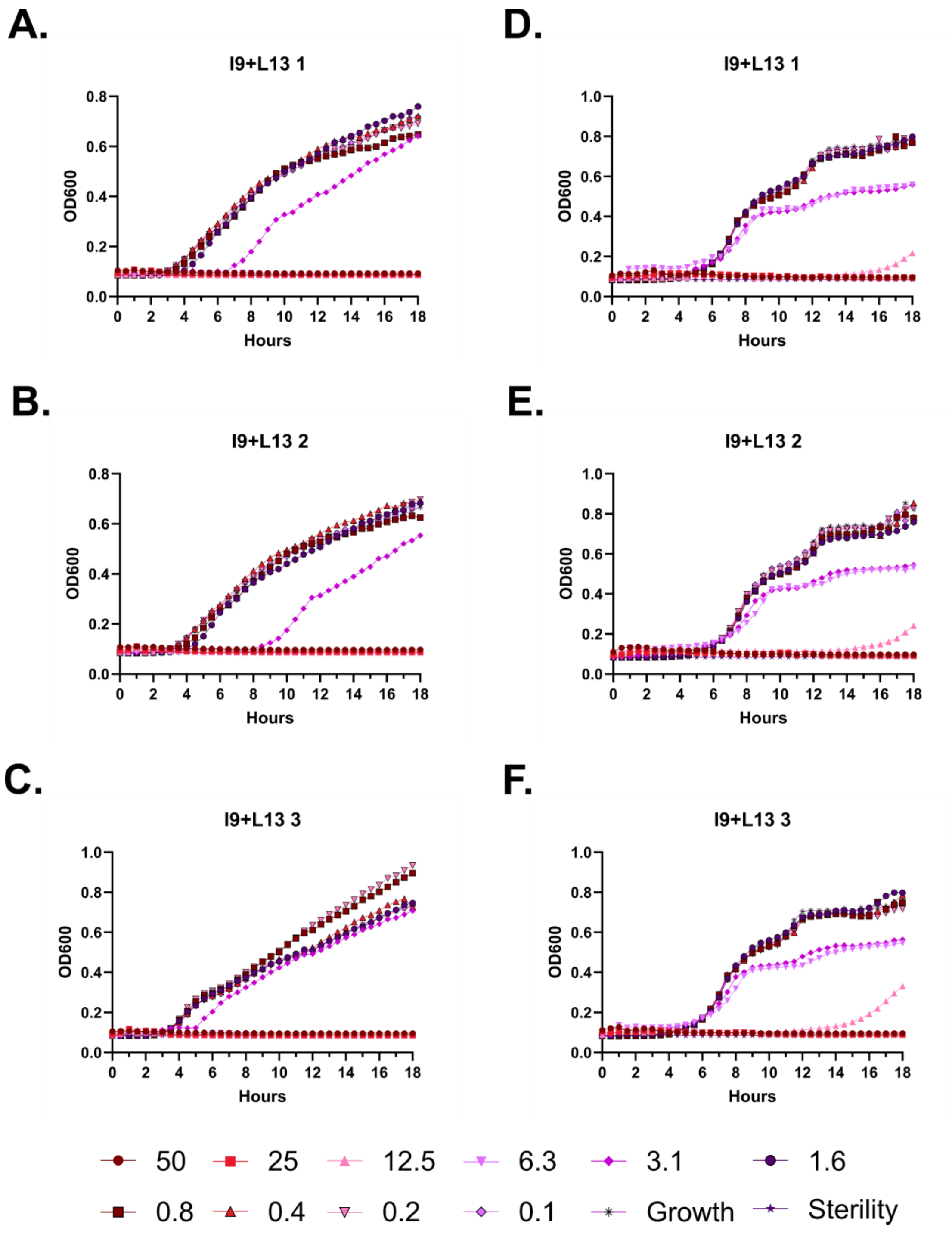
A.
I9+L13 1
B.
I9+L13 2
C.
I9+L13 3
D.
I9+L13 1
E.
I9+L13 2
F.
I9+L13 3
OD600
Hours
50
25
12.5
6.3
3.1
1.6
0.8
0.4
0.2
0.1
Growth
Sterility

**Figure S67 – Antimicrobial susceptibility curves for (#31) FF-14 I9+L13. A-C.** Independent experiments with *E. coli*. **D-F.** Independent experiments with *P. aeruginosa.* Growth over 18 hours at 37 °C with a gradient of peptide concentrations given in μM.

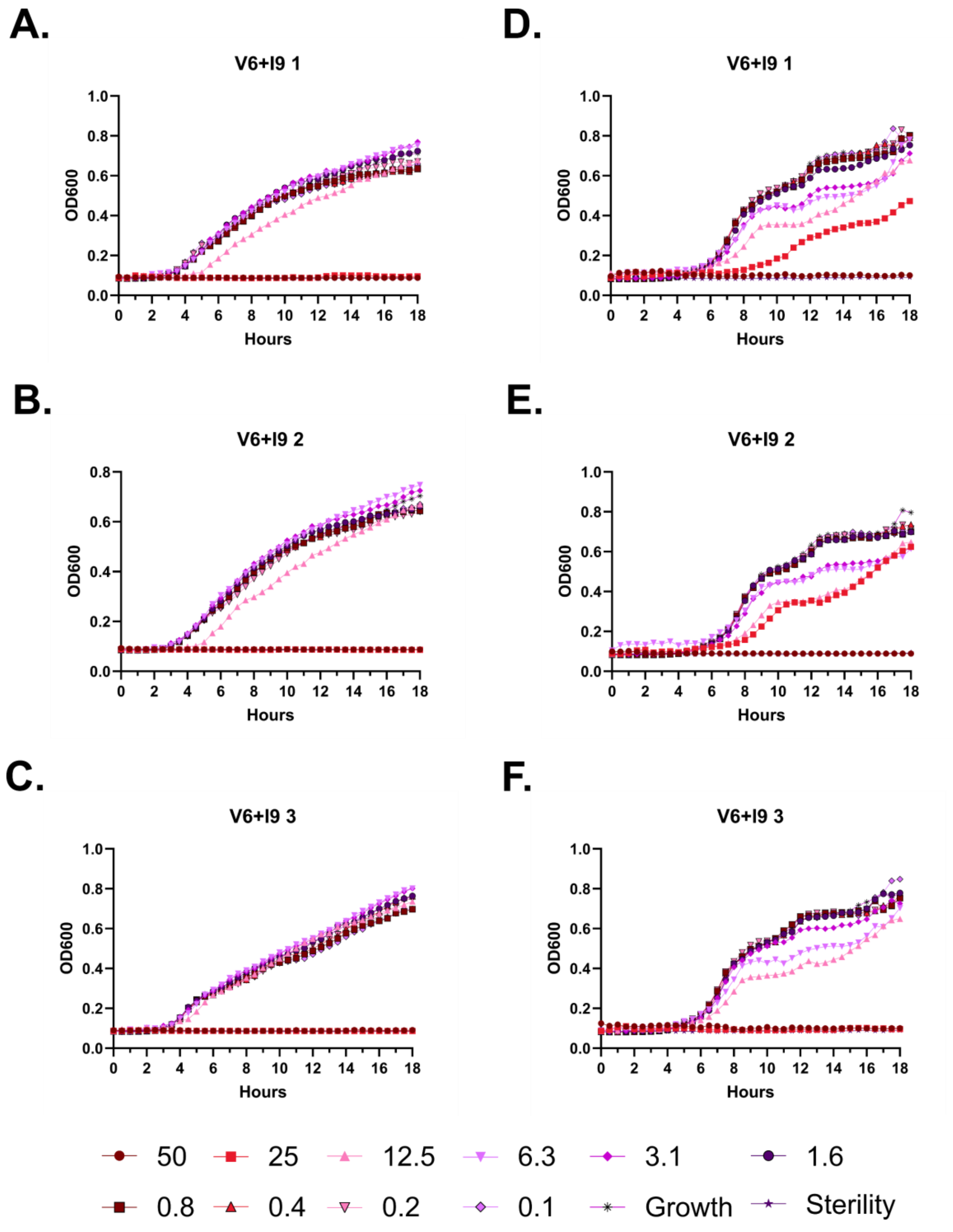
A.
V6+I9 1
OD600
Hours
B.
V6+I9 2
OD600
Hours
C.
V6+I9 3
OD600
Hours
D.
V6+I9 1
OD600
Hours
E.
V6+I9 2
OD600
Hours
F.
V6+I9 3
OD600
Hours
50
25
12.5
6.3
3.1
1.6
0.8
0.4
0.2
0.1
Growth
Sterility

**Figure S68 – Antimicrobial susceptibility curves for (#32) FF-14 V6+I9. A-C.** Independent experiments with *E. coli*. **D-F.** Independent experiments with *P. aeruginosa.* Growth over 18 hours at 37 °C with a gradient of peptide concentrations given in μM.

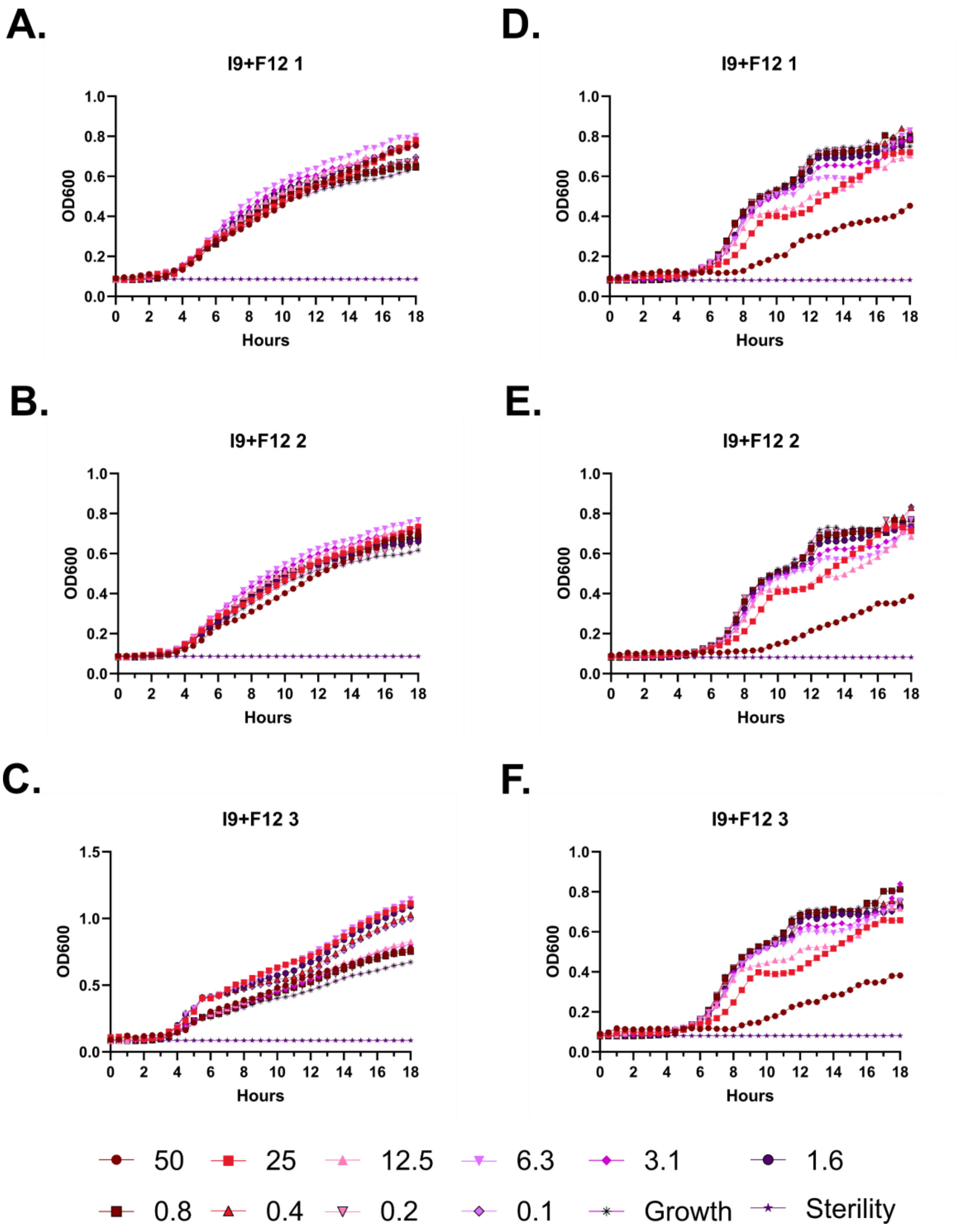
A.
I9+F12 1
D.
I9+F12 1
B.
I9+F12 2
E.
I9+F12 2
C.
I9+F12 3
F.
I9+F12 3
OD600
Hours
50
25
12.5
6.3
3.1
1.6
0.8
0.4
0.2
0.1
Growth
Sterility

**Figure S69 – Antimicrobial susceptibility curves for (#33) FF-14 I9+F12. A-C.** Independent experiments with *E. coli*. **D-F.** Independent experiments with *P. aeruginosa.* Growth over 18 hours at 37 °C with a gradient of peptide concentrations given in μM.

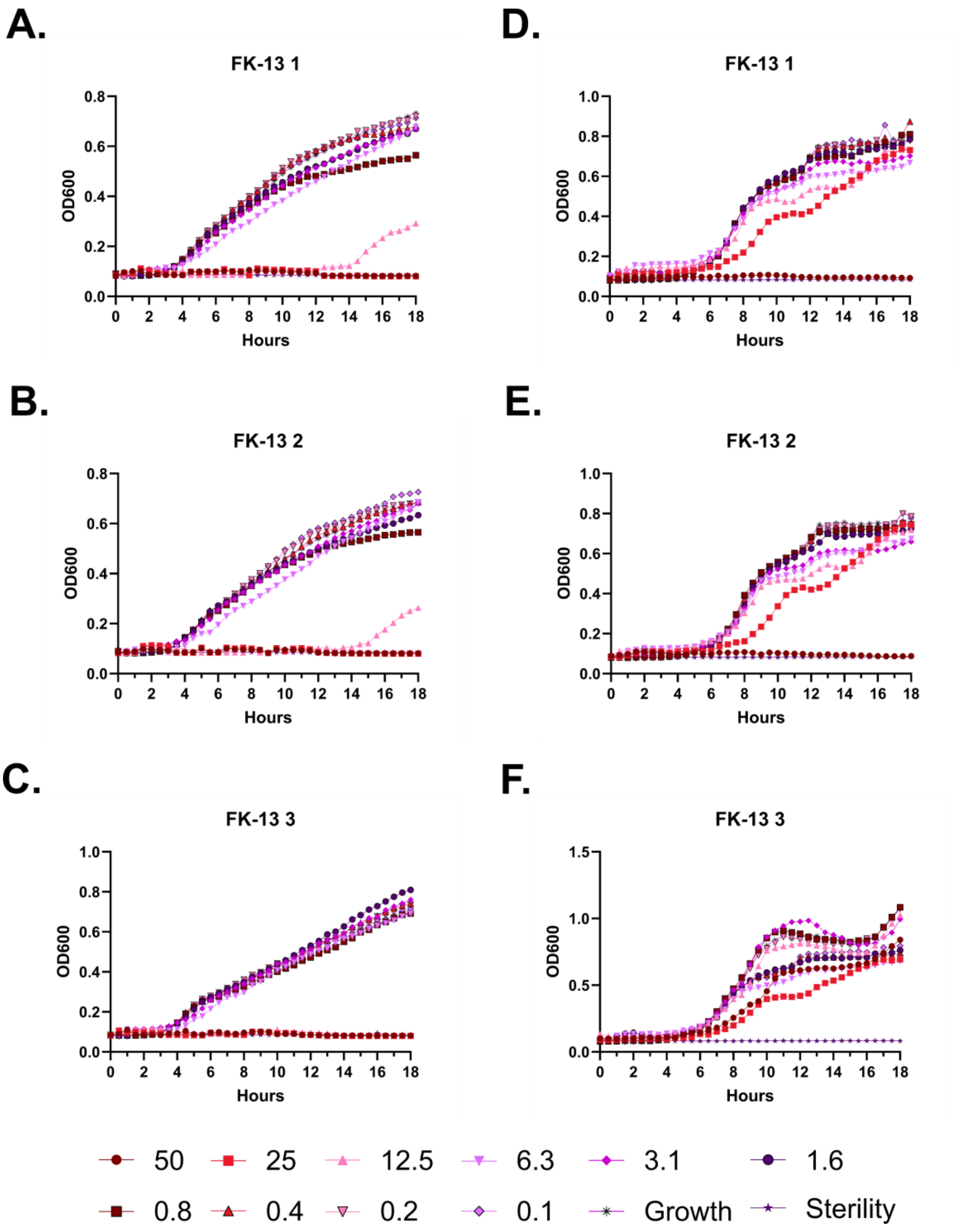
A.
FK-13 1
D.
FK-13 1
B.
FK-13 2
E.
FK-13 2
C.
FK-13 3
F.
FK-13 3
OD600
Hours
50
25
12.5
6.3
3.1
1.6
0.8
0.4
0.2
0.1
Growth
Sterility

**Figure S70 – Antimicrobial susceptibility curves for (#7) FK-13. A-C.** Independent experiments with *E. coli*. **D-F.** Independent experiments with *P. aeruginosa.* Growth over 18 hours at 37 °C with a gradient of peptide concentrations given in μM.

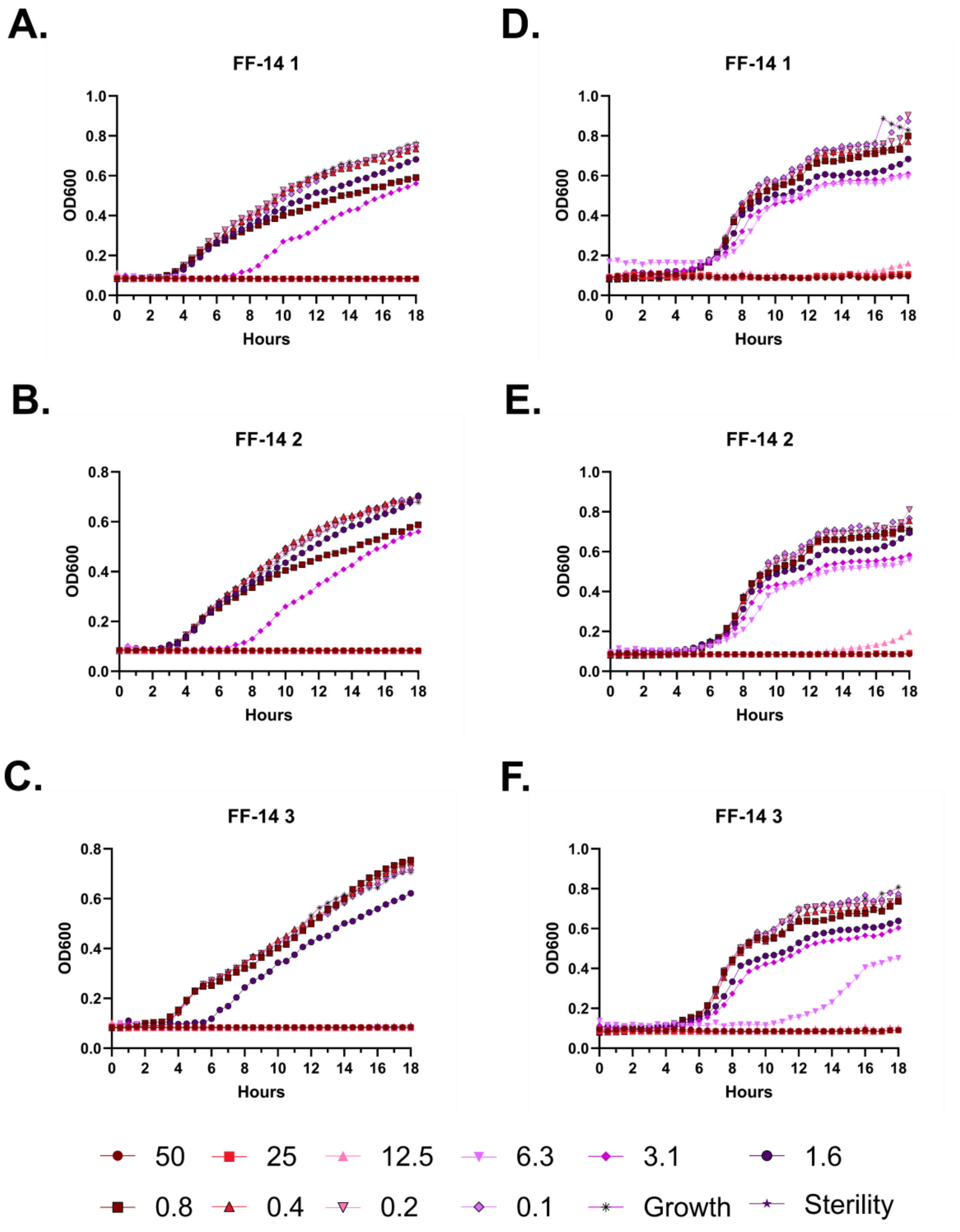
A.
FF-14 1
OD600
Hours
D.
FF-14 1
OD600
Hours
B.
FF-14 2
OD600
Hours
E.
FF-14 2
OD600
Hours
C.
FF-14 3
OD600
Hours
F.
FF-14 3
OD600
Hours
50
25
12.5
6.3
3.1
1.6
0.8
0.4
0.2
0.1
Growth
Sterility

**Figure S71 – Antimicrobial susceptibility curves for (#18) FF-14. A-C.** Independent experiments with *E. coli*. **D-F.** Independent experiments with *P. aeruginosa.* Growth over 18 hours at 37 °C with a gradient of peptide concentrations given in μM.

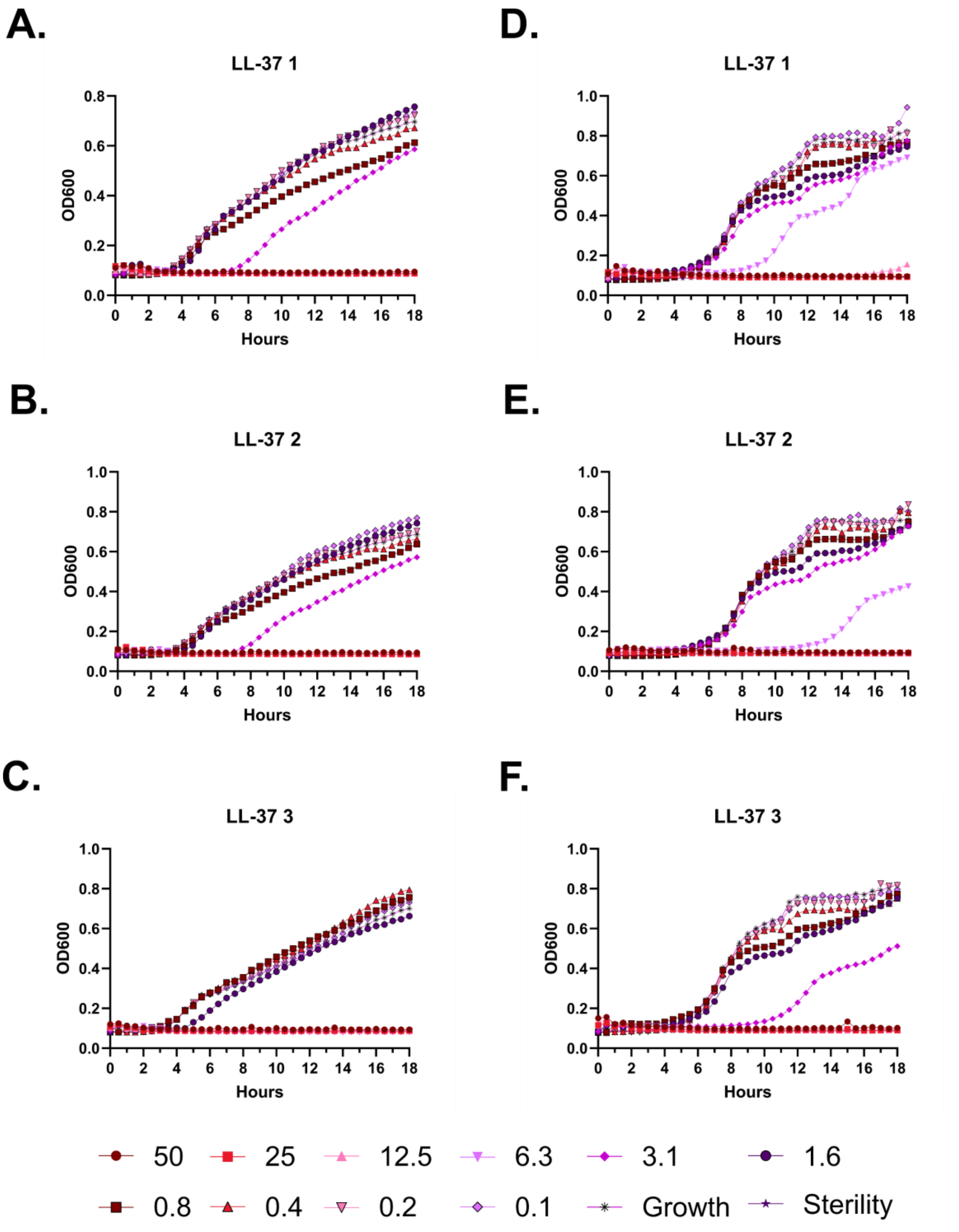
A.
LL-37 1
D.
LL-37 1
B.
LL-37 2
E.
LL-37 2
C.
LL-37 3
F.
LL-37 3
OD600
Hours
50
25
12.5
6.3
3.1
1.6
0.8
0.4
0.2
0.1
Growth
Sterility

**Figure S72 – Antimicrobial susceptibility curves for (#1) LL-37. A-C.** Independent experiments with *E. coli*. **D-F.** Independent experiments with *P. aeruginosa.* Growth over 18 hours at 37 °C with a gradient of peptide concentrations given in μM.

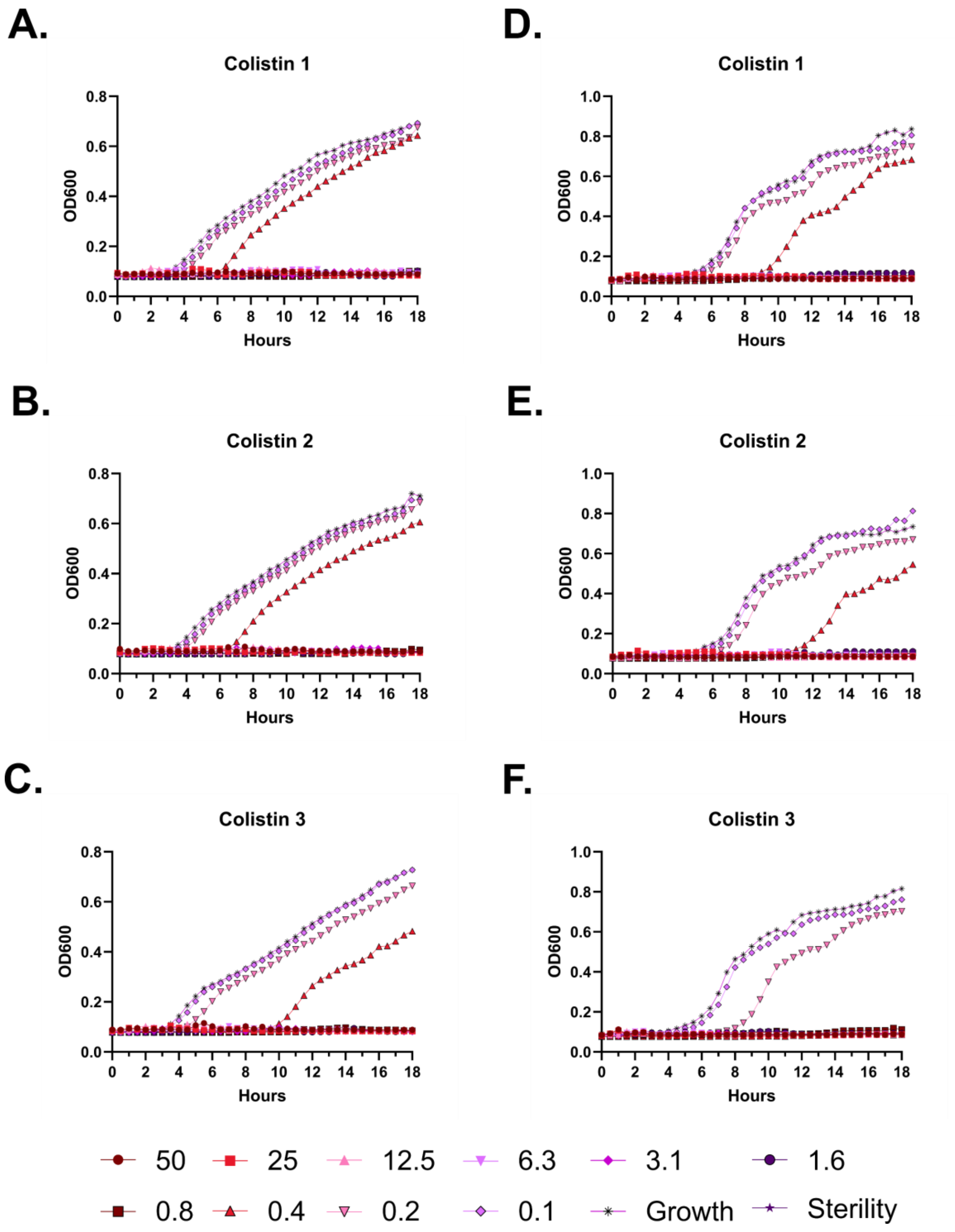

A.
Colistin 1
B.
Colistin 2
C.
Colistin 3
D.
Colistin 1
E.
Colistin 2
F.
Colistin 3
OD600
Hours
50
25
12.5
6.3
3.1
1.6
0.8
0.4
0.2
0.1
Growth
Sterility

**Figure S73 – Antimicrobial susceptibility curves for colistin. A-C.** Independent experiments with *E. coli*. **D-F.** Independent experiments with *P. aeruginosa.* Growth over 18 hours at 37 °C with a gradient of peptide concentrations given in μM.

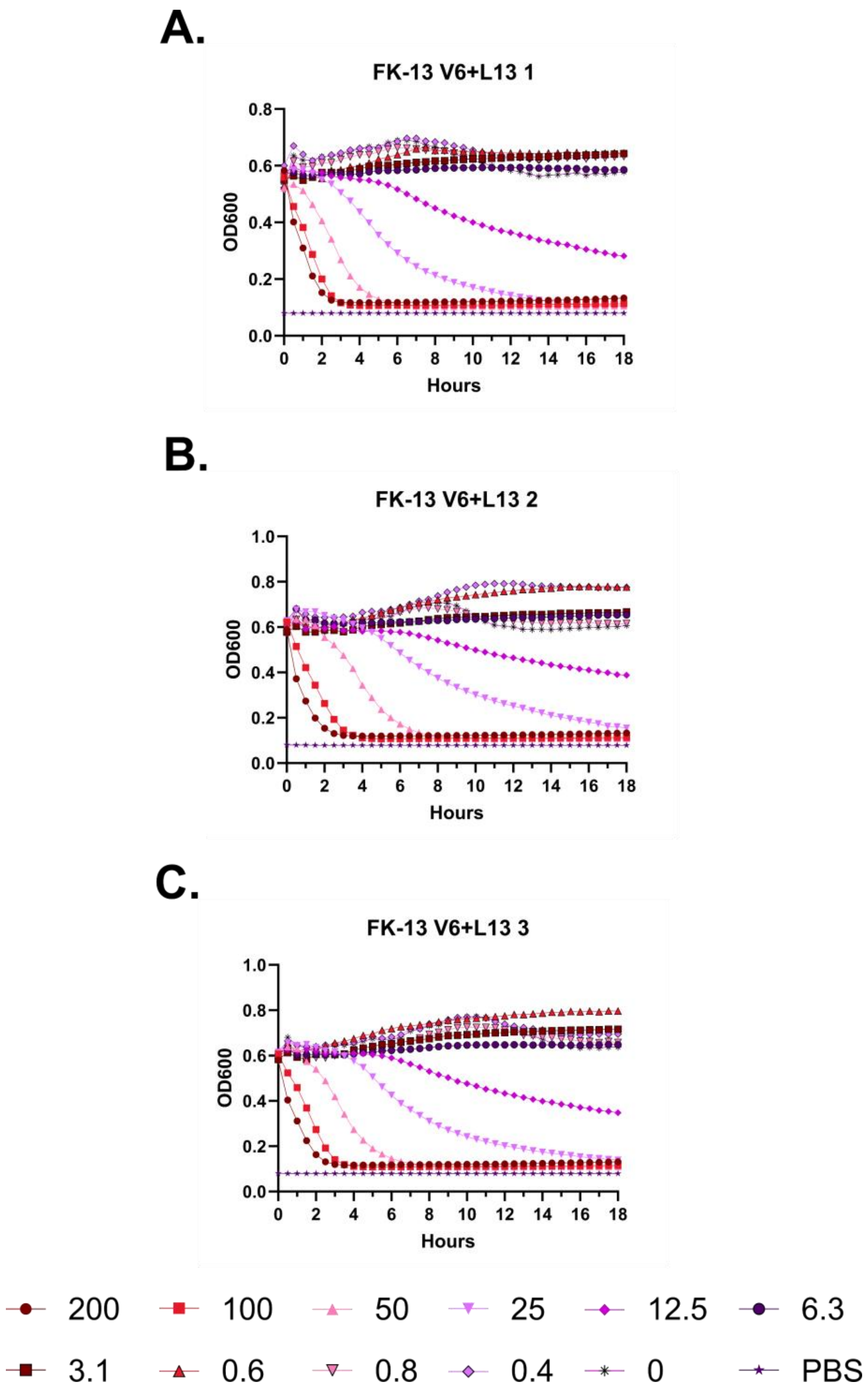
A.
FK-13 V6+L13 1
OD600
Hours
B.
FK-13 V6+L13 2
OD600
Hours
C.
FK-13 V6+L13 3
OD600
Hours
200
100
50
25
12.5
6.3
3.1
0.6
0.8
0.4
0
PBS

**Figure S74 – (#23) FK-13 V6+L13 kinetic hemolysis assay. A.** Defibrinated sheep blood was incubated in the presence of the indicated peptide concentrations at 37 °C with monitoring of OD600 over 18 hours, where decreasing OD600 indicates hemolysis and each panel indicates an independent experiment. Peptide concentrations are in μM.

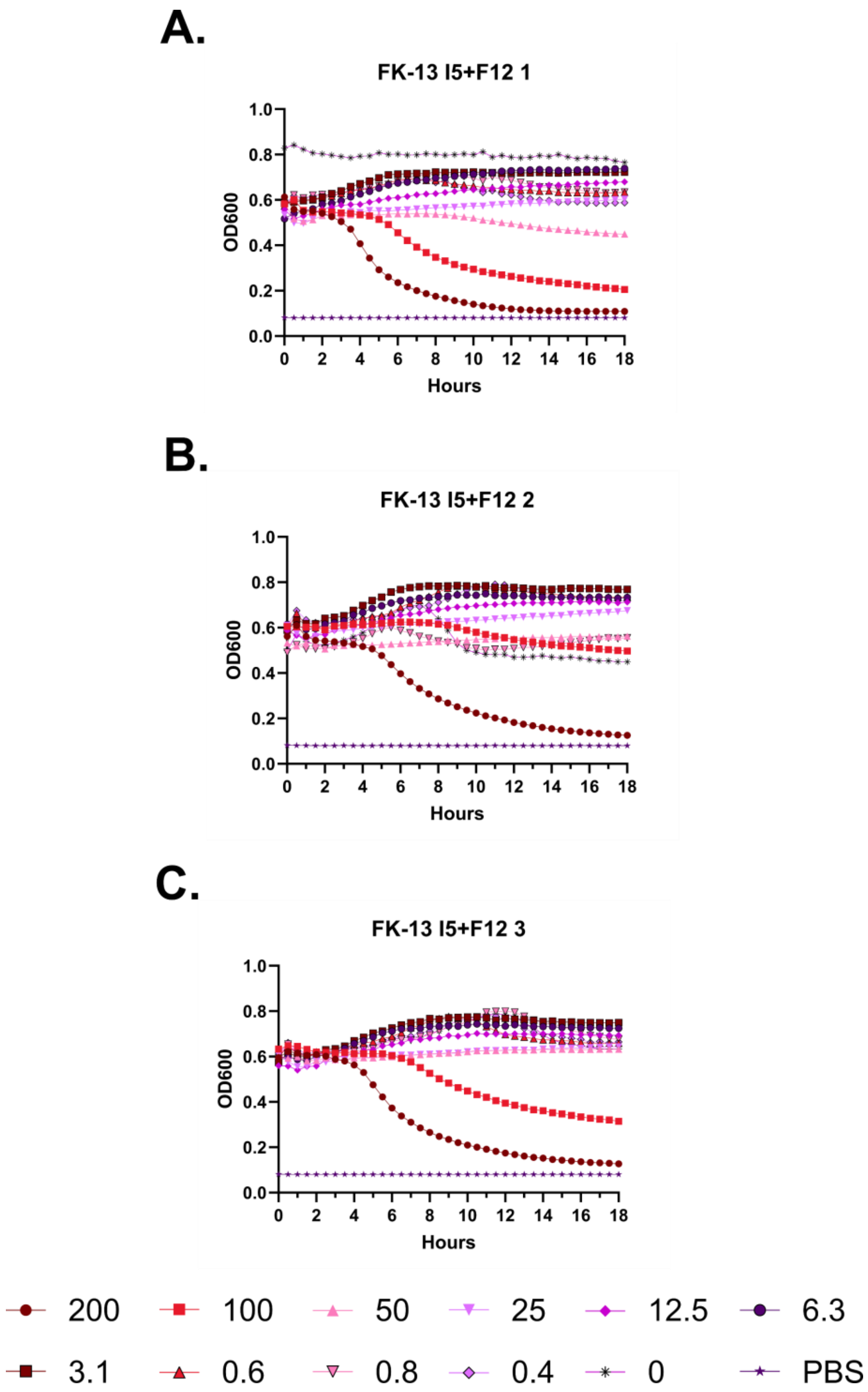
A.
FK-13 I5+F12 1
1.0
0.8
0.6
0.4
0.2
0.0
OD600
0 2 4 6 8 10 12 14 16 18
Hours
B.
FK-13 I5+F12 2
1.0
0.8
0.6
0.4
0.2
0.0
OD600
0 2 4 6 8 10 12 14 16 18
Hours
C.
FK-13 I5+F12 3
1.0
0.8
0.6
0.4
0.2
0.0
OD600
0 2 4 6 8 10 12 14 16 18
Hours
200
100
50
25
12.5
6.3
3.1
0.6
0.8
0.4
0
PBS

**Figure S75 – (#24) FK-13 I5+F12 kinetic hemolysis assay. A.** Defibrinated sheep blood was incubated in the presence of the indicated peptide concentrations at 37 °C with monitoring of OD600 over 18 hours, where decreasing OD600 indicates hemolysis and each panel indicates an independent experiment. Peptide concentrations are in µM.

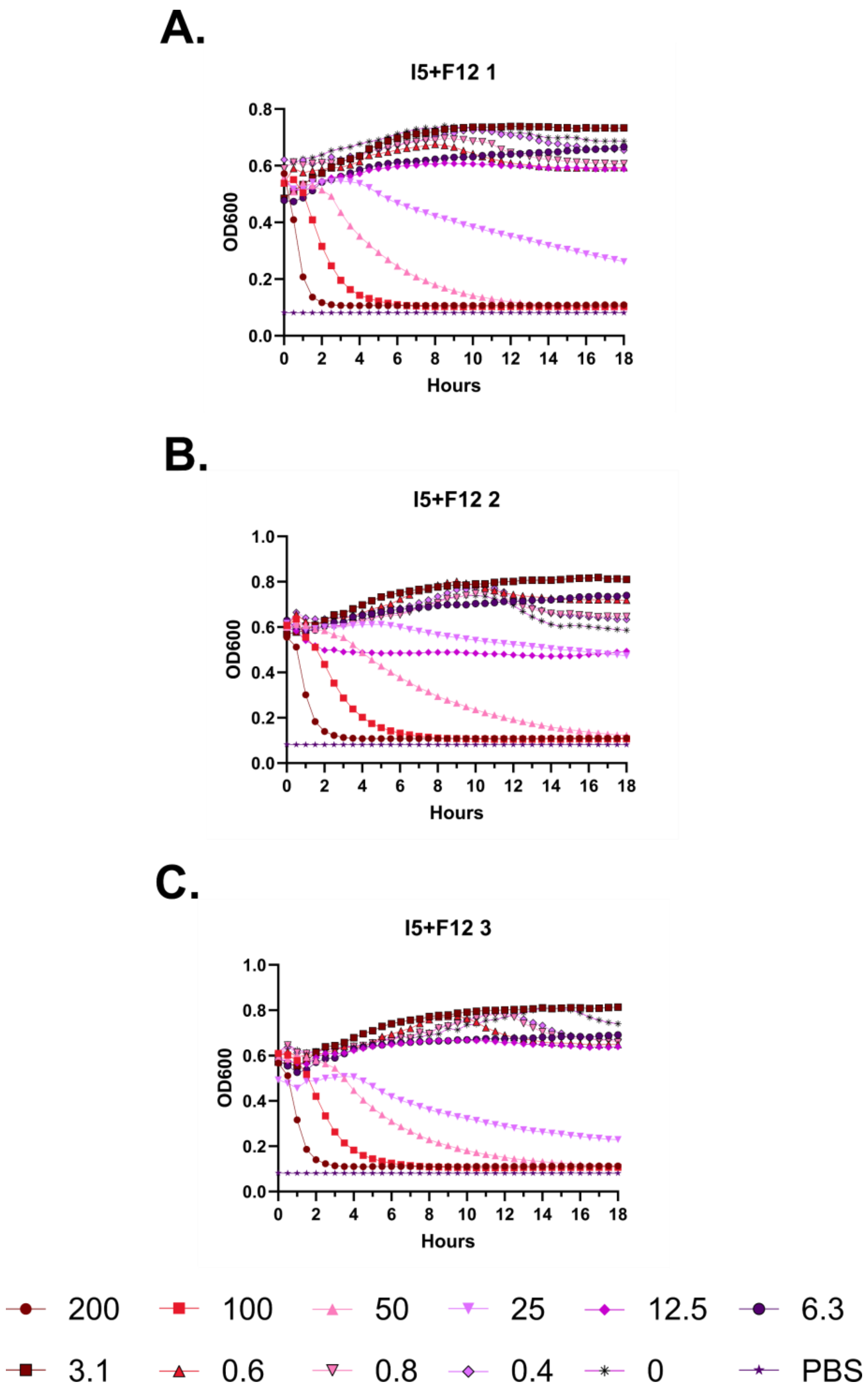
A.
I5+F12 1
OD600
Hours
0.0
0.2
0.4
0.6
0.8
0 2 4 6 8 10 12 14 16 18
B.
I5+F12 2
OD600
Hours
0.0
0.2
0.4
0.6
0.8
1.0
0 2 4 6 8 10 12 14 16 18
C.
I5+F12 3
OD600
Hours
0.0
0.2
0.4
0.6
0.8
1.0
0 2 4 6 8 10 12 14 16 18
200
100
50
25
12.5
6.3
3.1
0.6
0.8
0.4
0
PBS

**Figure S76 – (#25) FF-14 I5+F12 kinetic hemolysis assay. A.** Defibrinated sheep blood was incubated in the presence of the indicated peptide concentrations at 37 °C with monitoring of OD600 over 18 hours, where decreasing OD600 indicates hemolysis and each panel indicates an independent experiment. Peptide concentrations are in μM.

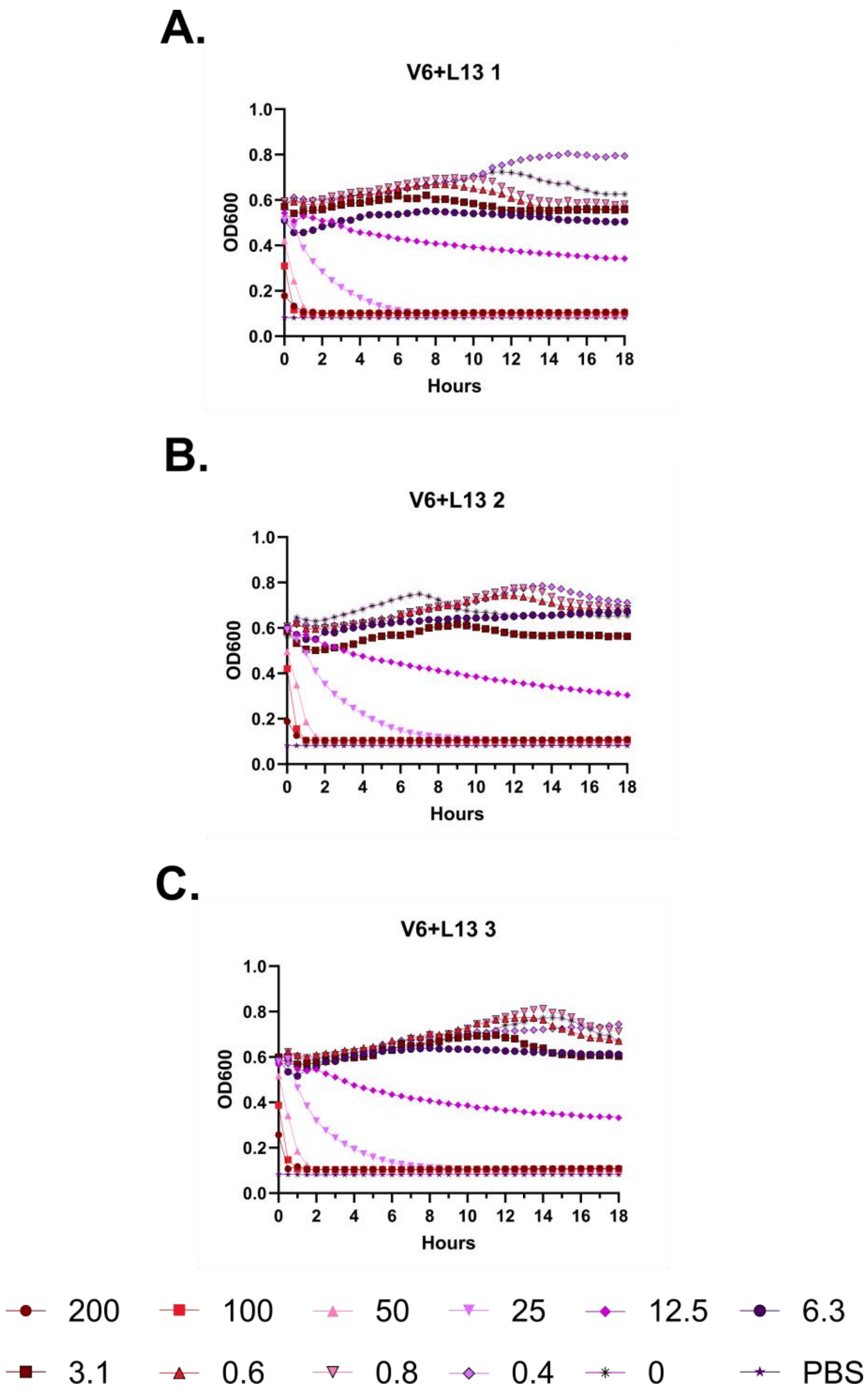
A.
V6+L13 1
B.
V6+L13 2
C.
V6+L13 3
OD600
Hours
200
100
50
25
12.5
6.3
3.1
0.6
0.8
0.4
0
PBS

**Figure S77 – (#26) FF-14 V6+L13 kinetic hemolysis assay. A.** Defibrinated sheep blood was incubated in the presence of the indicated peptide concentrations at 37 °C with monitoring of OD600 over 18 hours, where decreasing OD600 indicates hemolysis and each panel indicates an independent experiment. Peptide concentrations are in µM.

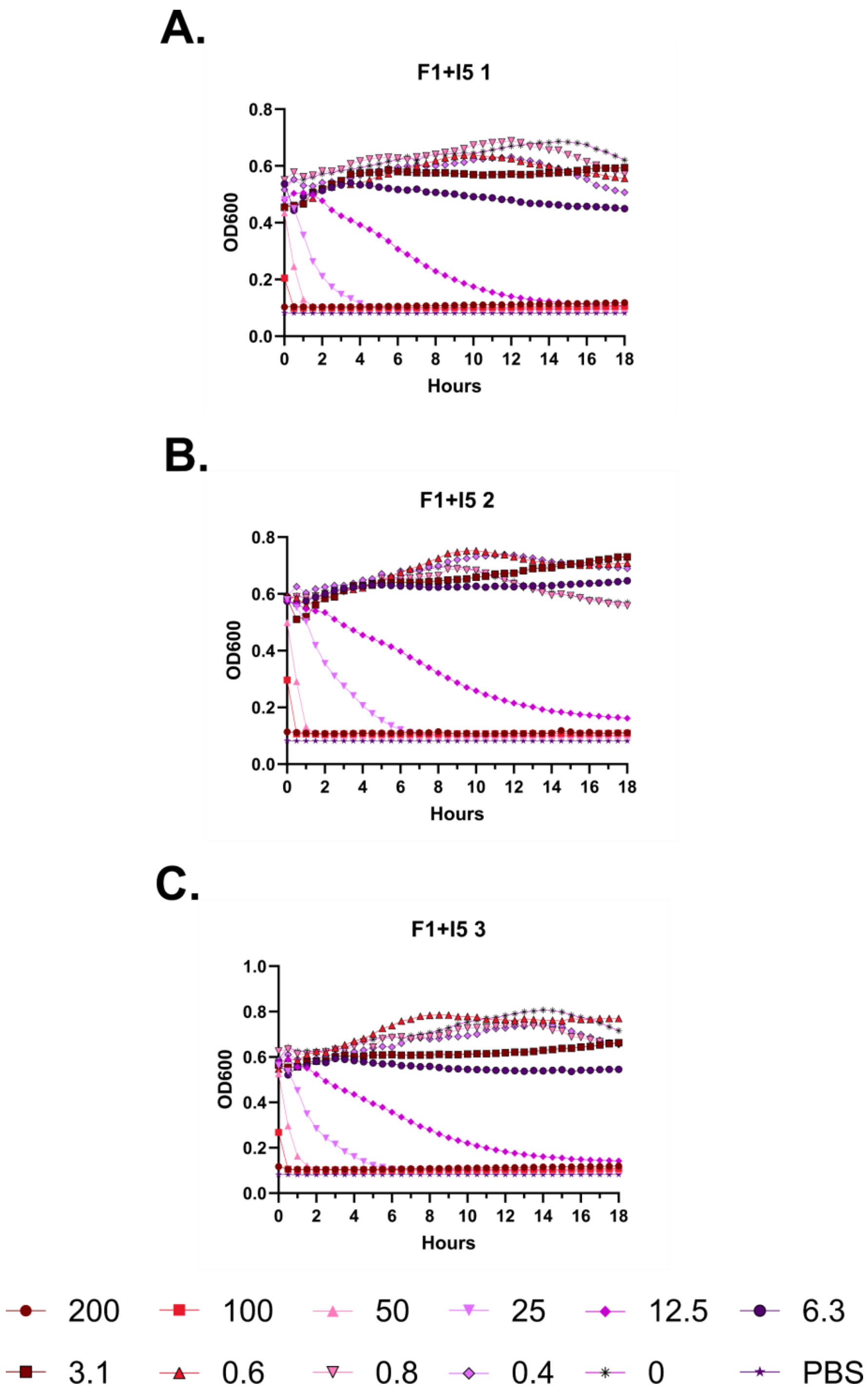
A.
F1+I5 1
OD600
Hours
B.
F1+I5 2
OD600
Hours
C.
F1+I5 3
OD600
Hours
200
100
50
25
12.5
6.3
3.1
0.6
0.8
0.4
0
PBS

**Figure S78 – (#27) FF-14 F1+I5 kinetic hemolysis assay. A.** Defibrinated sheep blood was incubated in the presence of the indicated peptide concentrations at 37 °C with monitoring of OD600 over 18 hours, where decreasing OD600 indicates hemolysis and each panel indicates an independent experiment. Peptide concentrations are in μM.

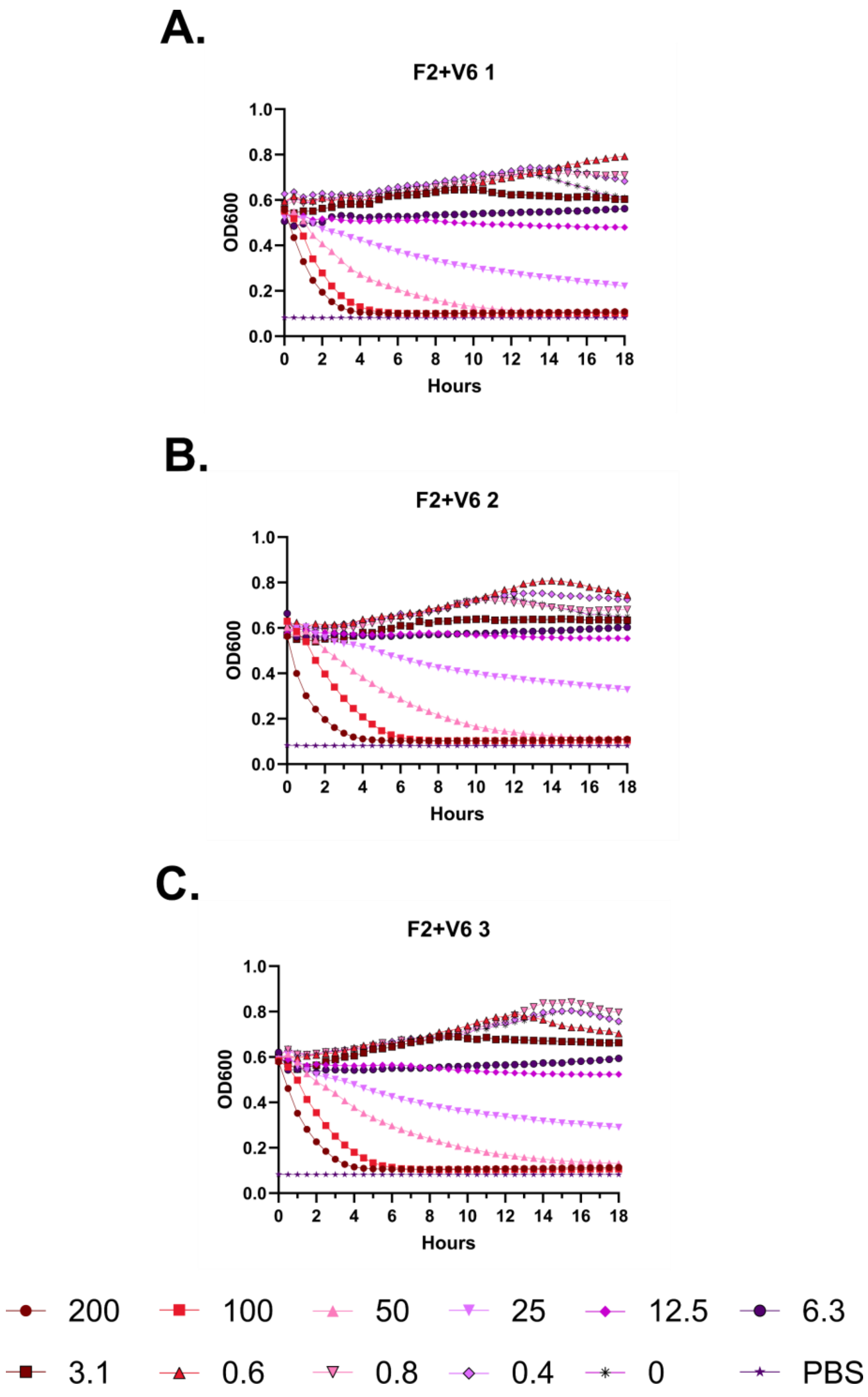
A.
F2+V6 1
OD600
Hours
B.
F2+V6 2
OD600
Hours
C.
F2+V6 3
OD600
Hours
200
100
50
25
12.5
6.3
3.1
0.6
0.8
0.4
0
PBS

**Figure S79 – (#28) FF-14 F2+V6 kinetic hemolysis assay. A.** Defibrinated sheep blood was incubated in the presence of the indicated peptide concentrations at 37 °C with monitoring of OD600 over 18 hours, where decreasing OD600 indicates hemolysis and each panel indicates an independent experiment. Peptide concentrations are in μM.

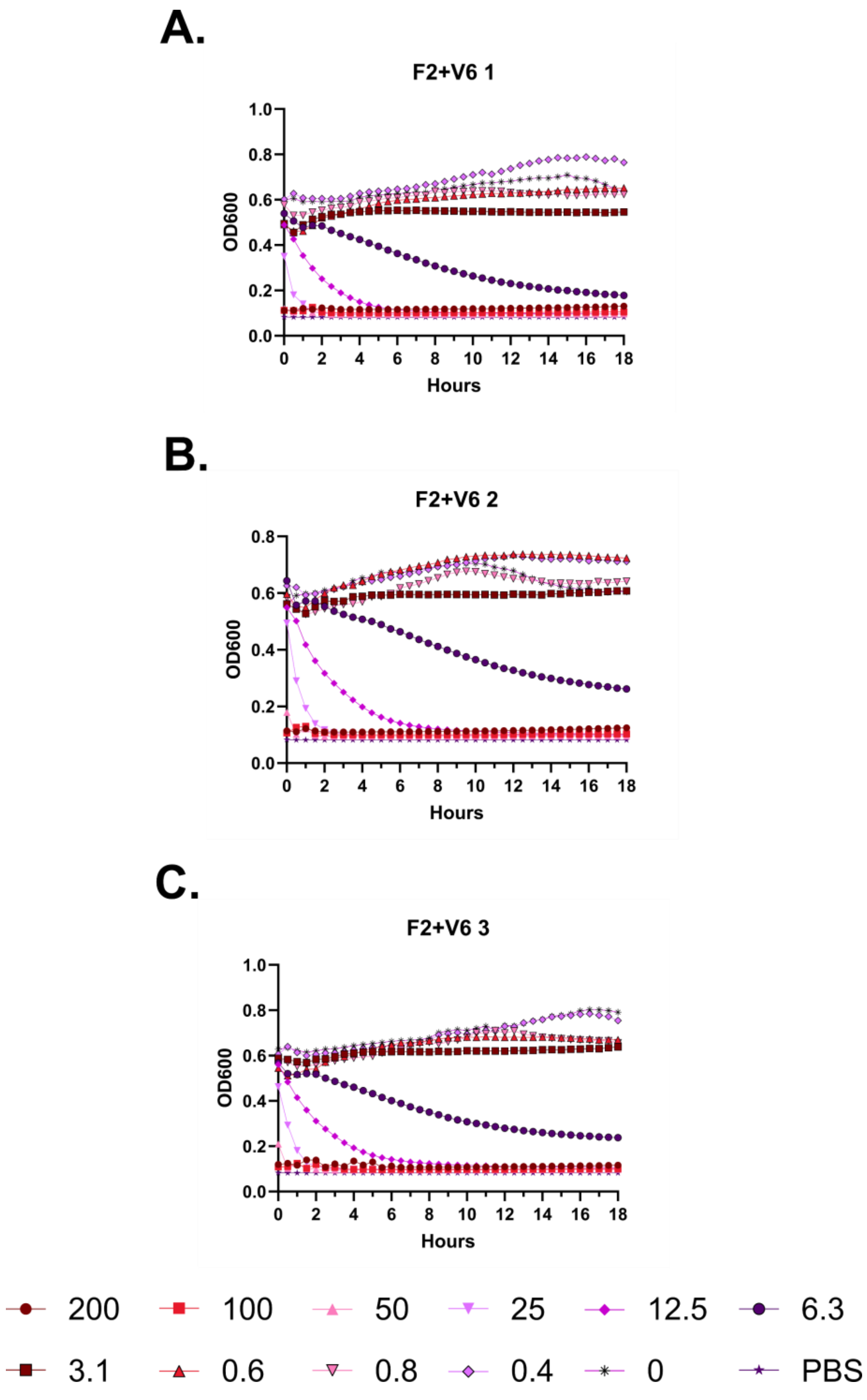
A.
F2+V6 1
OD600
Hours
B.
F2+V6 2
OD600
Hours
C.
F2+V6 3
OD600
Hours
200
100
50
25
12.5
6.3
3.1
0.6
0.8
0.4
0
PBS

**Figure S80 – (#29) FF-14 F2+V6 kinetic hemolysis assay. A.** Defibrinated sheep blood was incubated in the presence of the indicated peptide concentrations at 37 °C with monitoring of OD600 over 18 hours, where decreasing OD600 indicates hemolysis and each panel indicates an independent experiment. Peptide concentrations are in µM.

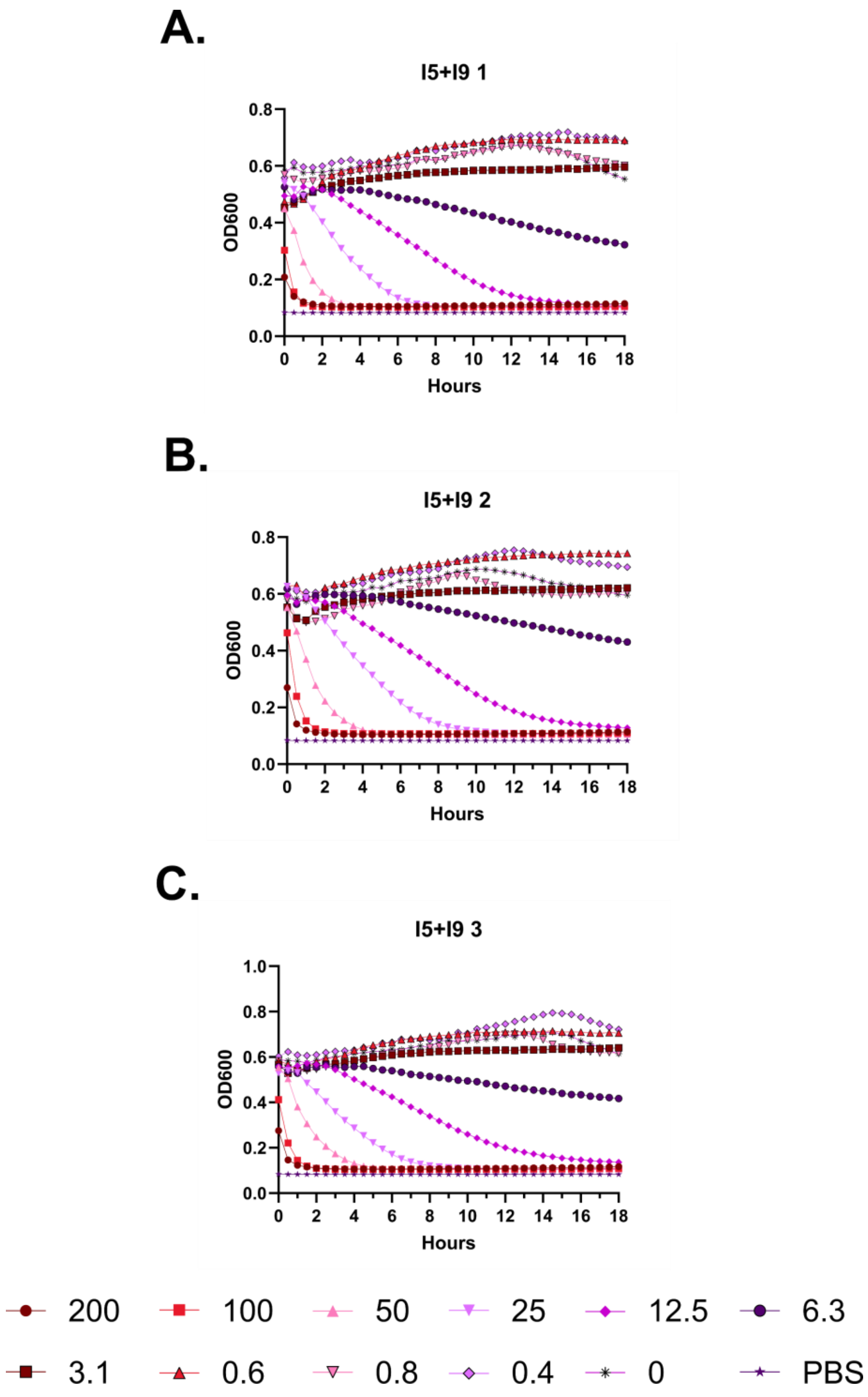
A.
I5+I9 1
0.8
0.6
0.4
0.2
0.0
OD600
0 2 4 6 8 10 12 14 16 18
Hours
B.
I5+I9 2
0.8
0.6
0.4
0.2
0.0
OD600
0 2 4 6 8 10 12 14 16 18
Hours
C.
I5+I9 3
1.0
0.8
0.6
0.4
0.2
0.0
OD600
0 2 4 6 8 10 12 14 16 18
Hours
200
100
50
25
12.5
6.3
3.1
0.6
0.8
0.4
0
PBS

**Figure S81 – (#30) FF-14 I5+I9 kinetic hemolysis assay. A.** Defibrinated sheep blood was incubated in the presence of the indicated peptide concentrations at 37 °C with monitoring of OD600 over 18 hours, where decreasing OD600 indicates hemolysis and each panel indicates an independent experiment. Peptide concentrations are in µM.

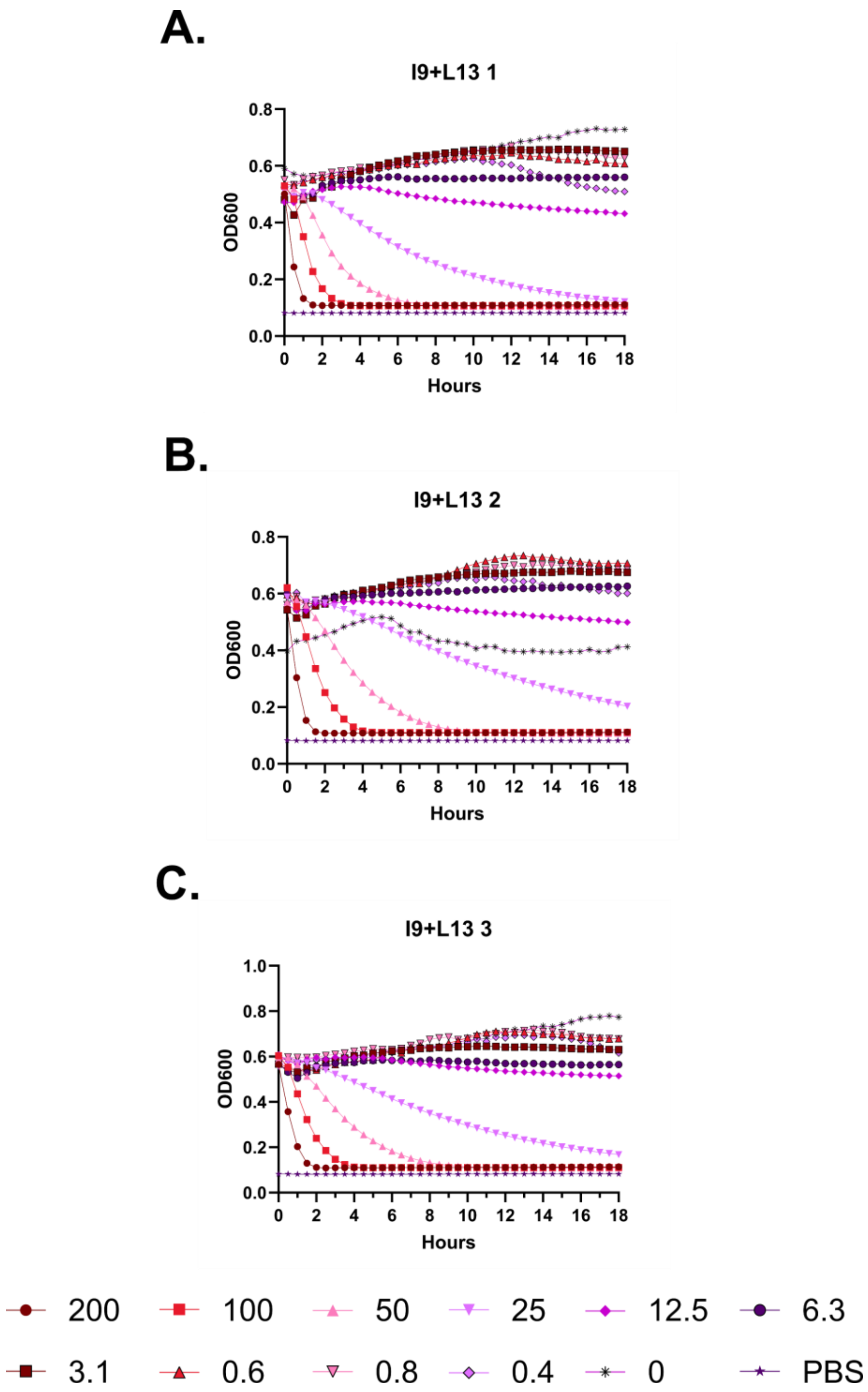
A.
I9+L13 1
OD600
Hours
B.
I9+L13 2
OD600
Hours
C.
I9+L13 3
OD600
Hours
200
100
50
25
12.5
6.3
3.1
0.6
0.8
0.4
0
PBS

**Figure S82 – (#31) FF-14 I9+L13 kinetic hemolysis assay. A.** Defibrinated sheep blood was incubated in the presence of the indicated peptide concentrations at 37 °C with monitoring of OD600 over 18 hours, where decreasing OD600 indicates hemolysis and each panel indicates an independent experiment. Peptide concentrations are in µM.

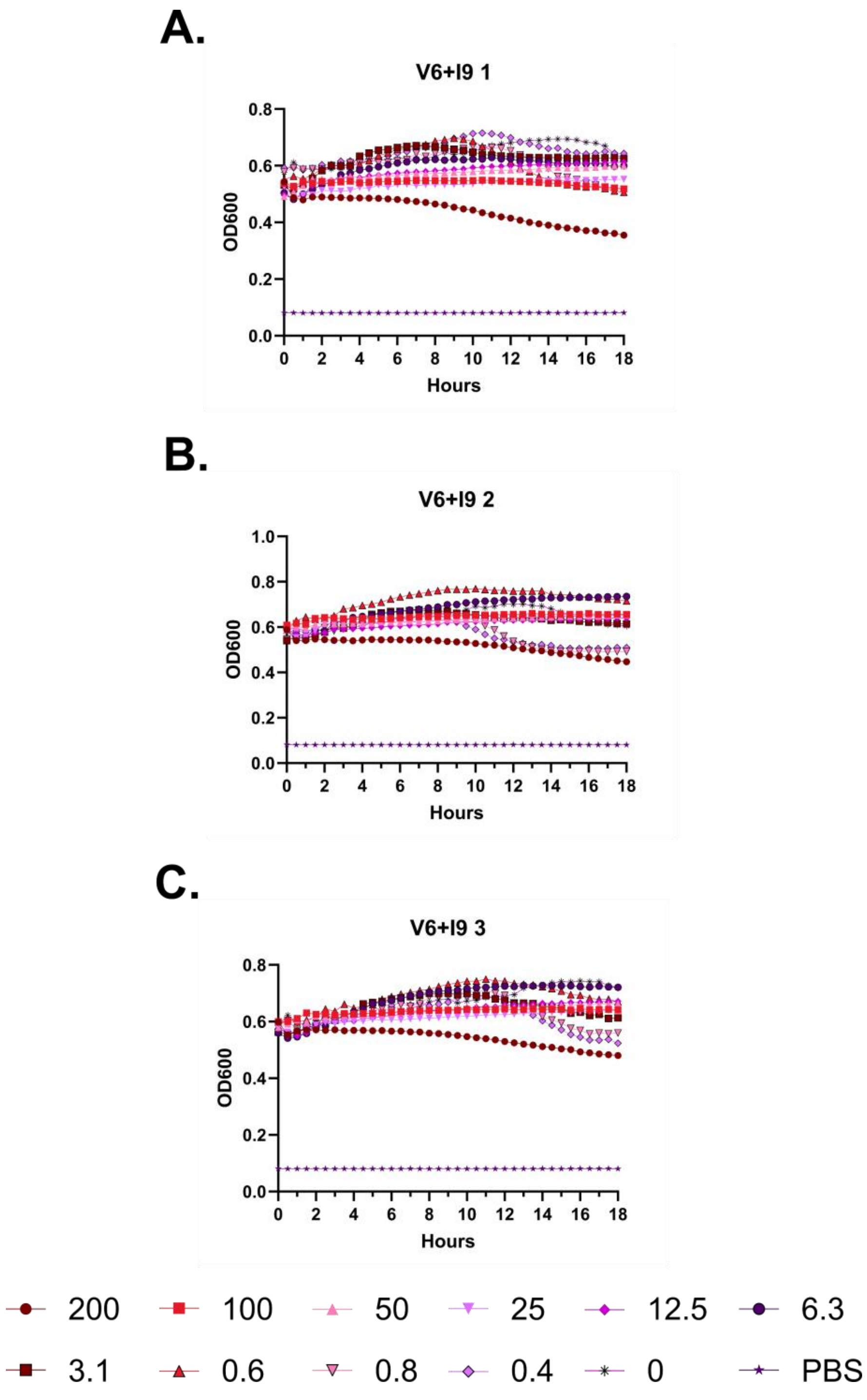
A.
V6+I9 1
OD600
Hours
B.
V6+I9 2
OD600
Hours
C.
V6+I9 3
OD600
Hours
200
100
50
25
12.5
6.3
3.1
0.6
0.8
0.4
0
PBS

**Figure S83 – (#32) FF-14 V6+I9 kinetic hemolysis assay. A.** Defibrinated sheep blood was incubated in the presence of the indicated peptide concentrations at 37 °C with monitoring of OD600 over 18 hours, where decreasing OD600 indicates hemolysis and each panel indicates an independent experiment. Peptide concentrations are in μM.

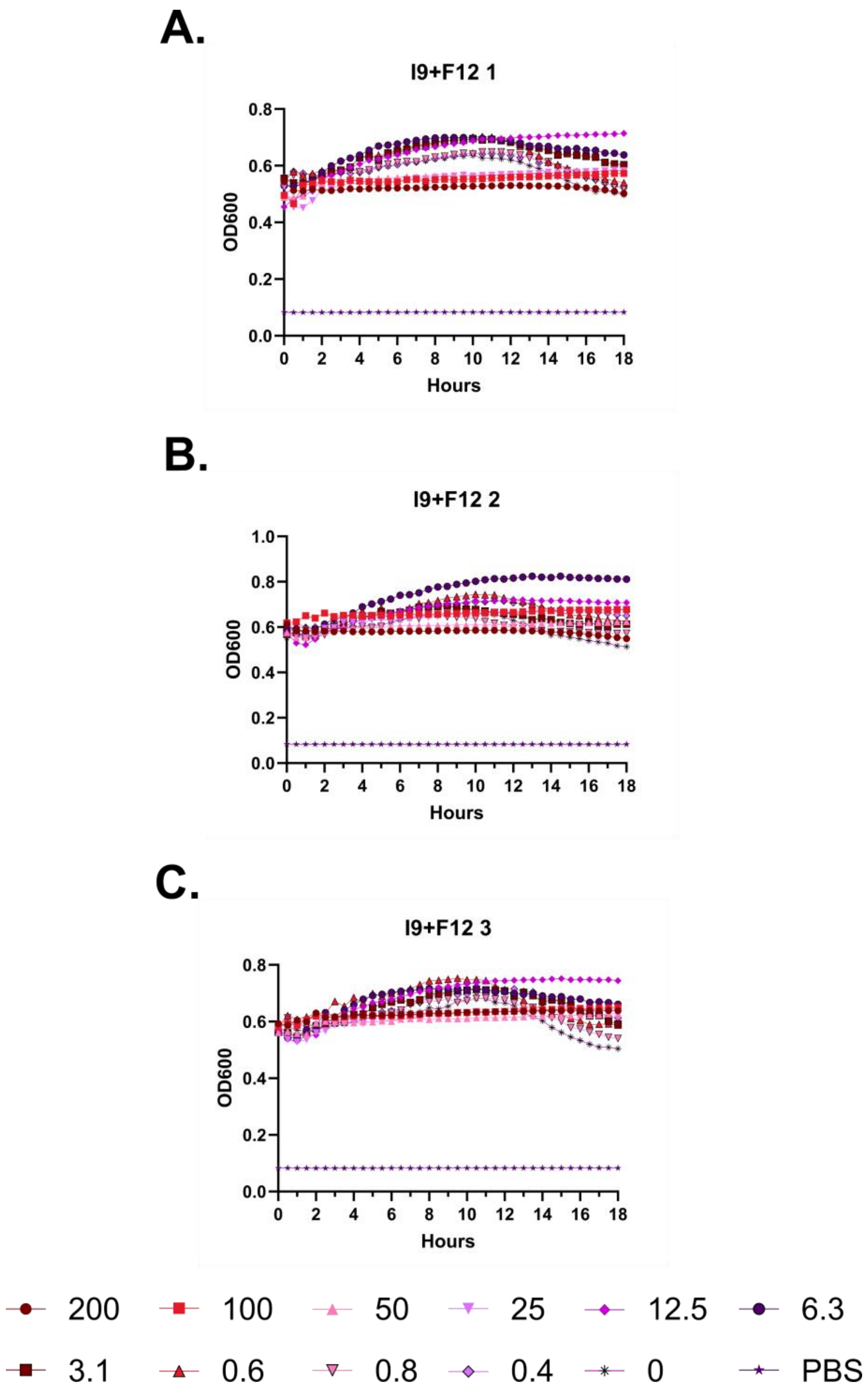
A.
I9+F12 1
OD600
Hours
B.
I9+F12 2
OD600
Hours
C.
I9+F12 3
OD600
Hours
200
100
50
25
12.5
6.3
3.1
0.6
0.8
0.4
0
PBS

**Figure S84 – (#33) FF-14 I9+F12 kinetic hemolysis assay. A.** Defibrinated sheep blood was incubated in the presence of the indicated peptide concentrations at 37 °C with monitoring of OD600 over 18 hours, where decreasing OD600 indicates hemolysis and each panel indicates an independent experiment. Peptide concentrations are in μM.

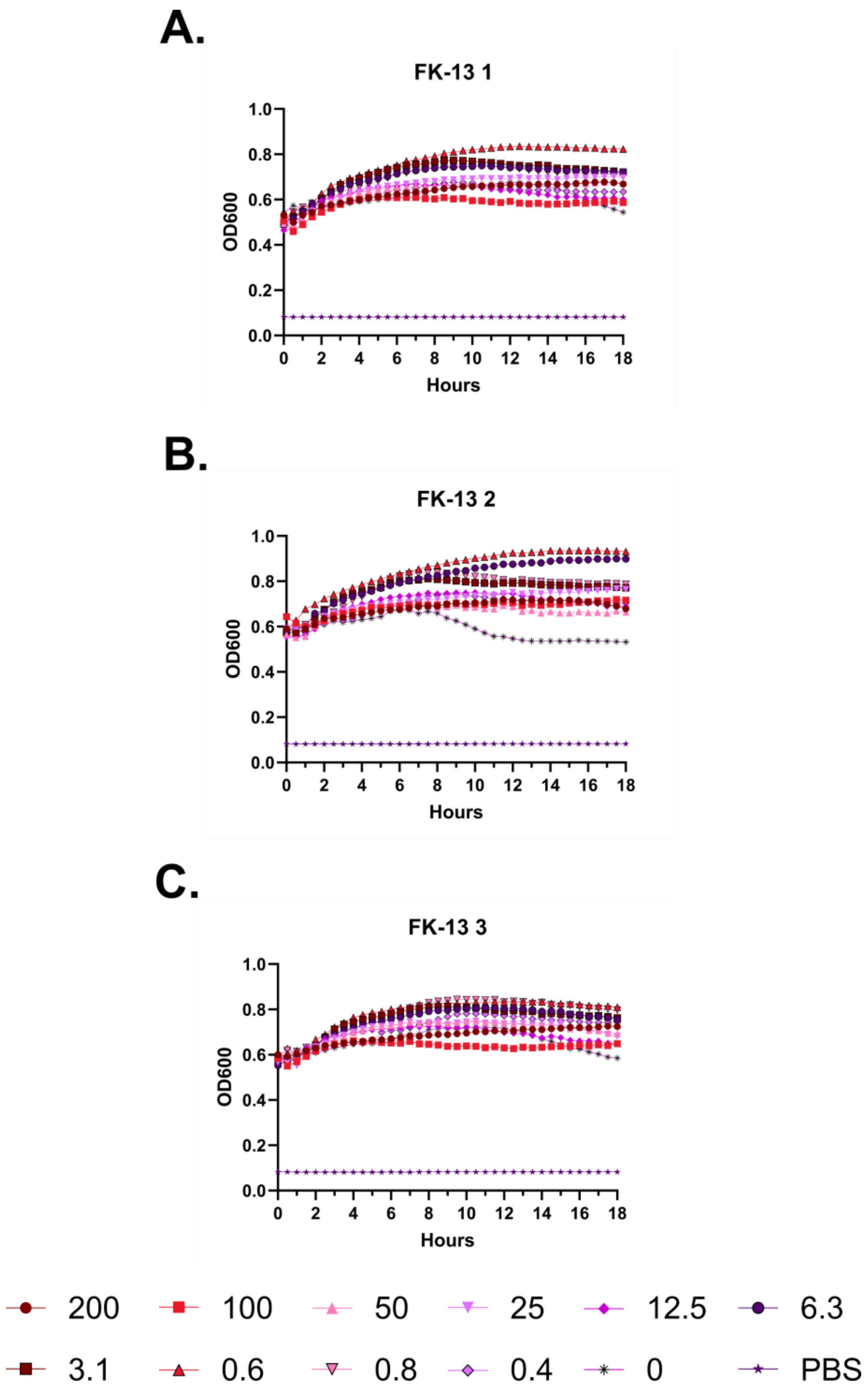
A.
FK-13 1
B.
FK-13 2
C.
FK-13 3
OD600
Hours
1.0
0.8
0.6
0.4
0.2
0.0
0
2
4
6
8
10
12
14
16
18
200
100
50
25
12.5
6.3
3.1
0.6
0.8
0.4
0
PBS

**Figure S85 – (#7) FK-13 kinetic hemolysis assay. A.** Defibrinated sheep blood was incubated in the presence of the indicated peptide concentrations at 37 °C with monitoring of OD600 over 18 hours, where decreasing OD600 indicates hemolysis and each panel indicates an independent experiment. Peptide concentrations are in μM.

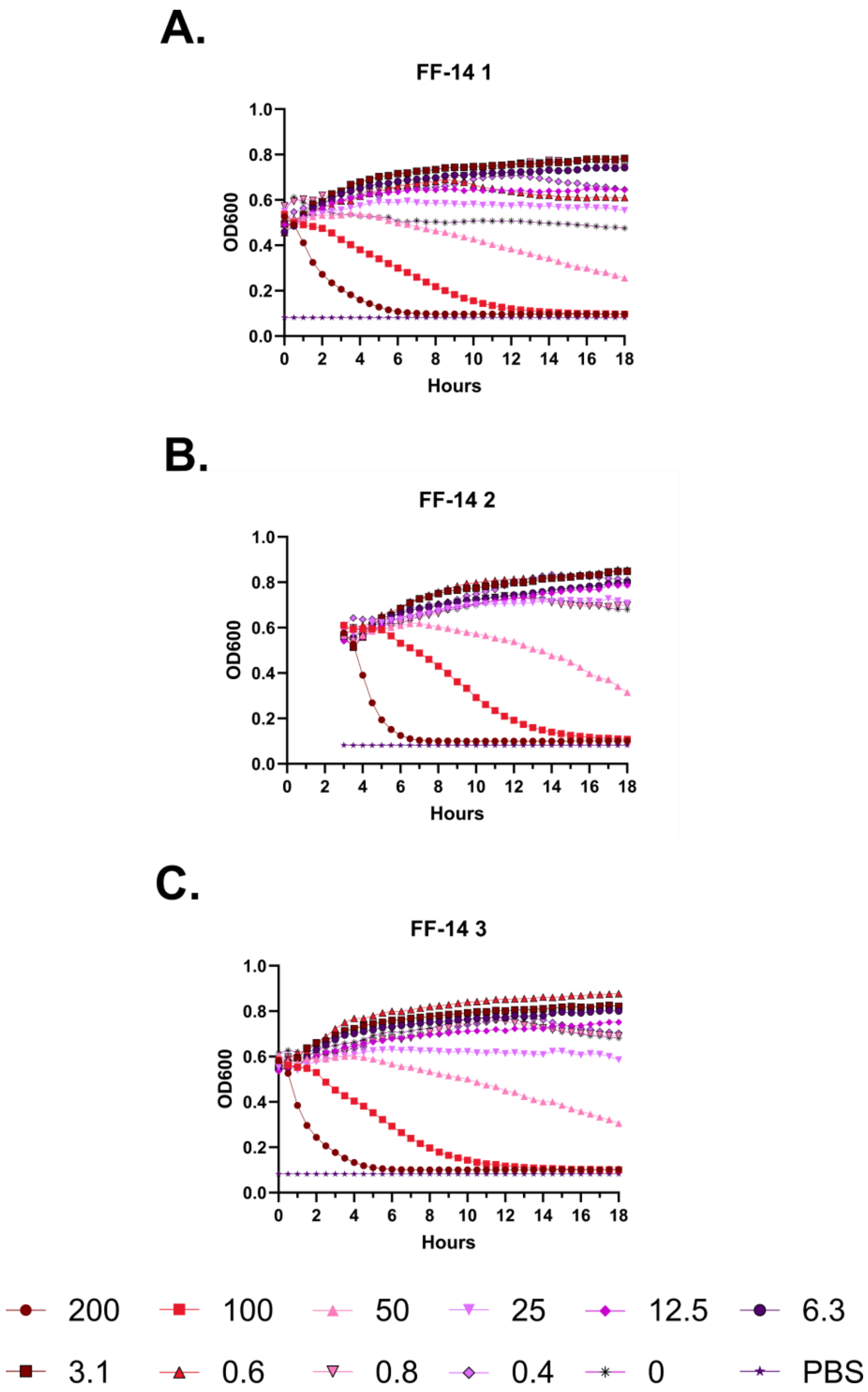
A.
FF-14 1
OD600
Hours
B.
FF-14 2
OD600
Hours
C.
FF-14 3
OD600
Hours
200
100
50
25
12.5
6.3
3.1
0.6
0.8
0.4
0
PBS

**Figure S86 – (#18) FF-14 kinetic hemolysis assay. A.** Defibrinated sheep blood was incubated in the presence of the indicated peptide concentrations at 37 °C with monitoring of OD600 over 18 hours, where decreasing OD600 indicates hemolysis and each panel indicates an independent experiment. Peptide concentrations are in µM.

A.

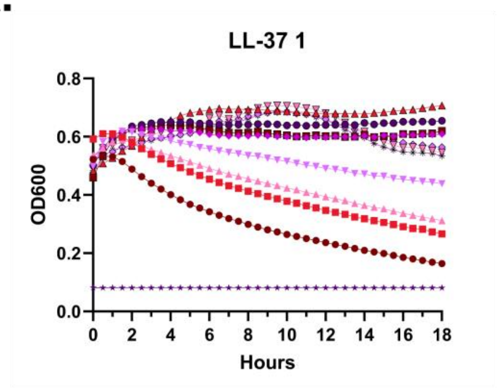
LL-37 1
0.8
0.6
0.4
0.2
0.0
OD600
0
2
4
6
8
10
12
14
16
18
Hours

B.

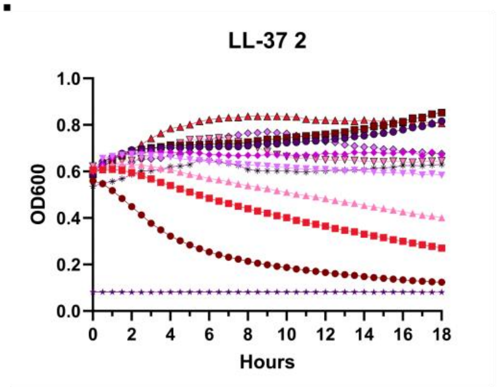
LL-37 2
1.0
0.8
0.6
0.4
0.2
0.0
OD600
0
2
4
6
8
10
12
14
16
18
Hours

C.

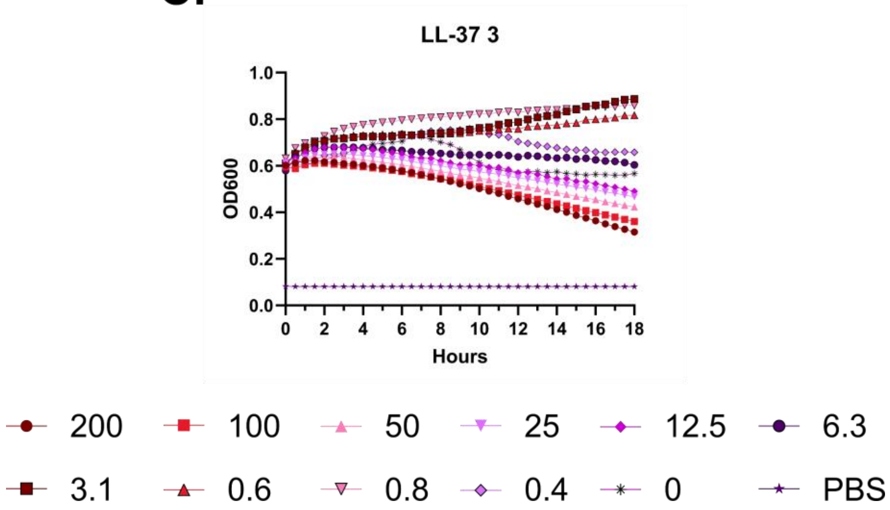
LL-37 3
1.0
0.8
0.6
0.4
0.2
0.0
OD600
0
2
4
6
8
10
12
14
16
18
Hours
200
100
50
25
12.5
6.3
3.1
0.6
0.8
0.4
0
PBS

**Figure S87 – (#1) LL-37 kinetic hemolysis assay. A.** Defibrinated sheep blood was incubated in the presence of the indicated peptide concentrations at 37 °C with monitoring of OD600 over 18 hours, where decreasing OD600 indicates hemolysis and each panel indicates an independent experiment. Peptide concentrations are in μM.

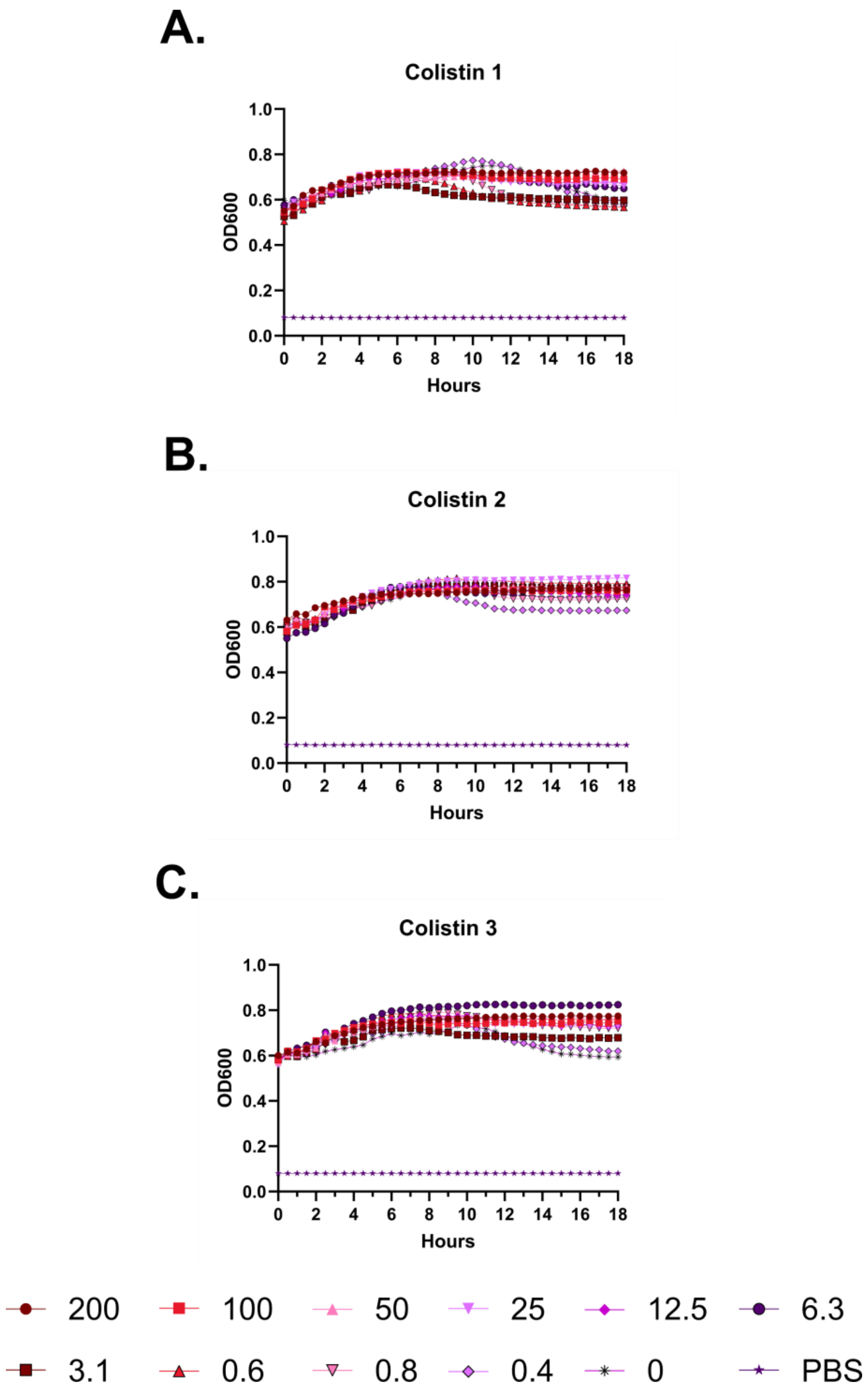
A.
Colistin 1
OD600
Hours
B.
Colistin 2
OD600
Hours
C.
Colistin 3
OD600
Hours
200
100
50
25
12.5
6.3
3.1
0.6
0.8
0.4
0
PBS

**Figure S88 – Colistin kinetic hemolysis assay. A.** Defibrinated sheep blood was incubated in the presence of the indicated peptide concentrations at 37 °C with monitoring of OD600 over 18 hours, where decreasing OD600 indicates hemolysis and each panel indicates an independent experiment. Peptide concentrations are in μM.

**A.**

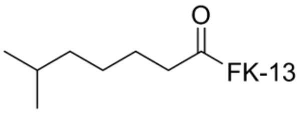


(#40) 6-methyl-heptanoic acid

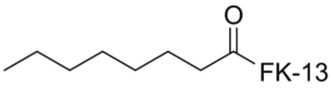


(#43) Octanoic Acid

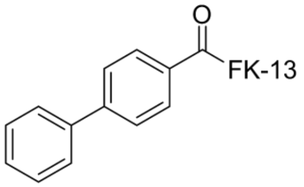


(#41) Biphenyl-4-carboxylic Acid

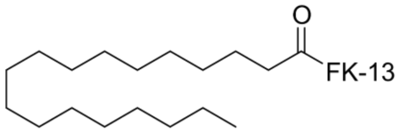


(#44) Stearic Acid

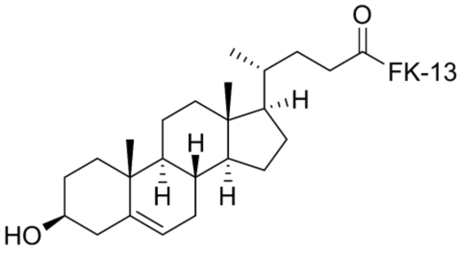


(#42) Cholenic Acid

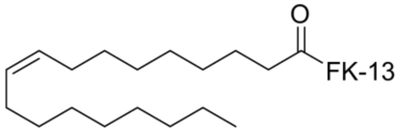


(#45) Oleic Acid

**B.**

| | MIC Ec | MIC Pa | < 20% Hemo | Hemo/MIC | Fold Δ |
|---|---|---|---|---|---|
| **(#40) 6-Me-Hepta** | 3.1 | 6.3 | 0.8 | 0.25 | 1 |
| **(#41) Biphenyl** | 3.1 | 6.3 | 1.6 | 0.5 | 2 |
| **(#42) Cholenic** | 25 | >50 | 0.4 | 0.016 | 0.063 |
| **(#43) Octanoic** | 1.6 | 3.1 | 0.8 | 0.5 | 2 |
| **(#44) Stearic** | >50 | >50 | 0.2 | 0.002 | 0.008 |
| **(#45) Oleic** | 12.5 | 50 | 0.2 | 0.016 | 0.063 |
| **(#10) KR-12** | >50 | >50 | >200 | 2 | 8 |
| **(#18) FF-14** | 3.1 | 12.5 | 6.3 | 2 | 8 |
| **(#1) LL-37** | 3.1 | 3.1 | 0.8 | 0.25 | 1 |
| **Colistin** | 0.4 | 0.4 | 100 | 256 | 1024 |

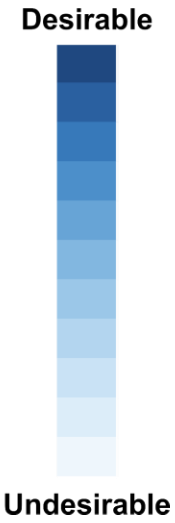

**Figure S89 – Modulation of (#7) FK-13 activities via N-lipidation. A.** Structures are shown of the lipid tails appended to the N-terminus of (#7) FK-13. 6-Me-Hepta derives from colistin, while the biphenyl moiety has been shown to be important in some preclinical antibiotics. Cholenic acid is taken as a sterol representative. Octanoic, stearic, and oleic acids are taken as short, long, and mono-unsaturated tails. **B.** Compound properties including the MICs of each compound against *E. coli* (ATCC 25922) (**MIC Ec**) and *P. aeruginosa* (ATCC 27853) (**MIC Pa**) in the left two columns and the highest concentration of peptide in which hemolysis remains below a 20% threshold (**< 20% Hemo**). Calculated values include **Hemo/MIC** = (**< 20% Hemo**) / **MIC Ec** and **Fold Δ** = (**Hemo/MIC** of compound) / (**Hemo/MIC** of LL-37), where larger numbers are better. Rationales for analysis strategies are discussed further under **Methods**. Shading of the values within each column follows a convention by which less desirable values are assigned lighter shades while more desirable values are assigned darker shades.

**A.**

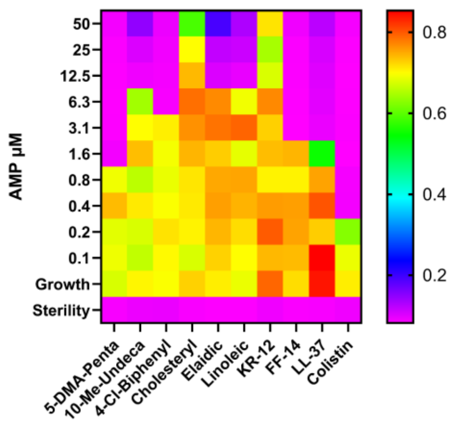

AMP µM
50
25
12.5
6.3
3.1
1.6
0.8
0.4
0.2
0.1
Growth
Sterility
5-DMA-Penta
10-Me-Undeca
4-Cl-Biphenyl
Cholesteryl
Elaidic
Linoleic
KR-12
FF-14
LL-37
Colistin
0.8
0.6
0.4
0.2


**B.**

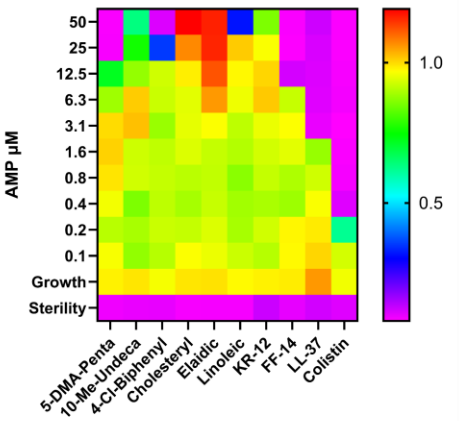

AMP µM
50
25
12.5
6.3
3.1
1.6
0.8
0.4
0.2
0.1
Growth
Sterility
5-DMA-Penta
10-Me-Undeca
4-Cl-Biphenyl
Cholesteryl
Elaidic
Linoleic
KR-12
FF-14
LL-37
Colistin
1.0
0.5


**C.**

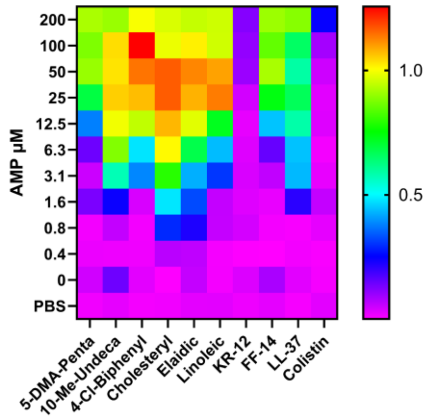

AMP µM
200
100
50
25
12.5
6.3
3.1
1.6
0.8
0.4
0
PBS
5-DMA-Penta
10-Me-Undeca
4-Cl-Biphenyl
Cholesteryl
Elaidic
Linoleic
KR-12
FF-14
LL-37
Colistin
1.0
0.5

**Figure S90 – Effects of N-terminal lipidation moieties on FF-14 activity. A.** Antimicrobial susceptibility testing showing the activity of each derivative as well as a colistin control against *E. coli* (ATCC 25922). In the heatmap plate depiction, cooler colors indicate low or absent bacterial growth as detected by the OD600 measure at 18 hours of incubation at 37 °C. **B.** As in **A.**, but with *P. aeruginosa* (ATCC 27853). **C.** Hemolysis assays in which sheep RBCs in PBS were incubated with the indicated peptide for 18 hours at 37 °C with subsequent quantification of hemoglobin released into the supernatant at 414 nm. Cool colors indicate little or no release of hemoglobin, where all levels are scaled to 100% hemolysis freeze-thaw controls. Levels throughout are the median across three independent experiments.

A.

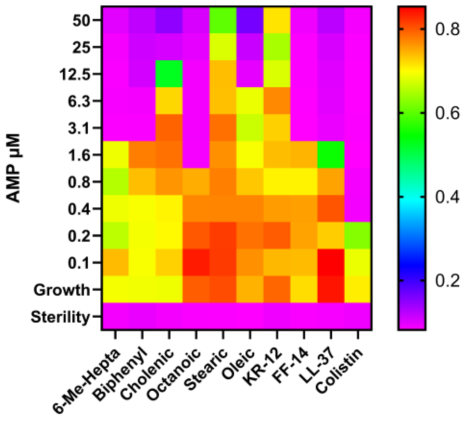
AMP µM
50
25
12.5
6.3
3.1
1.6
0.8
0.4
0.2
0.1
Growth
Sterility
6-Me-Hepta
Biphenyl
Cholenic
Octanoic
Stearic
Oleic
KR-12
FF-14
LL-37
Colistin
0.8
0.6
0.4
0.2

B.

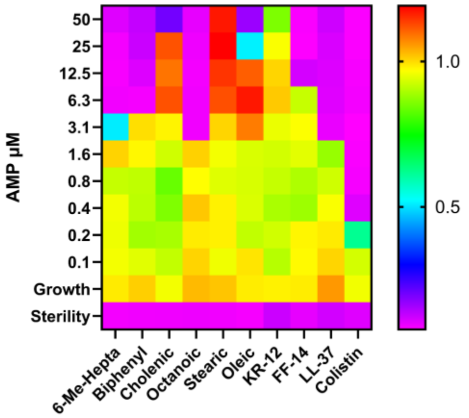
AMP µM
50
25
12.5
6.3
3.1
1.6
0.8
0.4
0.2
0.1
Growth
Sterility
6-Me-Hepta
Biphenyl
Cholenic
Octanoic
Stearic
Oleic
KR-12
FF-14
LL-37
Colistin
1.0
0.5

C.

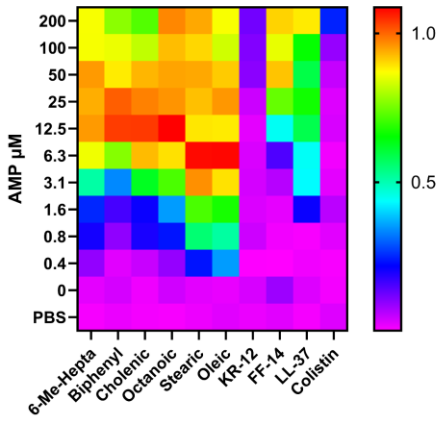
AMP µM
200
100
50
25
12.5
6.3
3.1
1.6
0.8
0.4
0
PBS
6-Me-Hepta
Biphenyl
Cholenic
Octanoic
Stearic
Oleic
KR-12
FF-14
LL-37
Colistin
1.0
0.5

**Figure S91 – Effects of N-terminal lipidation moieties on FK-13 activity. B.** Antimicrobial susceptibility testing showing the activity of each derivative as well as a colistin control against *E. coli* (ATCC 25922). In the heatmap plate depiction, cooler colors indicate low or absent bacterial growth as detected by the OD600 measure at 18 hours of incubation at 37 °C. **B.** As in **A.**, but with *P. aeruginosa* (ATCC 27853). **C.** Hemolysis assays in which sheep RBCs in PBS were incubated with the indicated peptide for 18 hours at 37 °C with subsequent quantification of hemoglobin released into the supernatant at 414 nm. Cool colors indicate little or no release of hemoglobin, where all levels are scaled to 100% hemolysis freeze-thaw controls. Levels throughout are the median across three independent experiments. Data shown in this figure were part of the same experiments shown in **Figures 5**, **S90**, and **S92**, and thus the data for control peptides (#10) KR-12, (#18) FF-14, (#1) LL-37, and colistin are identical to those shown in **Figures 5**, **S90**, and **S92**. (#7) FK-13 derivatives are shown separately as supplementary data due to lower purities following single-pass purification among these peptides; no additional purification or experimentation was undertaken due to lack of apparent phenotypically desirable effects in the first-pass data.

A.

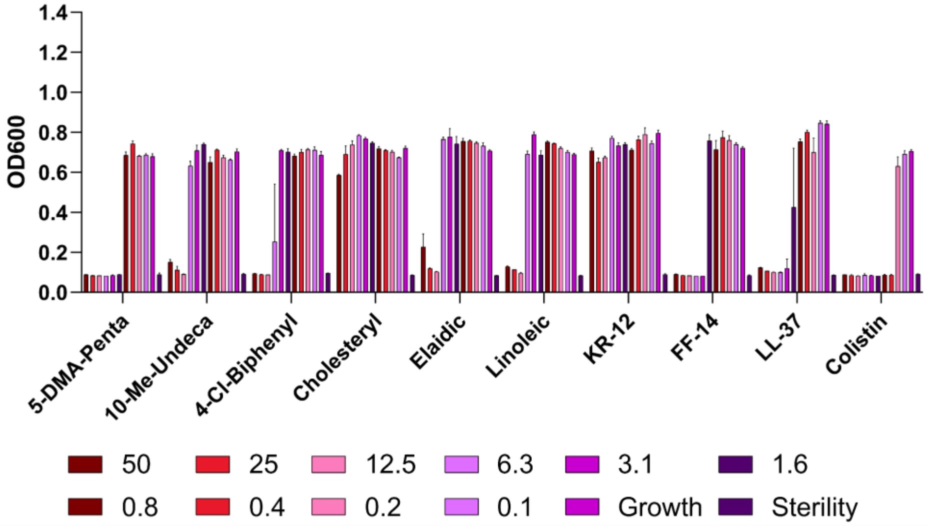
OD600
0.0
0.2
0.4
0.6
0.8
1.0
1.2
1.4
5-DMA-Penta
10-Me-Undeca
4-Cl-Biphenyl
Cholesteryl
Elaidic
Linoleic
KR-12
FF-14
LL-37
Colistin
50
25
12.5
6.3
3.1
1.6
0.8
0.4
0.2
0.1
Growth
Sterility

B.

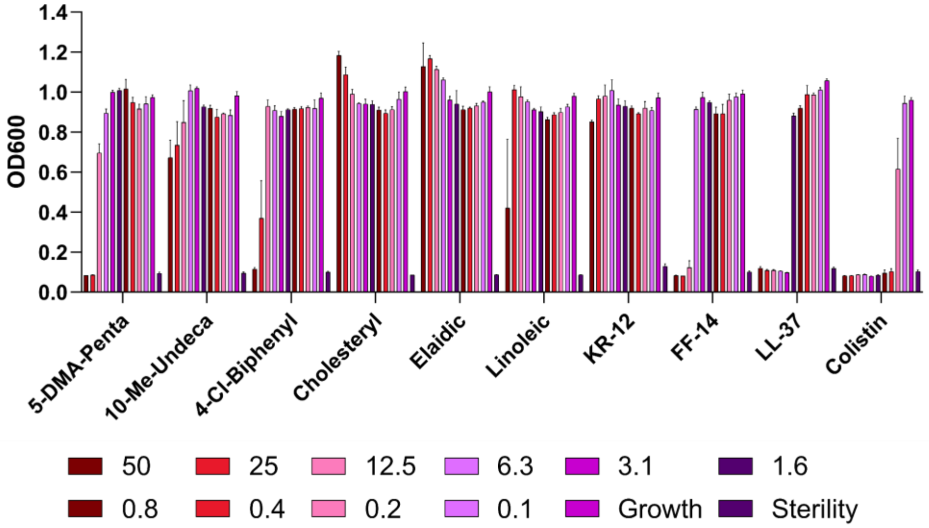
OD600
0.0
0.2
0.4
0.6
0.8
1.0
1.2
1.4
5-DMA-Penta
10-Me-Undeca
4-Cl-Biphenyl
Cholesteryl
Elaidic
Linoleic
KR-12
FF-14
LL-37
Colistin
50
25
12.5
6.3
3.1
1.6
0.8
0.4
0.2
0.1
Growth
Sterility

C.

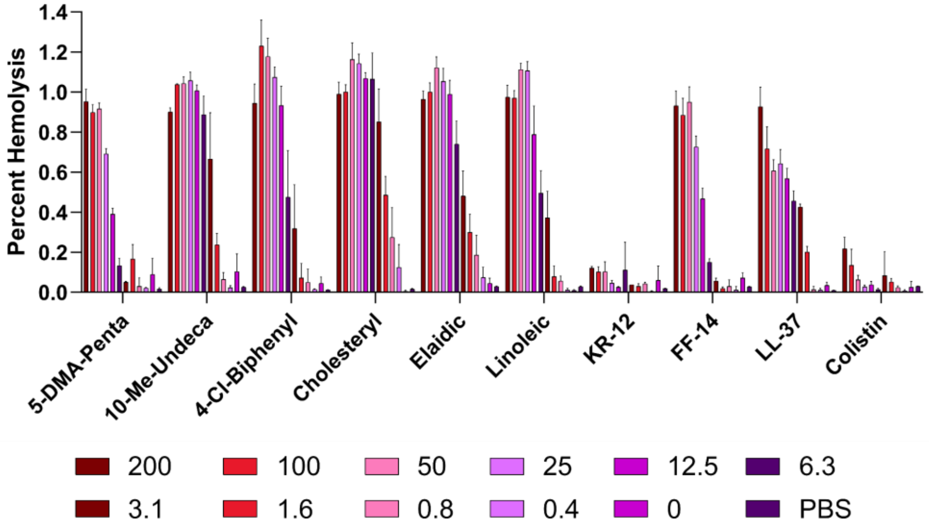
Percent Hemolysis
0.0
0.2
0.4
0.6
0.8
1.0
1.2
1.4
5-DMA-Penta
10-Me-Undeca
4-Cl-Biphenyl
Cholesteryl
Elaidic
Linoleic
KR-12
FF-14
LL-37
Colistin
200
100
50
25
12.5
6.3
3.1
1.6
0.8
0.4
0
PBS

**Figure S92 – Histograms corresponding to Figure 5**. **A.** Growth of *E. coli* (ATCC 25922) by OD600 at 18 hours at 37 °C. Error bars indicate the standard deviation of three independent experiments. **B.** As in **A.**, but with *P. aeruginosa* (ATCC 27853). **C.** Endpoint hemolysis assays following incubation with the indicated peptide for 18 hours at 37 °C. Error bars indicate the standard deviation of three independent experiments. All peptide levels are given in μM.

A.

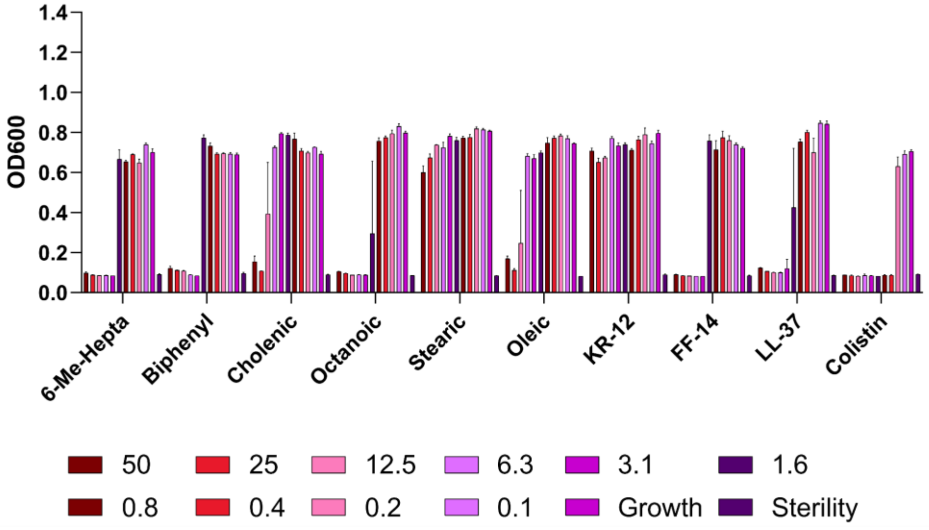
OD600
1.4
1.2
1.0
0.8
0.6
0.4
0.2
0.0
6-Me-Hepta
Biphenyl
Cholenic
Octanoic
Stearic
Oleic
KR-12
FF-14
LL-37
Colistin
50
25
12.5
6.3
3.1
1.6
0.8
0.4
0.2
0.1
Growth
Sterility

B.

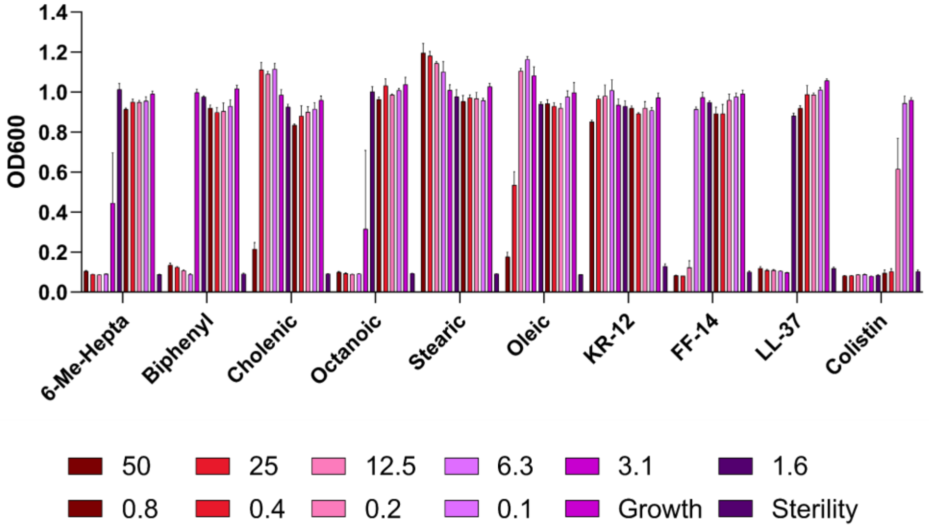
OD600
1.4
1.2
1.0
0.8
0.6
0.4
0.2
0.0
6-Me-Hepta
Biphenyl
Cholenic
Octanoic
Stearic
Oleic
KR-12
FF-14
LL-37
Colistin
50
25
12.5
6.3
3.1
1.6
0.8
0.4
0.2
0.1
Growth
Sterility

C.

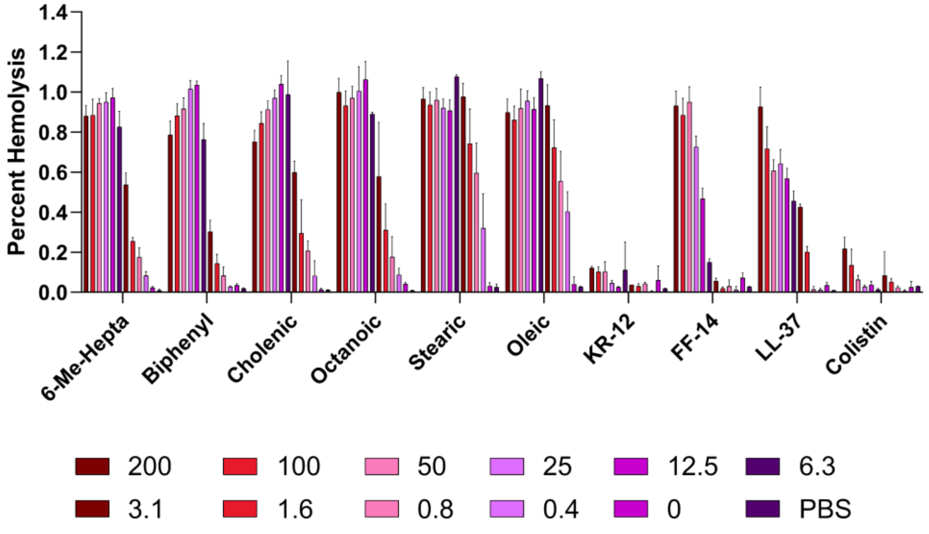
Percent Hemolysis
1.4
1.2
1.0
0.8
0.6
0.4
0.2
0.0
6-Me-Hepta
Biphenyl
Cholenic
Octanoic
Stearic
Oleic
KR-12
FF-14
LL-37
Colistin
200
100
50
25
12.5
6.3
3.1
1.6
0.8
0.4
0
PBS

**Figure S93 – Histograms corresponding to Figure S89**. **A.** Growth of *E. coli* (ATCC 25922) by OD600 at 18 hours at 37 °C. Error bars indicate the standard deviation of three independent experiments. **B.** As in **A.**, but with *P. aeruginosa* (ATCC 27853). **C.** Endpoint hemolysis assays following incubation with the indicated peptide for 18 hours at 37 °C. Error bars indicate the standard deviation of three independent experiments. All peptide levels are given in μM. Data shown in this figure were part of the same experiments shown in **Figures 5**, **S90**, and **S92**, and thus the data for control peptides (#10) KR-12, (#18) FF-14, (#1) LL-37, and colistin are identical to those shown in **Figures 5**, **S90**, and **S92**.

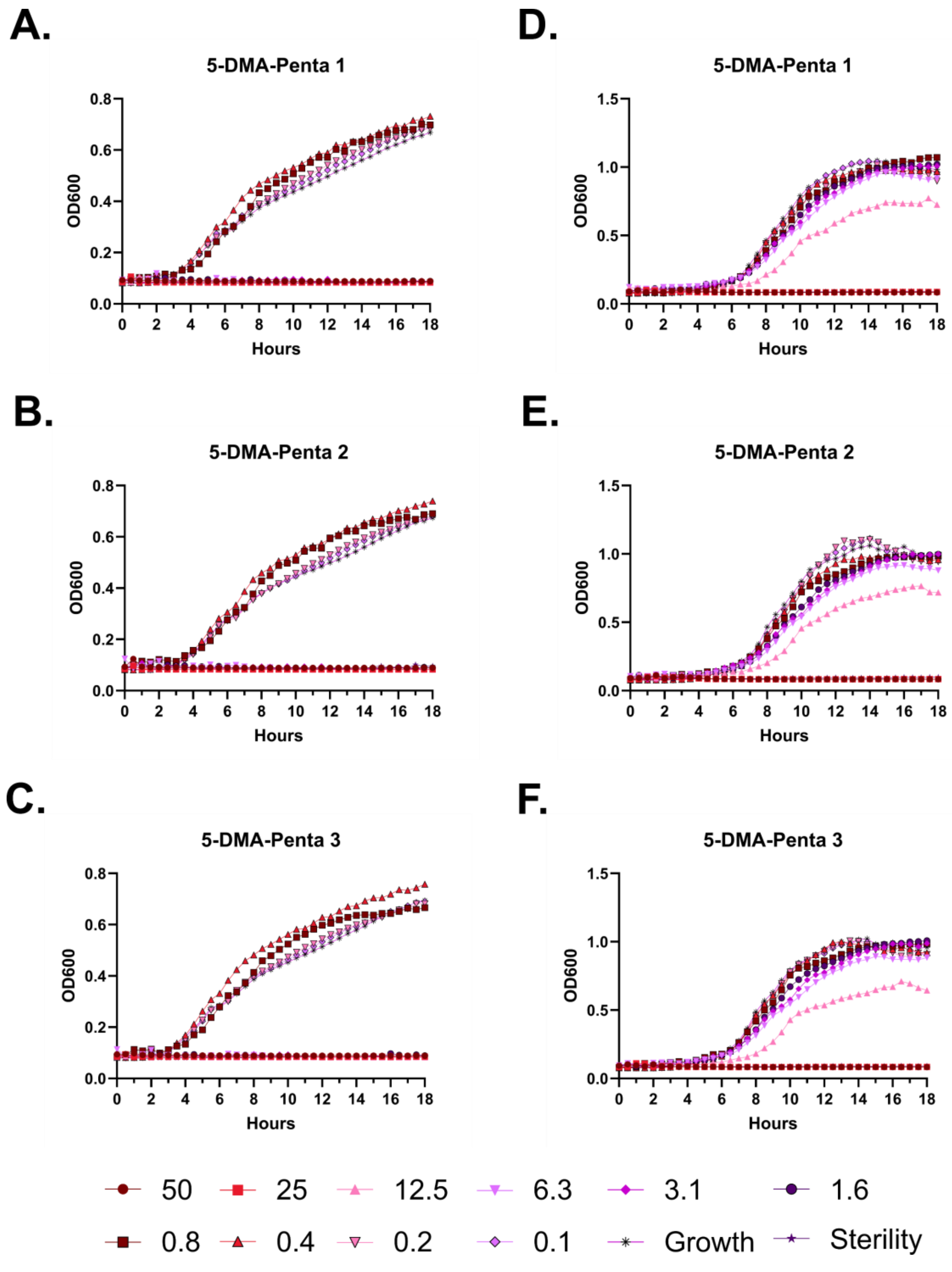

A.
5-DMA-Penta 1
B.
5-DMA-Penta 2
C.
5-DMA-Penta 3
D.
5-DMA-Penta 1
E.
5-DMA-Penta 2
F.
5-DMA-Penta 3
OD600
Hours
50
25
12.5
6.3
3.1
1.6
0.8
0.4
0.2
0.1
Growth
Sterility

**Figure S94 – Antimicrobial susceptibility curves for (#34) FF-14 N-lipidated with 5-dimethylamino pentanoic acid. A-C.** Independent experiments with *E. coli.* **D-F.** Independent experiments with *P. aeruginosa.* Growth over 18 hours at 37 °C with a gradient of peptide concentrations given in µM.

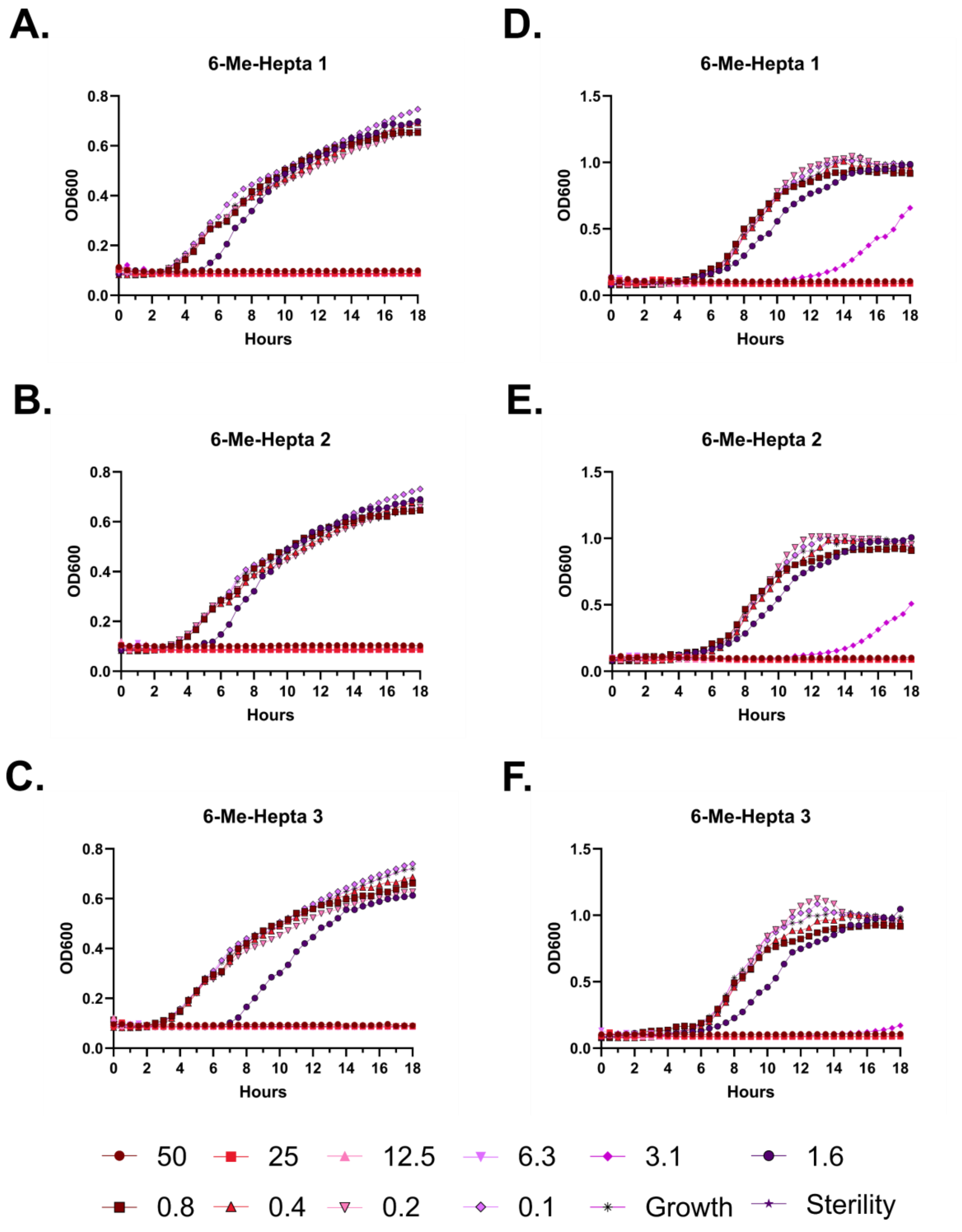
A.
6-Me-Hepta 1
B.
6-Me-Hepta 2
C.
6-Me-Hepta 3
D.
6-Me-Hepta 1
E.
6-Me-Hepta 2
F.
6-Me-Hepta 3
OD600
Hours
50
25
12.5
6.3
3.1
1.6
0.8
0.4
0.2
0.1
Growth
Sterility

**Figure S95 – Antimicrobial susceptibility curves for (#40) FK-13 N-lipidated with 6-methyl hepatnoic acid. A-C.** Independent experiments with *E. coli*. **D-F.** Independent experiments with *P. aeruginosa.* Growth over 18 hours at 37 °C with a gradient of peptide concentrations given in μM.

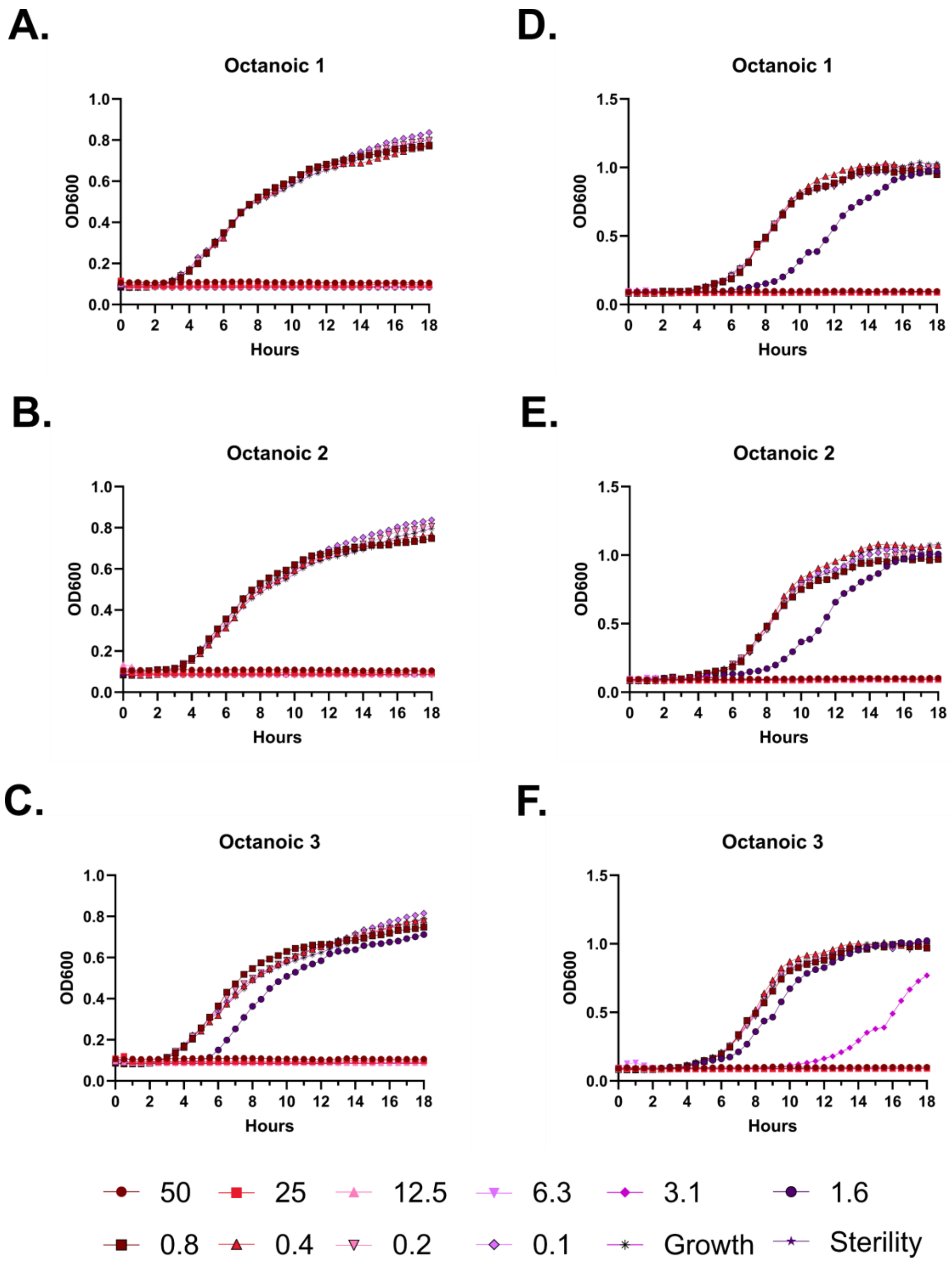
A.
Octanoic 1
D.
Octanoic 1
B.
Octanoic 2
E.
Octanoic 2
C.
Octanoic 3
F.
Octanoic 3
OD600
Hours
50
25
12.5
6.3
3.1
1.6
0.8
0.4
0.2
0.1
Growth
Sterility

**Figure S96 – Antimicrobial susceptibility curves for (#43) FK-13 N-lipidated with octanoic acid.** **A-C.** Independent experiments with *E. coli*. **D-F.** Independent experiments with *P. aeruginosa.* Growth over 18 hours at 37 °C with a gradient of peptide concentrations given in μM.

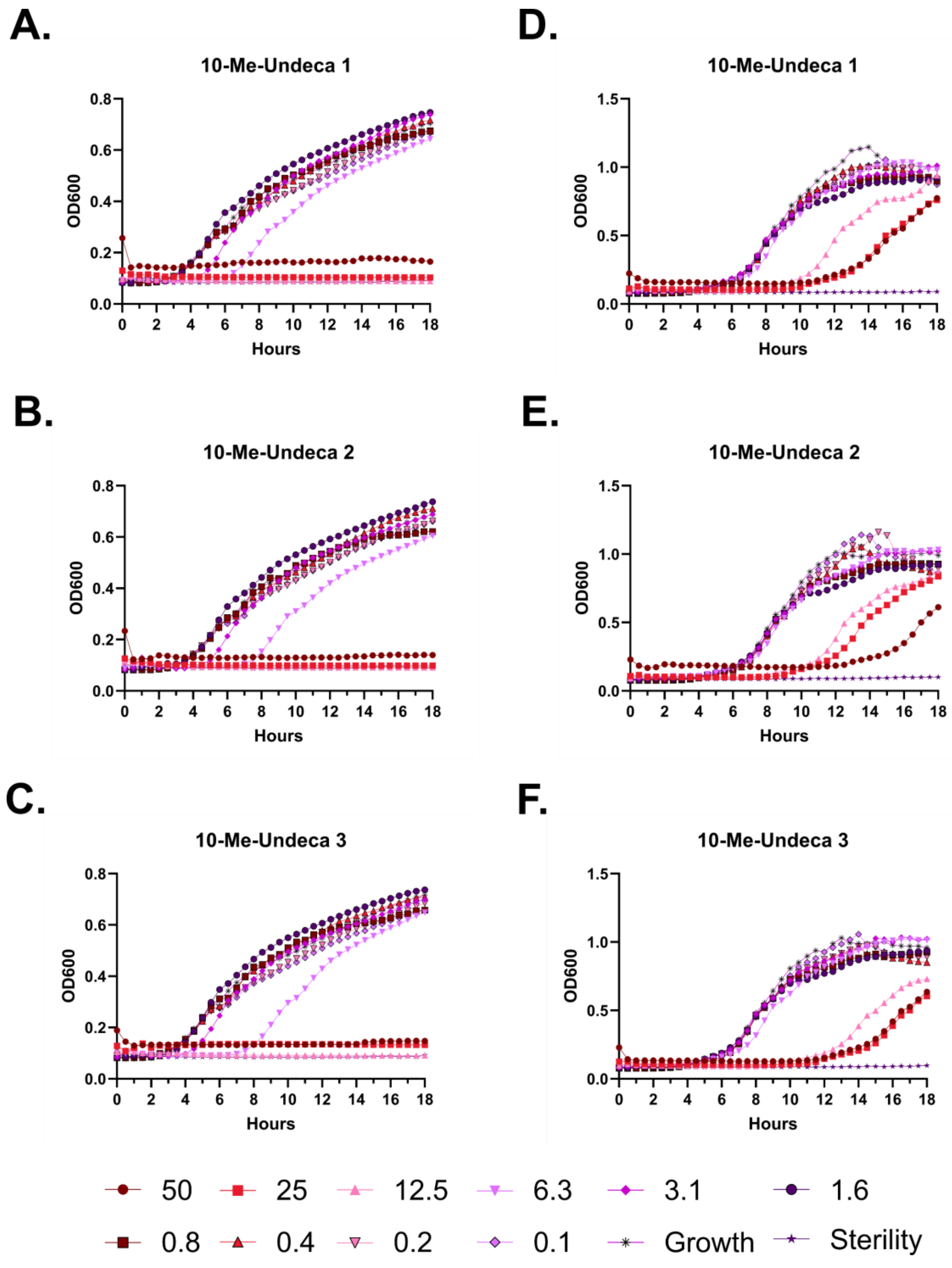
A.
10-Me-Undeca 1
B.
10-Me-Undeca 2
C.
10-Me-Undeca 3
D.
10-Me-Undeca 1
E.
10-Me-Undeca 2
F.
10-Me-Undeca 3
OD600
Hours
50
25
12.5
6.3
3.1
1.6
0.8
0.4
0.2
0.1
Growth
Sterility

**Figure S97 – Antimicrobial susceptibility curves for (#35) FF-14 N-lipidated with 10-methyl undecanoic acid. A-C.** Independent experiments with *E. coli*. **D-F.** Independent experiments with *P. aeruginosa.* Growth over 18 hours at 37 °C with a gradient of peptide concentrations given in μM.

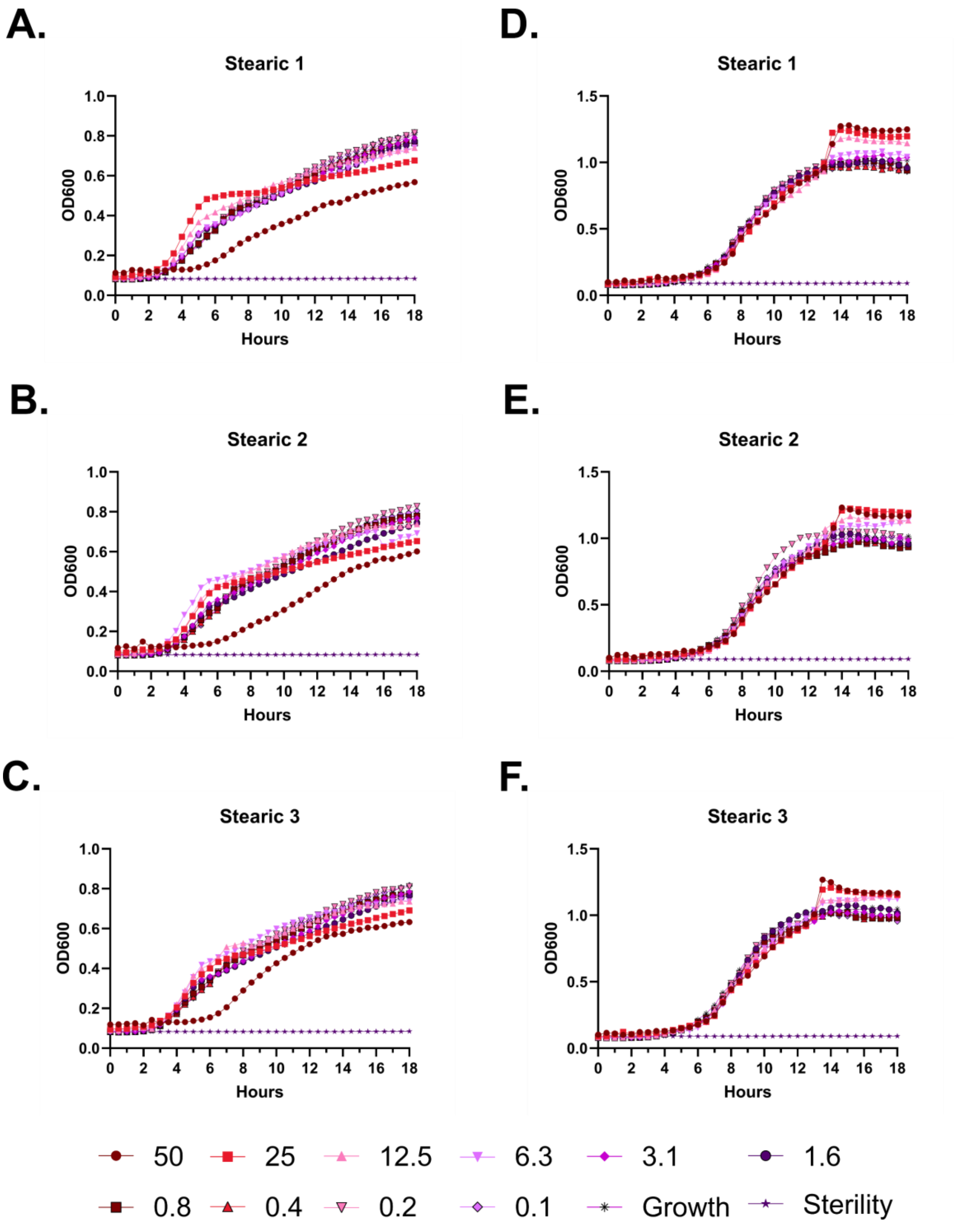
A.
Stearic 1
OD600
Hours
D.
Stearic 1
OD600
Hours
B.
Stearic 2
OD600
Hours
E.
Stearic 2
OD600
Hours
C.
Stearic 3
OD600
Hours
F.
Stearic 3
OD600
Hours
50
25
12.5
6.3
3.1
1.6
0.8
0.4
0.2
0.1
Growth
Sterility

**Figure S98 – Antimicrobial susceptibility curves for (#44) FK-13 N-lipidated with stearic acid.** **A-C.** Independent experiments with *E. coli*. **D-F.** Independent experiments with *P. aeruginosa.* Growth over 18 hours at 37 °C with a gradient of peptide concentrations given in μM.

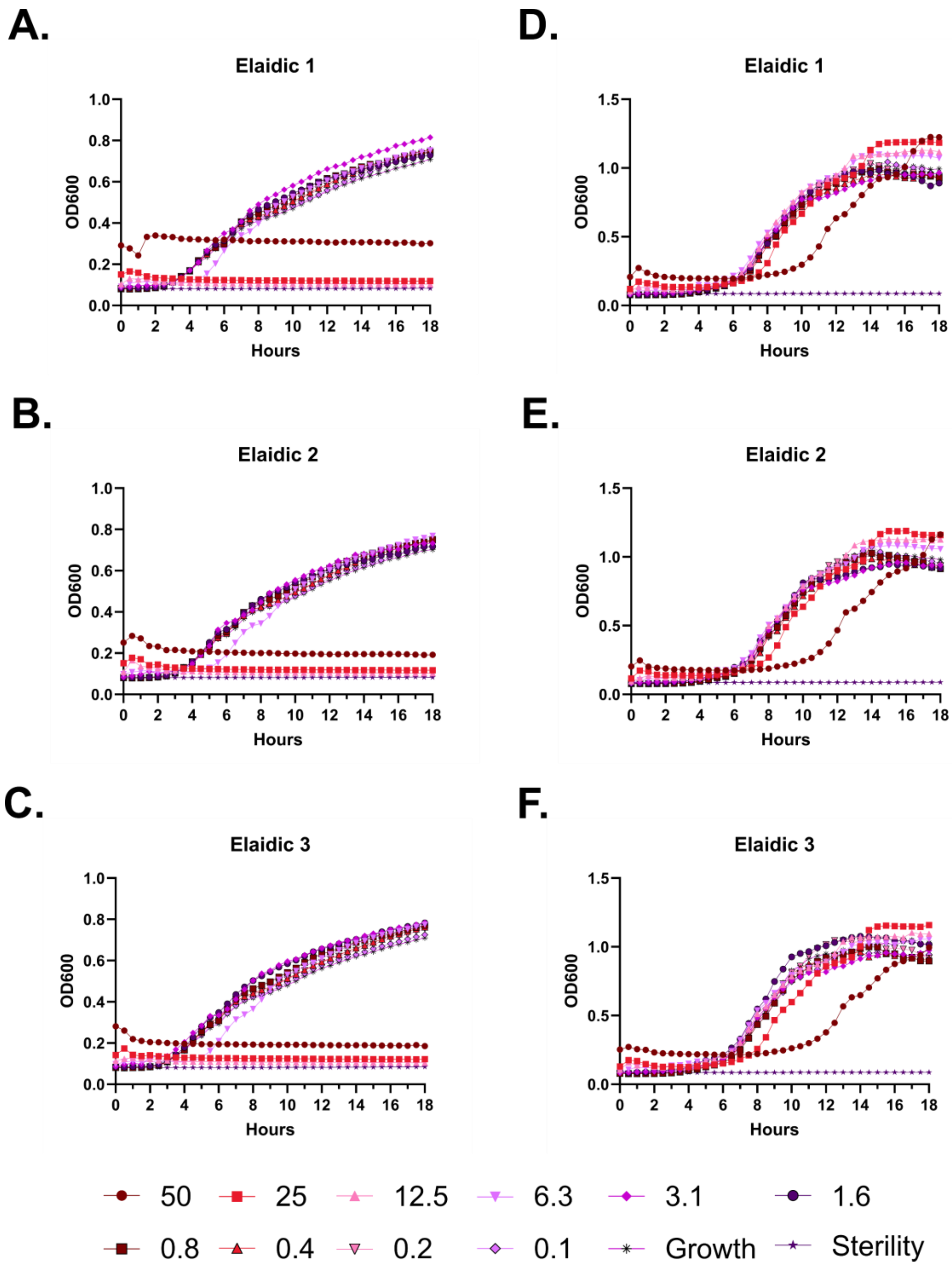
A.
Elaidic 1
D.
Elaidic 1
B.
Elaidic 2
E.
Elaidic 2
C.
Elaidic 3
F.
Elaidic 3
OD600
Hours
50
25
12.5
6.3
3.1
1.6
0.8
0.4
0.2
0.1
Growth
Sterility

**Figure S99 – Antimicrobial susceptibility curves for (#38) FF-14 N-lipidated with elaidic acid. A-C.** Independent experiments with *E. coli*. **D-F.** Independent experiments with *P. aeruginosa.* Growth over 18 hours at 37 °C with a gradient of peptide concentrations given in μM.

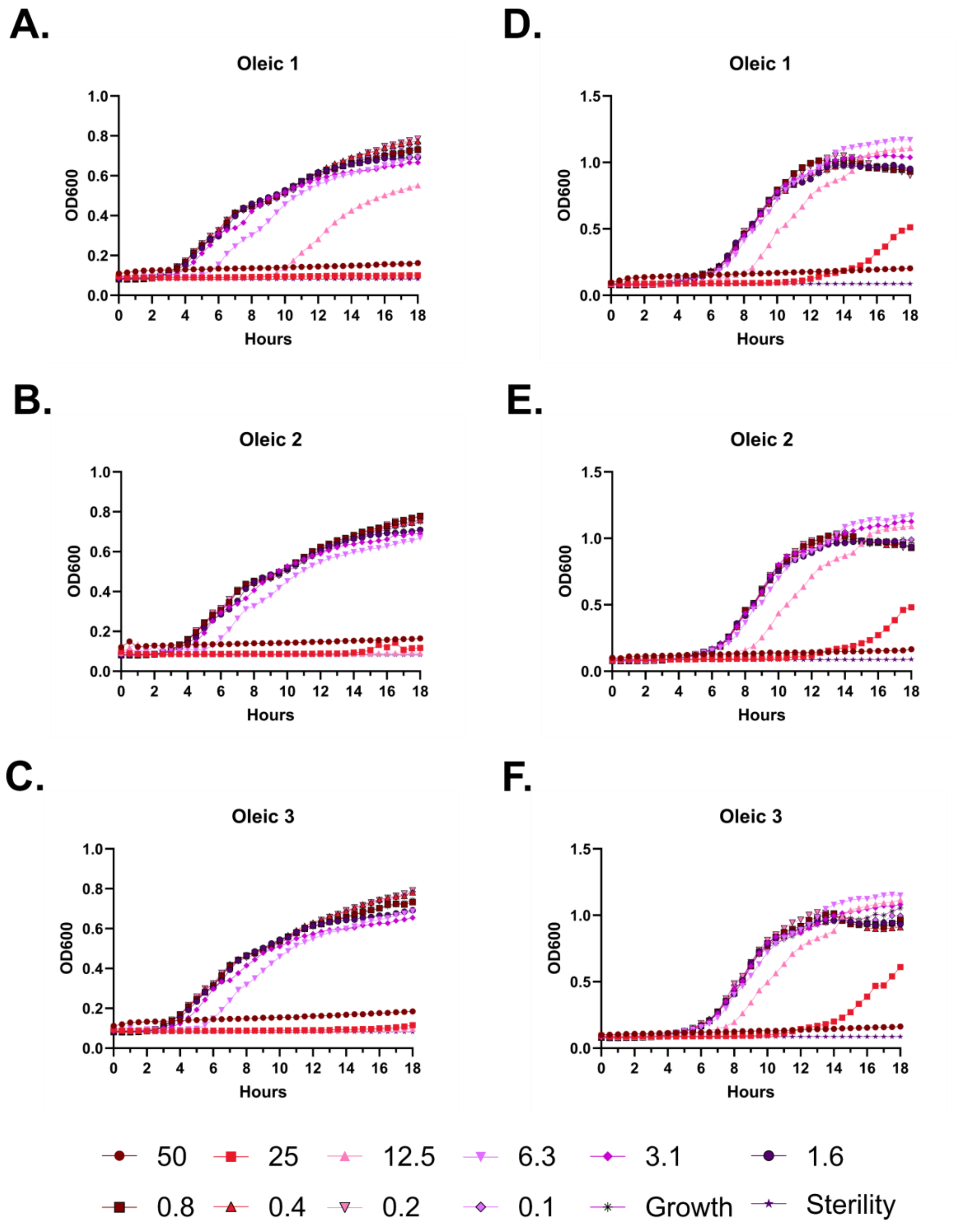
A.
Oleic 1
D.
Oleic 1
B.
Oleic 2
E.
Oleic 2
C.
Oleic 3
F.
Oleic 3
OD600
Hours
50
25
12.5
6.3
3.1
1.6
0.8
0.4
0.2
0.1
Growth
Sterility

**Figure S100 – Antimicrobial susceptibility curves for (#45) FK-13 N-lipidated with oleic acid. A-C.** Independent experiments with *E. coli*. **D-F.** Independent experiments with *P. aeruginosa.* Growth over 18 hours at 37 °C with a gradient of peptide concentrations given in μM.

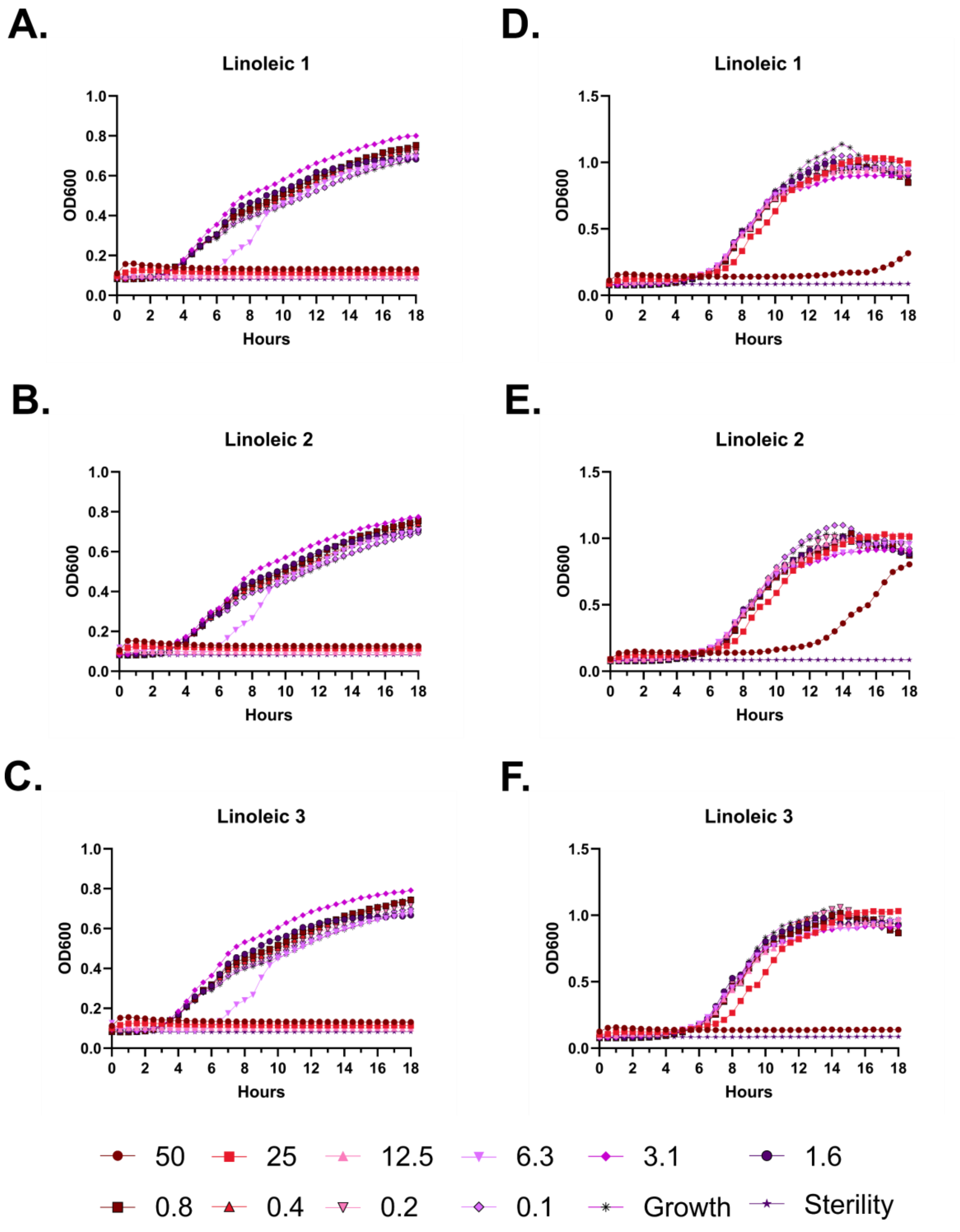
A.
Linoleic 1
B.
Linoleic 2
C.
Linoleic 3
D.
Linoleic 1
E.
Linoleic 2
F.
Linoleic 3
OD600
Hours
50
25
12.5
6.3
3.1
1.6
0.8
0.4
0.2
0.1
Growth
Sterility

**Figure S101 – Antimicrobial susceptibility curves for (#39) FF-14 N-lipidated with linoleic acid.** **A-C.** Independent experiments with *E. coli*. **D-F.** Independent experiments with *P. aeruginosa.* Growth over 18 hours at 37 °C with a gradient of peptide concentrations given in µM.

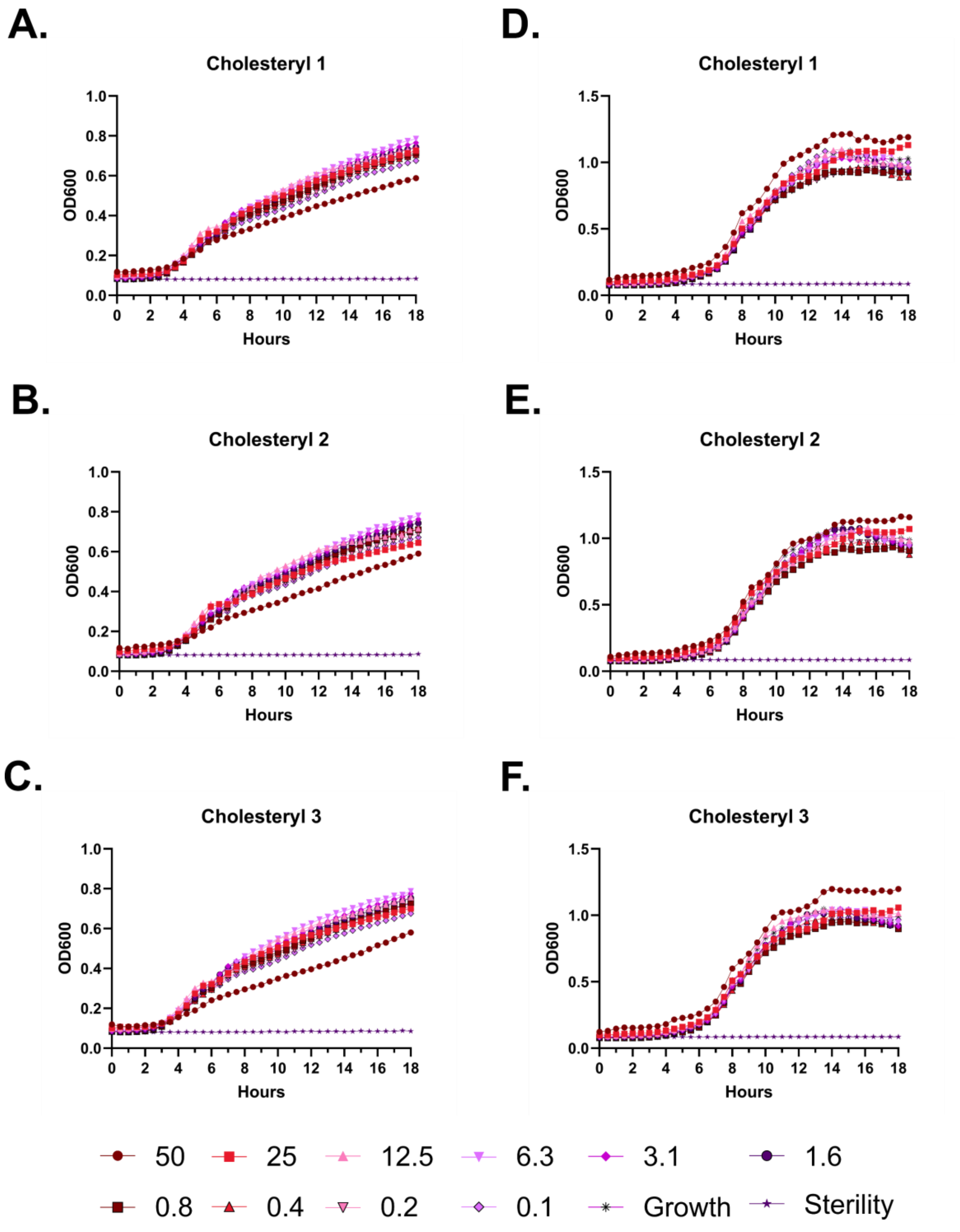
A.
Cholesteryl 1
OD600
Hours
B.
Cholesteryl 2
OD600
Hours
C.
Cholesteryl 3
OD600
Hours
D.
Cholesteryl 1
OD600
Hours
E.
Cholesteryl 2
OD600
Hours
F.
Cholesteryl 3
OD600
Hours
50
25
12.5
6.3
3.1
1.6
0.8
0.4
0.2
0.1
Growth
Sterility

**Figure S102 – Antimicrobial susceptibility curves for (#37) FF-14 N-lipidated with cholesteryl hemisuccinate. A-C.** Independent experiments with *E. coli*. **D-F.** Independent experiments with *P. aeruginosa.* Growth over 18 hours at 37 °C with a gradient of peptide concentrations given in μM.

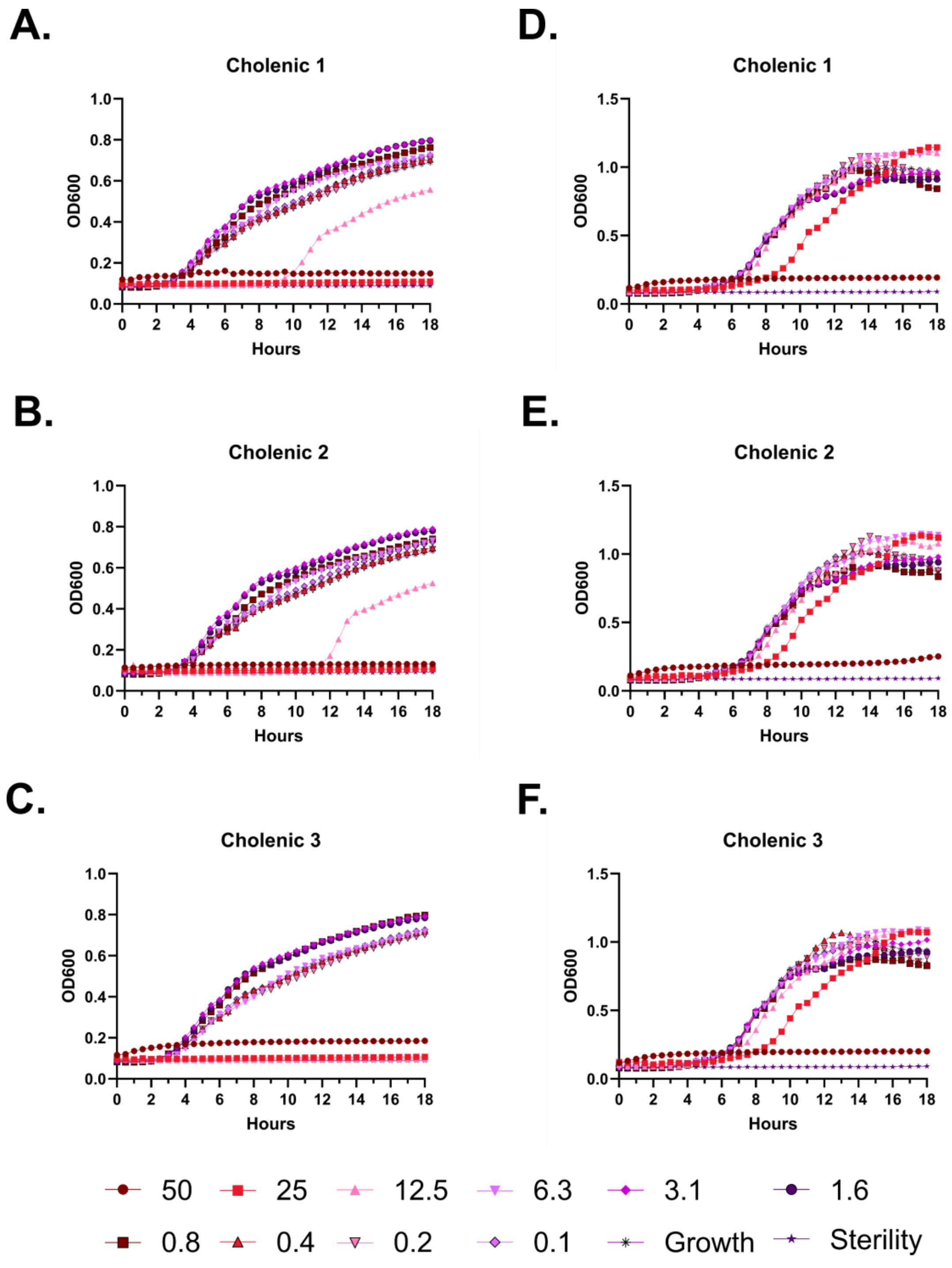
A.
Cholenic 1
D.
Cholenic 1
B.
Cholenic 2
E.
Cholenic 2
C.
Cholenic 3
F.
Cholenic 3
OD600
Hours
50
25
12.5
6.3
3.1
1.6
0.8
0.4
0.2
0.1
Growth
Sterility

**Figure S103 – Antimicrobial susceptibility curves for (#42) FK-13 N-lipidated with cholenic acid. A-C.** Independent experiments with *E. coli*. **D-F.** Independent experiments with *Pseudomonas.* Growth over 18 hours at 37 °C with a gradient of peptide concentrations given in μM.

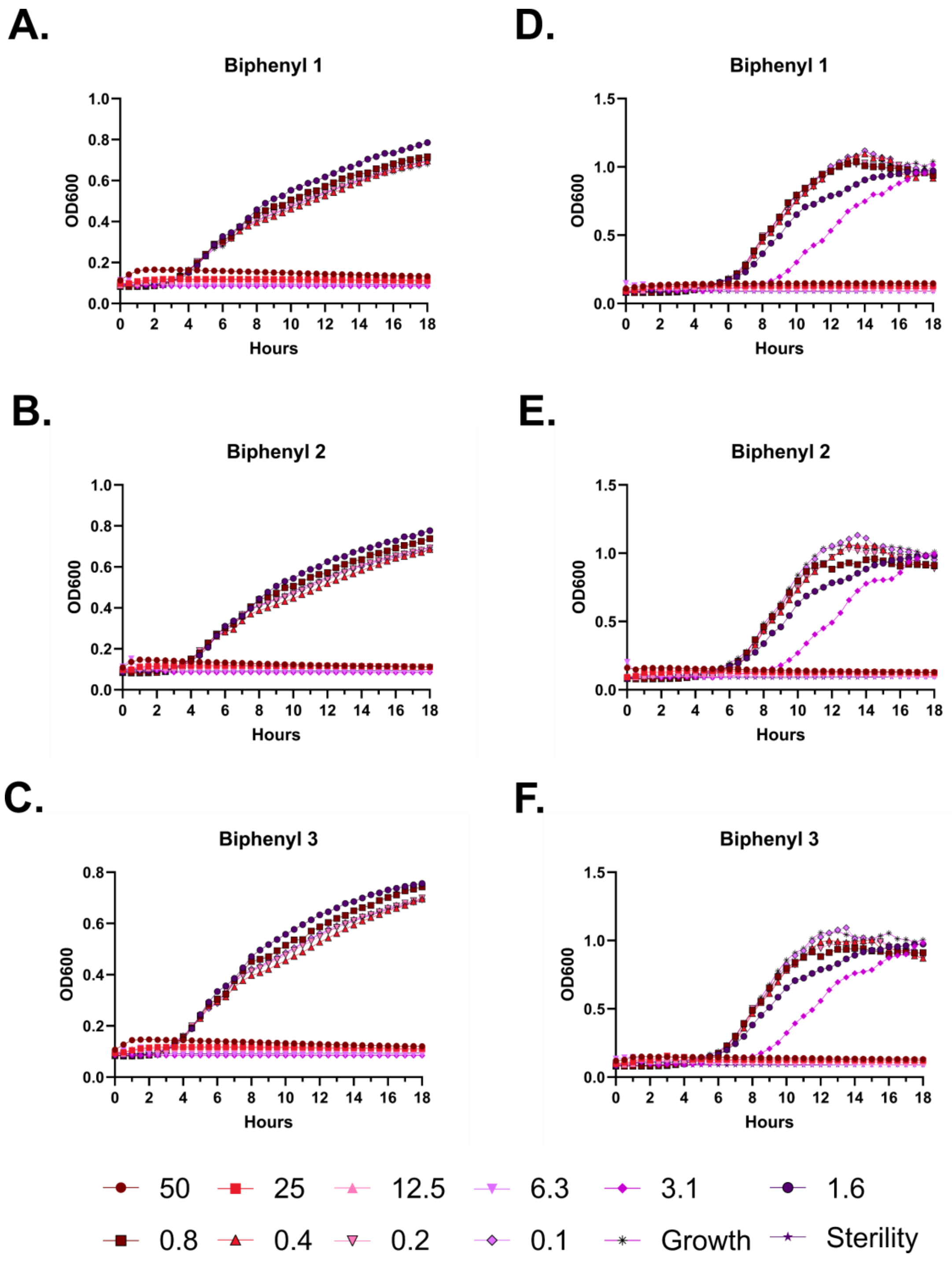
A.
Biphenyl 1
OD600
Hours
B.
Biphenyl 2
OD600
Hours
C.
Biphenyl 3
OD600
Hours
D.
Biphenyl 1
OD600
Hours
E.
Biphenyl 2
OD600
Hours
F.
Biphenyl 3
OD600
Hours
50
25
12.5
6.3
3.1
1.6
0.8
0.4
0.2
0.1
Growth
Sterility

**Figure S104 – Antimicrobial susceptibility curves for (#41) FK-13 N-lipidated with biphenyl-4-carboxylic acid. A-C.** Independent experiments with *E. coli.* **D-F.** Independent experiments with *P. aeruginosa.* Growth over 18 hours at 37 °C with a gradient of peptide concentrations given in μM.

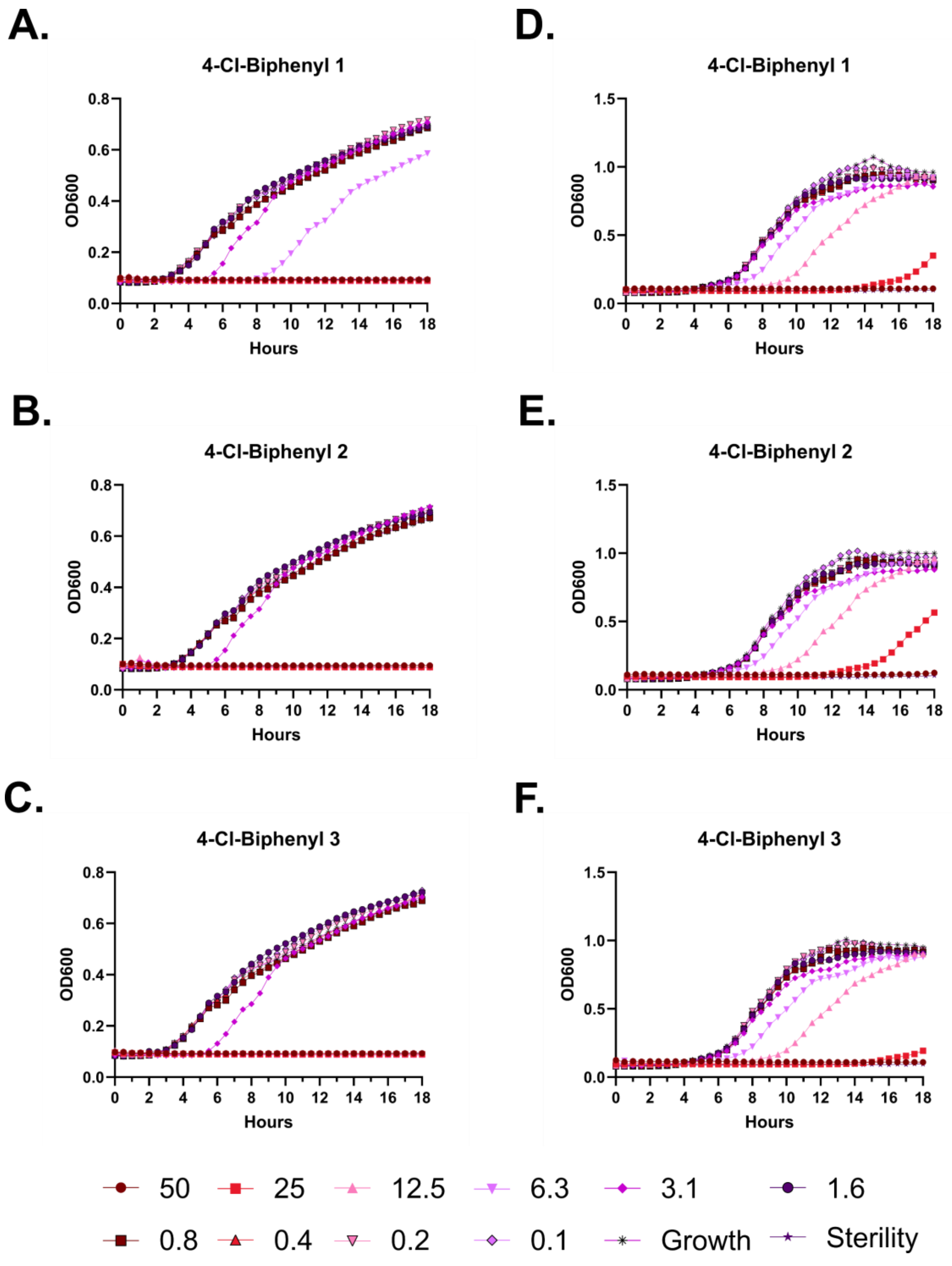
A.
4-Cl-Biphenyl 1
OD600
Hours
D.
4-Cl-Biphenyl 1
OD600
Hours
B.
4-Cl-Biphenyl 2
OD600
Hours
E.
4-Cl-Biphenyl 2
OD600
Hours
C.
4-Cl-Biphenyl 3
OD600
Hours
F.
4-Cl-Biphenyl 3
OD600
Hours
50
25
12.5
6.3
3.1
1.6
0.8
0.4
0.2
0.1
Growth
Sterility

**Figure S105 – Antimicrobial susceptibility curves for (#36) FF-14 N-lipidated with 4-chloro-biphenyl-4-carboxylic acid. A-C.** Independent experiments with *E. coli*. **D-F.** Independent experiments with *P. aeruginosa.* Growth over 18 hours at 37 °C with a gradient of peptide concentrations given in µM.

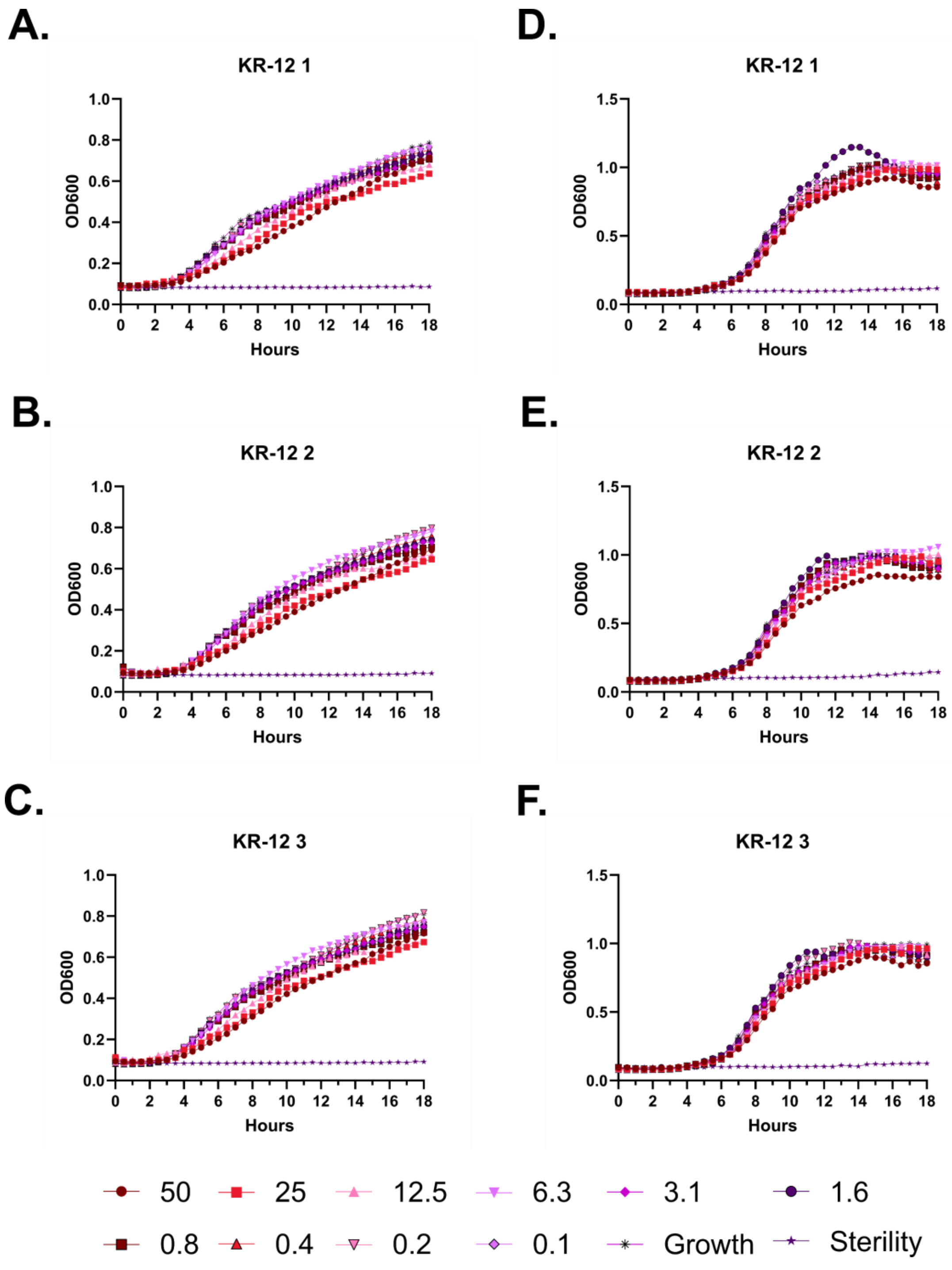

A.
KR-12 1
B.
KR-12 2
C.
KR-12 3
D.
KR-12 1
E.
KR-12 2
F.
KR-12 3
OD600
Hours
50
25
12.5
6.3
3.1
1.6
0.8
0.4
0.2
0.1
Growth
Sterility

**Figure S106 – Antimicrobial susceptibility curves for (#10) KR-12. A-C.** Independent experiments with *E. coli*. **D-F.** Independent experiments with *P. aeruginosa.* Growth over 18 hours at 37 °C with a gradient of peptide concentrations given in μM.

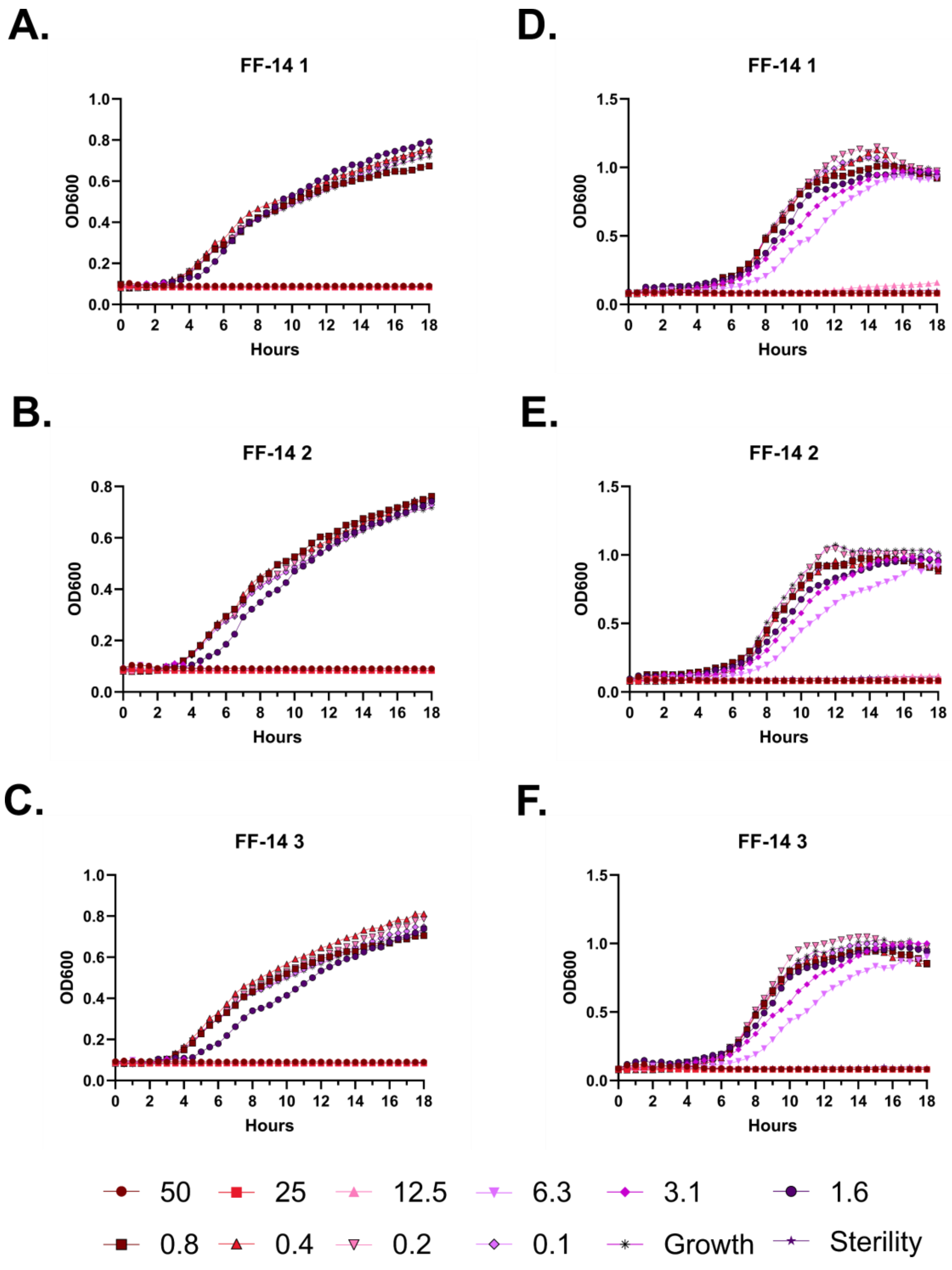

A.
FF-14 1
OD600
Hours
B.
FF-14 2
C.
FF-14 3
D.
E.
F.
50
25
12.5
6.3
3.1
1.6
0.8
0.4
0.2
0.1
Growth
Sterility

**Figure S107 – Antimicrobial susceptibility curves for (#18) FF-14. A-C.** Independent experiments with *E. coli*. **D-F.** Independent experiments with *P. aeruginosa.* Growth over 18 hours at 37 °C with a gradient of peptide concentrations given in μM.

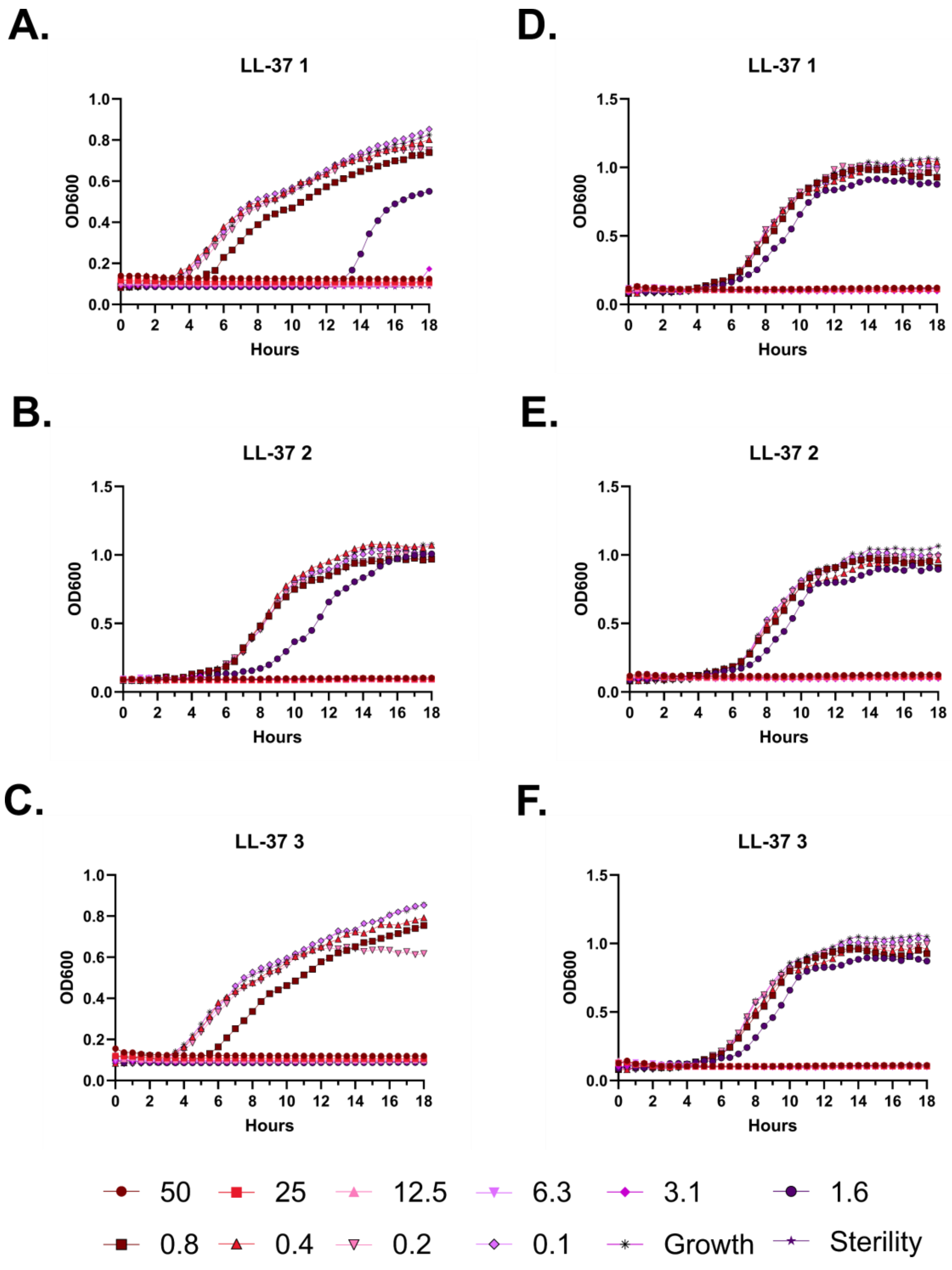

A.
LL-37 1
OD600
Hours
B.
LL-37 2
OD600
Hours
C.
LL-37 3
OD600
Hours
D.
LL-37 1
OD600
Hours
E.
LL-37 2
OD600
Hours
F.
LL-37 3
OD600
Hours
50
25
12.5
6.3
3.1
1.6
0.8
0.4
0.2
0.1
Growth
Sterility

**Figure S108 – Antimicrobial susceptibility curves for (#1) LL-37. A-C.** Independent experiments with *E. coli*. **D-F.** Independent experiments with *P. aeruginosa.* Growth over 18 hours at 37 °C with a gradient of peptide concentrations given in μM.

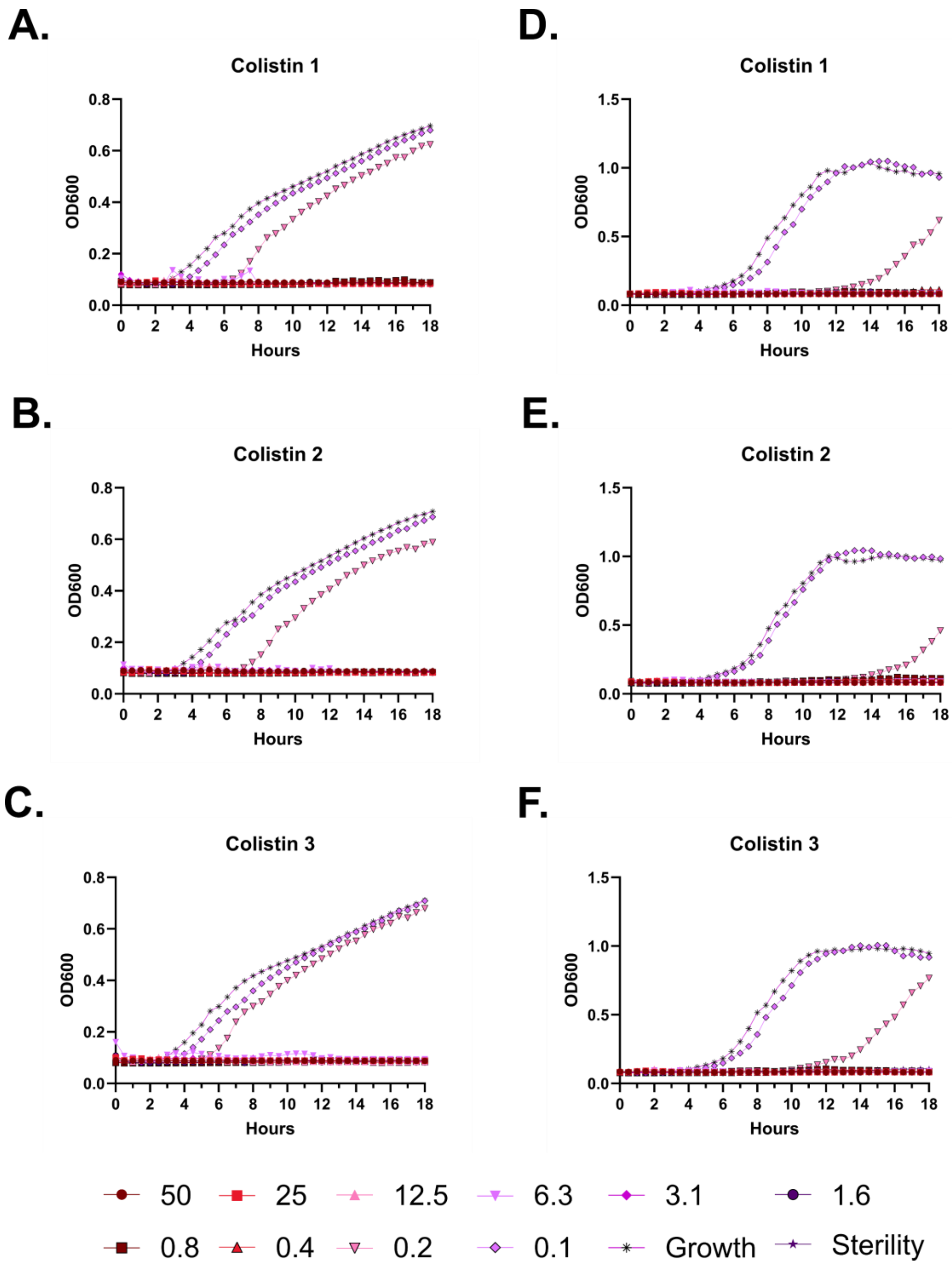

A.
Colistin 1
OD600
Hours
B.
Colistin 2
OD600
Hours
C.
Colistin 3
OD600
Hours
D.
Colistin 1
OD600
Hours
E.
Colistin 2
OD600
Hours
F.
Colistin 3
OD600
Hours
50
25
12.5
6.3
3.1
1.6
0.8
0.4
0.2
0.1
Growth
Sterility

**Figure S109 – Antimicrobial susceptibility curves for colistin. A-C.** Independent experiments with *E. coli*. **D-F.** Independent experiments with *P. aeruginosa.* Growth over 18 hours at 37 °C with a gradient of peptide concentrations given in μM.

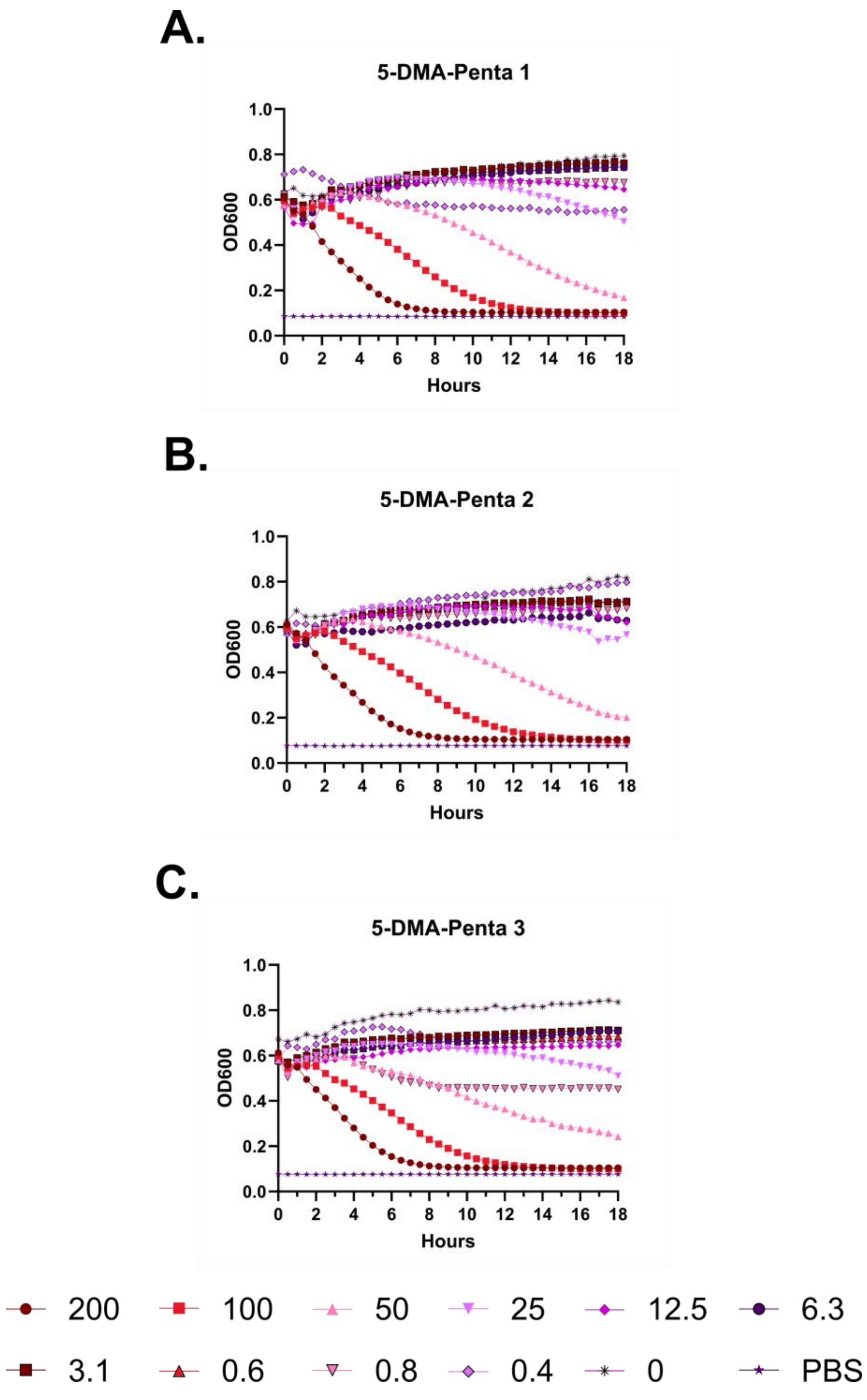

A.
5-DMA-Penta 1
OD600
Hours
B.
5-DMA-Penta 2
OD600
Hours
C.
5-DMA-Penta 3
OD600
Hours
200
100
50
25
12.5
6.3
3.1
0.6
0.8
0.4
0
PBS

**Figure S110 – (#34) FF-14 N-lipidated with 5-dimethylamino pentanoic acid kinetic hemolysis assay. A.** Defibrinated sheep blood was incubated in the presence of the indicated peptide concentrations at 37 °C with monitoring of OD600 over 18 hours, where decreasing OD600 indicates hemolysis and each panel indicates an independent experiment. Peptide concentrations are in μM.

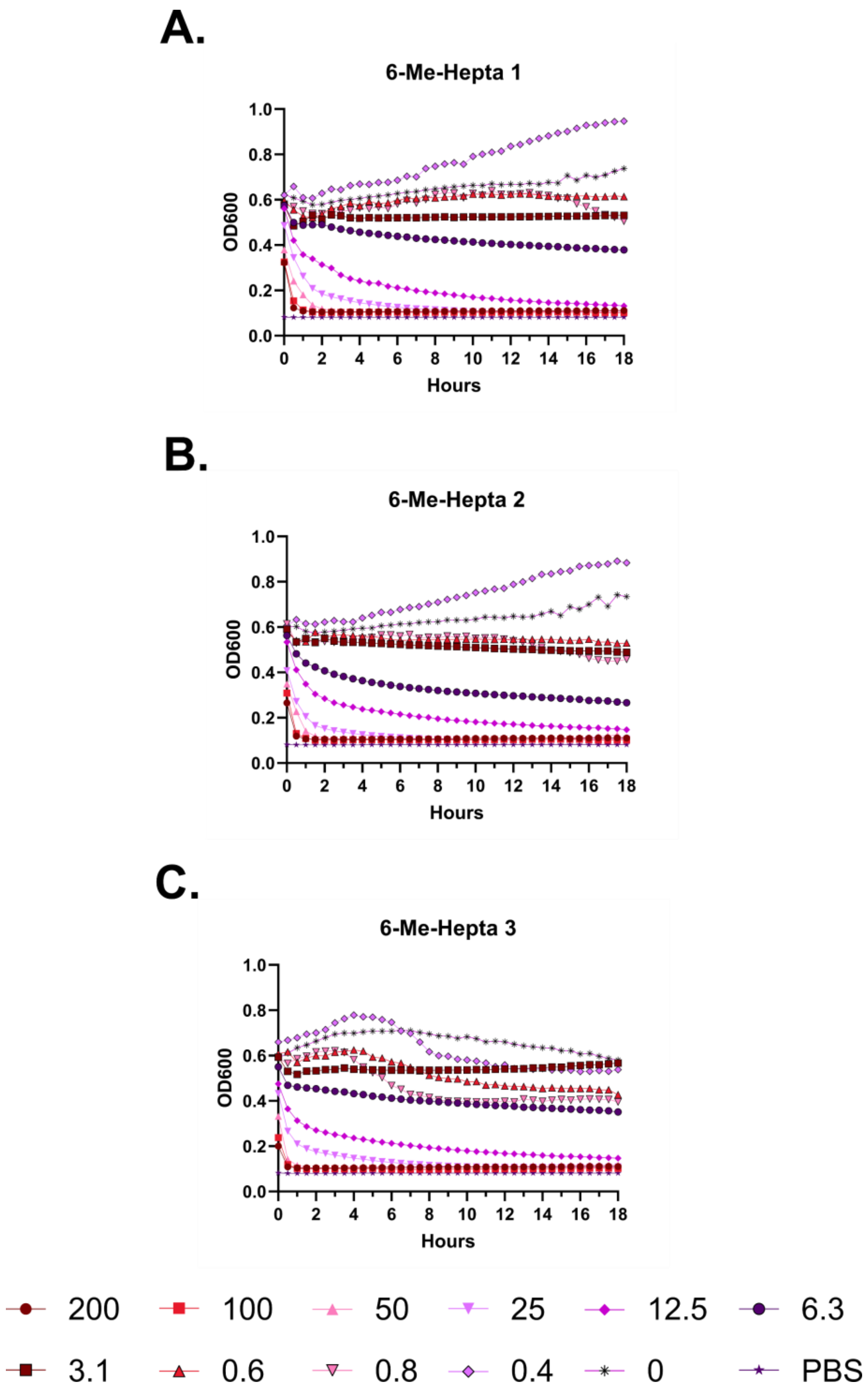
A.
6-Me-Hepta 1
B.
6-Me-Hepta 2
C.
6-Me-Hepta 3
OD600
Hours
1.0
0.8
0.6
0.4
0.2
0.0
0
2
4
6
8
10
12
14
16
18
200
100
50
25
12.5
6.3
3.1
0.6
0.8
0.4
0
PBS

**Figure S111 – (#40) FK-13 N-lipidated with 6-methyl heptanoic acid kinetic hemolysis assay. A.** Defibrinated sheep blood was incubated in the presence of the indicated peptide concentrations at 37 °C with monitoring of OD600 over 18 hours, where decreasing OD600 indicates hemolysis and each panel indicates an independent experiment. Peptide concentrations are in μM.

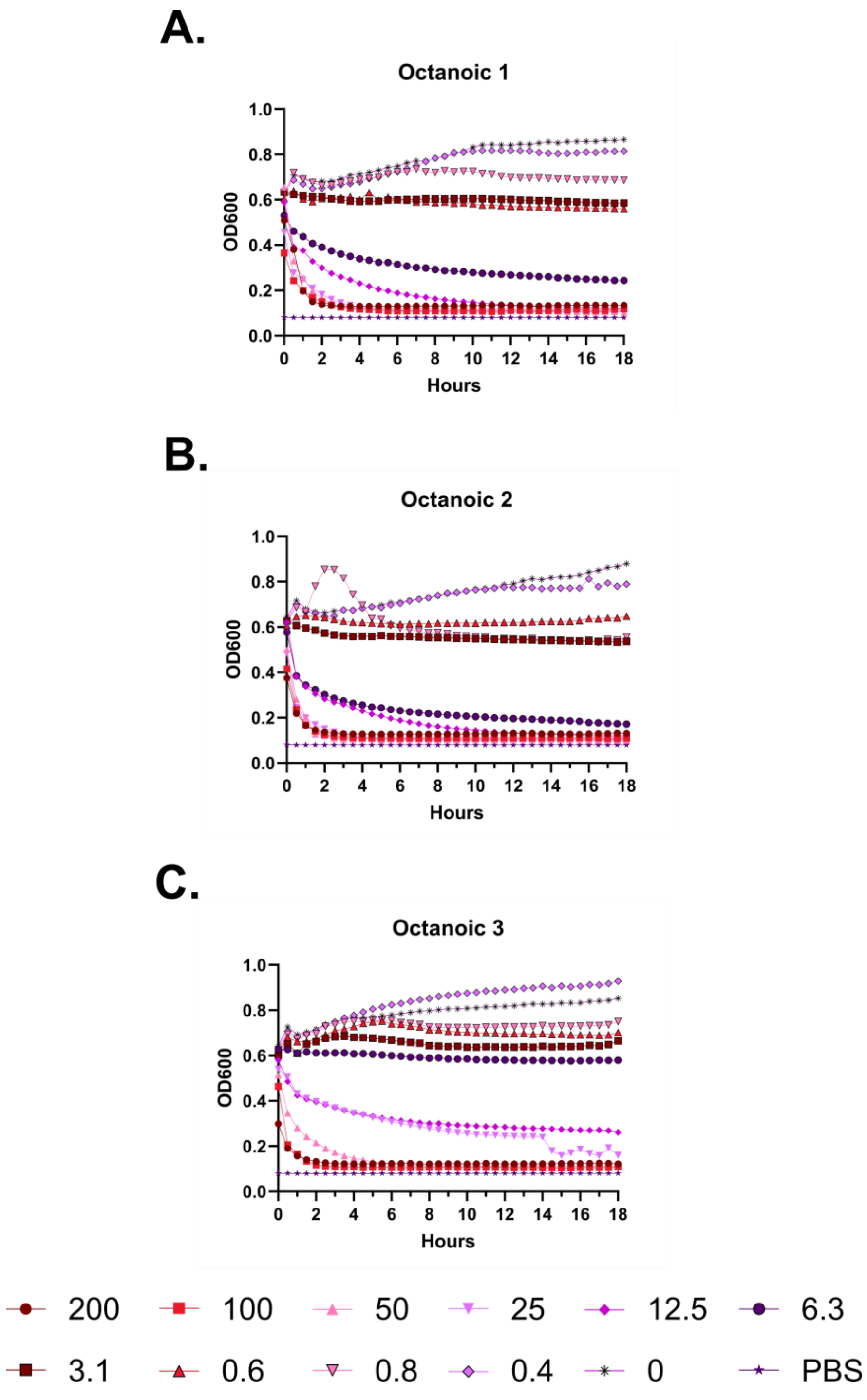
A.
Octanoic 1
OD600
Hours
B.
Octanoic 2
OD600
Hours
C.
Octanoic 3
OD600
Hours
200
100
50
25
12.5
6.3
3.1
0.6
0.8
0.4
0
PBS

**Figure S112 – (#43) FK-13 N-lipidated with octanoic acid kinetic hemolysis assay. A.** Defibrinated sheep blood was incubated in the presence of the indicated peptide concentrations at 37 °C with monitoring of OD600 over 18 hours, where decreasing OD600 indicates hemolysis and each panel indicates an independent experiment. Peptide concentrations are in μM.

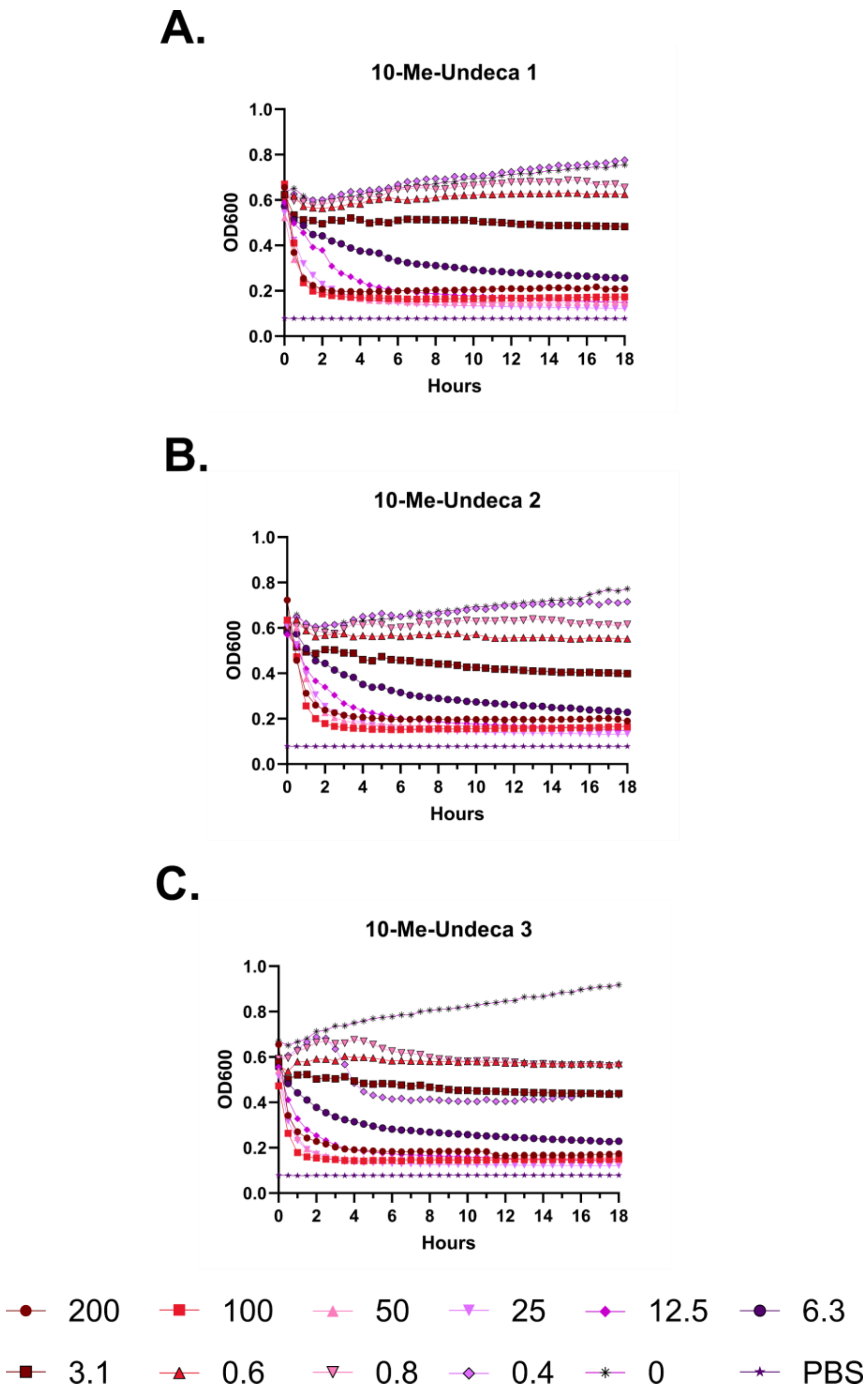
A.
10-Me-Undeca 1
OD600
Hours
B.
10-Me-Undeca 2
OD600
Hours
C.
10-Me-Undeca 3
OD600
Hours
200
100
50
25
12.5
6.3
3.1
0.6
0.8
0.4
0
PBS

**Figure S113 – (#35) FF-14 N-lipidated with 10-methyl undecanoic acid kinetic hemolysis assay. A.** Defibrinated sheep blood was incubated in the presence of the indicated peptide concentrations at 37 °C with monitoring of OD600 over 18 hours, where decreasing OD600 indicates hemolysis and each panel indicates an independent experiment. Peptide concentrations are in μM.

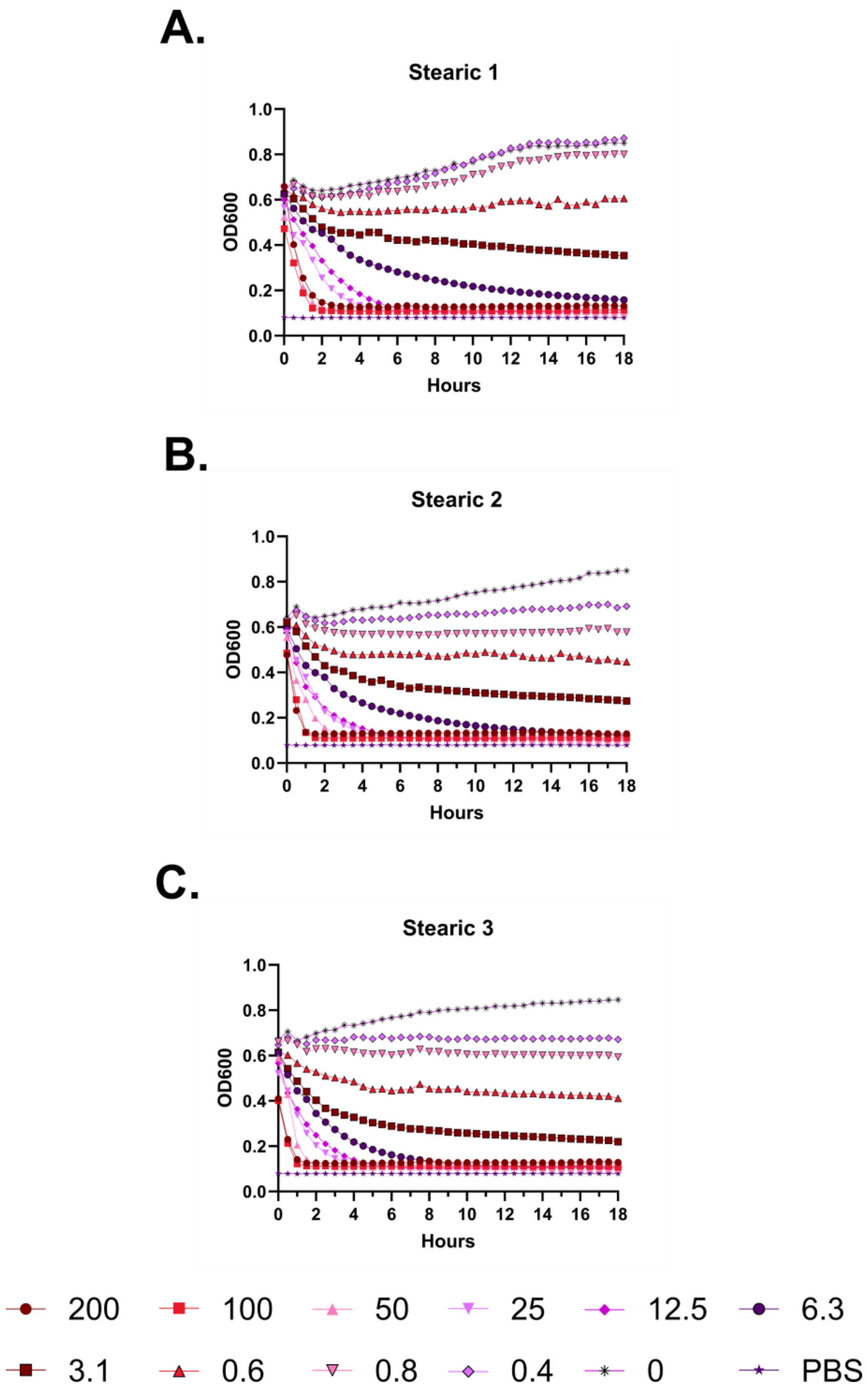
A.
Stearic 1
OD600
Hours
B.
Stearic 2
OD600
Hours
C.
Stearic 3
OD600
Hours
200
100
50
25
12.5
6.3
3.1
0.6
0.8
0.4
0
PBS

**Figure S114 – (#44) FK-13 N-lipidated with stearic acid kinetic hemolysis assay. A.** Defibrinated sheep blood was incubated in the presence of the indicated peptide concentrations at 37 °C with monitoring of OD600 over 18 hours, where decreasing OD600 indicates hemolysis and each panel indicates an independent experiment. Peptide concentrations are in µM.

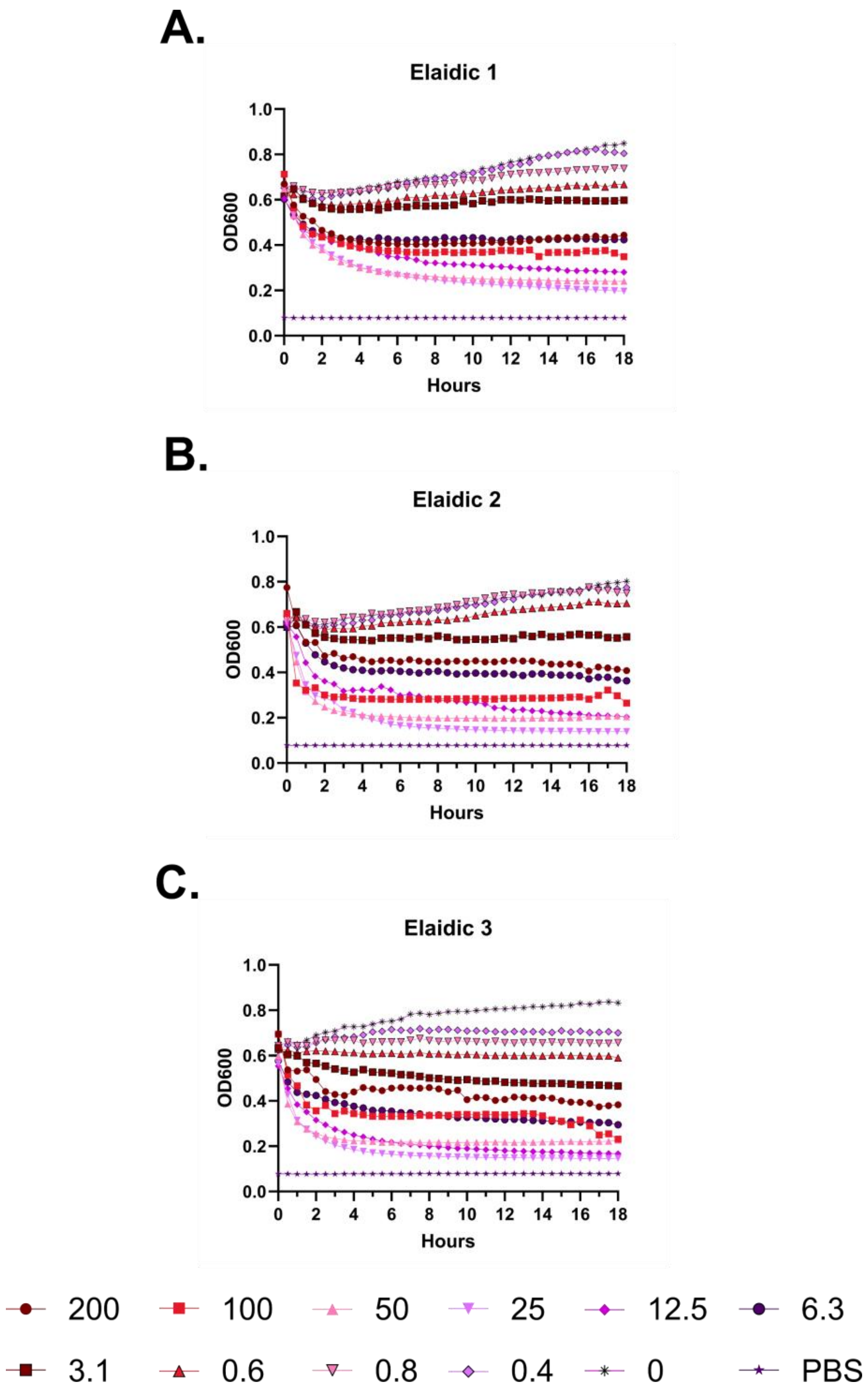
A.
Elaidic 1
OD600
Hours
B.
Elaidic 2
OD600
Hours
C.
Elaidic 3
OD600
Hours
200
100
50
25
12.5
6.3
3.1
0.6
0.8
0.4
0
PBS

**Figure S115 – (#38) FF-14 N-lipidated with elaidic acid kinetic hemolysis assay. A.** Defibrinated sheep blood was incubated in the presence of the indicated peptide concentrations at 37 °C with monitoring of OD600 over 18 hours, where decreasing OD600 indicates hemolysis and each panel indicates an independent experiment. Peptide concentrations are in µM.

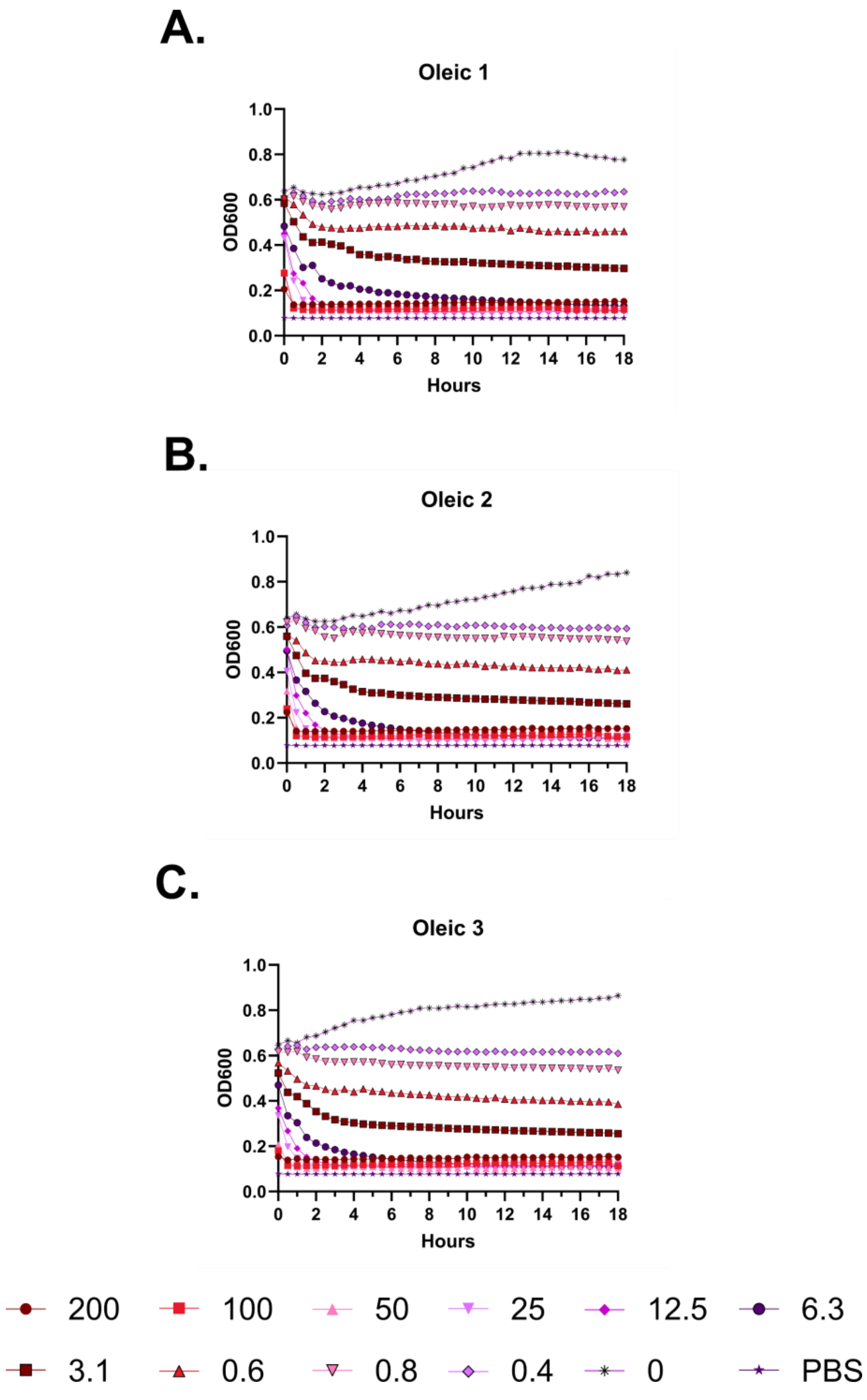
A.
Oleic 1
OD600
1.0
0.8
0.6
0.4
0.2
0.0
0 2 4 6 8 10 12 14 16 18
Hours
B.
Oleic 2
OD600
1.0
0.8
0.6
0.4
0.2
0.0
0 2 4 6 8 10 12 14 16 18
Hours
C.
Oleic 3
OD600
1.0
0.8
0.6
0.4
0.2
0.0
0 2 4 6 8 10 12 14 16 18
Hours
200
100
50
25
12.5
6.3
3.1
0.6
0.8
0.4
0
PBS

**Figure S116 – (#45) FK-13 N-lipidated with oleic acid kinetic hemolysis assay. A.** Defibrinated sheep blood was incubated in the presence of the indicated peptide concentrations at 37 °C with monitoring of OD600 over 18 hours, where decreasing OD600 indicates hemolysis and each panel indicates an independent experiment. Peptide concentrations are in µM.

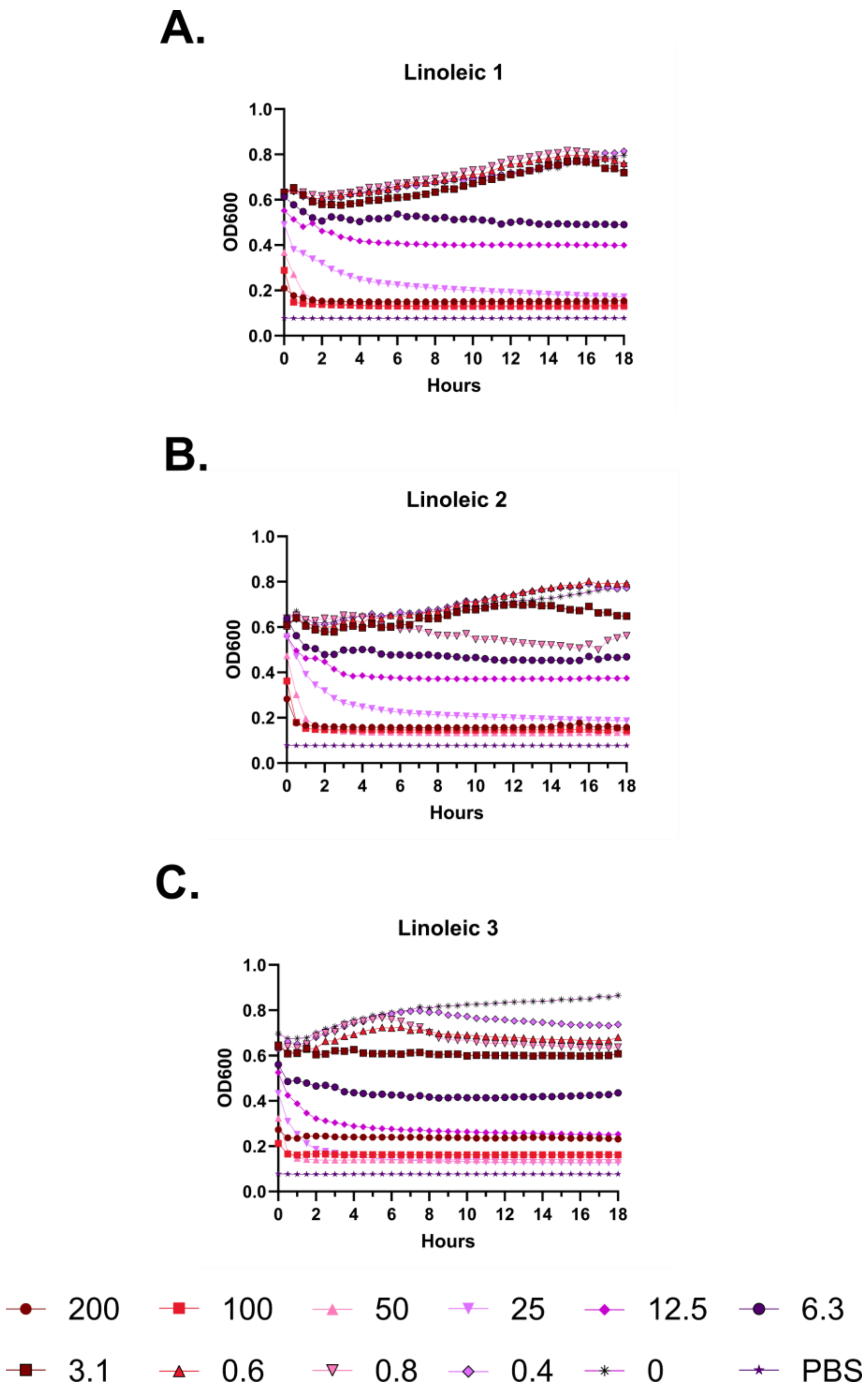
A.
Linoleic 1
OD600
Hours
B.
Linoleic 2
OD600
Hours
C.
Linoleic 3
OD600
Hours
200
100
50
25
12.5
6.3
3.1
0.6
0.8
0.4
0
PBS

**Figure S117 – (#39) FF-14 N-lipidated with linoleic acid kinetic hemolysis assay. A.** Defibrinated sheep blood was incubated in the presence of the indicated peptide concentrations at 37 °C with monitoring of OD600 over 18 hours, where decreasing OD600 indicates hemolysis and each panel indicates an independent experiment. Peptide concentrations are in µM.

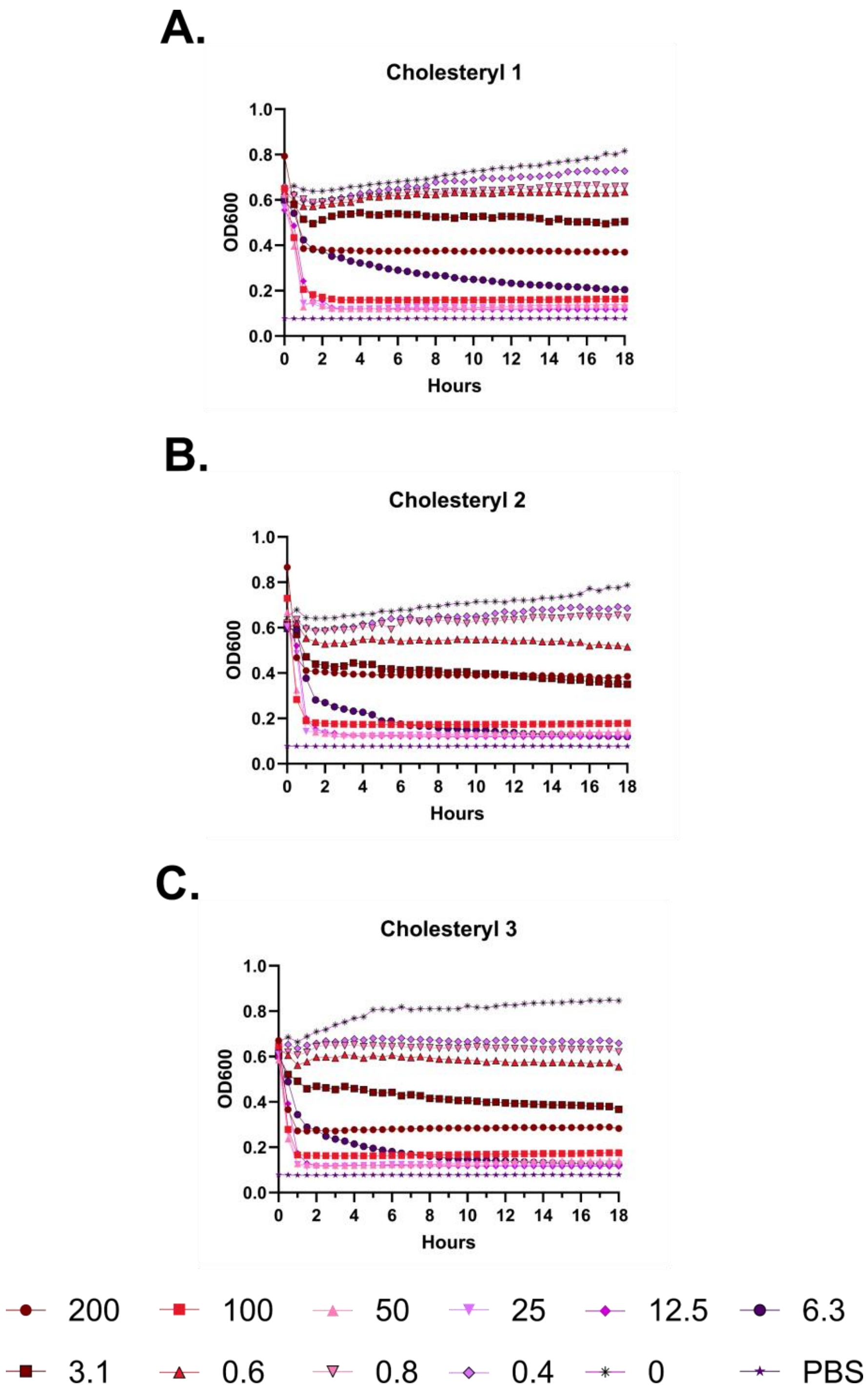
A.
Cholesteryl 1
OD600
Hours
B.
Cholesteryl 2
OD600
Hours
C.
Cholesteryl 3
OD600
Hours
200
100
50
25
12.5
6.3
3.1
0.6
0.8
0.4
0
PBS

**Figure S118 – (#37) FF-14 N-lipidated with cholesteryl hemisuccinate kinetic hemolysis assay. A.** Defibrinated sheep blood was incubated in the presence of the indicated peptide concentrations at 37 °C with monitoring of OD600 over 18 hours, where decreasing OD600 indicates hemolysis and each panel indicates an independent experiment. Peptide concentrations are in µM.

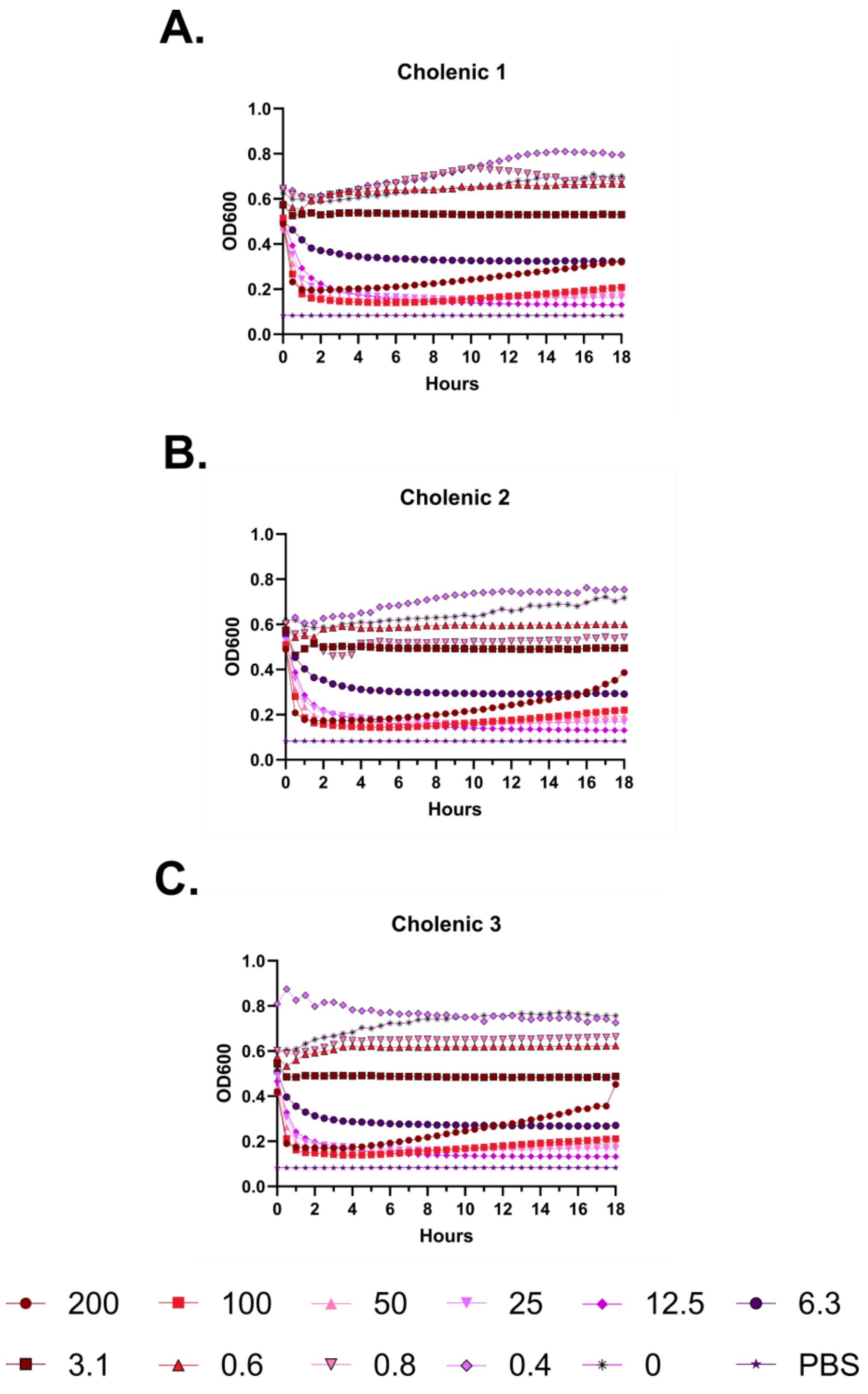
A.
Cholenic 1
OD600
Hours
B.
Cholenic 2
OD600
Hours
C.
Cholenic 3
OD600
Hours
200
100
50
25
12.5
6.3
3.1
0.6
0.8
0.4
0
PBS

**Figure S119 – (#42) FK-13 N-lipidated with cholenic acid kinetic hemolysis assay. A.** Defibrinated sheep blood was incubated in the presence of the indicated peptide concentrations at 37 °C with monitoring of OD600 over 18 hours, where decreasing OD600 indicates hemolysis and each panel indicates an independent experiment. Peptide concentrations are in µM.

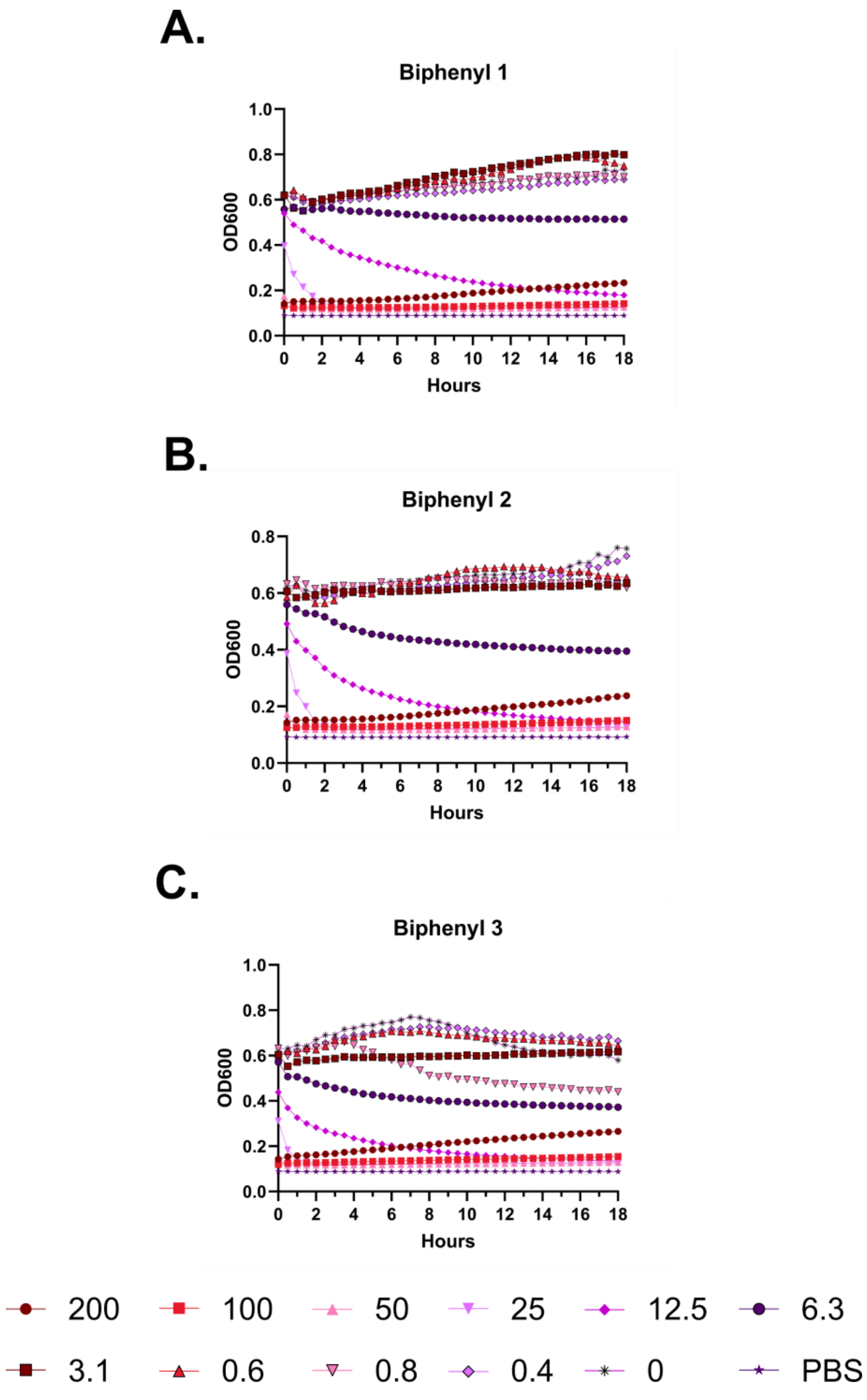
A.
Biphenyl 1
OD600
Hours
B.
Biphenyl 2
OD600
Hours
C.
Biphenyl 3
OD600
Hours
200
100
50
25
12.5
6.3
3.1
0.6
0.8
0.4
0
PBS

**Figure S120 – (#41) FK-13 N-lipidated with biphenyl-4-carboxylic acid kinetic hemolysis assay. A.** Defibrinated sheep blood was incubated in the presence of the indicated peptide concentrations at 37 °C with monitoring of OD600 over 18 hours, where decreasing OD600 indicates hemolysis and each panel indicates an independent experiment. Peptide concentrations are in μM.

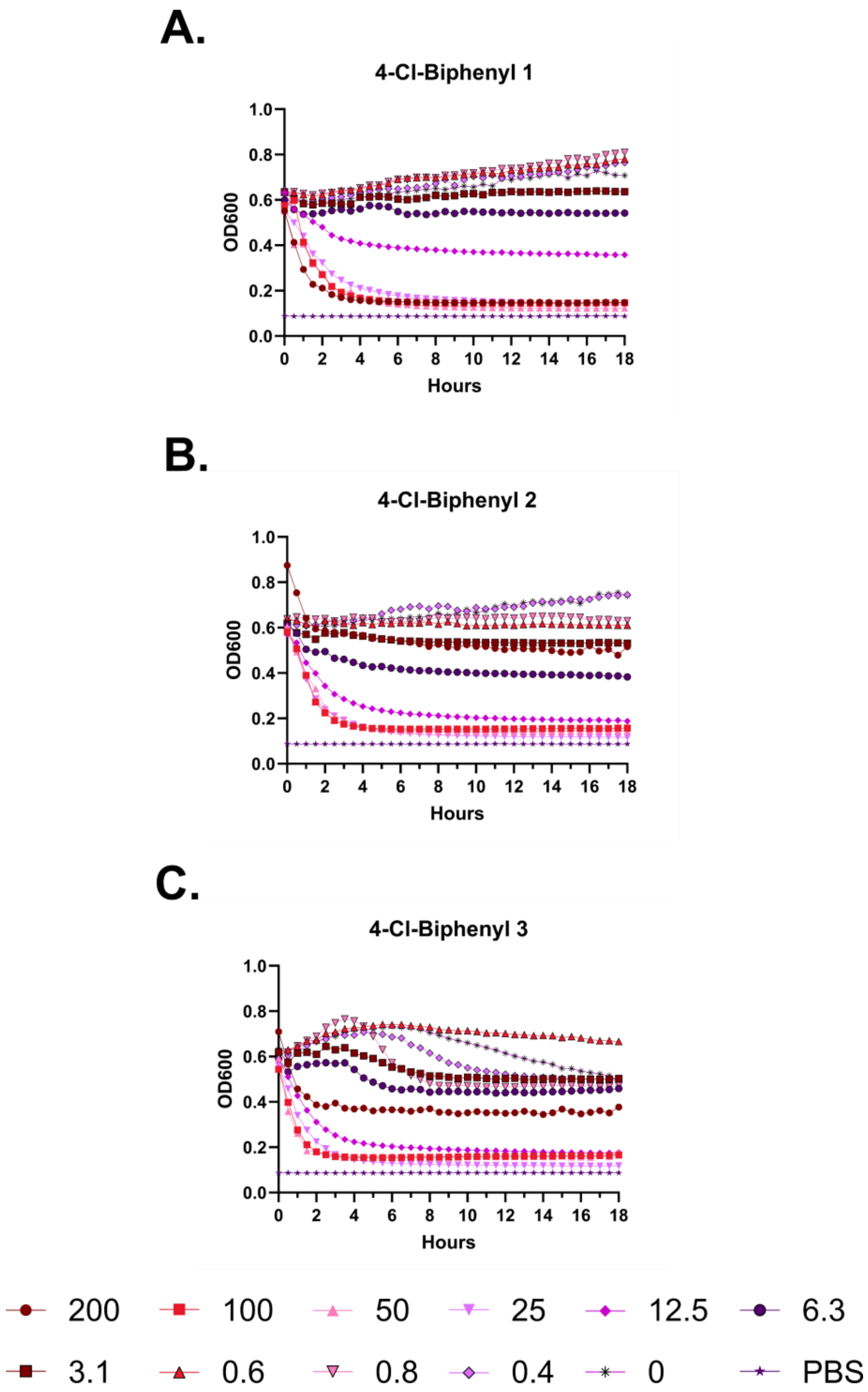
A.
4-Cl-Biphenyl 1
OD600
Hours
B.
4-Cl-Biphenyl 2
OD600
Hours
C.
4-Cl-Biphenyl 3
OD600
Hours
200
100
50
25
12.5
6.3
3.1
0.6
0.8
0.4
0
PBS

**Figure S121 – (#36) FF-14 N-lipidated with 4-chloro-biphenyl-4-carboxylic acid kinetic hemolysis assay. A.** Defibrinated sheep blood was incubated in the presence of the indicated peptide concentrations at 37 °C with monitoring of OD600 over 18 hours, where decreasing OD600 indicates hemolysis and each panel indicates an independent experiment. Peptide concentrations are in μM.

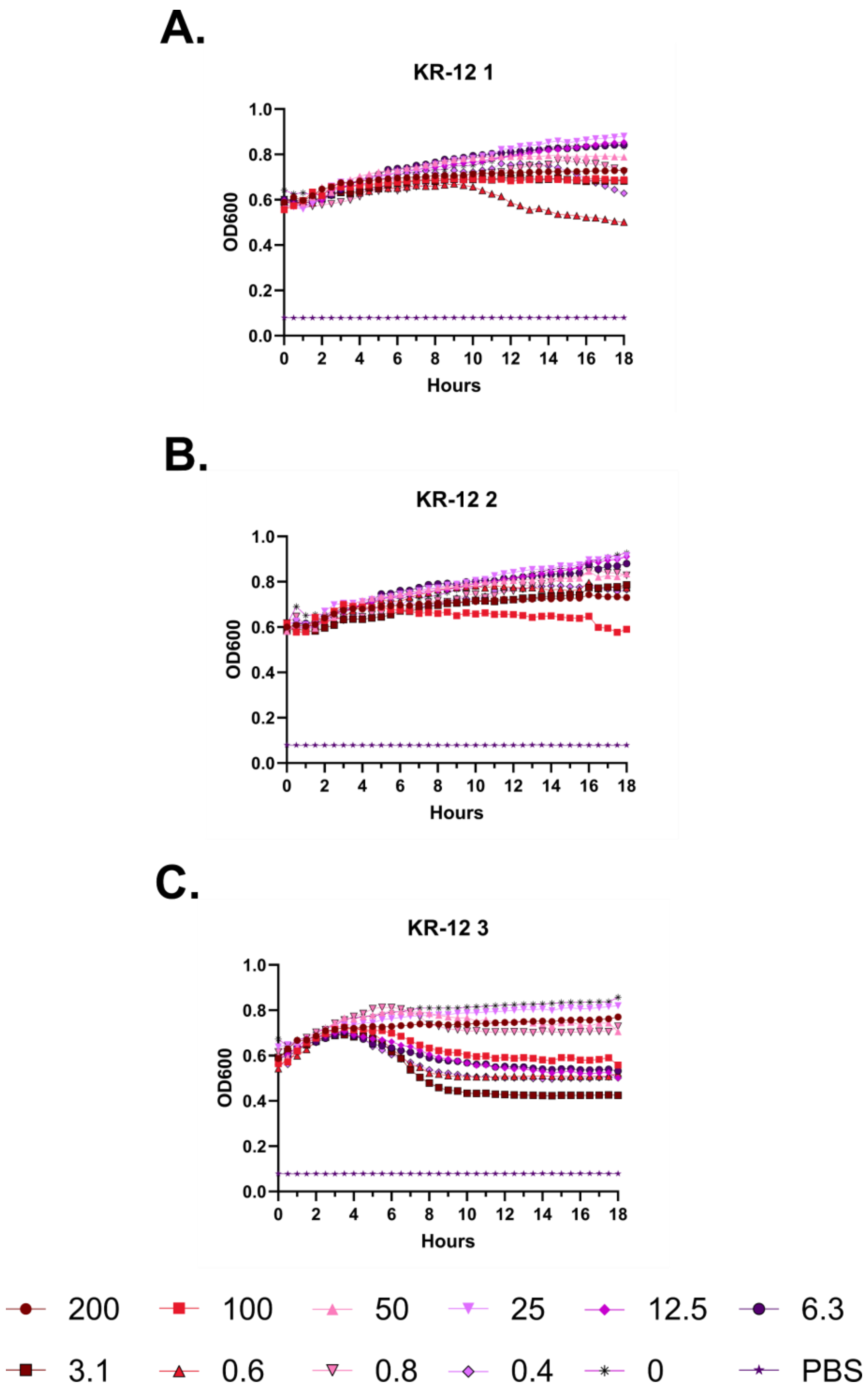
A.
KR-12 1
OD600
Hours
B.
KR-12 2
OD600
Hours
C.
KR-12 3
OD600
Hours
200
100
50
25
12.5
6.3
3.1
0.6
0.8
0.4
0
PBS

**Figure S122 – (#10) KR-12 kinetic hemolysis assay. A.** Defibrinated sheep blood was incubated in the presence of the indicated peptide concentrations at 37 °C with monitoring of OD600 over 18 hours, where decreasing OD600 indicates hemolysis and each panel indicates an independent experiment. Peptide concentrations are in µM.

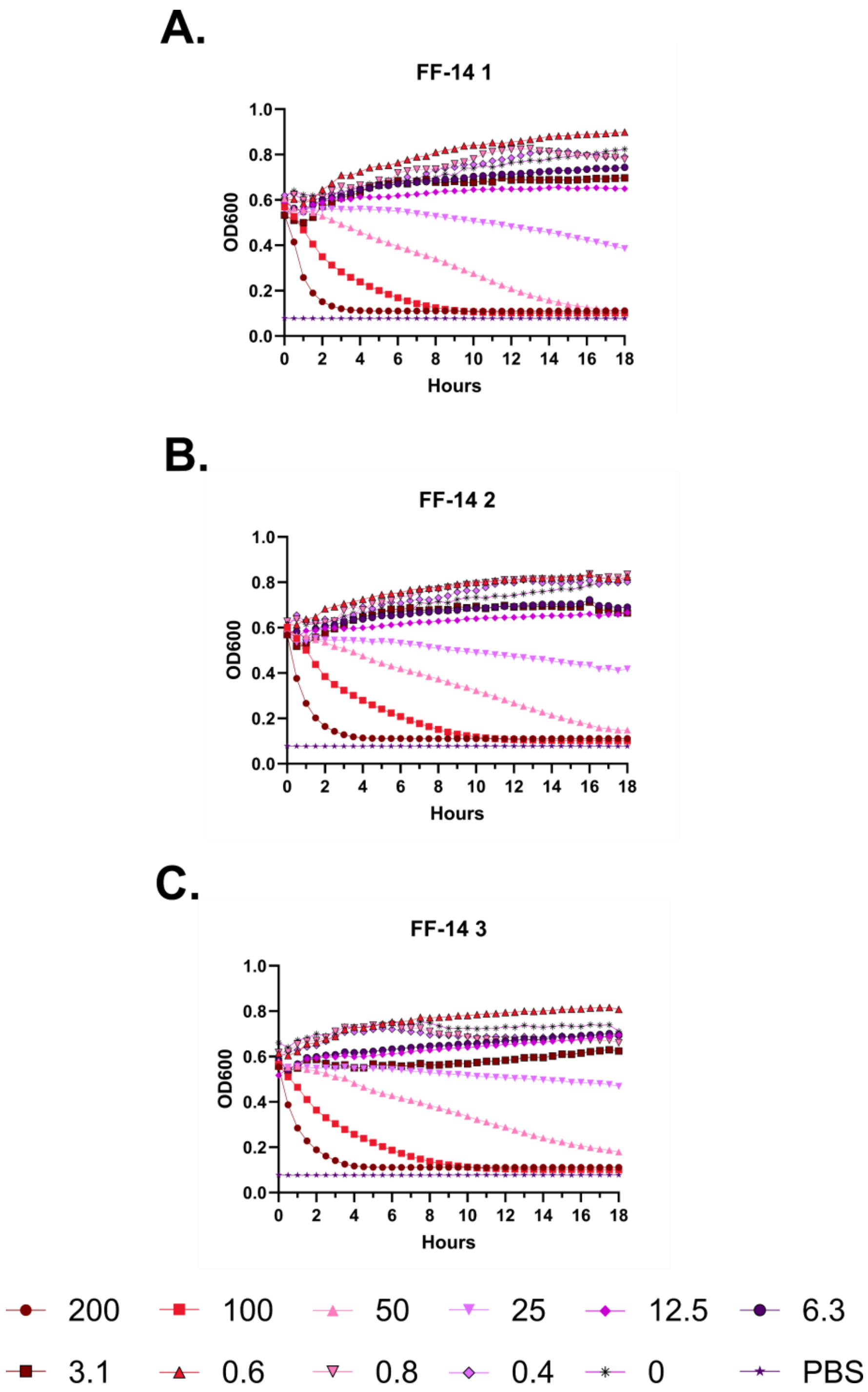
A.
FF-14 1
OD600
Hours
B.
FF-14 2
OD600
Hours
C.
FF-14 3
OD600
Hours
200
100
50
25
12.5
6.3
3.1
0.6
0.8
0.4
0
PBS

**Figure S123 – (#18) FF-14 kinetic hemolysis assay. A.** Defibrinated sheep blood was incubated in the presence of the indicated peptide concentrations at 37 °C with monitoring of OD600 over 18 hours, where decreasing OD600 indicates hemolysis and each panel indicates an independent experiment. Peptide concentrations are in μM.

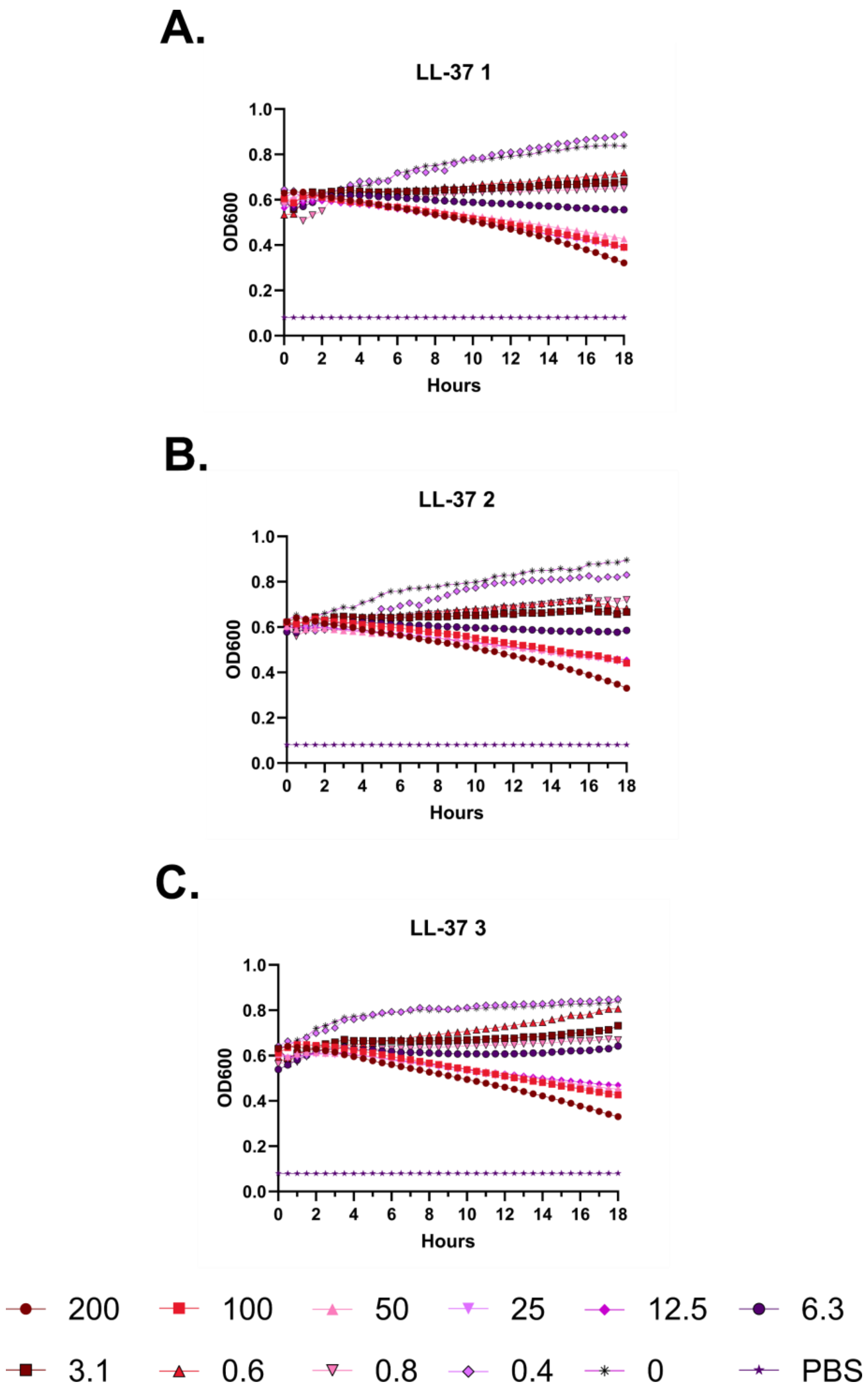
A.
LL-37 1
B.
LL-37 2
C.
LL-37 3
OD600
Hours
1.0
0.8
0.6
0.4
0.2
0.0
0 2 4 6 8 10 12 14 16 18
200
100
50
25
12.5
6.3
3.1
0.6
0.8
0.4
0
PBS

**Figure S124 – (#1) LL-37 kinetic hemolysis assay. A.** Defibrinated sheep blood was incubated in the presence of the indicated peptide concentrations at 37 °C with monitoring of OD600 over 18 hours, where decreasing OD600 indicates hemolysis and each panel indicates an independent experiment. Peptide concentrations are in μM.

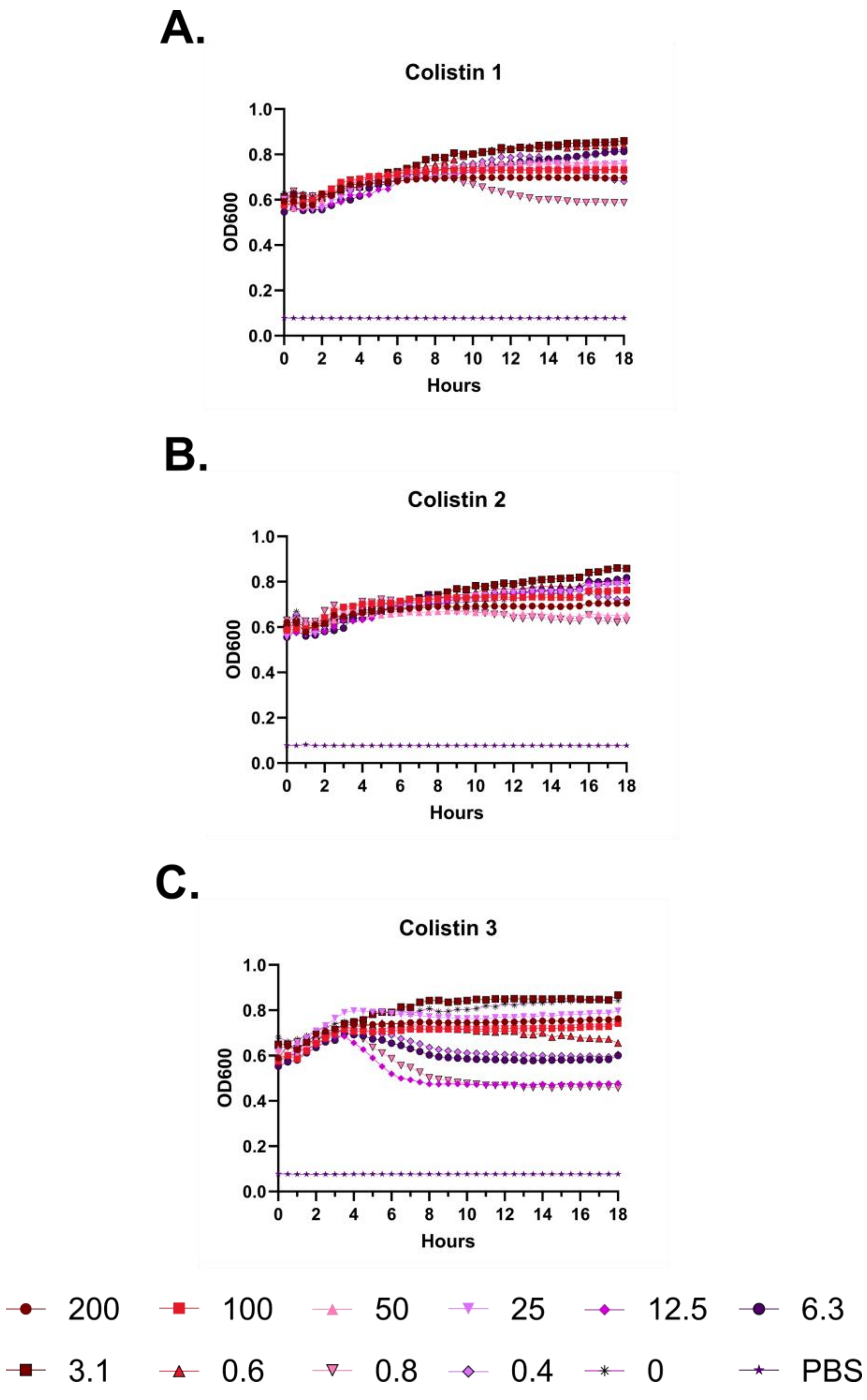
A.
Colistin 1
B.
Colistin 2
C.
Colistin 3
1.0
0.8
0.6
0.4
0.2
0.0
OD600
0 2 4 6 8 10 12 14 16 18
Hours
200
100
50
25
12.5
6.3
3.1
0.6
0.8
0.4
0
PBS

**Figure S125 – Colistin kinetic hemolysis assay. A.** Defibrinated sheep blood was incubated in the presence of the indicated peptide concentrations at 37 °C with monitoring of OD600 over 18 hours, where decreasing OD600 indicates hemolysis and each panel indicates an independent experiment. Peptide concentrations are in μM.

A.

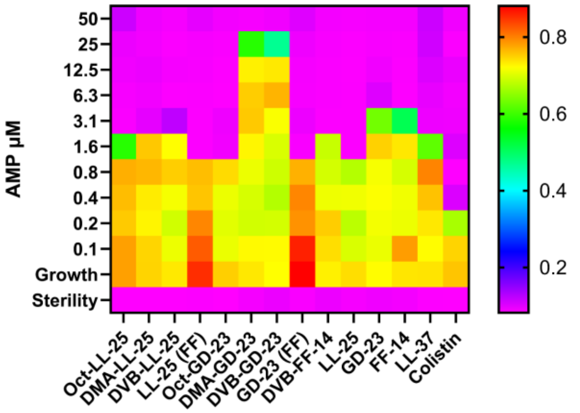
AMP µM
50
25
12.5
6.3
3.1
1.6
0.8
0.4
0.2
0.1
Growth
Sterility
Oct-LL-25
DMA-LL-25
DVB-LL-25
LL-25 (FF)
Oct-GD-23
DMA-GD-23
DVB-GD-23
GD-23 (FF)
DVB-FF-14
LL-25
GD-23
FF-14
LL-37
Colistin
0.8
0.6
0.4
0.2

B.

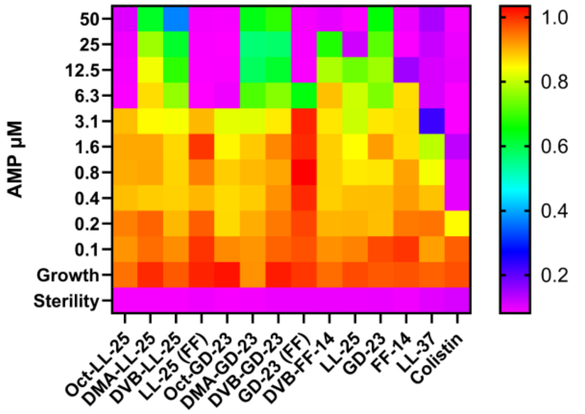
AMP µM
50
25
12.5
6.3
3.1
1.6
0.8
0.4
0.2
0.1
Growth
Sterility
Oct-LL-25
DMA-LL-25
DVB-LL-25
LL-25 (FF)
Oct-GD-23
DMA-GD-23
DVB-GD-23
GD-23 (FF)
DVB-FF-14
LL-25
GD-23
FF-14
LL-37
Colistin
1.0
0.8
0.6
0.4
0.2

C.

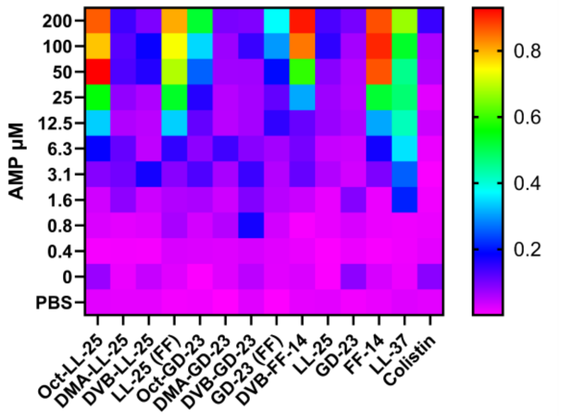
AMP µM
200
100
50
25
12.5
6.3
3.1
1.6
0.8
0.4
0
PBS
Oct-LL-25
DMA-LL-25
DVB-LL-25
LL-25 (FF)
Oct-GD-23
DMA-GD-23
DVB-GD-23
GD-23 (FF)
DVB-FF-14
LL-25
GD-23
FF-14
LL-37
Colistin
0.8
0.6
0.4
0.2

**Figure S126 – Effects of selected N-terminal lipidation moieties on short cathelicidin derivatives.** **A.** Antimicrobial susceptibility testing showing the activity of each derivative as well as a colistin control against *E. coli* (ATCC 25922). In the heatmap plate depiction, cooler colors indicate low or absent bacterial growth as detected by the OD600 measure at 18 hours of incubation at 37 °C. **B.** As in **A.**, but with *P. aeruginosa* (ATCC 27853). **C.** Hemolysis assays in which sheep RBCs in PBS were incubated with the indicated peptide for 18 hours at 37 °C with subsequent quantification of hemoglobin released into the supernatant at 414 nm. Cool colors indicate little or no release of hemoglobin, where all levels are scaled to 100% hemolysis freeze-thaw controls. Levels throughout are the median across three independent experiments.

A.

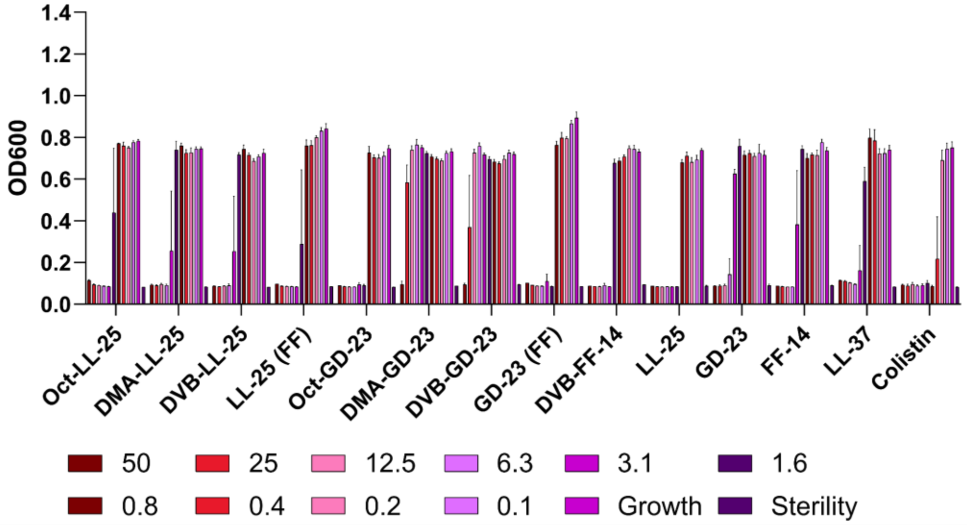

OD600
1.4
1.2
1.0
0.8
0.6
0.4
0.2
0.0
Oct-LL-25
DMA-LL-25
DVB-LL-25
LL-25 (FF)
Oct-GD-23
DMA-GD-23
DVB-GD-23
GD-23 (FF)
DVB-FF-14
LL-25
GD-23
FF-14
LL-37
Colistin
50
25
12.5
6.3
3.1
1.6
0.8
0.4
0.2
0.1
Growth
Sterility


B.

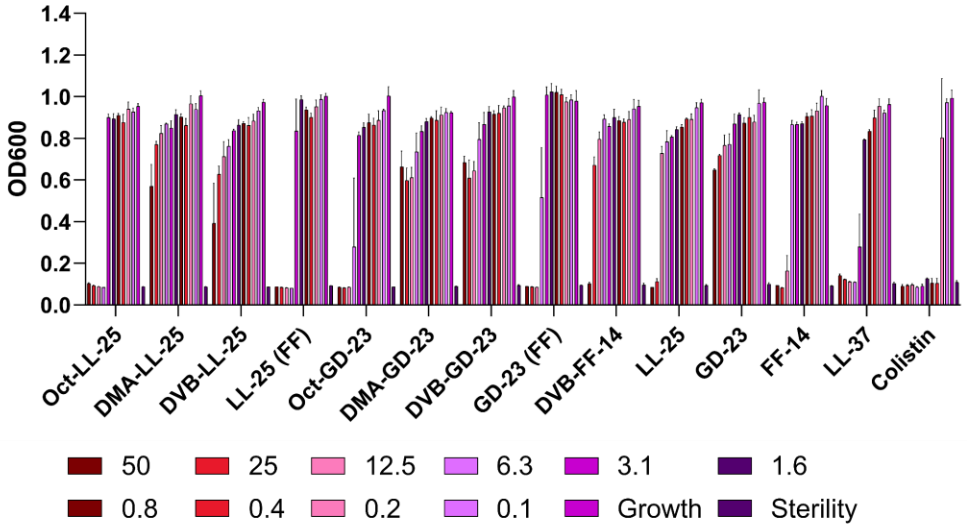

OD600
1.4
1.2
1.0
0.8
0.6
0.4
0.2
0.0
Oct-LL-25
DMA-LL-25
DVB-LL-25
LL-25 (FF)
Oct-GD-23
DMA-GD-23
DVB-GD-23
GD-23 (FF)
DVB-FF-14
LL-25
GD-23
FF-14
LL-37
Colistin
50
25
12.5
6.3
3.1
1.6
0.8
0.4
0.2
0.1
Growth
Sterility


C.

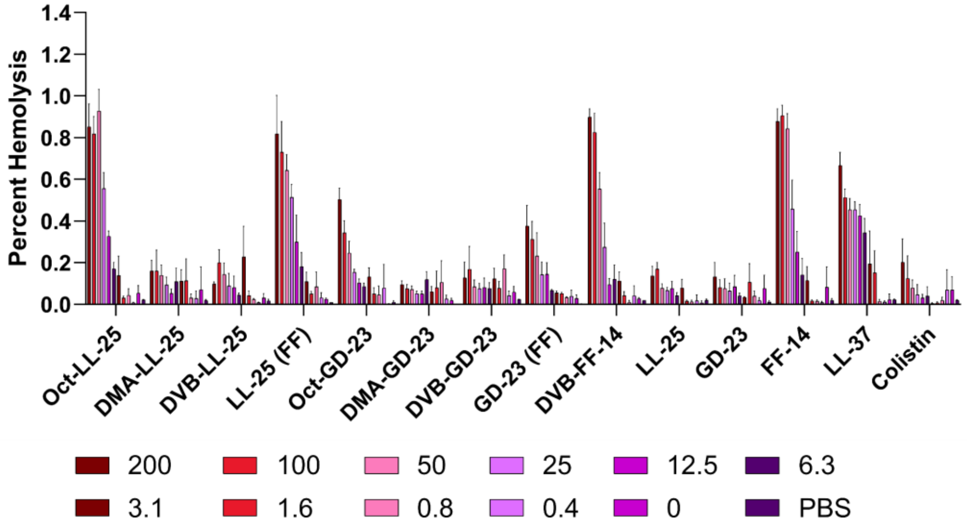

Percent Hemolysis
1.4
1.2
1.0
0.8
0.6
0.4
0.2
0.0
Oct-LL-25
DMA-LL-25
DVB-LL-25
LL-25 (FF)
Oct-GD-23
DMA-GD-23
DVB-GD-23
GD-23 (FF)
DVB-FF-14
LL-25
GD-23
FF-14
LL-37
Colistin
200
100
50
25
12.5
6.3
3.1
1.6
0.8
0.4
0
PBS

**Figure S127 – Histograms corresponding to Figure 6**. **A.** Growth of *E. coli* (ATCC 25922) by OD600 at 18 hours at 37 °C. Error bars indicate the standard deviation of three independent experiments. **B.** As in **A.**, but with *P. aeruginosa* (ATCC 27853). **C.** Endpoint hemolysis assays following incubation with the indicated peptide for 18 hours at 37 °C. Error bars indicate the standard deviation of three independent experiments. All peptide levels are given in μM.

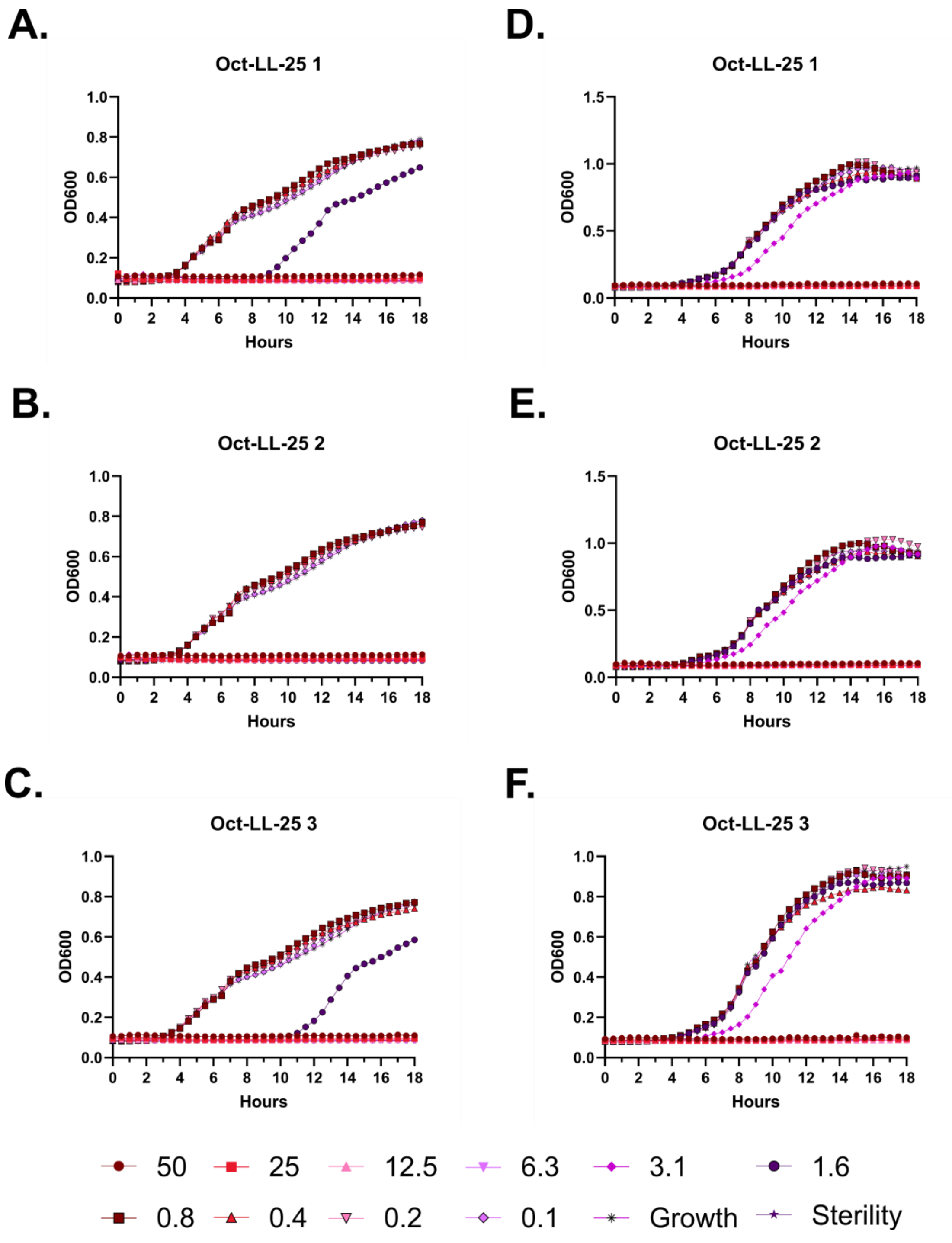
A.
Oct-LL-25 1
OD600
Hours
B.
Oct-LL-25 2
OD600
Hours
C.
Oct-LL-25 3
OD600
Hours
D.
Oct-LL-25 1
OD600
Hours
E.
Oct-LL-25 2
OD600
Hours
F.
Oct-LL-25 3
OD600
Hours
50
25
12.5
6.3
3.1
1.6
0.8
0.4
0.2
0.1
Growth
Sterility

**Figure S128 – Antimicrobial susceptibility curves for (#46) LL-25 N-lipidated with octanoic acid. A-C.** Independent experiments with *E. coli*. **D-F.** Independent experiments with *P. aeruginosa.* Growth over 18 hours at 37 °C with a gradient of peptide concentrations given in μM.

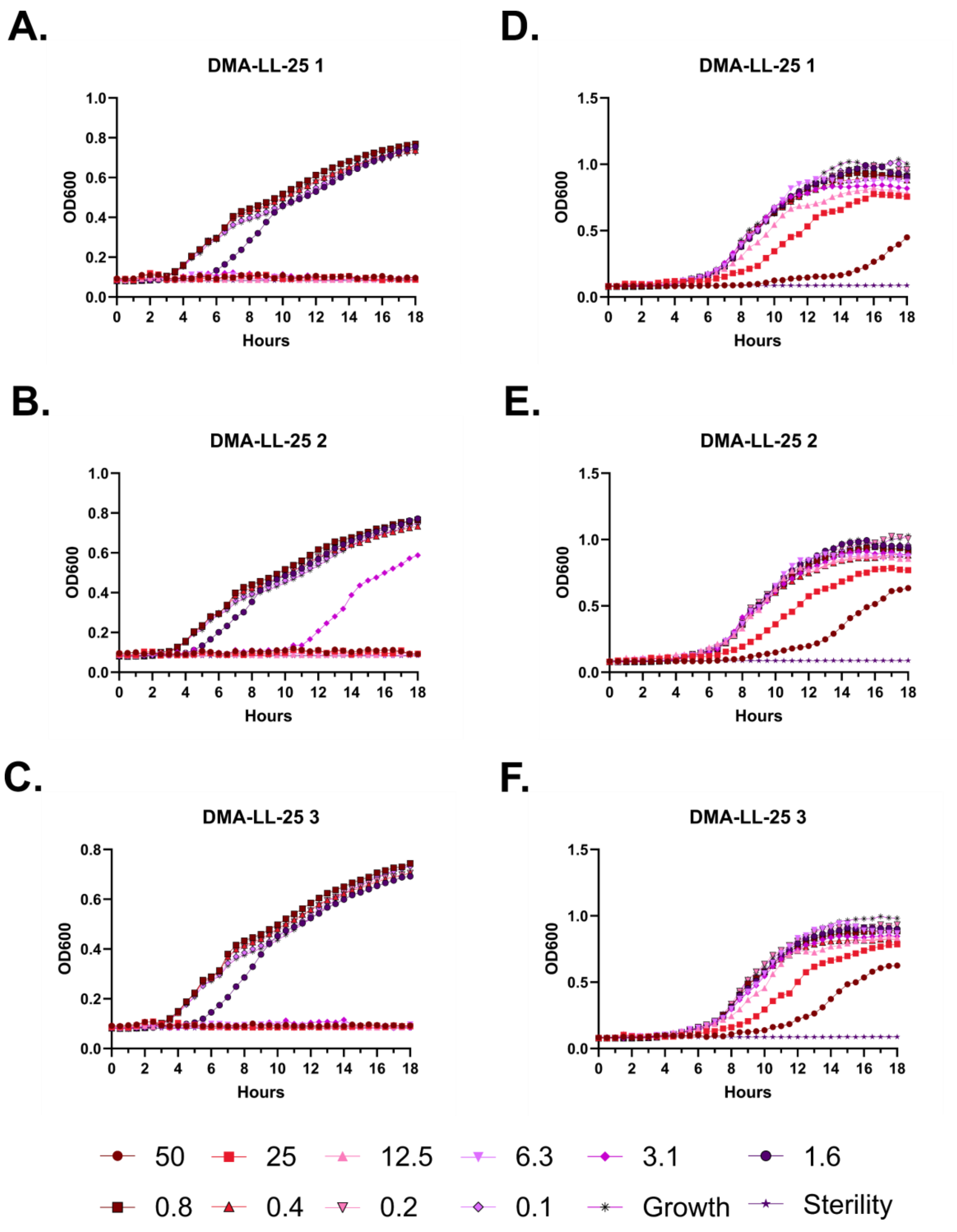
A.
DMA-LL-25 1
B.
DMA-LL-25 2
C.
DMA-LL-25 3
D.
DMA-LL-25 1
E.
DMA-LL-25 2
F.
DMA-LL-25 3
OD600
Hours
50
25
12.5
6.3
3.1
1.6
0.8
0.4
0.2
0.1
Growth
Sterility

**Figure S129 – Antimicrobial susceptibility curves for (#47) LL-25 N-lipidated with 5-dimethylamino pentanoic acid. A-C.** Independent experiments with *E. coli.* **D-F.** Independent experiments with *P. aeruginosa.* Growth over 18 hours at 37 °C with a gradient of peptide concentrations given in μM.

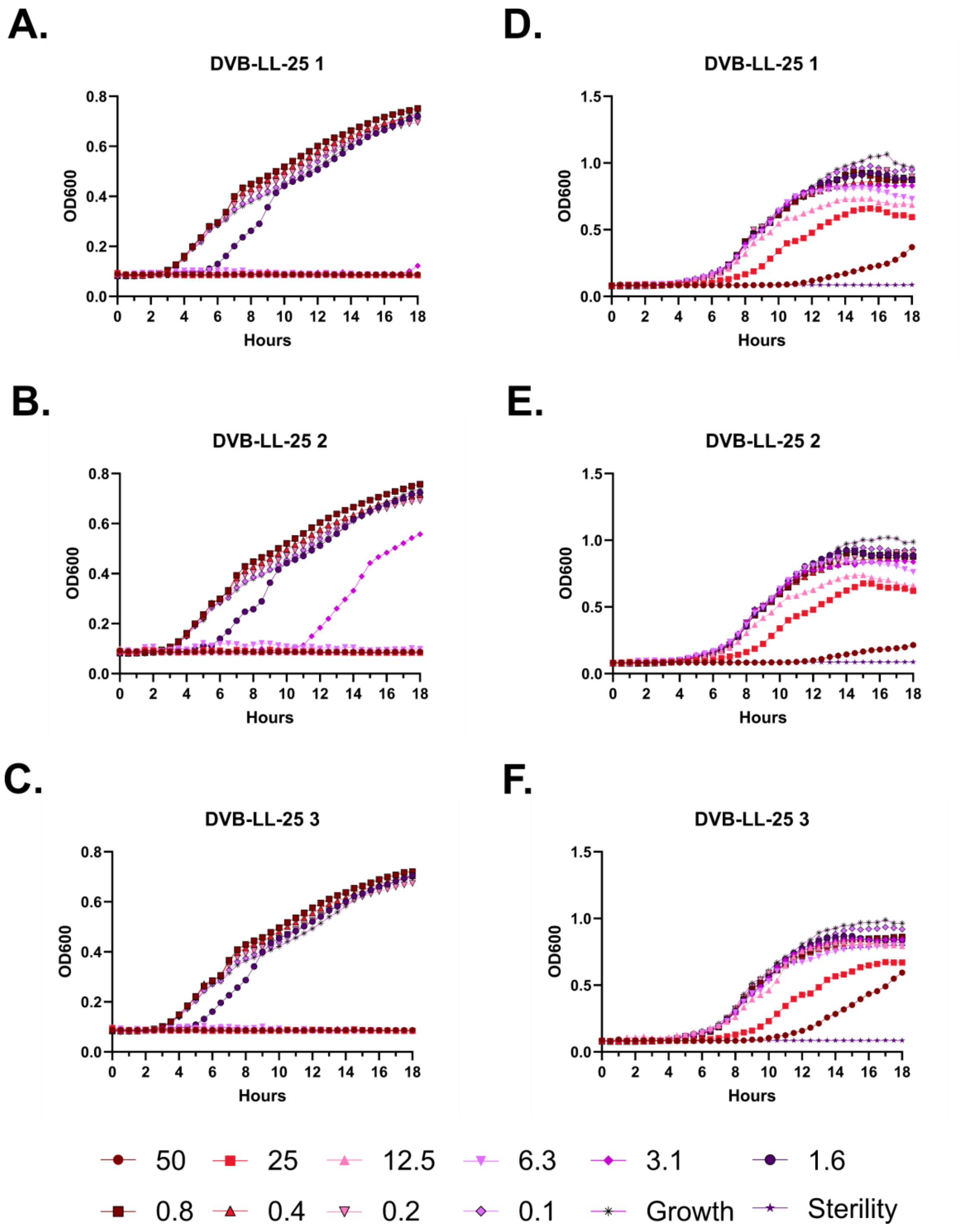

A.
DVB-LL-25 1
B.
DVB-LL-25 2
C.
DVB-LL-25 3
D.
DVB-LL-25 1
E.
DVB-LL-25 2
F.
DVB-LL-25 3
OD600
Hours
50
25
12.5
6.3
3.1
1.6
0.8
0.4
0.2
0.1
Growth
Sterility

**Figure S130 – Antimicrobial susceptibility curves for (#48) LL-25 N-lipidated with δ-valerobetaine. A-C.** Independent experiments with *E. coli*. **D-F.** Independent experiments with *P. aeruginosa.* Growth over 18 hours at 37 °C with a gradient of peptide concentrations given in μM.

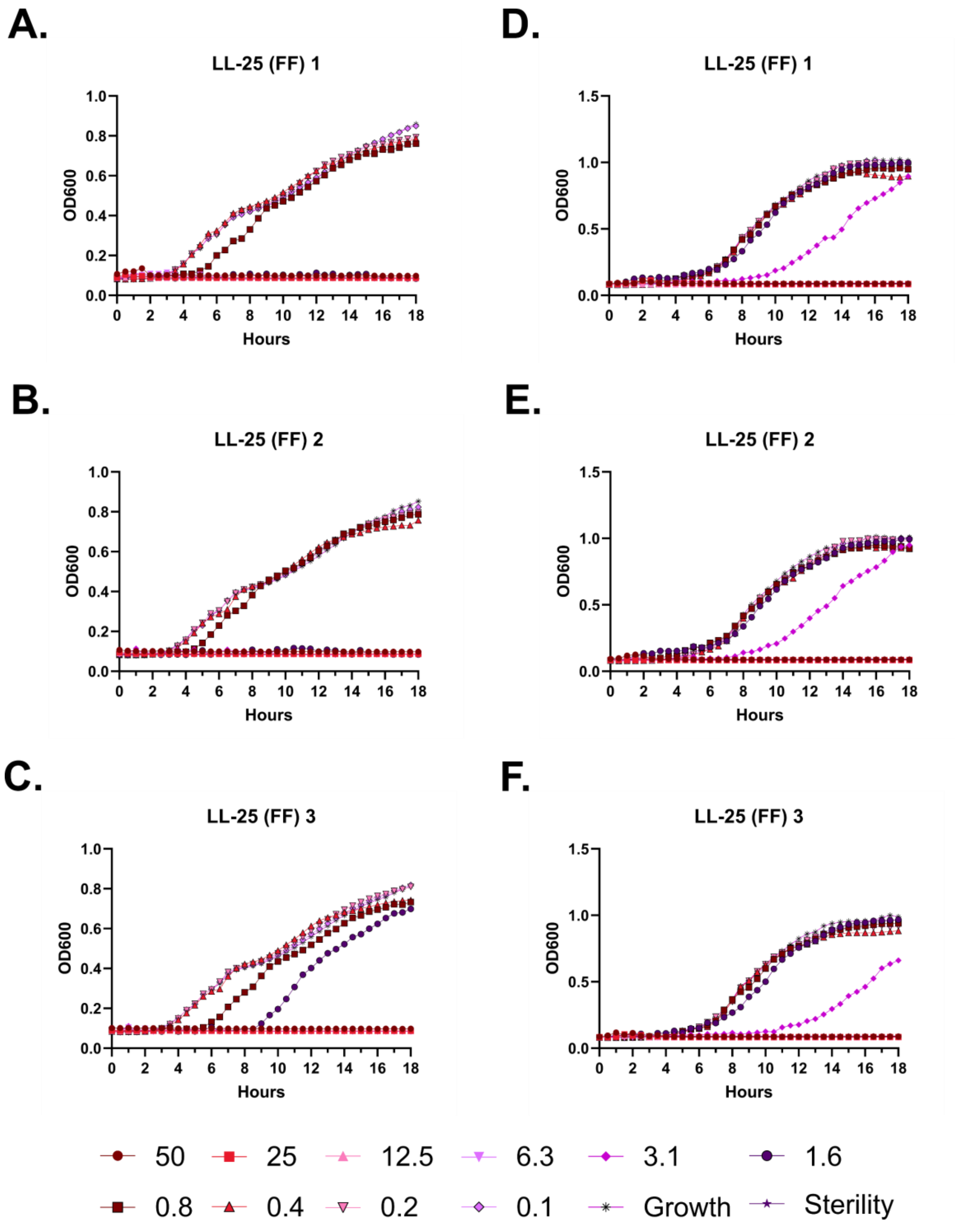
A.
LL-25 (FF) 1
OD600
Hours
B.
LL-25 (FF) 2
OD600
Hours
C.
LL-25 (FF) 3
OD600
Hours
D.
LL-25 (FF) 1
OD600
Hours
E.
LL-25 (FF) 2
OD600
Hours
F.
LL-25 (FF) 3
OD600
Hours
50
25
12.5
6.3
3.1
1.6
0.8
0.4
0.2
0.1
Growth
Sterility

**Figure S131 – Antimicrobial susceptibility curves for (#49) LL-25 (FF). A-C.** Independent experiments with *E. coli*. **D-F.** Independent experiments with *P. aeruginosa.* Growth over 18 hours at 37 °C with a gradient of peptide concentrations given in μM.

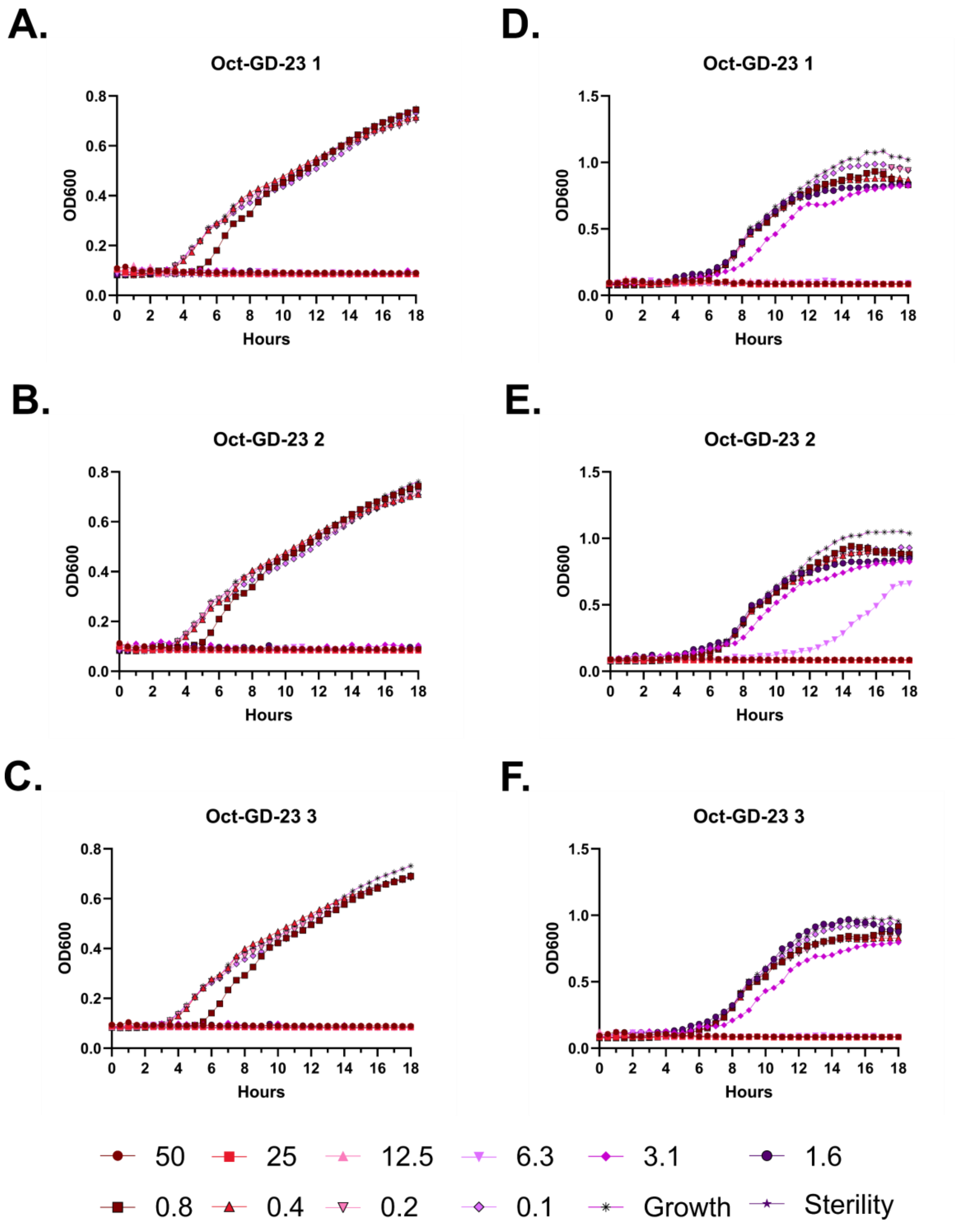
A.
Oct-GD-23 1
B.
Oct-GD-23 2
C.
Oct-GD-23 3
D.
Oct-GD-23 1
E.
Oct-GD-23 2
F.
Oct-GD-23 3
OD600
Hours
50
25
12.5
6.3
3.1
1.6
0.8
0.4
0.2
0.1
Growth
Sterility

**Figure S132 – Antimicrobial susceptibility curves for (#50) GD-23 N-lipidated with octanoic acid. A-C.** Independent experiments with *E. coli*. **D-F.** Independent experiments with *P. aeruginosa.* Growth over 18 hours at 37 °C with a gradient of peptide concentrations given in μM.

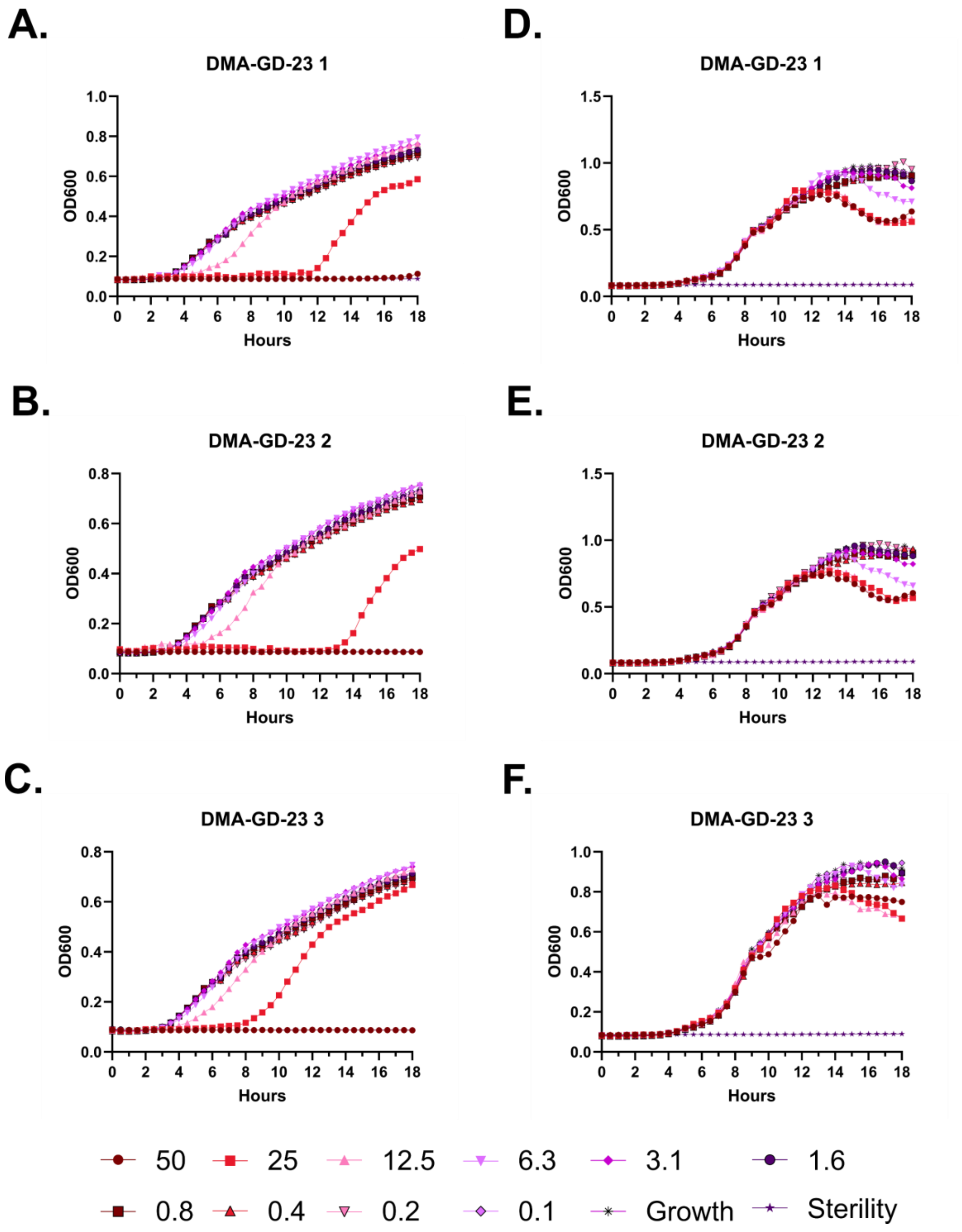
A.
DMA-GD-23 1
OD600
Hours
B.
DMA-GD-23 2
OD600
Hours
C.
DMA-GD-23 3
OD600
Hours
D.
DMA-GD-23 1
OD600
Hours
E.
DMA-GD-23 2
OD600
Hours
F.
DMA-GD-23 3
OD600
Hours
50
25
12.5
6.3
3.1
1.6
0.8
0.4
0.2
0.1
Growth
Sterility

**Figure S133 – Antimicrobial susceptibility curves for (#51) GD-23 N-lipidated with 5-dimethylamino pentanoic acid. A-C.** Independent experiments with *E. coli.* **D-F.** Independent experiments with *P. aeruginosa.* Growth over 18 hours at 37 °C with a gradient of peptide concentrations given in μM.

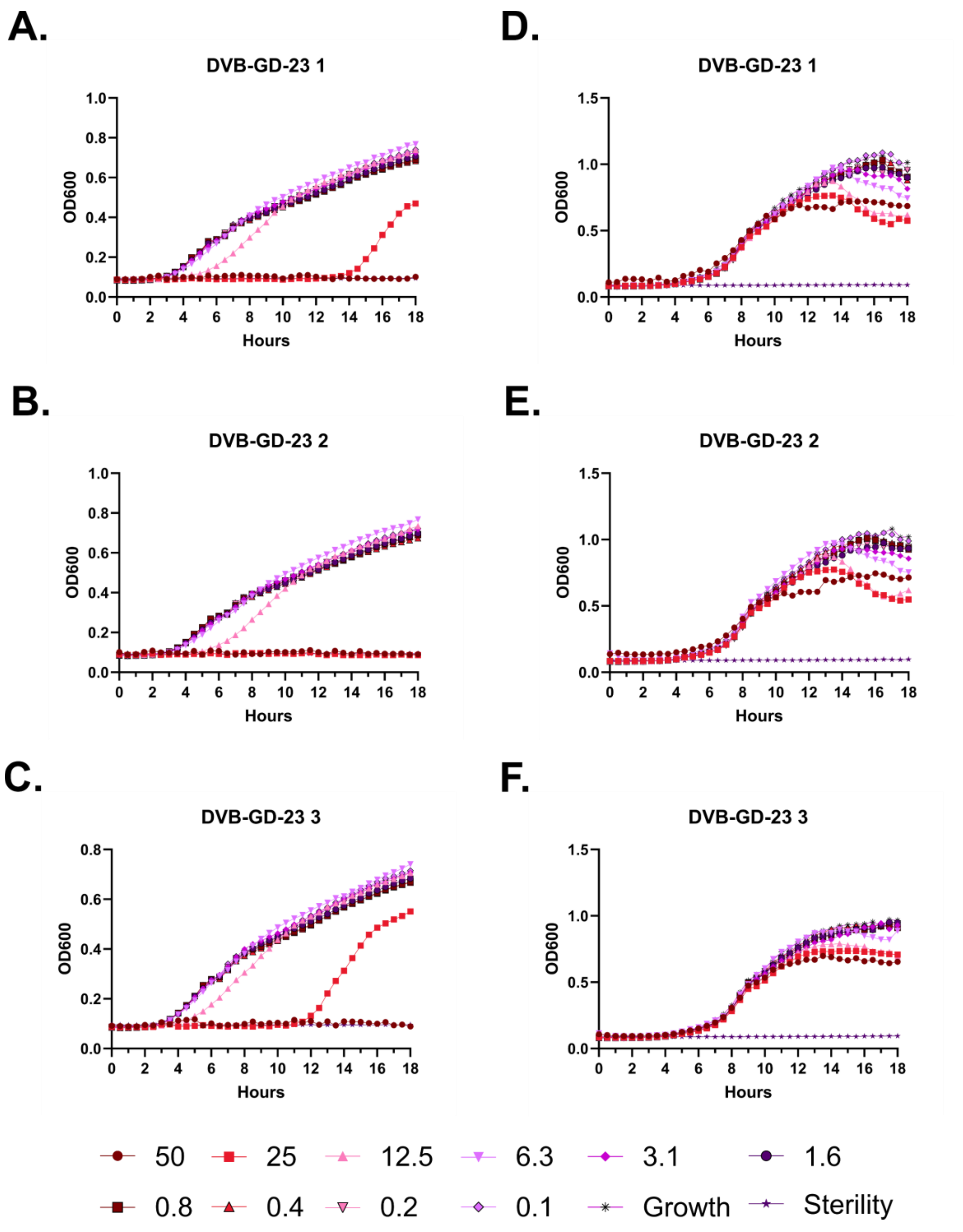
A.
DVB-GD-23 1
B.
DVB-GD-23 2
C.
DVB-GD-23 3
D.
DVB-GD-23 1
E.
DVB-GD-23 2
F.
DVB-GD-23 3
OD600
Hours
50
25
12.5
6.3
3.1
1.6
0.8
0.4
0.2
0.1
Growth
Sterility

**Figure S134 – Antimicrobial susceptibility curves for (#52) GD-23 N-lipidated with δ-valerobetaine. A-C.** Independent experiments with *E. coli*. **D-F.** Independent experiments with *P. aeruginosa.* Growth over 18 hours at 37 °C with a gradient of peptide concentrations given in μM.

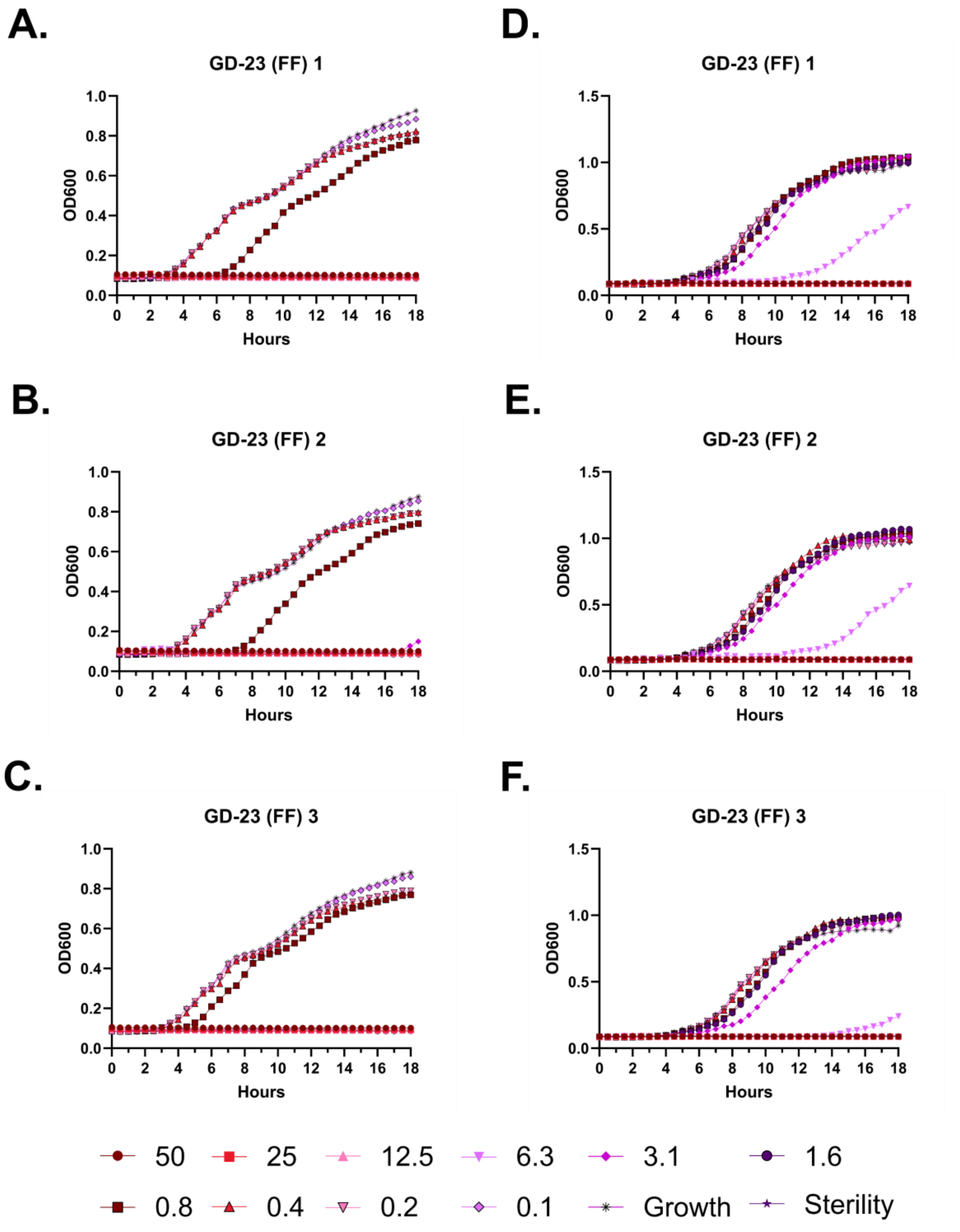

A.
GD-23 (FF) 1
B.
GD-23 (FF) 2
C.
GD-23 (FF) 3
D.
GD-23 (FF) 1
E.
GD-23 (FF) 2
F.
GD-23 (FF) 3
OD600
Hours
50
25
12.5
6.3
3.1
1.6
0.8
0.4
0.2
0.1
Growth
Sterility

**Figure S135 – Antimicrobial susceptibility curves for (#53) GD-23 (FF). A-C.** Independent experiments with *E. coli*. **D-F.** Independent experiments with *P. aeruginosa.* Growth over 18 hours at 37 °C with a gradient of peptide concentrations given in μM.

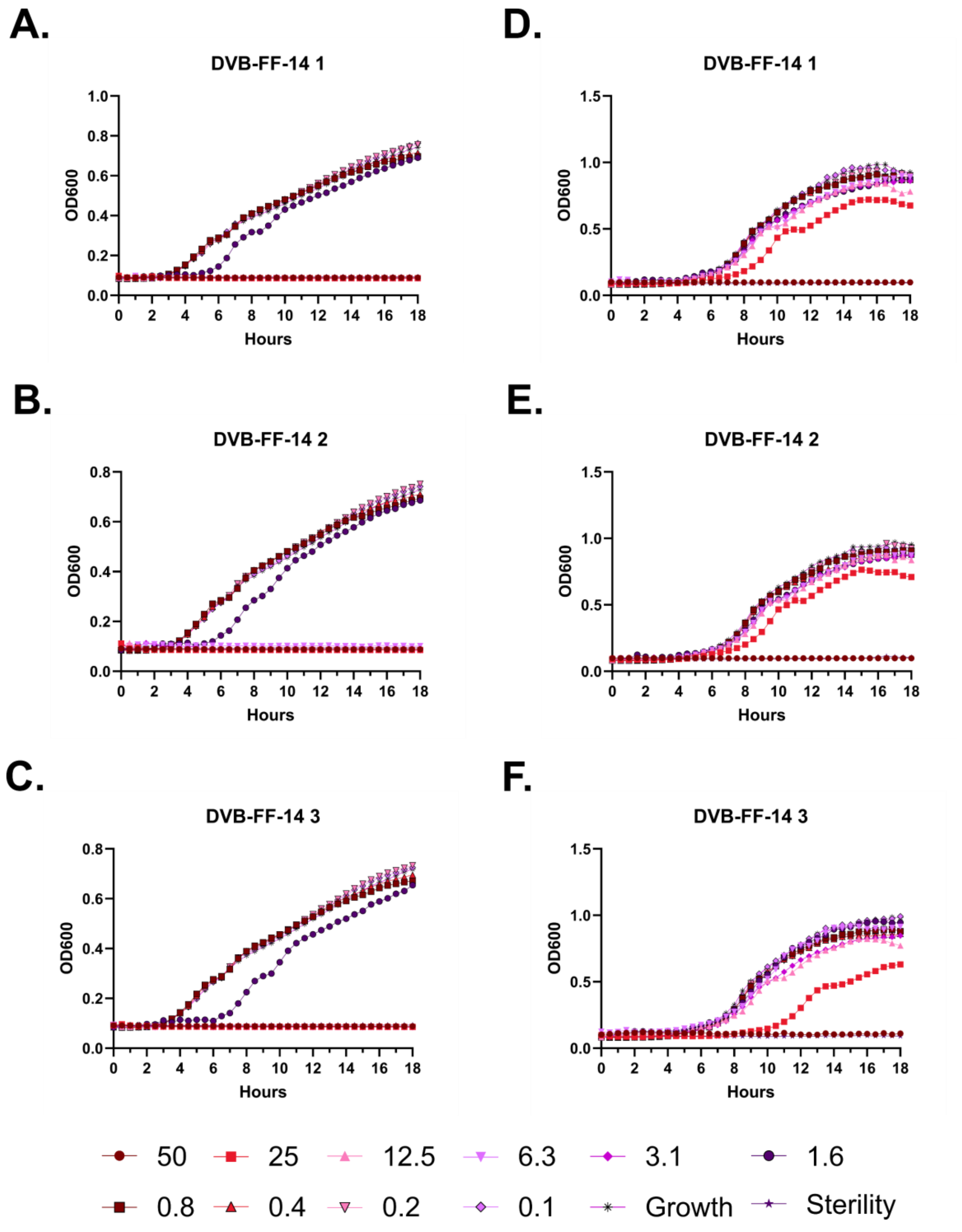
A.
DVB-FF-14 1
B.
DVB-FF-14 2
C.
DVB-FF-14 3
D.
DVB-FF-14 1
E.
DVB-FF-14 2
F.
DVB-FF-14 3
OD600
Hours
50
25
12.5
6.3
3.1
1.6
0.8
0.4
0.2
0.1
Growth
Sterility

**Figure S136 – Antimicrobial susceptibility curves for (#54) FF-14 N-lipidated with δ-valerobetaine. A-C.** Independent experiments with *E. coli*. **D-F.** Independent experiments with *P. aeruginosa.* Growth over 18 hours at 37 °C with a gradient of peptide concentrations given in μM.

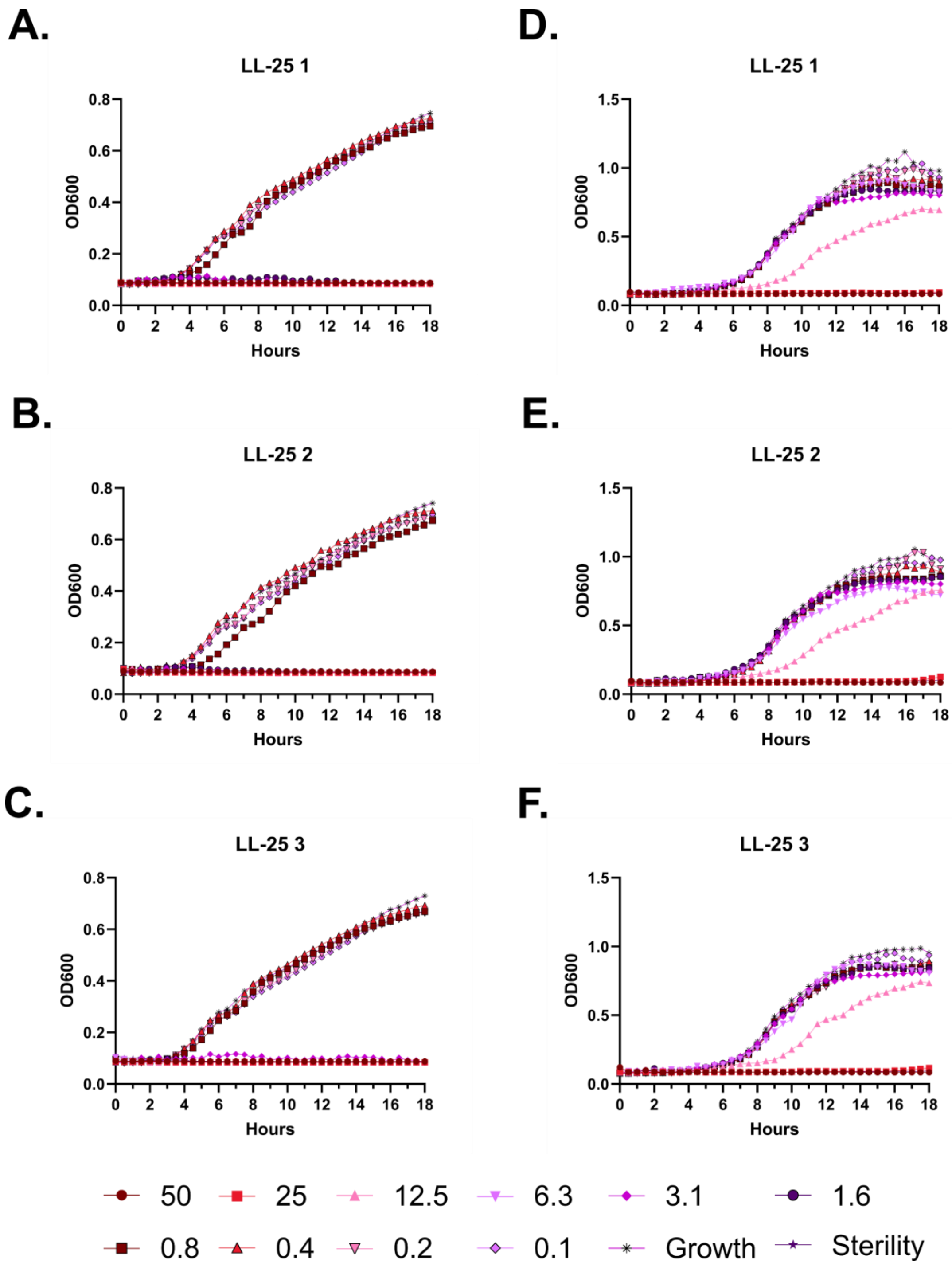
A.
LL-25 1
OD600
Hours
B.
LL-25 2
OD600
Hours
C.
LL-25 3
OD600
Hours
D.
LL-25 1
OD600
Hours
E.
LL-25 2
OD600
Hours
F.
LL-25 3
OD600
Hours
50
25
12.5
6.3
3.1
1.6
0.8
0.4
0.2
0.1
Growth
Sterility

**Figure S137 – Antimicrobial susceptibility curves for (#2) LL-25. A-C.** Independent experiments with *E. coli*. **D-F.** Independent experiments with *P. aeruginosa.* Growth over 18 hours at 37 °C with a gradient of peptide concentrations given in μM.

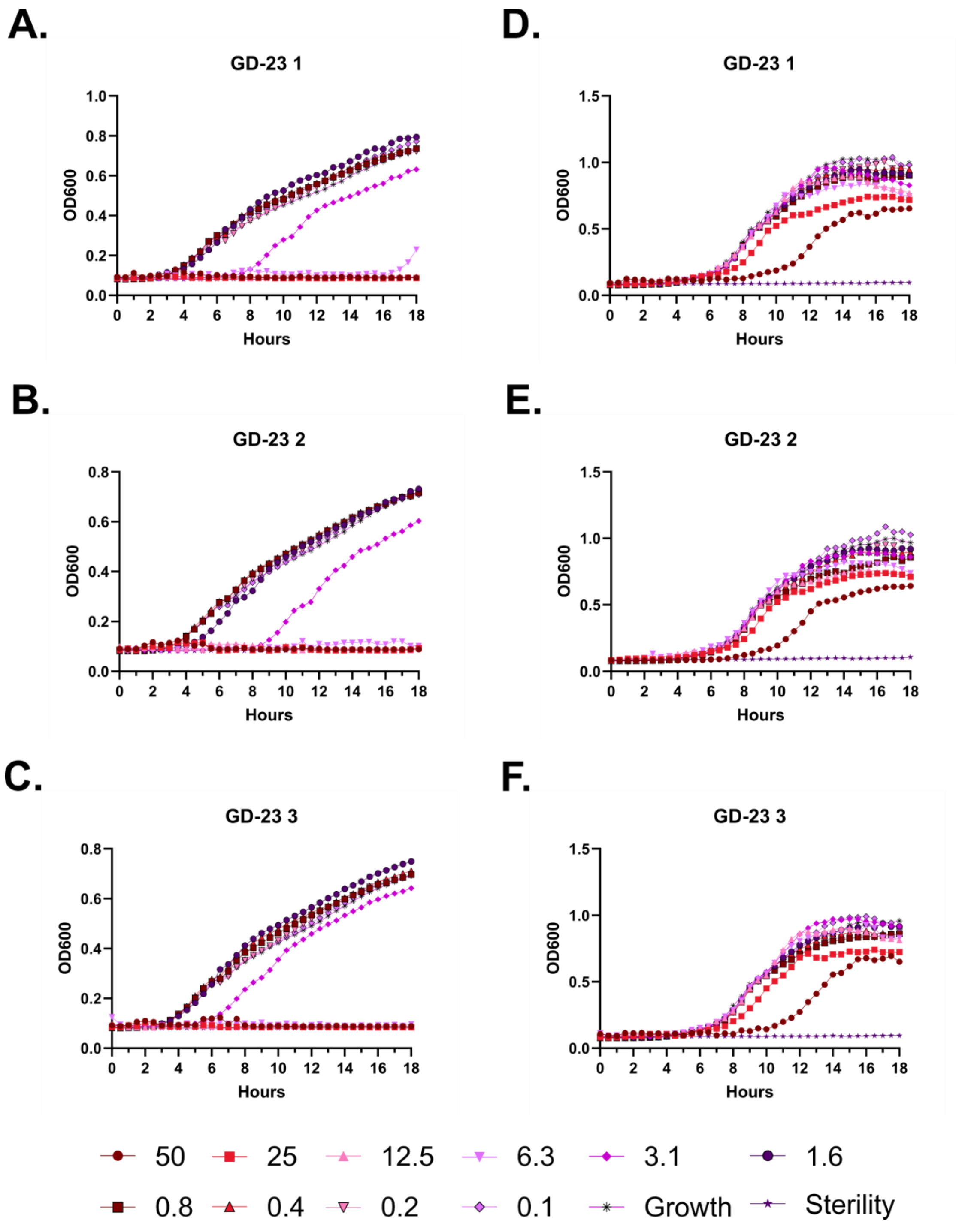
A.
GD-23 1
OD600
Hours
D.
GD-23 1
OD600
Hours
B.
GD-23 2
OD600
Hours
E.
GD-23 2
OD600
Hours
C.
GD-23 3
OD600
Hours
F.
GD-23 3
OD600
Hours
50
25
12.5
6.3
3.1
1.6
0.8
0.4
0.2
0.1
Growth
Sterility

**Figure S138 – Antimicrobial susceptibility curves for (#4) GD-23. A-C.** Independent experiments with *E. coli*. **D-F.** Independent experiments with *P. aeruginosa.* Growth over 18 hours at 37 °C with a gradient of peptide concentrations given in μM.

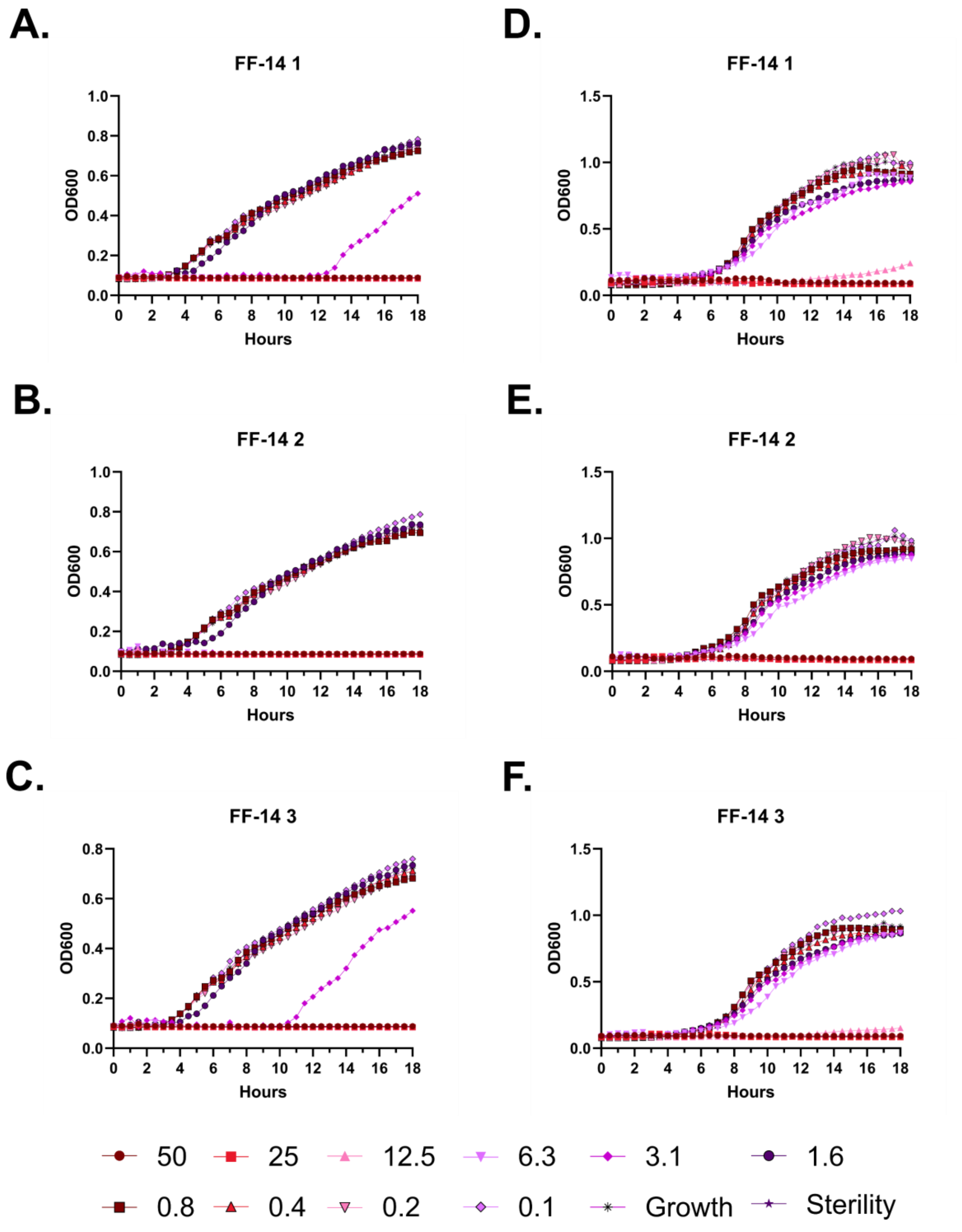
A.
FF-14 1
OD600
Hours
B.
FF-14 2
OD600
Hours
C.
FF-14 3
OD600
Hours
D.
FF-14 1
OD600
Hours
E.
FF-14 2
OD600
Hours
F.
FF-14 3
OD600
Hours
50
25
12.5
6.3
3.1
1.6
0.8
0.4
0.2
0.1
Growth
Sterility

**Figure S139 – Antimicrobial susceptibility curves for (#18) FF-14. A-C.** Independent experiments with *E. coli*. **D-F.** Independent experiments with *P. aeruginosa.* Growth over 18 hours at 37 °C with a gradient of peptide concentrations given in μM.

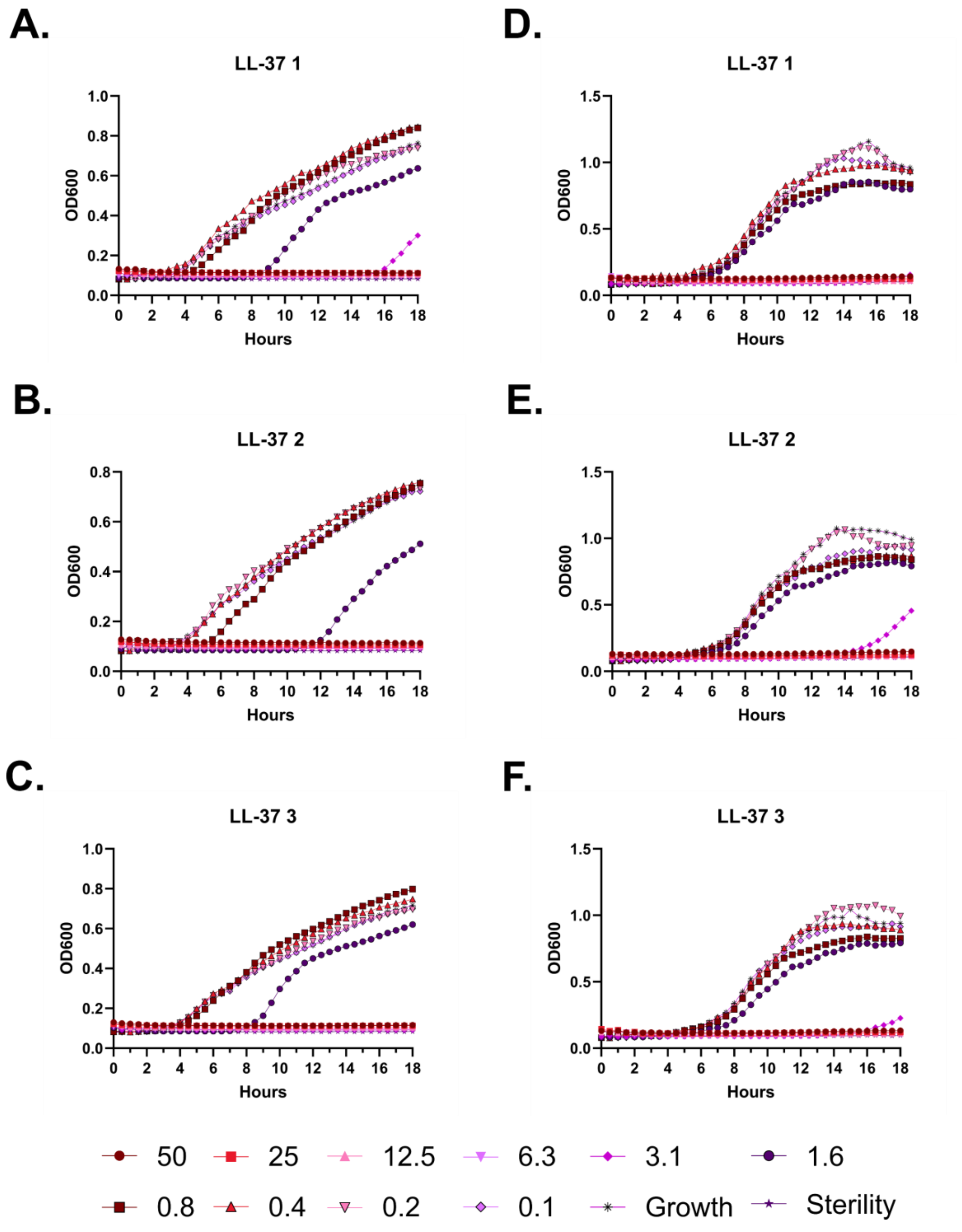
A.
LL-37 1
D.
LL-37 1
B.
LL-37 2
E.
LL-37 2
C.
LL-37 3
F.
LL-37 3
OD600
Hours
50
25
12.5
6.3
3.1
1.6
0.8
0.4
0.2
0.1
Growth
Sterility

**Figure S140 – Antimicrobial susceptibility curves for (#1) LL-37. A-C.** Independent experiments with *E. coli*. **D-F.** Independent experiments with *P. aeruginosa.* Growth over 18 hours at 37 °C with a gradient of peptide concentrations given in μM.

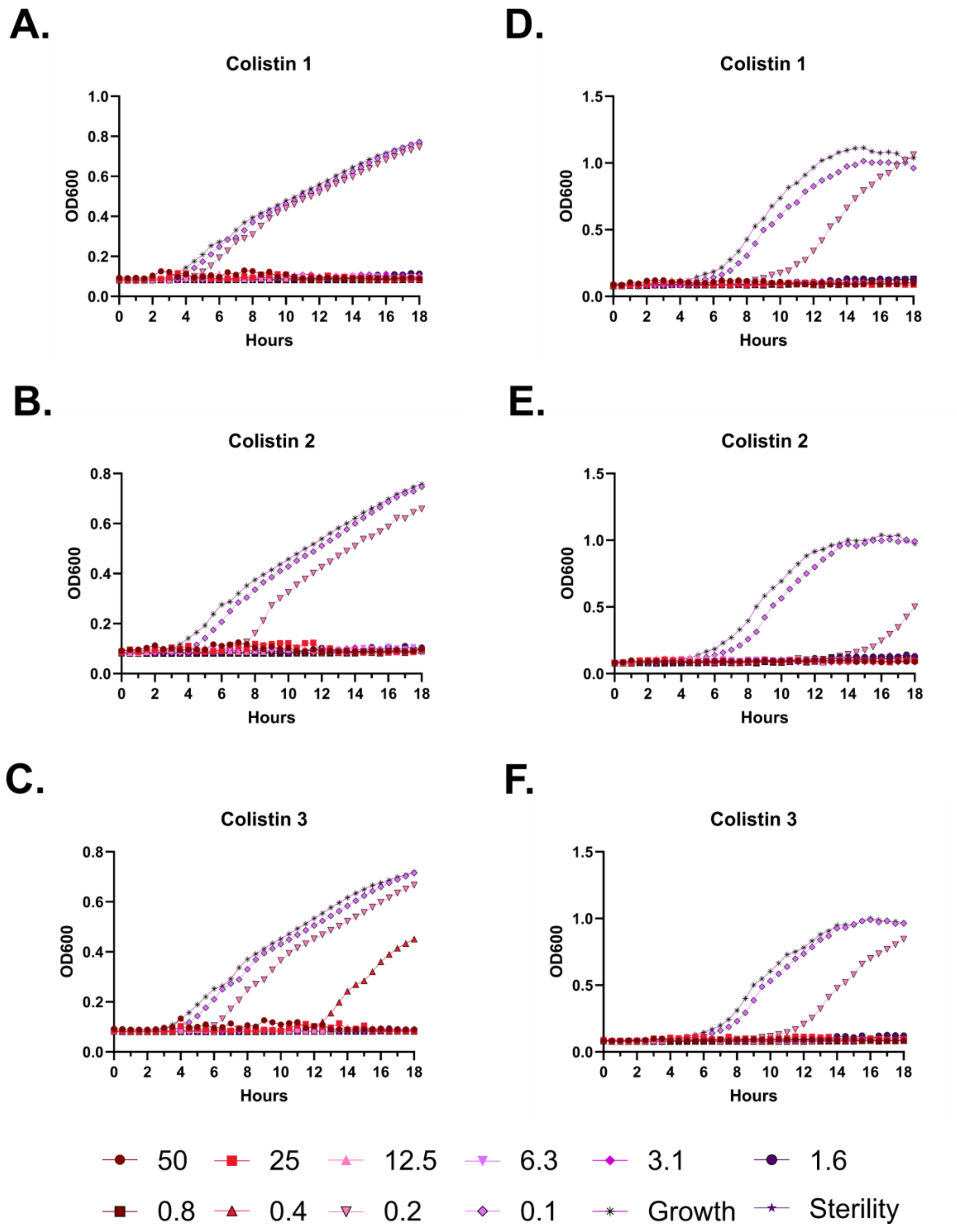
A.
Colistin 1
D.
Colistin 1
B.
Colistin 2
E.
Colistin 2
C.
Colistin 3
F.
Colistin 3
OD600
Hours
50
25
12.5
6.3
3.1
1.6
0.8
0.4
0.2
0.1
Growth
Sterility

**Figure S141 – Antimicrobial susceptibility curves for colistin. A-C.** Independent experiments with *E. coli*. **D-F.** Independent experiments with *P. aeruginosa.* Growth over 18 hours at 37 °C with a gradient of peptide concentrations given in μM.

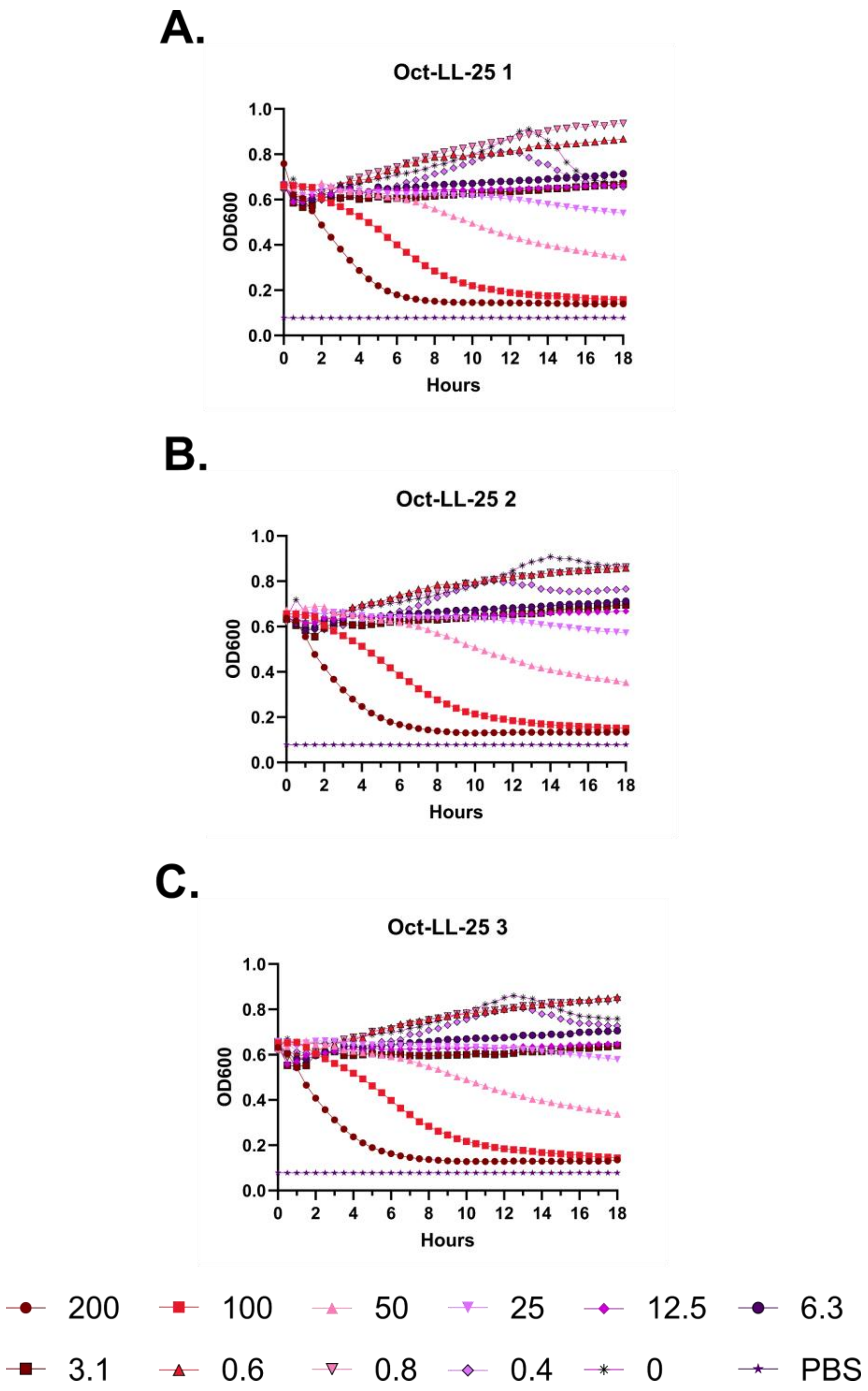
A.
Oct-LL-25 1
OD600
Hours
B.
Oct-LL-25 2
OD600
Hours
C.
Oct-LL-25 3
OD600
Hours
200
100
50
25
12.5
6.3
3.1
0.6
0.8
0.4
0
PBS

**Figure S142 – (#46) Oct-LL-25 kinetic hemolysis assay. A.** Defibrinated sheep blood was incubated in the presence of the indicated peptide concentrations at 37 °C with monitoring of OD600 over 18 hours, where decreasing OD600 indicates hemolysis and each panel indicates an independent experiment. Peptide concentrations are in µM.

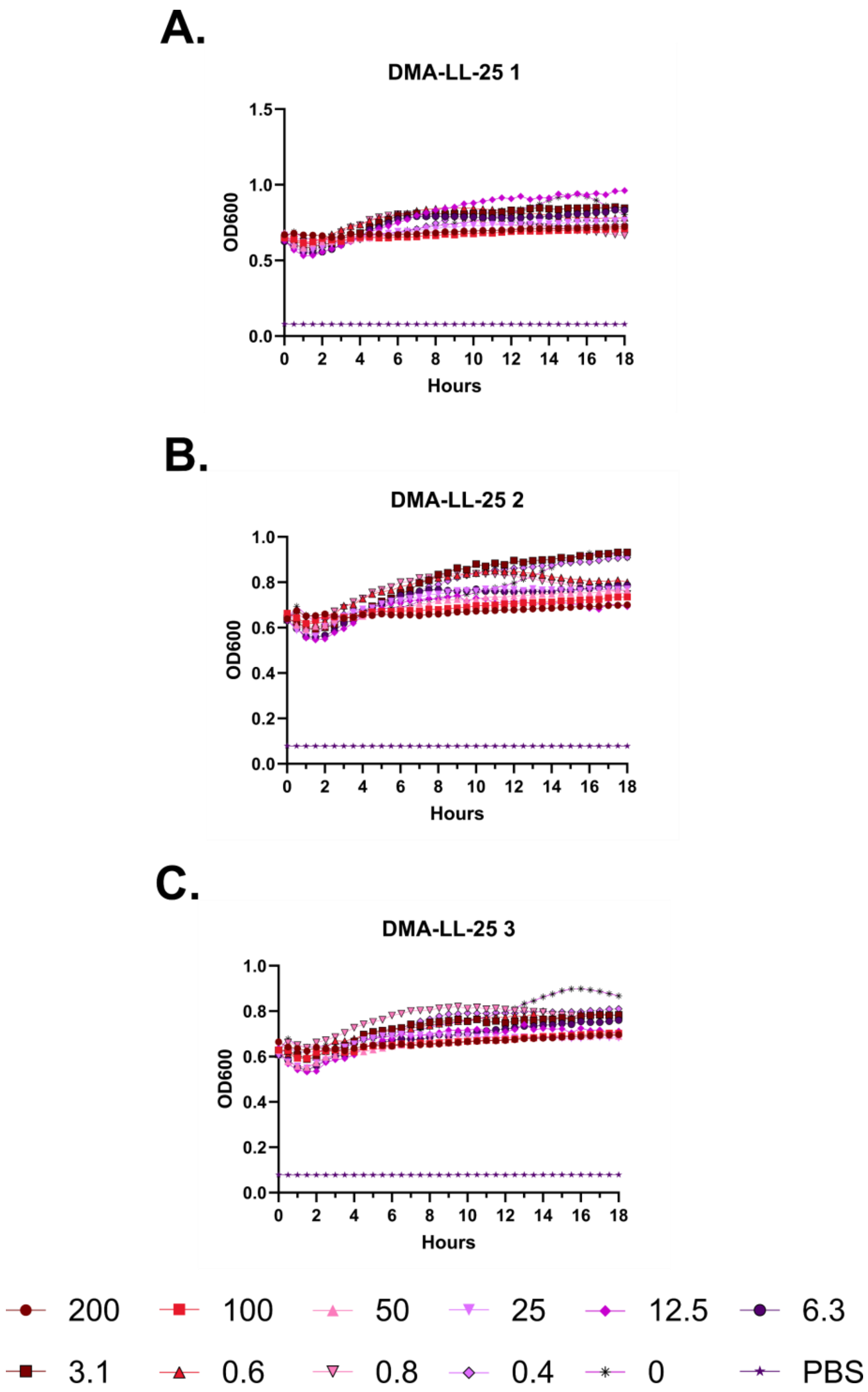
A.
DMA-LL-25 1
OD600
Hours
B.
DMA-LL-25 2
OD600
Hours
C.
DMA-LL-25 3
OD600
Hours
200
100
50
25
12.5
6.3
3.1
0.6
0.8
0.4
0
PBS

**Figure S143 – (#47) DMA-LL-25 kinetic hemolysis assay. A.** Defibrinated sheep blood was incubated in the presence of the indicated peptide concentrations at 37 °C with monitoring of OD600 over 18 hours, where decreasing OD600 indicates hemolysis and each panel indicates an independent experiment. Peptide concentrations are in μM.

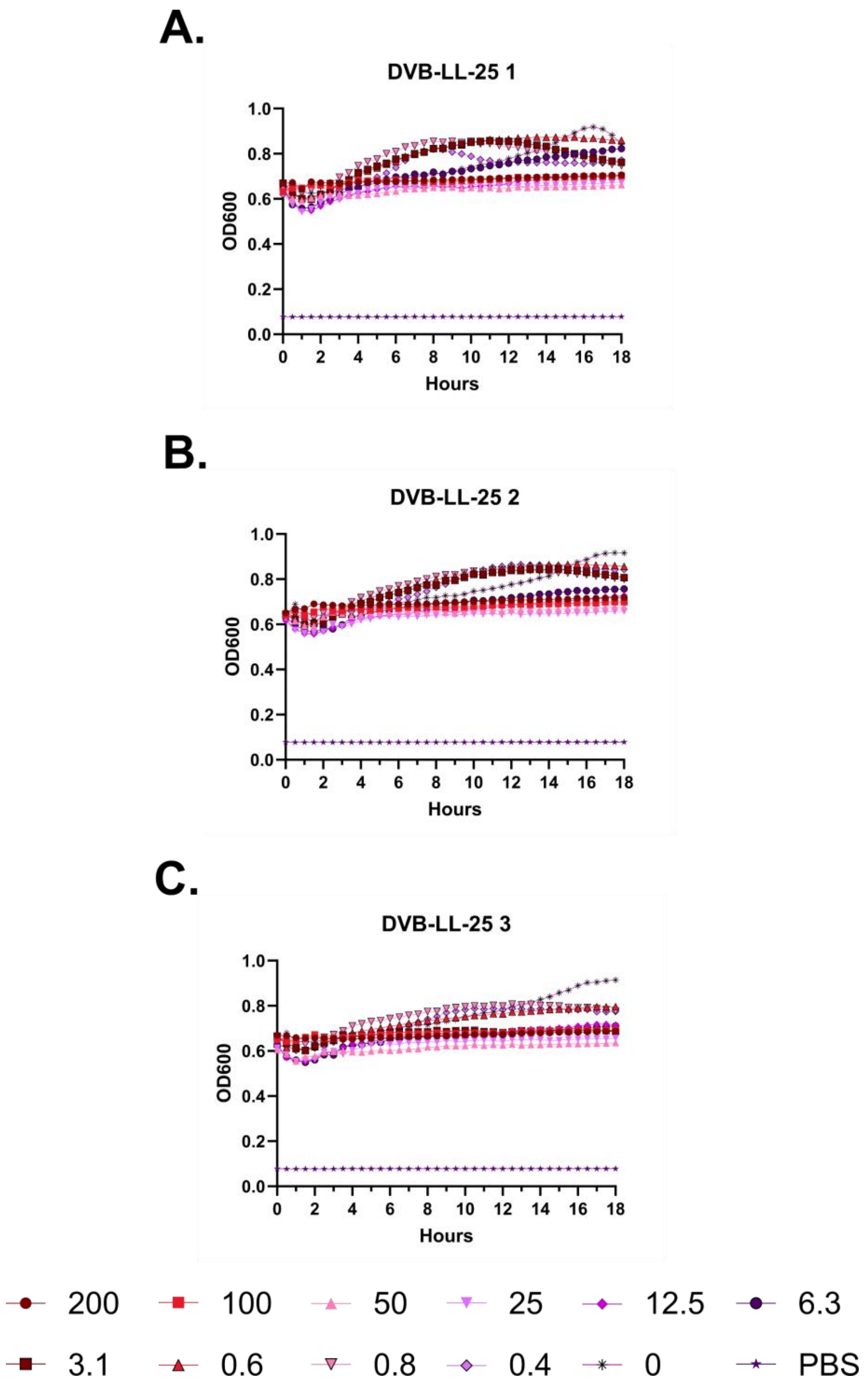
A.
DVB-LL-25 1
OD600
1.0
0.8
0.6
0.4
0.2
0.0
0 2 4 6 8 10 12 14 16 18
Hours
B.
DVB-LL-25 2
OD600
1.0
0.8
0.6
0.4
0.2
0.0
0 2 4 6 8 10 12 14 16 18
Hours
C.
DVB-LL-25 3
OD600
1.0
0.8
0.6
0.4
0.2
0.0
0 2 4 6 8 10 12 14 16 18
Hours
200
100
50
25
12.5
6.3
3.1
0.6
0.8
0.4
0
PBS

**Figure S144 – (#48) DVB-LL-25 kinetic hemolysis assay. A.** Defibrinated sheep blood was incubated in the presence of the indicated peptide concentrations at 37 °C with monitoring of OD600 over 18 hours, where decreasing OD600 indicates hemolysis and each panel indicates an independent experiment. Peptide concentrations are in μM.

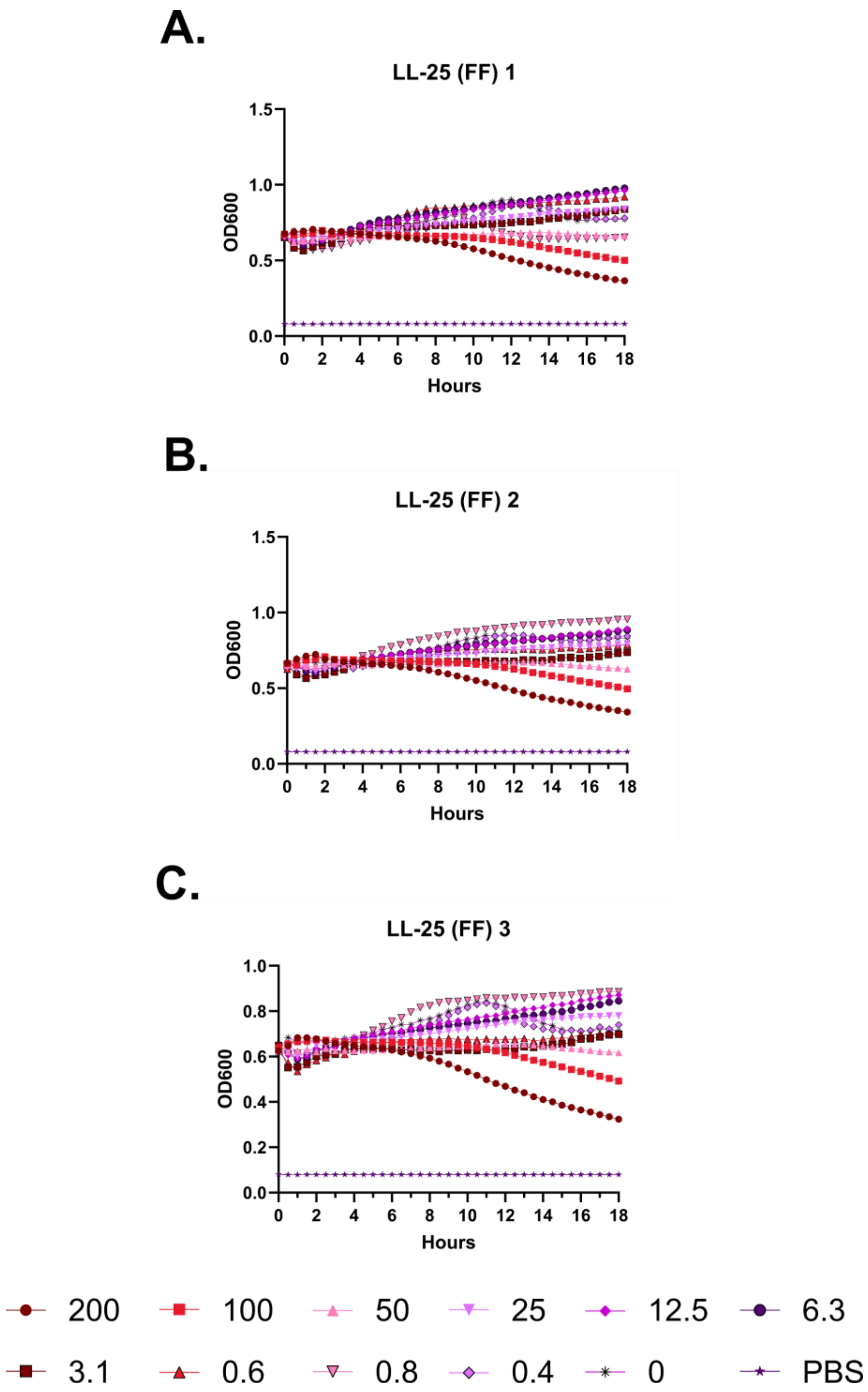
A.
LL-25 (FF) 1
OD600
Hours
B.
LL-25 (FF) 2
OD600
Hours
C.
LL-25 (FF) 3
OD600
Hours
200
100
50
25
12.5
6.3
3.1
0.6
0.8
0.4
0
PBS

**Figure S145 – (#49) LL-25 (FF) kinetic hemolysis assay. A.** Defibrinated sheep blood was incubated in the presence of the indicated peptide concentrations at 37 °C with monitoring of OD600 over 18 hours, where decreasing OD600 indicates hemolysis and each panel indicates an independent experiment. Peptide concentrations are in µM.

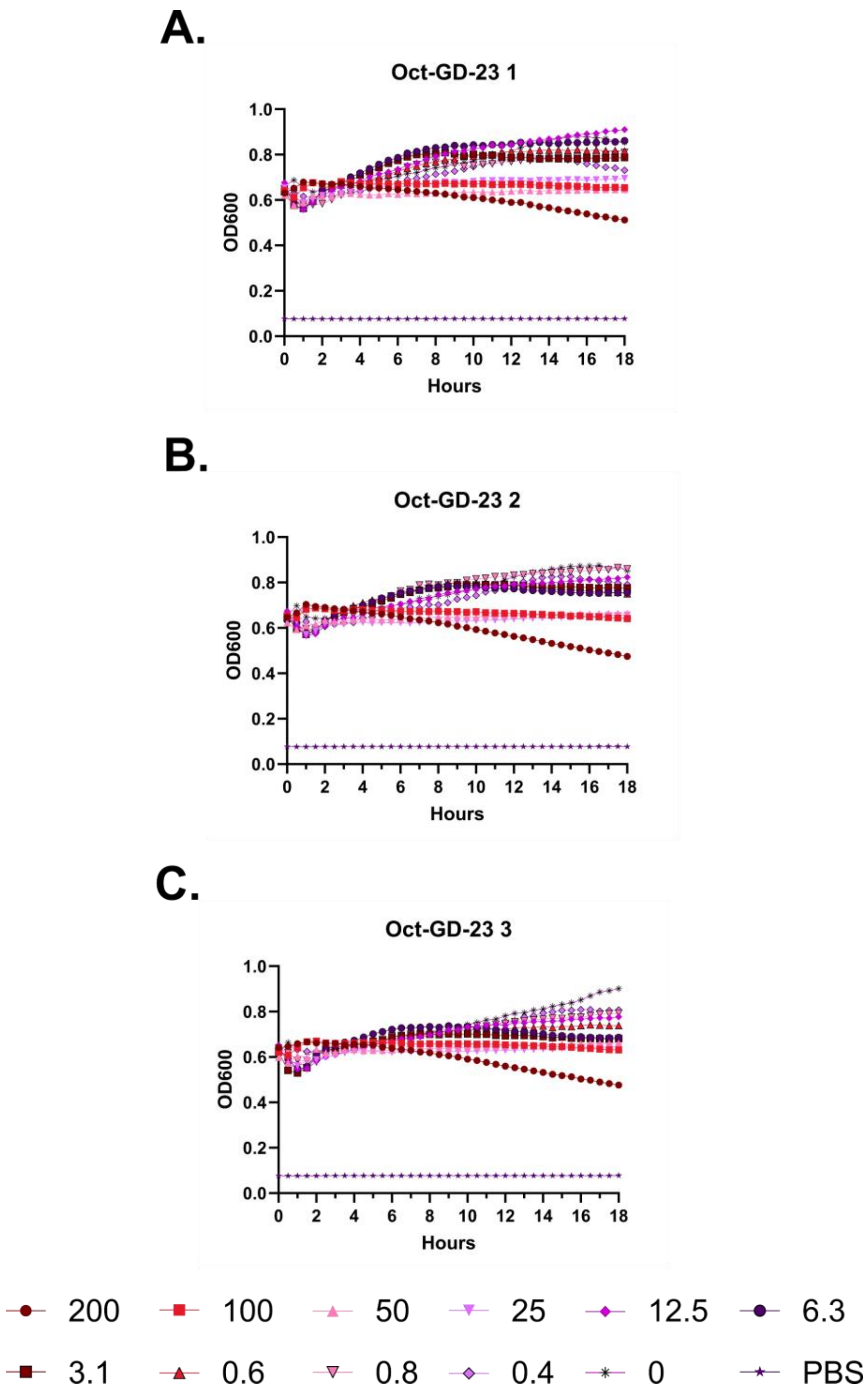
A.
Oct-GD-23 1
OD600
1.0
0.8
0.6
0.4
0.2
0.0
0 2 4 6 8 10 12 14 16 18
Hours
B.
Oct-GD-23 2
OD600
1.0
0.8
0.6
0.4
0.2
0.0
0 2 4 6 8 10 12 14 16 18
Hours
C.
Oct-GD-23 3
OD600
1.0
0.8
0.6
0.4
0.2
0.0
0 2 4 6 8 10 12 14 16 18
Hours
200
100
50
25
12.5
6.3
3.1
0.6
0.8
0.4
0
PBS

**Figure S146 – (#50) Oct-GD-23 kinetic hemolysis assay. A.** Defibrinated sheep blood was incubated in the presence of the indicated peptide concentrations at 37 °C with monitoring of OD600 over 18 hours, where decreasing OD600 indicates hemolysis and each panel indicates an independent experiment. Peptide concentrations are in μM.

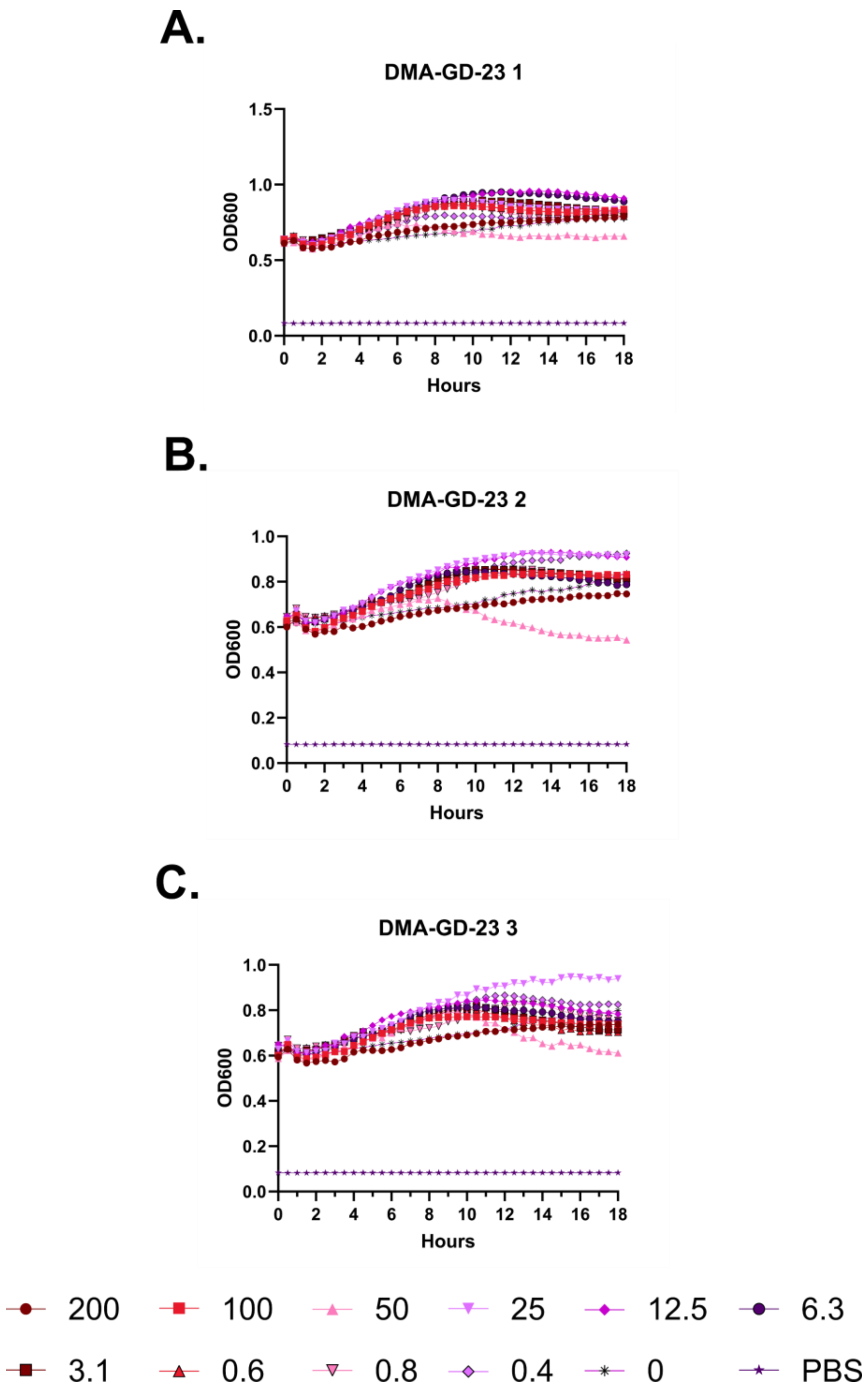
A.
DMA-GD-23 1
OD600
Hours
B.
DMA-GD-23 2
OD600
Hours
C.
DMA-GD-23 3
OD600
Hours
200
100
50
25
12.5
6.3
3.1
0.6
0.8
0.4
0
PBS

**Figure S147 – (#51) DMA-GD-23 kinetic hemolysis assay. A.** Defibrinated sheep blood was incubated in the presence of the indicated peptide concentrations at 37 °C with monitoring of OD600 over 18 hours, where decreasing OD600 indicates hemolysis and each panel indicates an independent experiment. Peptide concentrations are in μM.

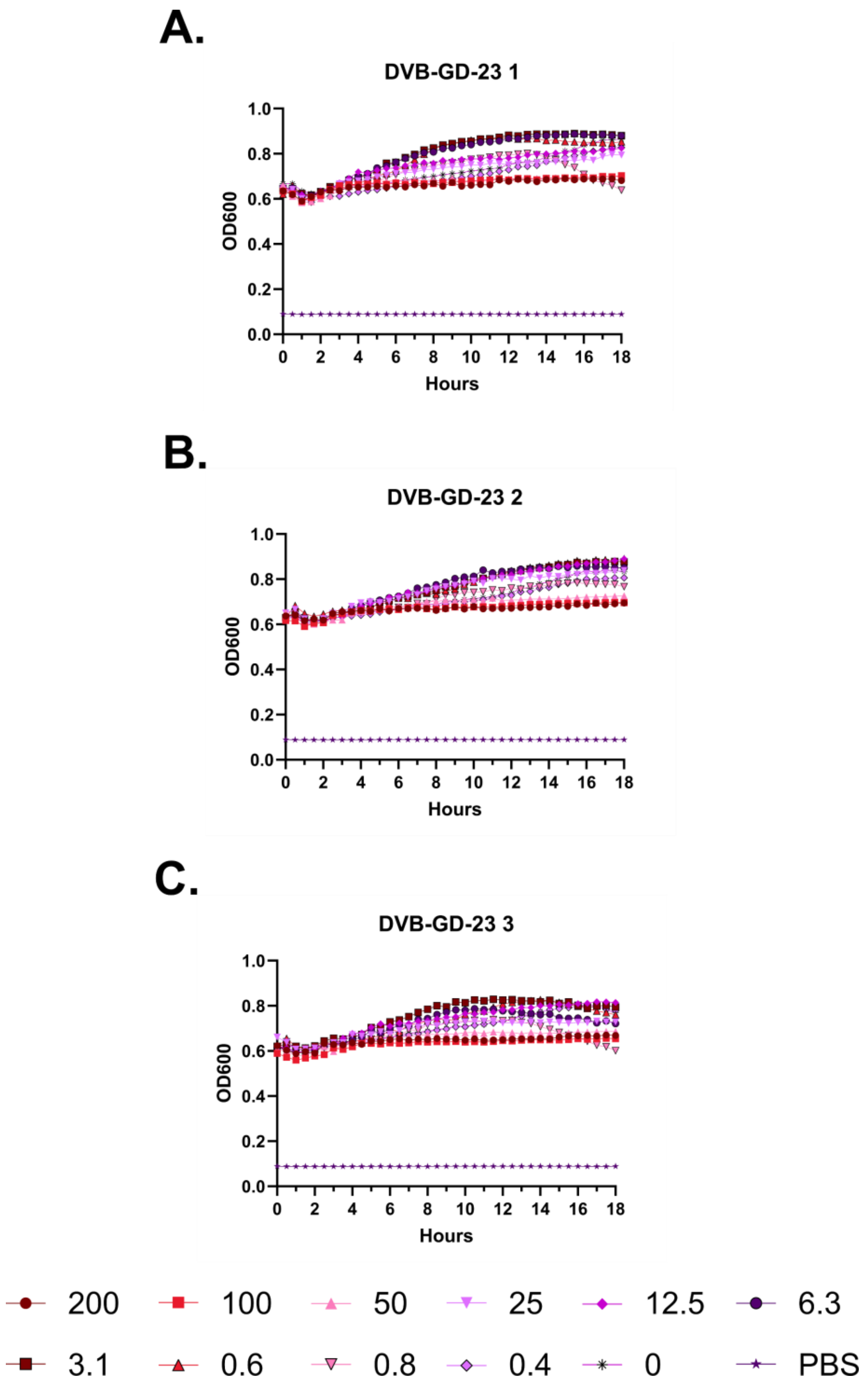
A.
DVB-GD-23 1
OD600
Hours
B.
DVB-GD-23 2
OD600
Hours
C.
DVB-GD-23 3
OD600
Hours
200
100
50
25
12.5
6.3
3.1
0.6
0.8
0.4
0
PBS

**Figure S148 – (#52) DVB-GD-23 kinetic hemolysis assay. A.** Defibrinated sheep blood was incubated in the presence of the indicated peptide concentrations at 37 °C with monitoring of OD600 over 18 hours, where decreasing OD600 indicates hemolysis and each panel indicates an independent experiment. Peptide concentrations are in μM.

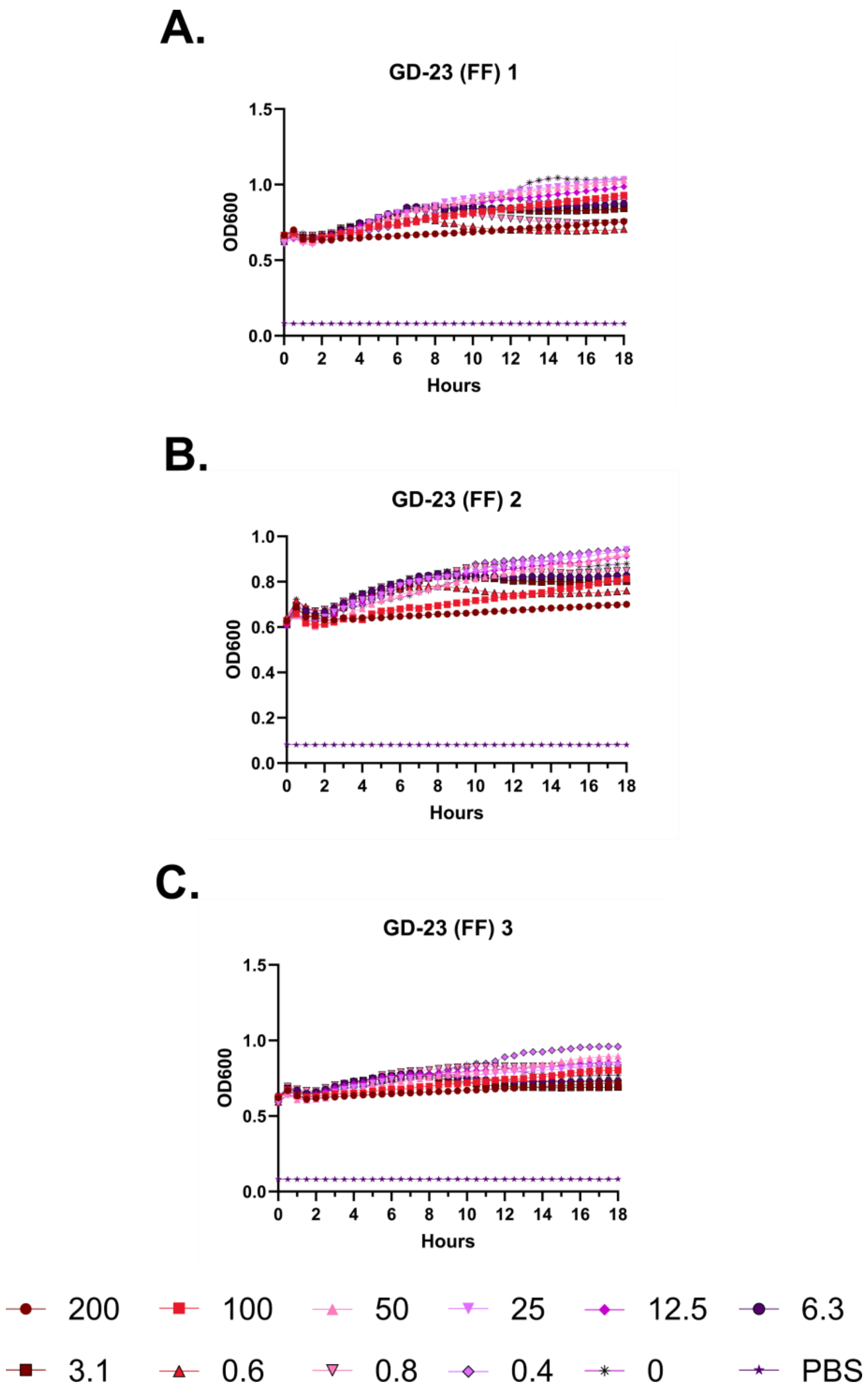
A.
GD-23 (FF) 1
OD600
Hours
B.
GD-23 (FF) 2
OD600
Hours
C.
GD-23 (FF) 3
OD600
Hours
200
100
50
25
12.5
6.3
3.1
0.6
0.8
0.4
0
PBS

**Figure S149 – (#53) GD-23 (FF) kinetic hemolysis assay. A.** Defibrinated sheep blood was incubated in the presence of the indicated peptide concentrations at 37 °C with monitoring of OD600 over 18 hours, where decreasing OD600 indicates hemolysis and each panel indicates an independent experiment. Peptide concentrations are in μM.

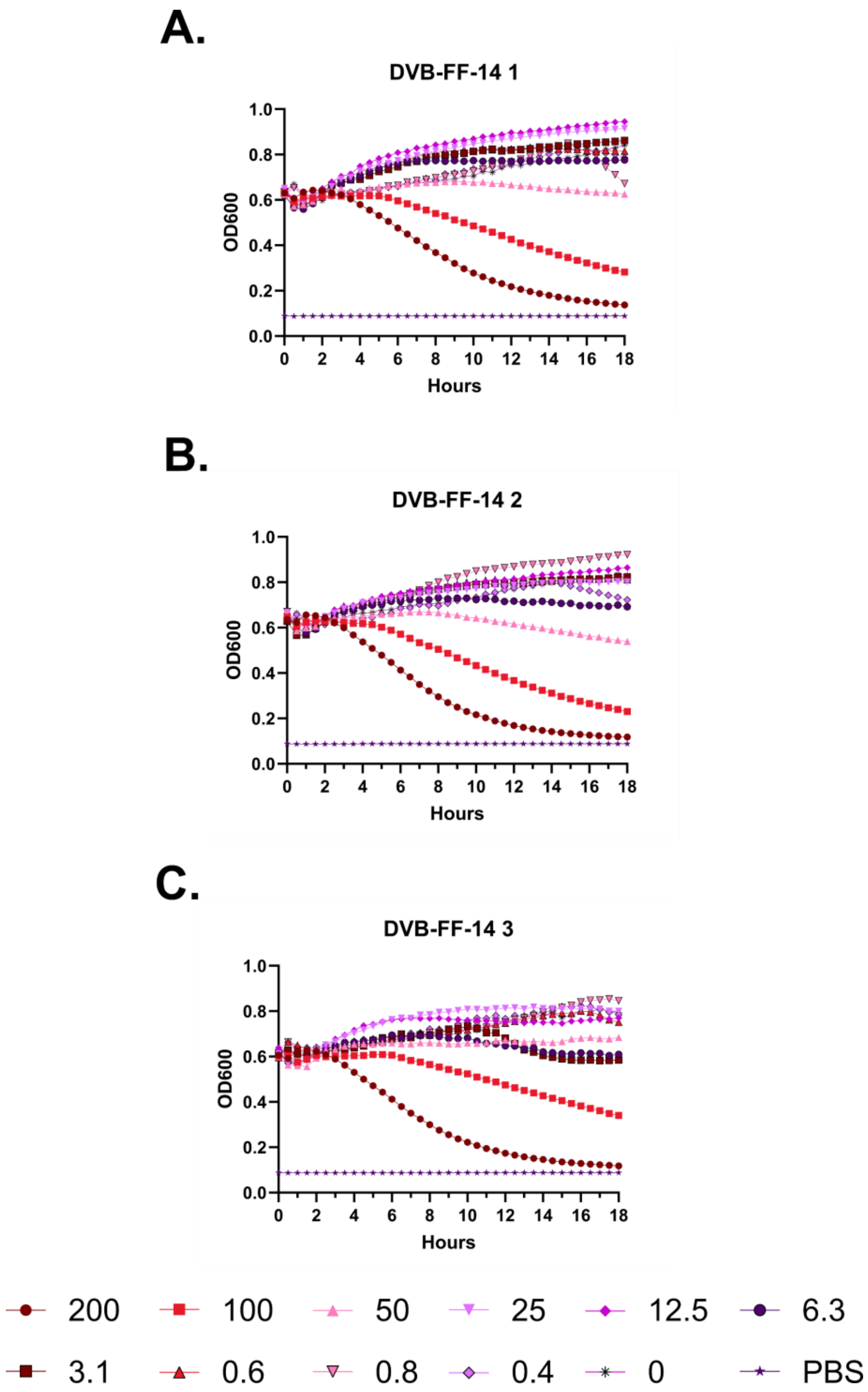
A.
DVB-FF-14 1
OD600
Hours
B.
DVB-FF-14 2
OD600
Hours
C.
DVB-FF-14 3
OD600
Hours
200
100
50
25
12.5
6.3
3.1
0.6
0.8
0.4
0
PBS

**Figure S150 – (#54) DVB-FF-14 kinetic hemolysis assay. A.** Defibrinated sheep blood was incubated in the presence of the indicated peptide concentrations at 37 °C with monitoring of OD600 over 18 hours, where decreasing OD600 indicates hemolysis and each panel indicates an independent experiment. Peptide concentrations are in µM.

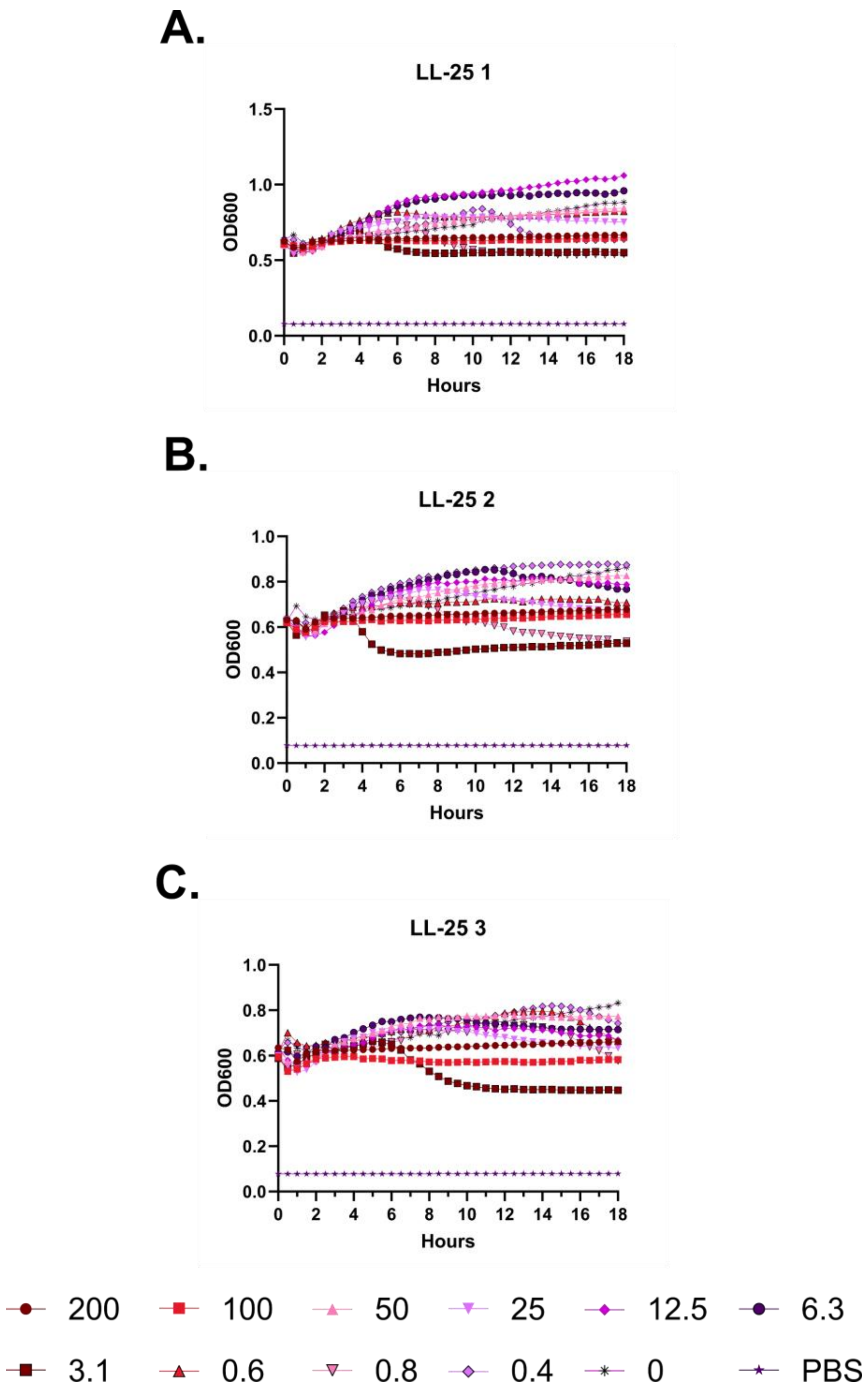
A.
LL-25 1
OD600
Hours
B.
LL-25 2
OD600
Hours
C.
LL-25 3
OD600
Hours
200
100
50
25
12.5
6.3
3.1
0.6
0.8
0.4
0
PBS

**Figure S151 – (#2) LL-25 kinetic hemolysis assay. A.** Defibrinated sheep blood was incubated in the presence of the indicated peptide concentrations at 37 °C with monitoring of OD600 over 18 hours, where decreasing OD600 indicates hemolysis and each panel indicates an independent experiment. Peptide concentrations are in μM.

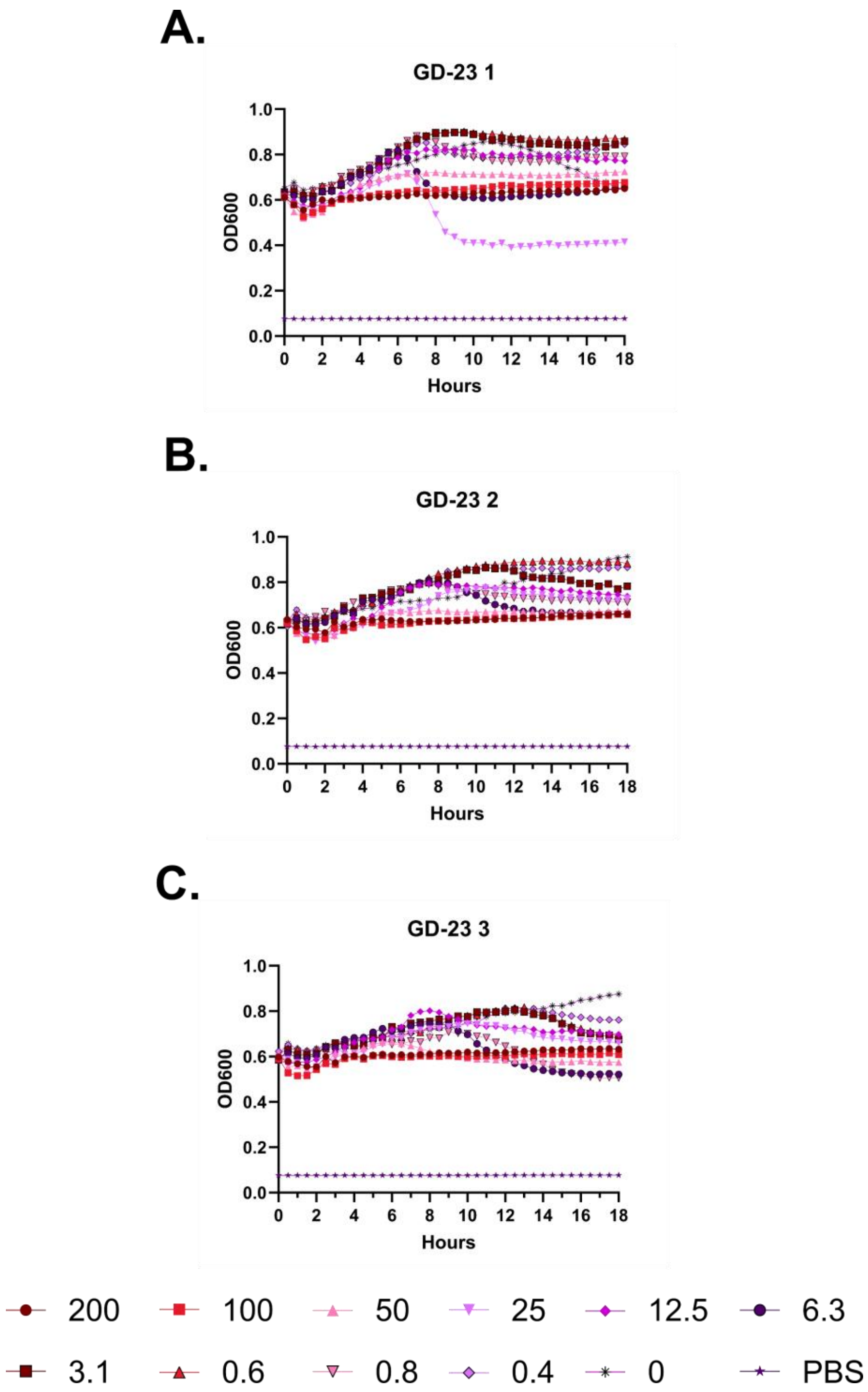
A.
GD-23 1
OD600
Hours
B.
GD-23 2
OD600
Hours
C.
GD-23 3
OD600
Hours
200
100
50
25
12.5
6.3
3.1
0.6
0.8
0.4
0
PBS

**Figure S152 – (#4) GD-23 kinetic hemolysis assay. A.** Defibrinated sheep blood was incubated in the presence of the indicated peptide concentrations at 37 °C with monitoring of OD600 over 18 hours, where decreasing OD600 indicates hemolysis and each panel indicates an independent experiment. Peptide concentrations are in μM.

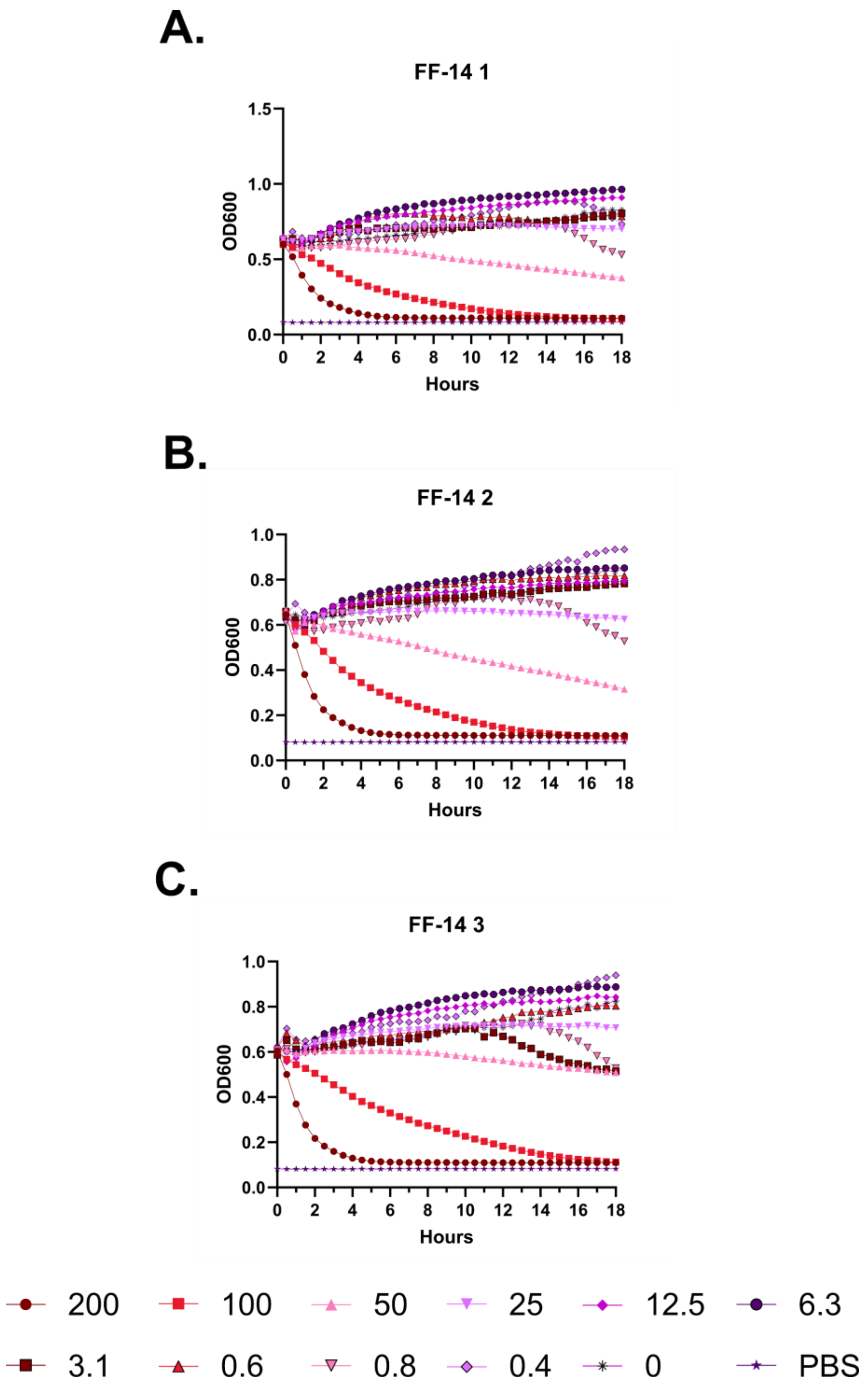
A.
FF-14 1
OD600
Hours
B.
FF-14 2
OD600
Hours
C.
FF-14 3
OD600
Hours
200
100
50
25
12.5
6.3
3.1
0.6
0.8
0.4
0
PBS

**Figure S153 – (#18) FF-14 kinetic hemolysis assay. A.** Defibrinated sheep blood was incubated in the presence of the indicated peptide concentrations at 37 °C with monitoring of OD600 over 18 hours, where decreasing OD600 indicates hemolysis and each panel indicates an independent experiment. Peptide concentrations are in μM.

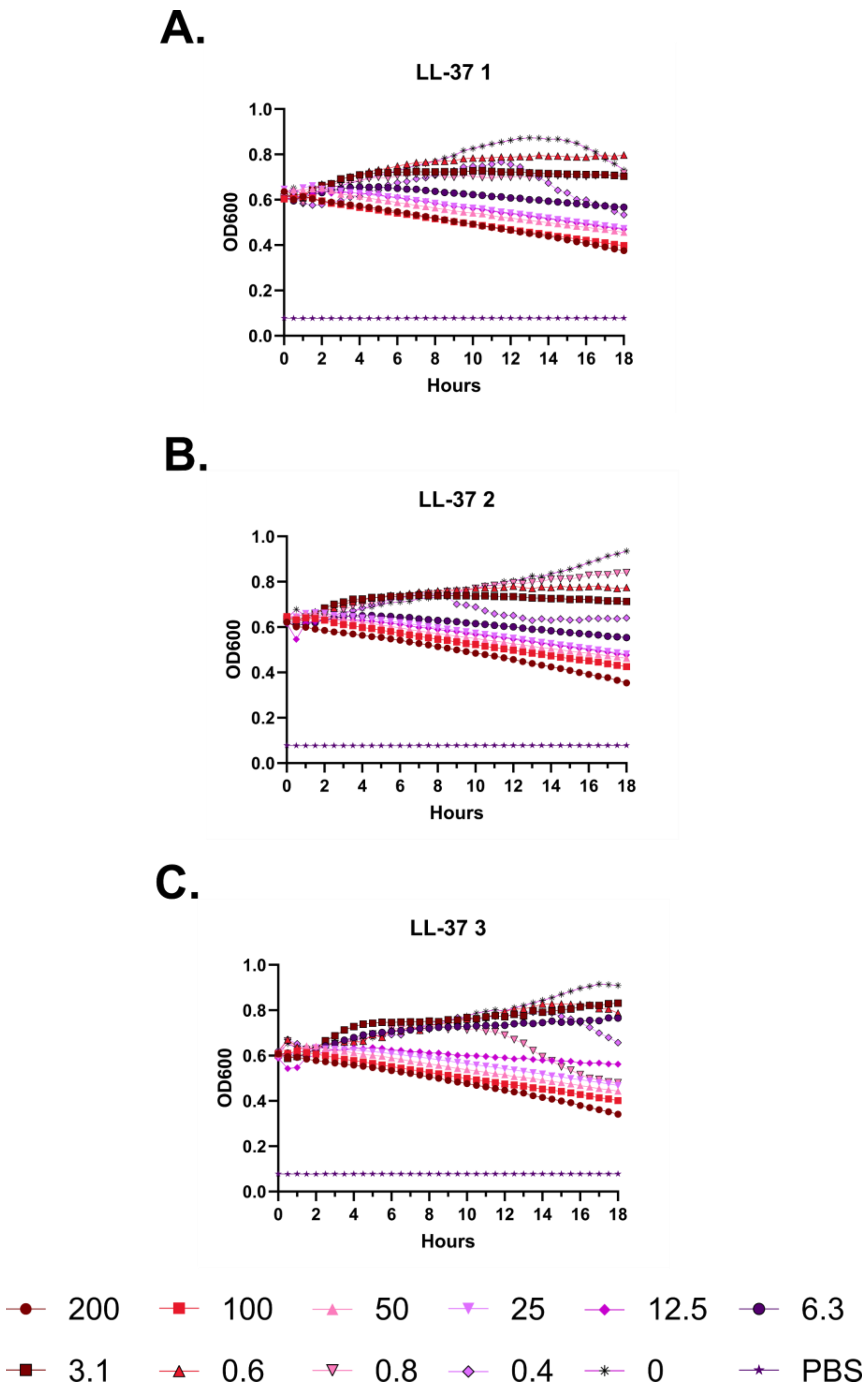
A.
LL-37 1
OD600
Hours
B.
LL-37 2
OD600
Hours
C.
LL-37 3
OD600
Hours
200
100
50
25
12.5
6.3
3.1
0.6
0.8
0.4
0
PBS

**Figure S154 – (#1) LL-37 kinetic hemolysis assay. A.** Defibrinated sheep blood was incubated in the presence of the indicated peptide concentrations at 37 °C with monitoring of OD600 over 18 hours, where decreasing OD600 indicates hemolysis and each panel indicates an independent experiment. Peptide concentrations are in μM.

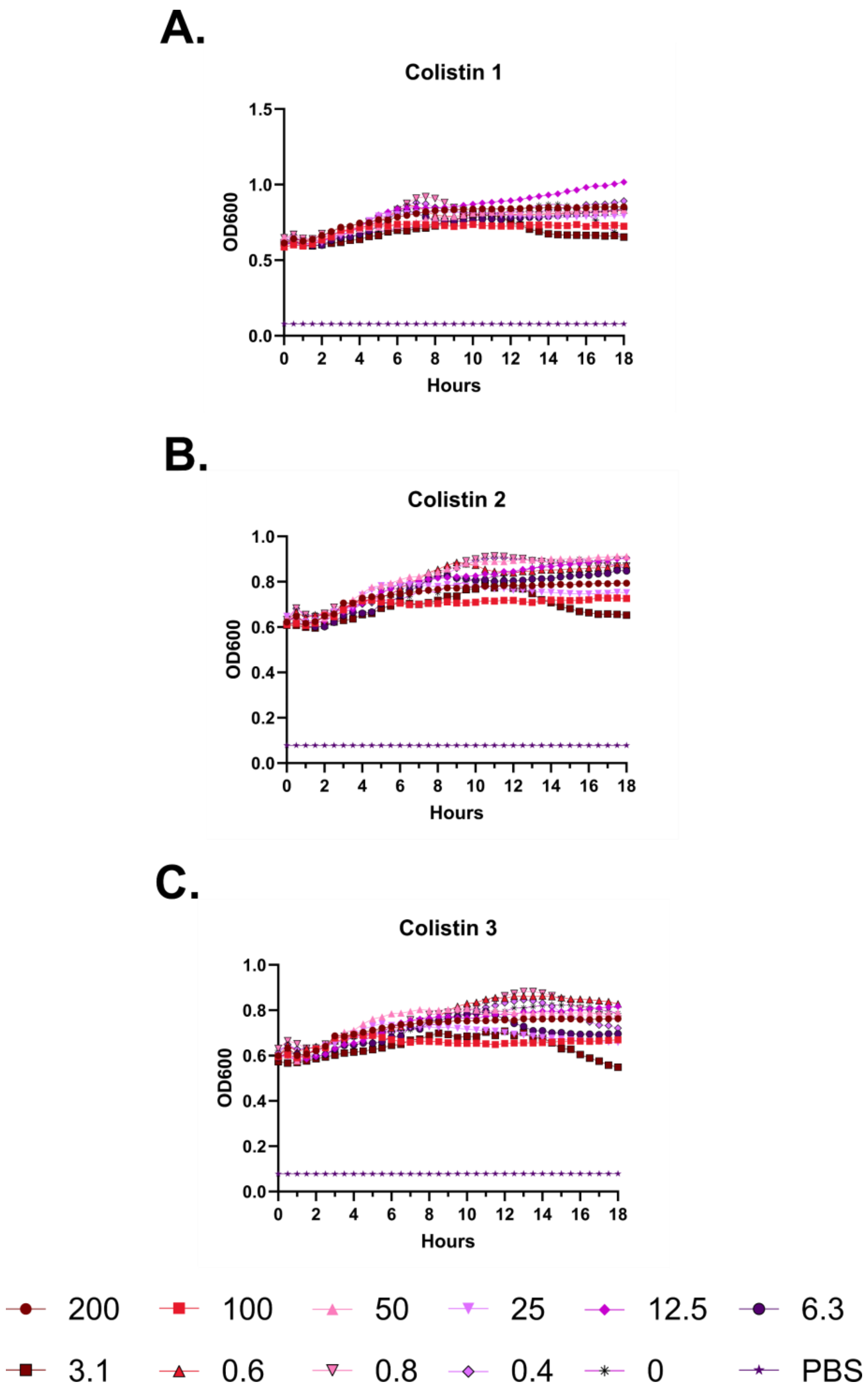
A.
Colistin 1
OD600
1.5
1.0
0.5
0.0
0 2 4 6 8 10 12 14 16 18
Hours
B.
Colistin 2
OD600
1.0
0.8
0.6
0.4
0.2
0.0
0 2 4 6 8 10 12 14 16 18
Hours
C.
Colistin 3
OD600
1.0
0.8
0.6
0.4
0.2
0.0
0 2 4 6 8 10 12 14 16 18
Hours
200
100
50
25
12.5
6.3
3.1
0.6
0.8
0.4
0
PBS

**Figure S155 – Colistin kinetic hemolysis assay. A.** Defibrinated sheep blood was incubated in the presence of the indicated peptide concentrations at 37 °C with monitoring of OD600 over 18 hours, where decreasing OD600 indicates hemolysis and each panel indicates an independent experiment. Peptide concentrations are in μM.

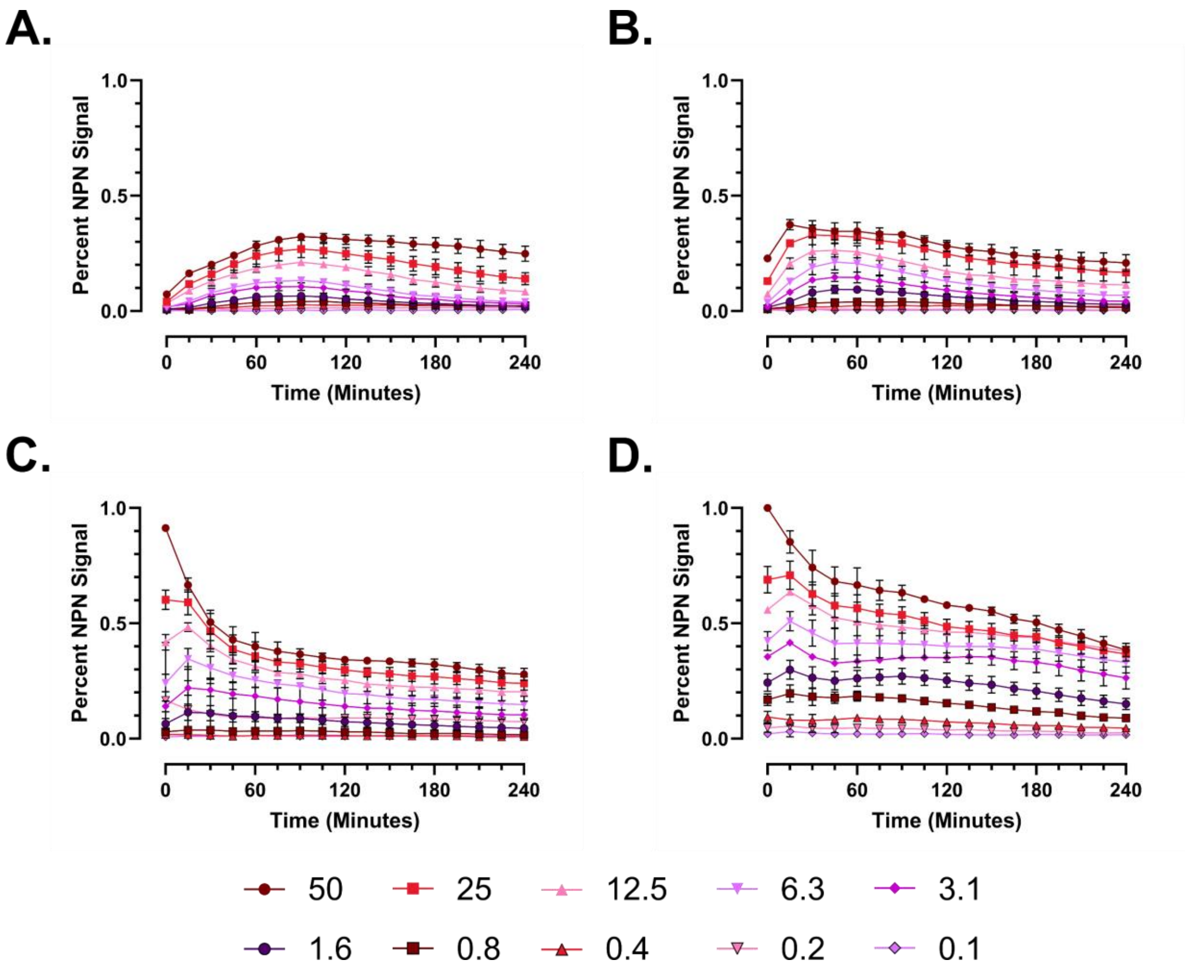
A.
B.
C.
D.
Percent NPN Signal
Time (Minutes)
1.0
0.5
0.0
0
60
120
180
240
50
25
12.5
6.3
3.1
1.6
0.8
0.4
0.2
0.1

**Figure S156 – Relative outer membrane permeabilization of short cathelicidin derivatives over time.** Dual membrane permeabilization assays were completed by incubating *E. coli* (ATCC 25922) in the presence of N-phenyl-naphthylamine (NPN) and propidium iodide (PI) as a markers of outer and inner membrane permeabilization, respectively. Results show NPN signal in the presence of a range of concentrations of (#10) KR-12 (**A**), (#7) FK-13 (**B**), (#18) FF-14 (**C**), or (#1) LL-37 (**D**) over 240 minutes, where all results are scaled to (#1) LL-37.

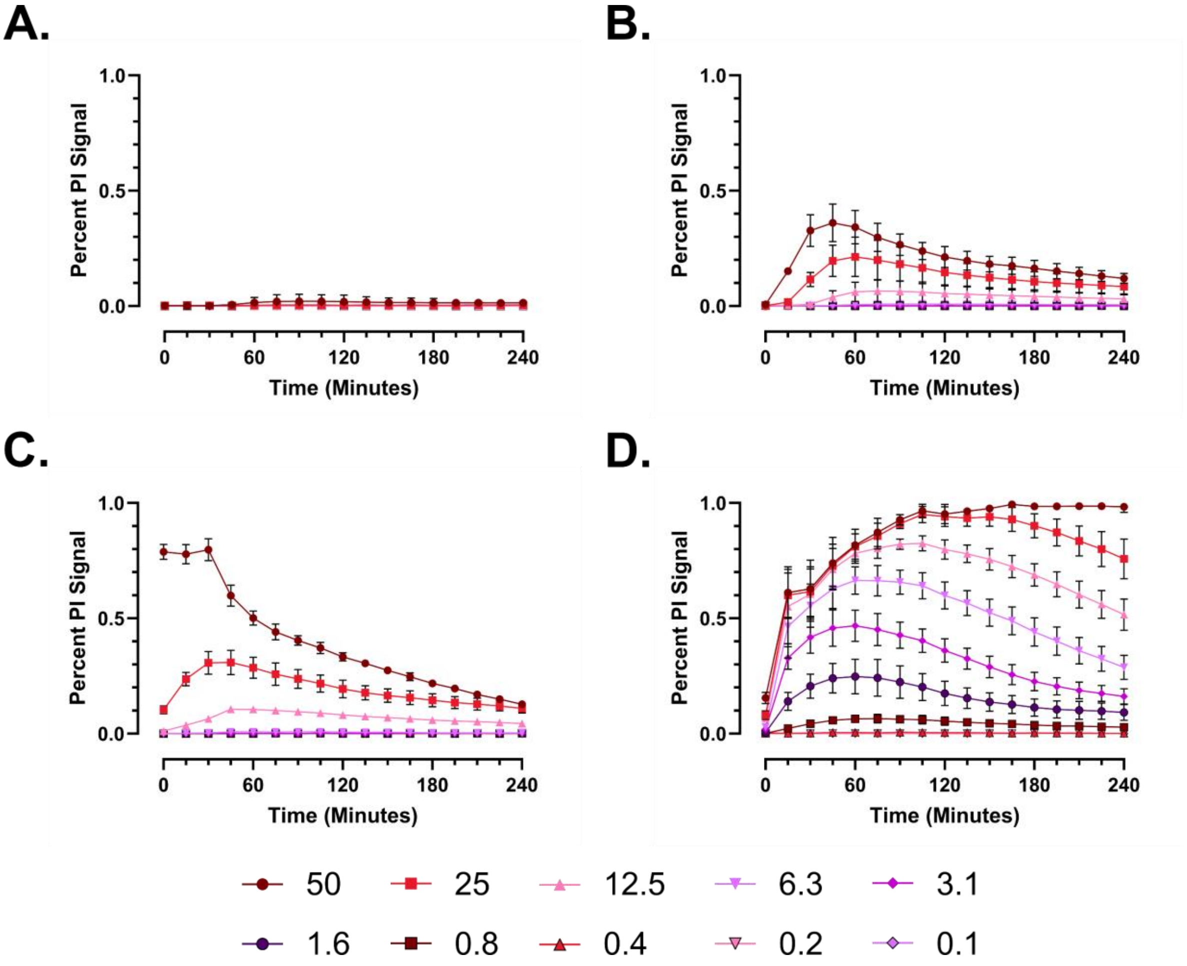
A.
B.
C.
D.
Percent PI Signal
1.0
0.5
0.0
0
60
120
180
240
Time (Minutes)
50
25
12.5
6.3
3.1
1.6
0.8
0.4
0.2
0.1

**Figure S157 – Relative inner membrane permeabilization of short cathelicidin derivatives over time.** Dual membrane permeabilization assays were completed by incubating *E. coli* (ATCC 25922) in the presence of N-phenyl-naphthylamine (NPN) and propidium iodide (PI) as a markers of outer and inner membrane permeabilization, respectively. Results show PI signal in the presence of a range of concentrations of (#10) KR-12 (**A**), (#7) FK-13 (**B**), (#18) FF-14 (**C**), or (#1) LL-37 (**D**) over 240 minutes, where all results are scaled to (#1) LL-37.

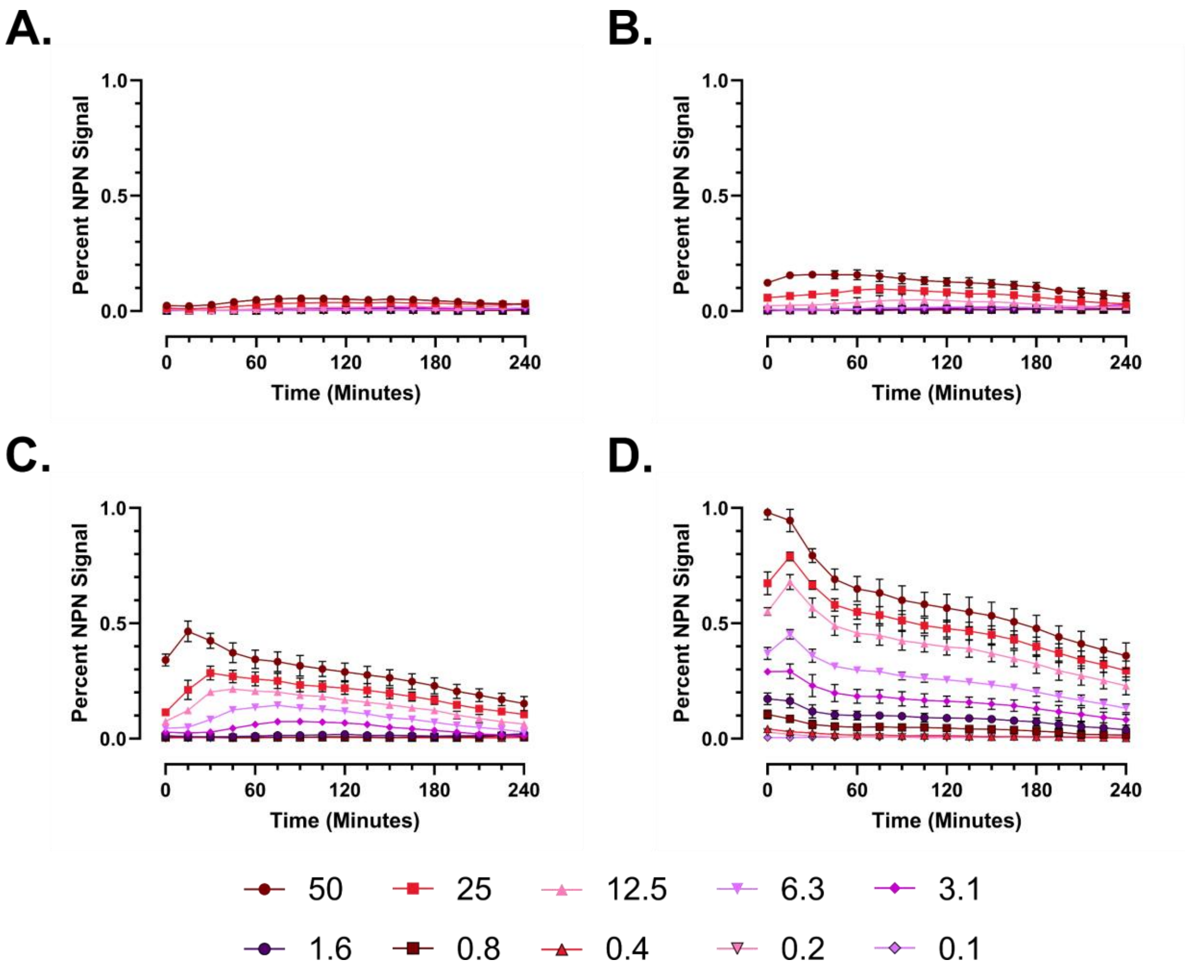

A.
B.
C.
D.
Percent NPN Signal
1.0
0.5
0.0
0
60
120
180
240
Time (Minutes)
50
25
12.5
6.3
3.1
1.6
0.8
0.4
0.2
0.1

**Figure S158 – Relative outer membrane permeabilization of short cathelicidin derivatives over time.** Dual membrane permeabilization assays were completed by incubating *P. aeruginosa* (ATCC 27853) in the presence of N-phenyl-naphthylamine (NPN) and propidium iodide (PI) as a markers of outer and inner membrane permeabilization, respectively. Results show NPN signal in the presence of a range of concentrations of (#10) KR-12 (**A**), (#7) FK-13 (**B**), (#18) FF-14 (**C**), or (#1) LL-37 (**D**) over 240 minutes, where all results are scaled to (#1) LL-37.

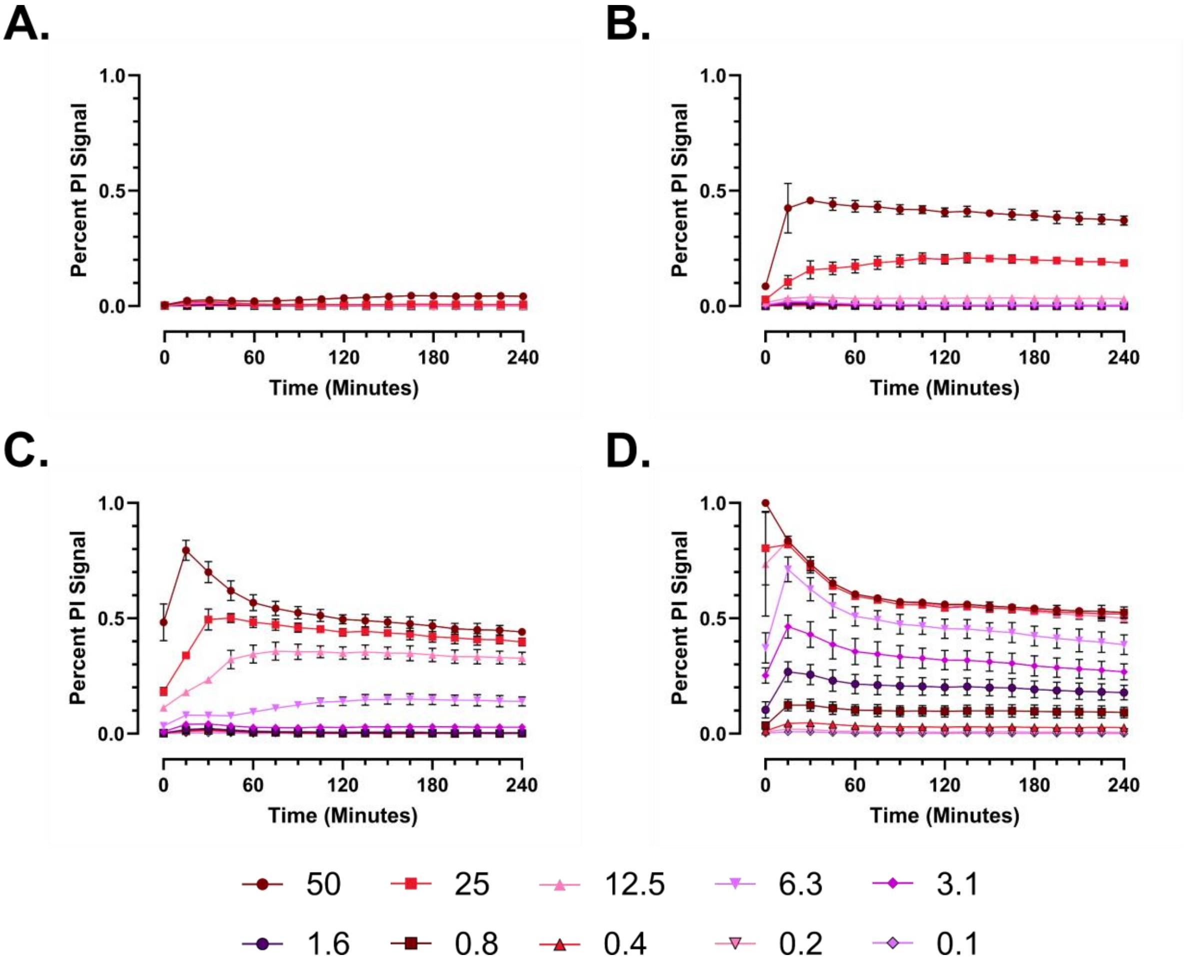
A.
B.
C.
D.
Percent PI Signal
1.0
0.5
0.0
0
60
120
180
240
Time (Minutes)
50
25
12.5
6.3
3.1
1.6
0.8
0.4
0.2
0.1

**Figure S159 – Relative inner membrane permeabilization of short cathelicidin derivatives over time.** Dual membrane permeabilization assays were completed by incubating *P. aeruginosa* (ATCC 27853) in the presence of N-phenyl-naphthylamine (NPN) and propidium iodide (PI) as a markers of outer and inner membrane permeabilization, respectively. Results show PI signal in the presence of a range of concentrations of (#10) KR-12 (**A**), (#7) FK-13 (**B**), (#18) FF-14 (**C**), or (#1) LL-37 (**D**) over 240 minutes, where all results are scaled to (#1) LL-37.

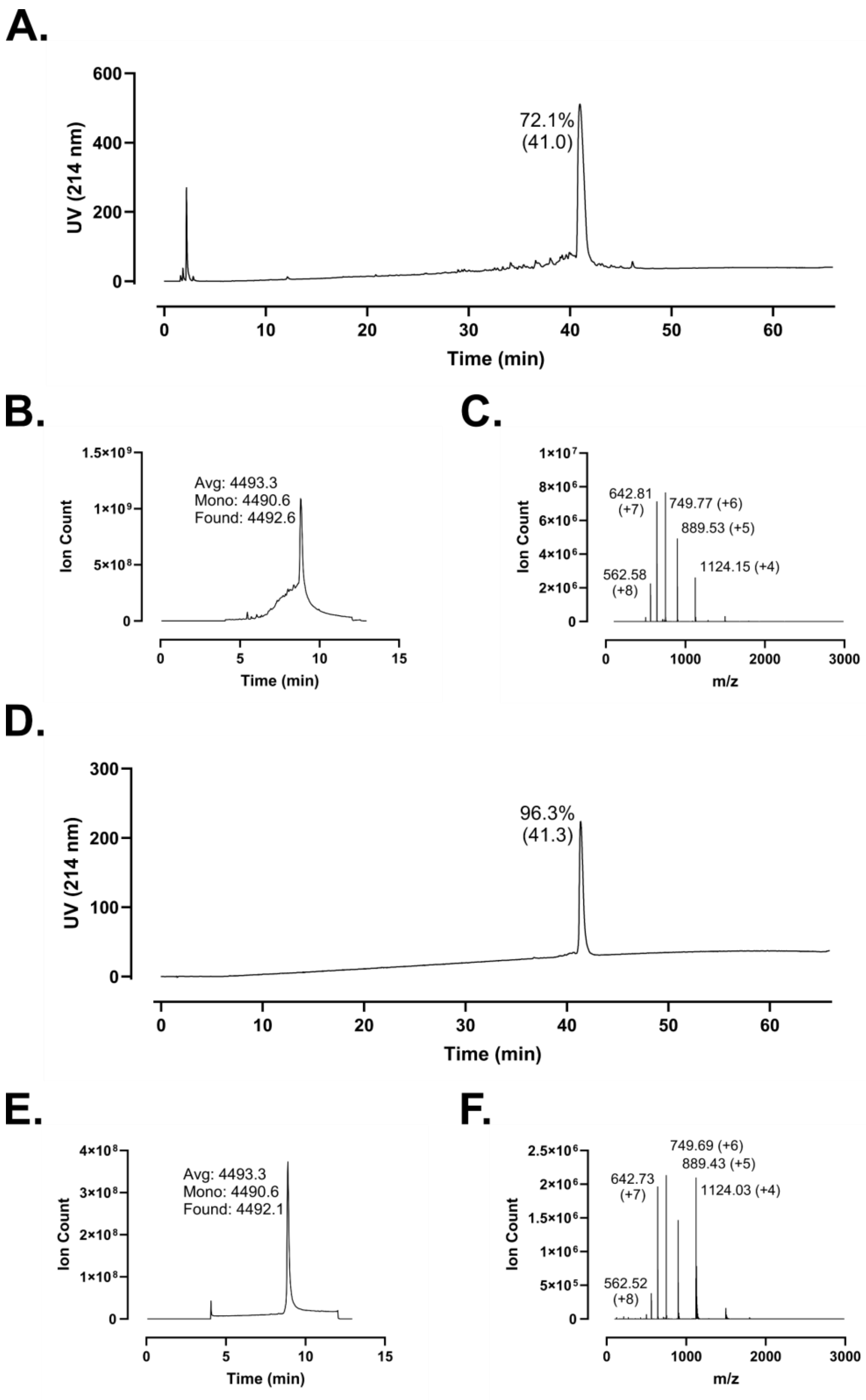
A.
UV (214 nm)
600
400
200
0
72.1%
(41.0)
0
10
20
30
40
50
60
Time (min)
B.
1.5×10⁹
1×10⁹
5×10⁸
0
Ion Count
Avg: 4493.3
Mono: 4490.6
Found: 4492.6
0
5
10
15
Time (min)
C.
1×10⁷
8×10⁶
6×10⁶
4×10⁶
2×10⁶
0
Ion Count
642.81
(+7)
749.77 (+6)
889.53 (+5)
562.58
(+8)
1124.15 (+4)
0
1000
2000
3000
m/z
D.
UV (214 nm)
300
200
100
0
96.3%
(41.3)
0
10
20
30
40
50
60
Time (min)
E.
4×10⁸
3×10⁸
2×10⁸
1×10⁸
0
Ion Count
Avg: 4493.3
Mono: 4490.6
Found: 4492.1
0
5
10
15
Time (min)
F.
2.5×10⁶
2×10⁶
1.5×10⁶
1×10⁶
5×10⁵
0
Ion Count
749.69 (+6)
889.43 (+5)
642.73
(+7)
1124.03 (+4)
562.52
(+8)
0
1000
2000
3000
m/z

**Figure S160 – (#1) LL-37 Analytical Data. A.** HPLC trace of crude peptide at 214 nm. Peak labeling indicates percent purity by trace integration. Retention time is given in parentheses. **B.** LCMS total ion chromatogram (TIC) of crude peptide. Peak labeling indicates the predicted average and monoisotopic masses as well as the observed peptide mass. **C.** Mass spectrum at peak of **B** with *m/z* labeling for selected peaks and the associated charge in parentheses. **D.** HPLC trace of peptide following single-pass purification. **E.** TIC of peptide following single-pass purification. **F.** Mass spectrum at peak of **E**.

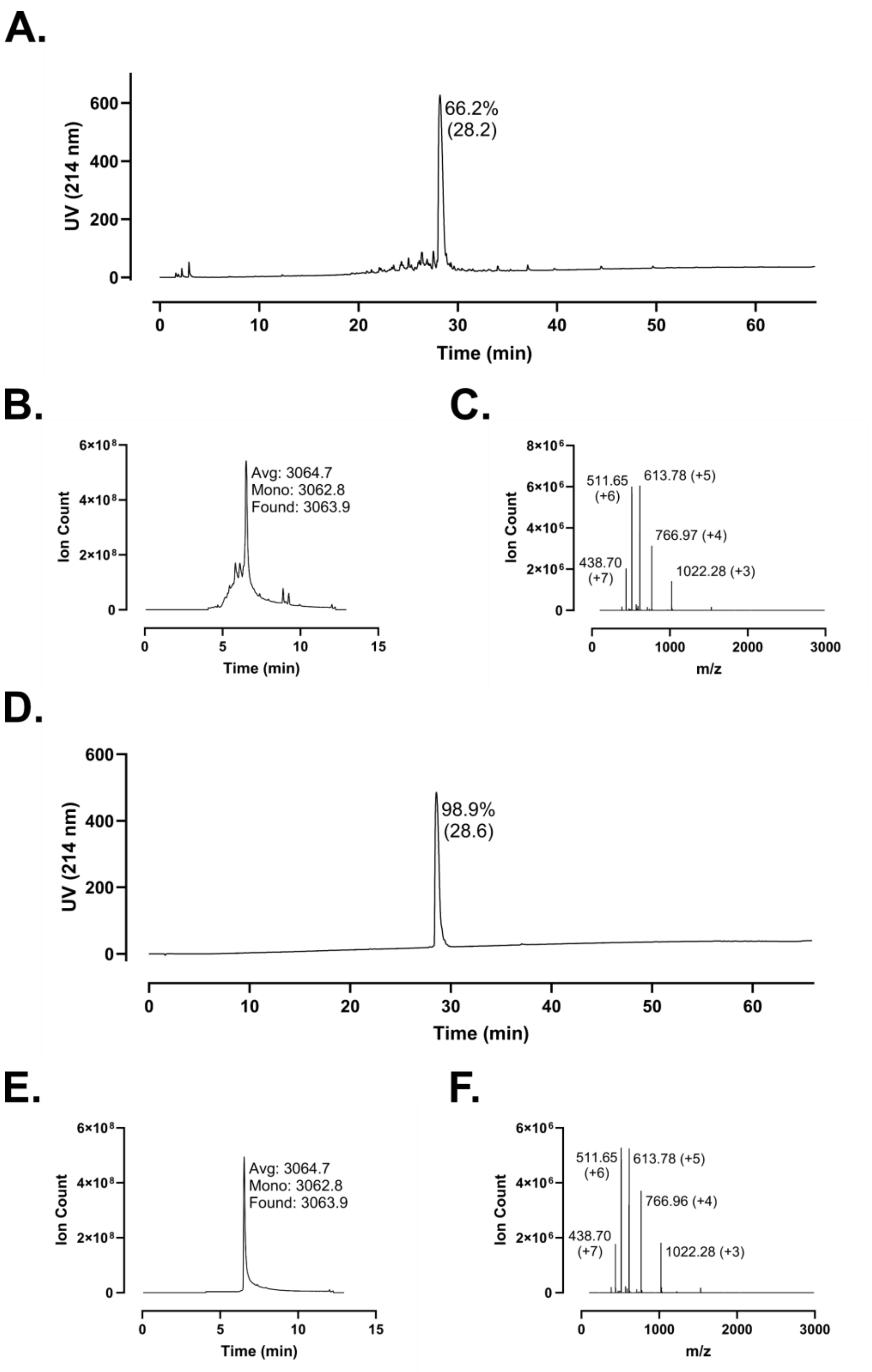
A.
66.2%
(28.2)
UV (214 nm)
Time (min)
B.
Avg: 3064.7
Mono: 3062.8
Found: 3063.9
Ion Count
Time (min)
C.
511.65
(+6)
613.78 (+5)
766.97 (+4)
438.70
(+7)
1022.28 (+3)
Ion Count
m/z
D.
98.9%
(28.6)
UV (214 nm)
Time (min)
E.
Avg: 3064.7
Mono: 3062.8
Found: 3063.9
Ion Count
Time (min)
F.
511.65
(+6)
613.78 (+5)
766.96 (+4)
438.70
(+7)
1022.28 (+3)
Ion Count
m/z

**Figure S161 – (#2) LL-25 Analytical Data. A.** HPLC trace of crude peptide at 214 nm. Peak labeling indicates percent purity by trace integration. Retention time is given in parentheses. **B.** LCMS total ion chromatogram (TIC) of crude peptide. Peak labeling indicates the predicted average and monoisotopic masses as well as the observed peptide mass. **C.** Mass spectrum at peak of **B** with *m/z* labeling for selected peaks and the associated charge in parentheses. **D.** HPLC trace of peptide following single-pass purification. **E.** TIC of peptide following single-pass purification. **F.** Mass spectrum at peak of **E**.

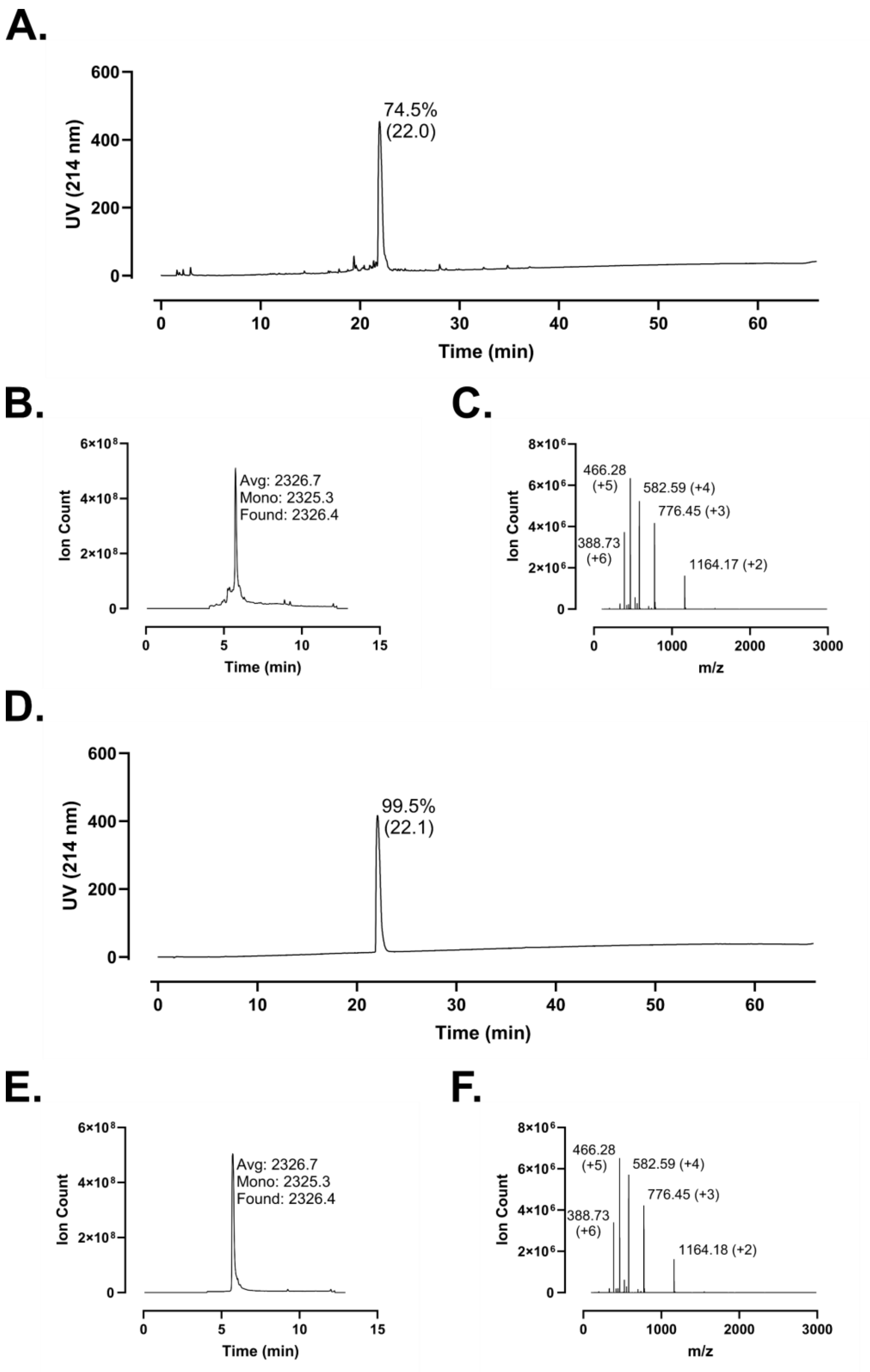
A.
UV (214 nm)
74.5%
(22.0)
Time (min)
B.
Ion Count
Avg: 2326.7
Mono: 2325.3
Found: 2326.4
Time (min)
C.
Ion Count
466.28
(+5)
582.59 (+4)
776.45 (+3)
388.73
(+6)
1164.17 (+2)
m/z
D.
UV (214 nm)
99.5%
(22.1)
Time (min)
E.
Ion Count
Avg: 2326.7
Mono: 2325.3
Found: 2326.4
Time (min)
F.
Ion Count
466.28
(+5)
582.59 (+4)
776.45 (+3)
388.73
(+6)
1164.18 (+2)
m/z

**Figure S162 – (#3) LL-19 Analytical Data. A.** HPLC trace of crude peptide at 214 nm. Peak labeling indicates percent purity by trace integration. Retention time is given in parentheses. **B.** LCMS total ion chromatogram (TIC) of crude peptide. Peak labeling indicates the predicted average and monoisotopic masses as well as the observed peptide mass. **C.** Mass spectrum at peak of **B** with *m/z* labeling for selected peaks and the associated charge in parentheses. **D.** HPLC trace of peptide following single-pass purification. **E.** TIC of peptide following single-pass purification. **F.** Mass spectrum at peak of **E**.

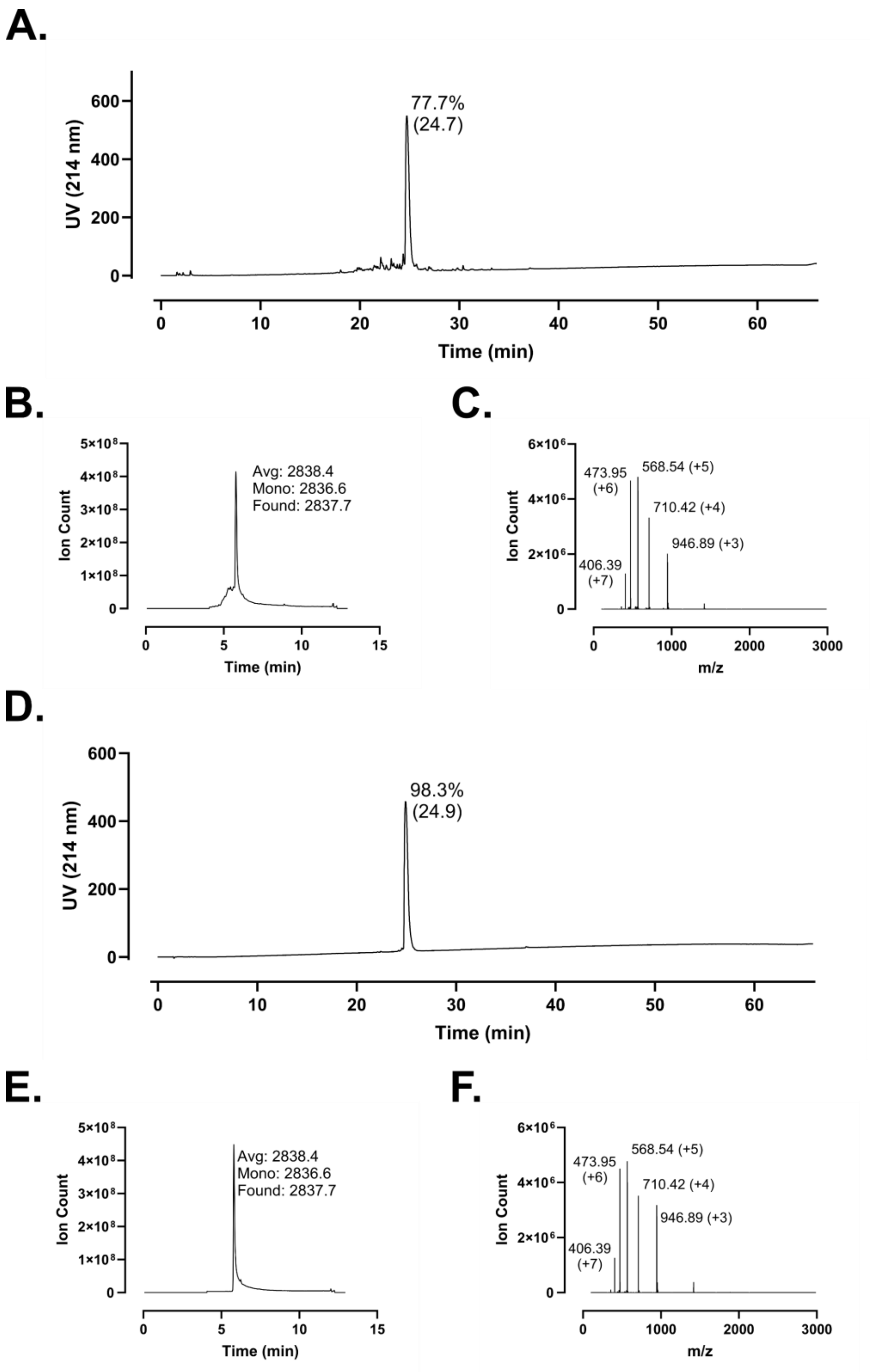
A.
77.7%
(24.7)
UV (214 nm)
Time (min)
B.
Avg: 2838.4
Mono: 2836.6
Found: 2837.7
Ion Count
Time (min)
C.
473.95
(+6)
568.54 (+5)
710.42 (+4)
946.89 (+3)
406.39
(+7)
Ion Count
m/z
D.
98.3%
(24.9)
UV (214 nm)
Time (min)
E.
Avg: 2838.4
Mono: 2836.6
Found: 2837.7
Ion Count
Time (min)
F.
473.95
(+6)
568.54 (+5)
710.42 (+4)
946.89 (+3)
406.39
(+7)
Ion Count
m/z

**Figure S163 – (#4) GD-23 Analytical Data. A.** HPLC trace of crude peptide at 214 nm. Peak labeling indicates percent purity by trace integration. Retention time is given in parentheses. **B.** LCMS total ion chromatogram (TIC) of crude peptide. Peak labeling indicates the predicted average and monoisotopic masses as well as the observed peptide mass. **C.** Mass spectrum at peak of **B** with *m/z* labeling for selected peaks and the associated charge in parentheses. **D.** HPLC trace of peptide following single-pass purification. **E.** TIC of peptide following single-pass purification. **F.** Mass spectrum at peak of **E**.

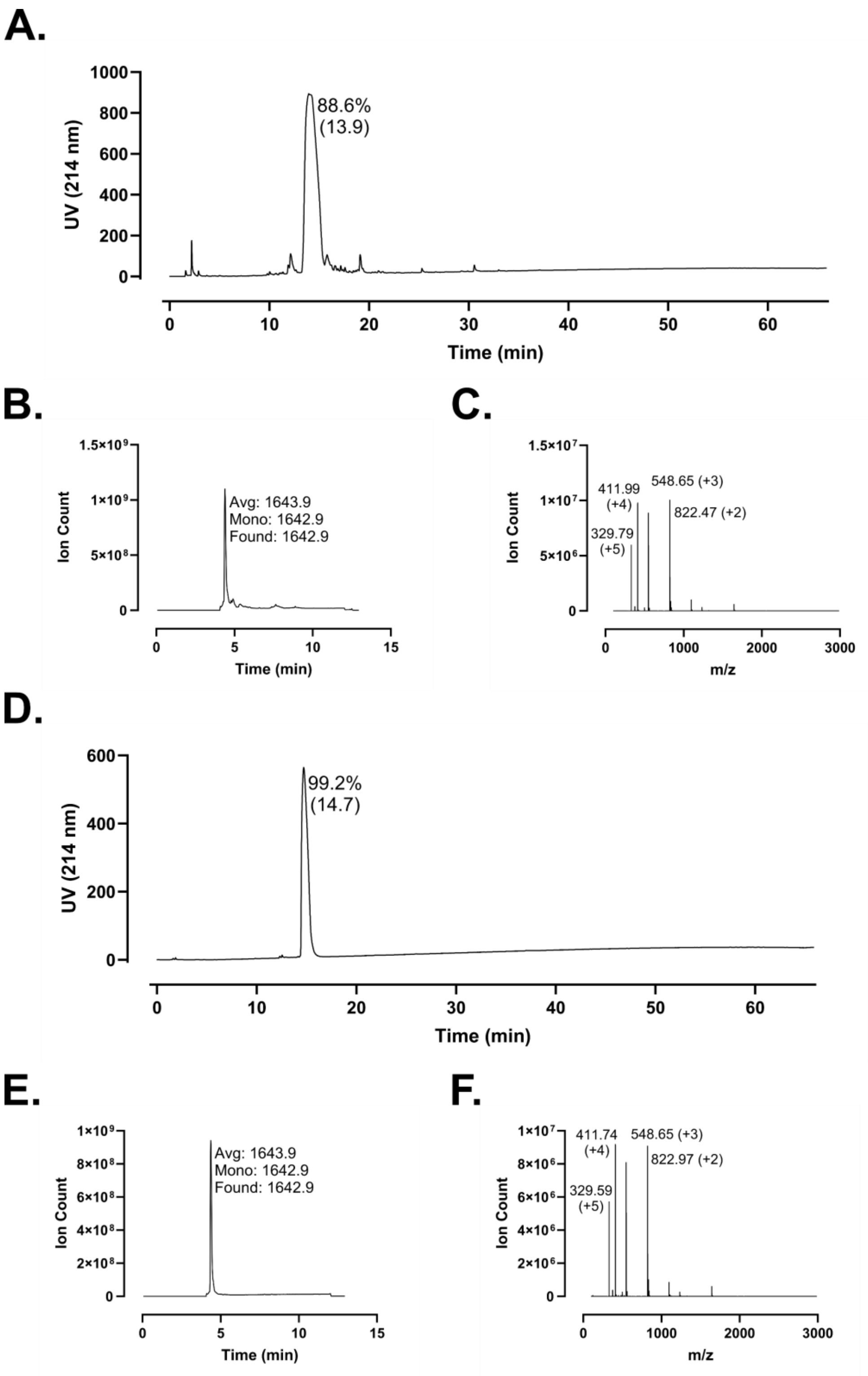
A.
88.6%
(13.9)
UV (214 nm)
Time (min)
B.
Avg: 1643.9
Mono: 1642.9
Found: 1642.9
Ion Count
Time (min)
C.
411.99
(+4)
548.65 (+3)
822.47 (+2)
329.79
(+5)
Ion Count
m/z
D.
99.2%
(14.7)
UV (214 nm)
Time (min)
E.
Avg: 1643.9
Mono: 1642.9
Found: 1642.9
Ion Count
Time (min)
F.
411.74
(+4)
548.65 (+3)
822.97 (+2)
329.59
(+5)
Ion Count
m/z

**Figure S164 – (#5) FF-13 Analytical Data. A.** HPLC trace of crude peptide at 214 nm. Peak labeling indicates percent purity by trace integration. Retention time is given in parentheses. **B.** LCMS total ion chromatogram (TIC) of crude peptide. Peak labeling indicates the predicted average and monoisotopic masses as well as the observed peptide mass. **C.** Mass spectrum at peak of **B** with *m/z* labeling for selected peaks and the associated charge in parentheses. **D.** HPLC trace of peptide following single-pass purification. **E.** TIC of peptide following single-pass purification. **F.** Mass spectrum at peak of **E**.

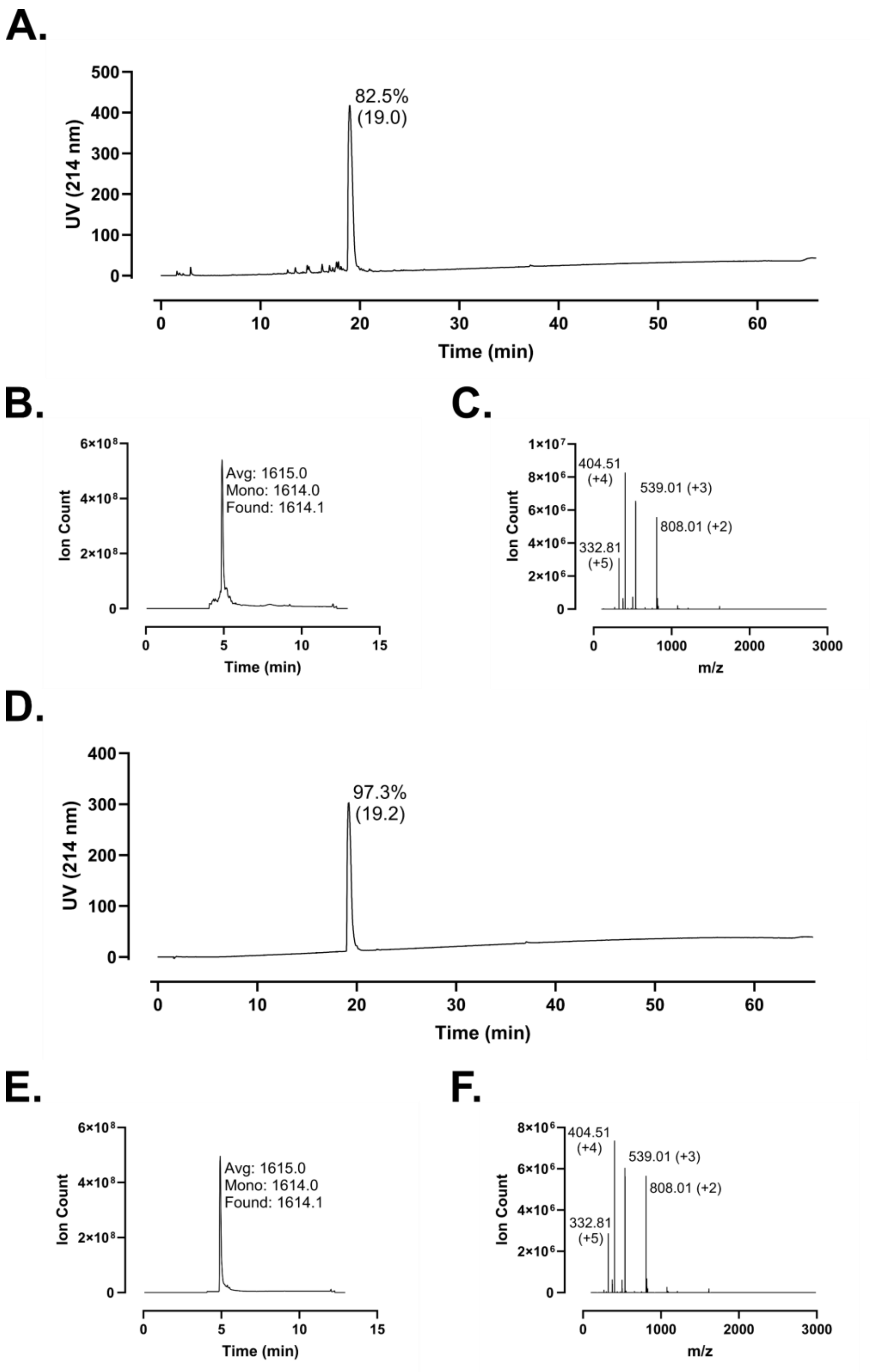
A.
82.5%
(19.0)
UV (214 nm)
Time (min)
B.
Avg: 1615.0
Mono: 1614.0
Found: 1614.1
Ion Count
Time (min)
C.
404.51
(+4)
539.01 (+3)
808.01 (+2)
332.81
(+5)
Ion Count
m/z
D.
97.3%
(19.2)
UV (214 nm)
Time (min)
E.
Avg: 1615.0
Mono: 1614.0
Found: 1614.1
Ion Count
Time (min)
F.
404.51
(+4)
539.01 (+3)
808.01 (+2)
332.81
(+5)
Ion Count
m/z

**Figure S165 – (#6) IG-13 Analytical Data. A.** HPLC trace of crude peptide at 214 nm. Peak labeling indicates percent purity by trace integration. Retention time is given in parentheses. **B.** LCMS total ion chromatogram (TIC) of crude peptide. Peak labeling indicates the predicted average and monoisotopic masses as well as the observed peptide mass. **C.** Mass spectrum at peak of **B** with *m/z* labeling for selected peaks and the associated charge in parentheses. **D.** HPLC trace of peptide following single-pass purification. **E.** TIC of peptide following single-pass purification. **F.** Mass spectrum at peak of **E**.

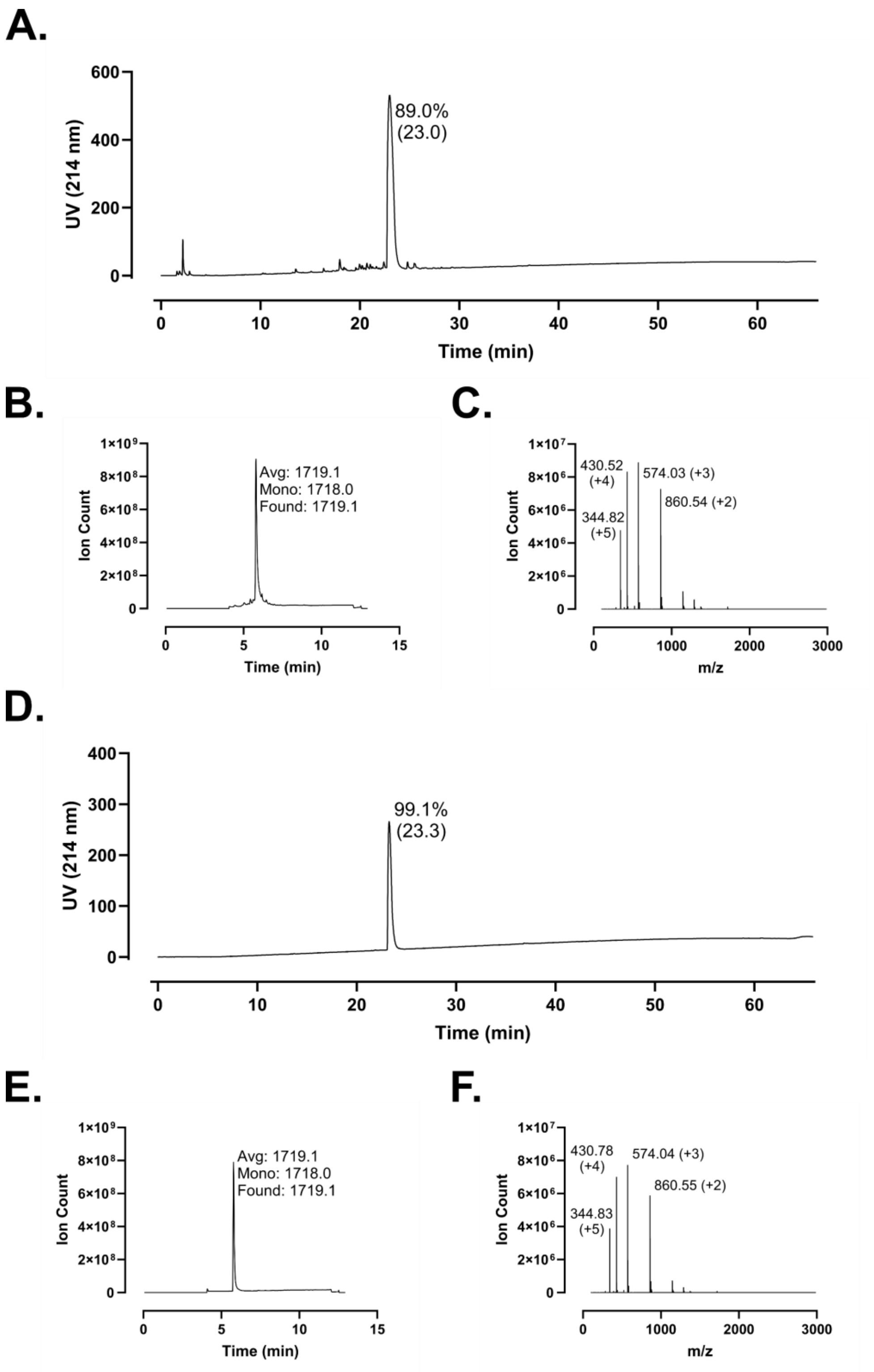
A.
UV (214 nm)
Time (min)
89.0%
(23.0)
B.
Ion Count
Time (min)
Avg: 1719.1
Mono: 1718.0
Found: 1719.1
C.
Ion Count
m/z
430.52
(+4)
574.03 (+3)
860.54 (+2)
344.82
(+5)
D.
UV (214 nm)
Time (min)
99.1%
(23.3)
E.
Ion Count
Time (min)
Avg: 1719.1
Mono: 1718.0
Found: 1719.1
F.
Ion Count
m/z
430.78
(+4)
574.04 (+3)
860.55 (+2)
344.83
(+5)

**Figure S166 – (#7) FK-13 Analytical Data. A.** HPLC trace of crude peptide at 214 nm. Peak labeling indicates percent purity by trace integration. Retention time is given in parentheses. **B.** LCMS total ion chromatogram (TIC) of crude peptide. Peak labeling indicates the predicted average and monoisotopic masses as well as the observed peptide mass. **C.** Mass spectrum at peak of **B** with *m/z* labeling for selected peaks and the associated charge in parentheses. **D.** HPLC trace of peptide following single-pass purification. **E.** TIC of peptide following single-pass purification. **F.** Mass spectrum at peak of **E**.

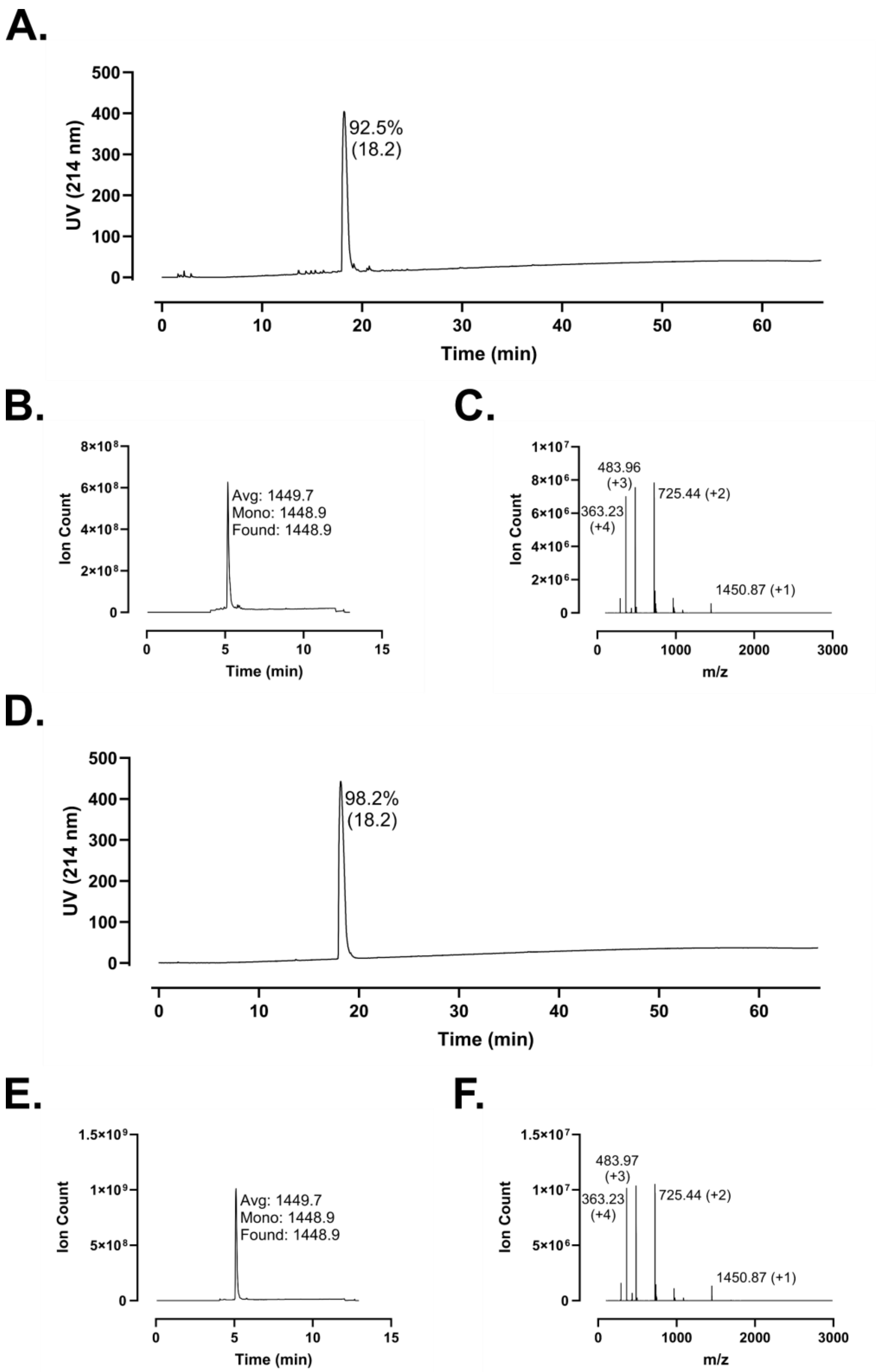
A.
UV (214 nm)
92.5%
(18.2)
Time (min)
B.
Ion Count
Avg: 1449.7
Mono: 1448.9
Found: 1448.9
Time (min)
C.
Ion Count
483.96
(+3)
363.23
(+4)
725.44 (+2)
1450.87 (+1)
m/z
D.
UV (214 nm)
98.2%
(18.2)
Time (min)
E.
Ion Count
Avg: 1449.7
Mono: 1448.9
Found: 1448.9
Time (min)
F.
Ion Count
483.97
(+3)
363.23
(+4)
725.44 (+2)
1450.87 (+1)
m/z

**Figure S167 – (#8) FK-11 Analytical Data. A.** HPLC trace of crude peptide at 214 nm. Peak labeling indicates percent purity by trace integration. Retention time is given in parentheses. **B.** LCMS total ion chromatogram (TIC) of crude peptide. Peak labeling indicates the predicted average and monoisotopic masses as well as the observed peptide mass. **C.** Mass spectrum at peak of **B** with *m/z* labeling for selected peaks and the associated charge in parentheses. **D.** HPLC trace of peptide following single-pass purification. **E.** TIC of peptide following single-pass purification. **F.** Mass spectrum at peak of **E**.

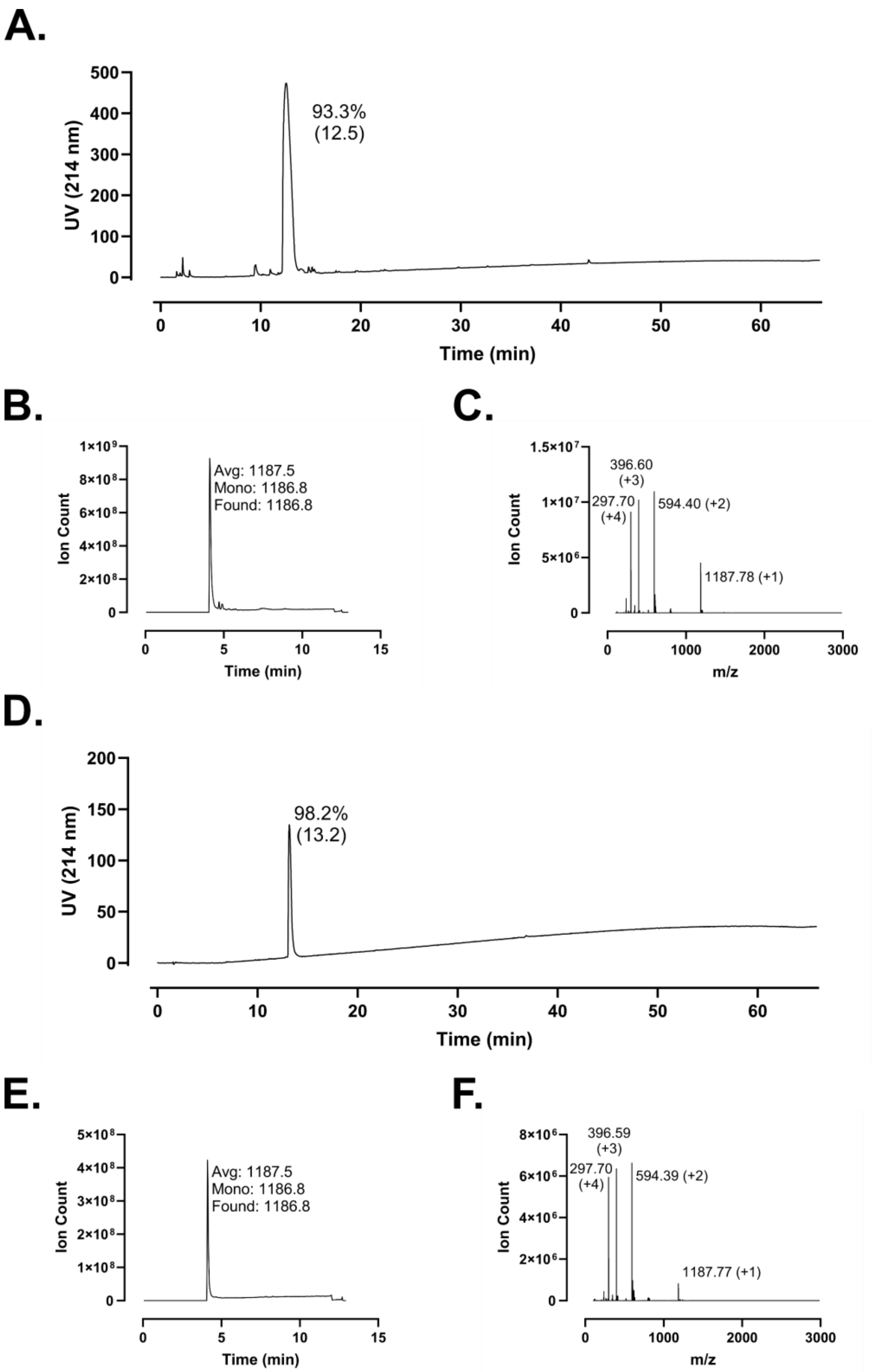
A.
UV (214 nm)
500
400
300
200
100
0
93.3%
(12.5)
0
10
20
30
40
50
60
Time (min)
B.
Ion Count
1×10⁹
8×10⁸
6×10⁸
4×10⁸
2×10⁸
0
Avg: 1187.5
Mono: 1186.8
Found: 1186.8
0
5
10
15
Time (min)
C.
Ion Count
1.5×10⁷
1×10⁷
5×10⁶
0
396.60
(+3)
297.70
(+4)
594.40 (+2)
1187.78 (+1)
0
1000
2000
3000
m/z
D.
UV (214 nm)
200
150
100
50
0
98.2%
(13.2)
0
10
20
30
40
50
60
Time (min)
E.
Ion Count
5×10⁸
4×10⁸
3×10⁸
2×10⁸
1×10⁸
0
Avg: 1187.5
Mono: 1186.8
Found: 1186.8
0
5
10
15
Time (min)
F.
Ion Count
8×10⁶
6×10⁶
4×10⁶
2×10⁶
0
396.59
(+3)
297.70
(+4)
594.39 (+2)
1187.77 (+1)
0
1000
2000
3000
m/z

**Figure S168 – (#9) FK-9 Analytical Data. A.** HPLC trace of crude peptide at 214 nm. Peak labeling indicates percent purity by trace integration. Retention time is given in parentheses. **B.** LCMS total ion chromatogram (TIC) of crude peptide. Peak labeling indicates the predicted average and monoisotopic masses as well as the observed peptide mass. **C.** Mass spectrum at peak of **B** with *m/z* labeling for selected peaks and the associated charge in parentheses. **D.** HPLC trace of peptide following single-pass purification. **E.** TIC of peptide following single-pass purification. **F.** Mass spectrum at peak of **E**.

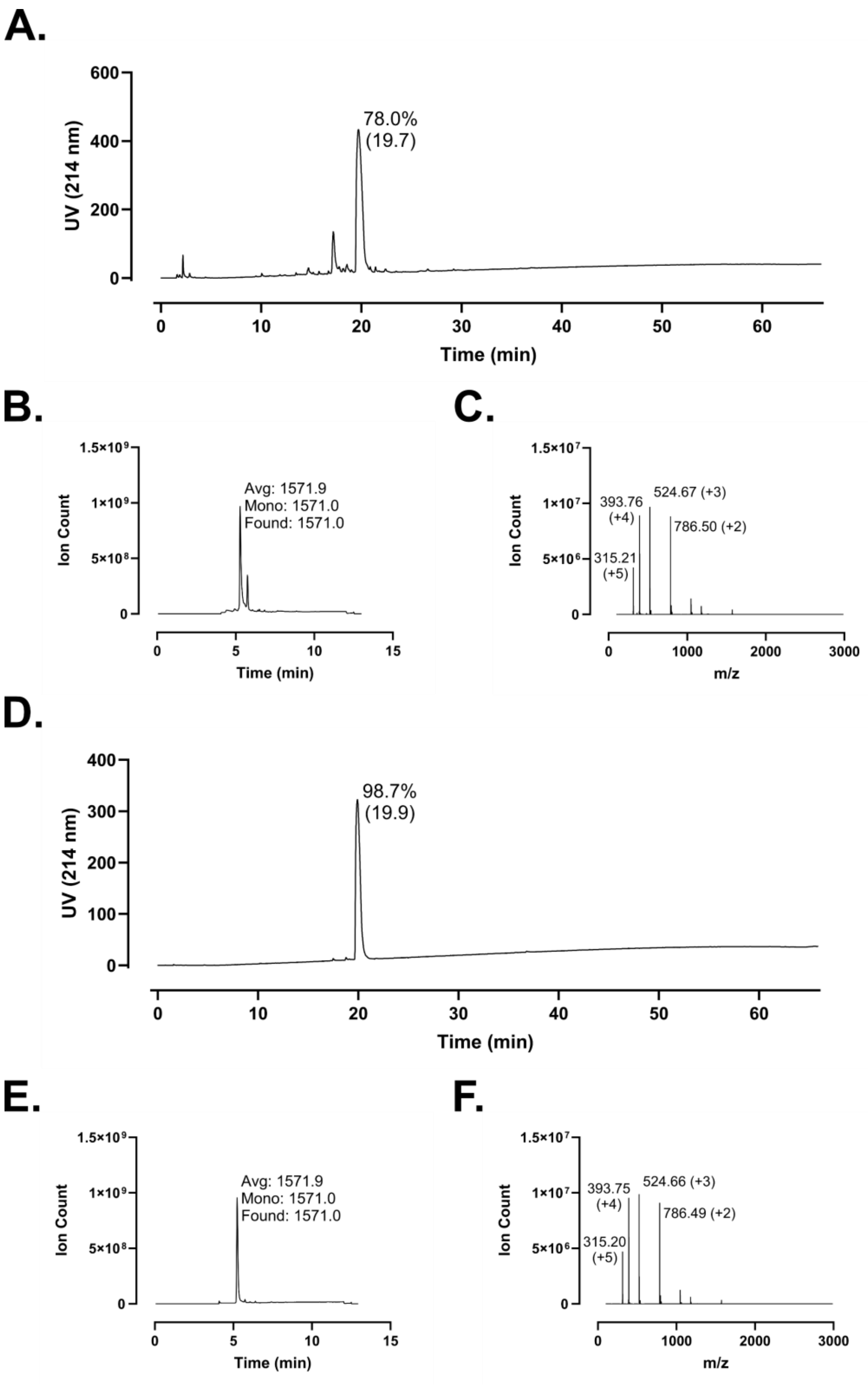
A.
UV (214 nm)
78.0%
(19.7)
Time (min)
B.
Ion Count
Avg: 1571.9
Mono: 1571.0
Found: 1571.0
Time (min)
C.
Ion Count
393.76
(+4)
524.67 (+3)
786.50 (+2)
315.21
(+5)
m/z
D.
UV (214 nm)
98.7%
(19.9)
Time (min)
E.
Ion Count
Avg: 1571.9
Mono: 1571.0
Found: 1571.0
Time (min)
F.
Ion Count
393.75
(+4)
524.66 (+3)
786.49 (+2)
315.20
(+5)
m/z

**Figure S169 – (#10) KR-12 Analytical Data. A.** HPLC trace of crude peptide at 214 nm. Peak labeling indicates percent purity by trace integration. Retention time is given in parentheses. **B.** LCMS total ion chromatogram (TIC) of crude peptide. Peak labeling indicates the predicted average and monoisotopic masses as well as the observed peptide mass. **C.** Mass spectrum at peak of **B** with *m/z* labeling for selected peaks and the associated charge in parentheses. **D.** HPLC trace of peptide following single-pass purification. **E.** TIC of peptide following single-pass purification. **F.** Mass spectrum at peak of **E**.

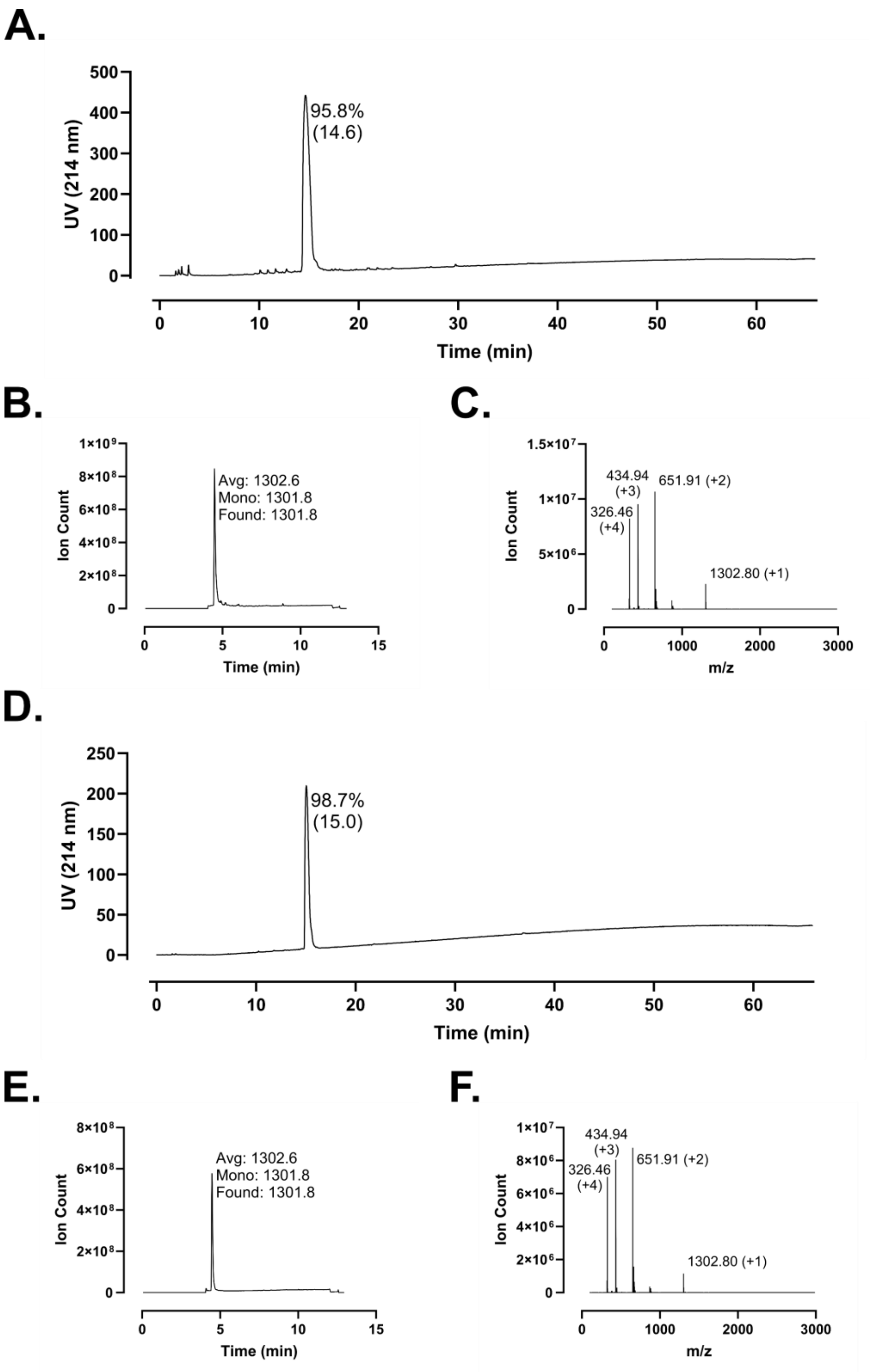
A.
UV (214 nm)
Time (min)
95.8%
(14.6)
B.
Ion Count
Time (min)
Avg: 1302.6
Mono: 1301.8
Found: 1301.8
C.
Ion Count
m/z
434.94
(+3)
651.91 (+2)
326.46
(+4)
1302.80 (+1)
D.
UV (214 nm)
Time (min)
98.7%
(15.0)
E.
Ion Count
Time (min)
Avg: 1302.6
Mono: 1301.8
Found: 1301.8
F.
Ion Count
m/z
434.94
(+3)
651.91 (+2)
326.46
(+4)
1302.80 (+1)

**Figure S170 – (#11) KR-10 Analytical Data. A.** HPLC trace of crude peptide at 214 nm. Peak labeling indicates percent purity by trace integration. Retention time is given in parentheses. **B.** LCMS total ion chromatogram (TIC) of crude peptide. Peak labeling indicates the predicted average and monoisotopic masses as well as the observed peptide mass. **C.** Mass spectrum at peak of **B** with *m/z* labeling for selected peaks and the associated charge in parentheses. **D.** HPLC trace of peptide following single-pass purification. **E.** TIC of peptide following single-pass purification. **F.** Mass spectrum at peak of **E**.

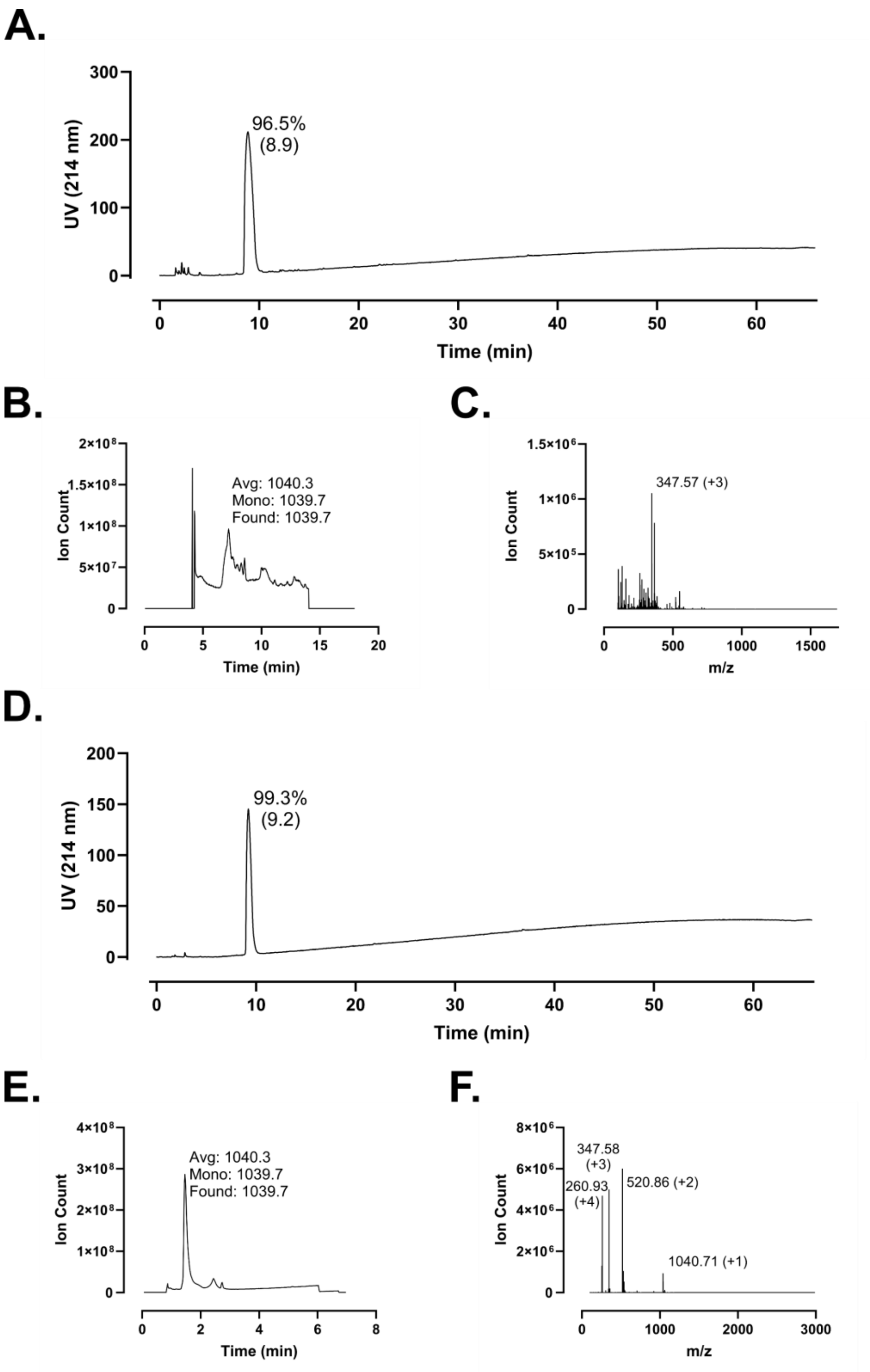
A.
300
200
100
0
UV (214 nm)
96.5%
(8.9)
0
10
20
30
40
50
60
Time (min)
B.
Ion Count
Avg: 1040.3
Mono: 1039.7
Found: 1039.7
Time (min)
C.
Ion Count
347.57 (+3)
m/z
D.
UV (214 nm)
99.3%
(9.2)
Time (min)
E.
Ion Count
Avg: 1040.3
Mono: 1039.7
Found: 1039.7
Time (min)
F.
Ion Count
347.58
(+3)
260.93
(+4)
520.86 (+2)
1040.71 (+1)
m/z

**Figure S171 – (#12) KR-8 Analytical Data. A.** HPLC trace of crude peptide at 214 nm. Peak labeling indicates percent purity by trace integration. Retention time is given in parentheses. **B.** LCMS total ion chromatogram (TIC) of crude peptide. Peak labeling indicates the predicted average and monoisotopic masses as well as the observed peptide mass. **C.** Mass spectrum at peak of **B** with *m/z* labeling for selected peaks and the associated charge in parentheses. **D.** HPLC trace of peptide following single-pass purification. **E.** TIC of peptide following single-pass purification. **F.** Mass spectrum at peak of **E**.

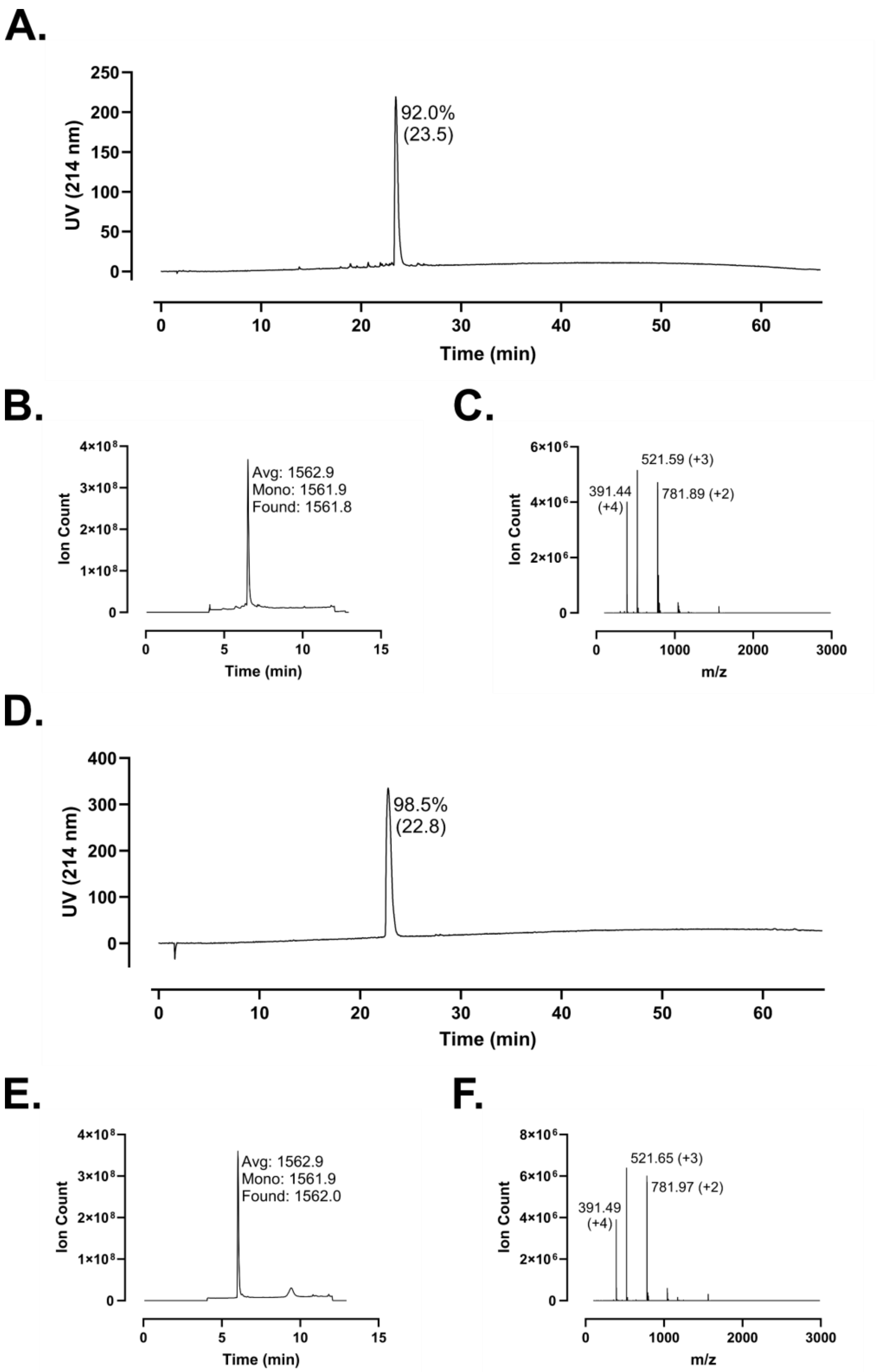
A.
UV (214 nm)
92.0%
(23.5)
Time (min)
B.
Ion Count
Avg: 1562.9
Mono: 1561.9
Found: 1561.8
Time (min)
C.
Ion Count
521.59 (+3)
391.44
(+4)
781.89 (+2)
m/z
D.
UV (214 nm)
98.5%
(22.8)
Time (min)
E.
Ion Count
Avg: 1562.9
Mono: 1561.9
Found: 1562.0
Time (min)
F.
Ion Count
521.65 (+3)
391.49
(+4)
781.97 (+2)
m/z

**Figure S172 – (#13) FK-12 Analytical Data. A.** HPLC trace of crude peptide at 214 nm. Peak labeling indicates percent purity by trace integration. Retention time is given in parentheses. **B.** LCMS total ion chromatogram (TIC) of crude peptide. Peak labeling indicates the predicted average and monoisotopic masses as well as the observed peptide mass. **C.** Mass spectrum at peak of **B** with *m/z* labeling for selected peaks and the associated charge in parentheses. **D.** HPLC trace of peptide following single-pass purification. **E.** TIC of peptide following single-pass purification. **F.** Mass spectrum at peak of **E**.

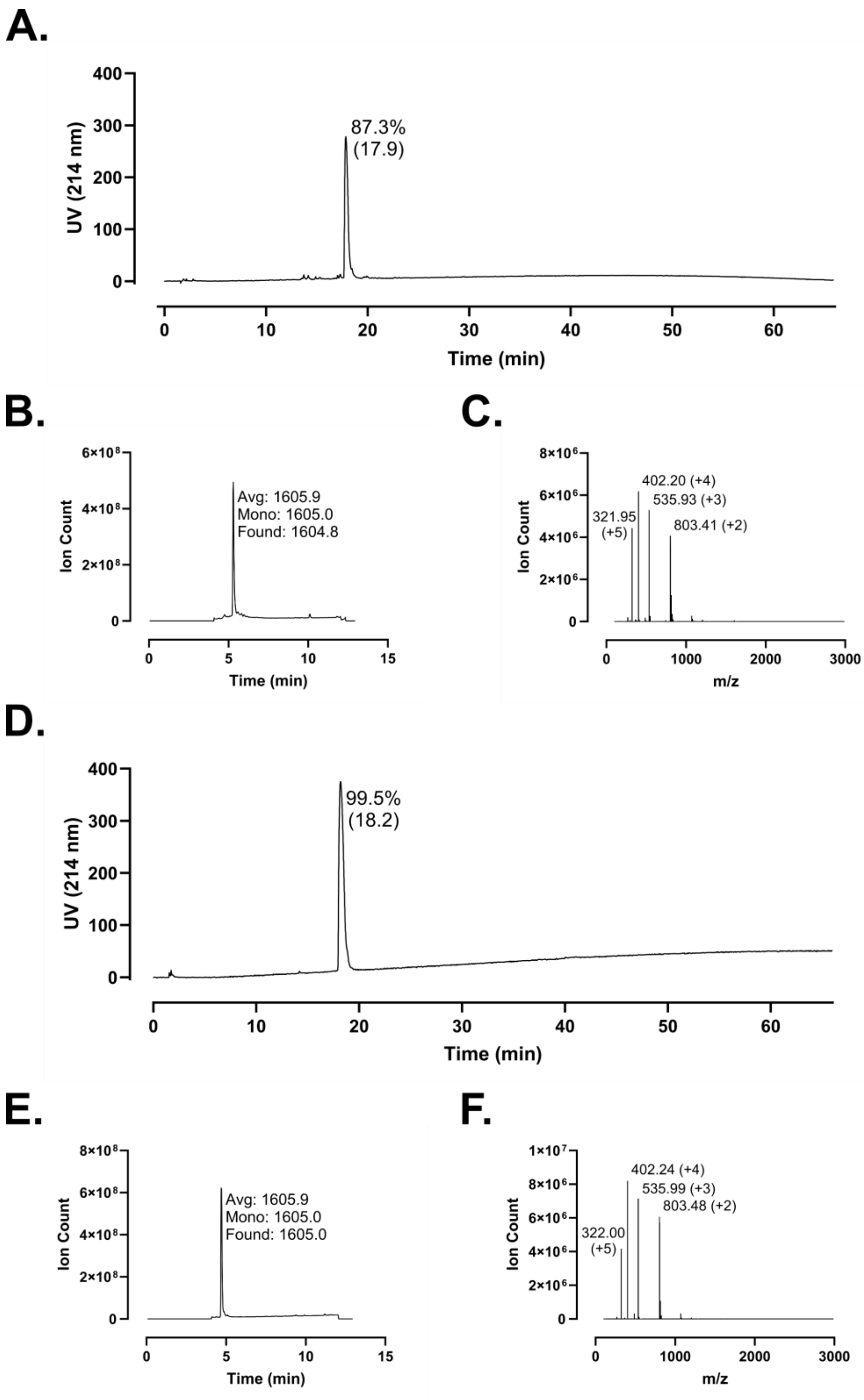
A.
UV (214 nm)
87.3%
(17.9)
Time (min)
B.
Ion Count
Avg: 1605.9
Mono: 1605.0
Found: 1604.8
Time (min)
C.
Ion Count
402.20 (+4)
535.93 (+3)
321.95
(+5)
803.41 (+2)
m/z
D.
UV (214 nm)
99.5%
(18.2)
Time (min)
E.
Ion Count
Avg: 1605.9
Mono: 1605.0
Found: 1605.0
Time (min)
F.
Ion Count
402.24 (+4)
535.99 (+3)
803.48 (+2)
322.00
(+5)
m/z

**Figure S173 – (#14) FK-12-R Analytical Data. A.** HPLC trace of crude peptide at 214 nm. Peak labeling indicates percent purity by trace integration. Retention time is given in parentheses. **B.** LCMS total ion chromatogram (TIC) of crude peptide. Peak labeling indicates the predicted average and monoisotopic masses as well as the observed peptide mass. **C.** Mass spectrum at peak of **B** with *m/z* labeling for selected peaks and the associated charge in parentheses. **D.** HPLC trace of peptide following single-pass purification. **E.** TIC of peptide following single-pass purification. **F.** Mass spectrum at peak of **E**.

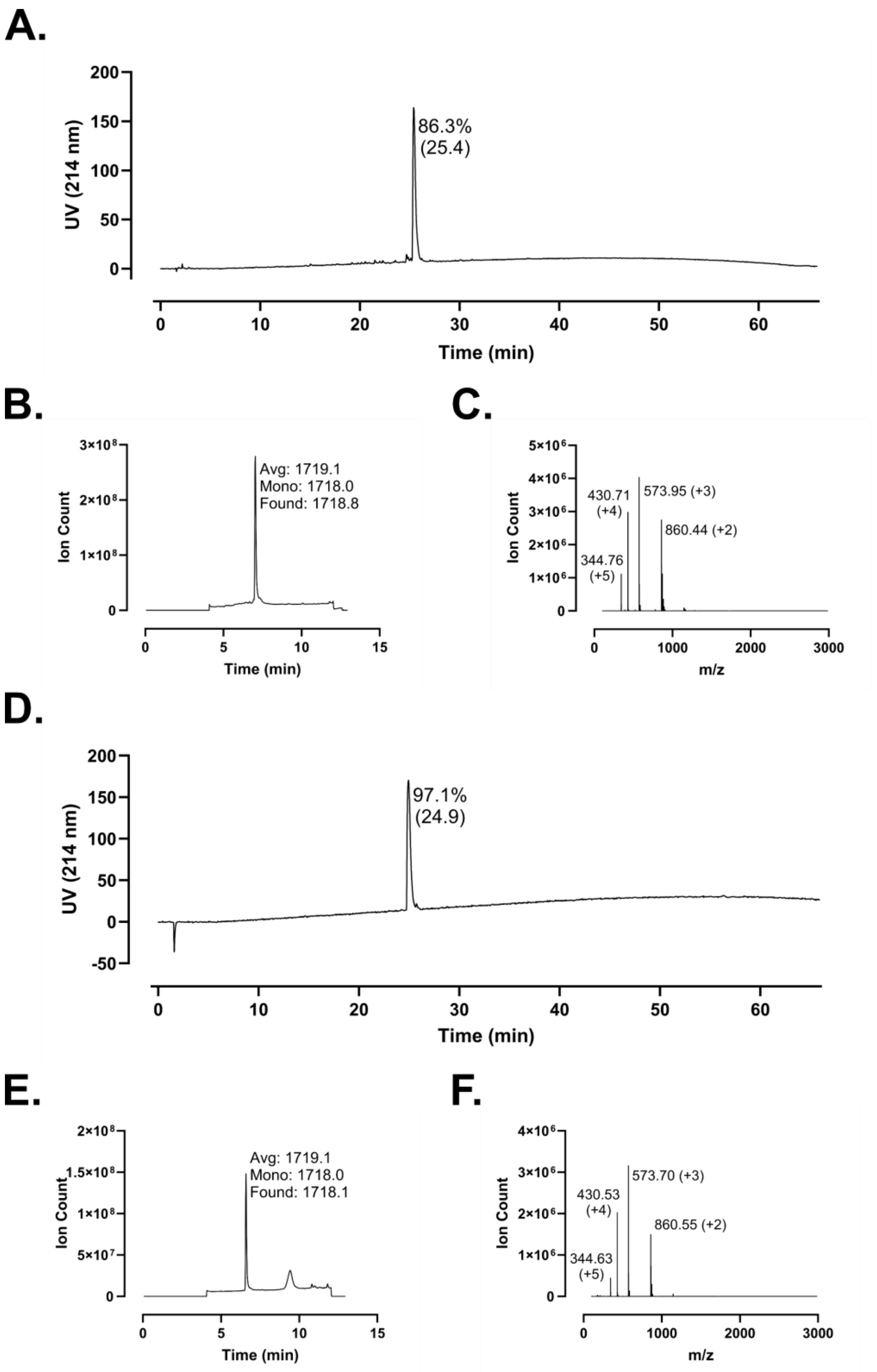
A.
200
150
100
50
0
UV (214 nm)
86.3%
(25.4)
0
10
20
30
40
50
60
Time (min)
B.
3×10⁸
2×10⁸
1×10⁸
0
Ion Count
Avg: 1719.1
Mono: 1718.0
Found: 1718.8
0
5
10
15
Time (min)
C.
5×10⁶
4×10⁶
3×10⁶
2×10⁶
1×10⁶
0
Ion Count
430.71
(+4)
573.95 (+3)
860.44 (+2)
344.76
(+5)
0
1000
2000
3000
m/z
D.
200
150
100
50
0
-50
UV (214 nm)
97.1%
(24.9)
0
10
20
30
40
50
60
Time (min)
E.
2×10⁸
1.5×10⁸
1×10⁸
5×10⁷
0
Ion Count
Avg: 1719.1
Mono: 1718.0
Found: 1718.1
0
5
10
15
Time (min)
F.
4×10⁶
3×10⁶
2×10⁶
1×10⁶
0
Ion Count
430.53
(+4)
573.70 (+3)
860.55 (+2)
344.63
(+5)
0
1000
2000
3000
m/z

**Figure S174 – (#15) Retro-FK-13 Analytical Data. A.** HPLC trace of crude peptide at 214 nm. Peak labeling indicates percent purity by trace integration. Retention time is given in parentheses. **B.** LCMS total ion chromatogram (TIC) of crude peptide. Peak labeling indicates the predicted average and monoisotopic masses as well as the observed peptide mass. **C.** Mass spectrum at peak of **B** with *m/z* labeling for selected peaks and the associated charge in parentheses. **D.** HPLC trace of peptide following single-pass purification. **E.** TIC of peptide following single-pass purification. **F.** Mass spectrum at peak of **E**.

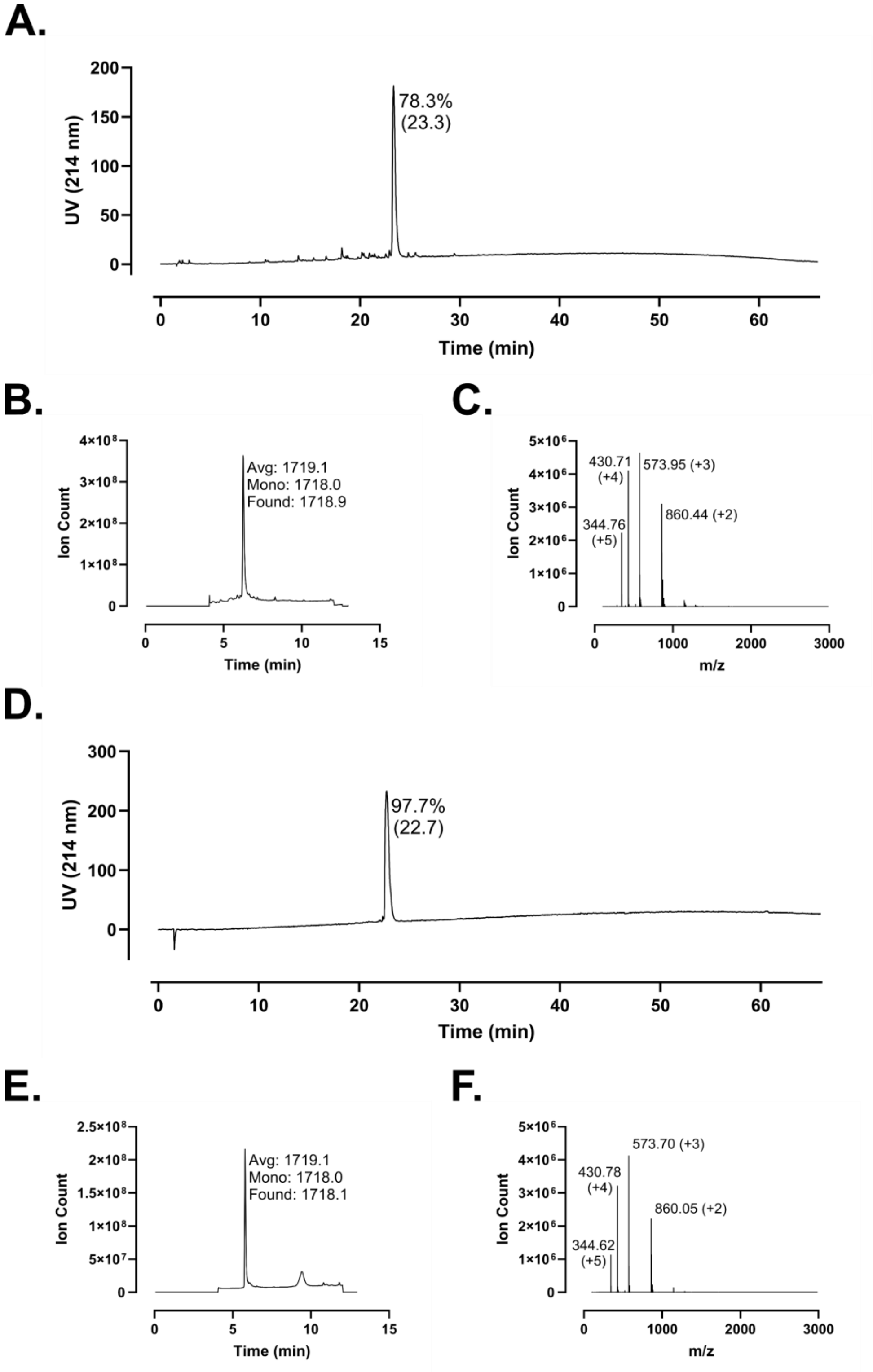
A.
78.3%
(23.3)
UV (214 nm)
Time (min)
B.
Avg: 1719.1
Mono: 1718.0
Found: 1718.9
Ion Count
Time (min)
C.
430.71
(+4)
573.95 (+3)
344.76
(+5)
860.44 (+2)
Ion Count
m/z
D.
97.7%
(22.7)
UV (214 nm)
Time (min)
E.
Avg: 1719.1
Mono: 1718.0
Found: 1718.1
Ion Count
Time (min)
F.
573.70 (+3)
430.78
(+4)
860.05 (+2)
344.62
(+5)
Ion Count
m/z

**Figure S175 – (#16) fk-13 Analytical Data. A.** HPLC trace of crude peptide at 214 nm. Peak labeling indicates percent purity by trace integration. Retention time is given in parentheses. **B.** LCMS total ion chromatogram (TIC) of crude peptide. Peak labeling indicates the predicted average and monoisotopic masses as well as the observed peptide mass. **C.** Mass spectrum at peak of **B** with *m/z* labeling for selected peaks and the associated charge in parentheses. **D.** HPLC trace of peptide following single-pass purification. **E.** TIC of peptide following single-pass purification. **F.** Mass spectrum at peak of **E**.

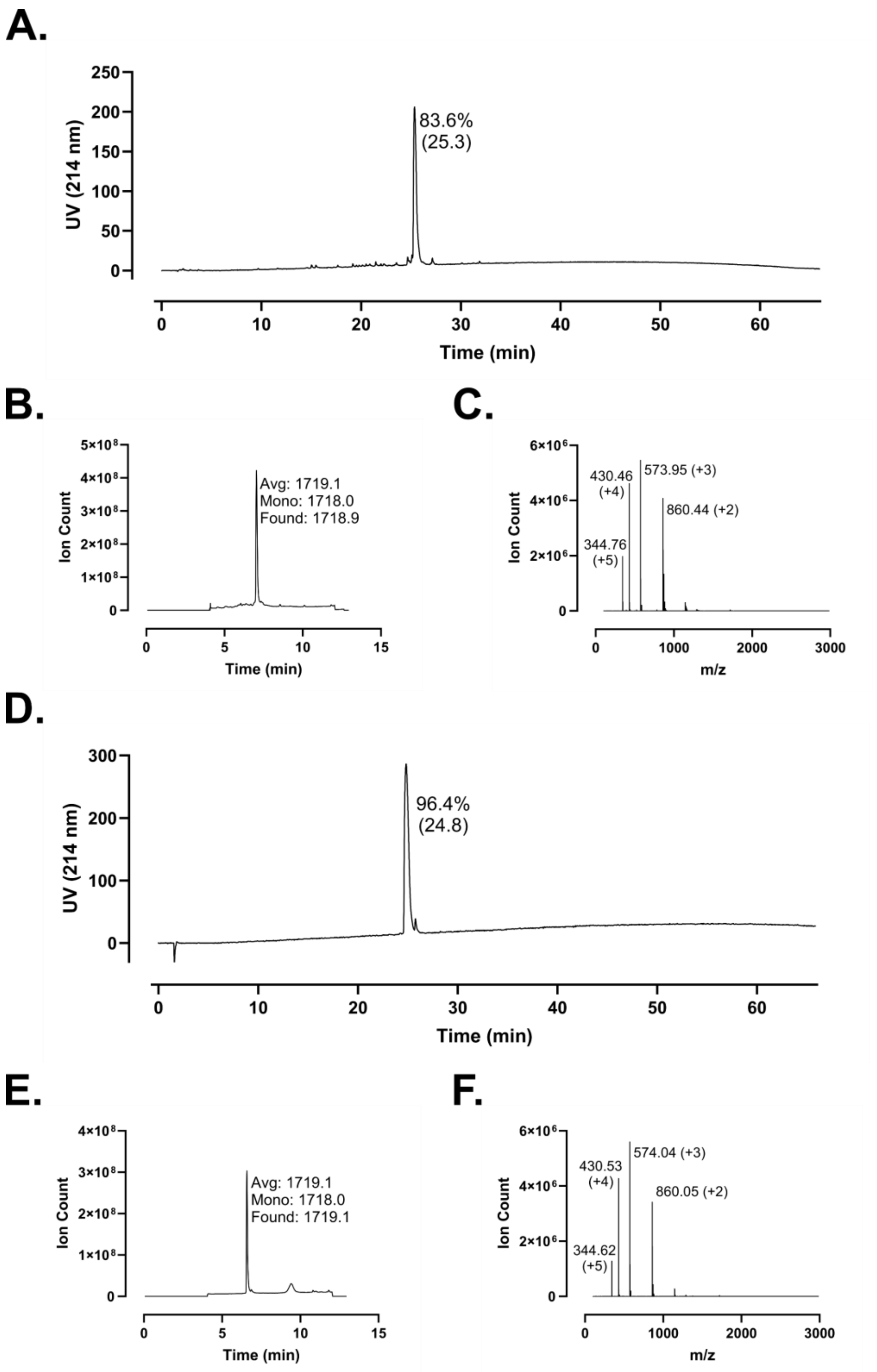
A.
250
200
150
100
50
0
UV (214 nm)
83.6%
(25.3)
0
10
20
30
40
50
60
Time (min)
B.
5×10⁸
4×10⁸
3×10⁸
2×10⁸
1×10⁸
0
Ion Count
Avg: 1719.1
Mono: 1718.0
Found: 1718.9
0
5
10
15
Time (min)
C.
6×10⁶
4×10⁶
2×10⁶
0
Ion Count
430.46
(+4)
573.95 (+3)
860.44 (+2)
344.76
(+5)
0
1000
2000
3000
m/z
D.
300
200
100
0
UV (214 nm)
96.4%
(24.8)
0
10
20
30
40
50
60
Time (min)
E.
4×10⁸
3×10⁸
2×10⁸
1×10⁸
0
Ion Count
Avg: 1719.1
Mono: 1718.0
Found: 1719.1
0
5
10
15
Time (min)
F.
6×10⁶
4×10⁶
2×10⁶
0
Ion Count
430.53
(+4)
574.04 (+3)
860.05 (+2)
344.62
(+5)
0
1000
2000
3000
m/z

**Figure S176 – (#17) Retro-fk-13 Analytical Data. A.** HPLC trace of crude peptide at 214 nm. Peak labeling indicates percent purity by trace integration. Retention time is given in parentheses. **B.** LCMS total ion chromatogram (TIC) of crude peptide. Peak labeling indicates the predicted average and monoisotopic masses as well as the observed peptide mass. **C.** Mass spectrum at peak of **B** with *m/z* labeling for selected peaks and the associated charge in parentheses. **D.** HPLC trace of peptide following single-pass purification. **E.** TIC of peptide following single-pass purification. **F.** Mass spectrum at peak of **E**.

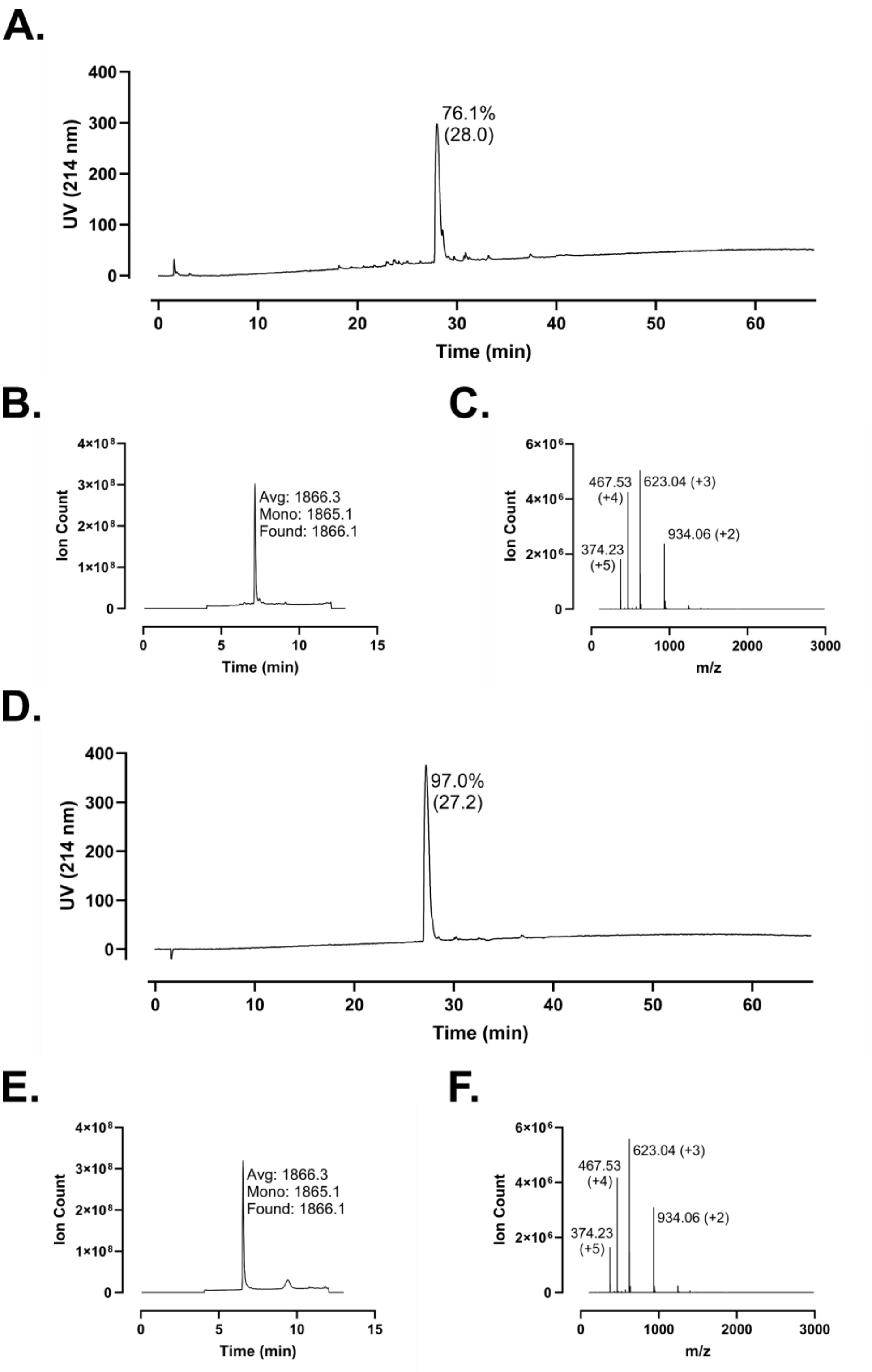
A.
76.1%
(28.0)
UV (214 nm)
Time (min)
B.
Avg: 1866.3
Mono: 1865.1
Found: 1866.1
Ion Count
Time (min)
C.
467.53
(+4)
623.04 (+3)
934.06 (+2)
374.23
(+5)
Ion Count
m/z
D.
97.0%
(27.2)
UV (214 nm)
Time (min)
E.
Avg: 1866.3
Mono: 1865.1
Found: 1866.1
Ion Count
Time (min)
F.
467.53
(+4)
623.04 (+3)
934.06 (+2)
374.23
(+5)
Ion Count
m/z

**Figure S177 – (#18) FF-14 Analytical Data. A.** HPLC trace of crude peptide at 214 nm. Peak labeling indicates percent purity by trace integration. Retention time is given in parentheses. **B.** LCMS total ion chromatogram (TIC) of crude peptide. Peak labeling indicates the predicted average and monoisotopic masses as well as the observed peptide mass. **C.** Mass spectrum at peak of **B** with *m/z* labeling for selected peaks and the associated charge in parentheses. **D.** HPLC trace of peptide following single-pass purification. **E.** TIC of peptide following single-pass purification. **F.** Mass spectrum at peak of **E**.

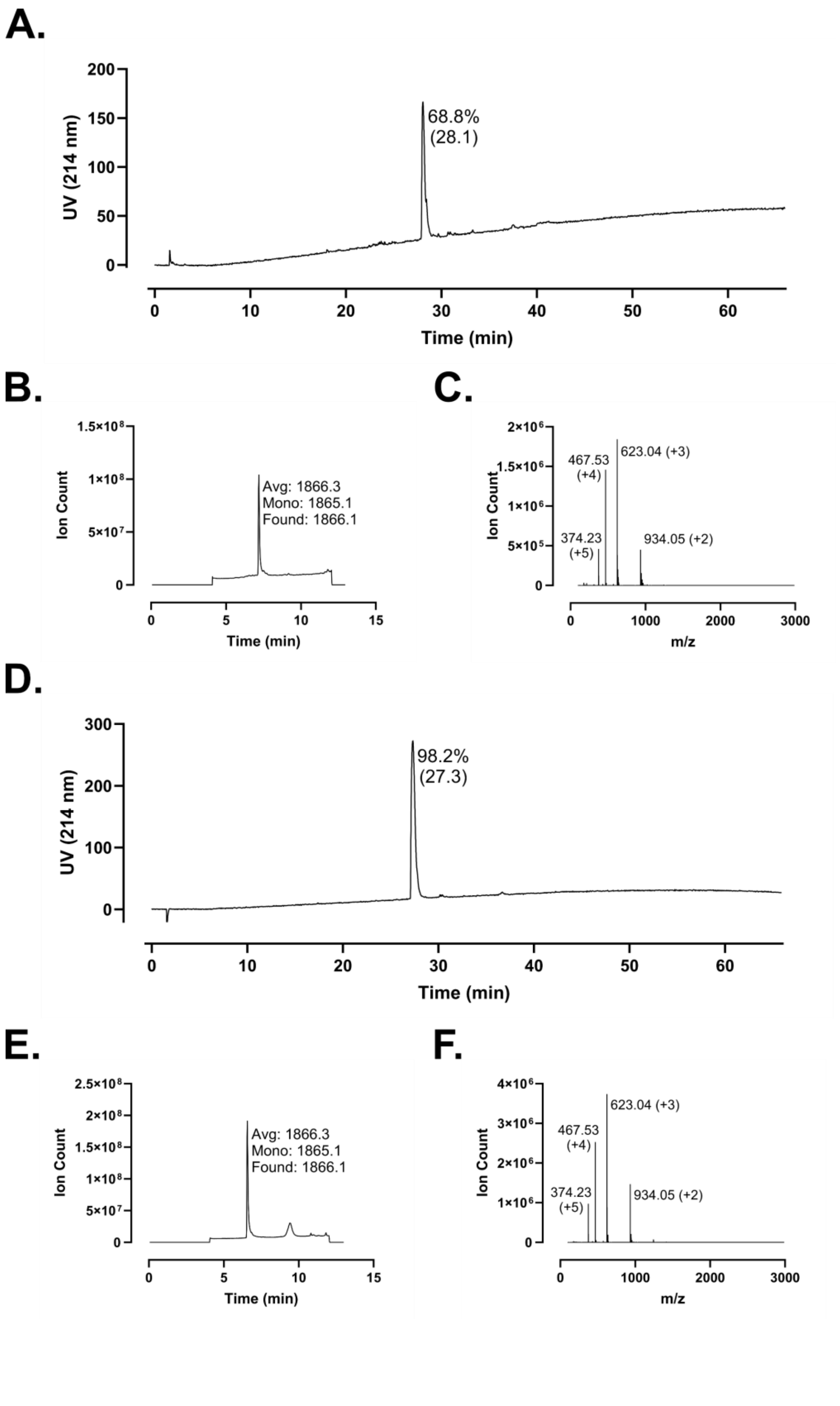
A.
UV (214 nm)
200
150
100
50
0
68.8%
(28.1)
0
10
20
30
40
50
60
Time (min)
B.
Ion Count
1.5×10^8
1×10^8
5×10^7
0
Avg: 1866.3
Mono: 1865.1
Found: 1866.1
0
5
10
15
Time (min)
C.
Ion Count
2×10^6
1.5×10^6
1×10^6
5×10^5
0
467.53
(+4)
623.04 (+3)
374.23
(+5)
934.05 (+2)
0
1000
2000
3000
m/z
D.
UV (214 nm)
300
200
100
0
98.2%
(27.3)
0
10
20
30
40
50
60
Time (min)
E.
Ion Count
2.5×10^8
2×10^8
1.5×10^8
1×10^8
5×10^7
0
Avg: 1866.3
Mono: 1865.1
Found: 1866.1
0
5
10
15
Time (min)
F.
Ion Count
4×10^6
3×10^6
2×10^6
1×10^6
0
623.04 (+3)
467.53
(+4)
374.23
(+5)
934.05 (+2)
0
1000
2000
3000
m/z

**Figure S178 – (#19) FF-14-RK Analytical Data. A.** HPLC trace of crude peptide at 214 nm. Peak labeling indicates percent purity by trace integration. Retention time is given in parentheses. **B.** LCMS total ion chromatogram (TIC) of crude peptide. Peak labeling indicates the predicted average and monoisotopic masses as well as the observed peptide mass. **C.** Mass spectrum at peak of **B** with *m/z* labeling for selected peaks and the associated charge in parentheses. **D.** HPLC trace of peptide following single-pass purification. **E.** TIC of peptide following single-pass purification. **F.** Mass spectrum at peak of **E**.

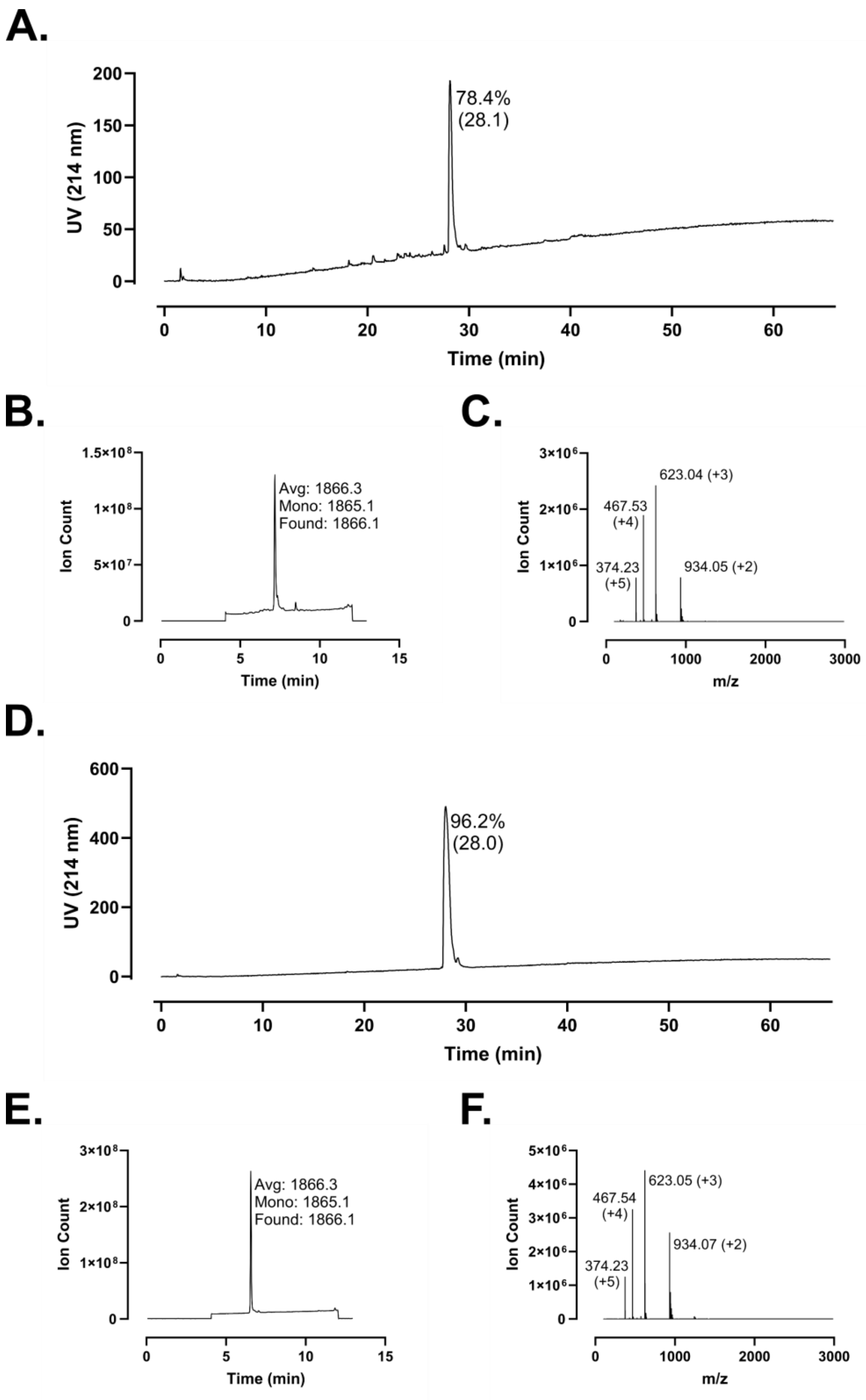
A.
78.4%
(28.1)
UV (214 nm)
Time (min)
B.
Avg: 1866.3
Mono: 1865.1
Found: 1866.1
Ion Count
Time (min)
C.
623.04 (+3)
467.53
(+4)
374.23
(+5)
934.05 (+2)
Ion Count
m/z
D.
96.2%
(28.0)
UV (214 nm)
Time (min)
E.
Avg: 1866.3
Mono: 1865.1
Found: 1866.1
Ion Count
Time (min)
F.
623.05 (+3)
467.54
(+4)
934.07 (+2)
374.23
(+5)
Ion Count
m/z

**Figure S179 – (#20) ff-14 Analytical Data. A.** HPLC trace of crude peptide at 214 nm. Peak labeling indicates percent purity by trace integration. Retention time is given in parentheses. **B.** LCMS total ion chromatogram (TIC) of crude peptide. Peak labeling indicates the predicted average and monoisotopic masses as well as the observed peptide mass. **C.** Mass spectrum at peak of **B** with *m/z* labeling for selected peaks and the associated charge in parentheses. **D.** HPLC trace of peptide following single-pass purification. **E.** TIC of peptide following single-pass purification. **F.** Mass spectrum at peak of **E**.

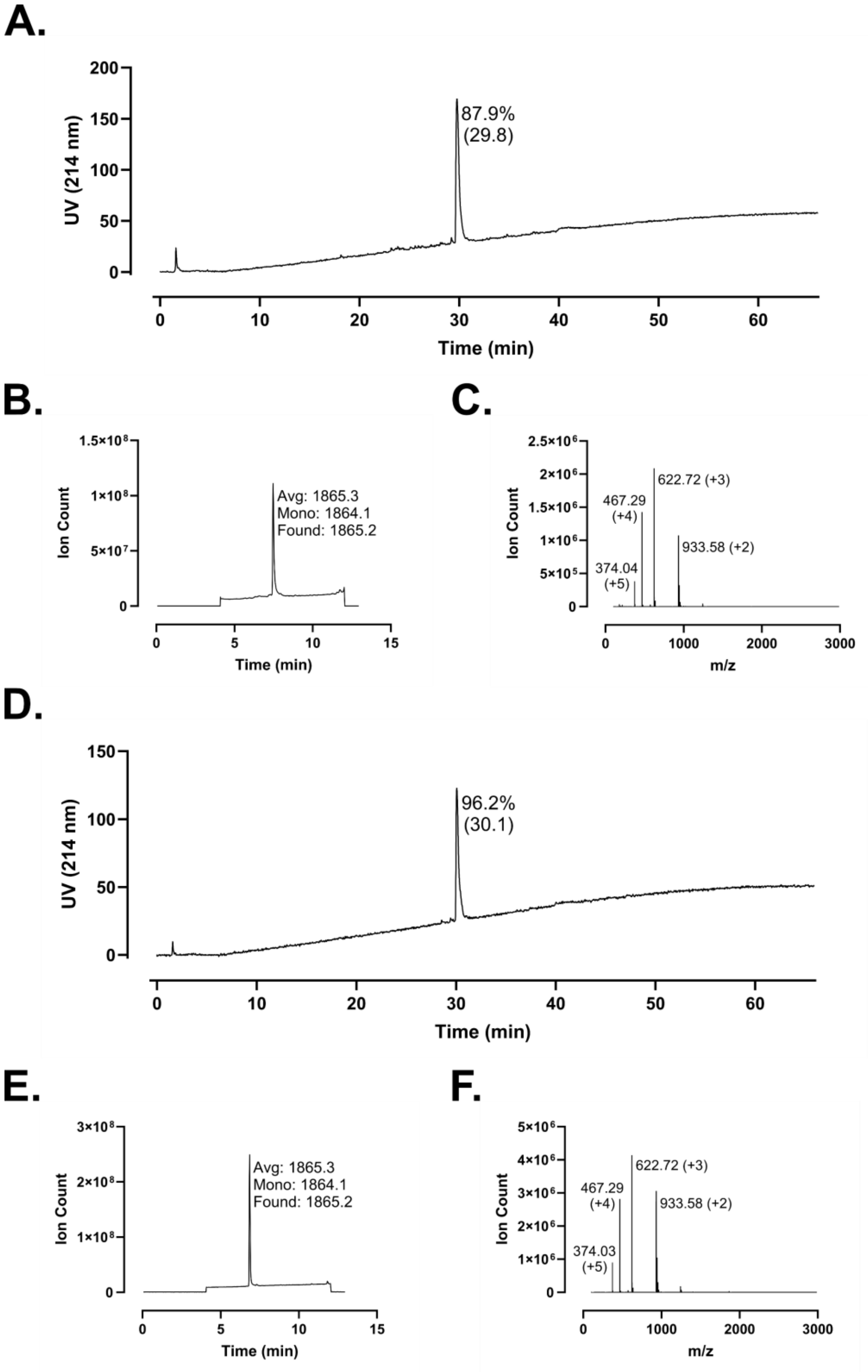
A.
87.9%
(29.8)
UV (214 nm)
Time (min)
B.
Avg: 1865.3
Mono: 1864.1
Found: 1865.2
Ion Count
Time (min)
C.
622.72 (+3)
467.29
(+4)
933.58 (+2)
374.04
(+5)
Ion Count
m/z
D.
96.2%
(30.1)
UV (214 nm)
Time (min)
E.
Avg: 1865.3
Mono: 1864.1
Found: 1865.2
Ion Count
Time (min)
F.
622.72 (+3)
467.29
(+4)
933.58 (+2)
374.03
(+5)
Ion Count
m/z

**Figure S180 – (#22) FF-14-$NH_2$ Analytical Data. A.** HPLC trace of crude peptide at 214 nm. Peak labeling indicates percent purity by trace integration. Retention time is given in parentheses. **B.** LCMS total ion chromatogram (TIC) of crude peptide. Peak labeling indicates the predicted average and monoisotopic masses as well as the observed peptide mass. **C.** Mass spectrum at peak of **B** with *m/z* labeling for selected peaks and the associated charge in parentheses. **D.** HPLC trace of peptide following single-pass purification. **E.** TIC of peptide following single-pass purification. **F.** Mass spectrum at peak of **E**.

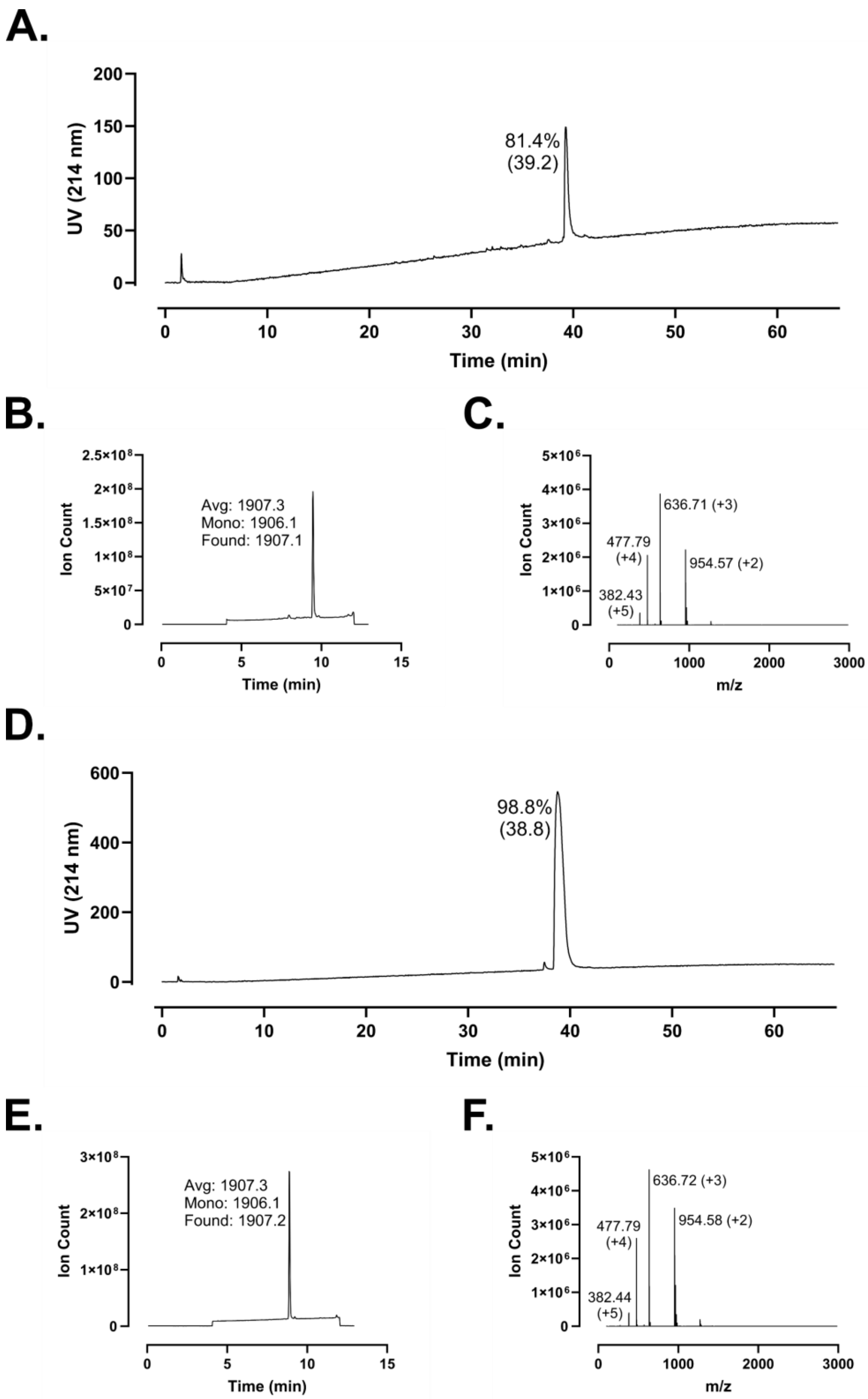
A.
UV (214 nm)
81.4%
(39.2)
Time (min)
B.
Ion Count
Avg: 1907.3
Mono: 1906.1
Found: 1907.1
Time (min)
C.
Ion Count
636.71 (+3)
477.79
(+4)
954.57 (+2)
382.43
(+5)
m/z
D.
UV (214 nm)
98.8%
(38.8)
Time (min)
E.
Ion Count
Avg: 1907.3
Mono: 1906.1
Found: 1907.2
Time (min)
F.
Ion Count
636.72 (+3)
477.79
(+4)
954.58 (+2)
382.44
(+5)
m/z

**Figure S181 – (#22) Ac-FF-14-$NH_2$ Analytical Data. A.** HPLC trace of crude peptide at 214 nm. Peak labeling indicates percent purity by trace integration. Retention time is given in parentheses. **B.** LCMS total ion chromatogram (TIC) of crude peptide. Peak labeling indicates the predicted average and monoisotopic masses as well as the observed peptide mass. **C.** Mass spectrum at peak of **B** with *m/z* labeling for selected peaks and the associated charge in parentheses. **D.** HPLC trace of peptide following single-pass purification. **E.** TIC of peptide following single-pass purification. **F.** Mass spectrum at peak of **E**.

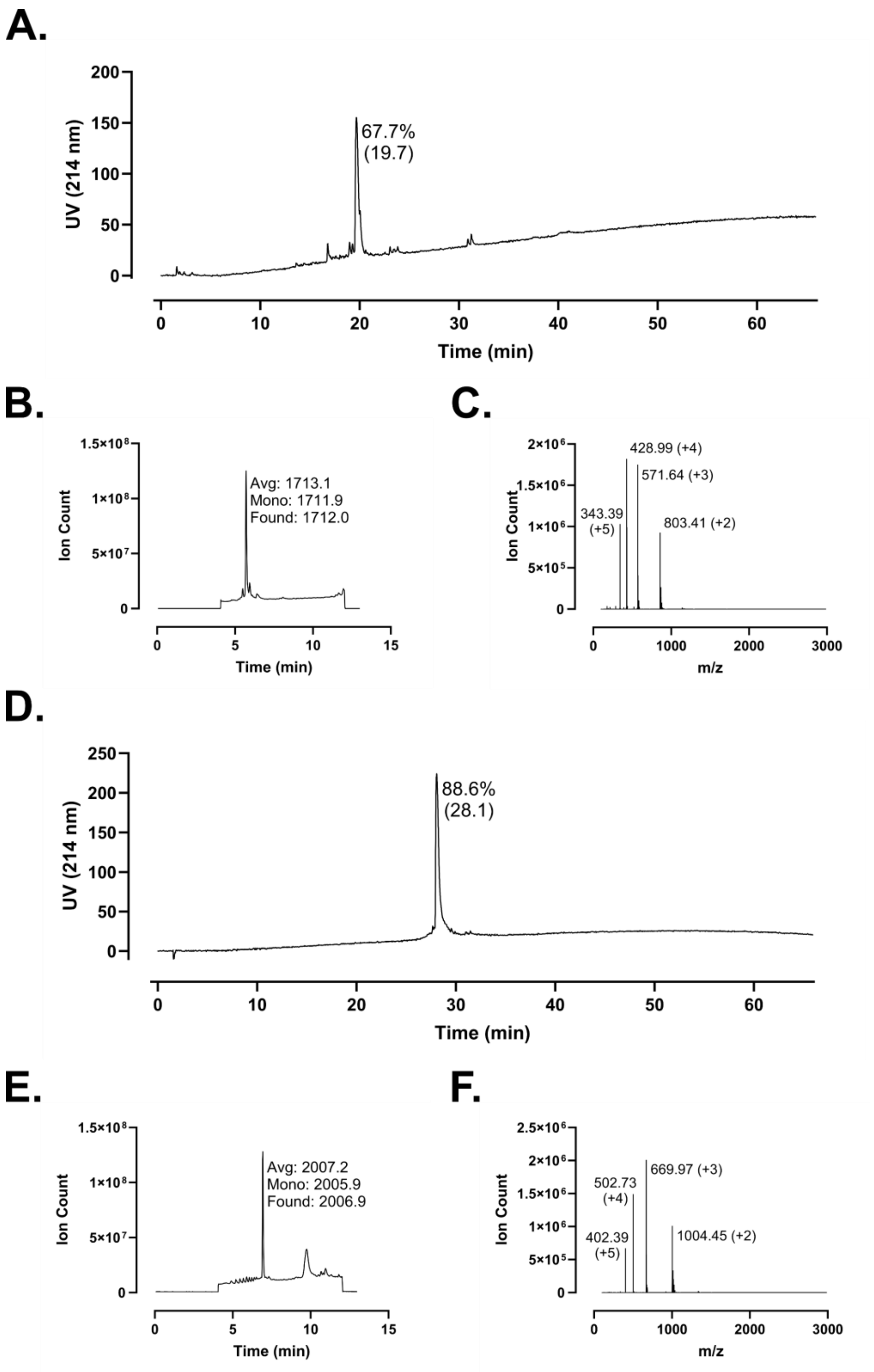
A.
UV (214 nm)
67.7%
(19.7)
Time (min)
B.
Ion Count
Avg: 1713.1
Mono: 1711.9
Found: 1712.0
Time (min)
C.
Ion Count
428.99 (+4)
571.64 (+3)
343.39
(+5)
803.41 (+2)
m/z
D.
UV (214 nm)
88.6%
(28.1)
Time (min)
E.
Ion Count
Avg: 2007.2
Mono: 2005.9
Found: 2006.9
Time (min)
F.
Ion Count
669.97 (+3)
502.73
(+4)
402.39
(+5)
1004.45 (+2)
m/z

**Figure S182 – (#23) FK-13 V6+L13 Perfluoroaryl Staple Analytical Data. A.** HPLC trace of crude peptide (FK-13 V6C+L13C) at 214 nm. Peak labeling indicates percent purity by trace integration. Retention time is given in parentheses. **B.** LCMS total ion chromatogram (TIC) of crude peptide. Peak labeling indicates the predicted average and monoisotopic masses as well as the observed peptide mass. **C.** Mass spectrum at peak of **B** with *m/z* labeling for selected peaks and the associated charge in parentheses. **D.** HPLC trace of perfluoroaryl stapled peptide following single-pass purification. **E.** TIC of peptide following single-pass purification. **F.** Mass spectrum at peak of **E**.

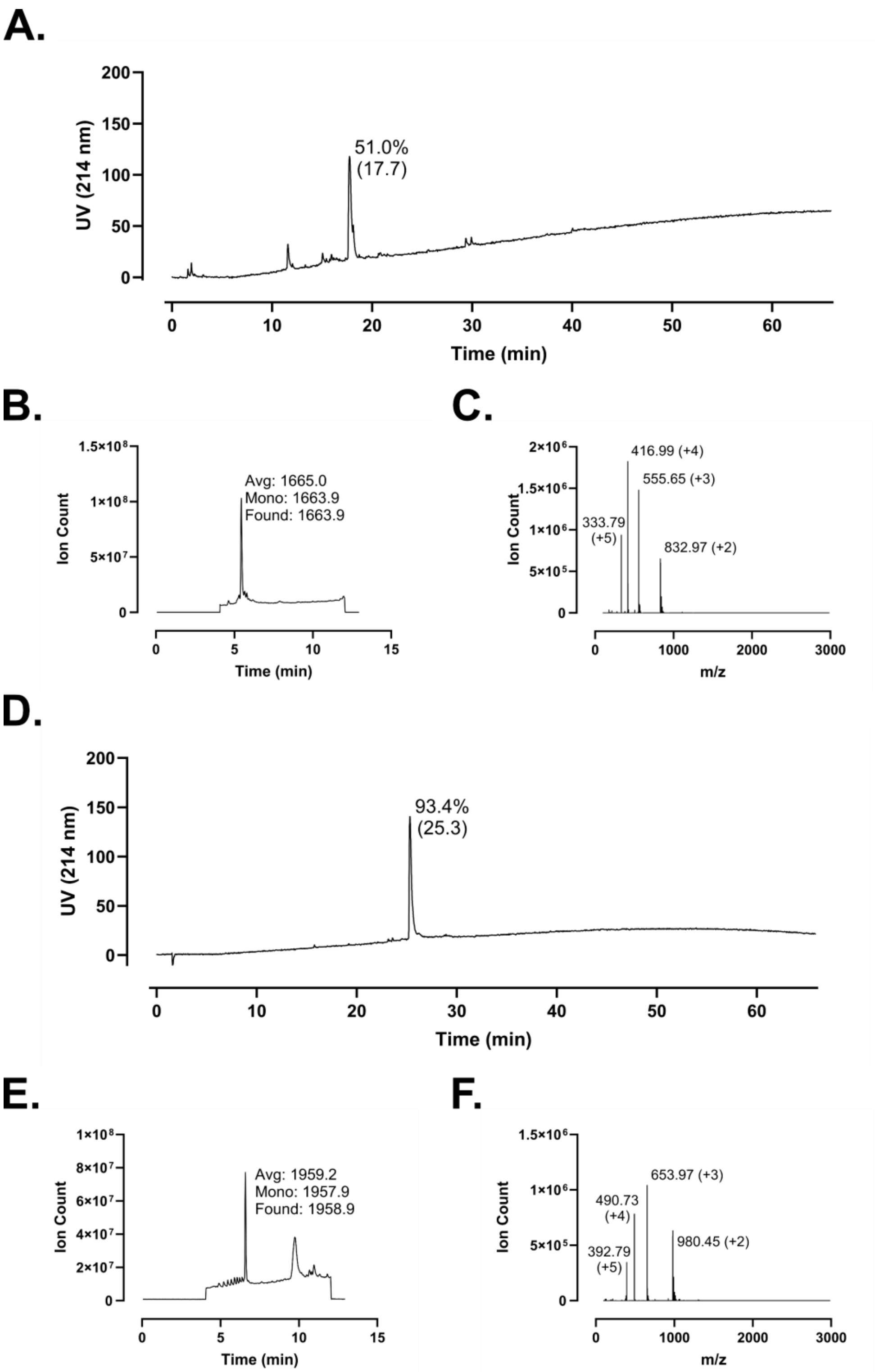
A.
UV (214 nm)
Time (min)
51.0%
(17.7)
B.
Ion Count
Time (min)
Avg: 1665.0
Mono: 1663.9
Found: 1663.9
C.
Ion Count
m/z
416.99 (+4)
555.65 (+3)
333.79
(+5)
832.97 (+2)
D.
UV (214 nm)
Time (min)
93.4%
(25.3)
E.
Ion Count
Time (min)
Avg: 1959.2
Mono: 1957.9
Found: 1958.9
F.
Ion Count
m/z
653.97 (+3)
490.73
(+4)
392.79
(+5)
980.45 (+2)

**Figure S183 – (#24) FK-13 I5+F12 Perfluoroaryl Staple Analytical Data. A.** HPLC trace of crude peptide (FK-13 I5C+F12C) at 214 nm. Peak labeling indicates percent purity by trace integration. Retention time is given in parentheses. **B.** LCMS total ion chromatogram (TIC) of crude peptide. Peak labeling indicates the predicted average and monoisotopic masses as well as the observed peptide mass. **C.** Mass spectrum at peak of **B** with *m/z* labeling for selected peaks and the associated charge in parentheses. **D.** HPLC trace of perfluoroaryl stapled peptide following single-pass purification. **E.** TIC of peptide following single-pass purification. **F.** Mass spectrum at peak of **E**.

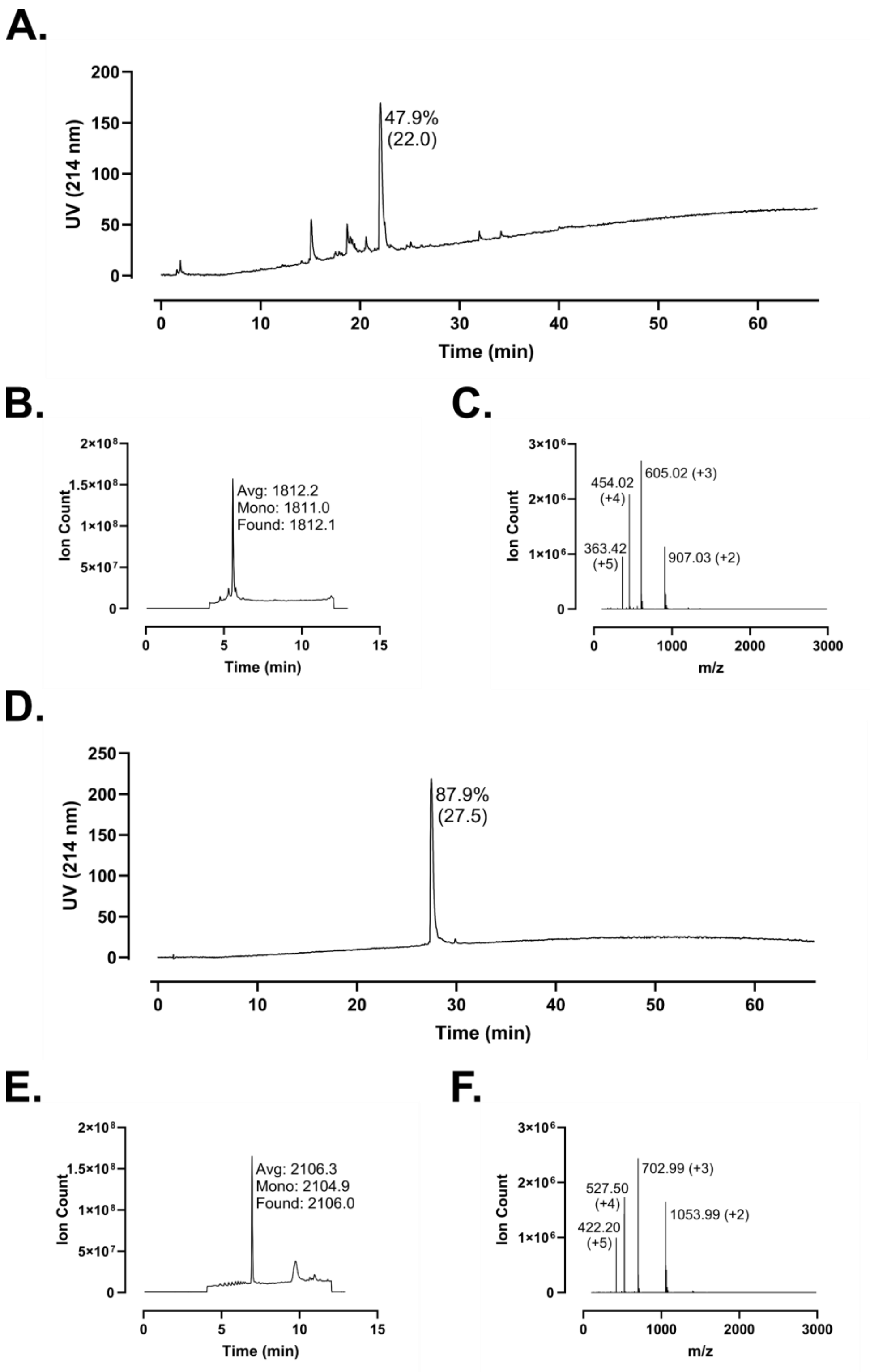
A.
UV (214 nm)
Time (min)
47.9%
(22.0)
B.
Ion Count
Time (min)
Avg: 1812.2
Mono: 1811.0
Found: 1812.1
C.
Ion Count
m/z
454.02
(+4)
605.02 (+3)
363.42
(+5)
907.03 (+2)
D.
UV (214 nm)
Time (min)
87.9%
(27.5)
E.
Ion Count
Time (min)
Avg: 2106.3
Mono: 2104.9
Found: 2106.0
F.
Ion Count
m/z
527.50
(+4)
702.99 (+3)
422.20
(+5)
1053.99 (+2)

**Figure S184 – (#25) FF-14 I5+F12 Perfluoroaryl Staple Analytical Data. A.** HPLC trace of crude peptide (FF-14 I5C+F12C) at 214 nm. Peak labeling indicates percent purity by trace integration. Retention time is given in parentheses. **B.** LCMS total ion chromatogram (TIC) of crude peptide. Peak labeling indicates the predicted average and monoisotopic masses as well as the observed peptide mass. **C.** Mass spectrum at peak of **B** with *m/z* labeling for selected peaks and the associated charge in parentheses. **D.** HPLC trace of perfluoroaryl stapled peptide following single-pass purification. **E.** TIC of peptide following single-pass purification. **F.** Mass spectrum at peak of **E**.

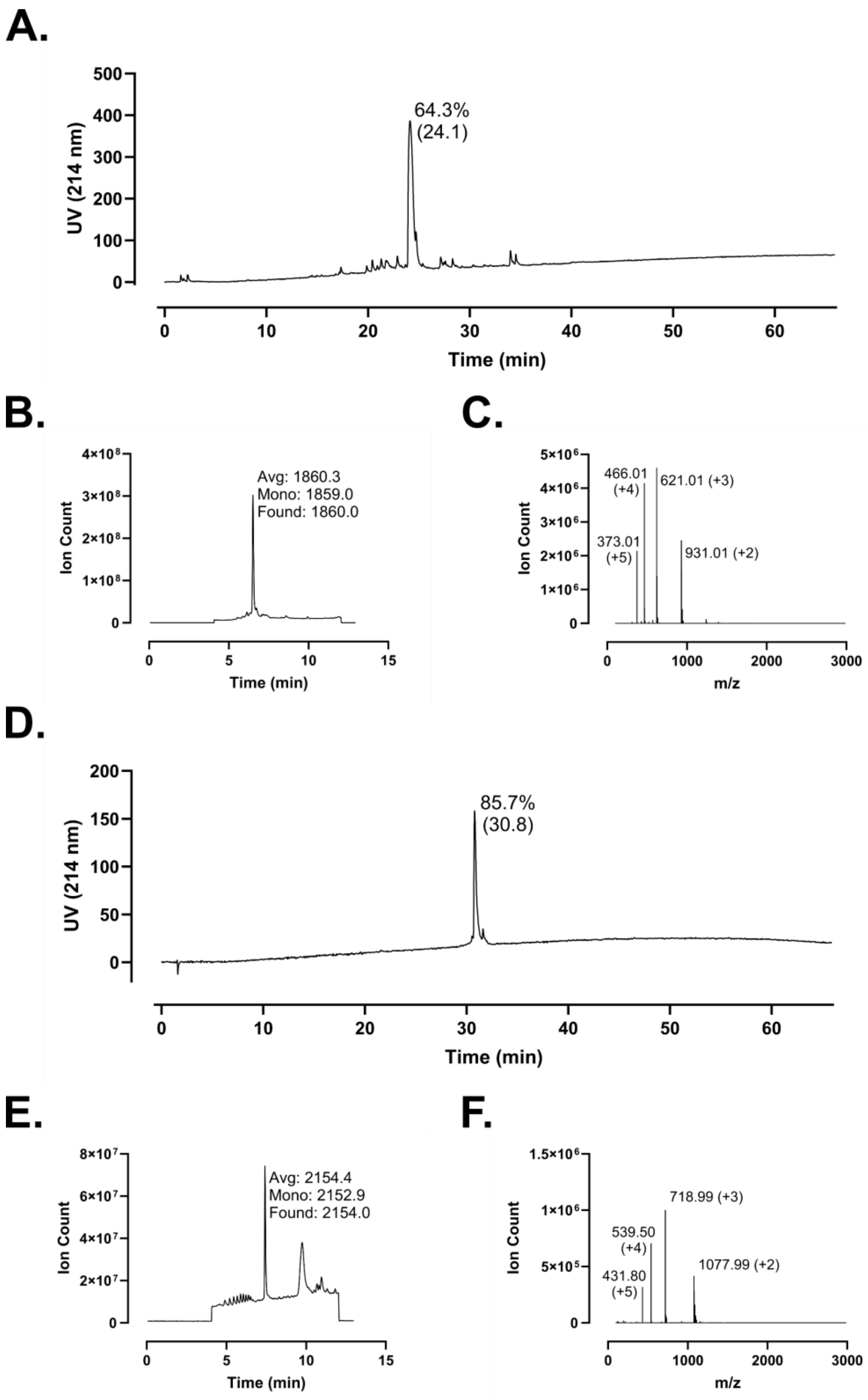
A.
UV (214 nm)
Time (min)
64.3%
(24.1)
B.
Ion Count
Time (min)
Avg: 1860.3
Mono: 1859.0
Found: 1860.0
C.
Ion Count
m/z
466.01
(+4)
621.01 (+3)
373.01
(+5)
931.01 (+2)
D.
UV (214 nm)
Time (min)
85.7%
(30.8)
E.
Ion Count
Time (min)
Avg: 2154.4
Mono: 2152.9
Found: 2154.0
F.
Ion Count
m/z
718.99 (+3)
539.50
(+4)
431.80
(+5)
1077.99 (+2)

**Figure S185 – (#26) FF-14 V6+L13 Perfluoroaryl Staple Analytical Data. A.** HPLC trace of crude peptide (FF-14 V6C+L13C) at 214 nm. Peak labeling indicates percent purity by trace integration. Retention time is given in parentheses. **B.** LCMS total ion chromatogram (TIC) of crude peptide. Peak labeling indicates the predicted average and monoisotopic masses as well as the observed peptide mass. **C.** Mass spectrum at peak of **B** with *m/z* labeling for selected peaks and the associated charge in parentheses. **D.** HPLC trace of perfluoroaryl stapled peptide following single-pass purification. **E.** TIC of peptide following single-pass purification. **F.** Mass spectrum at peak of **E**.

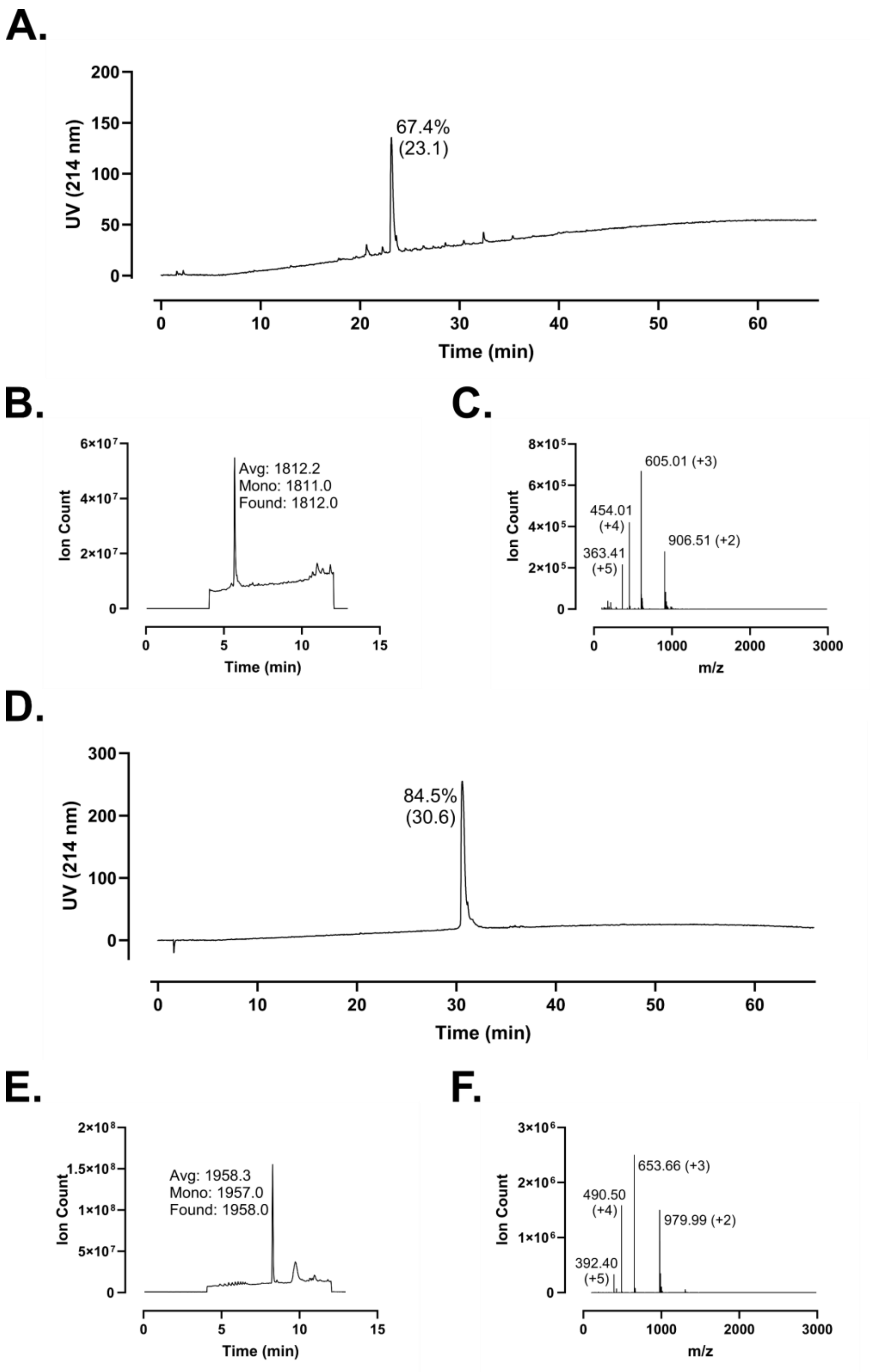
A.
67.4%
(23.1)
UV (214 nm)
Time (min)
B.
Avg: 1812.2
Mono: 1811.0
Found: 1812.0
Ion Count
Time (min)
C.
605.01 (+3)
454.01
(+4)
906.51 (+2)
363.41
(+5)
Ion Count
m/z
D.
84.5%
(30.6)
UV (214 nm)
Time (min)
E.
Avg: 1958.3
Mono: 1957.0
Found: 1958.0
Ion Count
Time (min)
F.
653.66 (+3)
490.50
(+4)
979.99 (+2)
392.40
(+5)
Ion Count
m/z

**Figure S186 – (#27) FF-14 F1+I5 Perfluoroaryl Staple Analytical Data. A.** HPLC trace of crude peptide (FF-14 F1C+I5C) at 214 nm. Peak labeling indicates percent purity by trace integration. Retention time is given in parentheses. **B.** LCMS total ion chromatogram (TIC) of crude peptide. Peak labeling indicates the predicted average and monoisotopic masses as well as the observed peptide mass. **C.** Mass spectrum at peak of **B** with *m/z* labeling for selected peaks and the associated charge in parentheses. **D.** HPLC trace of perfluoroaryl stapled peptide following single-pass purification. **E.** TIC of peptide following single-pass purification. **F.** Mass spectrum at peak of **E**.

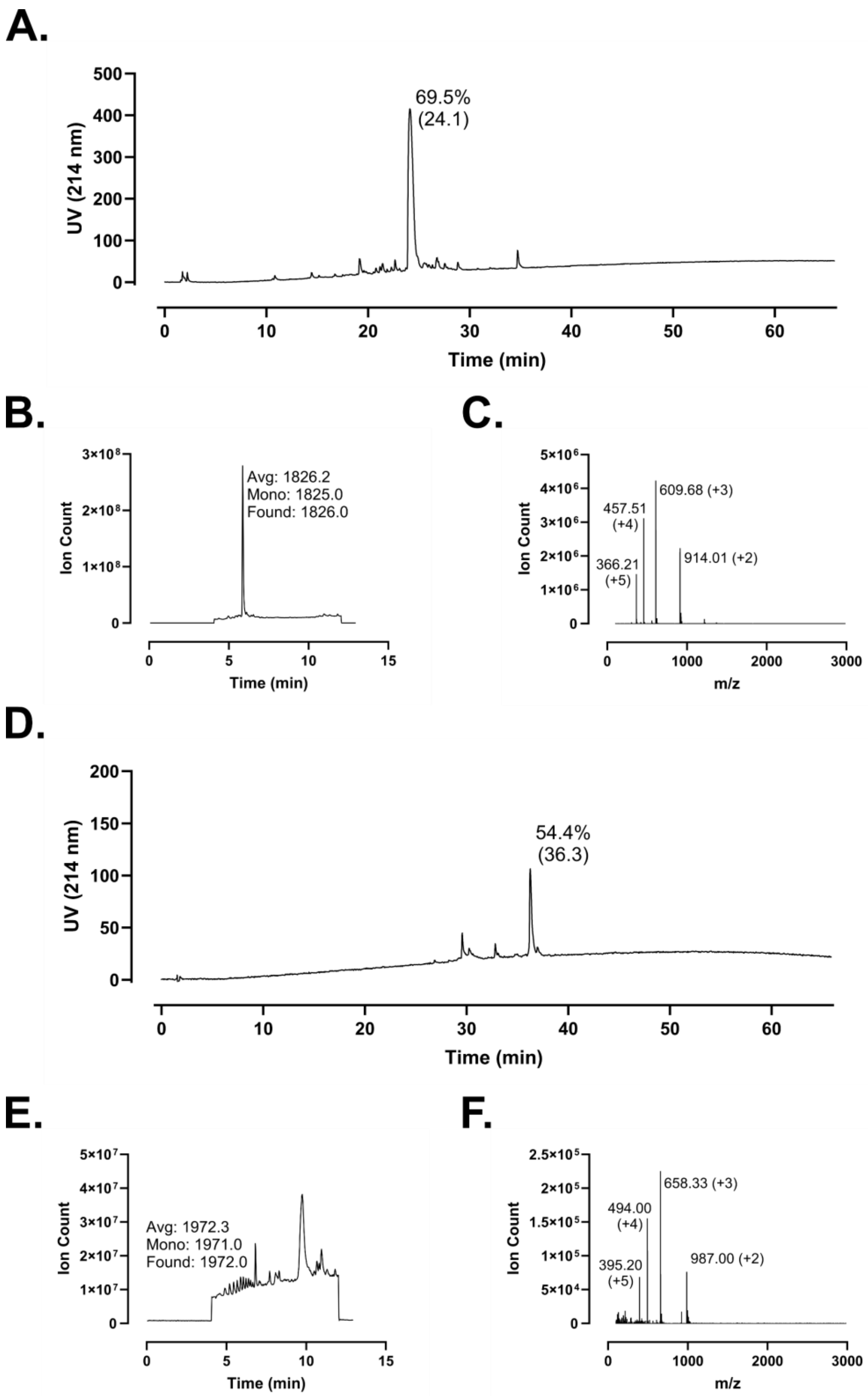
A.
69.5%
(24.1)
UV (214 nm)
Time (min)
B.
Avg: 1826.2
Mono: 1825.0
Found: 1826.0
Ion Count
Time (min)
C.
609.68 (+3)
457.51
(+4)
366.21
(+5)
914.01 (+2)
Ion Count
m/z
D.
54.4%
(36.3)
UV (214 nm)
Time (min)
E.
Avg: 1972.3
Mono: 1971.0
Found: 1972.0
Ion Count
Time (min)
F.
658.33 (+3)
494.00
(+4)
395.20
(+5)
987.00 (+2)
Ion Count
m/z

**Figure S187 – (#28) FF-14 F2+V6 Perfluoroaryl Staple Analytical Data. A.** HPLC trace of crude peptide (FF-14 F2C+V6C) at 214 nm. Peak labeling indicates percent purity by trace integration. Retention time is given in parentheses. **B.** LCMS total ion chromatogram (TIC) of crude peptide. Peak labeling indicates the predicted average and monoisotopic masses as well as the observed peptide mass. **C.** Mass spectrum at peak of **B** with *m/z* labeling for selected peaks and the associated charge in parentheses. **D.** HPLC trace of perfluoroaryl stapled peptide following single-pass purification; this is a minor product with a longer retention time but the same mass as the major product. Neither the minor nor the major product was characterized further due to lack of desirable biological properties. **E.** TIC of peptide following single-pass purification. **F.** Mass spectrum at peak of **E**.

A.

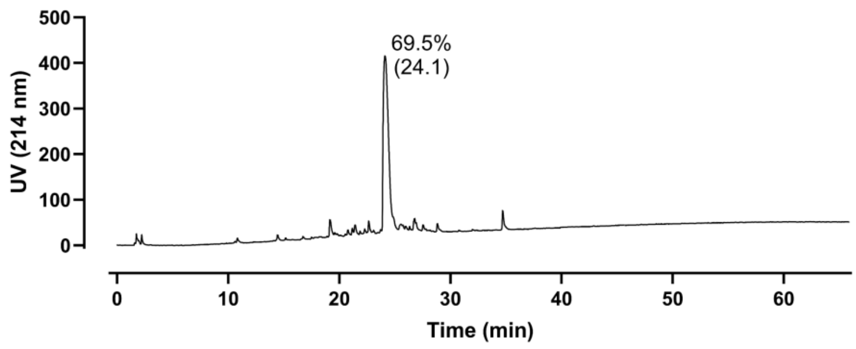

69.5%
(24.1)
UV (214 nm)
Time (min)


B.

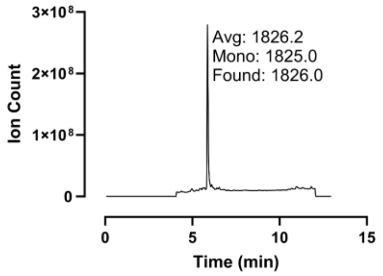

Avg: 1826.2
Mono: 1825.0
Found: 1826.0
Ion Count
Time (min)


C.

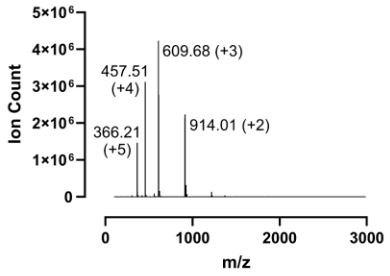

609.68 (+3)
457.51
(+4)
914.01 (+2)
366.21
(+5)
Ion Count
m/z


D.

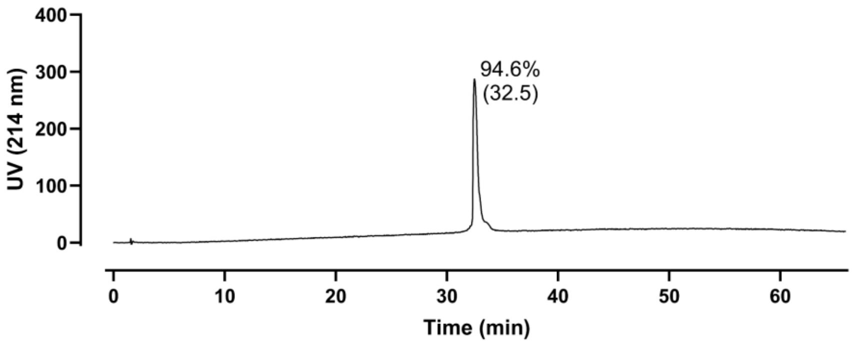

94.6%
(32.5)
UV (214 nm)
Time (min)


E.

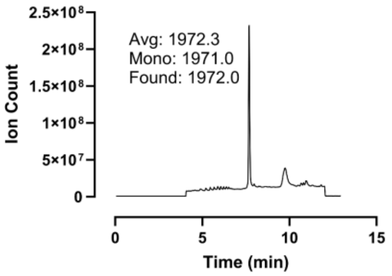

Avg: 1972.3
Mono: 1971.0
Found: 1972.0
Ion Count
Time (min)


F.

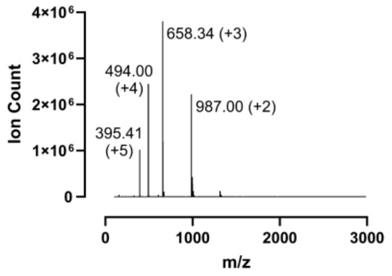

658.34 (+3)
494.00
(+4)
987.00 (+2)
395.41
(+5)
Ion Count
m/z

**Figure S188 – (#29) FF-14 F2+V6 Perfluoroaryl Staple Analytical Data. A.** HPLC trace of crude peptide (FF-14 F2C+V6C) at 214 nm. Peak labeling indicates percent purity by trace integration. Retention time is given in parentheses. These data are the same as those presented in **Figure S178 A-C** since the crude peptide used for stapling is the same in both instances. **B.** LCMS total ion chromatogram (TIC) of crude peptide. Peak labeling indicates the predicted average and monoisotopic masses as well as the observed peptide mass. **C.** Mass spectrum at peak of **B** with *m/z* labeling for selected peaks and the associated charge in parentheses. **D.** HPLC trace of perfluoroaryl stapled peptide following single-pass purification; this is the major product with a shorter retention time and the same mass as the minor product presented in **Figure S178**. Neither the minor nor the major product was characterized further due to lack of desirable biological properties. **E.** TIC of peptide following single-pass purification. **F.** Mass spectrum at peak of **E**.

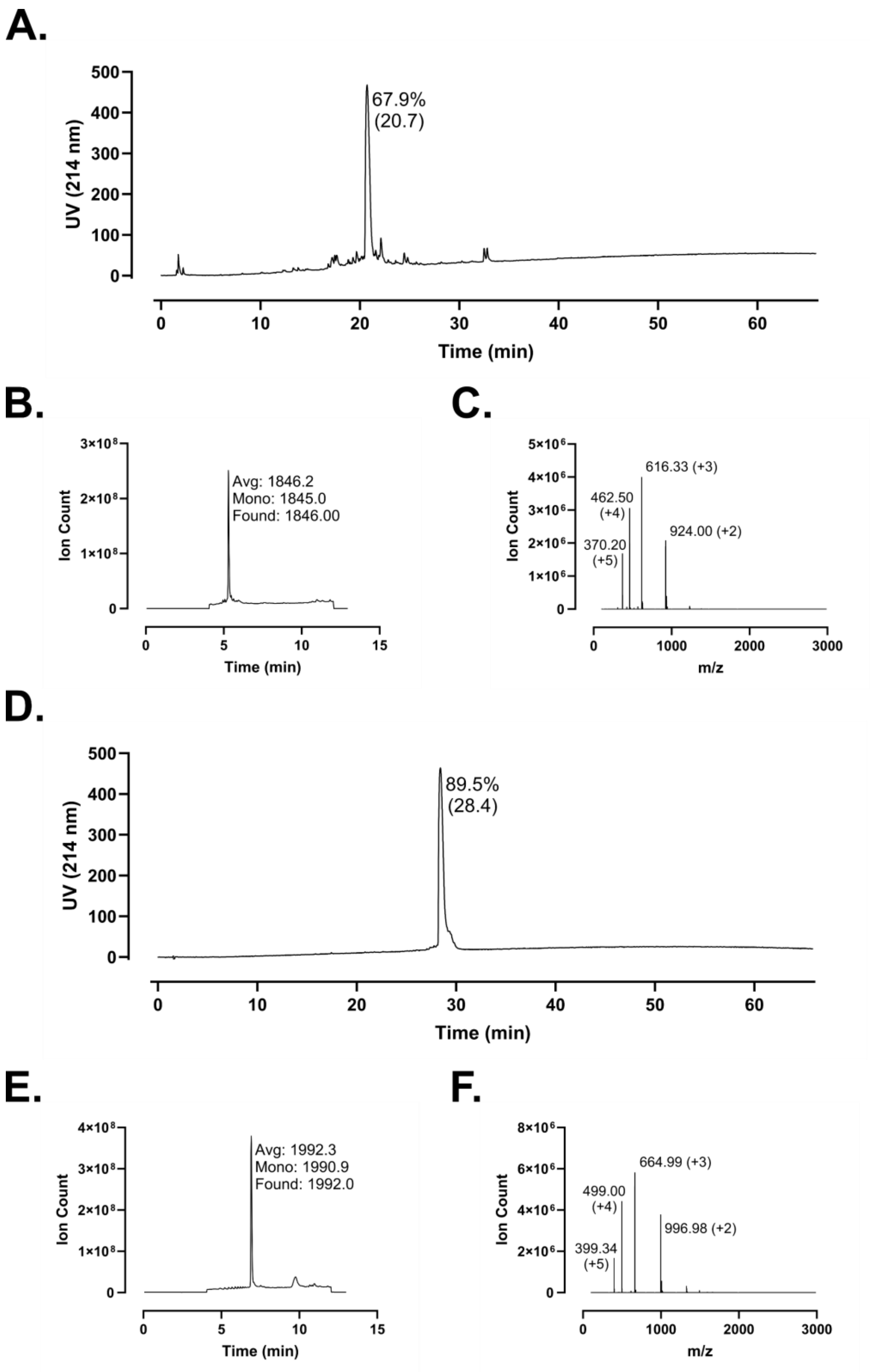
A.
500
400
300
200
100
0
UV (214 nm)
67.9%
(20.7)
0
10
20
30
40
50
60
Time (min)
B.
3×10⁸
2×10⁸
1×10⁸
0
Ion Count
Avg: 1846.2
Mono: 1845.0
Found: 1846.00
0
5
10
15
Time (min)
C.
5×10⁶
4×10⁶
3×10⁶
2×10⁶
1×10⁶
0
Ion Count
616.33 (+3)
462.50
(+4)
924.00 (+2)
370.20
(+5)
0
1000
2000
3000
m/z
D.
500
400
300
200
100
0
UV (214 nm)
89.5%
(28.4)
0
10
20
30
40
50
60
Time (min)
E.
4×10⁸
3×10⁸
2×10⁸
1×10⁸
0
Ion Count
Avg: 1992.3
Mono: 1990.9
Found: 1992.0
0
5
10
15
Time (min)
F.
8×10⁶
6×10⁶
4×10⁶
2×10⁶
0
Ion Count
664.99 (+3)
499.00
(+4)
996.98 (+2)
399.34
(+5)
0
1000
2000
3000
m/z

**Figure S189 – (#30) FF-14 I5+I9 Perfluoroaryl Staple Analytical Data. A.** HPLC trace of crude peptide (FF-14 I5C+I9C) at 214 nm. Peak labeling indicates percent purity by trace integration. Retention time is given in parentheses. **B.** LCMS total ion chromatogram (TIC) of crude peptide. Peak labeling indicates the predicted average and monoisotopic masses as well as the observed peptide mass. **C.** Mass spectrum at peak of **B** with *m/z* labeling for selected peaks and the associated charge in parentheses. **D.** HPLC trace of perfluoroaryl stapled peptide following single-pass purification. **E.** TIC of peptide following single-pass purification. **F.** Mass spectrum at peak of **E**.

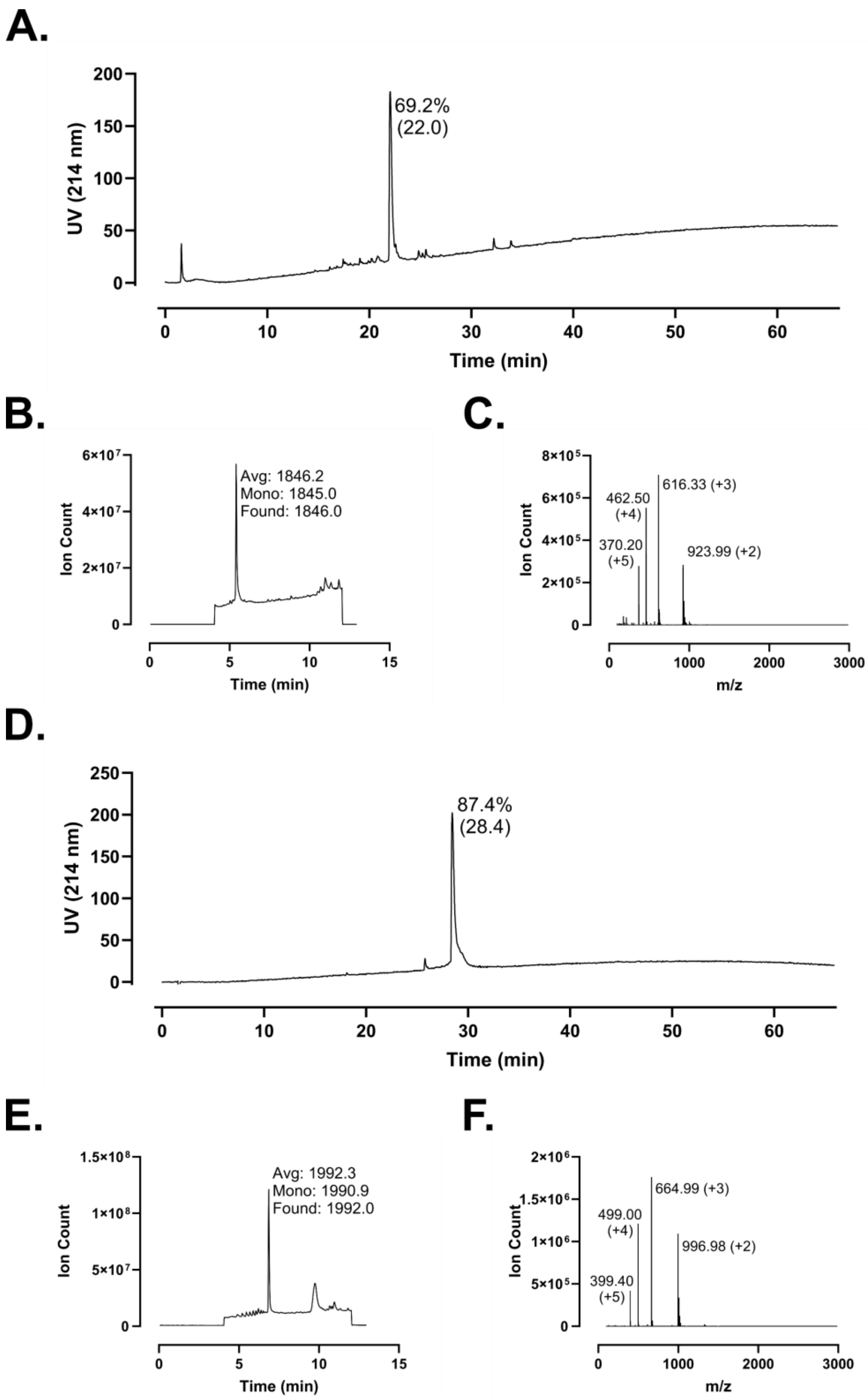
A.
UV (214 nm)
Time (min)
69.2%
(22.0)
B.
Ion Count
Time (min)
Avg: 1846.2
Mono: 1845.0
Found: 1846.0
C.
Ion Count
m/z
616.33 (+3)
462.50
(+4)
370.20
(+5)
923.99 (+2)
D.
UV (214 nm)
Time (min)
87.4%
(28.4)
E.
Ion Count
Time (min)
Avg: 1992.3
Mono: 1990.9
Found: 1992.0
F.
Ion Count
m/z
664.99 (+3)
499.00
(+4)
996.98 (+2)
399.40
(+5)

**Figure S190 – (#31) FF-14 I9+L13 Perfluoroaryl Staple Analytical Data. A.** HPLC trace of crude peptide (FF-14 I9C+L13C) at 214 nm. Peak labeling indicates percent purity by trace integration. Retention time is given in parentheses. **B.** LCMS total ion chromatogram (TIC) of crude peptide. Peak labeling indicates the predicted average and monoisotopic masses as well as the observed peptide mass. **C.** Mass spectrum at peak of **B** with *m/z* labeling for selected peaks and the associated charge in parentheses. **D.** HPLC trace of perfluoroaryl stapled peptide following single-pass purification. **E.** TIC of peptide following single-pass purification. **F.** Mass spectrum at peak of **E**.

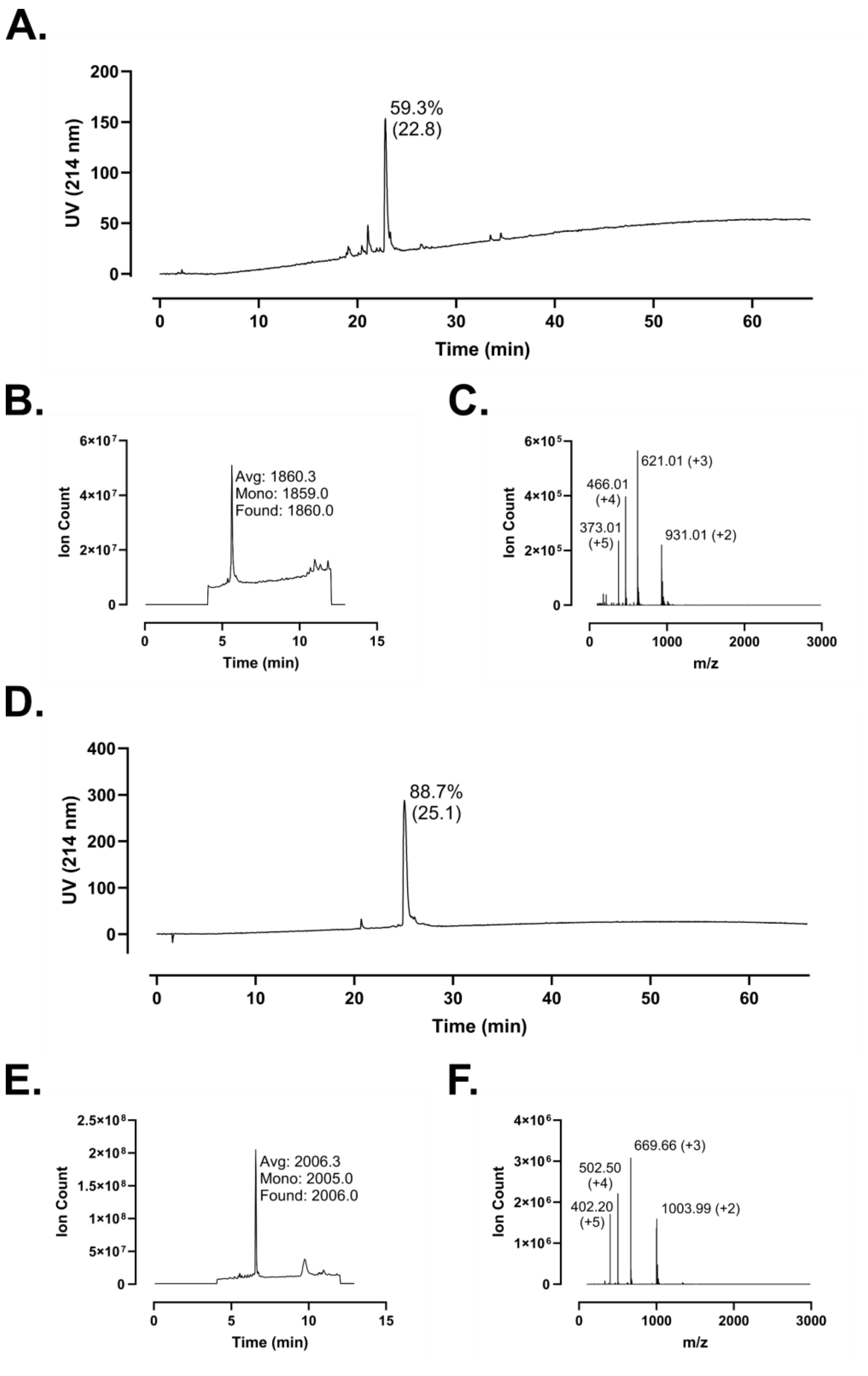
A.
59.3%
(22.8)
UV (214 nm)
Time (min)
B.
Avg: 1860.3
Mono: 1859.0
Found: 1860.0
Ion Count
Time (min)
C.
621.01 (+3)
466.01
(+4)
373.01
(+5)
931.01 (+2)
Ion Count
m/z
D.
88.7%
(25.1)
UV (214 nm)
Time (min)
E.
Avg: 2006.3
Mono: 2005.0
Found: 2006.0
Ion Count
Time (min)
F.
669.66 (+3)
502.50
(+4)
402.20
(+5)
1003.99 (+2)
Ion Count
m/z

**Figure S191 – (#32) FF-14 V6+I9 Perfluoroaryl Staple Analytical Data. A.** HPLC trace of crude peptide (FF-14 V6C+I9C) at 214 nm. Peak labeling indicates percent purity by trace integration. Retention time is given in parentheses. **B.** LCMS total ion chromatogram (TIC) of crude peptide. Peak labeling indicates the predicted average and monoisotopic masses as well as the observed peptide mass. **C.** Mass spectrum at peak of **B** with *m/z* labeling for selected peaks and the associated charge in parentheses. **D.** HPLC trace of perfluoroaryl stapled peptide following single-pass purification. **E.** TIC of peptide following single-pass purification. **F.** Mass spectrum at peak of **E**.

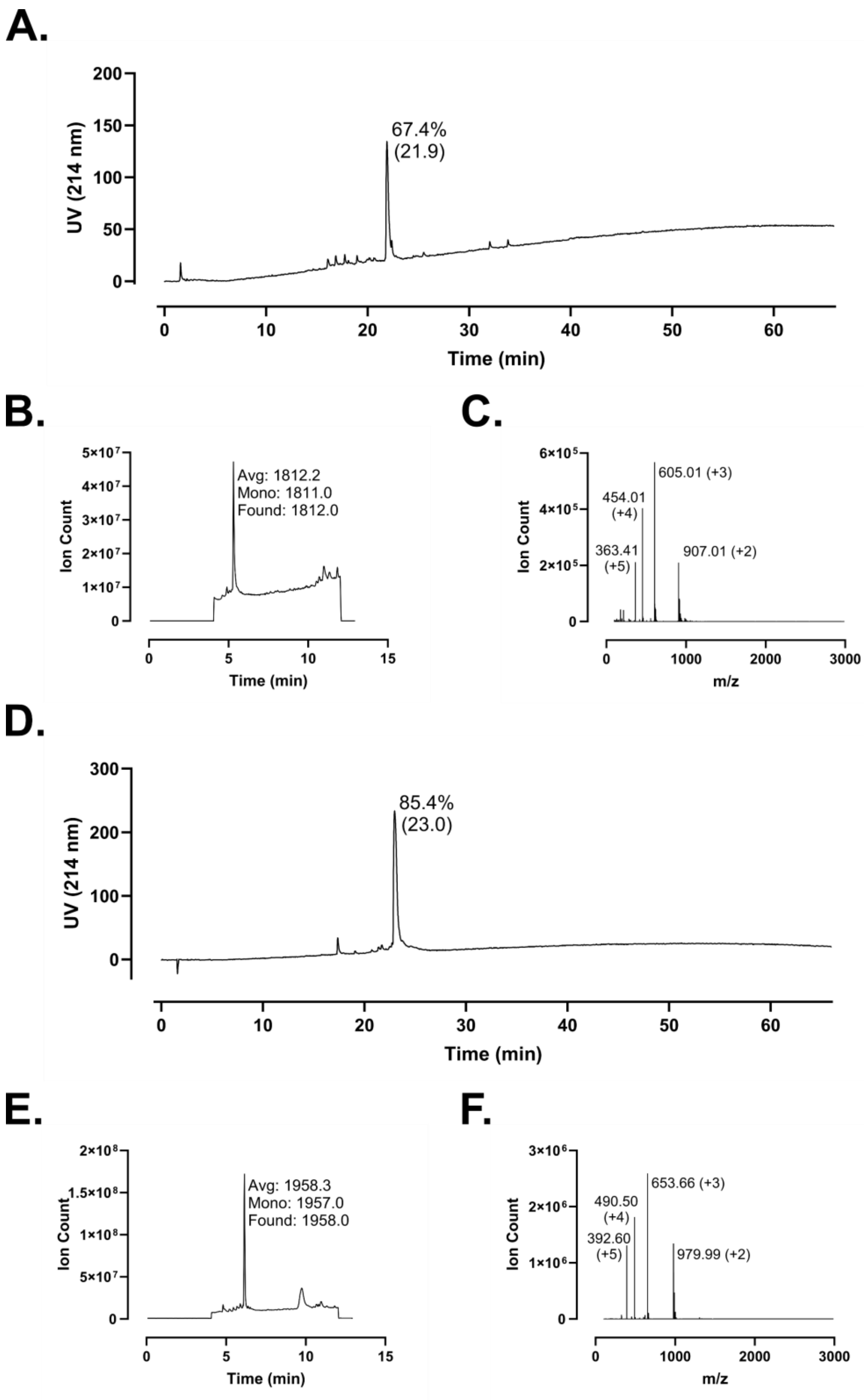
A.
UV (214 nm)
200
150
100
50
0
67.4%
(21.9)
0
10
20
30
40
50
60
Time (min)
B.
Ion Count
5×10⁷
4×10⁷
3×10⁷
2×10⁷
1×10⁷
0
Avg: 1812.2
Mono: 1811.0
Found: 1812.0
0
5
10
15
Time (min)
C.
Ion Count
6×10⁵
4×10⁵
2×10⁵
0
605.01 (+3)
454.01
(+4)
363.41
(+5)
907.01 (+2)
0
1000
2000
3000
m/z
D.
UV (214 nm)
300
200
100
0
85.4%
(23.0)
0
10
20
30
40
50
60
Time (min)
E.
Ion Count
2×10⁸
1.5×10⁸
1×10⁸
5×10⁷
0
Avg: 1958.3
Mono: 1957.0
Found: 1958.0
0
5
10
15
Time (min)
F.
Ion Count
3×10⁶
2×10⁶
1×10⁶
0
653.66 (+3)
490.50
(+4)
392.60
(+5)
979.99 (+2)
0
1000
2000
3000
m/z

**Figure S192 – (#33) FF-14 I9+F12 Perfluoroaryl Staple Analytical Data. A.** HPLC trace of crude peptide (FF-14 I9C+F12C) at 214 nm. Peak labeling indicates percent purity by trace integration. Retention time is given in parentheses. **B.** LCMS total ion chromatogram (TIC) of crude peptide. Peak labeling indicates the predicted average and monoisotopic masses as well as the observed peptide mass. **C.** Mass spectrum at peak of **B** with *m/z* labeling for selected peaks and the associated charge in parentheses. **D.** HPLC trace of perfluoroaryl stapled peptide following single-pass purification. **E.** TIC of peptide following single-pass purification. **F.** Mass spectrum at peak of **E**.

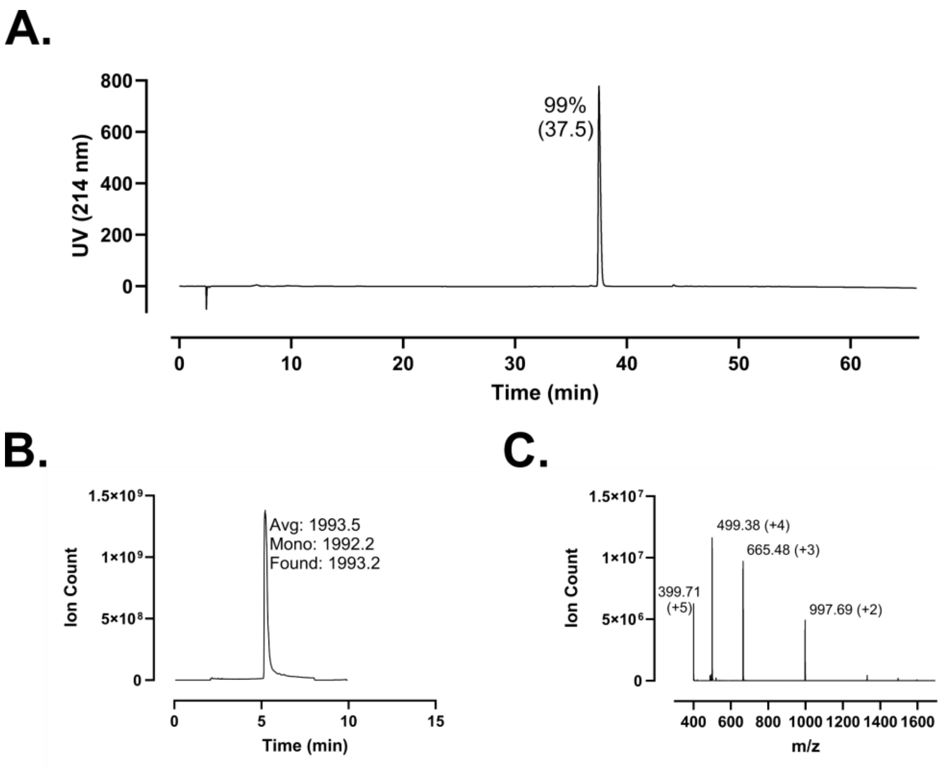
A.
99%
(37.5)
UV (214 nm)
800
600
400
200
0
0
10
20
30
40
50
60
Time (min)
B.
1.5×10⁹
1×10⁹
5×10⁸
0
Ion Count
Avg: 1993.5
Mono: 1992.2
Found: 1993.2
0
5
10
15
Time (min)
C.
1.5×10⁷
1×10⁷
5×10⁶
0
Ion Count
499.38 (+4)
665.48 (+3)
399.71
(+5)
997.69 (+2)
400
600
800
1000
1200
1400
1600
m/z

**Figure S193 – (#34) 5-DMA-Penta-FF-14 Analytical Data. A.** HPLC trace at 214 nm of N-lipidated peptide following single-pass purification. Peak labeling indicates percent purity by trace integration. Retention time is given in parentheses. **B.** LCMS total ion chromatogram (TIC) of peptide after single-pass purification. Peak labeling indicates the predicted average and monoisotopic masses as well as the observed peptide mass. **C.** Mass spectrum at peak of **B** with *m/z* labeling for selected peaks and the associated charge in parentheses.

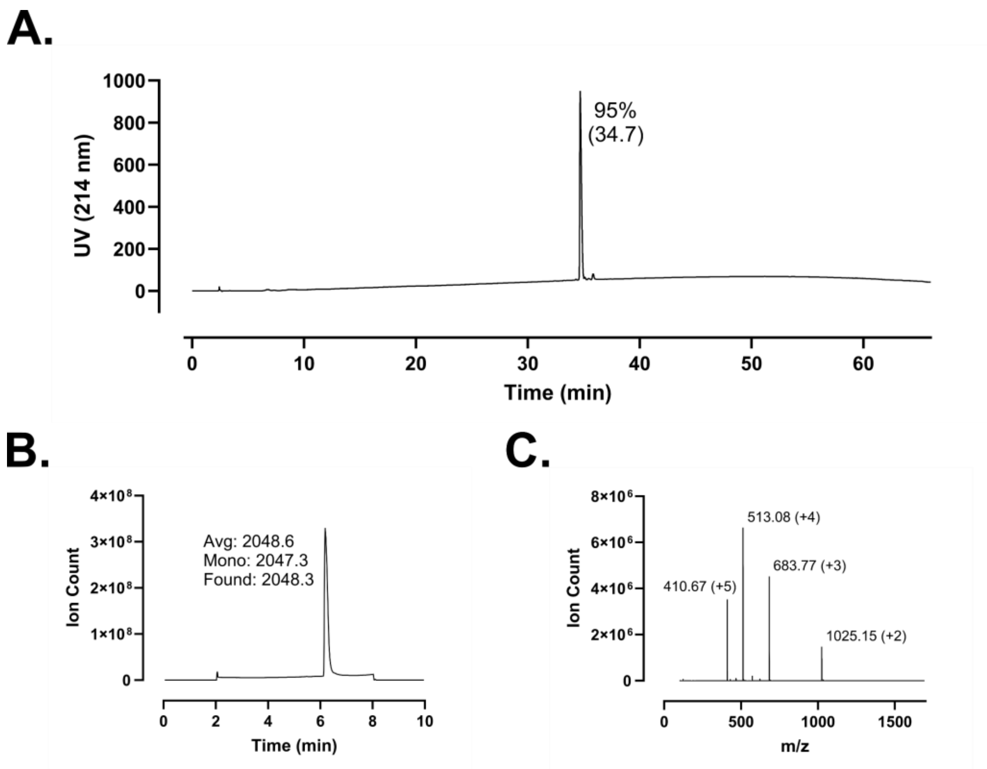
A.
UV (214 nm)
1000
800
600
400
200
0
95%
(34.7)
0
10
20
30
40
50
60
Time (min)
B.
Ion Count
4×10⁸
3×10⁸
2×10⁸
1×10⁸
0
Avg: 2048.6
Mono: 2047.3
Found: 2048.3
0
2
4
6
8
10
Time (min)
C.
Ion Count
8×10⁶
6×10⁶
4×10⁶
2×10⁶
0
513.08 (+4)
683.77 (+3)
410.67 (+5)
1025.15 (+2)
0
500
1000
1500
m/z

**Figure S194 – (#35) 10-Me-Undeca-FF-14 Analytical Data. A.** HPLC trace at 214 nm of N-lipidated peptide following single-pass purification. Peak labeling indicates percent purity by trace integration. Retention time is given in parentheses. **B.** LCMS total ion chromatogram (TIC) of peptide after single-pass purification. Peak labeling indicates the predicted average and monoisotopic masses as well as the observed peptide mass. **C.** Mass spectrum at peak of **B** with *m/z* labeling for selected peaks and the associated charge in parentheses.

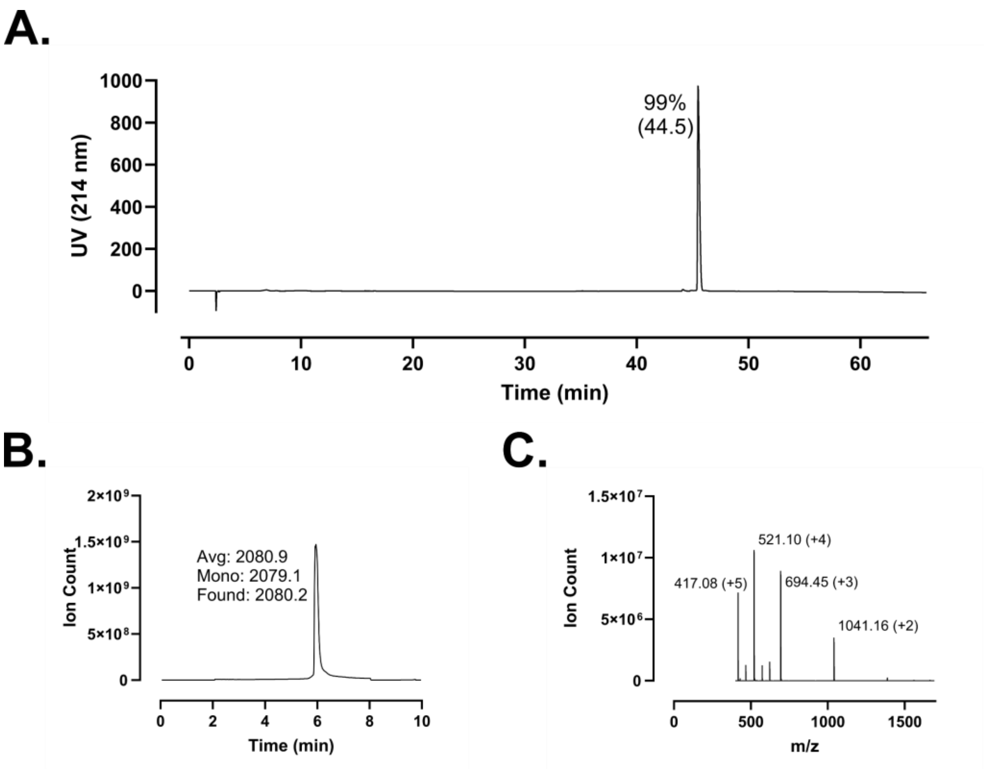
A.
99%
(44.5)
UV (214 nm)
Time (min)
B.
Avg: 2080.9
Mono: 2079.1
Found: 2080.2
Ion Count
Time (min)
C.
521.10 (+4)
417.08 (+5)
694.45 (+3)
1041.16 (+2)
Ion Count
m/z

**Figure S195 – (#36) 4-Cl-Biphenyl-FF-14 Analytical Data. A.** HPLC trace at 214 nm of N-lipidated peptide following single-pass purification. Peak labeling indicates percent purity by trace integration. Retention time is given in parentheses. **B.** LCMS total ion chromatogram (TIC) of peptide after single-pass purification. Peak labeling indicates the predicted average and monoisotopic masses as well as the observed peptide mass. **C.** Mass spectrum at peak of **B** with *m/z* labeling for selected peaks and the associated charge in parentheses.

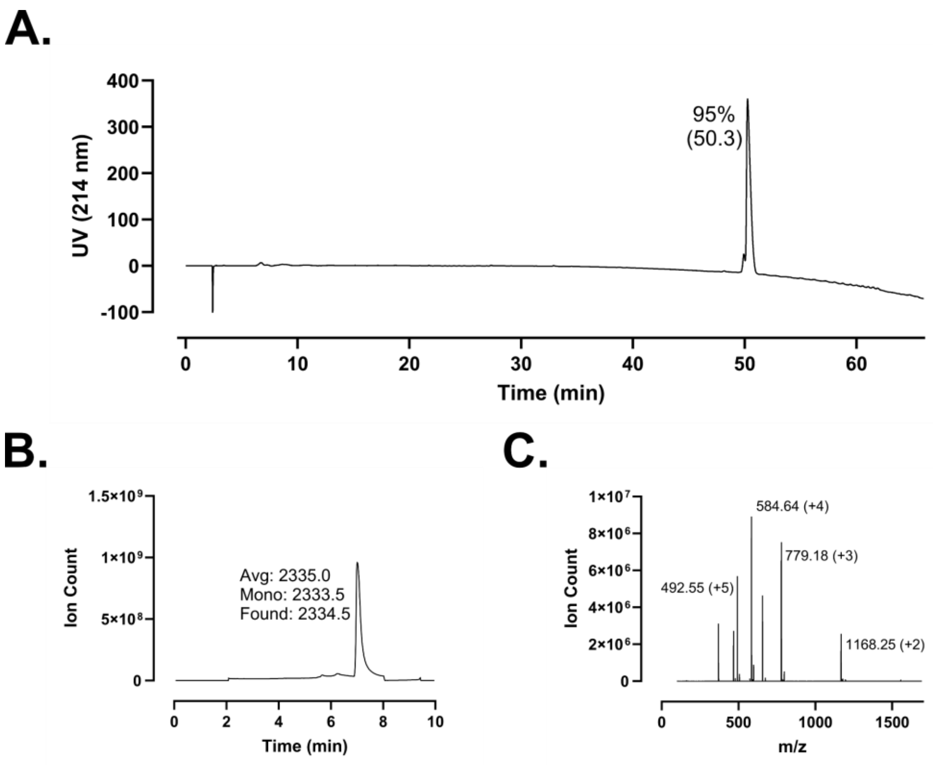

A.
400
300
200
100
0
-100
UV (214 nm)
95%
(50.3)
0
10
20
30
40
50
60
Time (min)
B.
1.5×10⁹
1×10⁹
5×10⁸
0
Ion Count
Avg: 2335.0
Mono: 2333.5
Found: 2334.5
0
2
4
6
8
10
Time (min)
C.
1×10⁷
8×10⁶
6×10⁶
4×10⁶
2×10⁶
0
Ion Count
584.64 (+4)
779.18 (+3)
492.55 (+5)
1168.25 (+2)
0
500
1000
1500
m/z

**Figure S196 – (#37) Cholesteryl-FF-14 Analytical Data. A.** HPLC trace at 214 nm of N-lipidated peptide following single-pass purification. Peak labeling indicates percent purity by trace integration. Retention time is given in parentheses. **B.** LCMS total ion chromatogram (TIC) of peptide after single-pass purification. Peak labeling indicates the predicted average and monoisotopic masses as well as the observed peptide mass. **C.** Mass spectrum at peak of **B** with *m/z* labeling for selected peaks and the associated charge in parentheses.

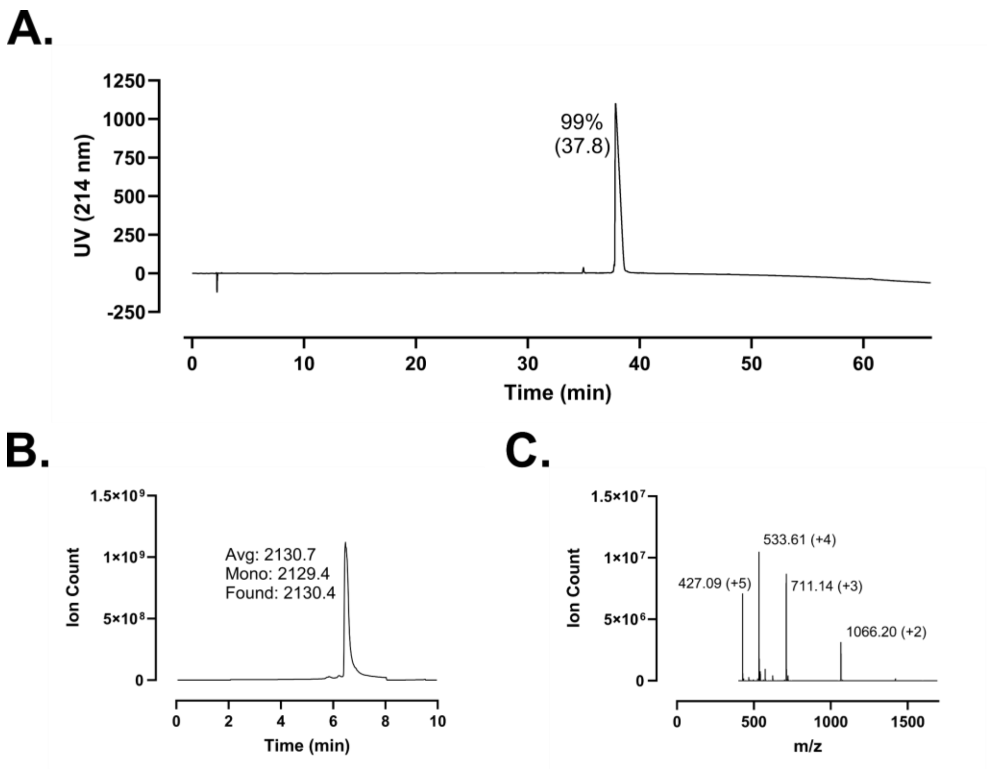
A.
UV (214 nm)
1250
1000
750
500
250
0
-250
99%
(37.8)
0
10
20
30
40
50
60
Time (min)
B.
Ion Count
1.5×10⁹
1×10⁹
5×10⁸
0
Avg: 2130.7
Mono: 2129.4
Found: 2130.4
0
2
4
6
8
10
Time (min)
C.
Ion Count
1.5×10⁷
1×10⁷
5×10⁶
0
533.61 (+4)
427.09 (+5)
711.14 (+3)
1066.20 (+2)
0
500
1000
1500
m/z

**Figure S197 – (#38) Elaidic-FF-14 Analytical Data. A.** HPLC trace at 214 nm of N-lipidated peptide following single-pass purification. Peak labeling indicates percent purity by trace integration. Retention time is given in parentheses. **B.** LCMS total ion chromatogram (TIC) of peptide after single-pass purification. Peak labeling indicates the predicted average and monoisotopic masses as well as the observed peptide mass. **C.** Mass spectrum at peak of **B** with *m/z* labeling for selected peaks and the associated charge in parentheses.

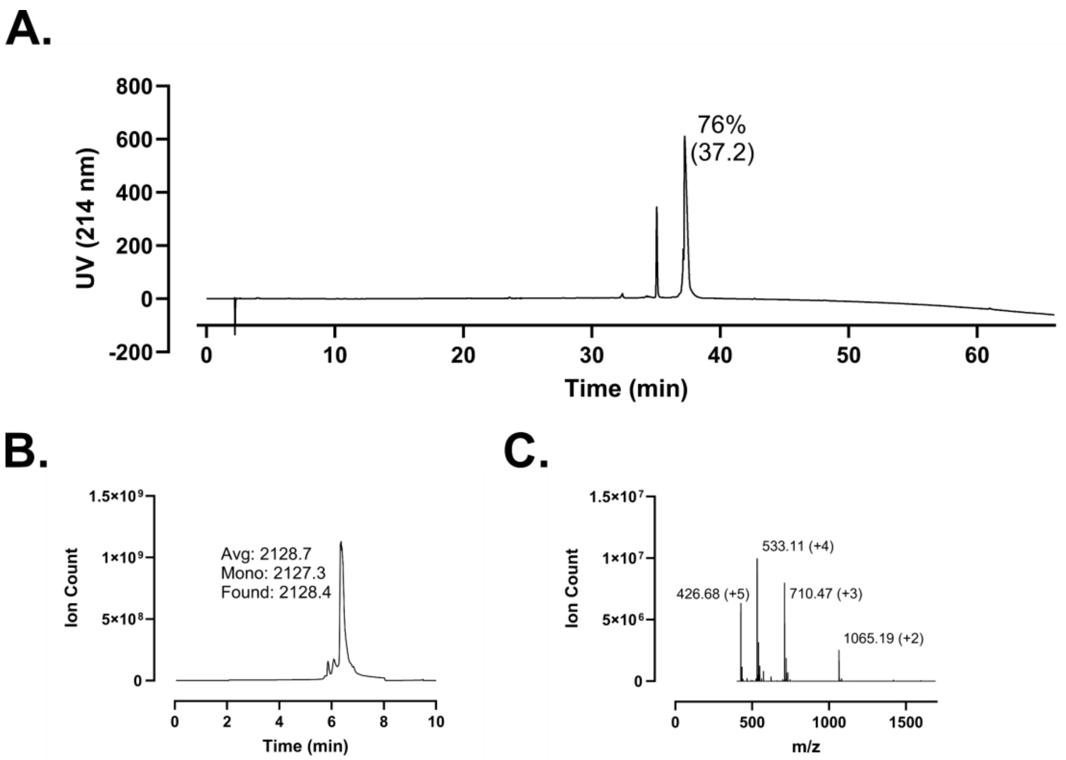
A.
800
600
400
200
0
-200
UV (214 nm)
76%
(37.2)
0
10
20
30
40
50
60
Time (min)
B.
1.5×10⁹
1×10⁹
5×10⁸
0
Ion Count
Avg: 2128.7
Mono: 2127.3
Found: 2128.4
0
2
4
6
8
10
Time (min)
C.
1.5×10⁷
1×10⁷
5×10⁶
0
Ion Count
533.11 (+4)
426.68 (+5)
710.47 (+3)
1065.19 (+2)
0
500
1000
1500
m/z

**Figure S198 – (#39) Linoleic-FF-14 Analytical Data. A.** HPLC trace at 214 nm of N-lipidated peptide following single-pass purification. Peak labeling indicates percent purity by trace integration. Retention time is given in parentheses. **B.** LCMS total ion chromatogram (TIC) of peptide after single-pass purification. Peak labeling indicates the predicted average and monoisotopic masses as well as the observed peptide mass. **C.** Mass spectrum at peak of **B** with *m/z* labeling for selected peaks and the associated charge in parentheses.

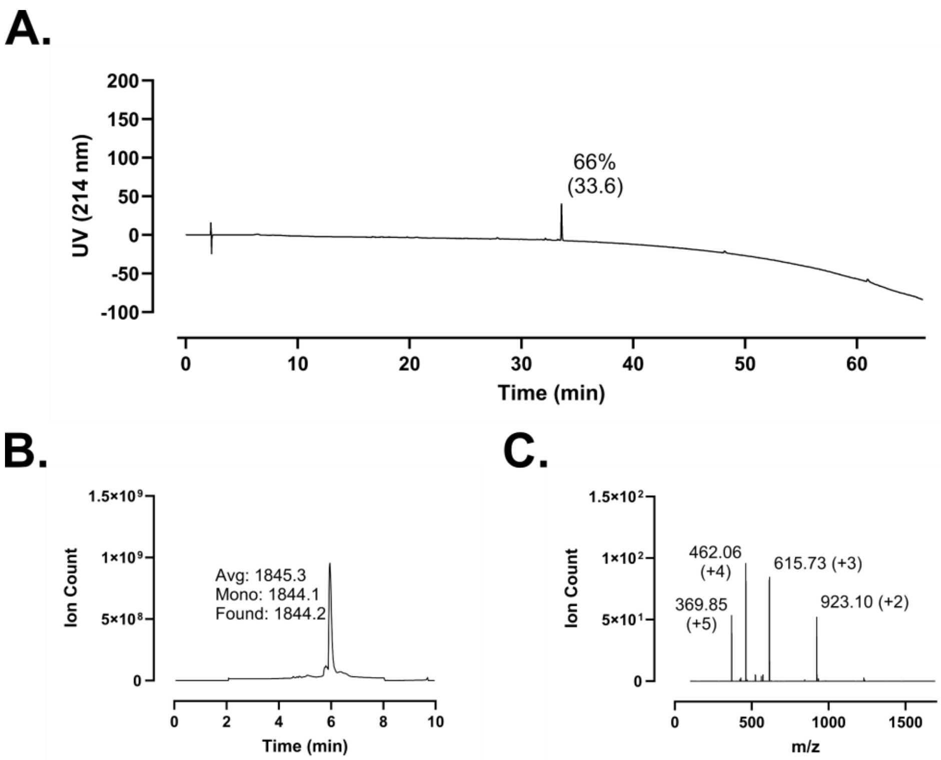
A.
200
150
100
50
0
-50
-100
UV (214 nm)
66%
(33.6)
0
10
20
30
40
50
60
Time (min)
B.
1.5×10⁹
1×10⁹
5×10⁸
0
Ion Count
Avg: 1845.3
Mono: 1844.1
Found: 1844.2
0
2
4
6
8
10
Time (min)
C.
1.5×10²
1×10²
5×10¹
0
Ion Count
462.06
(+4)
615.73 (+3)
369.85
(+5)
923.10 (+2)
0
500
1000
1500
m/z

**Figure S199 – (#40) 6-Me-Hepta-FK-13 Analytical Data. A.** HPLC trace at 214 nm of N-lipidated peptide following single-pass purification. Peak labeling indicates percent purity by trace integration. Retention time is given in parentheses. **B.** LCMS total ion chromatogram (TIC) of peptide after single-pass purification. Peak labeling indicates the predicted average and monoisotopic masses as well as the observed peptide mass. **C.** Mass spectrum at peak of **B** with *m/z* labeling for selected peaks and the associated charge in parentheses.

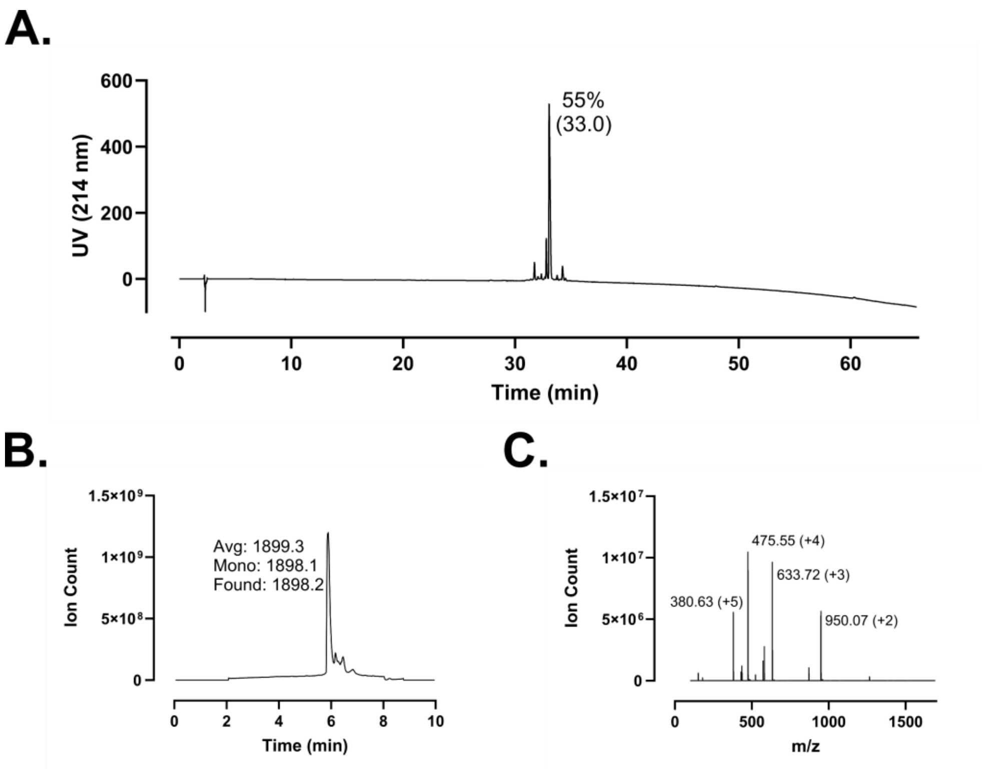
A.
55%
(33.0)
UV (214 nm)
Time (min)
B.
Avg: 1899.3
Mono: 1898.1
Found: 1898.2
Ion Count
Time (min)
C.
475.55 (+4)
633.72 (+3)
380.63 (+5)
950.07 (+2)
Ion Count
m/z

**Figure S200 – (#41) Biphenyl-FK-13 Analytical Data. A.** HPLC trace at 214 nm of N-lipidated peptide following single-pass purification. Peak labeling indicates percent purity by trace integration. Retention time is given in parentheses. **B.** LCMS total ion chromatogram (TIC) of peptide after single-pass purification. Peak labeling indicates the predicted average and monoisotopic masses as well as the observed peptide mass. **C.** Mass spectrum at peak of **B** with *m/z* labeling for selected peaks and the associated charge in parentheses.

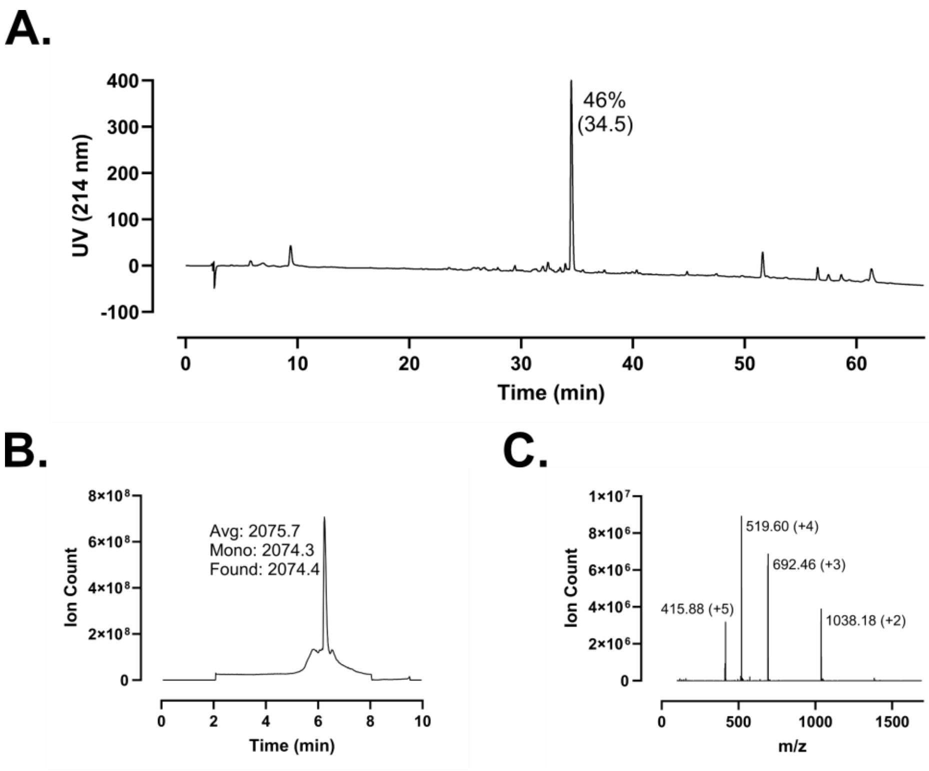
A.
46%
(34.5)
UV (214 nm)
Time (min)
B.
Avg: 2075.7
Mono: 2074.3
Found: 2074.4
Ion Count
Time (min)
C.
519.60 (+4)
692.46 (+3)
415.88 (+5)
1038.18 (+2)
Ion Count
m/z

**Figure S201 – (#42) Cholenic-FK-13 Analytical Data. A.** HPLC trace at 214 nm of N-lipidated peptide following single-pass purification. Peak labeling indicates percent purity by trace integration. Retention time is given in parentheses. **B.** LCMS total ion chromatogram (TIC) of peptide after single-pass purification. Peak labeling indicates the predicted average and monoisotopic masses as well as the observed peptide mass. **C.** Mass spectrum at peak of **B** with *m/z* labeling for selected peaks and the associated charge in parentheses.

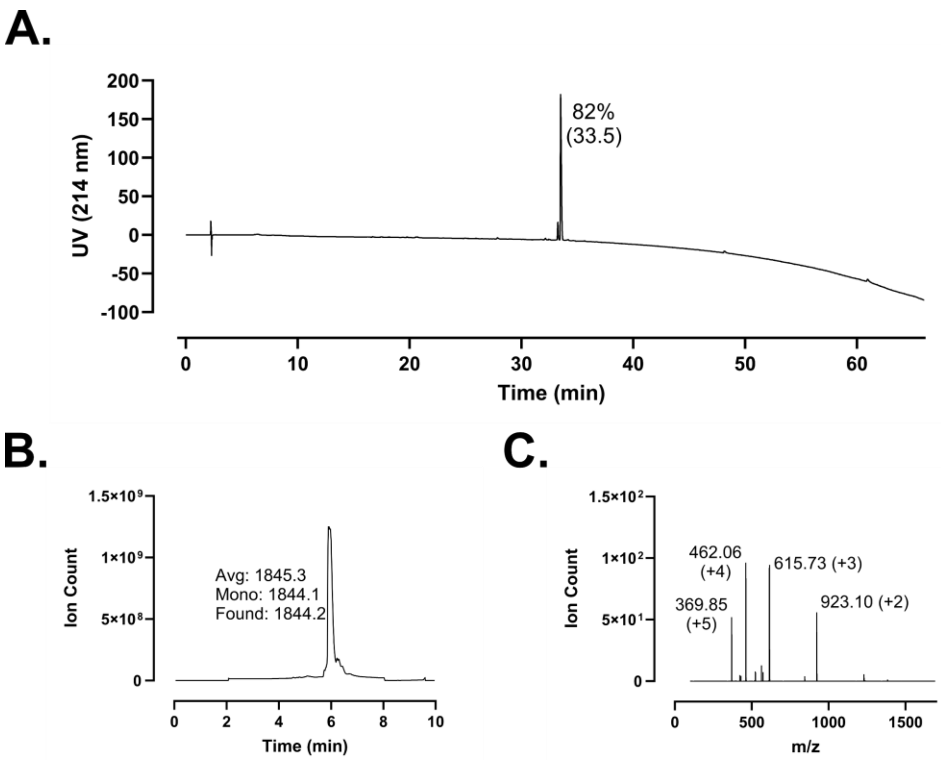
A.
UV (214 nm)
82%
(33.5)
Time (min)
B.
Ion Count
Avg: 1845.3
Mono: 1844.1
Found: 1844.2
Time (min)
C.
Ion Count
462.06
(+4)
615.73 (+3)
369.85
(+5)
923.10 (+2)
m/z

**Figure S202 – (#43) Octanoic-FK-13 Analytical Data. A.** HPLC trace at 214 nm of N-lipidated peptide following single-pass purification. Peak labeling indicates percent purity by trace integration. Retention time is given in parentheses. **B.** LCMS total ion chromatogram (TIC) of peptide after single-pass purification. Peak labeling indicates the predicted average and monoisotopic masses as well as the observed peptide mass. **C.** Mass spectrum at peak of **B** with *m/z* labeling for selected peaks and the associated charge in parentheses.

A.

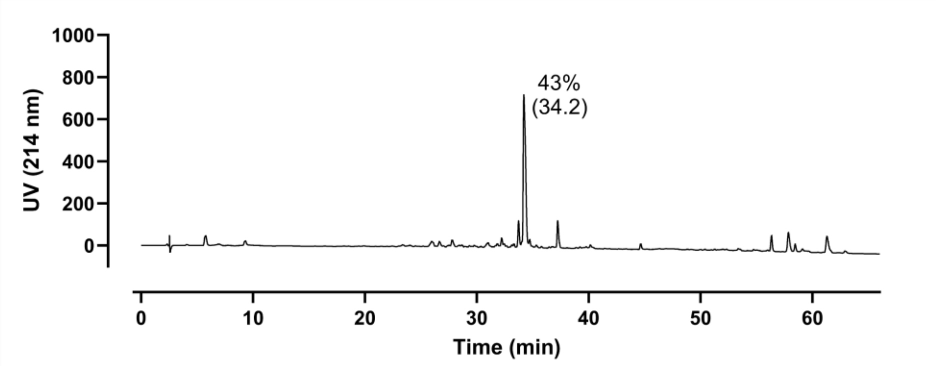

UV (214 nm)
1000
800
600
400
200
0
43%
(34.2)
0
10
20
30
40
50
60
Time (min)


B.

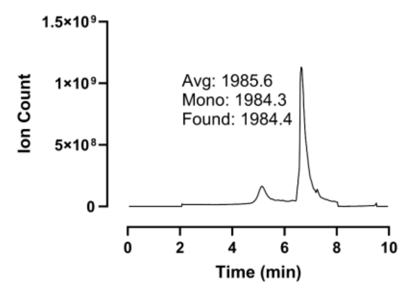

1.5×10⁹
1×10⁹
5×10⁸
0
Ion Count
Avg: 1985.6
Mono: 1984.3
Found: 1984.4
0
2
4
6
8
10
Time (min)


C.

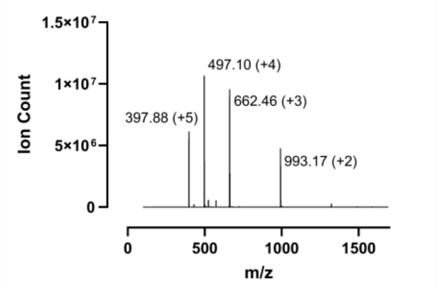

1.5×10⁷
1×10⁷
5×10⁶
0
Ion Count
497.10 (+4)
397.88 (+5)
662.46 (+3)
993.17 (+2)
0
500
1000
1500
m/z

**Figure S203 – (#44) Stearic-FK-13 Analytical Data. A.** HPLC trace at 214 nm of N-lipidated peptide following single-pass purification. Peak labeling indicates percent purity by trace integration. Retention time is given in parentheses. **B.** LCMS total ion chromatogram (TIC) of peptide after single-pass purification. Peak labeling indicates the predicted average and monoisotopic masses as well as the observed peptide mass. **C.** Mass spectrum at peak of **B** with *m/z* labeling for selected peaks and the associated charge in parentheses.

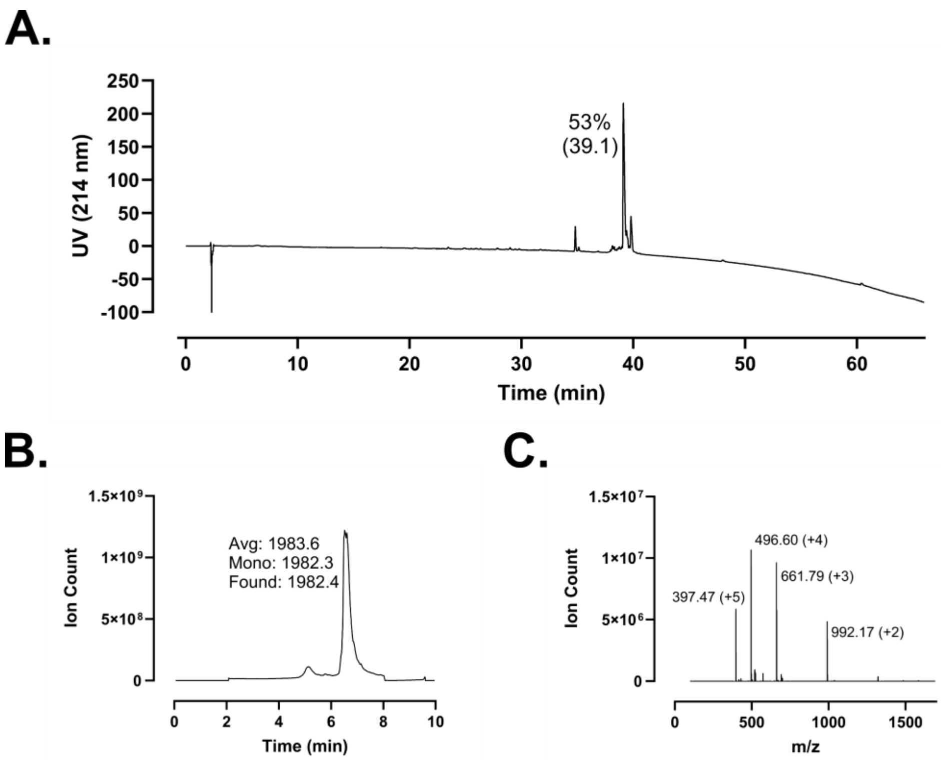
A.
53%
(39.1)
UV (214 nm)
Time (min)
B.
Avg: 1983.6
Mono: 1982.3
Found: 1982.4
Ion Count
Time (min)
C.
496.60 (+4)
661.79 (+3)
397.47 (+5)
992.17 (+2)
Ion Count
m/z

**Figure S204 – (#45) Oleic-FK-13 Analytical Data. A.** HPLC trace at 214 nm of N-lipidated peptide following single-pass purification. Peak labeling indicates percent purity by trace integration. Retention time is given in parentheses. **B.** LCMS total ion chromatogram (TIC) of peptide after single-pass purification. Peak labeling indicates the predicted average and monoisotopic masses as well as the observed peptide mass. **C.** Mass spectrum at peak of **B** with *m/z* labeling for selected peaks and the associated charge in parentheses.

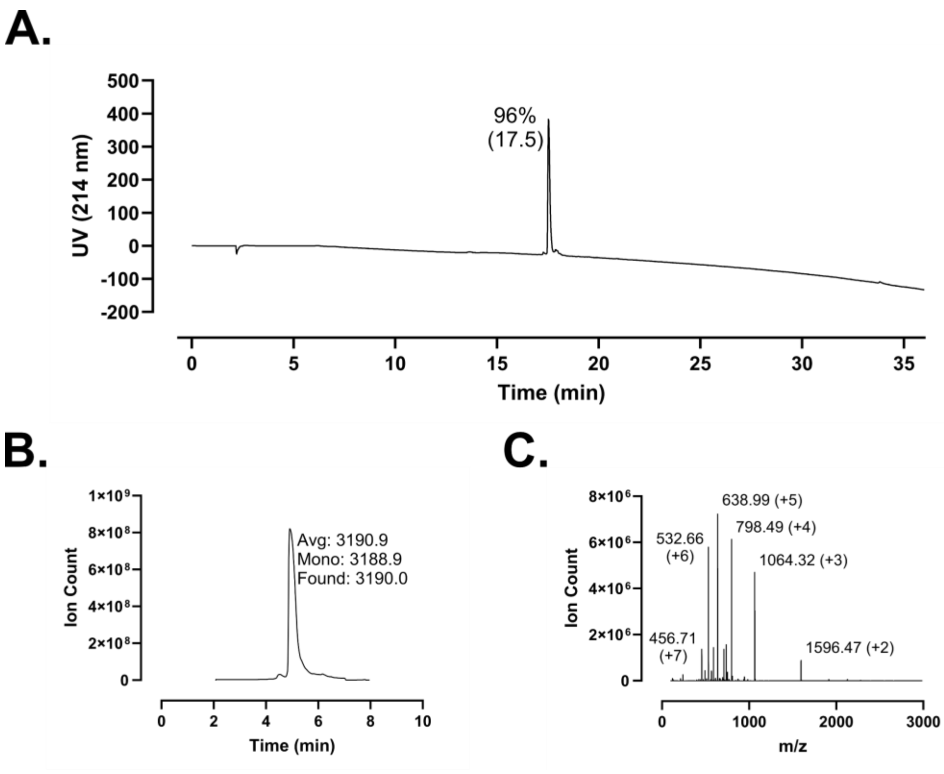
A.
96%
(17.5)
UV (214 nm)
500
400
300
200
100
0
-100
-200
0
5
10
15
20
25
30
35
Time (min)
B.
Ion Count
Avg: 3190.9
Mono: 3188.9
Found: 3190.0
Time (min)
C.
Ion Count
638.99 (+5)
798.49 (+4)
532.66
(+6)
1064.32 (+3)
456.71
(+7)
1596.47 (+2)
m/z

**Figure S205 – (#46) Oct-LL-25 Analytical Data. A.** HPLC trace at 214 nm of N-lipidated peptide following single-pass purification. Peak labeling indicates percent purity by trace integration. Retention time is given in parentheses. **B.** LCMS total ion chromatogram (TIC) of peptide after single-pass purification. Peak labeling indicates the predicted average and monoisotopic masses as well as the observed peptide mass. **C.** Mass spectrum at peak of **B** with *m/z* labeling for selected peaks and the associated charge in parentheses.

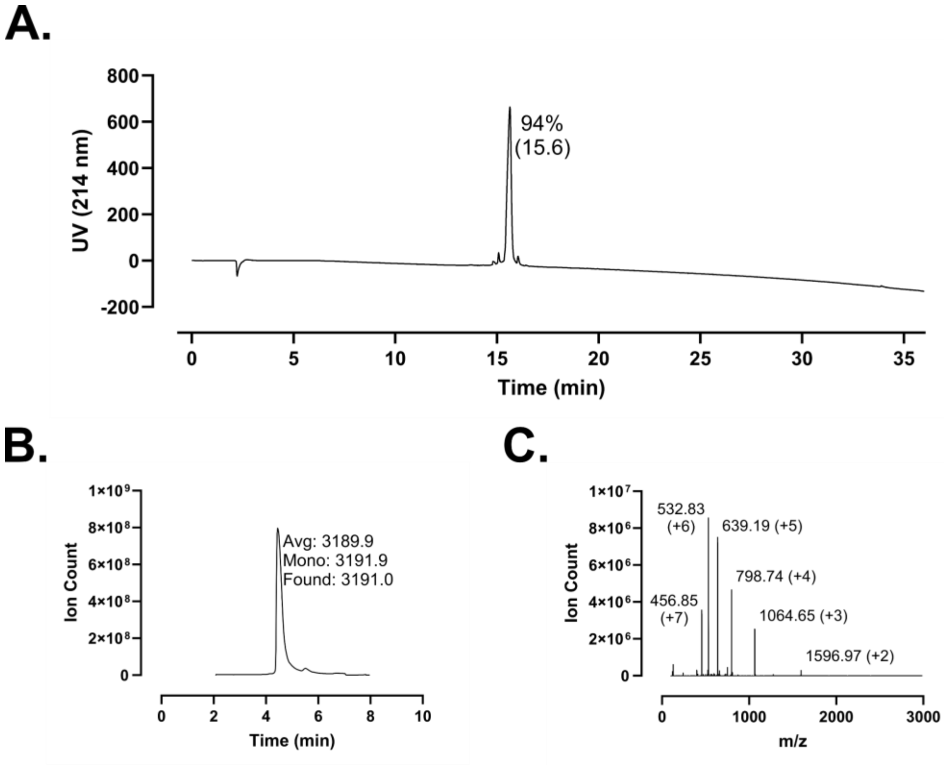
A.
800
600
400
200
0
-200
UV (214 nm)
94%
(15.6)
0
5
10
15
20
25
30
35
Time (min)
B.
1×10⁹
8×10⁸
6×10⁸
4×10⁸
2×10⁸
0
Ion Count
Avg: 3189.9
Mono: 3191.9
Found: 3191.0
0
2
4
6
8
10
Time (min)
C.
1×10⁷
8×10⁶
6×10⁶
4×10⁶
2×10⁶
0
Ion Count
532.83
(+6)
639.19 (+5)
798.74 (+4)
456.85
(+7)
1064.65 (+3)
1596.97 (+2)
0
1000
2000
3000
m/z

**Figure S206 – (#47) DMA-LL-25 Analytical Data. A.** HPLC trace at 214 nm of N-lipidated peptide following single-pass purification. Peak labeling indicates percent purity by trace integration. Retention time is given in parentheses. **B.** LCMS total ion chromatogram (TIC) of peptide after single-pass purification. Peak labeling indicates the predicted average and monoisotopic masses as well as the observed peptide mass. **C.** Mass spectrum at peak of **B** with *m/z* labeling for selected peaks and the associated charge in parentheses.

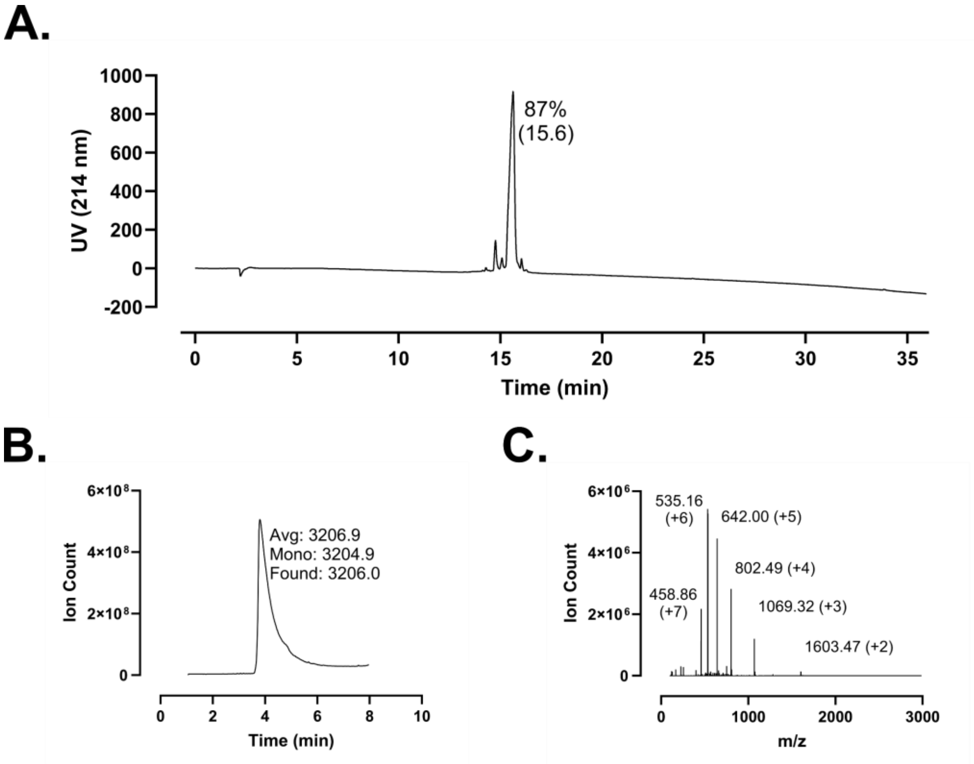
A.
B.
C.
UV (214 nm)
Time (min)
87%
(15.6)
Ion Count
Avg: 3206.9
Mono: 3204.9
Found: 3206.0
m/z
535.16
(+6)
642.00 (+5)
802.49 (+4)
458.86
(+7)
1069.32 (+3)
1603.47 (+2)

**Figure S207 – (#48) DVB-LL-25 Analytical Data. A.** HPLC trace at 214 nm of N-lipidated peptide following single-pass purification. Peak labeling indicates percent purity by trace integration. Retention time is given in parentheses. **B.** LCMS total ion chromatogram (TIC) of peptide after single-pass purification. Peak labeling indicates the predicted average and monoisotopic masses as well as the observed peptide mass. **C.** Mass spectrum at peak of **B** with *m/z* labeling for selected peaks and the associated charge in parentheses.

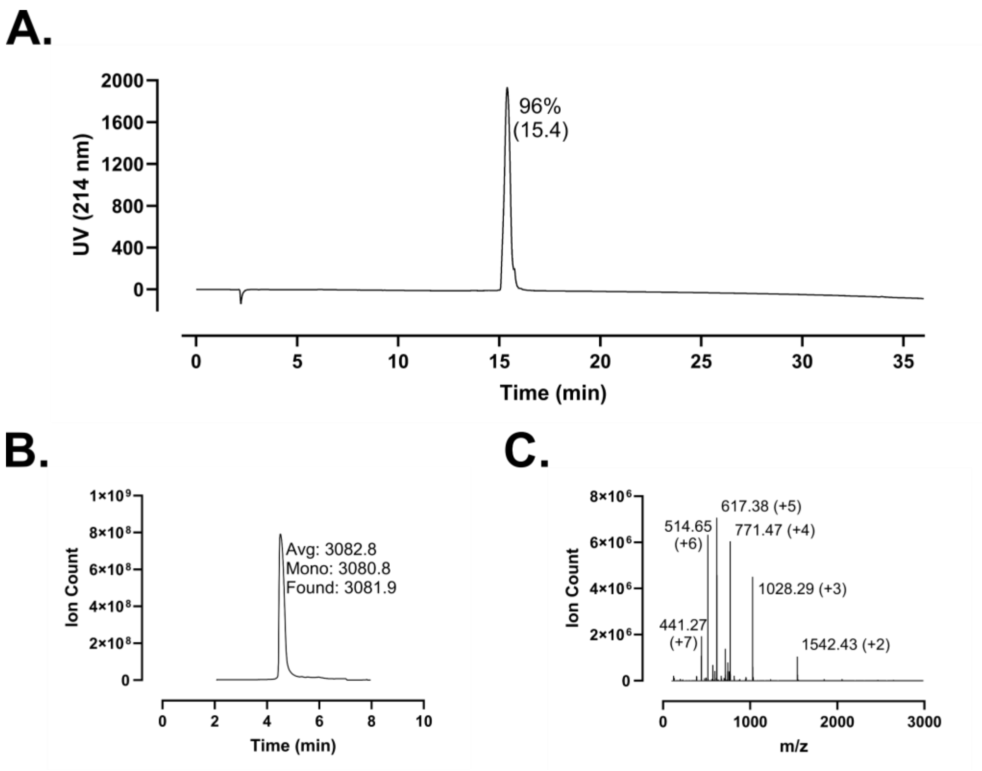
A.
96%
(15.4)
UV (214 nm)
Time (min)
B.
Avg: 3082.8
Mono: 3080.8
Found: 3081.9
Ion Count
Time (min)
C.
617.38 (+5)
514.65
(+6)
771.47 (+4)
1028.29 (+3)
441.27
(+7)
1542.43 (+2)
Ion Count
m/z

**Figure S208 – (#49) LL-25 (FF) Analytical Data. A.** HPLC trace at 214 nm of peptide following single-pass purification. Peak labeling indicates percent purity by trace integration. Retention time is given in parentheses. **B.** LCMS total ion chromatogram (TIC) of peptide after single-pass purification. Peak labeling indicates the predicted average and monoisotopic masses as well as the observed peptide mass. **C.** Mass spectrum at peak of **B** with *m/z* labeling for selected peaks and the associated charge in parentheses.

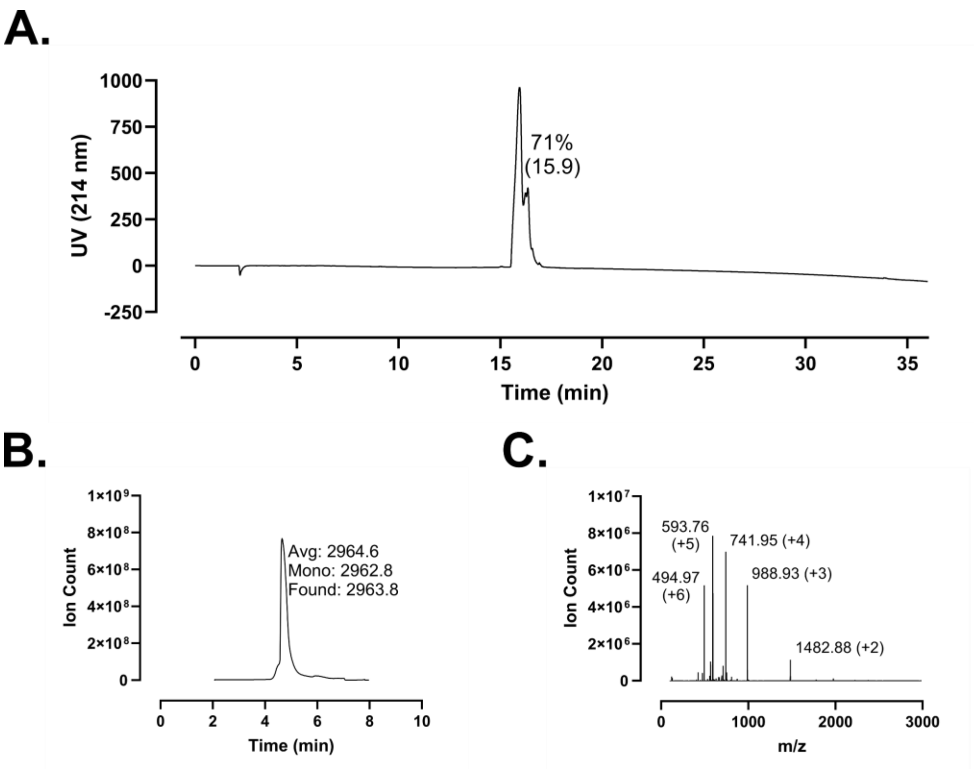
A.
UV (214 nm)
1000
750
500
250
0
-250
71%
(15.9)
0
5
10
15
20
25
30
35
Time (min)
B.
Ion Count
1×10⁹
8×10⁸
6×10⁸
4×10⁸
2×10⁸
0
Avg: 2964.6
Mono: 2962.8
Found: 2963.8
0
2
4
6
8
10
Time (min)
C.
Ion Count
1×10⁷
8×10⁶
6×10⁶
4×10⁶
2×10⁶
0
593.76
(+5)
741.95 (+4)
494.97
(+6)
988.93 (+3)
1482.88 (+2)
0
1000
2000
3000
m/z

**Figure S209 – (#50) Oct-GD-23 Analytical Data. A.** HPLC trace at 214 nm of N-lipidated peptide following single-pass purification. Peak labeling indicates percent purity by trace integration. Retention time is given in parentheses. **B.** LCMS total ion chromatogram (TIC) of peptide after single-pass purification. Peak labeling indicates the predicted average and monoisotopic masses as well as the observed peptide mass. **C.** Mass spectrum at peak of **B** with *m/z* labeling for selected peaks and the associated charge in parentheses.

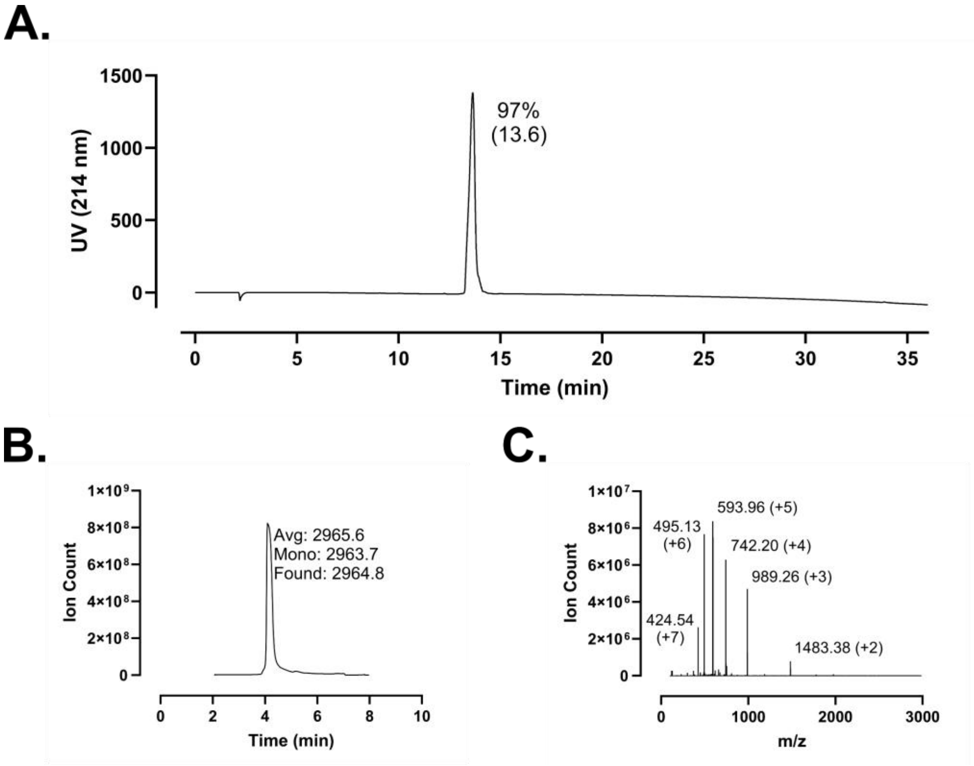
A.
1500
1000
500
0
UV (214 nm)
97%
(13.6)
0
5
10
15
20
25
30
35
Time (min)
B.
1×10⁹
8×10⁸
6×10⁸
4×10⁸
2×10⁸
0
Ion Count
Avg: 2965.6
Mono: 2963.7
Found: 2964.8
0
2
4
6
8
10
Time (min)
C.
1×10⁷
8×10⁶
6×10⁶
4×10⁶
2×10⁶
0
Ion Count
593.96 (+5)
495.13
(+6)
742.20 (+4)
989.26 (+3)
424.54
(+7)
1483.38 (+2)
0
1000
2000
3000
m/z

**Figure S210 – (#51) DMA-GD-23 Analytical Data. A.** HPLC trace at 214 nm of N-lipidated peptide following single-pass purification. Peak labeling indicates percent purity by trace integration. Retention time is given in parentheses. **B.** LCMS total ion chromatogram (TIC) of peptide after single-pass purification. Peak labeling indicates the predicted average and monoisotopic masses as well as the observed peptide mass. **C.** Mass spectrum at peak of **B** with *m/z* labeling for selected peaks and the associated charge in parentheses.

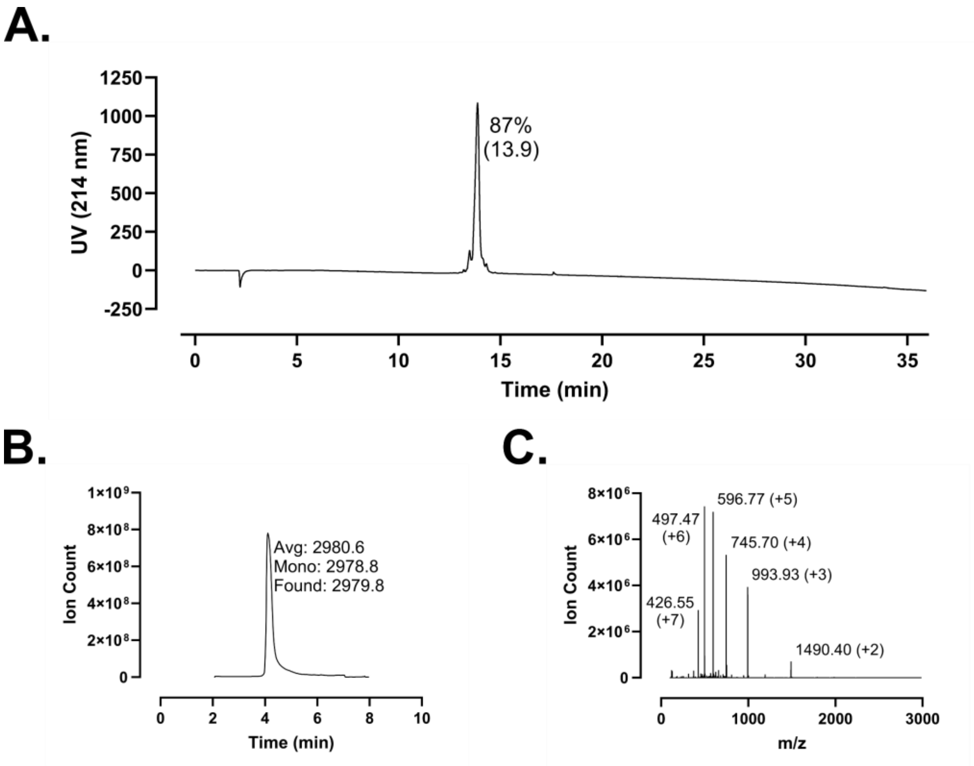
A.
UV (214 nm)
1250
1000
750
500
250
0
-250
87%
(13.9)
0
5
10
15
20
25
30
35
Time (min)
B.
Ion Count
1×10⁹
8×10⁸
6×10⁸
4×10⁸
2×10⁸
0
Avg: 2980.6
Mono: 2978.8
Found: 2979.8
0
2
4
6
8
10
Time (min)
C.
Ion Count
8×10⁶
6×10⁶
4×10⁶
2×10⁶
0
596.77 (+5)
497.47
(+6)
745.70 (+4)
993.93 (+3)
426.55
(+7)
1490.40 (+2)
0
1000
2000
3000
m/z

**Figure S211 – (#52) DVB-GD-23 Analytical Data. A.** HPLC trace at 214 nm of N-lipidated peptide following single-pass purification. Peak labeling indicates percent purity by trace integration. Retention time is given in parentheses. **B.** LCMS total ion chromatogram (TIC) of peptide after single-pass purification. Peak labeling indicates the predicted average and monoisotopic masses as well as the observed peptide mass. **C.** Mass spectrum at peak of **B** with *m/z* labeling for selected peaks and the associated charge in parentheses.

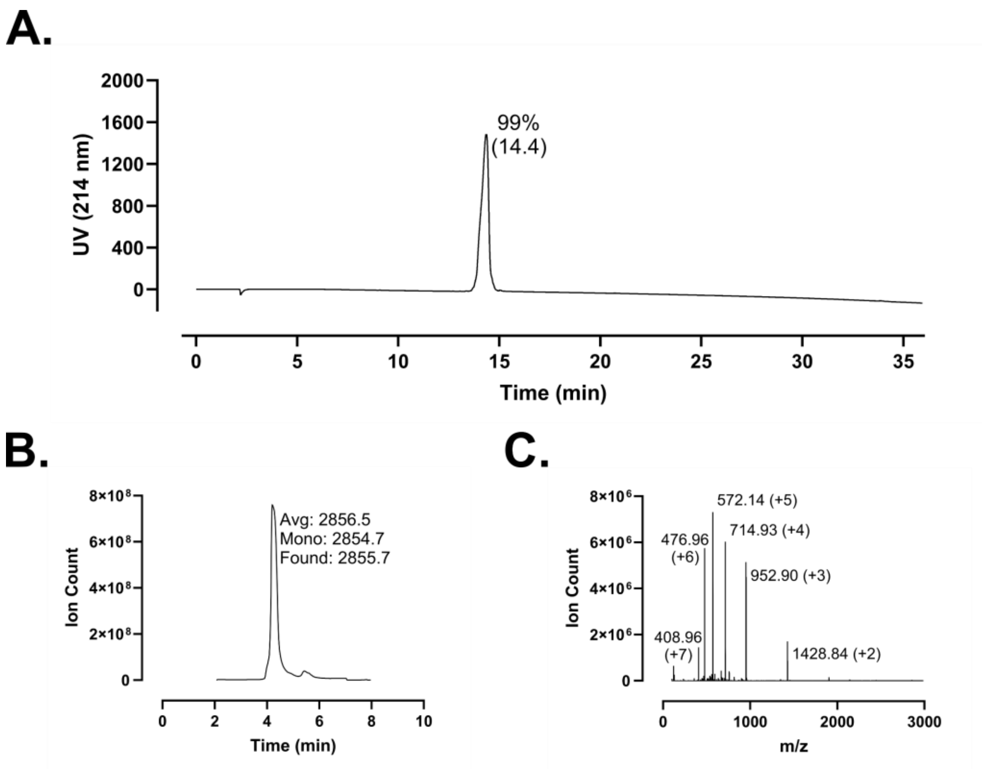
A.
99%
(14.4)
UV (214 nm)
Time (min)
B.
Avg: 2856.5
Mono: 2854.7
Found: 2855.7
Ion Count
Time (min)
C.
572.14 (+5)
714.93 (+4)
476.96
(+6)
952.90 (+3)
408.96
(+7)
1428.84 (+2)
Ion Count
m/z

**Figure S212 – (#53) GD-23 (FF) Analytical Data. A.** HPLC trace at 214 nm of peptide following single-pass purification. Peak labeling indicates percent purity by trace integration. Retention time is given in parentheses. **B.** LCMS total ion chromatogram (TIC) of peptide after single-pass purification. Peak labeling indicates the predicted average and monoisotopic masses as well as the observed peptide mass. **C.** Mass spectrum at peak of **B** with *m/z* labeling for selected peaks and the associated charge in parentheses.

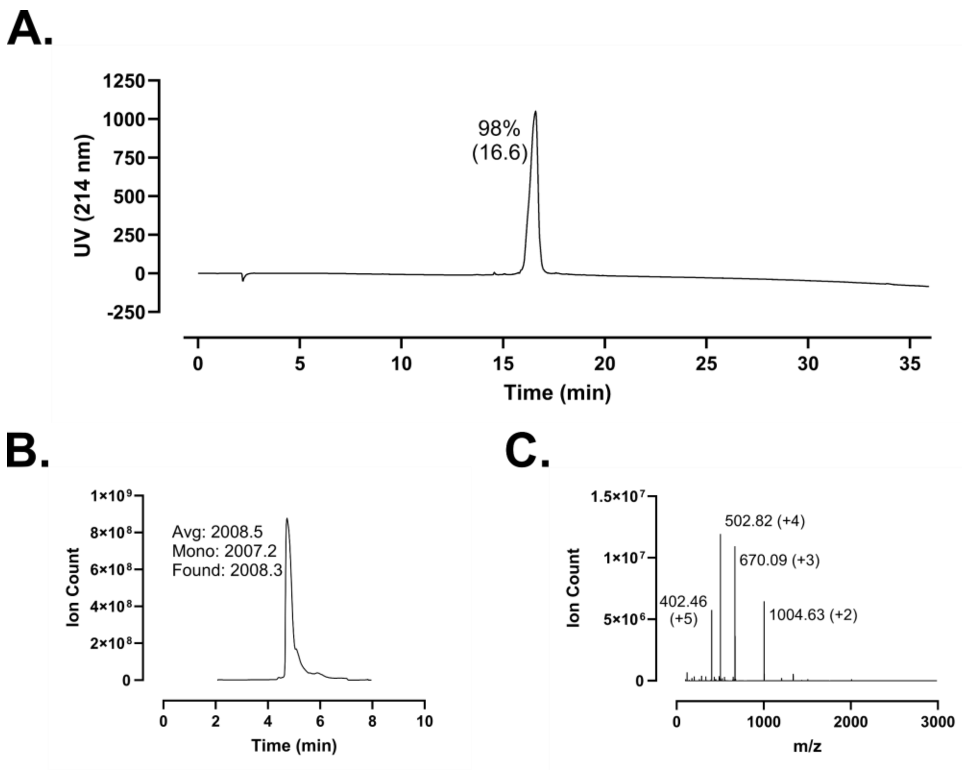
A.
98%
(16.6)
UV (214 nm)
1250
1000
750
500
250
0
-250
0
5
10
15
20
25
30
35
Time (min)
B.
Avg: 2008.5
Mono: 2007.2
Found: 2008.3
Ion Count
1×10⁹
8×10⁸
6×10⁸
4×10⁸
2×10⁸
0
0
2
4
6
8
10
Time (min)
C.
502.82 (+4)
670.09 (+3)
402.46
(+5)
1004.63 (+2)
Ion Count
1.5×10⁷
1×10⁷
5×10⁶
0
0
1000
2000
3000
m/z

**Figure S213 – (#54) DVB-FF-14 Analytical Data. A.** HPLC trace at 214 nm of N-lipidated peptide following single-pass purification. Peak labeling indicates percent purity by trace integration. Retention time is given in parentheses. **B.** LCMS total ion chromatogram (TIC) of peptide after single-pass purification. Peak labeling indicates the predicted average and monoisotopic masses as well as the observed peptide mass. **C.** Mass spectrum at peak of **B** with *m/z* labeling for selected peaks and the associated charge in parentheses.

A.

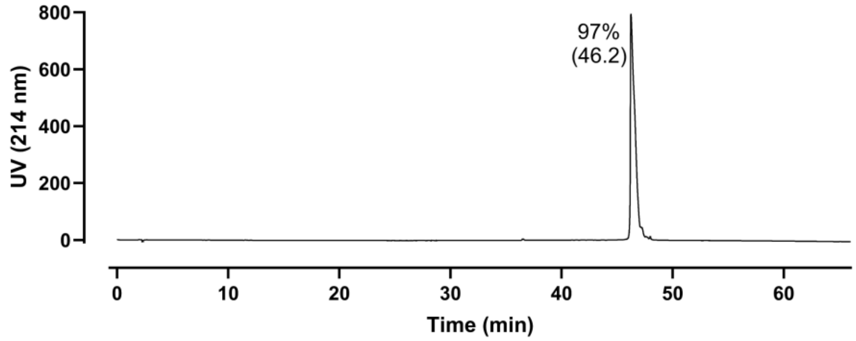

800
600
400
200
0
UV (214 nm)
97%
(46.2)
0
10
20
30
40
50
60
Time (min)


B.

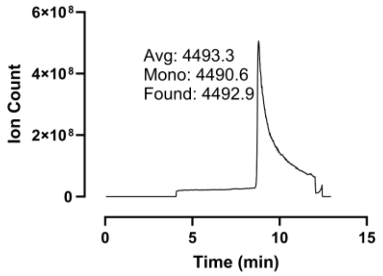

6×10⁸
4×10⁸
2×10⁸
0
Ion Count
Avg: 4493.3
Mono: 4490.6
Found: 4492.9
0
5
10
15
Time (min)


C.

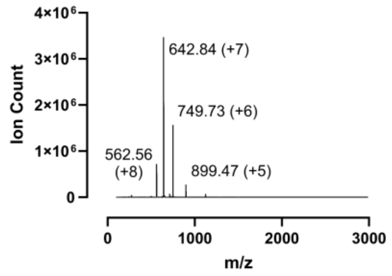

4×10⁶
3×10⁶
2×10⁶
1×10⁶
0
Ion Count
642.84 (+7)
749.73 (+6)
562.56
(+8)
899.47 (+5)
0
1000
2000
3000
m/z

**Figure S214 – (#1) LL-37 Analytical Data. A.** HPLC trace at 214 nm of peptide following single-pass purification. Peak labeling indicates percent purity by trace integration. Retention time is given in parentheses. The LL-37 preparation shown here was used in **Figure 6** and is distinct from the preparation used in **Figure S160**. **B.** LCMS total ion chromatogram (TIC) of peptide after single-pass purification. Peak labeling indicates the predicted average and monoisotopic masses as well as the observed peptide mass. **C.** Mass spectrum at peak of **B** with *m/z* labeling for selected peaks and the associated charge in parentheses.
